# Vitamin B_2_ Production by Vaginal Lactobacilli Promotes Symbiosis

**DOI:** 10.1101/2025.10.28.684342

**Authors:** Caroline E.M.K. Dricot, Denise M. Selegato, Tim Van Rillaer, Eline Cauwenberghs, Isabel Erreygers, Margo Hiel, Amber Brauer-Nikonow, Annelies Breynaert, Stefanie Wijnants, Isabel Pintelon, Sandra Condori, Sarah Ahannach, Thies Gehrmann, Sam Bakelants, Nina Hermans, Patrick Van Dijck, Michael France, Irina Spacova, Jacques Ravel, Michael Zimmermann, Sarah Lebeer

**Author notes:** These authors contributed equally to this work (Caroline E.M.K. Dricot, Denise M. Selegato). Shared senior authors (Michael Zimmermann, Sarah Lebeer).

## Abstract

The human vaginal microbiome, particularly with lactobacilli as the main inhabitants, plays a key role in maintaining women’s health. While lactic acid-mediated pathogen exclusion is well known, broader metabolic functions of vaginal lactobacilli remain underexplored. In this study, we analyzed the vaginal microbiome and metabolome of 258 healthy women from the Isala program. Using targeted metabolomics analysis, we detected a high prevalence with strong interpersonal differences of most B-vitamins, their precursors, and vitamin A in the vaginal microenvironment. Riboflavin (B_2_) and biotin (B_7_) showed strong associations with *Lactobacillus crispatus* and *Limosilactobacillus sp*. Comparative genomics, phenotypic assays, and *in vivo* metatranscriptomic data (VIRGO2) collectively confirmed riboflavin biosynthesis by these taxa. Using a riboflavin overproducing *Lim. reuteri* as a functional model, we showed that microbially derived riboflavin and its pathway intermediates are transported across the vaginal epithelium and modulate host redox balance, cytokine production, and activation of mucosal-associated invariant T (MAIT) cells via induction of MR1 (Major histocompatibility complex, class I-related protein receptor), revealing a potential immunometabolic interface between the vaginal microbiota and its host.

---

The human vagina hosts a unique microbial ecosystem that plays a central role in women’s health, reproduction, and well-being (*1*). Among mammals, humans exhibit a distinctive symbiosis with vaginal lactobacilli, which typically dominate the vaginal microbiota and contribute to protection against infections (*2*, *3*). In contrast, a shift toward a more taxonomically diverse microbial community, characterized by increased abundance of taxa such as *Gardnerella* and *Prevotella,* has been associated with conditions such as bacterial vaginosis, aerobic vaginitis, and pelvic inflammatory disease (*1*, *4–7*). The most prevalent and dominant vaginal *Lactobacillus* species, *Lactobacillus crispatus*, *Lactobacillus iners*, *Lactobacillus gasseri*, and *Lactobacillus jensenii,* contribute to colonization resistance largely through fermentation of epithelial glycogen into lactic acid, thereby maintaining a low pH environment unfavorable to several pathogens (*4*, *5*). While the role of lactic acid in maintaining vaginal health is well established, little is known about other metabolites produced by these microbes that may mediate host-microbe interactions. To address this knowledge gap, we established *Isala,* a large-scale citizen-science program that enables high-resolution characterization of the vaginal microbiome through integrated metagenomic, metabolomic, and culturomic analyses of vaginal swabs from healthy volunteers (*6*). This approach is especially valuable for mechanistic research, given the limitations of animal models in capturing human-specific vaginal ecology.

Notably, one of our first cultured isolates was a fluorescent, spontaneously riboflavin-overproducing *Limosilactobacillus reuteri* strain from a vaginal sample (*7*), raising the question of whether B-vitamin biosynthesis is a conserved feature of vaginal symbiosis. Riboflavin and its vitamers are recognized as key non-peptidic antigens for mucosal-associated invariant T (MAIT) cells (*8*), making them compelling candidates for exploring a host-microbe immunometabolic interface. In this study, we systematically investigated B-vitamin production and associated redox cofactors by vaginal *Lactobacillaceae,* leveraging the Isala dataset (*6*) and the publicly available VIRGO resource (*9*). First, using an integrated approach, combining microbiome sequencing and metabolome profiling, we traced the origin of B-vitamins, and evaluated if their production could be either food-derived or microbially produced. Following, we assessed the functional relevance of these vitamins by evaluating the riboflavin production capacity in bacteria isolates with predicted biosynthetic capacity, and their relation with key regulatory elements in riboflavin biosynthesis. Lastly, using cell culture analysis, we displayed the metabolic and immunomodulatory effects of riboflavin on the vaginal epithelium, including the activation of the highly specialized MR1-MAIT cell axis.

## Vaginal riboflavin (B_2_) and biotin (B_7_) vitamers are associated with *Lactobacillus crispatus*

The Isala participants included in this study (n = 258) were selected from the original program based on their age (31.6 ± 7.9 years), history of vaginal complaints, and contraceptive method (combination pill, intrauterine device, or no contraceptive). Participants provided self-collected vaginal samples at two distinct timepoints, approximately three months apart, and completed detailed questionnaires addressing dietary factors and lifestyle habits (Fig. S1).To further characterize this sub-cohort, we quantified 97 metabolites related to host-microbial metabolism (Data S1) in vaginal secretions using liquid chromatography coupled with mass spectrometry, with a specific focus on vitamins and associated metabolic variations. The 97 compounds included in our panel are relevant for the host and human microbiota and include vitamins, cofactors (NAD+, NADH, FMN), nucleotides, amino acids, tryptophan-like molecules (indoles and kynurenines), and bile acids (Fig. S2, Data S1). From this panel, 22 molecules were vitamers, of which 16 could be detected above the Limit of Detection (LOD) in at least 5% of the samples. Overall, pantothenic acid (B_5_), riboflavin (B_2_), the precursor of cyanocobalamine-B_12_ (5,6-dimethylbenzimidazole), PABA (B_9_ precursor), biotin (B_7_), retinoic acid (vitamin A), FMN (B2-derivative) and nicotinuric acid (a metabolite of nicotinic acid, one of the vitamers of vitamin B3) were most frequently detected (>80% prevalence) (Fig. 1A & B). However, these vitamins also showed strong interpersonal differences, indicating that their detection (and possibly also production) is environment-, host- and context-dependent. Of note, riboflavin and its intermediates had a prevalence between 29 and 84%, with most of its intermediates detected in more than a third of the samples analyzed.

**Fig. 1:**
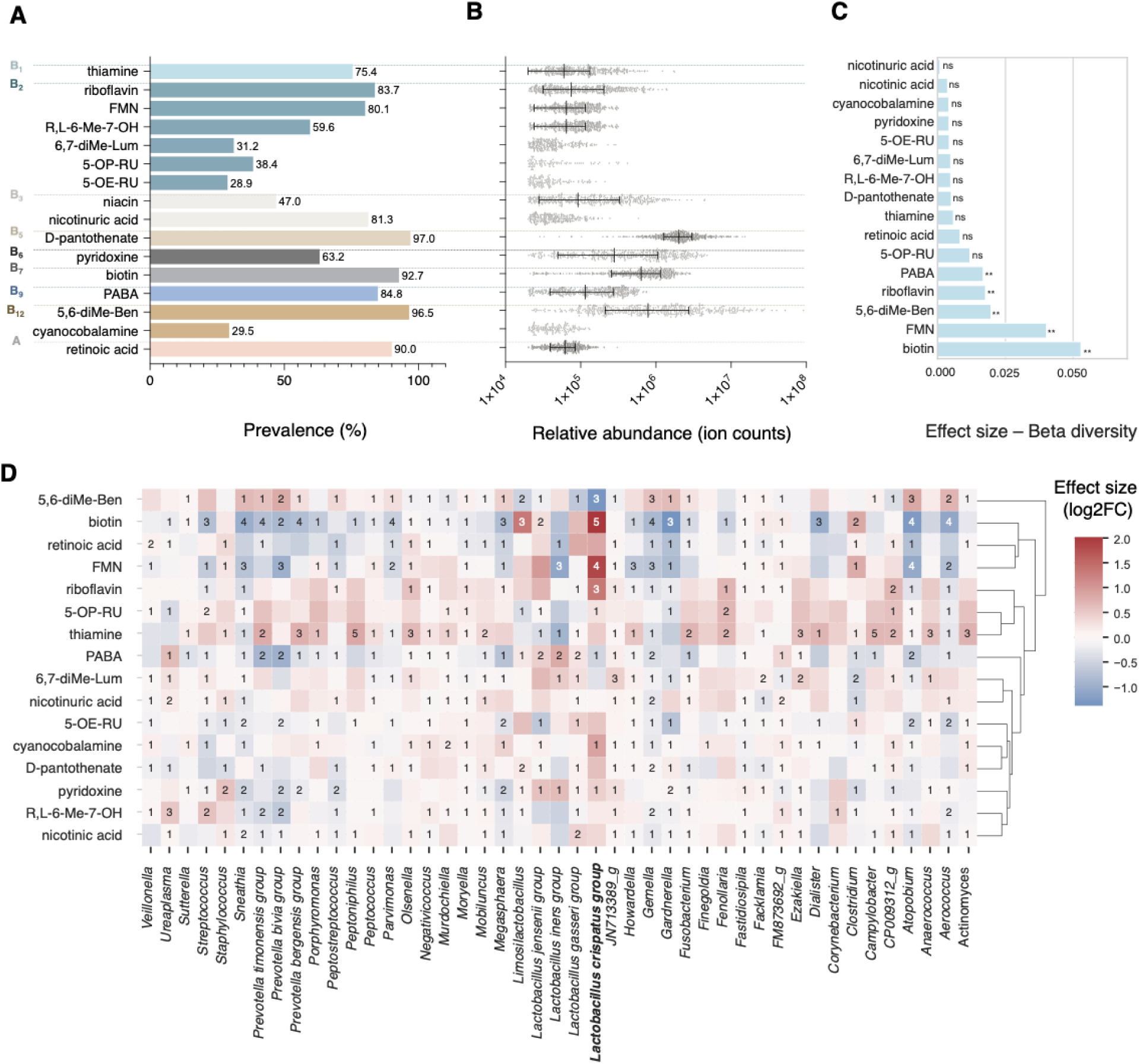
Prevalence of B-vitamins in the vaginal metabolome and their associations with ß-diversity, and individual taxa: **(A)** Barplot presenting the prevalence (%) of vitamins and derivatives (B-complex and vitamin A) in 491 paired vaginal samples of the Isala program (258 participants, n(t_1_) = 258, n(t_2_) = 233). Only those vitamers with prevalence >5% are taken into account. **(B)** Scatter plot (median, interquartile range) showing the distribution of the raw vitamer ion counts (relative abundance), in log-scale. Statistical associations of the vitamer ion counts with Bray-Curtis ß-diversity (permanova, significance p-value < 0.05 = *, p-value < 0.01 = **, p-value < 0.001 = ***). **(D)** Heatmap including all individual taxa with at least one significant association with one of the vitamers (linear regression the CLR-transformed abundance data, FDR adjusted and using a threshold of 0.05). The number refers to the number of pipelines indicating a significant association between the vitamin ion counts: ALDeX2, ANCOM-BC, DESeq2, limma, Maaslin2, Corncob, lmclr. The effect sizes (log2FC) are presented by the double-coloured gradient.

We subsequently tested whether the abundance of these compounds could be associated with self-reported dietary factors and lifestyle habits from Isala questionnaires. Although a direct dietary origin is less obvious for metabolites found in the vagina compared to the gastrointestinal tract, these vitamins could possibly reach the vagina following intestinal absorption from the diet into the bloodstream and then be released into the vagina through specialized estrogen-sensitive riboflavin binding proteins and transporters (*10–13*). Riboflavin, for example, is abundant in dairy, meat, eggs and green vegetables (*14*). However, we did not find any significant associations between the presence of B-vitamins in the vagina and vitamin-related data from the questionnaires, including vitamin intake from supplements, vegetable consumption, fish, grain and dairy consumption, fermentation products, probiotics, or other possible endogenous causes, such as vaginal discomfort, hormonal contraceptives and number of pregnancies (Fig. S3). These findings indicate that the observed differences in vaginal vitamin levels might be driven rather by interpersonal differences in the vaginal microbiome than by dietary habits or general health states. Next, we investigated the abundance of these vitamins in relation to vaginal microbiome composition, as we speculate that they, at least partially, originate from *in situ* microbial production, similar to what has been reported for the intestine (*15–18*). We explored associations between vitamin levels and various microbiome parameters, including α- and β-diversity (Shannon index and Adonis test), community structure analysis using Latent Dirichlet Allocation (LDA)-based topic modelling to identify subcommunities, and individual taxa using seven differential abundance methods (Fig. 1D, Fig. S4, Fig. S5, Data S2). Biotin and active vitamin B_2_ (flavin mononucleotide; FMN) were most strongly associated with α-(adj. p-val 2.35e-7 and 0.141 resp.) and β-diversity (p-values_FDRcor_ = 8.5e-3 and 8.5e-3 resp.), although other vitamers of B_2_, B_9_ and B_12_ (5-OP-RU, riboflavin, 5,6-diMe-Ben, PABA, all p-values_FDRcor_ = 0.0085) were also significantly associated with differences in β-diversity (Fig. 1C). Topic modelling (Fig. S5) identified robust positive associations between the occurrence of *L. crispatus, L. jensenii* and *Limosilactobacillus* taxa (especially *Lim. vaginalis* and *Lim. reuteri)*, in line with the existence of a *L. crispatus* subcommunity (*6*), and this subcommunity was positively associated with riboflavin, FMN, and biotin, along with a consistent negative association with PABA (4-aminobenzoic acid). These associations (with the exception of biotin) were stable (p-values_FDRcor_ < 0.05) across both sampling points, supporting their potential biological relevance within the vaginal ecosystem. At the individual taxon level, the significant positive associations were confirmed for biotin, FMN (active form of vitamin B_2_), and riboflavin with *L. crispatus* based on five, four, and three differential abundance methods, respectively (Fig. 1D and Data S2), whereas these vitamers were negatively associated with the taxa *Sneathia, Atopobium,* and *Gemella,* which are considered less optimal for the vaginal microbiota.

These findings suggest that the marked interpersonal differences in vaginal biotin and riboflavin levels may be influenced by specific microbiota members. However, it remained unclear whether vitamin availability shaped the microbiome or was instead determined by it. To address this question, we next examined the capacity of key vaginal taxa for vitamin biosynthesis.

## Microbial riboflavin production by vaginal lactobacilli: genomic potential, *in vitro* validation and *in vivo* activity

Although both riboflavin and biotin were significantly associated with *L. crispatus* and its subcommunity taxa *L. jensenii* and *Limosilactobacillus* spp., we focused on riboflavin and its potential microbial production due to its well-characterized and genomically tractable biosynthetic pathway (*16*, *17*) and our previous discovery of a vaginal isolate showing high riboflavin production (*7*). In contrast, *de novo* biotin production is genetically and phenotypically less straightforward to explore in bacteria and is uncommon in *Lactobacillales* (*18*).

First, we performed a large-scale comparative genomics analysis of the *rib* operon for riboflavin biosynthesis and transport across 1,862 bacterial genomes of species frequently found in the vagina, including 240 from bacterial isolates from the Isala program (DNBseq and Illumina HiSeq genome sequenced). All public genomes were downloaded from GTDB R220 and subsequently dereplicated to contain a maximum of 100 genomes per species with at most 99.98 ANI between two genomes in the same species (Data S3, S4). This analysis revealed that especially host-adapted species, including *L. crispatus* and its subcommunity taxa *L. jensenii*, and *Lim. reuteri*, consistently encode a complete riboflavin operon, in contrast to other taxa like *L. iners* and *L. gasseri*, lacking several synthesis genes (Fig. 2A). In addition, four other *Lactobacillaceae* species, which are commonly but not abundantly detected in the vagina (with prevalences 0.16% - 0.53% and relative abundances between 0.16% and 0.0017% genus-level in the Isala dataset (*6*)(i.e. *Lactobacillus mulieris, Lactobacillus delbrueckii, Lactiplantibacillus plantarum* and *Leuconostoc mesenteroides*) appeared to be able to produce riboflavin *de novo* based on their genomes (Fig. 2A). In contrast, *Lacticaseibacillus rhamnosus* and *Lacticaseibacillus paracasei* did not contain the full *rib* operon, while they have in general larger genomes (between 1.79 and 2.95 in Mbp) compared to *L. crispatus, L. jensenii, Lim. reuteri* and *Lim. fermentum* (average genome sizes between 1.61 Mbp and 2.06 Mbp) (*19*). This pattern suggests that, even when vaginal taxa such as *L. crispatus* and *L. jensenii* evolve toward a more symbiotic lifestyle characterized by genome reduction, selective pressures may favor the maintenance of essential metabolic functions such as vitamin B_2_ synthesis.

**Fig. 2.**
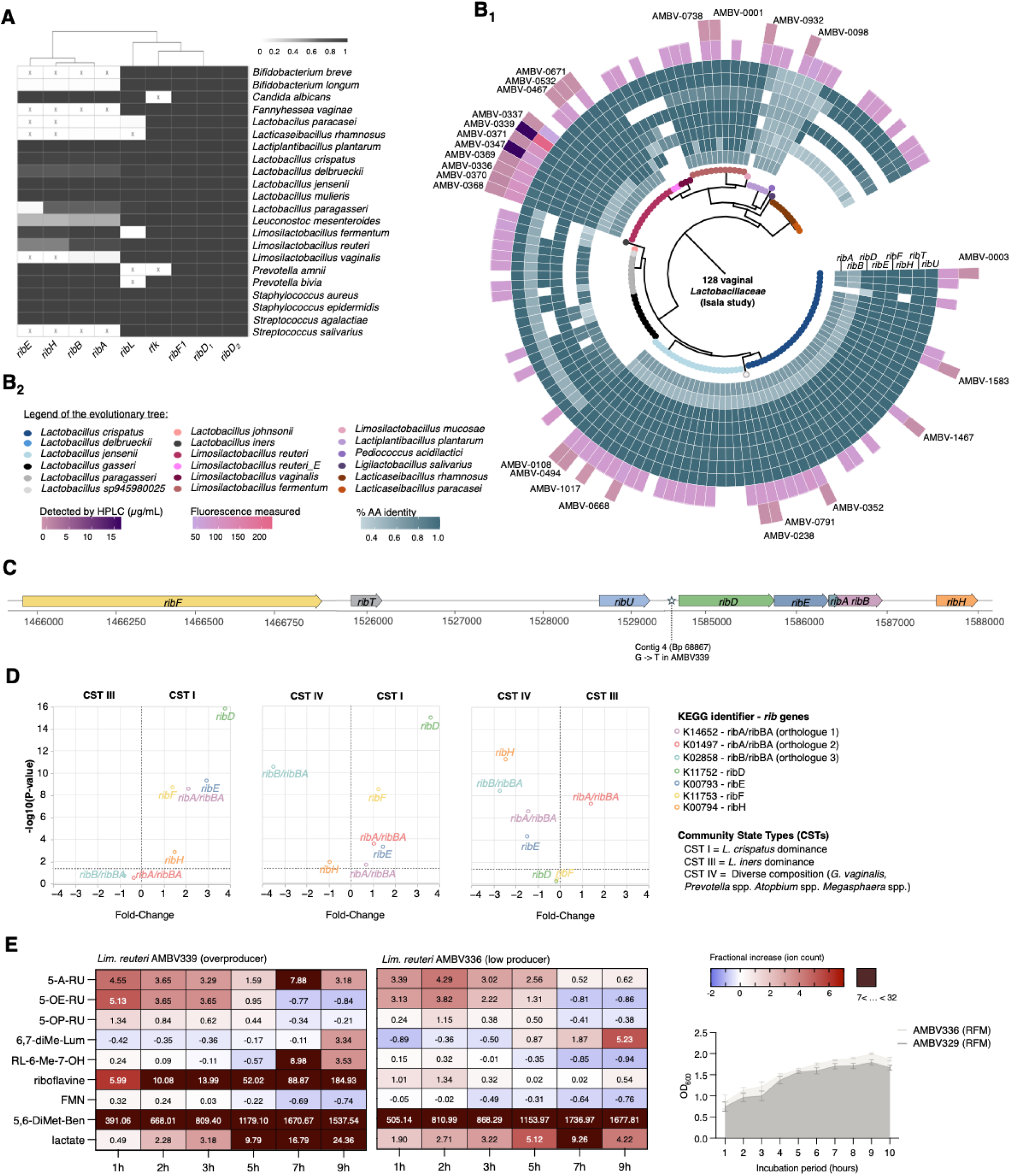
Genomic and phenotypic evidence for riboflavin biosynthesis in vaginal *Lactobacillaceae*. **(A)** Comparative genomics analysis of riboflavin biosynthesis genes across 1,862 publicly available genomes. Crossed boxes indicate a complete absence of gene hits. **(B)** Riboflavin operon structure in *Lactobacillaceae*, including the RFN riboswitch element. A G-to-T substitution in this element in *Lim. reuteri* AMBV339 and AMBV347 disrupts negative feedback regulation, resulting in riboflavin overproduction. **(C)** *In silico* and *in vitro* assessment of riboflavin production in 128 vaginal Isala isolates, visualized on a GTDB-based phylogenetic tree. Riboflavin overproducers *Lim. reuteri* AMBV339 and AMBV347 were identified. Volcano plots present differential gene expression (fold-change) of *rib* genes between vaginal microbiomes belonging to Community State Types (CSTs) I, II and IV (VIRGO, n = 180). **(E)** The impact of the G-to-T substitution on the negative feedback mechanism is demonstrated by the quantification of riboflavin (and its intermediates) in the supernatant of *Lim. reuteri* AMBV339 (overproducer) and *Lim. reuteri* AMBV336 (low producer) over a course of 9h. Fractional increase of the compounds to their original concentration in riboflavin-free medium (RFM) is presented by color intensities.

To confirm phenotypic riboflavin production capacity in the Isala isolates with predicted biosynthetic capacity (Fig. 2B_1_), we screened 128 *Lactobacillaceae* strains for riboflavin fluorescence using an absorbance UV assay (excitation at 485 nm, emission at 528 nm) in different growth media, including De Man-Rogosa-Sharpe (MRS), riboflavin-free medium (RFM) and simulated vaginal fluid (SVF). This screen identified 74 candidate riboflavin producers under the tested laboratory conditions (Fig. 2B_1_, second outer circle), while the remaining candidates did not produce detectable levels. Riboflavin production of the most interesting candidate producers was confirmed via High-Performance Liquid Chromatography (HPLC) (Fig. 2B_1_, outer circle) (Data 5). Among the tested strains, *Lim. reuteri* AMBV339 and *Lim. reuteri* AMBV347 exhibited markedly elevated riboflavin production compared to the other strains tested (ca. 18.36 μg/ml vs. 1.07 μg/ml of *Lim. reuteri* AMBV336). Their phenotype correlated with a guanine-to-thymine (G-to-T) substitution in the RFN riboswitch (Contig 4, BP 68867, T-tests p-value_FDRcor_ = 6.76E-08), a key regulatory element in riboflavin biosynthesis (Fig. 2C, Data S6) (*17*). Time-course analysis in RFM (Fig. 2E, and Fig. S8 for SVF and MRS) revealed sustained riboflavin accumulation in AMBV339 (T) over 9 hours, while levels plateaued in the control strain (low-producer) AMBV336 (G), which was isolated from the same women. Other B_2_ vitamers displayed distinct dynamics, with some intermediates declining over time, likely due to acidification by lactic acid production (*20*). This pattern is consistent with disrupted riboswitch-mediated feedback regulation in AMBV339 and suggests rapid within-host evolution toward enhanced vitamin B_2_ production. The presence of distinct clonal variants in the vaginal ecosystem of a single host supports the idea that selective pressures may favor mutualistic traits such as vitamin overproduction. To explore this further, we performed comparative genomics on seven closely related *Lim. reuteri* isolates from the same woman. This analysis only revealed 33 SNPs, with the G-to-T variant uniquely present in the two riboflavin overproducers. Metabolomic profiling across our host-microbe target panel of 97 metabolites (Data S1) linked this SNP to riboflavin overproduction and inversely to xanthine and methionine levels (Table S2), consistent with FAD-dependent enzymatic activity. AMBV339 and AMBV347 also produced significantly more indole-3-carboxaldehyde (I3A; FDR-corrected p = 6.76×10⁻⁸). This relation was further supported by negative Spearman correlations of I3A (FDR-corrected p = 9.67×10⁻³; Data S7 and Data S8), suggesting a possible link between riboflavin and tryptophan metabolism.

We next investigated whether vaginal lactobacilli are also capable of producing riboflavin *in vivo* within the human vaginal environment. To this end, we leveraged the VIRGO dataset (*9*) a non-redundant gene catalog constructed from 194 vaginal metagenomes and 180 metatranscriptomes (of 39 North American women, sampled at 5 timepoints, spaced 2 weeks apart). This resource enabled high-resolution taxonomic and functional profiling of the vaginal microbiome, including metatranscriptomic evidence of active riboflavin biosynthesis pathways in *Lactobacillaceae*-dominated communities. As can be observed in the volcano plots (Fig. 2D), all riboflavin genes are significantly upregulated in CST I, dominated by *L. crispatus*. In this CST, the riboflavin transcripts were mainly derived from *L. crispatus* and *L. jensenii*, in line with their genetic potential shown above (Fig. 2B_1_).

## Host uptake and immune modulation by riboflavin derived from vaginal lactobacilli

To investigate the biological relevance of riboflavin production by vaginal *Lactobacillaceae*, we examined its uptake, metabolic impact, and immunomodulatory potential in a 3D vaginal transwell model (Fig. S9–S10). We first assessed epithelial barrier integrity using a permeability marker mix (10 µM) (*21*) and found that the model maintained a stable barrier under control conditions, and upon exposure to *Lactobacillaceae* supernatant (Fig. S11). Classical radiotracer assays using [³H]- and [¹⁴C]-labeled riboflavin confirmed both apical uptake (Fig 3A.) and basolateral transport (Fig. 3B, Data S9) across the epithelial model within 2 hours, sampled at 15–20 min intervals. In parallel, we measured our host-microbe target panel of 97 metabolites in the cellular content of the epithelial cells after bacterial exposure, as well as the supernatants of the riboflavin-overproducing *Lim. reuteri* AMBV339, the 17 fold lower producer AMBV336, and MRS media control. Exposure to *Lim. reuteri* AMBV339 induced a significant, dose-dependent enhancement of vaginal epithelial redox balance when compared to riboflavin-producing *Lim. reuteri* AMBV336, evidenced by significantly increased intracellular levels of all-trans-retinal, hypoxanthine, PABA, 5,6-diMe-Ben, coenzyme Q10 and reduced glutathione (p-values_FDRcor_ < 2.02E-3, log2FC > 1.7) (Fig. 3C), as well as vitamers and their precursors listed on Fig. S12, Table S4 and S5. Together, these data support the bioavailability of microbial riboflavin to host cells and suggest that riboflavin modulates host redox metabolism, revealing an underappreciated metabolic interface in the vaginal niche.

**Fig. 3:**
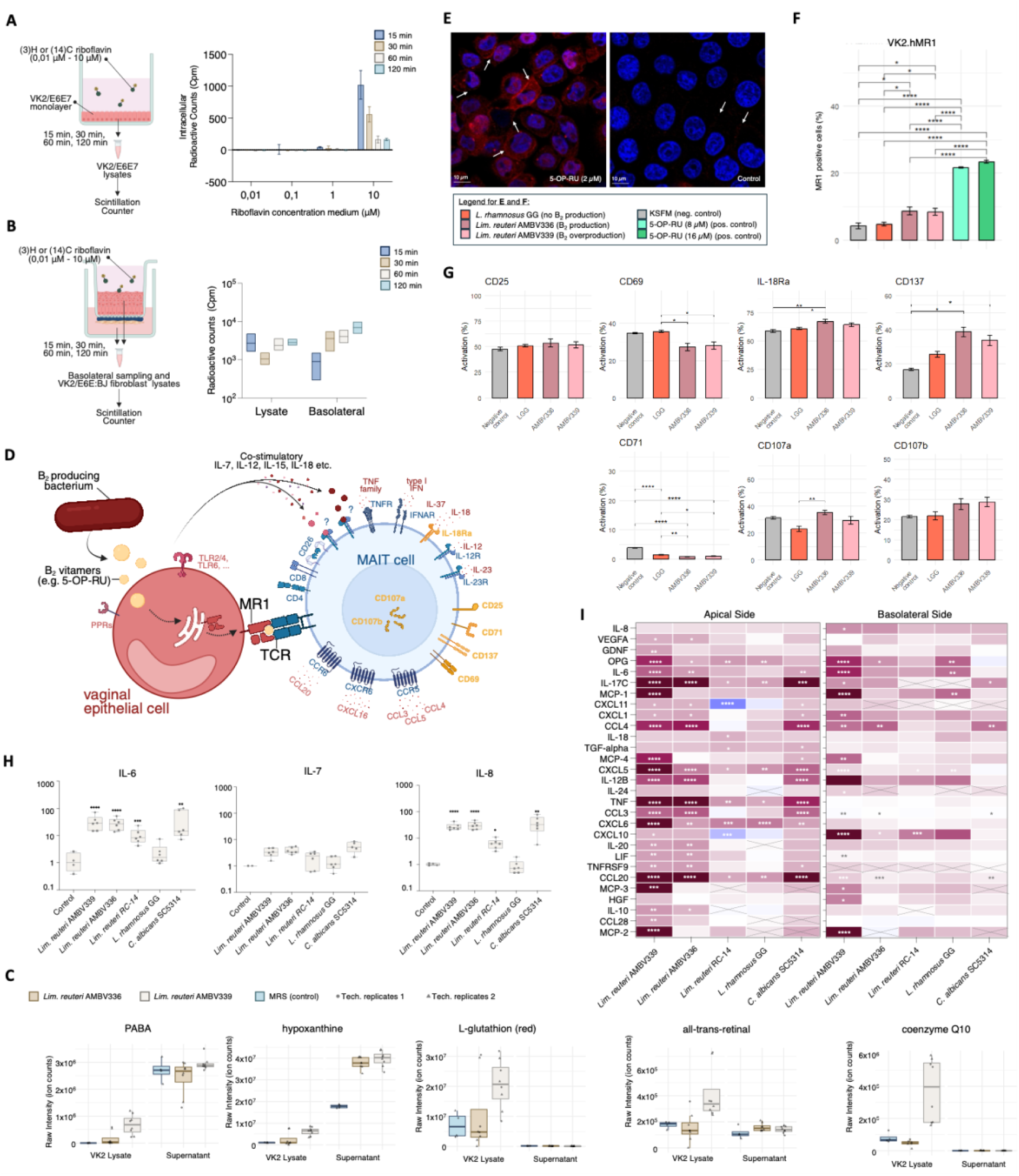
*Lactobacillaceae*-derived riboflavin is transported by the vaginal epithelium: **(A) & (B)** Uptake and transport assays of radioactive riboflavin in **(A)** a monolayer of VK2/E6E7 cells (n = 3, mean ±SD) and **(B)** the 3D vaginal transwell model (n = 3, mean, min-max). **(C)** The ion counts of redox-related molecules in VK2 monolayer lysates was significantly increased after a 3h exposure period to riboflavin overproducer *Lim. reuteri* AMBV339 compared to *Lim. reuteri* AMBV336 (presented as biological triplicates and technical duplicates, mean, 10-90 percentiles). **(D)** MR1-dependent MAIT cell activation by riboflavin intermediates, such as 5-OP-RU. The nature of the MAIT cell response is believed to be dependent on co-stimulatory signals from neighbouring cells in reaction to the riboflavin-producing microbe. MAIT cell surface and intracellular markers quantified during flow cytometry in (**G**) are highlighted in yellow. **(E)** Confocal fluorescence Microscopy images (92.3×92.3 microns) of VK2/E6E7.hMR1 cells stained for the MR1:5-OP-RU complex in red (anti-MR1 and anti-IgG2a Alexa Fluor™ 594) and nuclei in blue (DAPI) after 2h incubation with 5-OP-RU (2 µM). Images were processed in Fiji brightness/contrast blue channel (2-116) and red channel (6-100). White arrows indicate cell surface areas with enhanced red fluorescence. **(F)** Flow cytometry analysis of the percentage of MR1+ VK2/E6E7.hMR1 cells after incubation with 5-OP-RU or (riboflavin producing) microbes. **(G)** Flow cytometric analysis of MAIT cell activation and proliferation markers after exposure to (riboflavin-producing) microbes. **(H)** qPCR analysis of VK2/E6E7 monolayer lysates after 3h incubation 10% ON supernatant in KSFM (riboflavin-producing) microbes. **(I)** Olink® 96 inflammation panel on supernatant derived from both apical and basolateral sides from the vaginal 3D transwell model after 24 h exposure to (riboflavin-producing) microbes.

We subsequently assessed whether riboflavin-producing *Lactobacillaceae* could activate the MR1–MAIT cell axis in the vaginal environment (Fig. 3D), thereby possibly contributing to epithelial homeostasis, as previously described in other mucosal tissues (*22*, *23*). Although wild-type VK2/E6E7 cells endogenously express MR1, we engineered an MR1-overexpressing variant (VK2/E6E7.hMR1) via lentiviral transduction to improve detection sensitivity (Fig. S13). Using fluorescence microscopy (Fig. 3E) and flow cytometry (Fig. 3F, Fig. S14, Data 10), we observed a significant increase in MR1 expression in VK2/E6E7.hMR1 cells following exposure to the potent MR1 ligand 5-OP-RU and riboflavin-overproducing *Lim. reuteri* AMBV339 and low producer AMBV336, but not to the non-riboflavin-producing lactobacilli tested.

Similarly, peripheral MAIT cells responded to the riboflavin-producing *Lim. reuteri* strains (Fig. 3G) or their cell extract (Fig. S15, Data 10) with increased expression of activation/proliferation markers (CD25, IL-18Ra, CD137), a modest reduction in cytotoxicity-associated markers (CD69, CD71) and a variable change in CD107a (degranulation factor), indicating a homeostatic rather than pro-inflammatory activation profile. In line with this, we observed induction of the homeostatic TLR2/6 couple, but lack of induction of the more pro-inflammatory TLR4 by *Lim. reuteri* AMBV339 (Fig. S16). However, the role of TLRs in skewing the MAIT cell response is currently unclear (*8*).

To delineate the epithelial signals underlying this apparent homeostatic MAIT cell activation, we profiled the secretary immune response against a range of riboflavin producing and non-producing microbes in VK2/E6E7 monolayers and a 3D vaginal transwell model (Fig. 3H–I, Fig. S17–S19, Data S11, S12). By leveraging our genetic variants *Lim. reuteri* AMBV339 (overproducer) and AMBV336 (low producer), we aimed to identify cytokine signals linked to the riboflavin dose, while differences between these *Lim. reuteri* strains and *Candida albicans* SC5314 potentially indicate signals linked to the microbial nature, being either ‘commensal’ or ‘pathobiont’.

qPCR showed that besides the canonical vaginal cytokines IL-6 and IL-8, only IL-7 was upregulated among MAIT-relevant cytokines (Fig. 3H, Fig. S12, Data S11). Proteomic analysis revealed strong upregulation of MAIT-activating cytokines and chemokines (CCL3, CCL4, CCL20, IL12B, TNF) by all riboflavin-producing microbes (*Lim. reuteri* strains and *C. albicans)*, but not by non-producers (Fig. 3I, Data S12).

In the 3D model, AMBV339 induced anti-inflammatory IL-10 and barrier-supporting factors (e.g., VEGFA, GDNF, OPG, HGF), as well as chemoattractants (e.g., MCP1–4, CXCL10–11, CCL28) and pleiotropic cytokines (e.g. IL-6, IL-20, LIF, IL-17C, and TNFRSF9) (Fig. 3I, Data 3). The latter, TNFRSF9 (or CD137 or 4-1BB), is of special importance, as it is an established MAIT cell activation marker (*24*). Riboflavin dose-dependent differences between AMBV339 and AMBV336 were observed for MCP1, MCP4, OPG, CASP8, CDCP1, Flt3, and LAP-TGFβ. Conversely, several pro-inflammatory mediators, such as IL-1α and EN-RAGE, IL-18 (involved in MR1-independent MAIT activation) (*25*, *26*), DNER and ADA (a ligand for CD26 T lymphocyte stimulation), were induced by *C. albicans* in VK2/E6E7 monolayers, but suppressed by the *Lim. reuteri* strains (Fig. S18, Data 4). Together, these findings suggest that riboflavin-producing *Lim. reuteri* strains elicit a distinct, homeostatic immune profile compared to the pro-inflammatory response triggered by *C. albicans*, supporting protective MAIT cell activation and epithelial integrity in the vaginal niche.

## Discussion

Beyond classical roles such as lactic acid production (*27*), the broader metabolic potential of vaginal lactobacilli remains underexplored. In this study, we leveraged the Isala citizen-science program and integrated metabolomic, genomic, and biochemical approaches to investigate B-vitamin dynamics in the vaginal ecosystem. We found that vaginal *Lactobacillaceae* abundance correlated with biotin and riboflavin levels, with riboflavin biosynthesis genetically encoded and active in *L. crispatus*-dominated communities, where *L. crispatus*, *L. jensenii*, and *Limosilactobacillus* species typically co-occur in modules (*28*). Riboflavin was bioavailable to host cells, modulated epithelial redox metabolism, and triggered MR1-dependent MAIT cell activation. Vaginal epithelial cells responded by upregulating MR1, producing co-stimulatory cytokines, and releasing chemokines and repair factors, while peripheral MAIT cells were activated without a cytotoxic profile.

Riboflavin is well known for its essential role in cellular energy metabolism and oxidative stress regulation in humans. It functions either as a direct antioxidant in its reduced form (dihydroriboflavin) or as a co-factor for key antioxidative enzymes (*29*, *30*). Riboflavin plays an important role in women’s health due to increased physiological demands linked to menstruation, fertility, pregnancy, and pelvic tissue integrity and is implicated in neurological and psychiatric conditions such as postpartum depression, and migraine (*31*). In the gut, microbial riboflavin supports nutrient sharing and community stability (*32–35*), yet its biosynthesis by the vaginal microbiota was until now largely unexplored.

Our observation that exposure to riboflavin-overproducing lactobacilli significantly enhances 5,6-diMe-Ben, all trans retinal, coenzyme Q10, PABA, β-NAD and reduced glutathione in vaginal epithelial cells suggests a direct contribution of *Lactobacillaceae* to vaginal host redox homeostasis. In addition, we found that the microbial production levels of I3A were strongly upregulated due to riboflavin overproduction. Although the role of microbial indoles like I3A in the vaginal environment is still emerging, they are known to regulate barrier function and immunity via the aryl hydrocarbon receptor and protect against vulvovaginal candidiasis (*36*, *37*). We show that *Lim. reuteri* AMBV339 produces the riboflavin intermediate 5-A-RU and its immunomodulatory derivatives 5-OP-RU and 5-OE-RU, which activate MAIT cells via MR1 presentation, in line with the literature (*8*). While MAIT cells have been implicated in mucosal immune regulation in the gut and respiratory tract (*22*, *23*, *38*), their role in the vaginal environment remains largely unexplored, even though they have been found there (*39*). Our findings reveal that riboflavin-producing *Lactobacillaceae* induce MR1 surface expression and moderately activate MAIT cells, accompanied by epithelial release of cytokines and repair factors. Differences in IL-18, IL-10, IL-1α, EN-RAGE, and DNER responses between vaginal commensals and pathobionts support a model in which vaginal riboflavin production by lactobacilli contributes to epithelial homeostasis through MAIT cell modulation.

A key limitation of this study was the inability to perform transcriptomic profiling within the Isala self-sampling campaign, due to the use of non-invasive swabs and mRNA instability during room-temperature transport. To partially overcome this, we integrated metatranscriptomic data from the VIRGO dataset, though its metabolomics platform was not optimized for detecting unstable riboflavin intermediates. In contrast, Isala’s targeted metabolomics approach enabled sensitive detection of these compounds and highlighted the robustness of self-sampling for vaginal metabolome research. Another challenge is the lack of experimental models that accurately reflect the human vaginal microbiome, as conventional animal models lack *Lactobacillus* dominance (*40*). Advances in 3D culture systems (*40*), including the vagina-on-a-chip-model (*41*), offer promising platforms for upcoming mechanistic studies. However, *Lactobacillaceae* are quite refractory to genetic engineering (due to thickness of their Gram-positive peptidoglycan layer), and therefore we failed in this work to generate specific point mutations in the *rib* operon. In this study, we have circumvented this through screening of a unique collection of natural overexpressing isolates, some isolated from the same donor. Future research should refine culture models and engineering methods, explore inter-individual variation in microbial vitamin production, and expand in vivo sampling across diverse populations to validate microbial functions in both health and disease.

Aside from these limitations, our findings stimulate a new understanding of an immunometabolic axis between vaginal *Lactobacillaceae* and the host, centered on riboflavin biosynthesis and exchange. This work not only broadens the functional landscape of the vaginal microbiome but also lays the groundwork for precision microbiome-based strategies aimed at enhancing mucosal health through targeted delivery of beneficial microbial metabolites and their associated producer strain(s).

## Acknowledgements

We thank all Isala volunteers for their participation. The following colleagues and students were instrumental for completing this work: Inas Rahou, Leonore Vander Donck, Tom Eilers, Camille Allonsius, Vanessa Croatti, Sam Bakelants, Nele Van de Vliet, Ines Tuyaerts (LAMB), Camille Gepts, Erin Fierens, Feline De Coninck, Nathan Donies (four previous students at LAMB), Mathias Wolfgang Gross and Nikita Denisov (EMBL) as well as the Flow Cytometry and Cell Sorting Core facility of the University of Antwerp (FACSUA) and Chemical Synthesis Core Facility at EMBL, specifically Dr. Murat Kucukdisli. We further also acknowledge the use of ChatGPT for spelling and grammar correction.

## Author contributions

Conceptualization: CD, SL, DS, MZ, JR

Methodology: CD, DS, TVR, EC, MH, ND, ABN, AB, SW, IP, TG, IE, SA, SB, MF

Investigation: CD, DS, TVR, EC, MF, IS, SL

Visualization: CD, DS, TVR, MH, ABN, IS, MF

Funding acquisition: SL, IS, MZ, CD, EC, SA, IE, SC

Project administration: CD, SA, SC, DS, SL,

MZ Supervision: SL, IS, MZ, NH, PVD, JR

Writing – original draft: CD, DS, SL, MZ

Writing – review & editing: all authors

## Diversity, equity, ethics, and inclusion

Our team actively cultivates an inclusive environment where researchers from different cultural, disciplinary, and demographic backgrounds are encouraged to contribute their unique viewpoints. We also embrace diversity across gender identities, life stages, and cultural backgrounds, acknowledging that the vaginal microbiome is shaped by lived experiences as much as by biology. Through participatory research initiatives, such as citizen science projects on female microbiota, we strive to bridge scientific discovery with society awareness, ensuring that our research contributes to more equitable diagnostics, therapeutics, and health literacy for all women.

## Data and materials availability

All metabolomics datasets are available at MetaboLights Database (https://www.ebi.ac.uk/metabolights/) under accession number MTBLS7389. The sequencing data of the vaginal microbiome communities (Isala samples) was previously deposited at the European Nucleotide Archive (ENA) under bioproject PRJEB50407 (for the first timepoint) and PRJEB86101 (for the second timepoint). Whole genome sequences of Isala *Lactobacillaceae* isolates will be available at NCBI upon publication, and can be linked to their isolation source using the ENA IDs in Data S4 (file update upon publication). All other datasets are included in the Supplementary Materials.

New codes (.r, .py, .sh) are deposited at https://github.com/LebeerLab/vitamins-paper-CD. Other, reused codes, are publically available:

i. https://doi.org/10.1093/molbev/msab293
ii. https://doi.org/10.21105/joss.06313
iii. https://github.com/ravel-lab/MTMG_longitudinal
iv. https://doi.org/10.1093/bioinformatics/btae735,
v. https://github.com/thiesgehrmann/multidiffabundanc

## Supplementary Materials

## Materials and Methods

### Cohort

The citizen-science initiative Isala was approved by the Central Ethical Committee of Antwerp University Hospitals and the University of Antwerp (B300201942076) and registered at ClinicalTrials.gov (NCT04319536). In total, 3345 healthy, non-pregnant women donated two vaginal swabs for microbiome and metabolome profiling and culturomics (*6*). In addition, data on lifestyle and health were collected via questionnaires. From this large Isala cohort, 258 participants of reproductive age (average age = 31.6 ±7.9) were invited, based on their contraceptive use (combination pill, intrauterine device, no contraceptive) and general health, to donate a second pair of swabs after a few months. Additional metabolomics analyses were performed on the samples from this smaller cohort.

### Data collection and processing

Each participant from the Isala sub-cohort (n = 258) donated two vaginal swabs at two timepoints: an eNat^TM^ (Copan, Brescia, Italy), intended for microbiome profiling, and an ESwab^TM^ (Copan, Brescia, Italy) intended for culturomics and metabolomics. All swabs were stored at home in the fridge until transport at room temperature with prepaid services by the national parcel service (Bpost). Upon arrival at the lab, the eNat^TM^ swabs were immediately stored at -20°C until further processing, while the ESwabs were stored shortly at 4°C. After a maximal storage time of 6 hours, the ESwabs were vortexed for approximately 15 seconds and separated in two aliquots of 500μL. The first aliquot was stored at -80°C in a 96 tube Micronic plate with 500μL 50% glycerol (for culturomics), the other was centrifuged for 3 min at 13,000 g, and its supernatant was subsequently divided in a 96 tube Micronic plate at -80°C. These supernatants were stored for approximately 6-12 months before shipping them on dry ice to EMBL (Heidelberg, Germany) for metabolomics analysis. Upon arrival, samples were directly put at -80°C until further analysis in the lab.

### Metabolome analysis

#### Sample preparation and Data acquisition

Untargeted metabolomic analysis of the biological samples was performed using liquid chromatography and mass spectrometry. Biofluids were prepared by cold organic solvent (ACN:MeOH 1:1) extraction after the addition of an internal standard mix (phenylalanine-*d5*, tryptophan-*d5*, ibuprofen-*d4*, tolfenamic acid-*d4*, estriol-*d3*, diclofenac-*d4*, warfarin-*d5*, oxfendazole-*d3*, chloramphenicol-*d5*, nafcillin-*d5* and caffeine-*d9)*, each to a final concentration of 80 nM. One the one hand, chromatographic separation of the vaginal samples and bacterial cultures (described below) was performed by normal-phase methodology using an Agilent 1200 Infinity UHPLC system and an InfinityLab Poroshell 120 HILIC-Z (Agilent, 2.1 x 150 mm, 2.7 Micron particle size) column. Aqueous mobile phase (solvent A) was MiliQ water with 0.05% Agilent deactivator (v/v) and 5 mM of ammonium acetate pH 9 buffer and organic solvent B was acetonitrile with 0.05% Agilent deactivator (v/v) and 5 mM of ammonium acetate pH 9 buffer. Column operated at 50 °C. 3 µL of sample were injected at the following gradients: 0 min 96% B; 2 min 96% B; 5.5 min 88% B; 8.5 min 88% B; 9 min 86% B; 14 min 86% B; 17 min 82% B; 23 min 65% B; 24 min 65% B; 24.5 min 96% B; 26 min 96% B. The time between injections was 3 min. The qTOF instrument (Agilent 6550) was operated in negative scanning mode (50–1,700 *m/z*) at the following source parameters: VCap: 3500 V, nozzle voltage: 0 V, gas temp: 225 °C; drying gas 13 L/min; nebulizer: 35 psi; sheath gas temp 350 °C; sheath gas flow 12 L/min. Online mass calibration was performed using a second ionization source and a constant flow of reference solution containing the ions of ammonium TFA and hexakis (*m/z* 112.9857 and *m/z* 1033.9881, respectively). On the other hand, chromatographic separation of samples from the transport experiment (described below) was performed by reversed-phase chromatography (InfinityLab Poroshell HPH-C18, 2,1×100 mm, 1,9 um) using an Agilent 1200 Infinity UHPLC system and mobile phase A (H2O, 0.1% formic acid) and B (methanol, 0.1% formic acid), and the column compartment was kept at 45°C. 5 µL of sample was injected at 100% A and 0.6 mL/min flow, followed by a linear gradient to 95% B over 5.5 min and 0.4 ml/min. The qTOF instrument (Agilent 6546) was operated in positive scanning mode (50–1,000 m/z) with the following source parameters: VCap, 3,500 V; nozzle voltage, 2,000 V; gas temperature, 225°C; drying gas 13 l/min; nebulizer, 20 psig; sheath gas temperature 225°C; sheath gas flow 12 l/min. Online mass calibration was performed using a second ionization source and a constant flow (5 μl/min) of reference solution (121.0509 and 922.0098 m/z). All metabolomics data is available in the MetaboLights Database (https://www.ebi.ac.uk/metabolights/) under accession number MTBLS7389.

#### Targeted metabolomics analysis: peak identification and extraction

A library of metabolites of interest was made based on a subset of standards of the ‘Human metabolite library’ from MetaSci ^16^ (Data S1). This library mainly consists of B vitamins and derivates, cofactors and some primary metabolites of interest for the vaginal environment, including lactate. In addition, the list was extended with microbial riboflavin pathway intermediates and derivatives (Table S5), which were freshly synthesized at the EMBL Chemical Synthesis Core Facility ^17^ (Heidelberg, Germany) and stored at -80°C until MS acquisition. 1 µM solutions (in DMSO, H2O, or 70% ethanol) were acquired using the LC-MS method described above to determine accurate mass and retention times (Data S1) of all standards.

The spectral information from the in-house library was used on MassHunter Quantitative Analysis Software (Agilent, version 7.0) for the targeted extraction of the area under the curve (AUC) of each metabolite using the untargeted metabolomic data of (a) vaginal samples, (b) bacterial supernatant samples (both stationary and longitudinal experiment), and (c) the samples from the transwell assay, allowing tolerances for mass of 0.002 amu or 20 ppm and for a retention time of 0.3 minutes. The feature table was further processed using the statistical package R 4.1.0 in R 4.4.1 using RStudio/2023.06.1. In short, peaks with AUC background below 30,000 were imputed, and features that were detected in fewer than 5% of samples were excluded.

### Chemical synthesis of riboflavin vitamers

All reagents and solvents, including anhydrous solvents, were obtained from Sigma Aldrich, TCI, BLDpharm, and Thermo Fischer, and used without further purification. Air and water-sensitive reagents and reactions were handled under argon atmosphere. The reaction progress was monitored by UHPLC-MS. Flash chromatographic purification was performed on a Biotage® Isolera One purification system using Biotage® SFär silica or C18 flash cartridges. Nuclear magnetic resonance spectra were recorded on a Bruker Avance (400 MHz) NMR System at 298 K. UHPLC/MS analyses were performed on Agilent 1290 series equipment consisting of an Agilent 1290 quaternary pump, a 1290 sampler, a 1290 thermostated column compartment and a 1290 Diode array detector VL+ equipped with a quadrupole LC/MS 6120 and an Infinity 1260 ELSD. The analytical column used was a Titan C18 UHPLC Column (2.1 × 30 mm, 1.9 µm) operated at 40 °C and 1.5 ml/min flow rate with a gradient (10% to 15% B in 0.4 min, 15% to 100% B in 1.6 min, 100% B for 0.5 min) using water (A) and acetonitrile (B), both containing 0.1% TFA as solvents. Compound purity was determined by ELSD monitoring. 5-A-RU.HCl(*42*, *43*) and R,L-6-Me-7OH(*44*) were synthesized according to the literature. Unstable metabolites 5-OP-RU and 5-OE-RU were prepared similar to the literature (*45*) To a solution of 5-A-RU.HCl in degassed DMSO (4 mL, 20 mM) were added methyl glyoxal (5 equiv) or glyoxal (5 equiv). The mixture was stirred overnight for 5-OP-RU and 5 min for 5-OE-RU. The reaction mixtures were then directly analyzed by UHPLC-MS without further purification due to the instability of compounds.

#### Metabolite identification

All metabolites described in this manuscript were assigned a Level 1 annotation according to the Metabolomics Standards Initiative (MSI) classification system. Level 1 represents the highest degree of confidence in compound identification and requires direct comparison of the experimental data to an authentic reference standard analyzed under the same experimental conditions. This includes a match in retention time, accurate mass, and MS/MS fragmentation pattern (Data S1), ensuring unequivocal identification of the compounds.

### Microbiome analysis of vaginal samples

#### 16S rRNA amplicon sequencing

The vaginal microbial community composition of the selected 258 samples was determined from the eNAT swabs using 16S rRNA amplicon sequencing according to Ahannach *et al.*(*46*) and Lebeer *et al* (2023) (*6*). Sequencing data are available at the European Nucleotide Archive (ENA) under bioproject PRJEB50407 (for the first timepoint) and PRJEB86101 (for the second timepoint).

#### Isolation and identification of lactobacilli

From each Isala participant, 10 μL of the ESwab glycerol stock was inoculated on a small Petri dish (10 mL) with growth media (MRS, MRS + vancomycin, MRS + vit K + hem, YPD, BHI, TH, TSB/TSA or CB) and grown for 24-48h at 37°C and 5% CO2. After 24h, the plates were checked for colonies and, if present, one colony per plate was selected at random (max three isolates/woman). Part of this colony was inoculated in 10 mL MRS and grown overnight (ON) in 37°C and 5% CO2. Of the overnight culture, 800 μL was mixed with 800 μL 50% glycerol in labelled cryovials (Greiner Bio-one Cryo.STM) and stored in -80°C for future purposes. The rest of the colony was used for colony PCR for taxonomic identification with 16S rRNA gene Sanger sequencing, using universal primers 27 (5′-AGAGTTTGATCCTGGCTCAG-3′) and 1492R (5′-CTACGGCTACCTTGTTACGA-3′). The success of the colony PCR was checked with gel electrophoresis. Sanger sequencing was performed at VIB Genomics Core.

#### DNA extraction and Whole Genome Sequencing of isolates

Whole genome sequences were acquired by extracting DNA from overnight monocultures (10 mL MRS at 37°C, 5% CO_2_) following the P3 protocol in Alimolaei and Golchin(*47*). Illumina-based genomes were sequenced using the NexteraXT DNA Sample Preparation kit (Illumina, United States of America) and the MiSeq platform (Illumina, United States of America) using 2×250 cycles at the Laboratory of Medical Microbiology (University of Antwerp, Belgium) as previously described Spacova *et al.* (2022)(*7*). The genomes sequenced with DNBseq were sent for sequencing to BGI (location Hong Kong). Genome sequencing data was assembled using Shovill (https://github.com/tseemann/shovill). Genome completeness and contamination were assessed using checkM v1.0.13https://github.com/grp-bork/gunc) and GUNCusing v1.0.13 (https://github.com/grp-bork/gunc)^18^ using GTDBTk version Bakta version 1.9.4^19^ and classified using GTDBTk version 2.4.0^19^. Whole genome sequences are deposted in NCBI, together with their isolation source. ENA IDs will be made available upon publication in Data S3.

### In silico, in vitro and in vivo assessment of riboflavin production by lactobacilli

#### In silico: Identification of riboflavin synthesis genes

Key genes for riboflavin metabolism were identified in 1,862 publically available genomes (Data 2) and in-house Isala isolates (Data 3) via the Kyoto Encyclopedia of Genes and Genomes database (KEGG) PATHWAY database (https://www.genome.jp/kegg/pathway.html), after dereplication to contain a maximum of 100 genomes per species with at most 99.98 ANI between two genomes in the same species (https://github.com/bluenote-1577/skani)From the assembled vaginal isolate genomes, a selection was made of those belonging to the *Lactobacillaceae* family and with at least 70% completeness and at most 10% contamination. Those 240 genomes were mapped to the EGGNOG database V5.0^20^ using eggNOG-mapper version 2.1.5 (https://github.com/eggnogdb/eggnog-mapper). A core genome of the genomes was constructed using SCARAP (https://github.com/swittouck/scarap) and used to construct a phylogenetic tree with iqtree after gap removal with trimal.

#### In vitro: Quantification of the riboflavin production by isolates

For the high-throughput screening, *Lactobacillaceae* isolates (Data S3) were grown overnight (ON) at 37°C and 5% CO2 in MRS. The riboflavin synthesis capacity of the WGS isolates was investigated using riboflavin autofluorescence measurements (excitation at 485 nm and emission at 528 nm) and HPLC-UV analysis of whole cultures, following Spacova *et al.* (2022) (*7*). For the longitudinal experiments, overnight MRS cultures of riboflavin (over)producers *Limosilactobacillus reuteri* AMBV339 (18,36 μg/ml) and AMBV336 (1,07 μg/ml) were diluted 1:1000 in MRS. After a second overnight incubation, the cultures were centrifugated (10 min, 4°C, 2500 rpm) and washed in 10 mL of fresh medium (MRS, Riboflavin Assay Medium (RAM)(BD Difco) or Simulated Vaginal Fluid (SVF) (Table S3), before resuspending in a final 5 mL. Then, 20 μL of the cultures were added to 180 μL of the respective medium in a 96-well plate (10 plates, with 6 biological repetitions/plate, singular experiments) and incubated for 10 hours at 37°C. Every hour, OD_600_ was measured, the plate was centrifugated the supernatant was stored at at -80°C before targeted metabolomics analysis.

#### In vivo: Metatranscriptomics VIRGO dataset

Differential gene espression of the *rib* genes was asssed by reanalysis of the publically available VIRGO datatset (*9*, *48*). In short, the dream analsysis (https://github.com/ravel-lab/MTMG_longitudinal) was filtered using riboflavin KEGG identifiers for riboflavin synthesis.

### Cell culture models

#### Maintenance of VK2/E6E7 cells and BJ fibroblasts

The VK2/E6E7 (CRL-2616^TM^) cell line, originally derived from the normal vaginal mucosal tissue of a healthy premenopausal woman, and the BJ fibroblast (CRL-2522^TM^) cell line, taken from the foreskin of a male neonate, were cultured according to the manufacturer’s guidelines (ATCC^®^). In detail, both cell lines were grown in T75 flasks (CELLSTAR® by Greiner Bio-One) at 37°C, 5% CO2 and 100% humidity. Respectively, complete Keratinocyt Serum Free Medium (KSFM) (Gibco^TM^) (= supplemented with 25 mg Bovine Pituitary Extract (Gibco^TM^), CaCl_2_ (Promocell®) and 50 ng human Epidermal Growth Factor (Gibco^TM^)) and complete Eagles’ Minimum Essential Medium (EMEM) ( = with 10% Heat Inactivated Fetal Bovine Serum (FBS) (Gibco^TM^)) was used for VK2/E6E7 and BJ fibroblasts. For maintenance, cells were split at 70-95% confluency (approx. twice a week) at a ratio of +/- 1/4 respectively. For this purpose, cells were detached with 0.25% prewarmed trypsin/EDTA (Gibco^TM^). Trypsin was inactivated by adding 10 mL of prewarmed Dulbecco’s Modified Eagle Medium (DMEM) (GibcoTM), supplemented with 10% FBS (for VK2E6E7’s) or complete EMEM (for BJ’s). Cells were centrifugated (10 min, 300 x g) (Eppendorf^TM^), split and resuspended in complete medium. For monolayer experiments, VK2/E6E7 cells were seeded at a density of 1.5 × 10^5^ cells/mL in 12-well culture plates (CELLSTAR® by Greiner Bio-One) and used at 100% confluency (after 72h growth at 37°C, 5% CO2 and 100% humidity).

#### Initiation of the 3D model

To better replicate *in vivo* like conditions, we adapted the 3D transwell model of Edwards *et al.* (2022) with minor modifications (Fig. S9) (*49*). Briefly, 12-well ThinCert™ inserts (0.4 μm pores, Greiner Bio-One) were inverted and coated with 140–200 μL of chilled collagen mixture (200 μL of 10X Roswell Park Memorial Institute (RPMI) medium (Sigma^®^), 200 µL tissue culture water (Invitrogen), 50 μL of 1 M NaOH and 1600 μL of rat tail collagen (Corning^®^) at pH 6.5). The mixture was gelled for 30 min at room temperature (RT) in an inverted position, then for 3 h upright in a biosafety hood, and stored at 4°C for 48–72 h. BJ fibroblasts (ATCC #CRL 2522) were seeded onto the collagen-coated basal side (150 μL of 7×10⁵ cells/mL) and incubated at 37°C, 5% CO₂ for 6 h, followed by 48 h in EMEM supplemented with 10% FBS (1 mL basolateral, 500 μL apical). VK2/E6E7 cells (7×10⁵ cells/mL) were then seeded apically in KSFM, with media refreshed after 2 days. Four days post-VK2/E6E7 seeding, air–liquid interface cultures were established by removing the apical medium. The basolateral medium was changed every 2 days for 8–10 days, until a polarized multilayer was confirmed via confocal microscopy (4% paraformaldehyde fixation) and TEM (2.5% glutaraldehyde).

#### Microscopy imaging

For microscopy imaging of the 3D vaginal epithelial model, all inserts containing cells were washed twice with 1 mL Dulbecco’s phosphate buffered saline (DPBS, Gibco^TM^) (apical side) and 2 mL (basolateral side) and fixated with 4% paraformaldehyde (Light Microscopy and Fluorescence Microscopy) or 2.5 % glutaraldehyde in0.1M sodium cacodylate (Transmission Electron Microscopy) for a few days at 4°C. After transport to ACAM, the microscopy core facility of the University of Antwerp, the inserts containing the 3D vaginal epithelial multilayers were further processed. For light microscopy (LM), inserts were handled by a STP120 spin tissue processor (Epredia, Machelen, Belgium) and embedded in paraffin. 5 µm transverse sections were made, stained with hematoxylin & eosin (HE) and imaged on a Nikon Ti microscope (Nikon Europe B.V., Amstelveen, The Netherlands). For transmission electron microscopy (TEM), inserts were rinsed in 0.1 M sodium cacodylate-buffered (pH 7.4) 0.05 % CaCl2.2H2O, 7.5 % sucrose solution at room temperature (RT), post-fixed in 1 % osmium tetroxide in 0.033M veronal acetate buffer containing 4% sucrose. Samples were then rinsed and dehydrated in graded ethanol series (50%, 70%, 90%, 95%, and 2 x 100%) followed by a final dehydration step with propylene oxide. Then, transwell membranes were embedded in EM-bed812 and 50 nm transverse sections were made using an ultramicrotome (Ultracut EM UC7, Leica Microsystems, Wetzlar, Germany). Afterwards, the transverse sections were stained with lead citrate and examined with a Tecnai Spirit BioTwin electron microscope (FEI Europe B.V., Zaventem, Belgium) at 120 kV.

### Transport experiments

#### Barrier control

To assess the barrier integrity of the vaginal epithelial transwell model, 500 μL of an in-house developed drug control mix, consisting of 1 mM propranolol, antipyrine and haloperidol (with high permeability) and terbutaline, nadolol, etopside (with low permeability) in prewarmed Hanks Buffered Salt Solution (HBSS), was added to the apical compartment. Every hour (for 3 hours), 5 μL samples were taken from the apical and basolateral side to monitor transport of the drug compounds. Samples were analyzed by LC-QTOF using a HPH-C18-column in positive MS scanning mode described above. The area under the curve (AUC) for each compound was previously determined (*21*). Threshold values of the AUC were determined based on negative control samples, containing only water. Here, a threshold of 1e5 a.u. was determined for all compounds but etoposide, for which the threshold was set to 1e4. In addition, the impact of lactobacilli (*Limosilactobacillus reuteri* AMBV339, *Limosilactobacillus reuteri* AMBV336 and *L. rhamnosus* GG) on the epithelial barrier was also evaluated (3 biological replicates, singular experiments). Hereto, overnight bacterial cultures (in MRS) were diluted to an OD_600_ = 0.1, using 0.25 (OD) = 1 × 10^8^ CFU/mL, and incubated for an additional 9 hours at 37°C. The supernatant of every strain was harvested by centrifuging bacterial cultures at 2000g, 4°C for 10 min, and subsequently diluted 1:10 in prewarmed Hanks Buffered Salt Solution (HBSS).

#### Exposure to radioactive riboflavin

Uptake of [^14^C]-riboflavin and [^3^H]-riboflavin was conducted following Patel *et al.* (2012) on a fully confluent monolayer of VK2/E6E7 cells (4 biological replicates) or 3D vaginal epithelium model (5 biological replicates), respectively. Singular experiments were executed at the Laboratory of Molecular Cell Biology (KULeuven). In short, seeded plates were transported from Antwerp to Leuven on the day of the experiment. After half an hour of recuperation at 37°C, 5% CO_2_ and 100% humidity, cell surfaces were washed twice with 1 mL of prewarmed DPBS. Cells were subsequently incubated with 0.5 mL solution of [^14^C]-riboflavin (monolayer) (34531.8 cpm/nmol) or [^3^H]-riboflavin (100000 cpm/nmol) (3D model, apical compartment) in DPBS at 37 °C, in a CO_2_ producing sachet (brand), for 15 min, 30 min, 60 min or 120 min. For the 3D model specifically, 1 mL of DPBS was also added to the basolateral compartment. Samples (200 µL) were taken (apically and basolaterally) from each well right before the uptake of radioactive riboflavin was terminated by aspiration of the buffer and the addition of ice-cold stop solution (210 mM KCl, 2 mM HEPES, pH of 7.4). Cells were washed one more time in ice-cold stop solution, after which 1 mL commercial lysis buffer (from RNeasy Qiagen kit) was added to each well. After overnight incubation of the plates at room temperature, 500 μL lysate and 100 μL of the apical (and basolateral samples) were transferred to scintillation vials containing 3 mL of scintillation fluid (Lumagel Safe; PerkinElmer). Radioactivity of the respective solutions was determined using a scintillation counter (Hidex) (Data 9).

### Surface MR1 upregulation & MAIT cell activation

#### Generation of a MR1-overexpressing VK2/E6E7 cell line

The transduction of VK2/E6E7 cells with a pCMV-MR1-IRES2-GaussiaLuciferase-pEf1a-puromycin (GeneCopoeia^TM^, Fig. S12) was performed following the manufacturer’s guidelines. More specifically, VK2/E6E7 cells, seeded in 24-wel plates (VWR^®^ Tissue Culture Plate), were transduced overnight using Polybrene Transfection Reagent (EMD Millipore, 6 μg/mL) at Multiplicity Of Infection (MOI) rates (0, 0.5, 1, 1.5, 2 and 5). Succesfull integrants were selected using puromycin (Gibco^TM^, 1 μg/mL) for a total of 9 days, while refreshing the medium every 3 days. The supernatant was subsequently examined for Gaussia Luciferase (GLuc) activity using the Secrete-PairTM Gaussia Luciferase Assay Kit (GeneCopoiea^TM^).

#### MR1 staining, flow cytometry and fluorescence microscopy

In vitro stimulation (1h-3h) with either 5-OP-RU (1.6 μM to 16 μM) or (riboflavin producing) bacteria (ca. 1 × 10^7^ CFU/mL), VK2/E6E7 and hMR1.VK2/E6E7 were washed twice with DPBS. Cells were then either directly stained in Cell Staining Buffer (CSTB, Biolegend) with Hoechst (Thermofisher scientific) for nucleic acid visualization (fluorescence microscopy) or detached (0.25% trypsine/EDTA) and stained with LIVE/DEAD™ Fixable Aqua Dead Cell Stain Kit (Thermofisher Scientific) for dead cell exclusion (Flow Cytometry). Human TruStain FcX™ was used to block aspecific Fc receptor binding, before proceeding with cell surface staining for MR1 using Purified anti-human/mouse/rat MR1 Antibody (Biolegend, Cat. No 361102)) and Goat anti-Mouse IgG2a Cross-Adsorbed Secondary Antibody, Alexa Fluor™ 594 (500 μg), A-21135 (invitrogen) at the end concentration of 20 μg/mL. Cells were subsequently fixed using Inside fix solution (Miltenyi Biotek) and kept overnight at 4°C. For flow cytometry, samples were acquired using the NovoCyte Quanteon 621181110427 (ACEA Biosciences, Inc) and NovoExpress 1.6.0 software (Data S10) and analyzed with NovoExpress 1.6.0 and Prism software (Graphpad). For fluorescent confocal microscopy, cells were imaged using a Leica SP8 laser scanning confocal microscope (Leica Microsystems, Machelen, Belgium), at 20x magnification with N.A. 0.75, stacked and edited in Fiji (Brightness and Contrast level of min = 60 and max = 275).

#### MAIT cell activation

PBMCs were extracted from buffy coats (Rode Kruis), after 1:2 dilution in autoMACS Rinsing Solution (5% MACS BSA, Miltenyi Biotech)), by density gradient centrifugation using Lymphoprep and SepMate tubes (Stemcell^TM^ Technologies). PBMC’s (app. 1 × 10^6^ cells/500 μL MACS Freezing Solution (Miltenyi Biotech)) were gradually frozen (Mr. Frosty, Thermo Scientific™) at -80°C before long-term storage in liquid nitrogen (vapor phase). One day prior to the experiment, cells were thawed in RPMI-1640 medium (ATCC^®^) supplemented with 10% FCS (Gibco^TM^) and transferred to a 96 U-bottom plate (Greiner Bio-One) at a concentration of maximum 10^6^ cells/200 µL complete RPMI medium (Scepter^TM^ 3.0 (Millipore)) for overnight incubation at 37 °C. On the day of the experiment bacterial cultures were prepared at a starting OD₆₀₀ of 0.1 and incubated for 5 h at 37 °C to maximize the production of the riboflavin intermediate 5-OP-RU. Then, crude cell extracts (CE) were generated on ice by sonification (6 min, with 10:15 s on/off intervals and amplitude = 25%), centrifugation of 30 min (at max. 8603 g) and filter sterilization (3.0 μm). PBMCs were subsequently incubated for 6 h with 5-OP-RU (4-16 uM) or with a 1:10 mixture of CE and complete KSFM, supplemented with 5 living BpC of the respective strain. Brefeldin A (3.0 ug/mL, eBioscience^TM^) was added after the first 2 h to internalize cytokine production. The co-cultures were subsequently stained with Viobility 405/520 (live/dead) and MAIT cell antibody cocktail(*50*) (CD3, TCR Va7.2, IL18-Ra (CD218), CD25, CD69, CD137, and CD71) in CSTB (Table S4), following the manufacturers guidelines (Miltenyi Biotech). Cells were additionally treated with the Inside Stain Kit (Miltenyi Biotech), prior to intracellular staining of CD107a and CD107b. Finally, cells were resuspended in 50 µL CSTB and stored at 2-8°C in the dark until flow cytometric acquisition on the Novocyte Quanteon at the Laboratory of Experimental Hematology (FACSUA) (Data S10).

### Immune and redox response against riboflavin producing lactobacillaceae

For the following experiments VK2/E6E7 monolayers were incubated in triplicate with supernatant:complete KSFM 1:10 (derived after ON growth in MRS) or 1 × 10^7^ CFU in complete KSFM for 3h at 37°C, 5% CO2 and 100% humidity. Bacterial cultures included *Lim. reuteri* AMBV336 (Data S4), *Lim. reuteri* AMBV339 (Data S4), *Lim. reuteri* RC-14 (*51*), *Lactocaseibacillus rhamnosus* GG ((*52*)), *Candida albicans* SC5314 (*53*)(Data S3).

#### The intracellular redox status

After co-incubation, monolayers were washed twice with DPBS, lysed using ACN:MeOH (1:1) and supplemented with internal standard mix. Intracellular metabolites (Data S1) were measured in HILIC neg. mode as earlier described (3 biological replicates, 2 technical replicates, singular experiments).

#### qPCR

After co-incubation, cells were lysed by adding 600 μL RLT Buffer (RNeasy^®^ Mini Kit by Qiagen) and immediately stored at -80°C until further processing. RNA extraction was performed upon thawing, using the RNeasy^®^ Mini Kit by Qiagen according to the manufacturer’s protocol. RNA concentrations were measured using the NanoDrop^TM^ One (Thermo Scientific). cDNA was synthesized using the ReadyScript^TM^ cDNA Synthesis Mix protocol (Sigma-Aldrich^®^) and qRT-PCR machine (Eppendorf Mastercycler). qPCR was performed on 1:5 diluted cDNA samples with the StepOnePlus^TM^ Real-Time PCR System (Applied Biosystems) with StepOne^TM^ Software (version 2.3) (Applied Biosystems) (Supplementary Table S6 for master mix composition and instrument settings) (Data S11).

#### Proteomic analysis

For this analysis, the co-incubation period was prolonged to 24 h, the bacterial load lowered to 1 × 10^6^ CFU and the number of biological replicates increased to 6 for the VK2/E6E7 monolayer (and 3 for 3D transwell model). After co-incubation, co-culture supernatant was harvested by centrifugation (10 min, 2000g) and stored at -80°C until further analysis with Olink® Target 96 inflammation (Thermofisher Scientific). Analogously, the 3D transwell epithelial model was co-incubated at the apical side and supernatant was harvested at both apical and basolateral sides. The data on singular experiments is available in Data S12.

### Statistical analyses

#### Metabolomics results, Microbiome and FFQ’s

The semi-targeted feature table was merged with the 16S amplicon sequencing dataset, where the associations of the ion count of the vitamins with alpha diversity, beta diversity and genus level abundances were tested, respectively. In addition, associations between ion counts of the vitamins and vitamin-related data from the questionnaires, including vitamin intake from supplements, vegetable consumption, fermentation products, probiotics, or other possible endogenous sources, were tested. All four analyses were performed using the multidiffabundance R package (version 0.0.1) using linear models to model the shannon and Bray-Curtis index for alpha and beta diversity respectively and an ensemble of differential abundance tests in both batches separately due to the batch effect.

A linear model, corrected for age, BMI, previous pregnancies and contraception use, was used to look for associations between ion counts and vitamin-related data. Multiple testing correction using the Benjamini-Hochberg (BH) principle was performed within batches. Unsupervised clustering using Latent Dirichlet Allocation (LDA) topic modeling with k=9 was performed (using the topicmodels R package, version 0.2.17), and the resulting association of the vitamins with the resulting topics was tested using linear models with multiple testing correction (BH).

Statistical analysis and plotting were performed in R 4.4.1 using RStudio/2023.06.1. The statistical significance of molecular features in the SNP and transwell assay experiments were assessed with a t-test (t.test function in R), and p-values were FDR-corrected for multiple hypothesis testing using the Benjamini-Hochberg procedure (p.adjust function in R with ‘fdr’ parameter). Significant metabolites were selected when their intensity had a corrected p-value < 0.05.

#### Immune results

Data were initially processed using Microsoft Excel. Graphs and statistical analyses were performed in GraphPad Prism (version 9.5.1). Depending on the number of independent variables and the distribution of the data, assessed using the Shapiro–Wilk test, either a t-test or ANOVA (one-way, two-way, or mixed) was applied. For data not meeting assumptions of normality, non-parametric alternatives (Wilcoxon test or Kruskal–Wallis test) were used. Where applicable, multiple comparisons were corrected using Dunnett’s, Tukey’s, or Šidák’s post hoc tests. For qPCR analysis, simple linear regression was applied to calibration data in Microsoft Excel. Statistical significance was defined and indicated with p-value < 0.05 = *, p-value < 0.01 = **, p-value < 0.001 = ***).

**Data S1. (separate file)**

**List of metabolites and their analytical information:** Metabolite identification was done by level-1.

**Data S1. (separate file)**

**Clustermaps:** Statistical analysis of the associations between the B-vitamer ion counts and individual vaginal microbiome taxa have been performed using seven pipelines (ALDeX2, ANCOM-BC, DESeq2, limma, Maaslin2, Corncob, lmclr and a linear regression on the CLR-transformed abundance data). Only the ALDeX2-results for timepoint 1results shown in Fig. 1, the additional heatmaps can be found in this file.

**Data S3. (separate file)**

**Publically available genomes:** List of the ENA accession numbers of the publically available genomes used in Figure 2A.

**Data S4. (separate file)**

**List of microbial strains:** All strains used, together with their isolation source (mostly vaginal sample) and whole genome sequence (ENA IDs).

**Data S5. (separate file)**

**Riboflavin production capacity:** An overview of the 128 WGS vaginal isolates, together with the the presence/absence of riboflavin synthesis genes, autofluorescence measurements and HPLC quantification is given in this separate file.

**Data S6. (separate file)**

**Metabolite statistics of contig:**

**Data S7. (separate file)**

**Metabolite correlation analysis for the bacterial cultures: correlation matrix:**

**Data S8. (separate file)**

**Metabolite correlation analysis for the bacterial cultures: p-value matrix:**

**Data S9. (separate file)**

**Radioactive riboflavin transport:** Radioactive counts (Cpm) for VK2/E6E7 monolayers and 3D vaginal epithelial model.

**Data S10. (separate file)**

**Flow Cytometry:** Collection of low cytometric reports.

**Data S11. (separate file)**

**qPCR files:** This file includes raw Ct values and fold-change calculations (2^-ΔΔCt^).

**Data S12. (separate file)**

**Olink:** The raw data (npx values) can be find in this table. For every target marker it is indicated whether it surpasses the LOD threshold, which is defined as npx > LOD in 50% of replicates of at least ne condition.

---

**Fig. S1.**
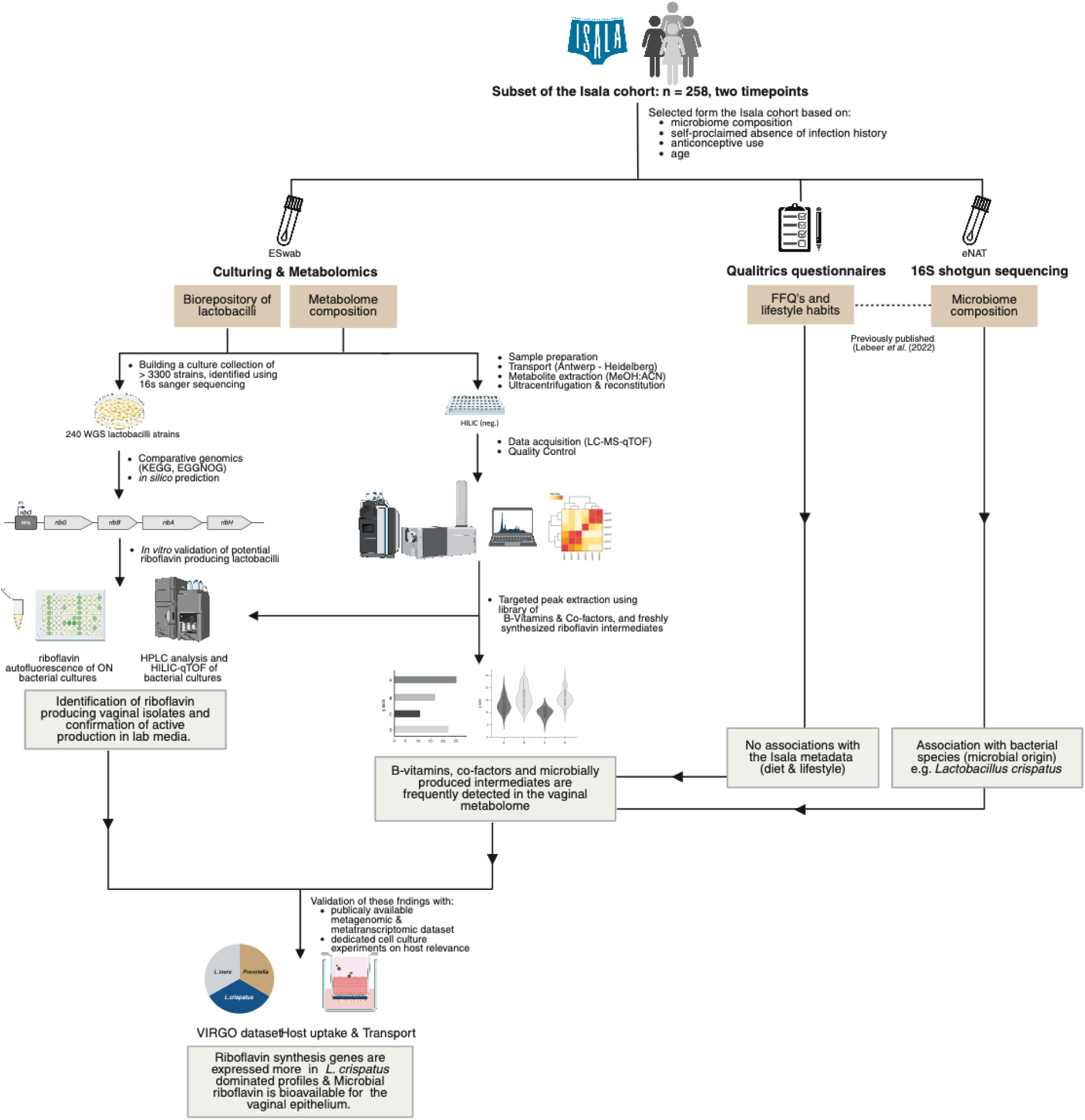
Graphical abstract: The samples and metadata of an Isala sub-cohort (n = 258) is leveraged in this work. The metadata (FFQs and lifestyle habits) and vaginal microbiome composition (16s data), previously processed in Lebeer *et al.* (2022) (*6*) is linked to the targeted metabolomic data on vitamers and co-factors (semi-quantification using ion-counts). The associations with vitamin B_2_ were confirmed with the in-house biobank of vaginal Isala isolates (*in silico* and *in vitro*), as well as externally validated using the VIRGO database (*54*). The functional relevance of vitamin B_2_ production by *Lactobacillaceae* in the vagina is investigated using a 3D vaginal epithelial model.

**Fig. S2.**
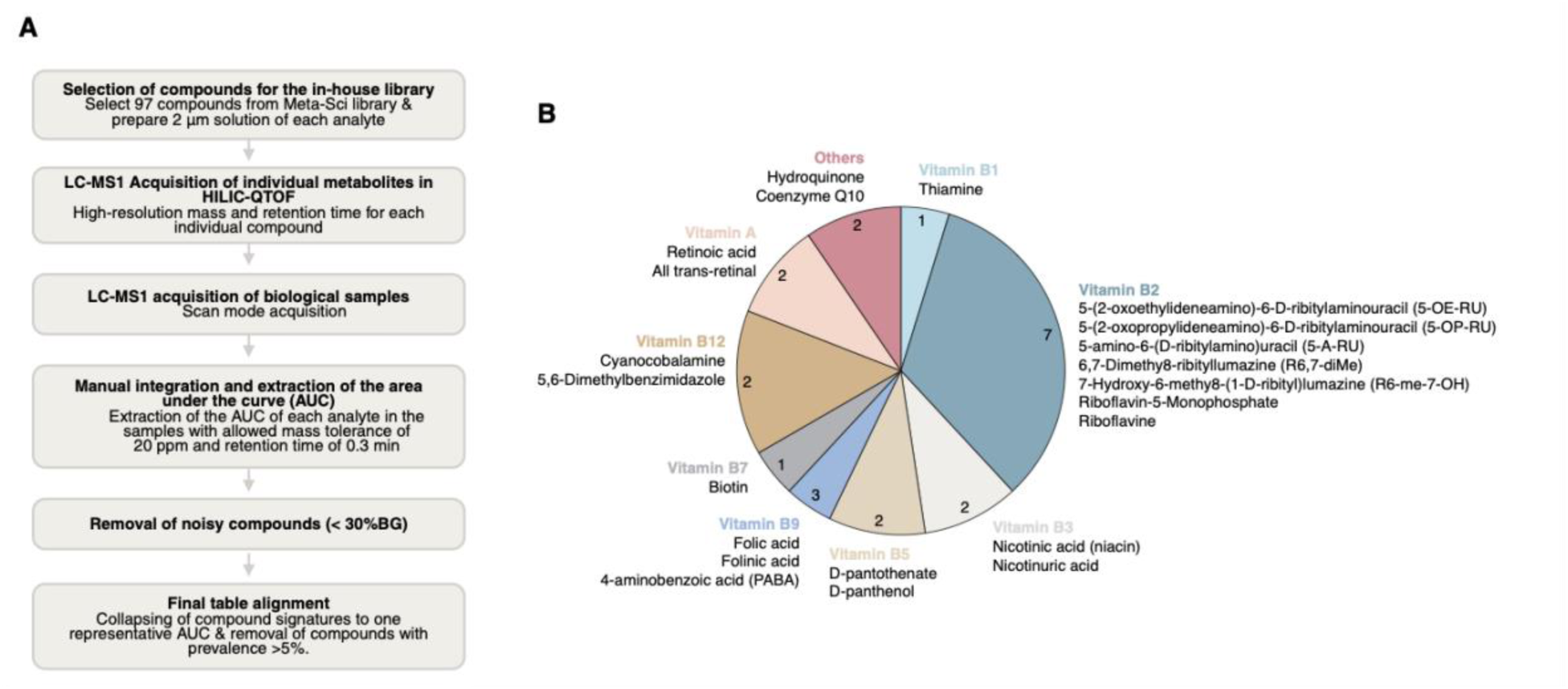
Method of generating a targeted list: **(A)** Flowchart describing the method behind building a HILIC neg. targeted library of compounds related to host-microbe interactions, specifically focusing on vitamers and other co-factors. In this flowchart, the term ‘biological sample’ refers to all types of samples used in this study, including vaginal swabs, bacterial cultures, cellular lysates as well as cellular and bacterial supernatant. **(B)** Pie chart representing the vitamers and co-factors that could be successfully detected in any of the biological samples and used in downstream statistical analyses.

**Fig. S3.**
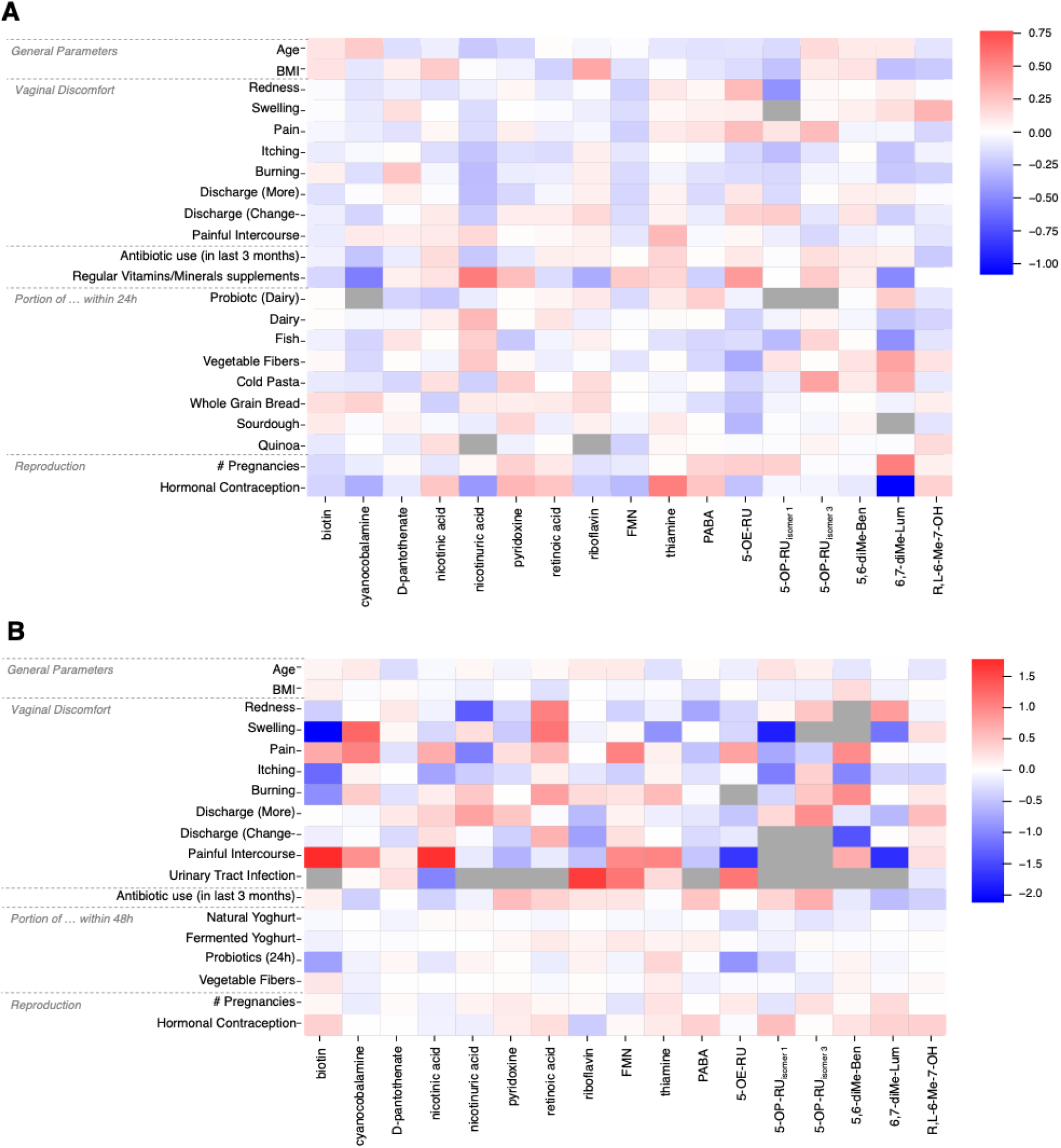
Metadata associations: Statistical associations of vitamer ion counts with the Isala lifestyle factors that have previously been reported as ‘having the largest impact on the vaginal microbiome composition’, including age, BMI, number of previous pregnancies and contraceptive method for **A)** sampling timepoint 1 and **B)** timepoint 2 (*6*). In addition, vaginal complaints and vitamin-related FFQ’s are included as well. No statistically significant associations were found, hence no numbers are included within the cells of the heatmap. Effect sizes are represented by the colored gradients. Gray cells indicate those associations that could not be calculated due to minimal variability in the data.

**Fig. S4.**
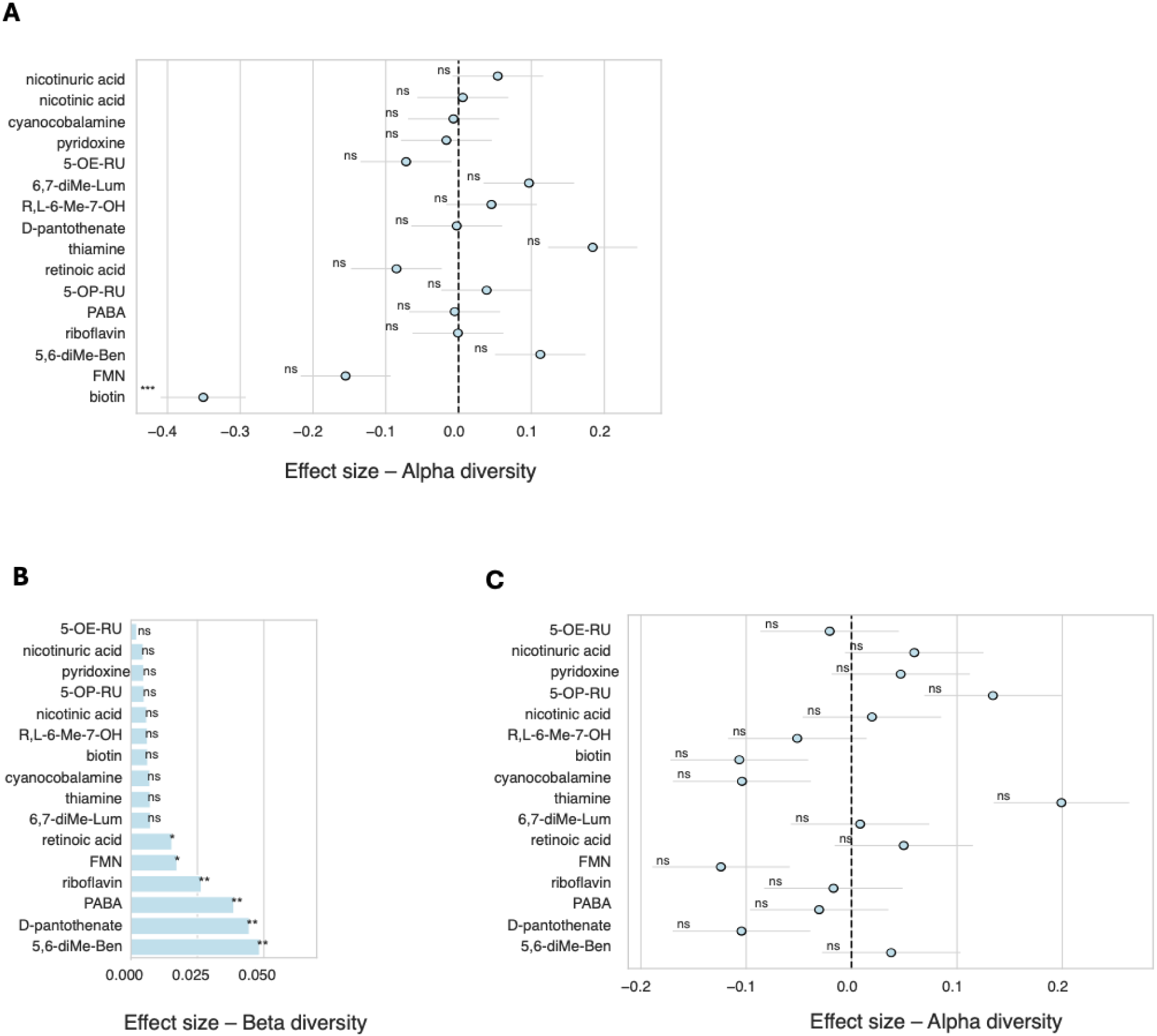
Associations between diversity and vitamer levels: Statistical associations of the vitamer ion counts with Shannon α-diversity (linear model) in **(A)** for timepoint 1 and **(C)** for timepoint 2, and bray-curtis β-diversity (permanova) **(B)** for timepoint 2. Significance (p-value < 0.05 = *, p-value < 0.01 = **, p-value < 0.001 = ***) (*6*).

**Fig. S5.**
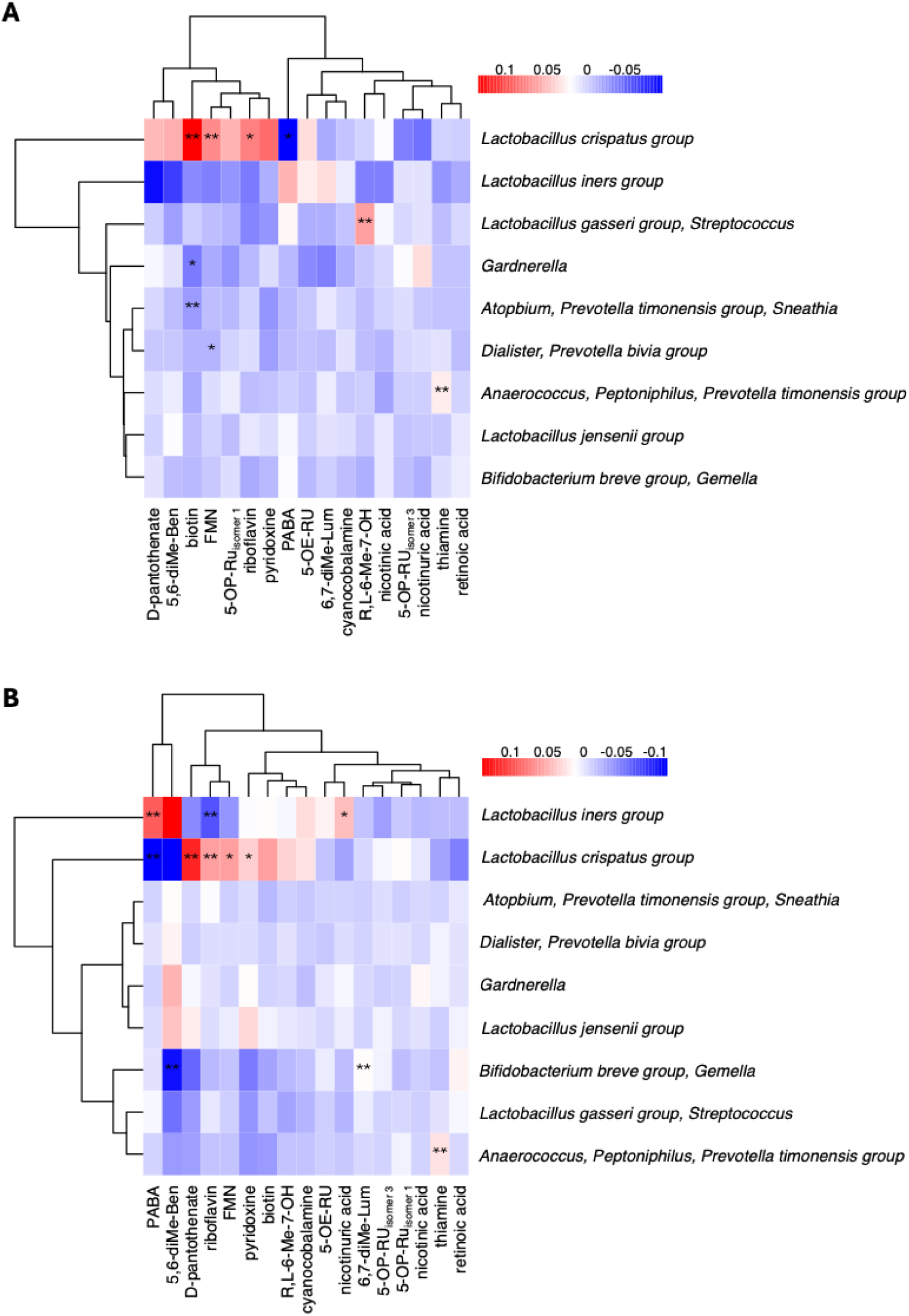
Associations between topics and vitamer levels: Statistical associations of each 9 topics with the vitamer ion counts for **(A)** sampling timepoint 1 and **(B)** timepoint 2 using a linear model (modelled by the scaled normalized metabolites). Significances (p-value < 0.05 = *, p-value < 0.01 = **, p-value < 0.001 = ***) are adjusted for multiple testing using the FDR method.

**Fig. S6.**
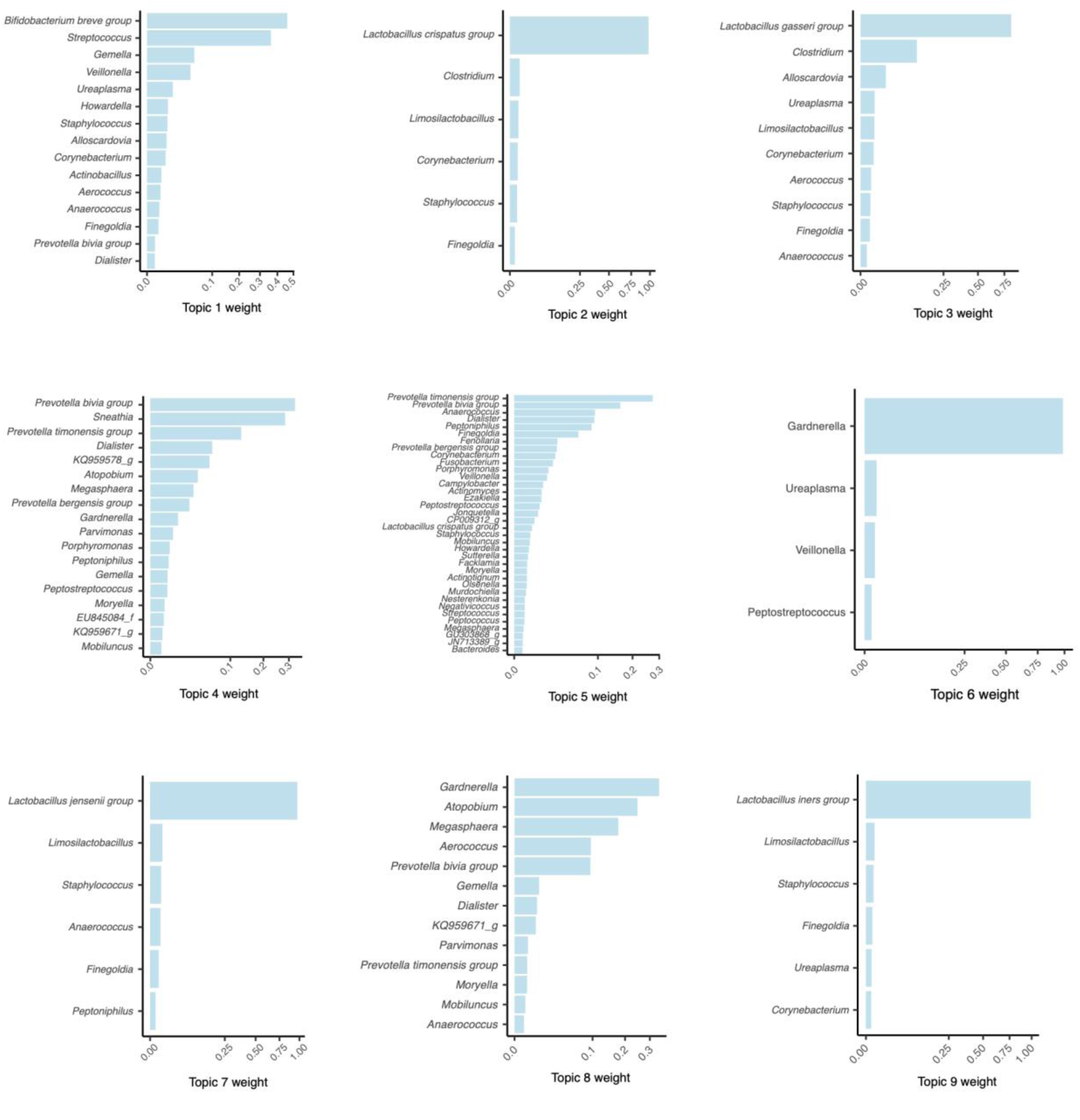
Topic modelling: Each microbiome is treated as a mixture of topics which contain a mixture of taxa (genera in this case) which are allowed to overlap to an extent, whereas with clustering a taxa either belongs to one group or another. The following plots show the largest probabilities of each taxa belonging to the 9 topics obtained using LDA (*55*).

**Fig. S7.**
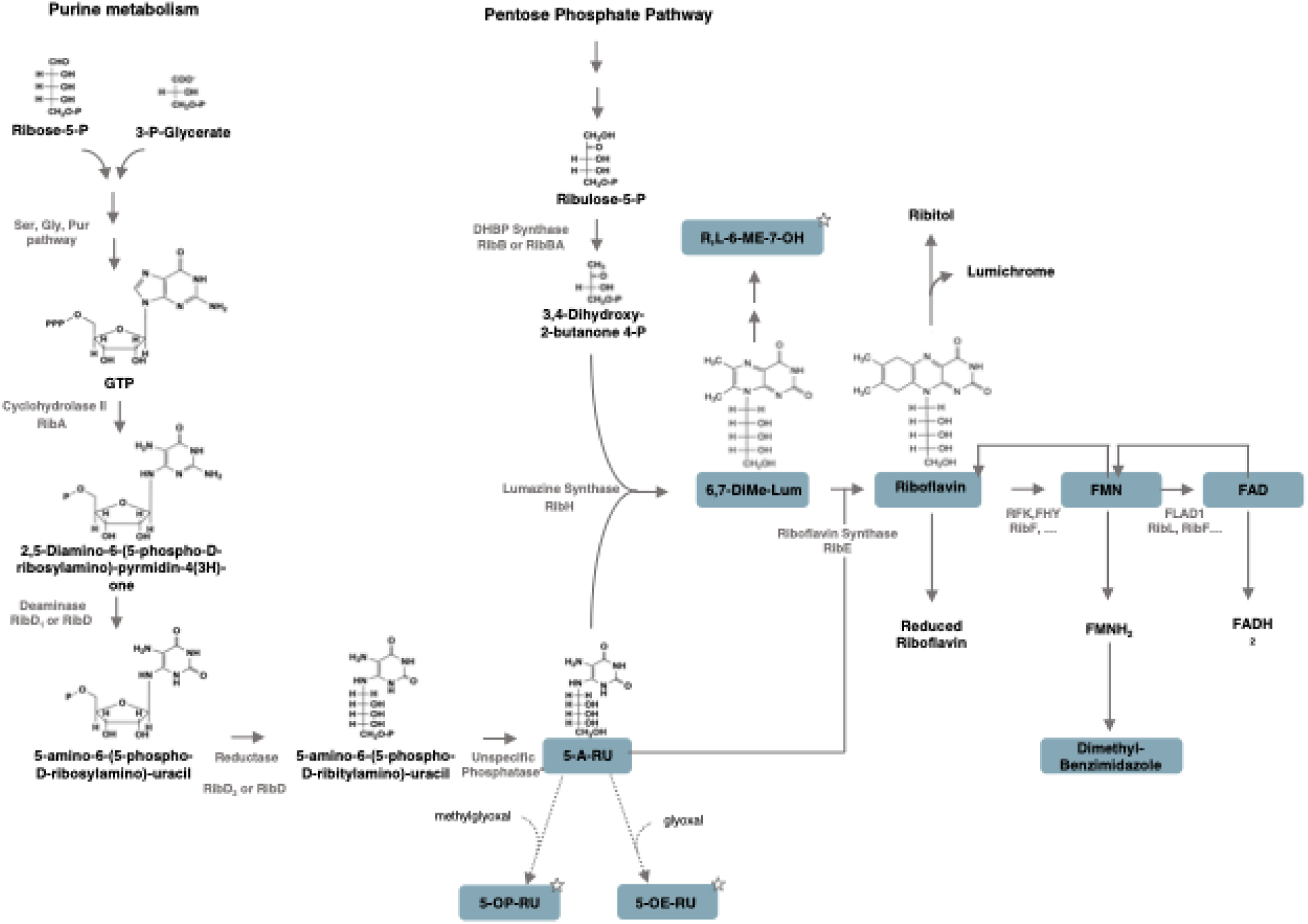
Microbial riboflavin biosynthesis pathway: Riboflavin synthesis originates from purine metabolism and the pentose phosphate pathway. Enzymatic reactions are indicated by solid arrows, non-enzymatic steps by dashed arrows. Pathway intermediates with MAIT cell-activating potential are marked with a star.

**Fig. S8.**
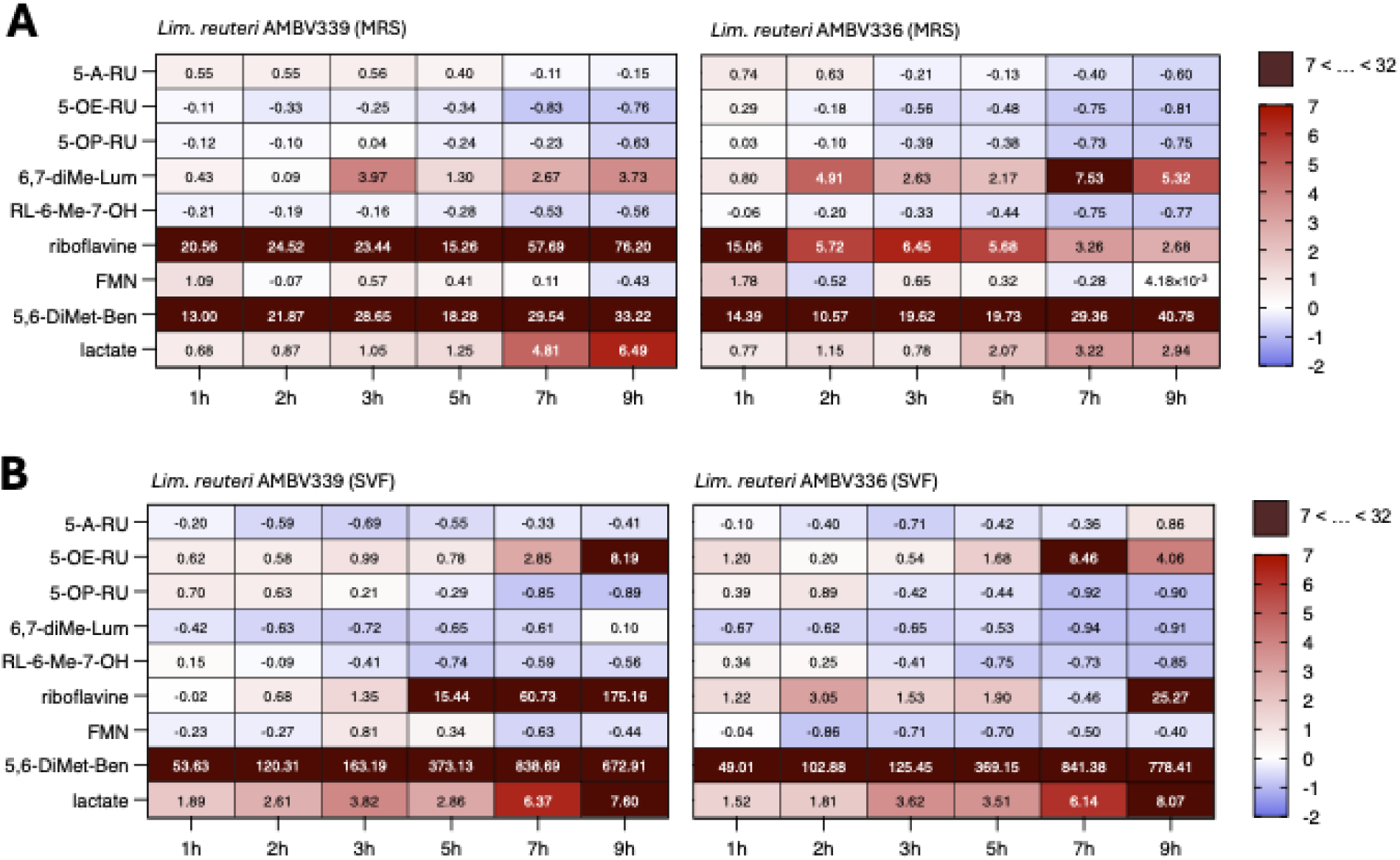
Quantification of riboflavin and its intermediates: Quantification (HILIC (neg.)) of riboflavin and its (immunomodulatory) pathway derivatives during *Lim. reuteri* AMBV339 and *Lim. reuteri* AMBV336 growth in lab media (MRS and SVF) over the course of 9h.

**Fig. S9.**
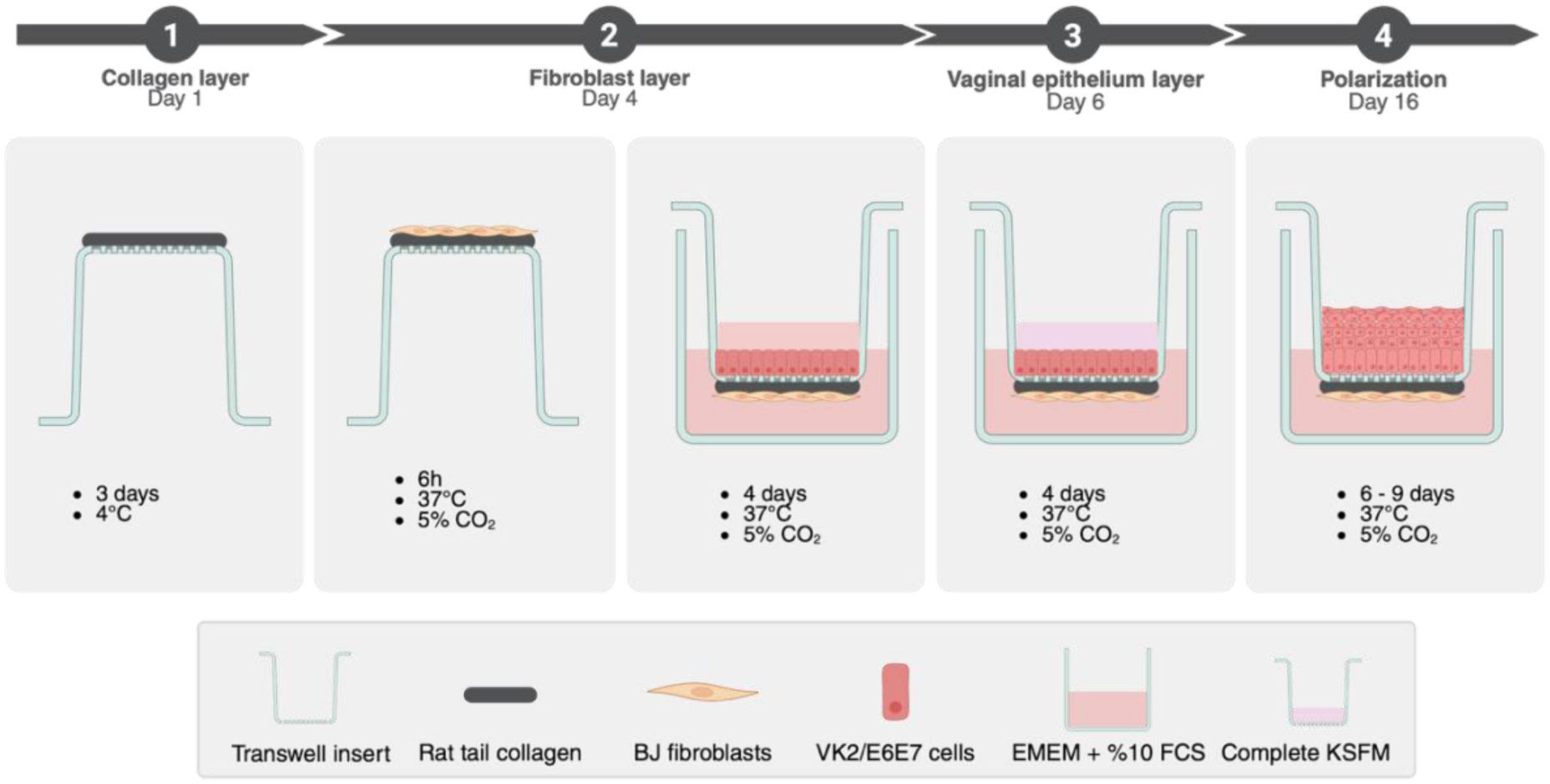
Experimental setup of the 3D transwell model of the vaginal epithelium. (*49*): On day 1, the collagen layer is seeded on the basolateral side of the transwell, followed by addition of a BJ fibroblast layer on day 4. On day 6, a monolayer of VK2 E6/E7 cells is added, which should polarize to a multilayer after the air-liquid interface creation on day 8.

**Fig. S10.**
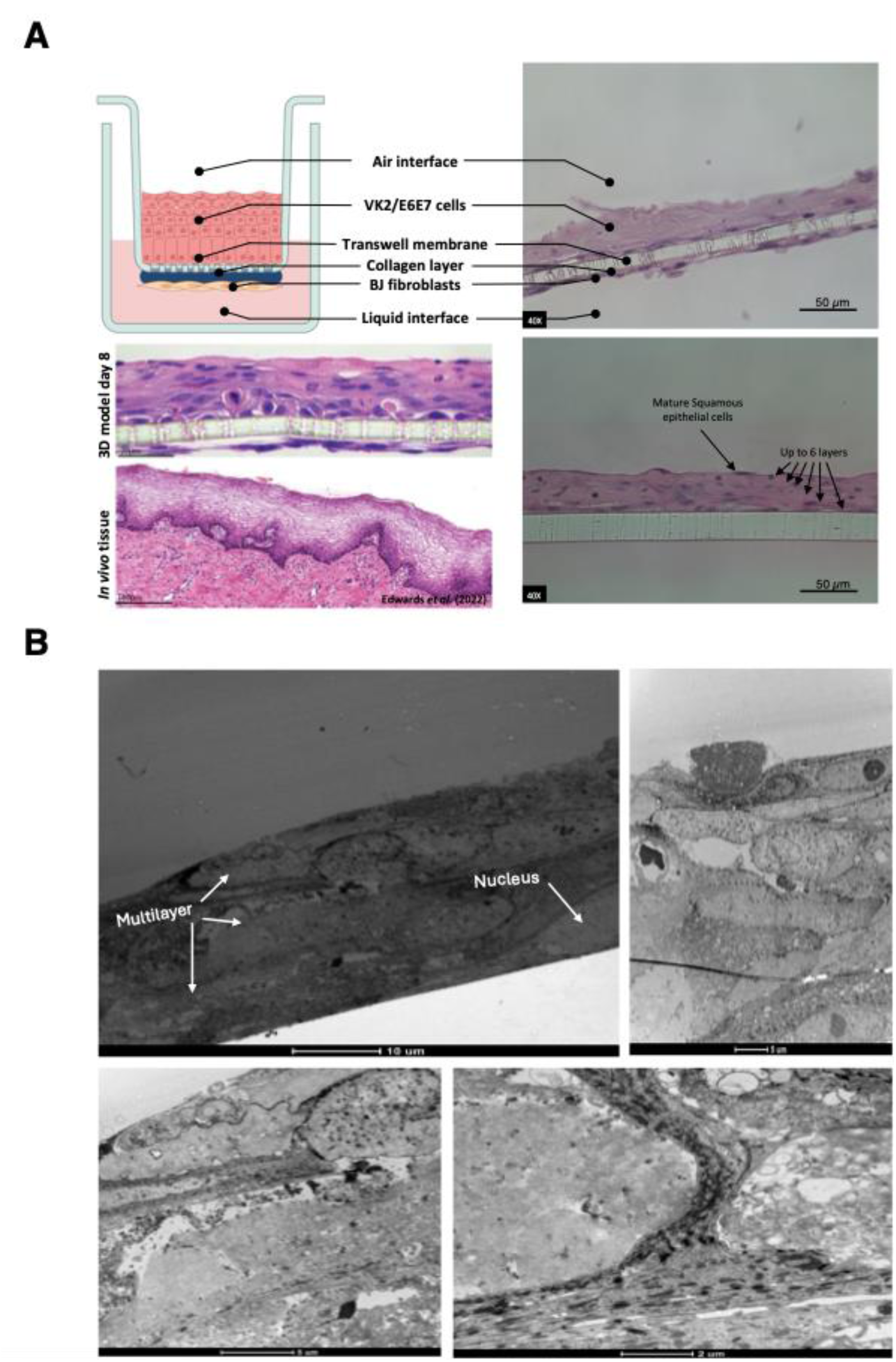
Functionalization of the 3D vaginal epithelium model: **(A) & (B)** Validation of epithelial stratification in the 3D vaginal model of Edwards *et al.* (2022) (*49*) using Light Microscopy **(A)** and Transmission Electron Microscopy **(B)** on insert cross-sections. Transmission electron microscopy images highlight the multiple layers and tight barrier.

**Fig. S11.**
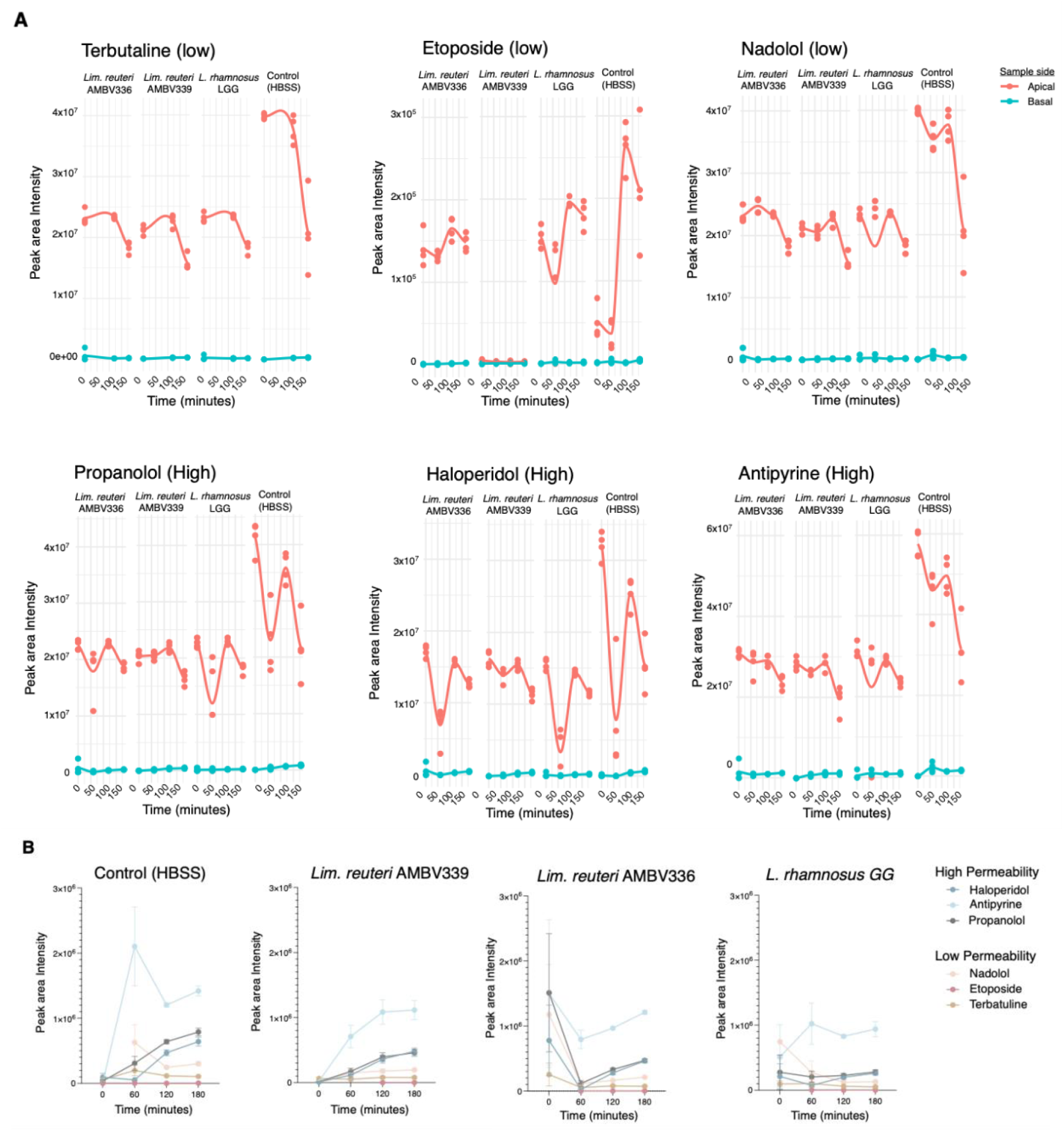
Assessment of epithelial barrier integrity in the 3D model: The barrier integrity of the vaginal transwell model is evaluated by following apical-to-basolateral transport of a mix of compounds with variable permeability over time (previously established for the gut epithelium (*21*) **(A)** For all compounds, both apical (orange) and basolateral (blue) raw ion counts are presented (6 biological replicates). **(B)** Ion counts for the basolateral side are shown as mean ± SD. These results highlight the strong barrier of the 3D model (ca. 5 cell layers): over time the highly permeable molecules (haloperidol, antipyrine and propoanolol) seem to increase, while the low permeable ones (nadolol, etoposide and terbatuline) do not.

**Fig. S12.**
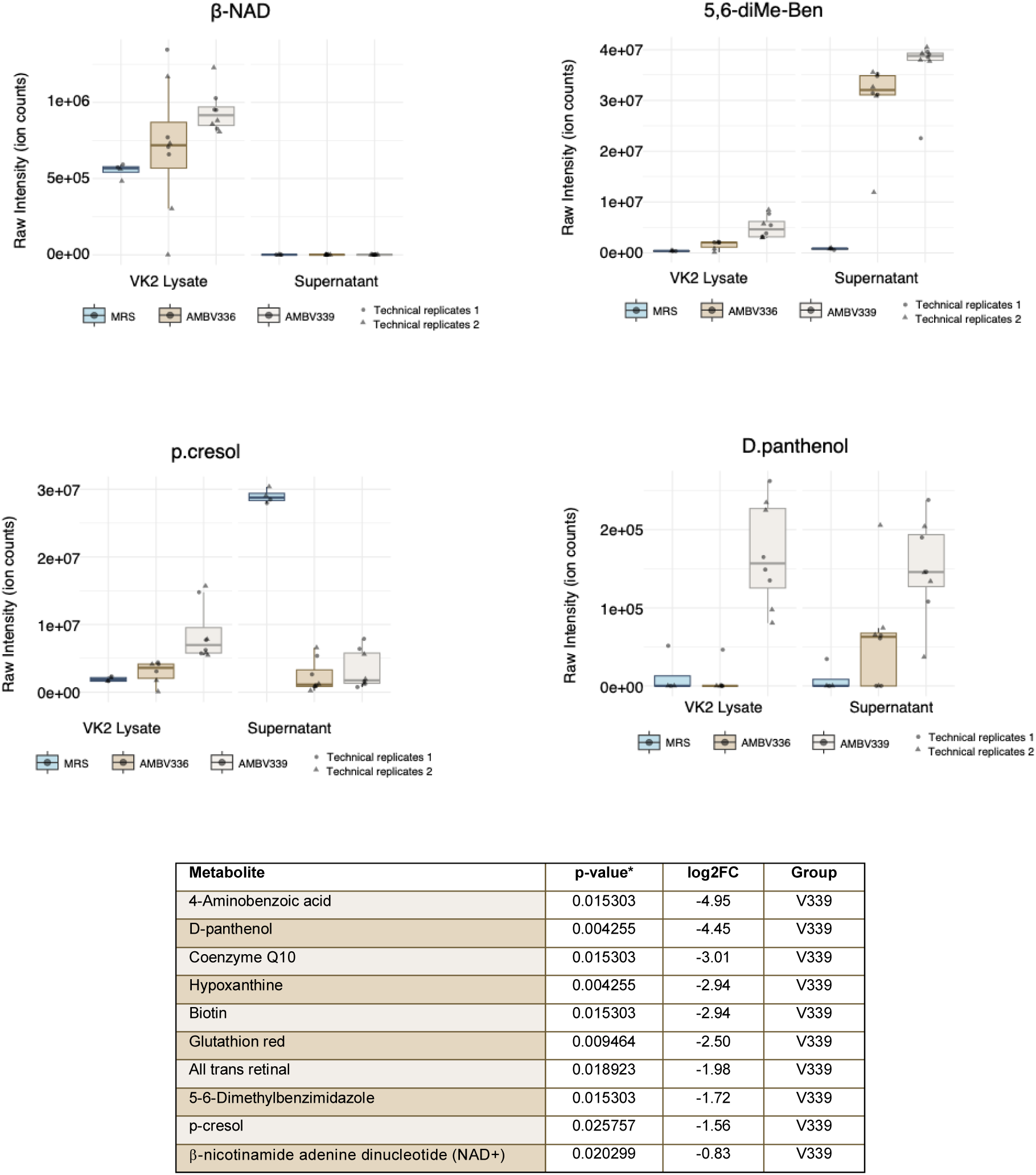
Quantification of redox-related molecules in VK2/E6E7 lysates after co-incubation with *Lim. reuteri* AMBV339 and *Lim. reuteri* AMBV336: VK2E6E7 monolayers were exposed for 3 h to 10% ON bacterial supernatant of riboflavin producer *Lim. reuteri* AMBV336 and its genetic variant *Lim. reuteri* AMBV339, which overproduces riboflavin. Vitamins, co-factors and other compounds related to cellular (redox) metabolism were quantified (with HILIC neg.) in the culture medium (10% supernatant) and in the VK2/E6E7 cellular lysates after bacterial co-incubation. The compounds significantly impacted (P_FDRcor_ < 0.05 and Log2FC > 0.8) by exposure to *Lim. reuteri* AMBV339, but not *Lim. reuteri* AMBV336, are reported above.

**Fig. S13.**
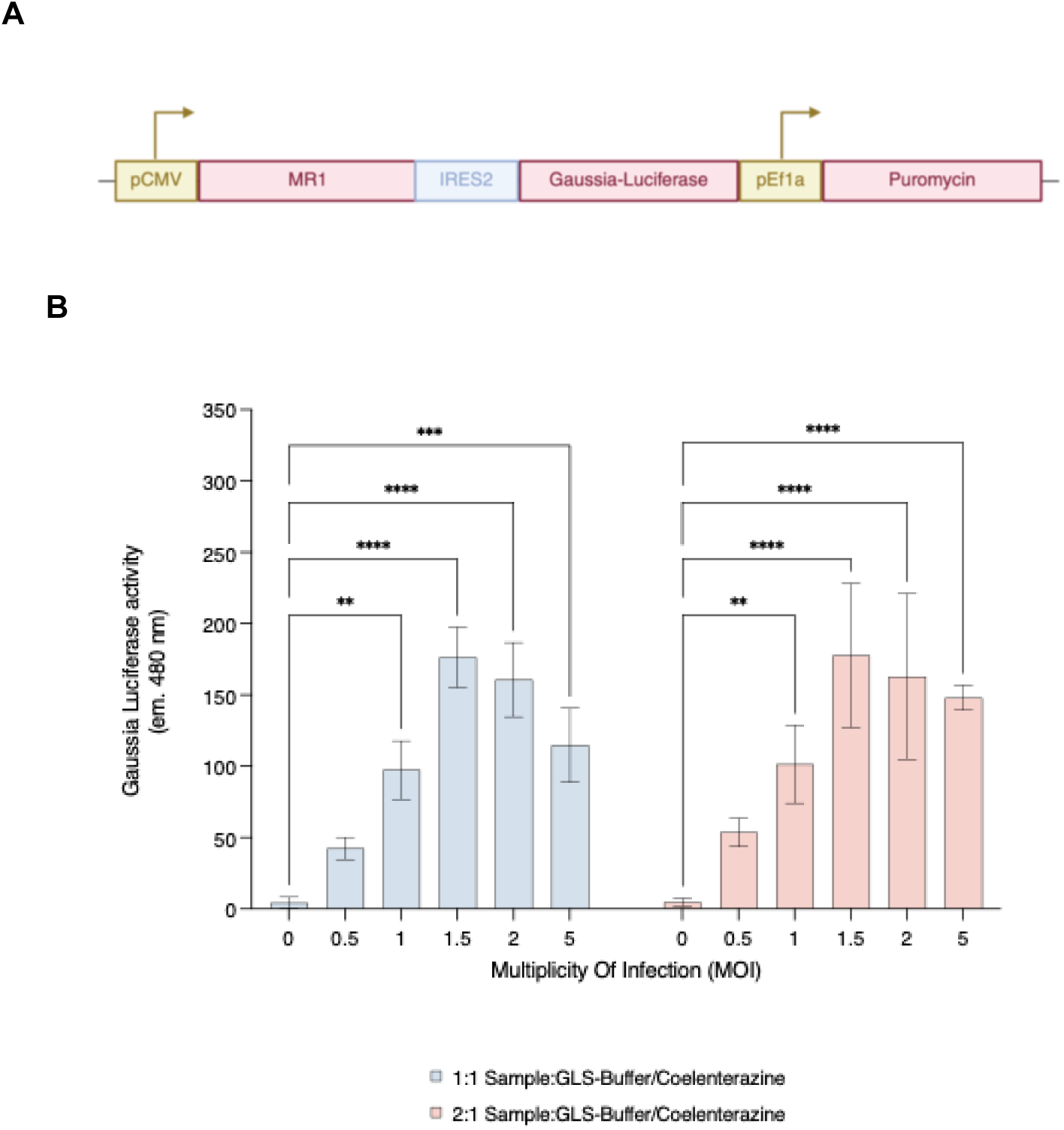
Generation of the VK2/E6E7.hMR1 cell line using lentiviral transduction: VK2/E6E7 cells were transduced with the lentiviral construct in **(A)** form GeneCopoiea^TM^. The Ires2 (internal ribosome entry site) sequence leads to co-expression of MR1 (NM_001531.2) (Major Histocompatibility Complex (MHC) related protein-1) and Gaussia Luciferase under the control of the constitutive pCMV (cytomegalovirus) promotor. The puromycin selection cassette is under control of the constitutive pEf1a (eukaryotic translation factor 1) promotor. **(B)** Transduction was performed at a viral load or Multiplicity Of Infection (MOI) of 0, 0.5, 1, 1.5, 2 and 5. Once expressed, the GLuc was secreted in the supernatant (‘sample’) and produced luminescence in presence of different ratios of the substrate coelenterazine and buffer. Data depicted as mean ± SD. Statistics were performed on technical replicates with a two-way ANOVA test, followed by a Dunnett’s multiple comparisons test compared to the blank (MOI = 0), with p-value < 0.05 = *, p-value < 0.01 = **, p-value < 0.001 = ***).

**Fig. S14.**
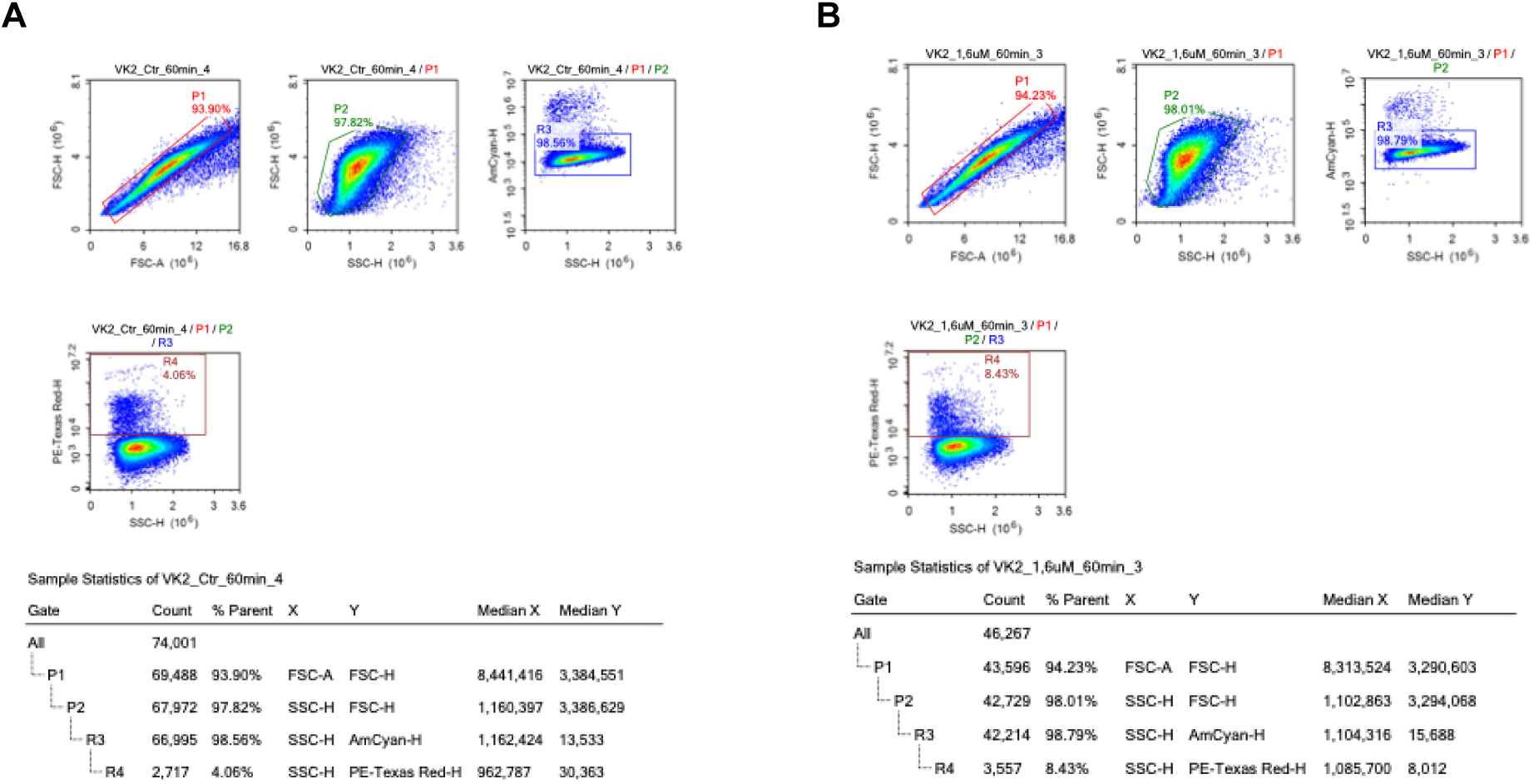
Flow Cytometry gating strategy for MR1 positive K2/E6E7.hMR1 cells: Density plots of VK2/E6E7.hMR1 cells co-incubated with **A)** KSFM and **B)** 5-OP-RU (1.6 uM) in KSFM. In both **A)** and **B)**, P1.) FSC-A vs. FSC-H: exclusion of doublets P2.) SSC-H vs. FSCH: Identification of homogenous cell population R3.) SSC-H vs. Amcyan-H: Exclusion of dead cells R4.) SSC-H vs. PE-Texas Red: MR1 surface expression (FSC = forward scatter, SSC = Side scatter, A = Area, H = Height, Count = number of selected cells, %Parent percentage of cells within a gate to the cells from the parent (previous) gate).

**Fig. S15.**
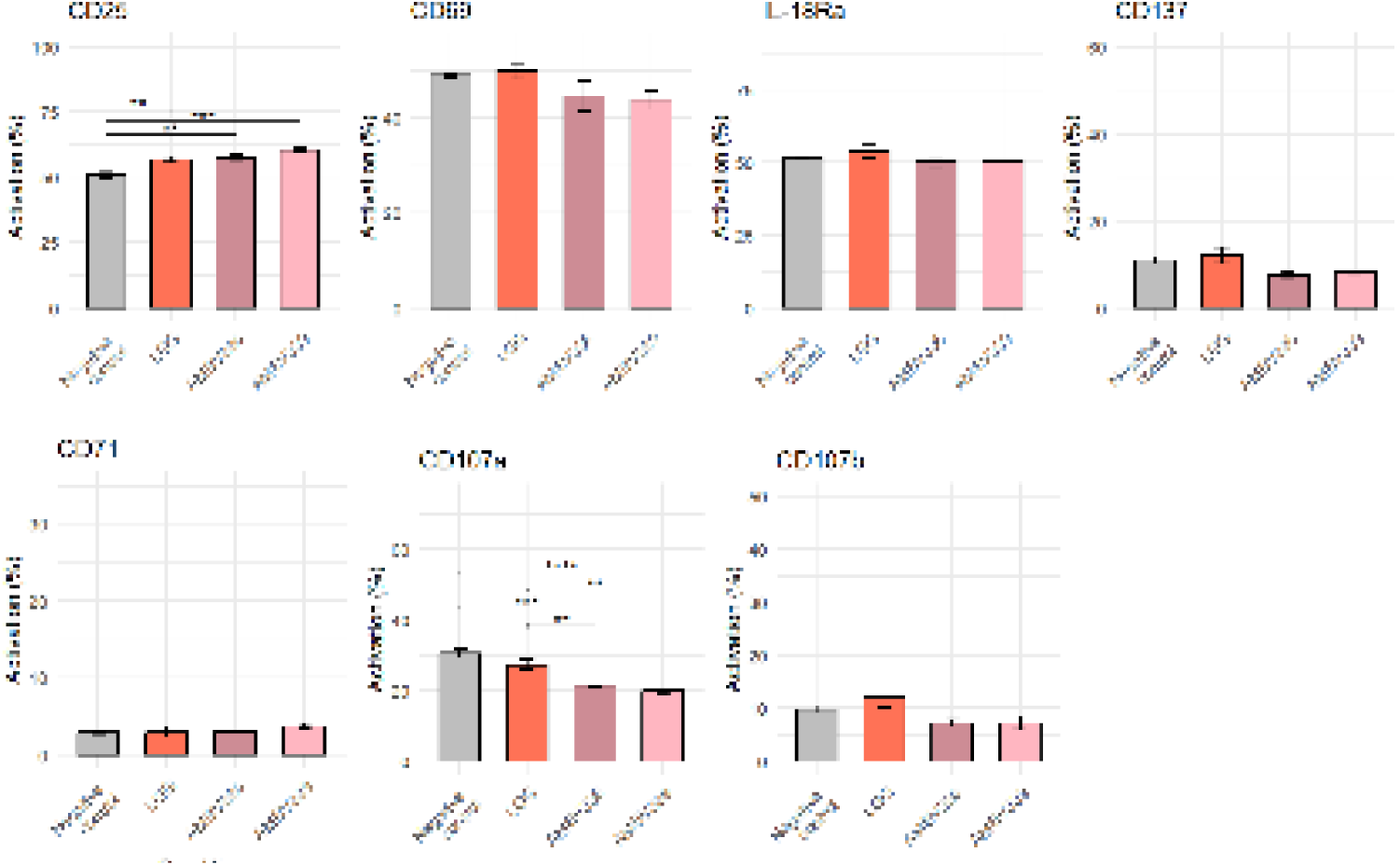
Flow cytometric analysis of MAIT cell activation and proliferation markers after exposure to the cell-extract of (riboflavin-producing) microbes: PBMCs were incubated for 3h with 1:10 sonicated cell extract:KSFM. MAIT cells were identified (CD3 and TCRVa7.2) and expression of various surface markers (CD25, CD69, IL-18Ra, CD137, CD71) and intracellular markers (CD107a, CD107b) was assessed. The impact of exposure to the riboflavin producing *Lim. reuteri* AMBV336, overproducer *Lim. reuteri* AMBV339 and non-producer *L. rhamnosus* GG was statistically compared with the negative control condition (regular KSFM) using a Kruskal-Wallis followed by a Dunnett’s multiple comparisons test with p-value < 0.05 = *, p-value < 0.01 = **, p-value < 0.001 = ***).

**Fig. S16.**
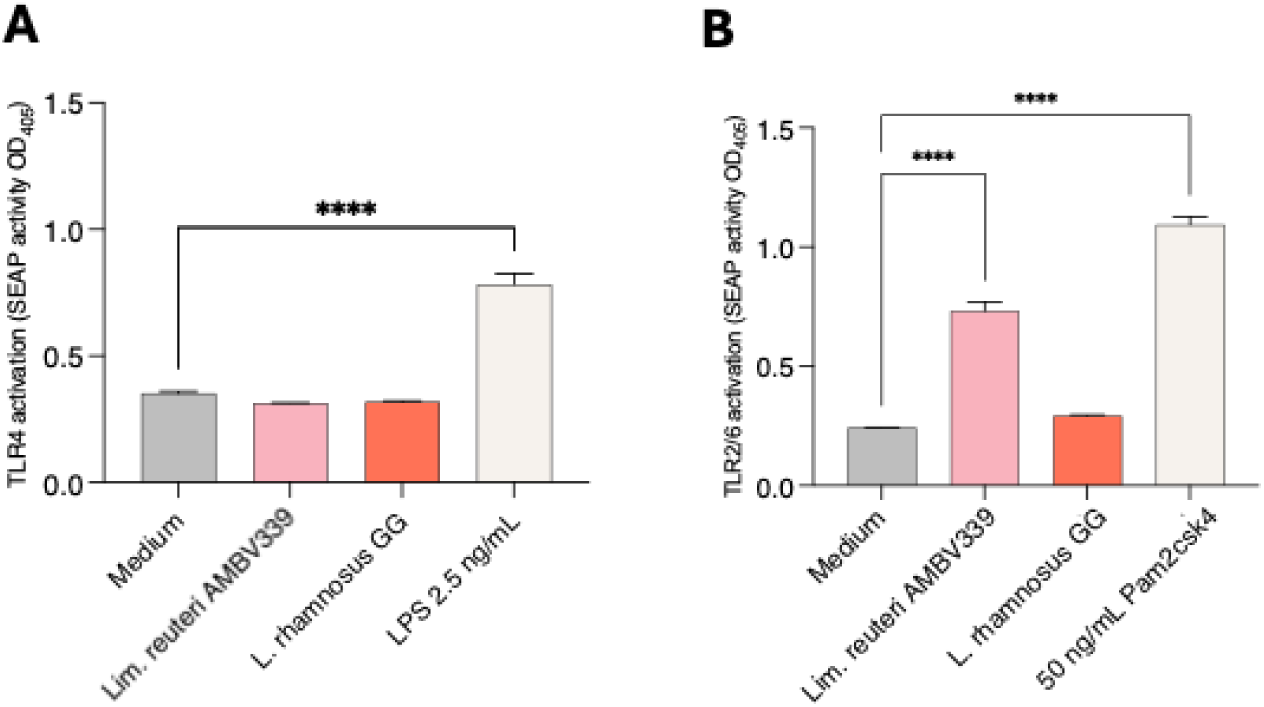
TLR2/6 and TLR4 activation by *Lim. reuteri* AMBV339 and *L. rhamnosus* GG: UV-inactivated bacteria were co-incubated with HEKreporter cell lines (in triplicate) for assessing **A)** TLR4 activation and **B)** TLR2/6 activation. Data is depicted as mean ± SD. Statistical significance is assessed with one-way ANOVA, and indicated with p-value < 0.05 = *, p-value < 0.01 = **, p-value < 0.001 = ***.

**Fig. S17.**
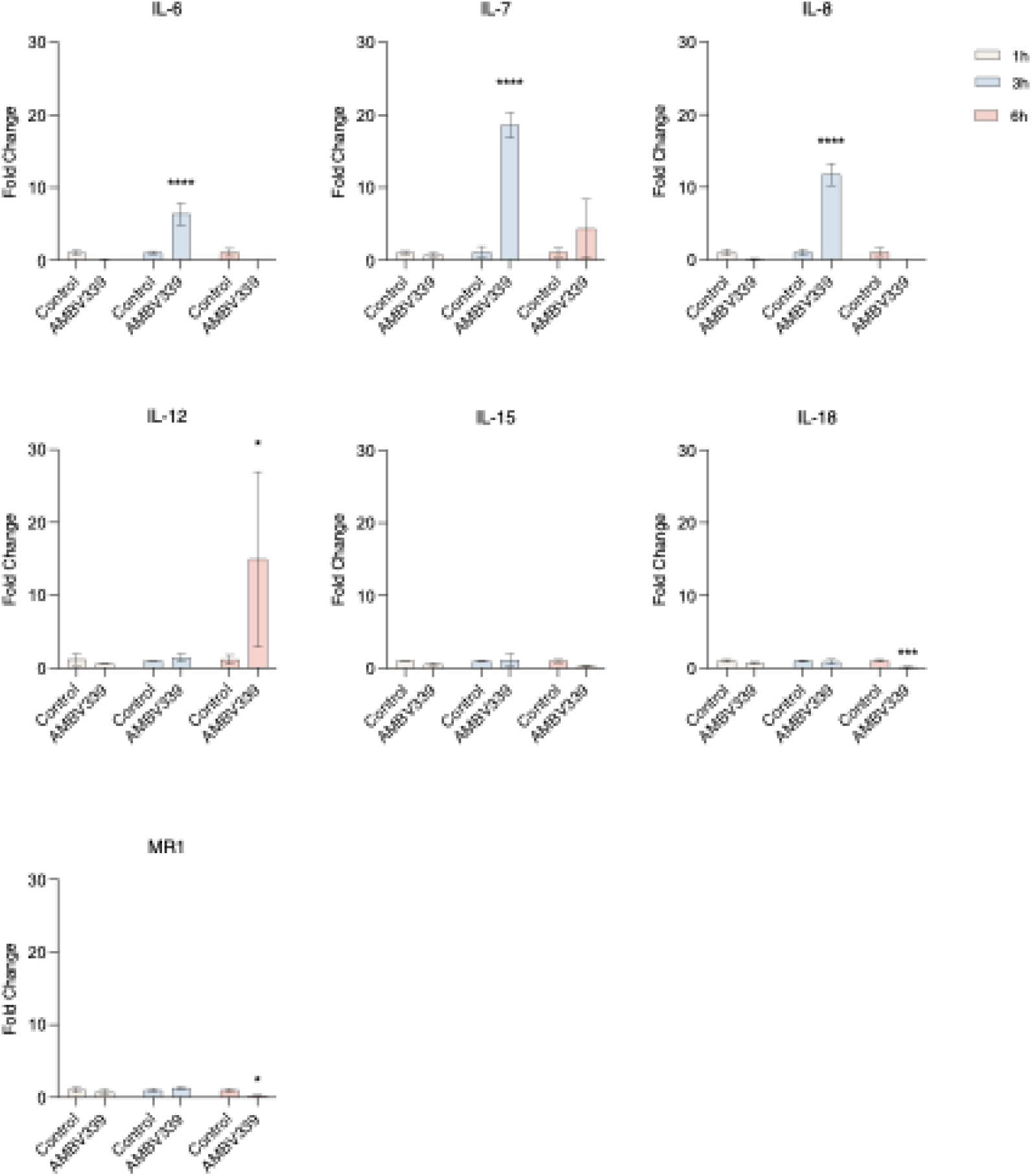
Optimization of the co-incubation time (1h, 3h or 6h) of VK2/E6E7 cells with *L. reuteri* AMBV339 for qPCR: Cytokine and MR1 transcription levels induced by *L. reuteri* AMBV339 are presented as fold changes (Mean^2-ΔΔ(mean^ ^Ct)^) compared to the expression level at control condition (KSFM + 30% MRS) and normalized for housekeeping gene transcription of GAPDH and CYC1 (Statistics: Two-way ANOVA, Sidak’s multiple comparisons correction, p-value < 0.05 = *, p-value < 0.01 = **, p-value < 0.001 = ***).

**Fig. S18.**
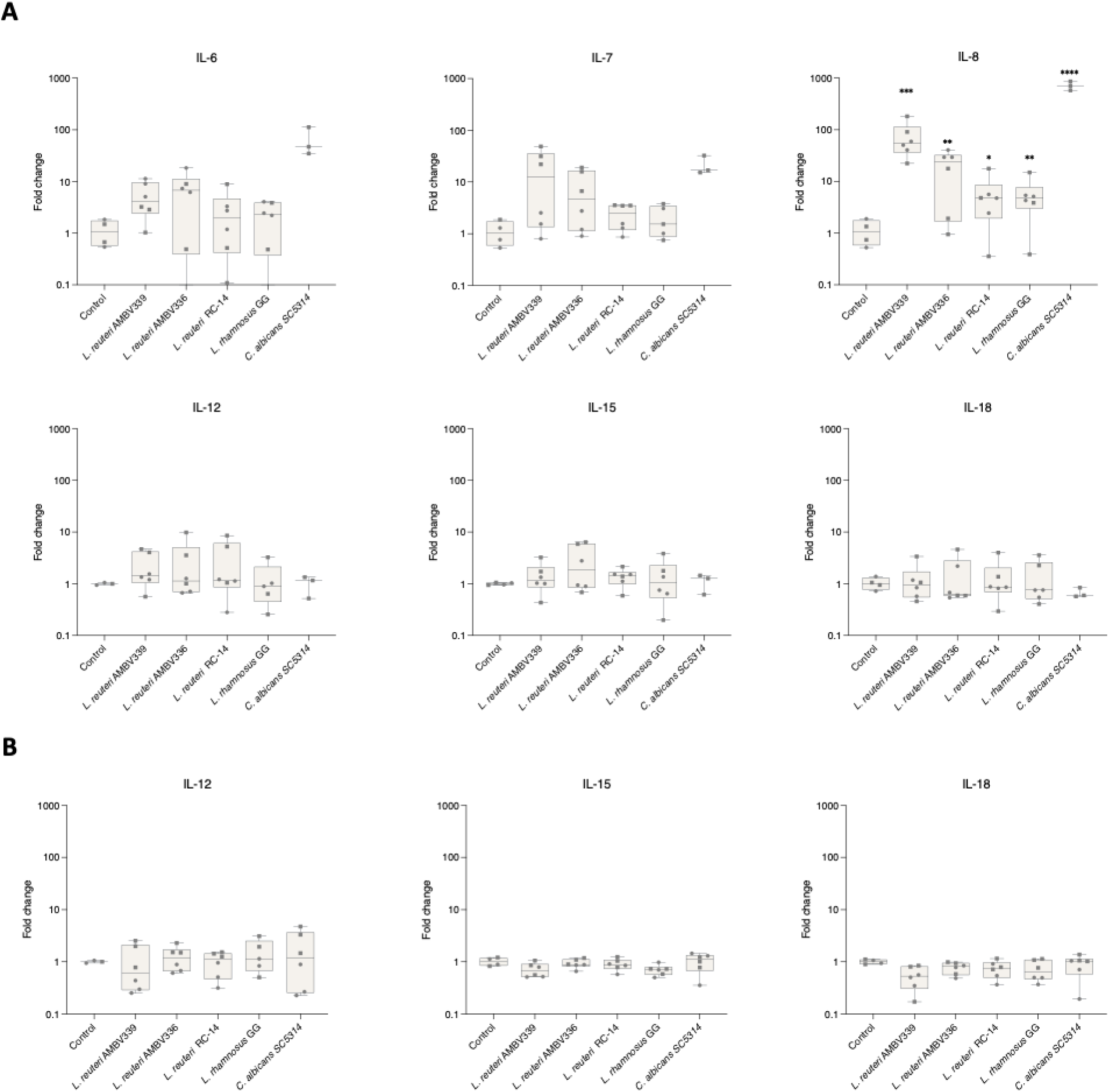
Optimization of the co-incubation time (1h, 3h or 6h) of VK2/E6E7 cells with *L. reuteri* AMBV339 for qPCR: Cytokine transcription levels induced by commensals *L. reuteri* AMBV339, *L. reuteri* AMBV336, *L. reuteri* RC14, *L. rhamnosus* GG and pathobiont *C. albicans* SC5314 are presented as fold changes (Mean^2-ΔΔ(mean^ ^Ct)^) compared to the expression level at control condition (KSFM) and normalized for housekeeping gene transcription of GAPDH and CYC1. In **(A)** VK2/E6E7 monolayers were co-incubated with 1 × 10^7^ CFU/mL, while in **(B)** the monolayers were incubated with 10% ON supernatant in KSFM. Biological replicates (2 x n = 3) originate from two VK2/E6E7 cell culture plates (plate 1 = circular, plate 2 = squared). (Statistics: Two-way ANOVA, Sidak’s multiple comparisons correction, p-value < 0.05 = *, p-value < 0.01 = **, p-value < 0.001 = ***.)

**Fig. S19.**
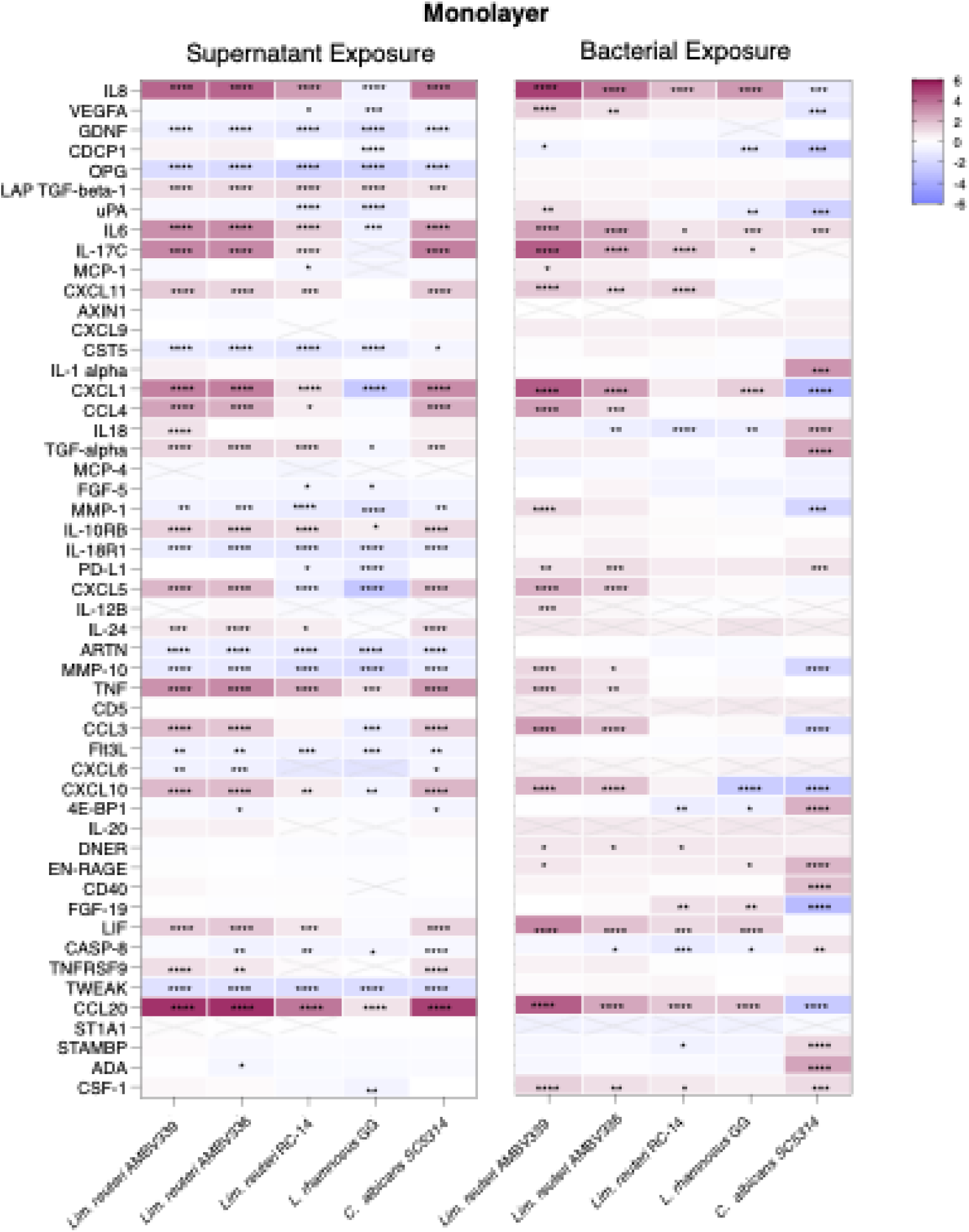
Immunoprofiling of VK2/E6E7 monolayer responses against microbes and their supernatant using the Olink 96 inflammation panel: Proteome levels induced by commensals *L. reuteri* AMBV339, *L. reuteri* AMBV336, *L. reuteri* RC14, *L. rhamnosus* GG and pathobiont *C. albicans* SC5314 are presented as fold changes (NpX differences compared to the control medium). Statistics are performed against the control medium (Statistics: Two-way ANOVA, Dunnet’s multiple comparisons correction, p-value < 0.05 = *, p-value < 0.01 = **, p-value < 0.001 = ***)

**Table S1.**
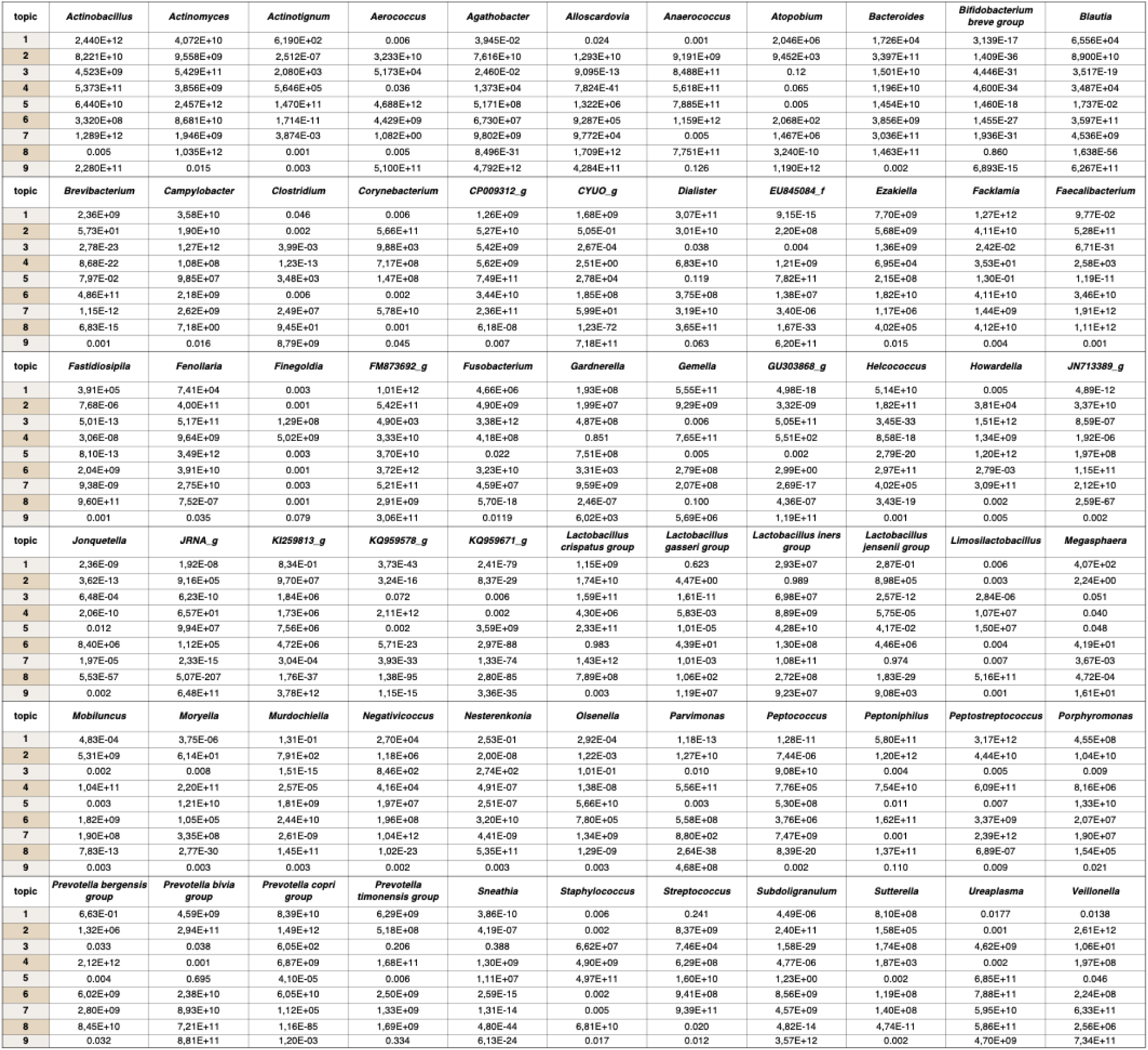
Topic modelling: Detailed contribution (weight) of vaginal microbial taxa to each of the 9 topics(*55*).

**Table S2.**
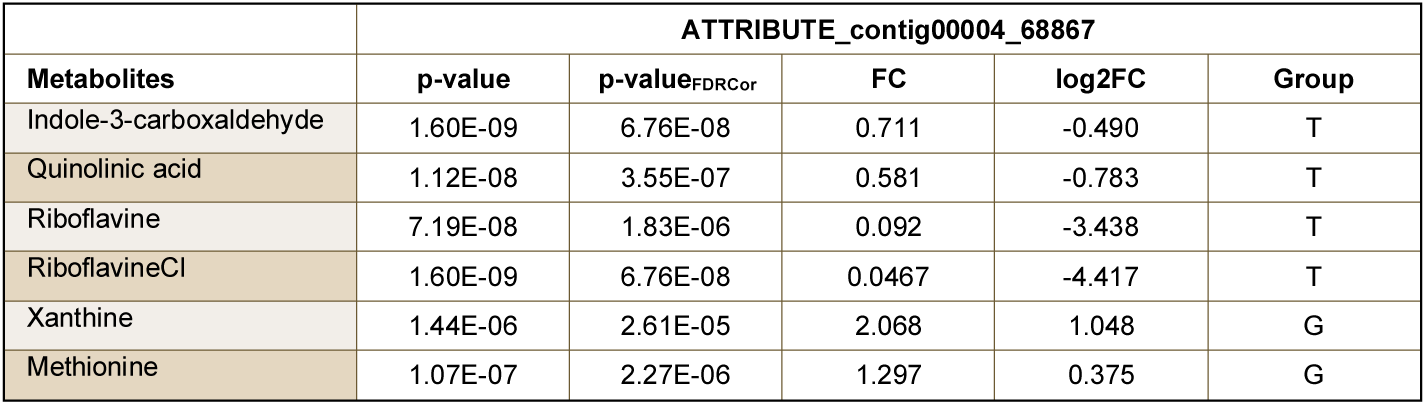
Impact of the G-to-T (contig 4, bp 68867) substitution on the metabolic production of *Lim. reuteri*: Semi-targeted metabolomics analysis was performed on overnight MRS cultures of 7 highly related *Lim. reuteri* strains (sharing only 33 SNPs amongst them). Differential abundance analysis was performed for all SNPs, and those highly significantly associated (p < 0.00001 with the G-to-T substitution in the RFN element of the *rib* operon are presented.

|  | ATTRIBUTE_contig00004_68867 |  |  |  |  |
| --- | --- | --- | --- | --- | --- |
| Metabolites | p-value | p-value<sub>FDRCor</sub> | FC | log2FC | Group |
| Indole-3-carboxaldehyde | 1.60E-09 | 6.76E-08 | 0.711 | -0.490 | T |
| Quinolinic acid | 1.12E-08 | 3.55E-07 | 0.581 | -0.783 | T |
| Riboflavine | 7.19E-08 | 1.83E-06 | 0.092 | -3.438 | T |
| RiboflavineCl | 1.60E-09 | 6.76E-08 | 0.0467 | -4.417 | T |
| Xanthine | 1.44E-06 | 2.61E-05 | 2.068 | 1.048 | G |
| Methionine | 1.07E-07 | 2.27E-06 | 1.297 | 0.375 | G |

**Table S3.** Composition of Simulated Vaginal Fluid. (*56*)

| Compound | Amount | Company |
| --- | --- | --- |
| Tween 80 | 0.1 % | Sigma-Aldrich |
| Ammonium citrate ($C_6H_{17}N_3O_7$) | 0.2 % | Sigma-Aldrich |
| Sodium acetate ($C_2H_3NaO_2$) | 0.5 % | Sigma-Aldrich |
| $MgSO_4 \cdot 7H_2O$ | 0.01 % | Sigma-Aldrich |
| $MnSO_4 \cdot H_2O$ | 0.005 % | Sigma-Aldrich |
| $K_2HPO_4$ | 0.2 % | Sigma-Aldrich |
| Proteose peptone N°3 | 0.3 % | BD Difco |
| Urea | 0.05 % | Sigma-Aldrich |
| D/L-lactic acid | 88 mM | Sigma-Aldrich |
| Mucin | 0.025 % | Sigma-Aldrich |
| Albumin (bovine) | 0.4 % | Roche |
| Vitamin solution (100x) | 1x | Carl Roth |
| Glucose | 1 % | Fischer Chemical |
| Glycogen | 1 % | Sigma-Aldrich |
| α-amylase | 2 mU/mL | Carl Roth |
| L-cysteine | 0.3 % | Carl Roth |

**Table S4.** Differential metabolic response between VK2 monolayers exposed to *Lim. reuteri* AMBV336 and *Lim. reuteri* AMBV339: This table gives an overview of all metabolites that are significantly differentially abundant (p < 0.05 after FDR correction) in VK2 monolayer lysates after a 3h exposure to 1:10 diluted bacterial supernatant of *Lim. reuteri* AMBV336 and *Lim. reuteri* AMBV339.

| Metabolite | p-value | p-value<sub>FDRcor</sub> | FC | log2FC | Group |
| --- | --- | --- | --- | --- | --- |
| 4-Aminobenzoic acid | 0.001506 | 0.015303 | 0.03 | -4.95 | V339 |
| 5-(2-oxopropylideneamino)-6-D-ribitylamino-uracil (5-OP-RU) | 0.008438 | 0.034359 | 0.04 | -4.74 | V339 |
| D-panthenol | 0.000142 | 0.004255 | 0.05 | -4.45 | V339 |
| Coenzyme Q10 | 0.00204 | 0.015303 | 0.12 | -3.01 | V339 |
| Hypoxanthine | 4.74E-05 | 0.004255 | 0.13 | -2.94 | V339 |
| Biotin | 0.001257 | 0.015303 | 0.13 | -2.94 | V339 |
| Glutathion red | 0.000421 | 0.009464 | 0.18 | -2.5 | V339 |
| All-trans retinal | 0.002733 | 0.018923 | 0.25 | -1.98 | V339 |
| Folic acid | 0.010073 | 0.034359 | 0.28 | -1.86 | V339 |
| 5-6-Dimethylbenzimidazole | 0.001995 | 0.015303 | 0.3 | -1.72 | V339 |
| Ethyl-3-Indoleacetate | 0.004034 | 0.021354 | 0.31 | -1.69 | V339 |
| p-cresol | 0.005724 | 0.025757 | 0.34 | -1.56 | V339 |
| Arginine | 0.000723 | 0.011372 | 0.35 | -1.53 | V339 |
| Indole-3-lactic acid | 0.00013 | 0.004255 | 0.38 | -1.39 | V339 |
| 5-amino-6-(D-ribitylamino)uracil (5-A-RU) | 0.004505 | 0.022278 | 0.44 | -1.2 | V339 |
| Tryptophan | 0.001741 | 0.015303 | 0.44 | -1.2 | V339 |
| Phenylalanine | 0.003567 | 0.021354 | 0.46 | -1.11 | V339 |
| Retinoic acid | 0.010308 | 0.034359 | 0.47 | -1.07 | V339 |
| Nicotinuric acid | 0.017049 | 0.04038 | 0.48 | -1.04 | V339 |
| Threonine | 0.004032 | 0.021354 | 0.49 | -1.04 | V339 |
| Skatole | 0.008357 | 0.034359 | 0.51 | -0.97 | V339 |
| Methionine | 0.004703 | 0.022278 | 0.51 | -0.97 | V339 |
| Glycine | 0.011105 | 0.035693 | 0.52 | -0.95 | V339 |
| Thiamine | 0.011833 | 0.036724 | 0.52 | -0.94 | V339 |
| Valine | 0.009423 | 0.034359 | 0.52 | -0.94 | V339 |
| Aspartic acid | 0.013391 | 0.038878 | 0.53 | -0.92 | V339 |
| Acetylcholine | 0.009519 | 0.034359 | 0.53 | -0.92 | V339 |
| Isoleucine-leucine | 0.0133 | 0.038878 | 0.53 | -0.92 | V339 |
| Alanine | 0.017014 | 0.04038 | 0.54 | -0.9 | V339 |
| D-pantothenic acid | 0.015792 | 0.03948 | 0.54 | -0.89 | V339 |
| Serine | 0.014605 | 0.03948 | 0.55 | -0.87 | V339 |
| Proline | 0.015779 | 0.03948 | 0.56 | -0.85 | V339 |
| Beta nicotinamide adenine dinucleotide (NAD+) | 0.020299 | 0.046844 | 0.56 | -0.83 | V339 |
| Taurine | 0.015003 | 0.03948 | 0.56 | -0.83 | V339 |
| Adenosine | 0.015751 | 0.03948 | 0.58 | -0.78 | V339 |
| Indole-3-acetic acid | 0.010248 | 0.034359 | 9.51 | 3.25 | **V336** |

**Table S5.** Metabolic response of VK2E6E7 monolayers against *Lim. reuteri* AMBV339: This table gives an overview of all metabolites that are significantly differentially abundant (p < 0.05 after FDR correction) in VK2 monolayer lysates in response to a 3h exposure to *Lim. reuteri* AMBV339.

| Metabolite | p-value | p-value<sub>FDRcor</sub> | FC | log2FC | Group |
| --- | --- | --- | --- | --- | --- |
| Mannitol | 0.017665 | 0.04676 | 0.04 | -4.62 | V339 |
| 5-6-Dimethylbenzimidazole | 0.00048 | 0.003323 | 0.07 | -3.74 | V339 |
| D-panthenol | 0.000181 | 0.002033 | 0.08 | -3.71 | V339 |
| Hypoxanthine | 0.000102 | 0.002033 | 0.16 | -2.64 | V339 |
| Valine | 0.008023 | 0.025788 | 0.18 | -2.45 | V339 |
| Coenzyme Q10 | 0.003445 | 0.014765 | 0.21 | -2.24 | V339 |
| p-cresol | 0.002481 | 0.011163 | 0.22 | -2.17 | V339 |
| Indole-3-acrylic acid | 0.00244 | 0.011163 | 0.22 | -2.17 | V339 |
| Taurocholic acid | 0.005477 | 0.018959 | 0.25 | -1.98 | V339 |
| Indole-3-propionic acid | 0.003782 | 0.0148 | 0.31 | -1.68 | V339 |
| 5-amino-6-(D-ribitylamino)uracil (5-A-RU) | 0.000839 | 0.005391 | 0.33 | -1.61 | V339 |
| Glutathion oxi | 0.003753 | 0.0148 | 0.34 | -1.55 | V339 |
| Ethy3-Indoleacetate | 0.004782 | 0.017931 | 0.35 | -1.53 | V339 |
| All-trans retinal | 0.011626 | 0.032699 | 0.43 | -1.23 | V339 |
| Indole-3-lactic acid | 0.000162 | 0.002033 | 0.53 | -0.91 | V339 |
| Tryptophan | 0.000228 | 0.002282 | 0.58 | -0.79 | V339 |
| Proline | 0.000165 | 0.002033 | 0.58 | -0.78 | V339 |
| Beta nicotinamide adenine dinucleotide (NAD<sup>+</sup>) | 3.85E-05 | 0.002033 | 0.59 | -0.77 | V339 |
| Arginine | 0.017542 | 0.04676 | 0.63 | -0.67 | V339 |
| Threonine | 0.000101 | 0.002033 | 0.68 | -0.56 | V339 |
| Phenylalanine | 0.000289 | 0.002418 | 0.69 | -0.53 | V339 |
| Taurine | 0.005332 | 0.018959 | 0.69 | -0.53 | V339 |
| Skatole | 0.000131 | 0.002033 | 0.69 | -0.53 | V339 |
| Methionine | 0.000295 | 0.002418 | 0.72 | -0.47 | V339 |
| Acetylcholine | 0.01132 | 0.032699 | 0.73 | -0.46 | V339 |
| Glycine | 0.001262 | 0.006767 | 0.74 | -0.44 | V339 |
| Lysine | 0.010615 | 0.031845 | 0.75 | -0.41 | V339 |
| Thiamine | 0.000136 | 0.002033 | 0.78 | -0.35 | V339 |
| Serine | 0.009179 | 0.028487 | 0.84 | -0.25 | V339 |
| 4-Aminobenzoic acid | 0.001278 | 0.006767 | Presence/Absence | Presence/Absence | V339 |
| 3-hydroxyanthranilic acid | 0.006283 | 0.020942 | 1.56 | 0.64 | MRS |
| Inosine | 0.001005 | 0.006029 | 3.65 | 1.87 | MRS |

**Table S6.** MAIT cell identification: Overview of all antibodies and dyes from Miltenyi Biotek for the identification of MAT cells and their activation markers.

| Marker | Fluorochrome | REA | Cat. No. | Location |
| --- | --- | --- | --- | --- |
| CD3 | Viobright V600 |  | 130-129-580 | Cell surface |
| TCR Va7.2 | PerCP-Vio700 | REA179 | 130-123-906 | Cell surface |
| IL18-Ra (CD218) | VioBright V423 | REA1095 | 130-130-465 | Cell surface |
| CD25 | PE | REA570 | 130-113-286 | Cell surface |
| CD69 | PE-Vio770 | REA824 | 130-112-615 | Cell surface |
| CD137 | Viobright FITC |  |  | Cell surface |
| CD71 | VioBright R720 | REA902 | 130-131-950 | Cell surface |
| CD107a (LAMP-1) | APC-Vio770 | REA792 | 130-111-701 | Intracellular |
| CD107b (LAMP-2) | APC | REA1073 | 130-118-819 | Intracellular |
| Live/dead | Viability 405/520 |  | 130-130-404 |  |

**Table S7.** Overview of the chemically synthesized riboflavin vitamers with their retention time and mass.

| **Riboflavin intermediate** | **Abreviation** | **Molecular Formula** | **RT (min)** | **Massm/z [M-H]** |
| --- | --- | --- | --- | --- |
| 5-amino-6-(D-ribitylamino)uracil | 5-A-RU | C<sub>9</sub>H<sub>16</sub>N<sub>4</sub>O<sub>6</sub> | 1.5 | 275.0992 |
| 5-(2-oxopropylideneamino)-6-D-ribitylaminouracil | 5-OP-RU | C<sub>12</sub>H<sub>18</sub>N<sub>4</sub>O<sub>7</sub> | 4.6 | 329.1098 |
| 5-(2-oxoethylideneamino)-6-D-ribitylaminouracil | 5-OE-RU | C<sub>11</sub>H<sub>15</sub>N<sub>4</sub>O<sub>7</sub> | 3.6 | 315.0941 |
| 7-hydroxy-6-methyl-8-(1-D-ribityl)lumazine | R,L-6-Me-7-OH | C<sub>12</sub>H<sub>16</sub>N<sub>4</sub>O<sub>7</sub> | 5 | 327.0941 |
| 6,7-Dimethylribityl Lumazine | 6,7-diMe-Lum | C<sub>13</sub>H<sub>18</sub>N<sub>4</sub>O<sub>6</sub> | 4.13 | 325.1148 |

**Table S8.**
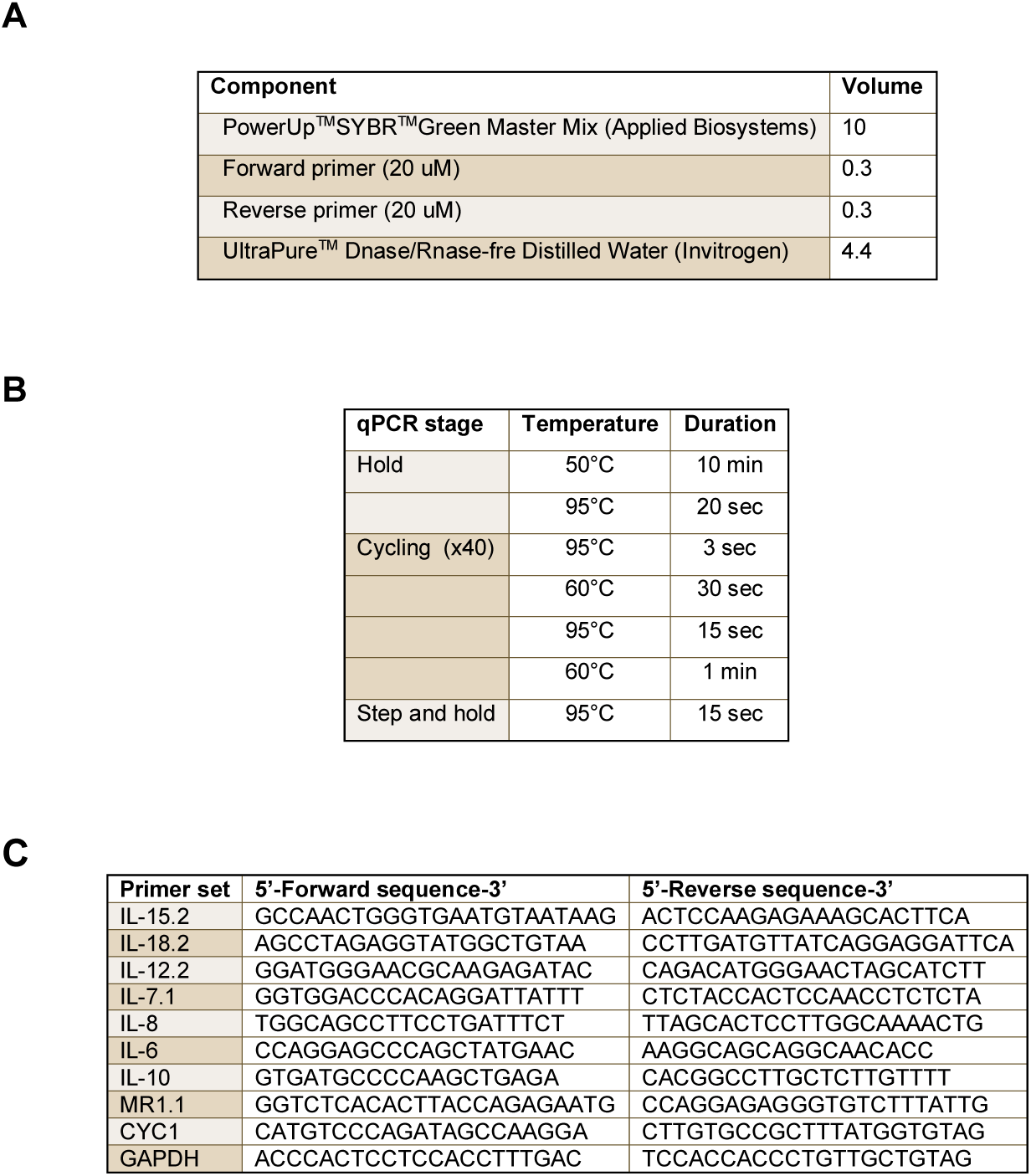
qRT-PCR overview: **(A)** mastermix composition **(B)** instrument settings and **(C)** primers

