## Supplementary material for "Vitamin B_2_ Production by Vaginal Lactobacilli Promotes Symbiosis": Data S10: 250912 Only CE.pdf

### Report of PBMC 12/09/25

Specimen Name: PBMC 12/09/25

Run Time: 12-Sep-25 1:26 PM

Cytometer: NovoCyte Quanteon 621181110427

Software: NovoExpress 1.6.0

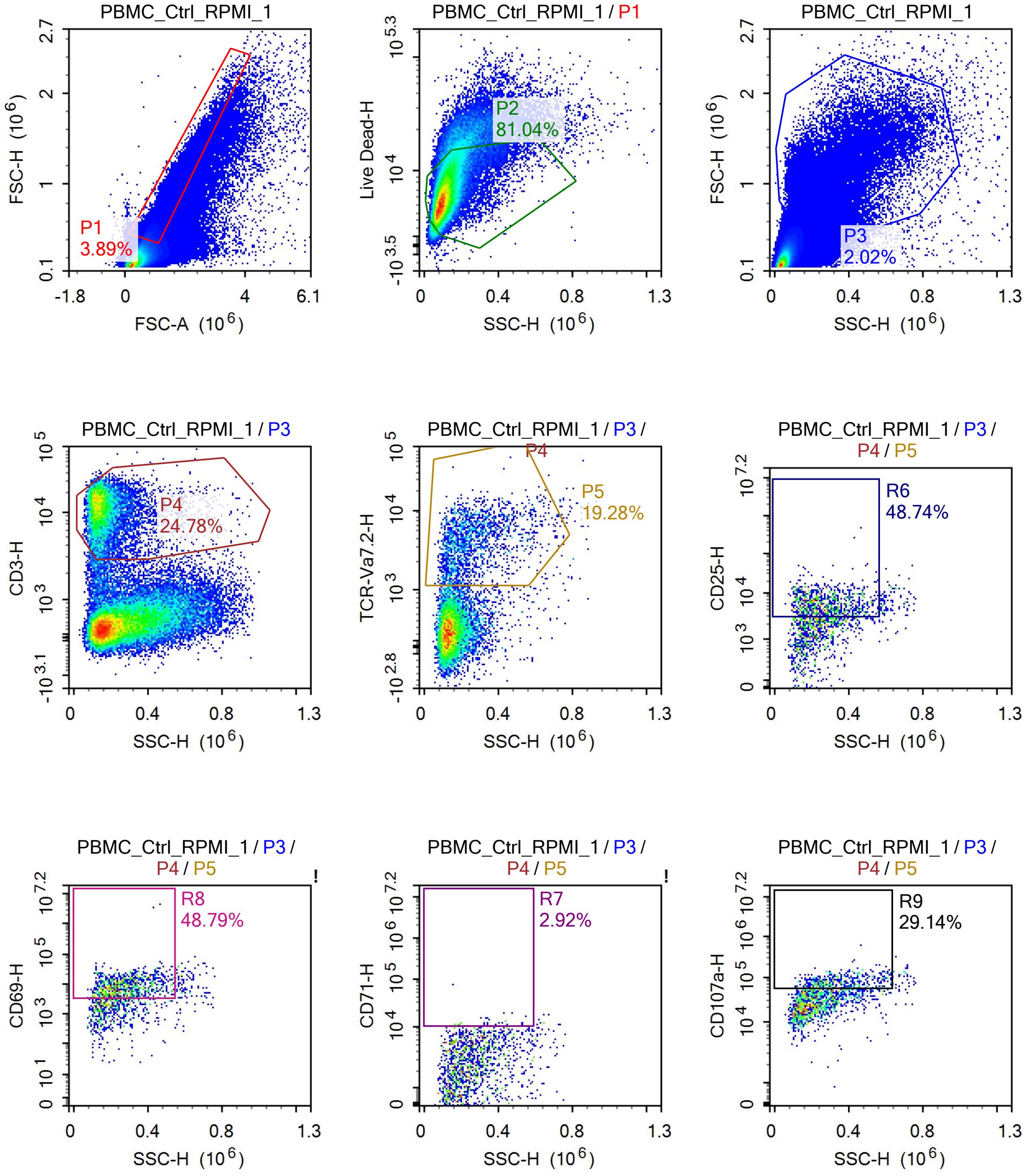

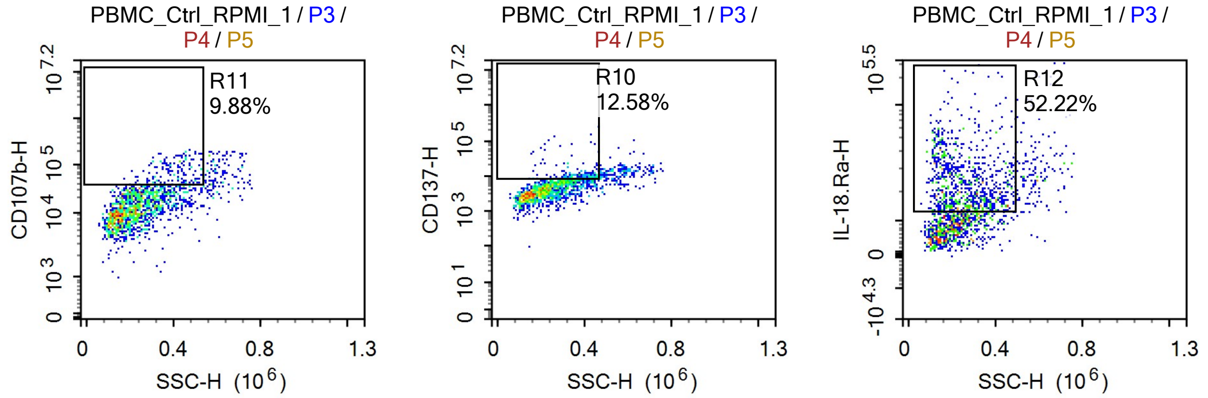


Sample Statistics of PBMC_Ctrl_RPMI_1

| Gate | Count | % Parent | X | Y | Median X | Median Y |
|---|---|---|---|---|---|---|
| All | 1,842,411 | | | | | |
| P1 | 71,739 | 3.89% | FSC-A | FSC-H | 1,014,041 | 489,640 |
| P2 | 58,137 | 81.04% | SSC-H | Live Dead-H | 107,806 | 6,169 |
| P3 | 37,278 | 2.02% | SSC-H | FSC-H | 204,862 | 954,967 |
| P4 | 9,237 | 24.78% | SSC-H | CD3-H | 146,861 | 11,368 |
| P5 | 1,781 | 19.28% | SSC-H | TCR-Va7.2-H | 231,900 | 3,823 |
| R6 | 868 | 48.74% | SSC-H | CD25-H | 258,153 | 5,571 |
| R7 | 52 | 2.92% | SSC-H | CD71-H | 327,375 | 11,127 |
| R8 | 869 | 48.79% | SSC-H | CD69-H | 257,849 | 8,180 |
| R9 | 519 | 29.14% | SSC-H | CD107a-H | 320,524 | 77,327 |
| R10 | 224 | 12.58% | SSC-H | CD137-H | 366,419 | 9,693 |
| R11 | 176 | 9.88% | SSC-H | CD107b-H | 400,009 | 66,444 |
| R12 | 930 | 52.22% | SSC-H | IL-18.Ra-H | 241,819 | 21,868 |

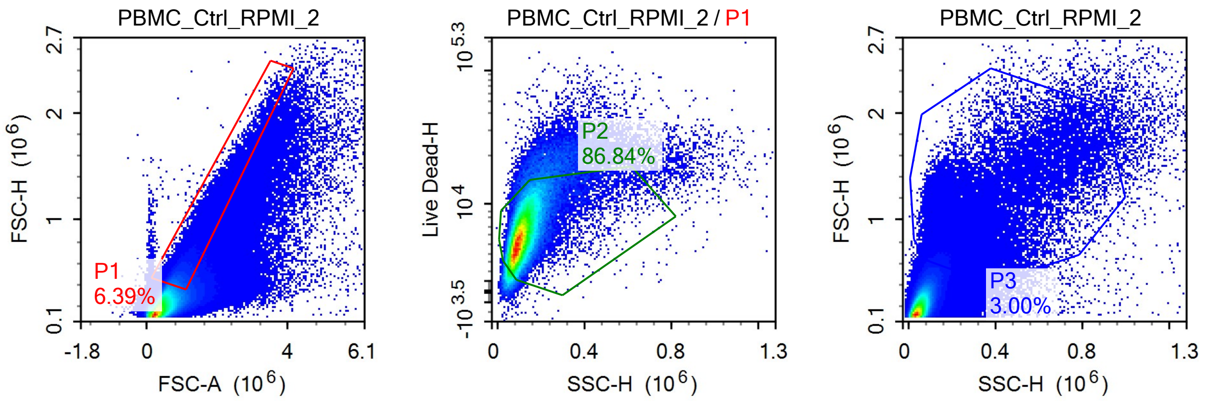

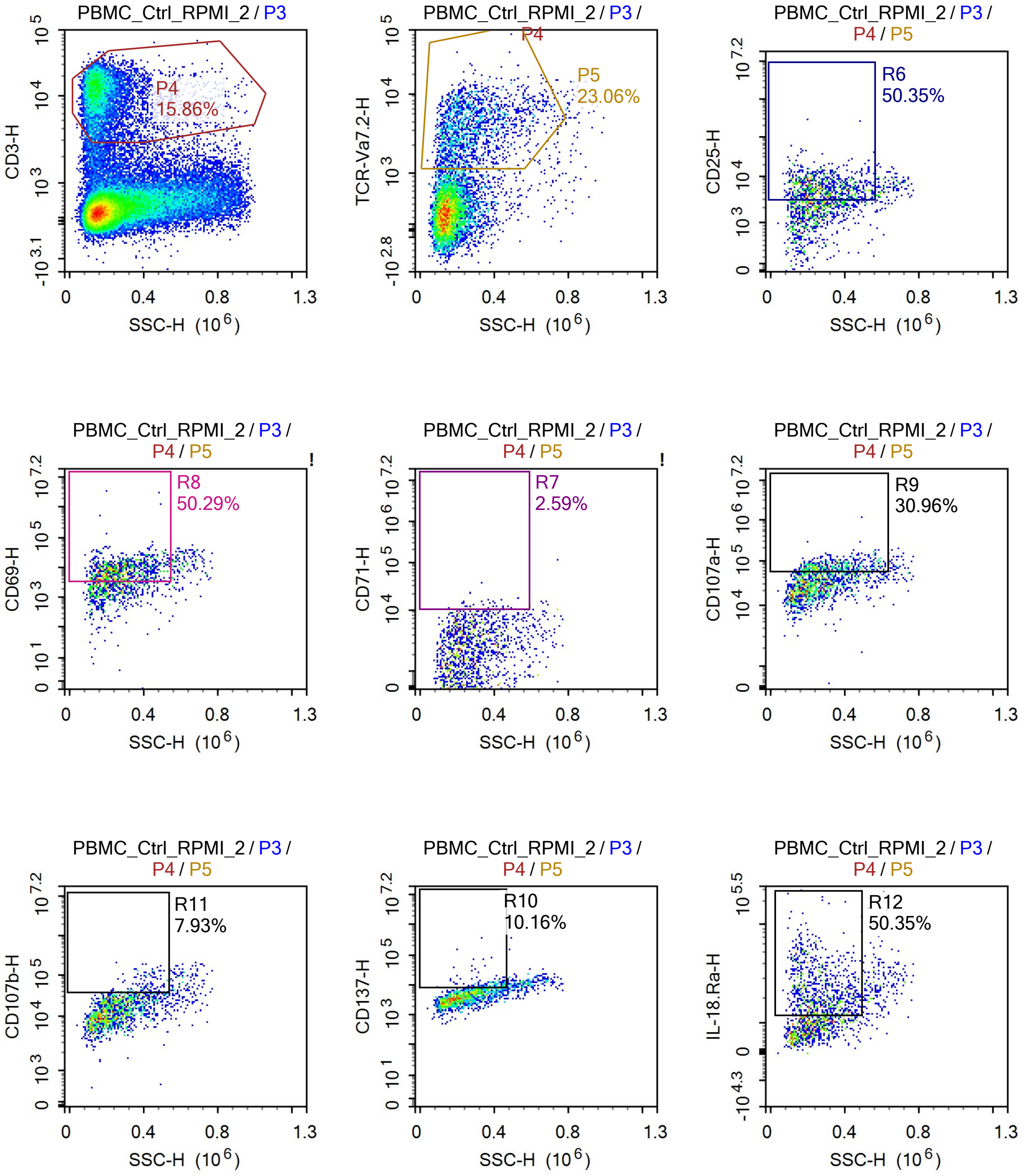
PBMC_Ctrl_RPMI_2 / P3
P4
15.86%
CD3-H
SSC-H (10^6)
PBMC_Ctrl_RPMI_2 / P3 / P4
P5
23.06%
TCR-Va7.2-H
PBMC_Ctrl_RPMI_2 / P3 / P4 / P5
R6
50.35%
CD25-H
R8
50.29%
CD69-H
R7
2.59%
CD71-H
R9
30.96%
CD107a-H
R11
7.93%
CD107b-H
R10
10.16%
CD137-H
R12
50.35%
IL-18.Ra-H

Sample Statistics of PBMC_Ctrl_RPMI_2

| Gate | Count | % Parent | X | Y | Median X | Median Y |
|---|---|---|---|---|---|---|
| All | 1,549,068 | | | | | |
| P1 | 99,042 | 6.39% | FSC-A | FSC-H | 990,278 | 472,055 |
| P2 | 86,010 | 86.84% | SSC-H | Live Dead-H | 105,827 | 5,831 |
| P3 | 46,549 | 3.00% | SSC-H | FSC-H | 196,318 | 825,779 |
| P4 | 7,382 | 15.86% | SSC-H | CD3-H | 161,027 | 10,793 |
| P5 | 1,702 | 23.06% | SSC-H | TCR-Va7.2-H | 252,958 | 3,481 |
| R6 | 857 | 50.35% | SSC-H | CD25-H | 275,174 | 5,458 |
| R7 | 44 | 2.59% | SSC-H | CD71-H | 347,088 | 11,927 |
| R8 | 856 | 50.29% | SSC-H | CD69-H | 274,817 | 8,325 |
| R9 | 527 | 30.96% | SSC-H | CD107a-H | 334,835 | 73,179 |
| R10 | 173 | 10.16% | SSC-H | CD137-H | 387,671 | 9,663 |
| R11 | 135 | 7.93% | SSC-H | CD107b-H | 406,445 | 53,203 |
| R12 | 857 | 50.35% | SSC-H | IL-18.Ra-H | 256,673 | 21,928 |

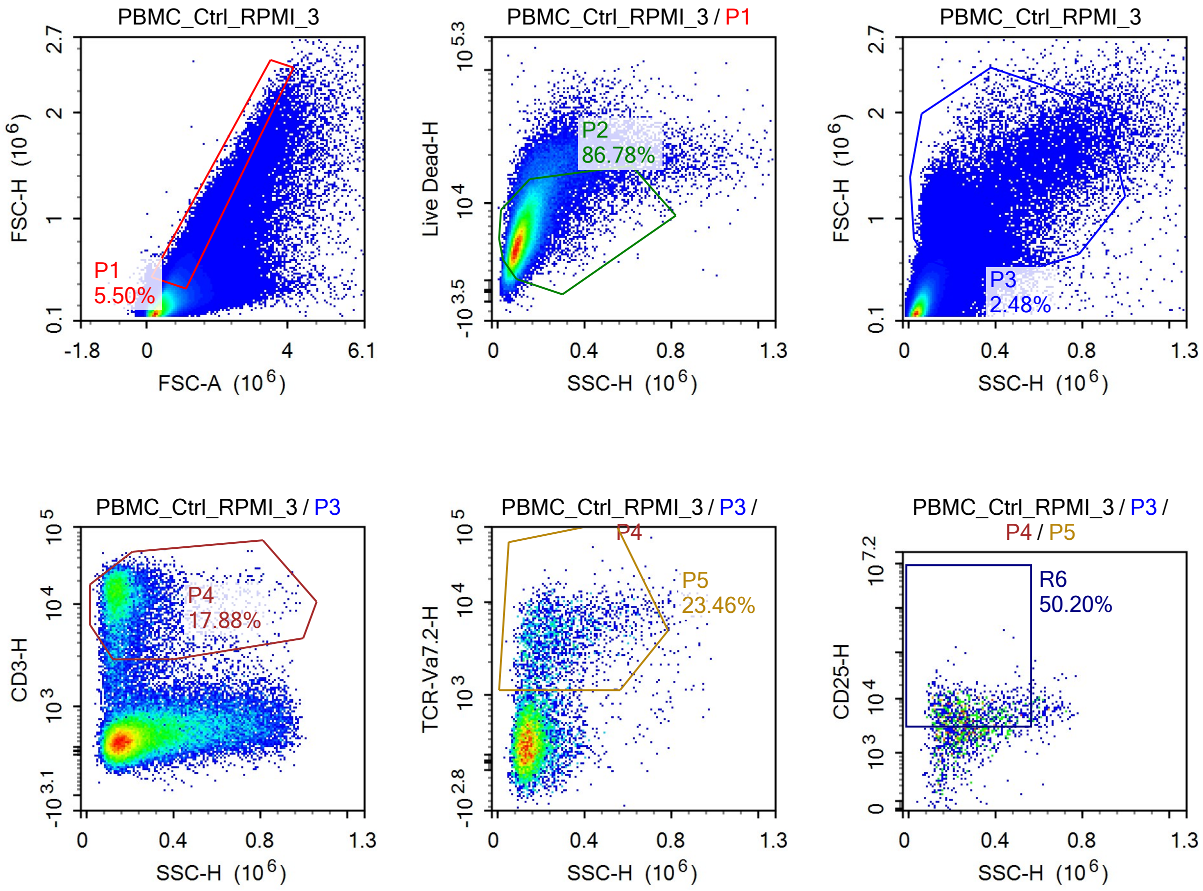

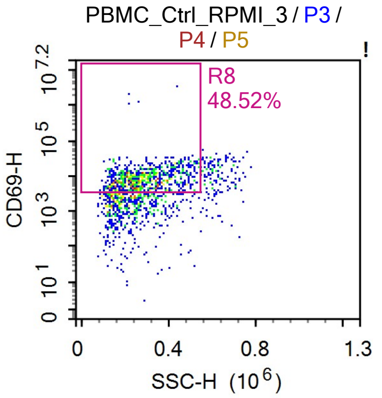


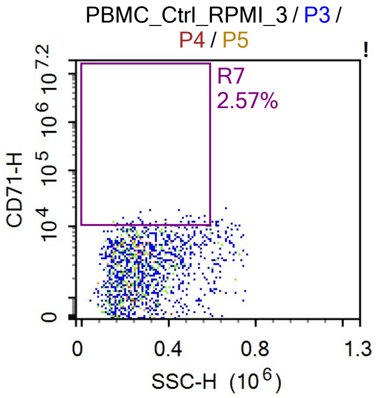


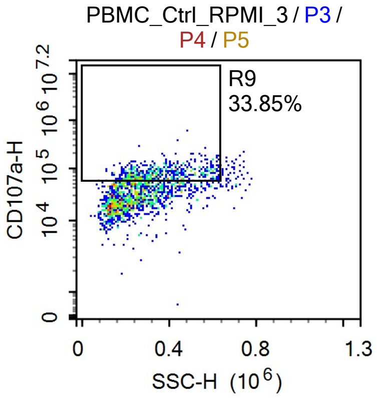


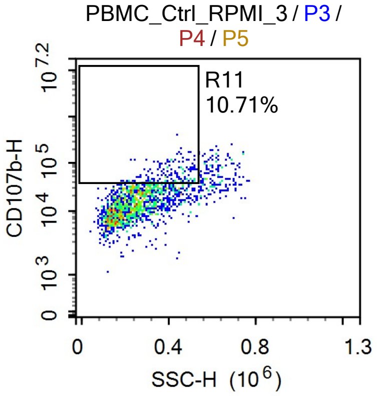


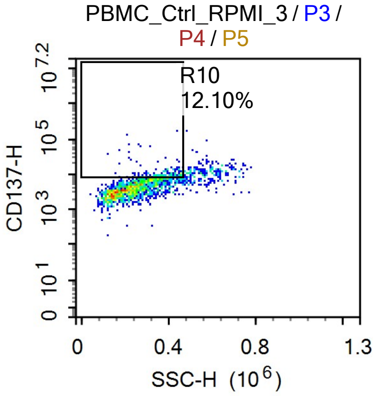


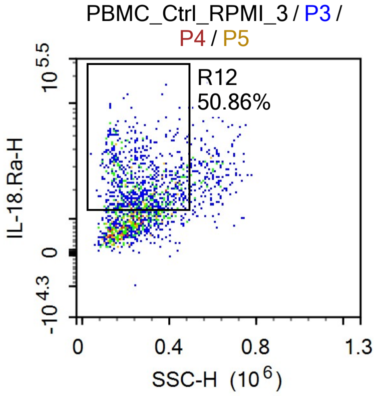


Sample Statistics of PBMC_Ctrl_RPMI_3

| Gate | Count | % Parent | X | Y | Median X | Median Y |
|---|---|---|---|---|---|---|
| All | 1,725,636 | | | | | |
| P1 | 94,859 | 5.50% | FSC-A | FSC-H | 986,720 | 469,540 |
| P2 | 82,321 | 86.78% | SSC-H | Live Dead-H | 104,129 | 5,650 |
| P3 | 42,742 | 2.48% | SSC-H | FSC-H | 195,647 | 840,369 |
| P4 | 7,643 | 17.88% | SSC-H | CD3-H | 157,558 | 11,567 |
| P5 | 1,793 | 23.46% | SSC-H | TCR-Va7.2-H | 253,278 | 3,799 |
| R6 | 900 | 50.20% | SSC-H | CD25-H | 261,064 | 5,612 |
| R7 | 46 | 2.57% | SSC-H | CD71-H | 363,562 | 11,789 |
| R8 | 870 | 48.52% | SSC-H | CD69-H | 274,720 | 8,280 |
| R9 | 607 | 33.85% | SSC-H | CD107a-H | 338,306 | 75,685 |
| R10 | 217 | 12.10% | SSC-H | CD137-H | 373,034 | 9,498 |
| R11 | 192 | 10.71% | SSC-H | CD107b-H | 402,920 | 48,095 |
| R12 | 912 | 50.86% | SSC-H | IL-18.Ra-H | 261,667 | 21,548 |

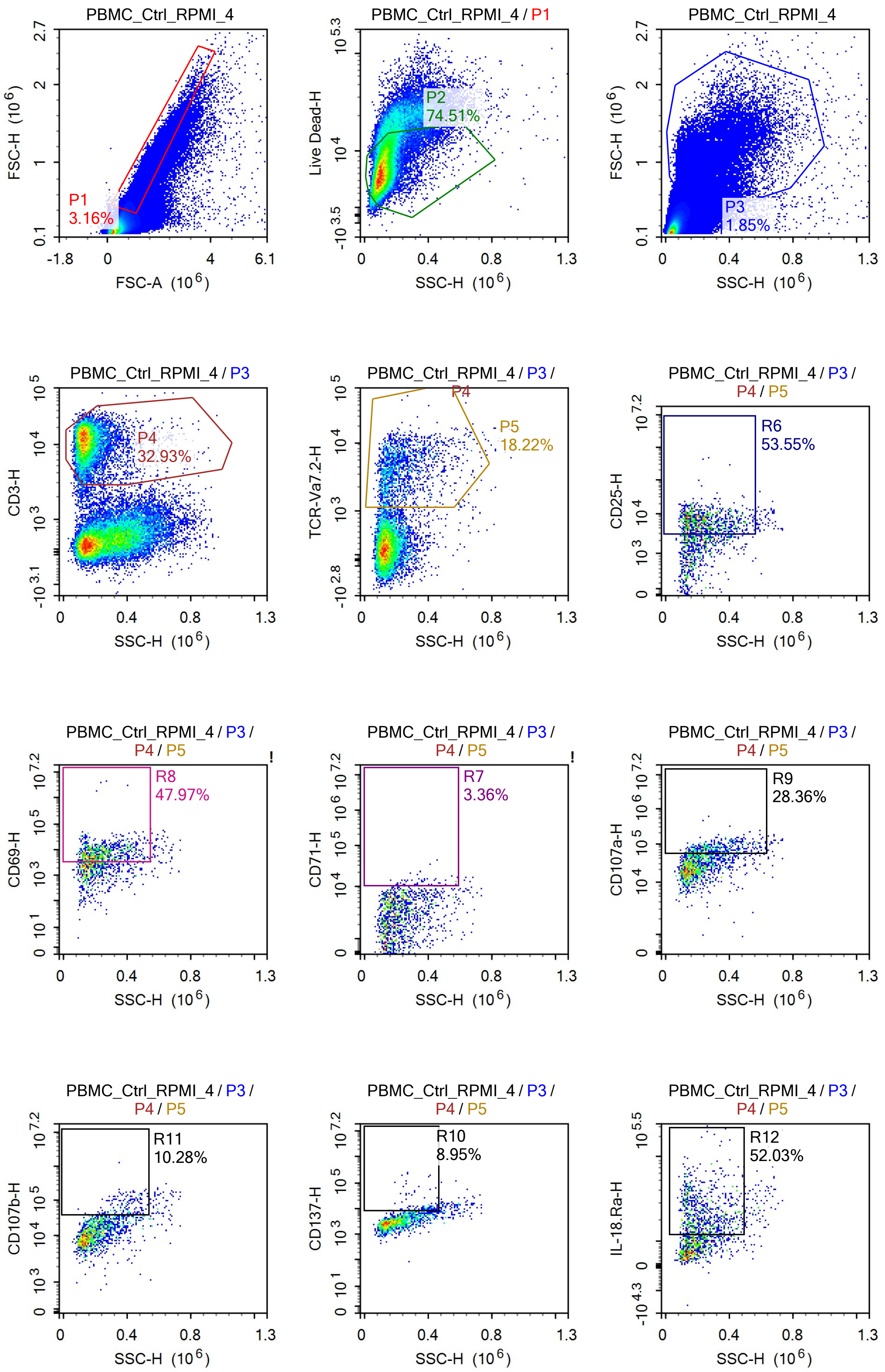
PBMC_Ctrl_RPMI_4
FSC-H (10^6)
FSC-A (10^6)
P1
3.16%
PBMC_Ctrl_RPMI_4 / P1
Live Dead-H
SSC-H (10^6)
P2
74.51%
PBMC_Ctrl_RPMI_4
FSC-H (10^6)
P3
1.85%
PBMC_Ctrl_RPMI_4 / P3
CD3-H
P4
32.93%
PBMC_Ctrl_RPMI_4 / P3 /
TCR-Va7.2-H
P4
P5
18.22%
PBMC_Ctrl_RPMI_4 / P3 /
P4 / P5
CD25-H
R6
53.55%
PBMC_Ctrl_RPMI_4 / P3 /
P4 / P5
CD69-H
R8
47.97%
PBMC_Ctrl_RPMI_4 / P3 /
P4 / P5
CD71-H
R7
3.36%
PBMC_Ctrl_RPMI_4 / P3 /
P4 / P5
CD107a-H
R9
28.36%
PBMC_Ctrl_RPMI_4 / P3 /
P4 / P5
CD107b-H
R11
10.28%
PBMC_Ctrl_RPMI_4 / P3 /
P4 / P5
CD137-H
R10
8.95%
PBMC_Ctrl_RPMI_4 / P3 /
P4 / P5
IL-18.Ra-H
R12
52.03%

Sample Statistics of PBMC_Ctrl_RPMI_4

| Gate | Count | % Parent | X | Y | Median X | Median Y |
|---|---|---|---|---|---|---|
| All | 1,423,509 | | | | | |
| P1 | 45,021 | 3.16% | FSC-A | FSC-H | 1,060,403 | 531,699 |
| P2 | 33,543 | 74.51% | SSC-H | Live Dead-H | 114,934 | 6,584 |
| P3 | 26,265 | 1.85% | SSC-H | FSC-H | 195,199 | 995,281 |
| P4 | 8,648 | 32.93% | SSC-H | CD3-H | 140,461 | 11,458 |
| P5 | 1,576 | 18.22% | SSC-H | TCR-Va7.2-H | 202,106 | 3,883 |
| R6 | 844 | 53.55% | SSC-H | CD25-H | 223,597 | 6,304 |
| R7 | 53 | 3.36% | SSC-H | CD71-H | 350,403 | 12,388 |
| R8 | 756 | 47.97% | SSC-H | CD69-H | 226,637 | 7,345 |
| R9 | 447 | 28.36% | SSC-H | CD107a-H | 303,845 | 75,946 |
| R10 | 141 | 8.95% | SSC-H | CD137-H | 366,146 | 9,936 |
| R11 | 162 | 10.28% | SSC-H | CD107b-H | 393,861 | 61,424 |
| R12 | 820 | 52.03% | SSC-H | IL-18.Ra-H | 223,340 | 22,299 |

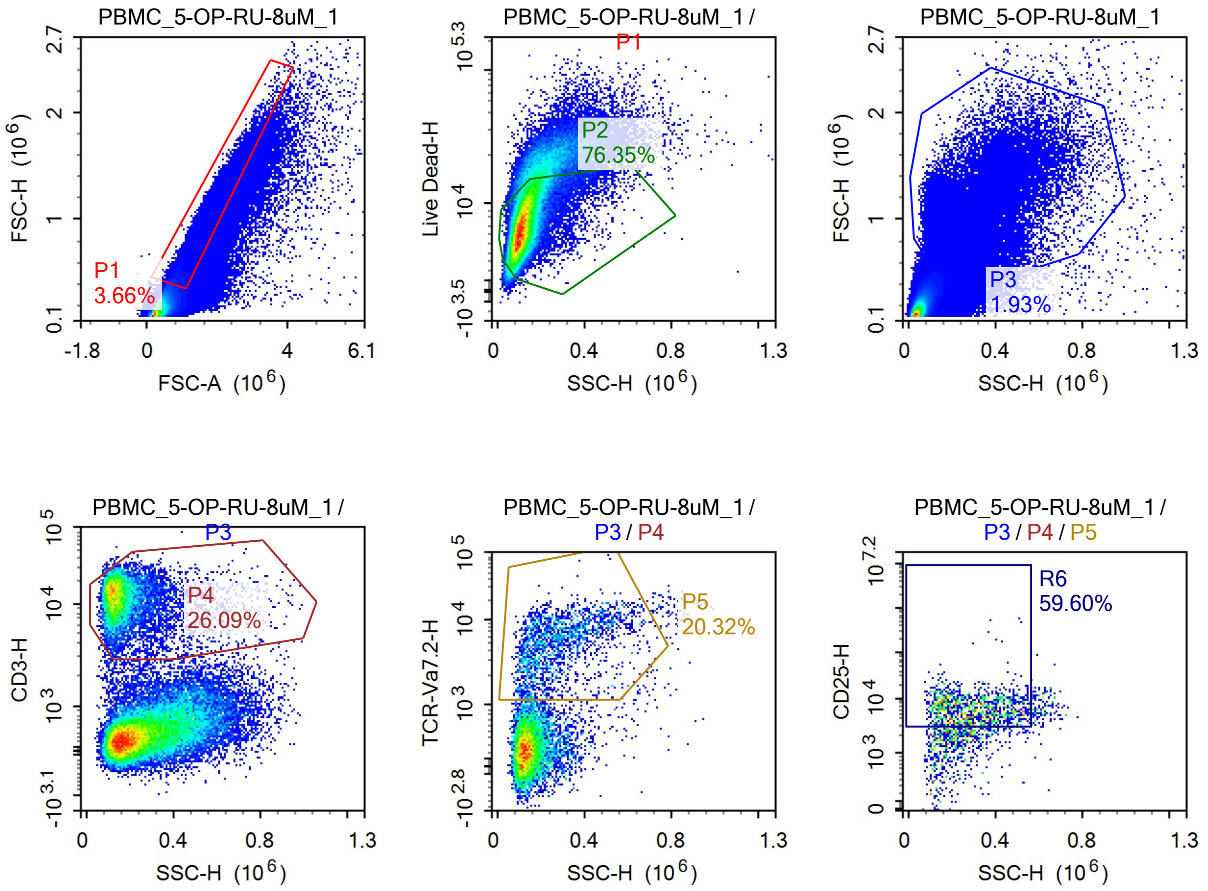

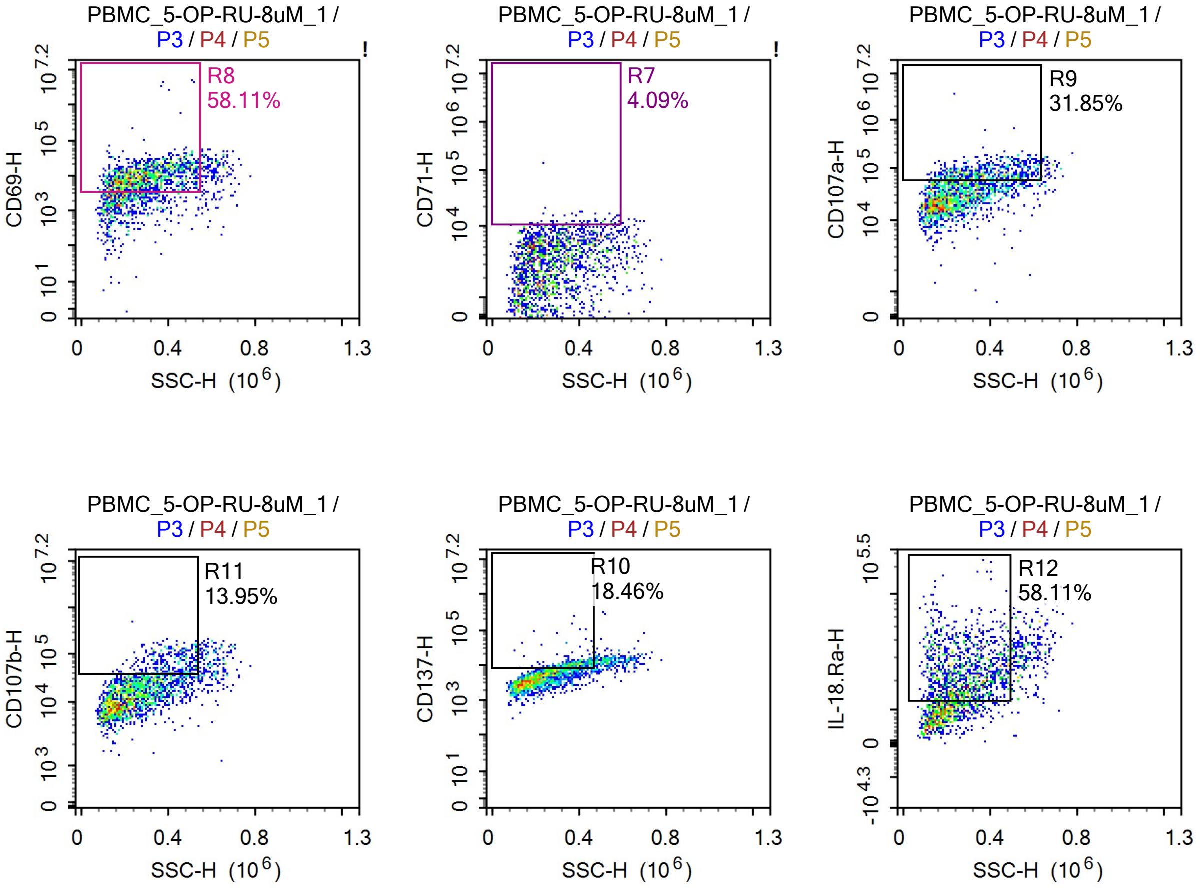


Sample Statistics of PBMC_5-OP-RU-8uM_1

| Gate | Count | % Parent | X | Y | Median X | Median Y |
|---|---|---|---|---|---|---|
| All | 2,101,112 | | | | | |
| P1 | 76,918 | 3.66% | FSC-A | FSC-H | 1,005,279 | 501,881 |
| P2 | 58,728 | 76.35% | SSC-H | Live Dead-H | 117,049 | 7,135 |
| P3 | 40,574 | 1.93% | SSC-H | FSC-H | 211,422 | 936,969 |
| P4 | 10,587 | 26.09% | SSC-H | CD3-H | 144,061 | 12,225 |
| P5 | 2,151 | 20.32% | SSC-H | TCR-Va7.2-H | 254,947 | 5,249 |
| R6 | 1,282 | 59.60% | SSC-H | CD25-H | 287,998 | 6,204 |
| R7 | 88 | 4.09% | SSC-H | CD71-H | 381,966 | 11,129 |
| R8 | 1,250 | 58.11% | SSC-H | CD69-H | 284,115 | 9,526 |
| R9 | 685 | 31.85% | SSC-H | CD107a-H | 367,158 | 75,784 |
| R10 | 397 | 18.46% | SSC-H | CD137-H | 366,421 | 10,112 |
| R11 | 300 | 13.95% | SSC-H | CD107b-H | 408,812 | 66,626 |
| R12 | 1,250 | 58.11% | SSC-H | IL-18.Ra-H | 277,017 | 22,659 |

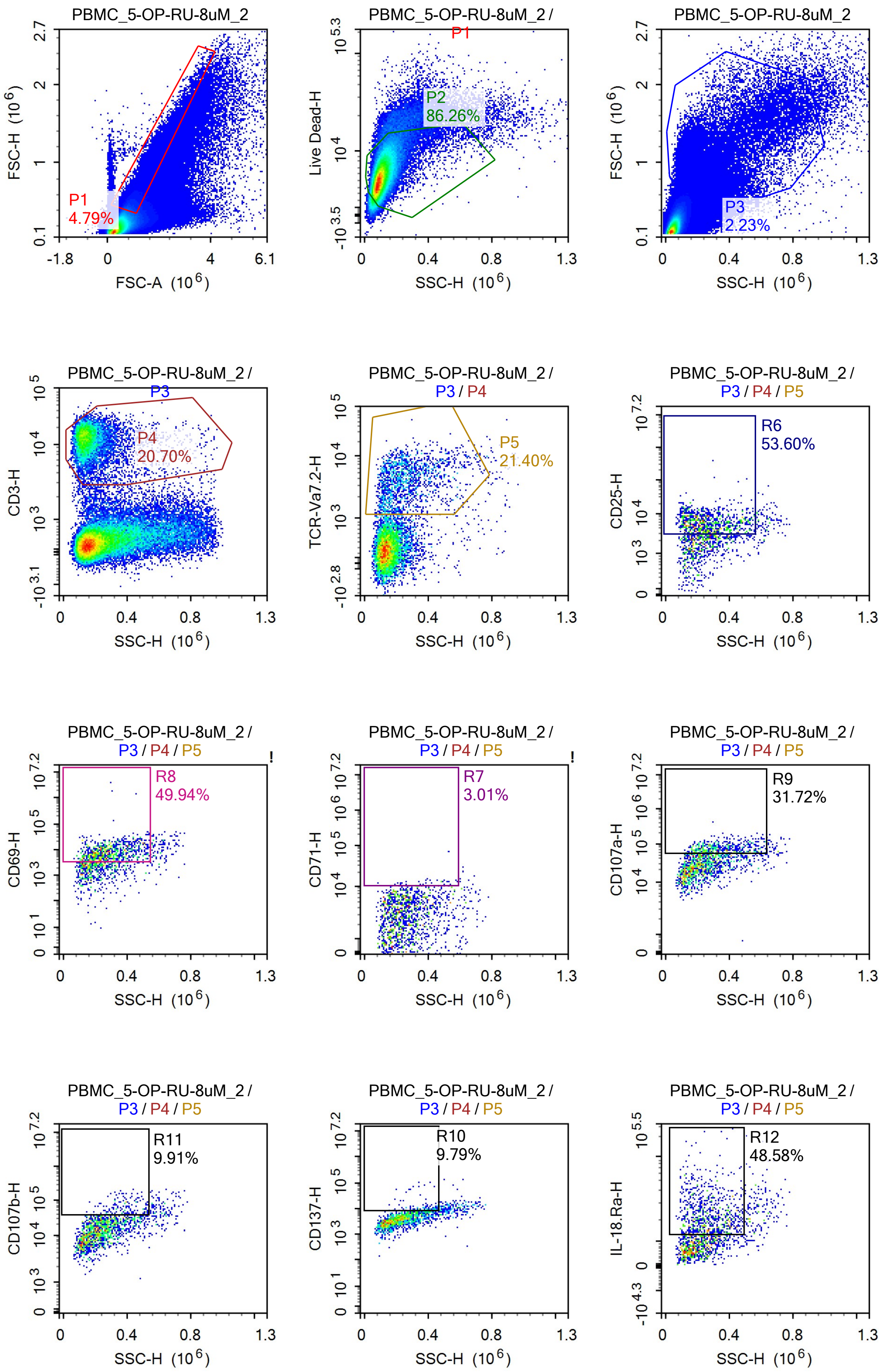
PBMC_5-OP-RU-8uM_2
FSC-H (10^6)
FSC-A (10^6)
P1
4.79%
PBMC_5-OP-RU-8uM_2 / P1
Live Dead-H
SSC-H (10^6)
P2
86.26%
PBMC_5-OP-RU-8uM_2
FSC-H (10^6)
P3
2.23%
PBMC_5-OP-RU-8uM_2 / P3
CD3-H
P4
20.70%
PBMC_5-OP-RU-8uM_2 / P3 / P4
TCR-Va7.2-H
P5
21.40%
PBMC_5-OP-RU-8uM_2 / P3 / P4 / P5
CD25-H
R6
53.60%
CD69-H
R8
49.94%
CD71-H
R7
3.01%
CD107a-H
R9
31.72%
CD107b-H
R11
9.91%
CD137-H
R10
9.79%
IL-18.Ra-H
R12
48.58%

Sample Statistics of PBMC_5-OP-RU-8uM_2

| Gate | Count | % Parent | X | Y | Median X | Median Y |
|---|---|---|---|---|---|---|
| All | 1,718,594 | | | | | |
| P1 | 82,332 | 4.79% | FSC-A | FSC-H | 992,647 | 463,118 |
| P2 | 71,021 | 86.26% | SSC-H | Live Dead-H | 100,626 | 5,494 |
| P3 | 38,291 | 2.23% | SSC-H | FSC-H | 194,743 | 862,481 |
| P4 | 7,926 | 20.70% | SSC-H | CD3-H | 150,916 | 12,251 |
| P5 | 1,696 | 21.40% | SSC-H | TCR-Va7.2-H | 239,696 | 3,798 |
| R6 | 909 | 53.60% | SSC-H | CD25-H | 249,408 | 5,735 |
| R7 | 51 | 3.01% | SSC-H | CD71-H | 383,468 | 11,153 |
| R8 | 847 | 49.94% | SSC-H | CD69-H | 269,442 | 7,874 |
| R9 | 538 | 31.72% | SSC-H | CD107a-H | 323,647 | 72,542 |
| R10 | 166 | 9.79% | SSC-H | CD137-H | 376,920 | 9,466 |
| R11 | 168 | 9.91% | SSC-H | CD107b-H | 387,057 | 58,422 |
| R12 | 824 | 48.58% | SSC-H | IL-18.Ra-H | 257,053 | 21,046 |

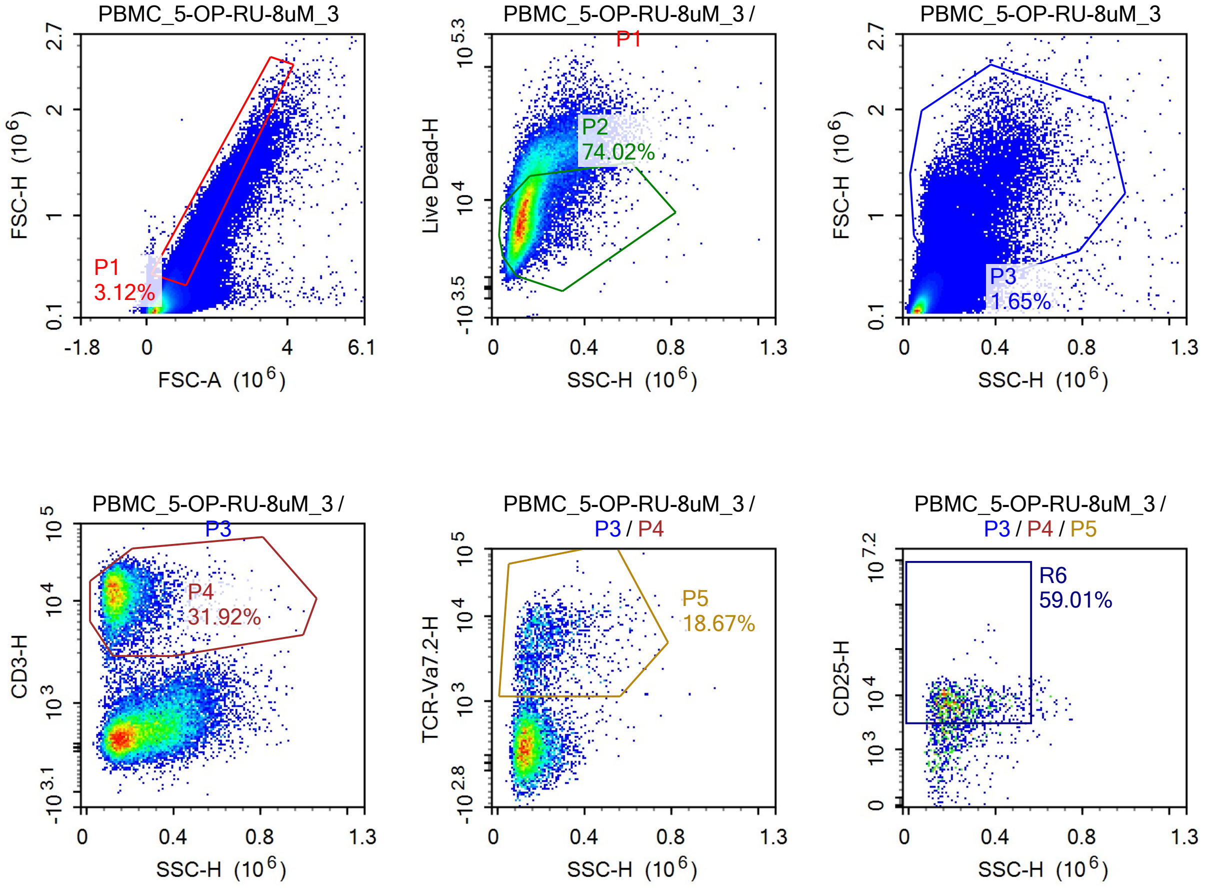

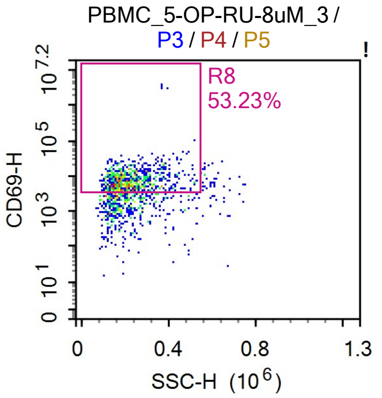


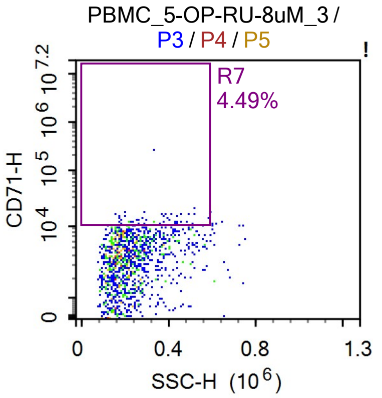


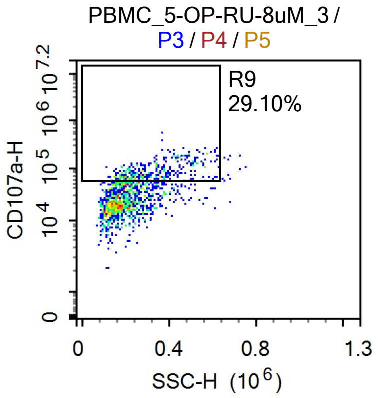


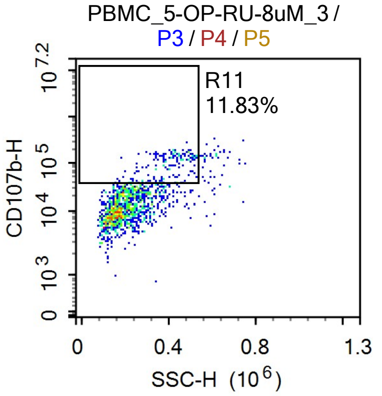


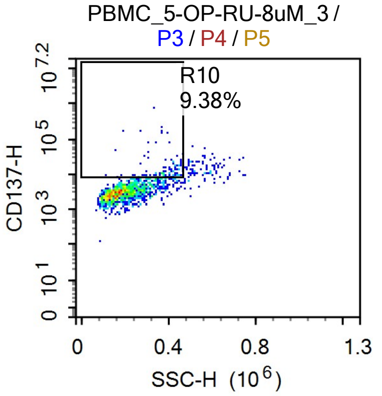


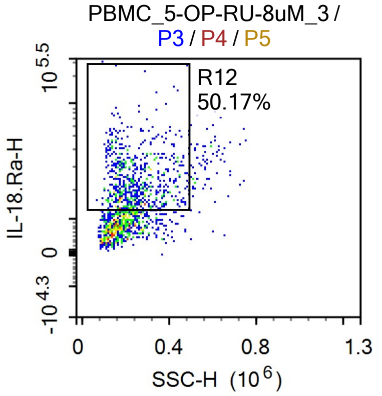


Sample Statistics of PBMC_5-OP-RU-8uM_3

| Gate | Count | % Parent | X | Y | Median X | Median Y |
|---|---|---|---|---|---|---|
| All | 1,493,888 | | | | | |
| P1 | 46,535 | 3.12% | FSC-A | FSC-H | 1,014,305 | 508,650 |
| P2 | 34,444 | 74.02% | SSC-H | Live Dead-H | 119,118 | 7,528 |
| P3 | 24,689 | 1.65% | SSC-H | FSC-H | 193,958 | 963,657 |
| P4 | 7,880 | 31.92% | SSC-H | CD3-H | 142,962 | 12,035 |
| P5 | 1,471 | 18.67% | SSC-H | TCR-Va7.2-H | 204,296 | 4,503 |
| R6 | 868 | 59.01% | SSC-H | CD25-H | 216,002 | 6,723 |
| R7 | 66 | 4.49% | SSC-H | CD71-H | 316,834 | 12,409 |
| R8 | 783 | 53.23% | SSC-H | CD69-H | 212,605 | 7,412 |
| R9 | 428 | 29.10% | SSC-H | CD107a-H | 289,994 | 80,220 |
| R10 | 138 | 9.38% | SSC-H | CD137-H | 371,737 | 10,096 |
| R11 | 174 | 11.83% | SSC-H | CD107b-H | 402,415 | 99,604 |
| R12 | 738 | 50.17% | SSC-H | IL-18.Ra-H | 229,624 | 21,863 |

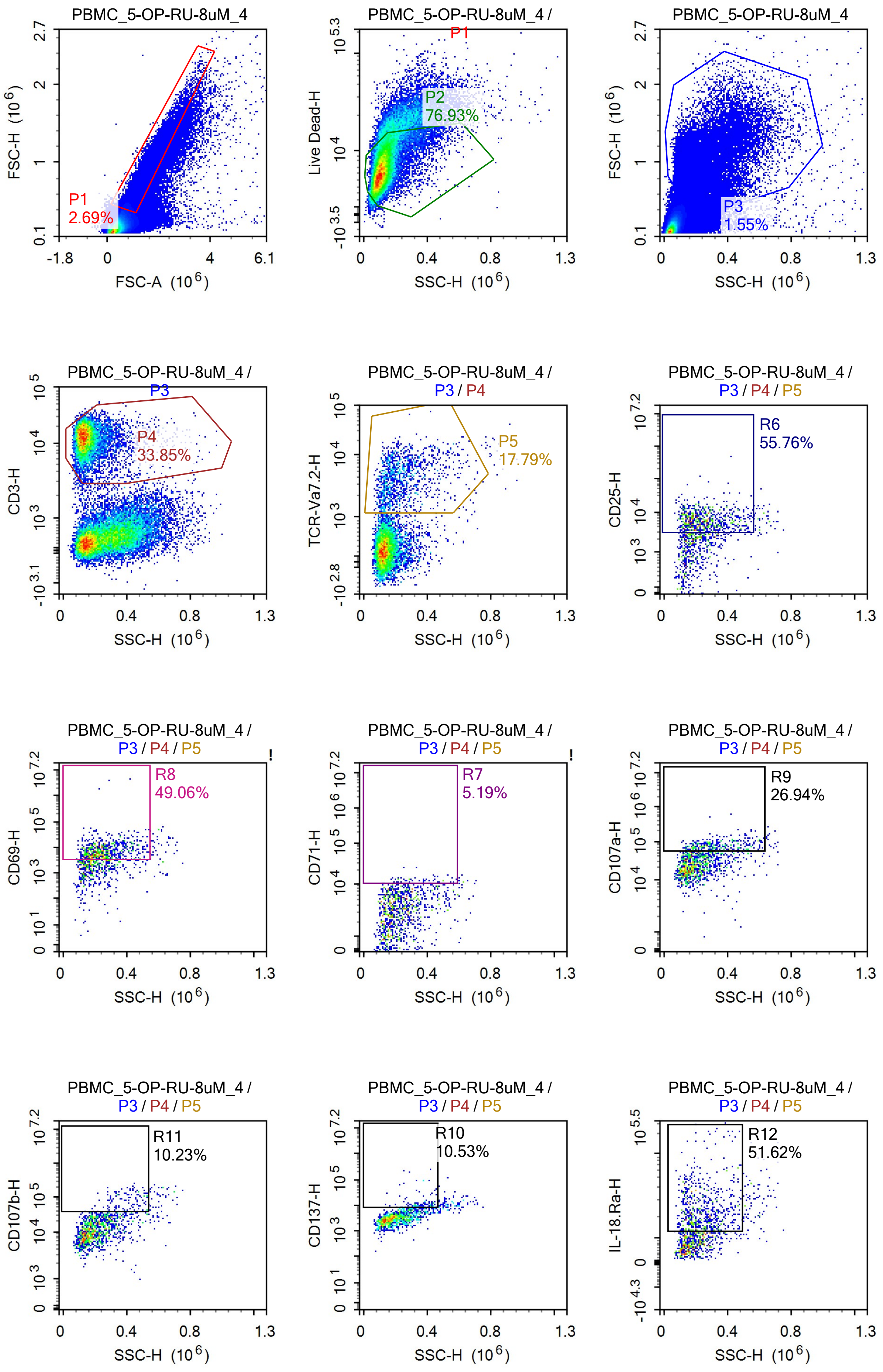

PBMC_5-OP-RU-8uM_4
FSC-H (10^6)
FSC-A (10^6)
P1
2.69%
PBMC_5-OP-RU-8uM_4 / P1
Live Dead-H
SSC-H (10^6)
P2
76.93%
PBMC_5-OP-RU-8uM_4
FSC-H (10^6)
SSC-H (10^6)
P3
1.55%
PBMC_5-OP-RU-8uM_4 / P3
CD3-H
SSC-H (10^6)
P4
33.85%
PBMC_5-OP-RU-8uM_4 / P3 / P4
TCR-Va7.2-H
SSC-H (10^6)
P5
17.79%
PBMC_5-OP-RU-8uM_4 / P3 / P4 / P5
CD25-H
SSC-H (10^6)
R6
55.76%
PBMC_5-OP-RU-8uM_4 / P3 / P4 / P5
CD69-H
SSC-H (10^6)
R8
49.06%
PBMC_5-OP-RU-8uM_4 / P3 / P4 / P5
CD71-H
SSC-H (10^6)
R7
5.19%
PBMC_5-OP-RU-8uM_4 / P3 / P4 / P5
CD107a-H
SSC-H (10^6)
R9
26.94%
PBMC_5-OP-RU-8uM_4 / P3 / P4 / P5
CD107b-H
SSC-H (10^6)
R11
10.23%
PBMC_5-OP-RU-8uM_4 / P3 / P4 / P5
CD137-H
SSC-H (10^6)
R10
10.53%
PBMC_5-OP-RU-8uM_4 / P3 / P4 / P5
IL-18.Ra-H
SSC-H (10^6)
R12
51.62%

Sample Statistics of PBMC_5-OP-RU-8uM_4

| Gate | Count | % Parent | X | Y | Median X | Median Y |
|---|---|---|---|---|---|---|
| All | 1,426,487 | | | | | |
| P1 | 38,330 | 2.69% | FSC-A | FSC-H | 1,050,687 | 534,440 |
| P2 | 29,488 | 76.93% | SSC-H | Live Dead-H | 112,987 | 6,486 |
| P3 | 22,076 | 1.55% | SSC-H | FSC-H | 194,702 | 1,013,852 |
| P4 | 7,472 | 33.85% | SSC-H | CD3-H | 138,011 | 12,366 |
| P5 | 1,329 | 17.79% | SSC-H | TCR-Va7.2-H | 209,283 | 3,987 |
| R6 | 741 | 55.76% | SSC-H | CD25-H | 232,335 | 6,008 |
| R7 | 69 | 5.19% | SSC-H | CD71-H | 365,825 | 11,885 |
| R8 | 652 | 49.06% | SSC-H | CD69-H | 239,030 | 7,261 |
| R9 | 358 | 26.94% | SSC-H | CD107a-H | 297,335 | 74,597 |
| R10 | 140 | 10.53% | SSC-H | CD137-H | 364,671 | 10,602 |
| R11 | 136 | 10.23% | SSC-H | CD107b-H | 397,808 | 80,248 |
| R12 | 686 | 51.62% | SSC-H | IL-18.Ra-H | 228,150 | 21,839 |

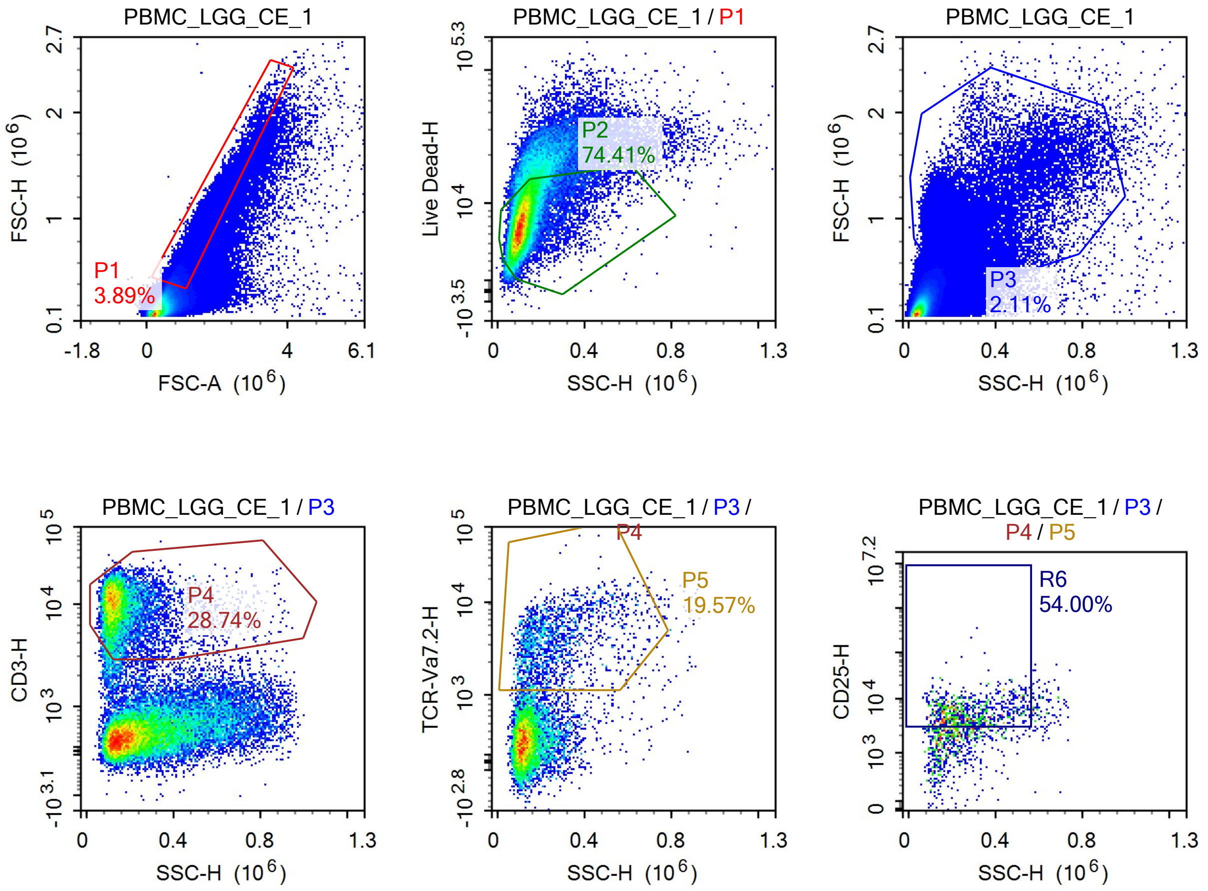

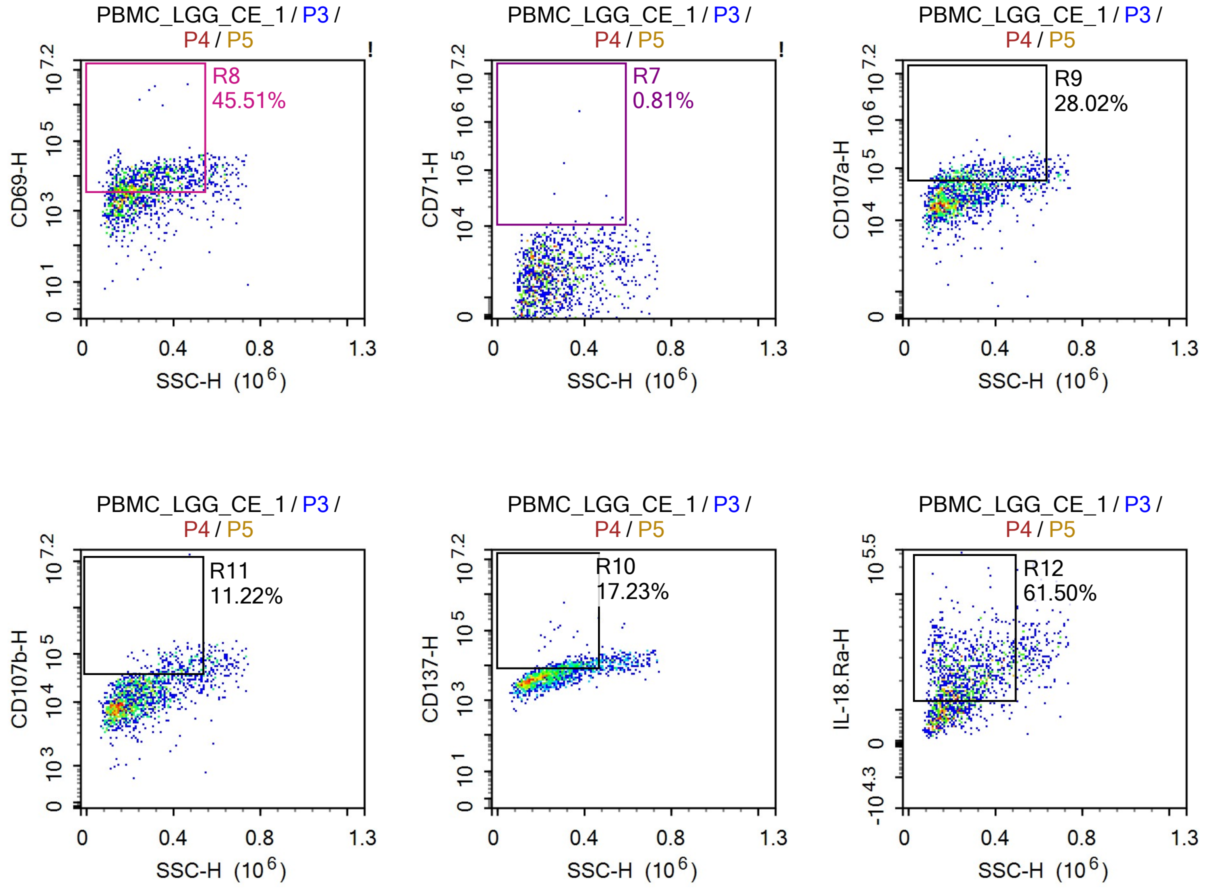


Sample Statistics of PBMC_LGG_CE_1

| Gate | Count | % Parent | X | Y | Median X | Median Y |
|---|---|---|---|---|---|---|
| All | 1,356,889 | | | | | |
| P1 | 52,799 | 3.89% | FSC-A | FSC-H | 1,033,187 | 505,238 |
| P2 | 39,290 | 74.41% | SSC-H | Live Dead-H | 111,967 | 7,458 |
| P3 | 28,684 | 2.11% | SSC-H | FSC-H | 201,735 | 961,711 |
| P4 | 8,243 | 28.74% | SSC-H | CD3-H | 137,616 | 9,614 |
| P5 | 1,613 | 19.57% | SSC-H | TCR-Va7.2-H | 232,277 | 3,520 |
| R6 | 871 | 54.00% | SSC-H | CD25-H | 259,447 | 5,398 |
| R7 | 13 | 0.81% | SSC-H | CD71-H | 416,473 | 13,403 |
| R8 | 734 | 45.51% | SSC-H | CD69-H | 268,451 | 8,116 |
| R9 | 452 | 28.02% | SSC-H | CD107a-H | 320,684 | 75,155 |
| R10 | 278 | 17.23% | SSC-H | CD137-H | 356,073 | 10,127 |
| R11 | 181 | 11.22% | SSC-H | CD107b-H | 416,318 | 58,835 |
| R12 | 992 | 61.50% | SSC-H | IL-18.Ra-H | 245,212 | 22,724 |

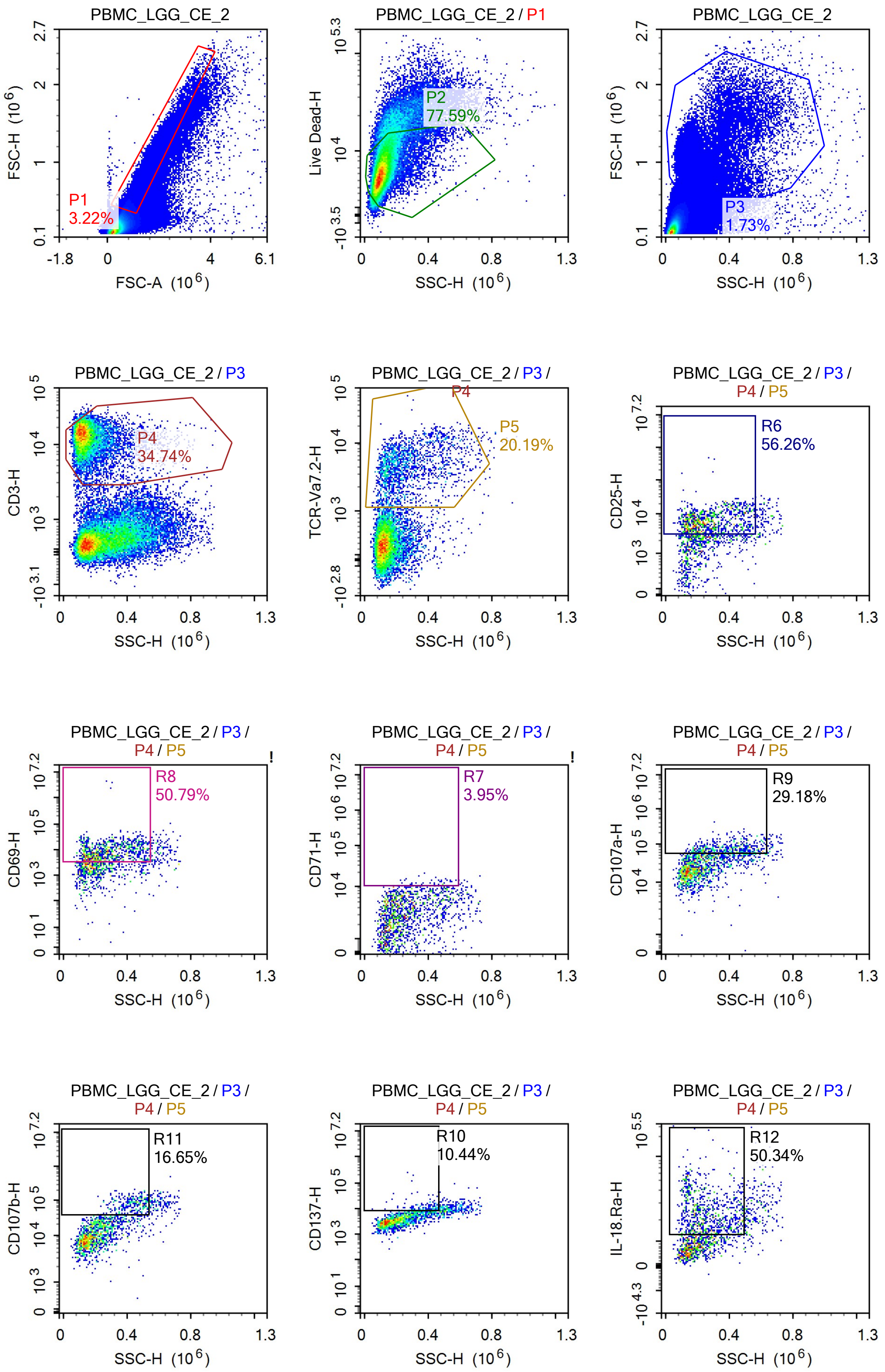
PBMC_LGG_CE_2
P1
3.22%
FSC-H (10^6)
FSC-A (10^6)
PBMC_LGG_CE_2 / P1
P2
77.59%
Live Dead-H
SSC-H (10^6)
PBMC_LGG_CE_2
P3
1.73%
FSC-H (10^6)
SSC-H (10^6)
PBMC_LGG_CE_2 / P3
P4
34.74%
CD3-H
SSC-H (10^6)
PBMC_LGG_CE_2 / P3 / P4
P5
20.19%
TCR-Va7.2-H
SSC-H (10^6)
PBMC_LGG_CE_2 / P3 / P4 / P5
R6
56.26%
CD25-H
SSC-H (10^6)
PBMC_LGG_CE_2 / P3 / P4 / P5
R8
50.79%
CD69-H
SSC-H (10^6)
PBMC_LGG_CE_2 / P3 / P4 / P5
R7
3.95%
CD71-H
SSC-H (10^6)
PBMC_LGG_CE_2 / P3 / P4 / P5
R9
29.18%
CD107a-H
SSC-H (10^6)
PBMC_LGG_CE_2 / P3 / P4 / P5
R11
16.65%
CD107b-H
SSC-H (10^6)
PBMC_LGG_CE_2 / P3 / P4 / P5
R10
10.44%
CD137-H
SSC-H (10^6)
PBMC_LGG_CE_2 / P3 / P4 / P5
R12
50.34%
IL-18.Ra-H
SSC-H (10^6)

Sample Statistics of PBMC_LGG_CE_2

| Gate | Count | % Parent | X | Y | Median X | Median Y |
|---|---|---|---|---|---|---|
| All | 1,459,237 | | | | | |
| P1 | 47,023 | 3.22% | FSC-A | FSC-H | 1,028,974 | 510,880 |
| P2 | 36,484 | 77.59% | SSC-H | Live Dead-H | 108,284 | 6,381 |
| P3 | 25,262 | 1.73% | SSC-H | FSC-H | 189,458 | 1,024,603 |
| P4 | 8,775 | 34.74% | SSC-H | CD3-H | 135,613 | 13,359 |
| P5 | 1,772 | 20.19% | SSC-H | TCR-Va7.2-H | 228,394 | 4,174 |
| R6 | 997 | 56.26% | SSC-H | CD25-H | 248,570 | 6,101 |
| R7 | 70 | 3.95% | SSC-H | CD71-H | 385,804 | 10,949 |
| R8 | 900 | 50.79% | SSC-H | CD69-H | 257,739 | 7,951 |
| R9 | 517 | 29.18% | SSC-H | CD107a-H | 388,081 | 71,632 |
| R10 | 185 | 10.44% | SSC-H | CD137-H | 371,264 | 9,320 |
| R11 | 295 | 16.65% | SSC-H | CD107b-H | 437,240 | 78,268 |
| R12 | 892 | 50.34% | SSC-H | IL-18.Ra-H | 242,549 | 22,286 |

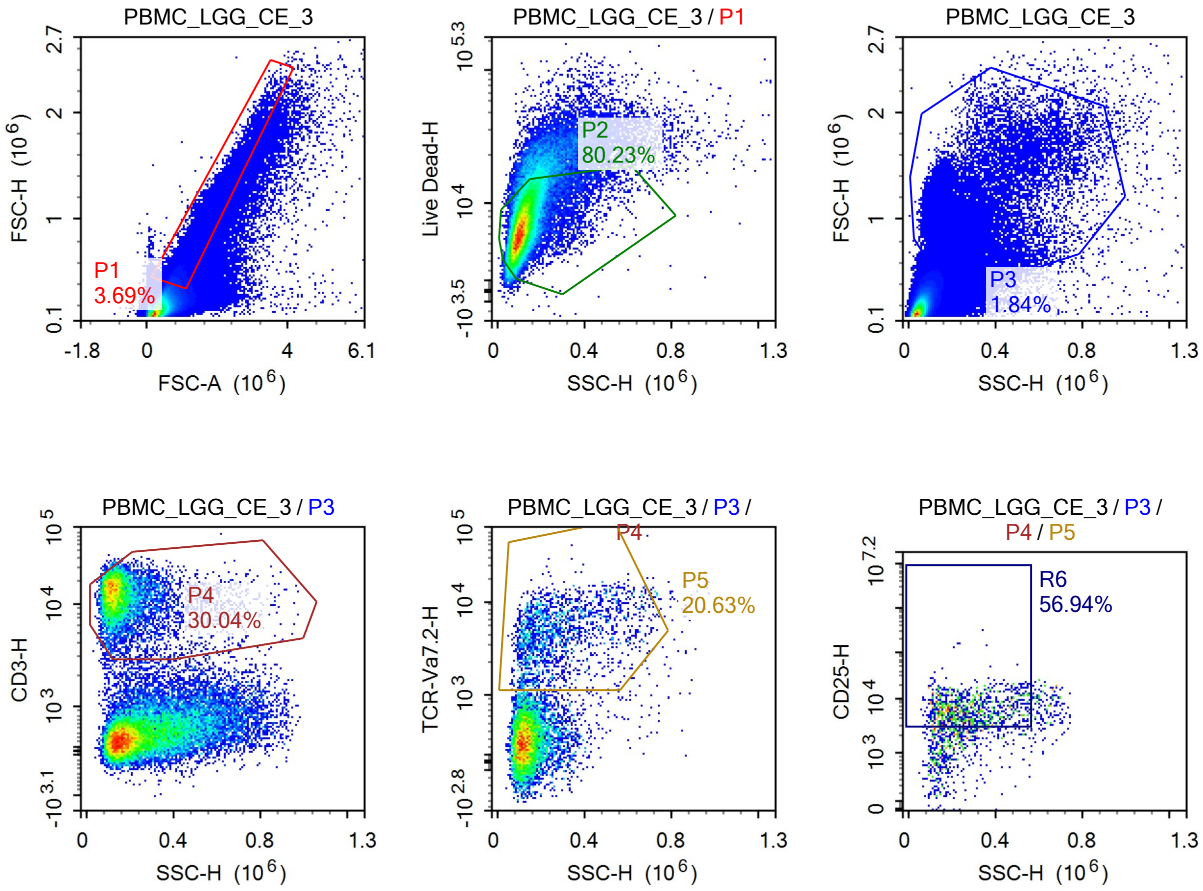

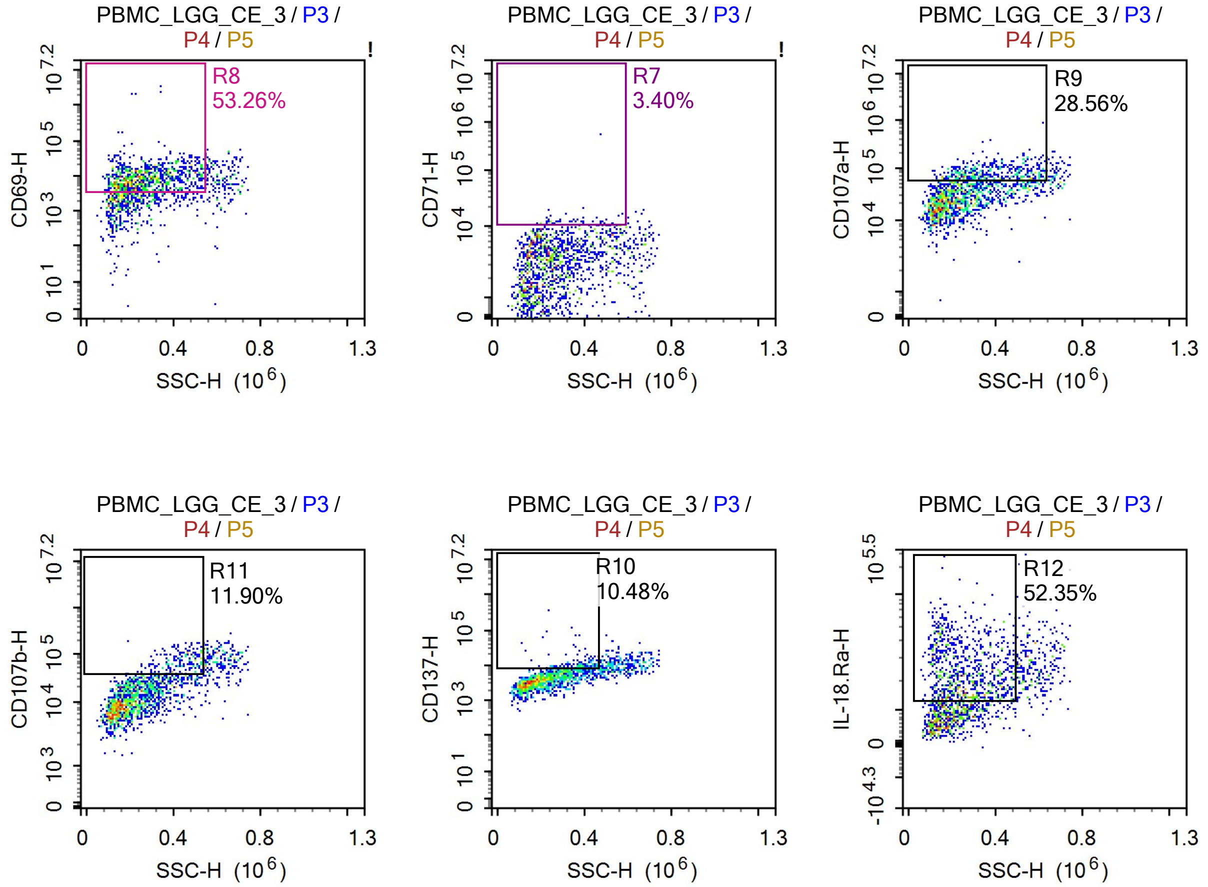


Sample Statistics of PBMC_LGG_CE_3

| Gate | Count | % Parent | X | Y | Median X | Median Y |
|---|---|---|---|---|---|---|
| All | 1,544,283 | | | | | |
| P1 | 56,961 | 3.69% | FSC-A | FSC-H | 1,001,051 | 493,706 |
| P2 | 45,701 | 80.23% | SSC-H | Live Dead-H | 111,200 | 6,855 |
| P3 | 28,473 | 1.84% | SSC-H | FSC-H | 192,141 | 963,794 |
| P4 | 8,554 | 30.04% | SSC-H | CD3-H | 137,900 | 12,974 |
| P5 | 1,765 | 20.63% | SSC-H | TCR-Va7.2-H | 241,230 | 4,288 |
| R6 | 1,005 | 56.94% | SSC-H | CD25-H | 261,272 | 6,265 |
| R7 | 60 | 3.40% | SSC-H | CD71-H | 387,281 | 11,803 |
| R8 | 940 | 53.26% | SSC-H | CD69-H | 258,529 | 8,679 |
| R9 | 504 | 28.56% | SSC-H | CD107a-H | 360,482 | 73,682 |
| R10 | 185 | 10.48% | SSC-H | CD137-H | 371,748 | 9,528 |
| R11 | 210 | 11.90% | SSC-H | CD107b-H | 432,002 | 67,874 |
| R12 | 924 | 52.35% | SSC-H | IL-18.Ra-H | 244,086 | 22,310 |

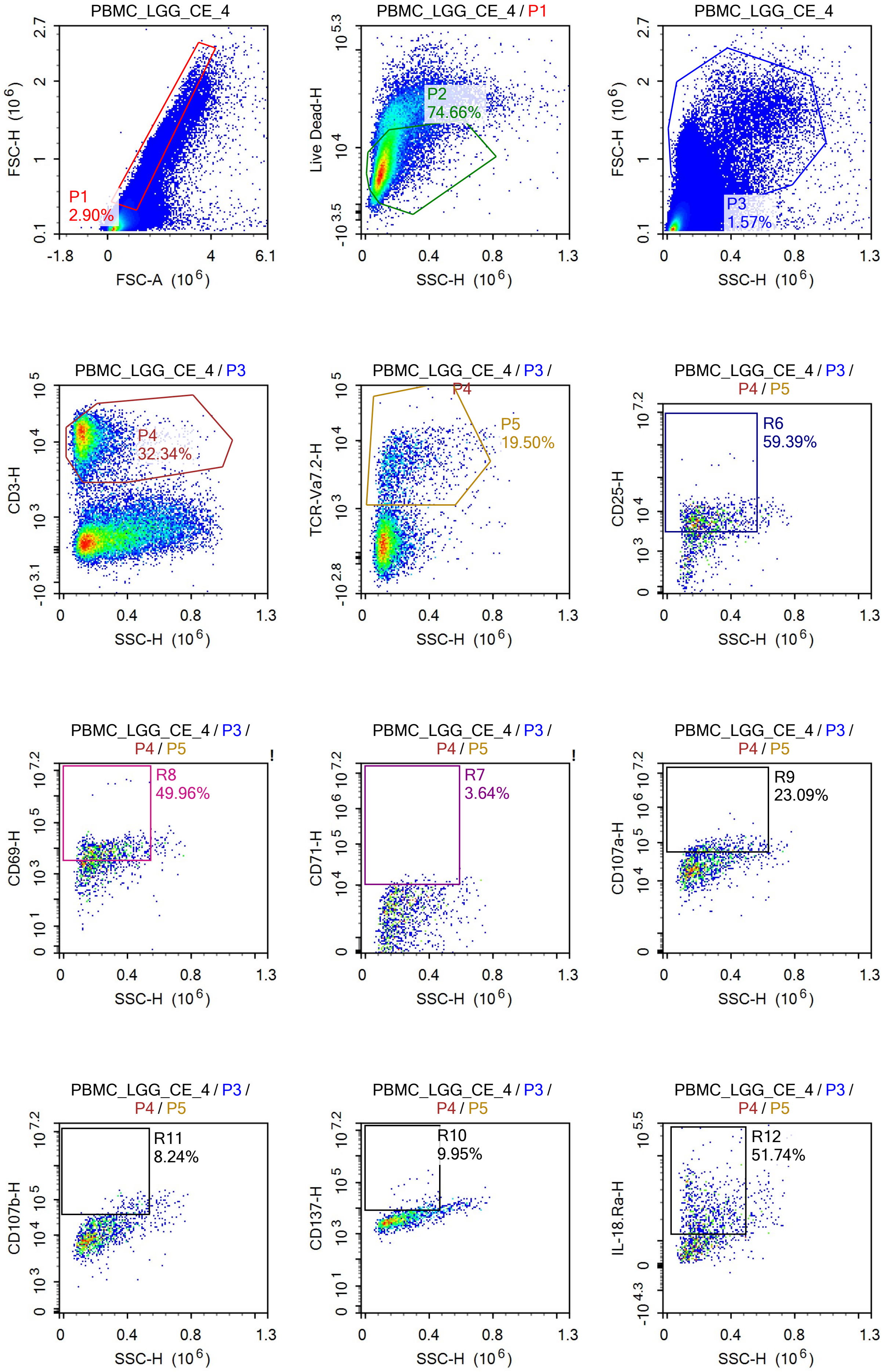
PBMC_LGG_CE_4
P1
2.90%
FSC-H (10^6)
FSC-A (10^6)
PBMC_LGG_CE_4 / P1
P2
74.66%
Live Dead-H
SSC-H (10^6)
PBMC_LGG_CE_4
P3
1.57%
FSC-H (10^6)
SSC-H (10^6)
PBMC_LGG_CE_4 / P3
P4
32.34%
CD3-H
SSC-H (10^6)
PBMC_LGG_CE_4 / P3 /
P4
P5
19.50%
TCR-Va7.2-H
SSC-H (10^6)
PBMC_LGG_CE_4 / P3 / P4 / P5
R6
59.39%
CD25-H
SSC-H (10^6)
PBMC_LGG_CE_4 / P3 / P4 / P5
R8
49.96%
CD69-H
SSC-H (10^6)
PBMC_LGG_CE_4 / P3 / P4 / P5
R7
3.64%
CD71-H
SSC-H (10^6)
PBMC_LGG_CE_4 / P3 / P4 / P5
R9
23.09%
CD107a-H
SSC-H (10^6)
PBMC_LGG_CE_4 / P3 / P4 / P5
R11
8.24%
CD107b-H
SSC-H (10^6)
PBMC_LGG_CE_4 / P3 / P4 / P5
R10
9.95%
CD137-H
SSC-H (10^6)
PBMC_LGG_CE_4 / P3 / P4 / P5
R12
51.74%
IL-18.Ra-H
SSC-H (10^6)

Sample Statistics of PBMC_LGG_CE_4

| Gate | Count | % Parent | X | Y | Median X | Median Y |
|---|---|---|---|---|---|---|
| All | 1,361,752 | | | | | |
| P1 | 39,521 | 2.90% | FSC-A | FSC-H | 1,028,901 | 512,722 |
| P2 | 29,508 | 74.66% | SSC-H | Live Dead-H | 110,543 | 6,626 |
| P3 | 21,363 | 1.57% | SSC-H | FSC-H | 194,178 | 1,030,468 |
| P4 | 6,909 | 32.34% | SSC-H | CD3-H | 131,781 | 12,193 |
| P5 | 1,347 | 19.50% | SSC-H | TCR-Va7.2-H | 212,371 | 4,156 |
| R6 | 800 | 59.39% | SSC-H | CD25-H | 233,403 | 6,106 |
| R7 | 49 | 3.64% | SSC-H | CD71-H | 312,712 | 11,123 |
| R8 | 673 | 49.96% | SSC-H | CD69-H | 243,804 | 7,974 |
| R9 | 311 | 23.09% | SSC-H | CD107a-H | 310,368 | 70,165 |
| R10 | 134 | 9.95% | SSC-H | CD137-H | 367,985 | 9,913 |
| R11 | 111 | 8.24% | SSC-H | CD107b-H | 413,259 | 60,696 |
| R12 | 697 | 51.74% | SSC-H | IL-18.Ra-H | 234,608 | 22,728 |

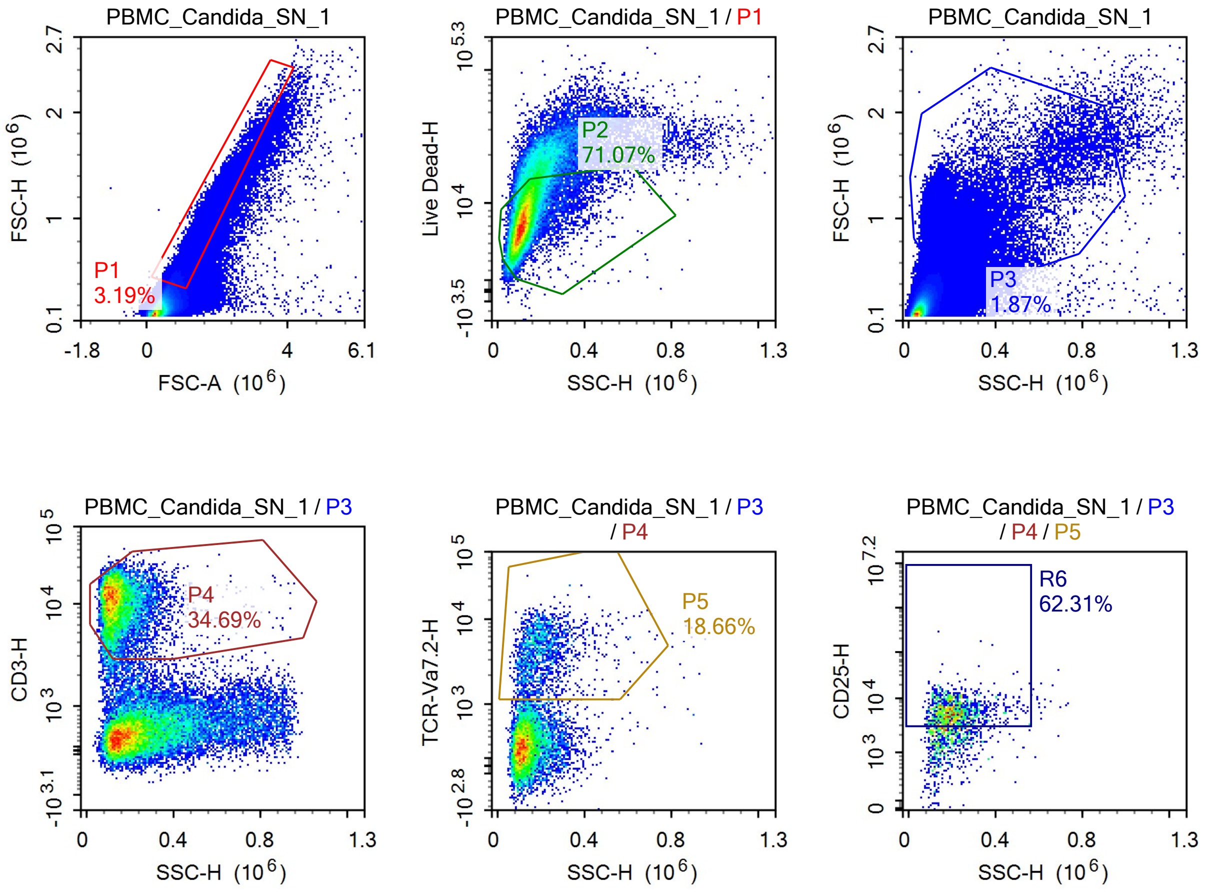

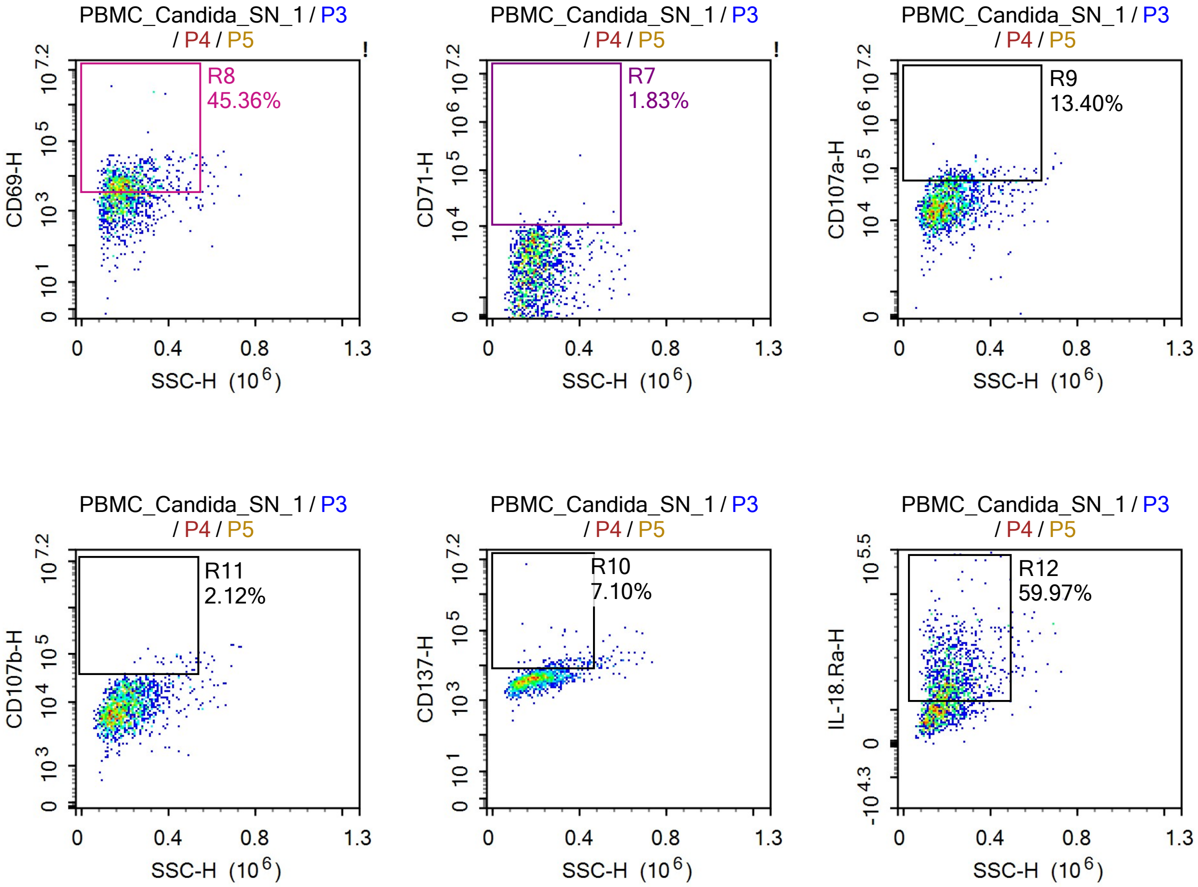


Sample Statistics of PBMC_Candida_SN_1

| Gate | Count | % Parent | X | Y | Median X | Median Y |
|---|---|---|---|---|---|---|
| All | 1,438,412 | | | | | |
| P1 | 45,877 | 3.19% | FSC-A | FSC-H | 1,082,465 | 556,281 |
| P2 | 32,604 | 71.07% | SSC-H | Live Dead-H | 117,114 | 7,623 |
| P3 | 26,968 | 1.87% | SSC-H | FSC-H | 186,591 | 978,540 |
| P4 | 9,355 | 34.69% | SSC-H | CD3-H | 132,041 | 10,808 |
| P5 | 1,746 | 18.66% | SSC-H | TCR-Va7.2-H | 189,047 | 3,909 |
| R6 | 1,088 | 62.31% | SSC-H | CD25-H | 200,699 | 5,480 |
| R7 | 32 | 1.83% | SSC-H | CD71-H | 270,098 | 10,844 |
| R8 | 792 | 45.36% | SSC-H | CD69-H | 202,468 | 6,411 |
| R9 | 234 | 13.40% | SSC-H | CD107a-H | 252,658 | 63,939 |
| R10 | 124 | 7.10% | SSC-H | CD137-H | 323,810 | 11,591 |
| R11 | 37 | 2.12% | SSC-H | CD107b-H | 418,688 | 67,879 |
| R12 | 1,047 | 59.97% | SSC-H | IL-18.Ra-H | 210,046 | 20,923 |

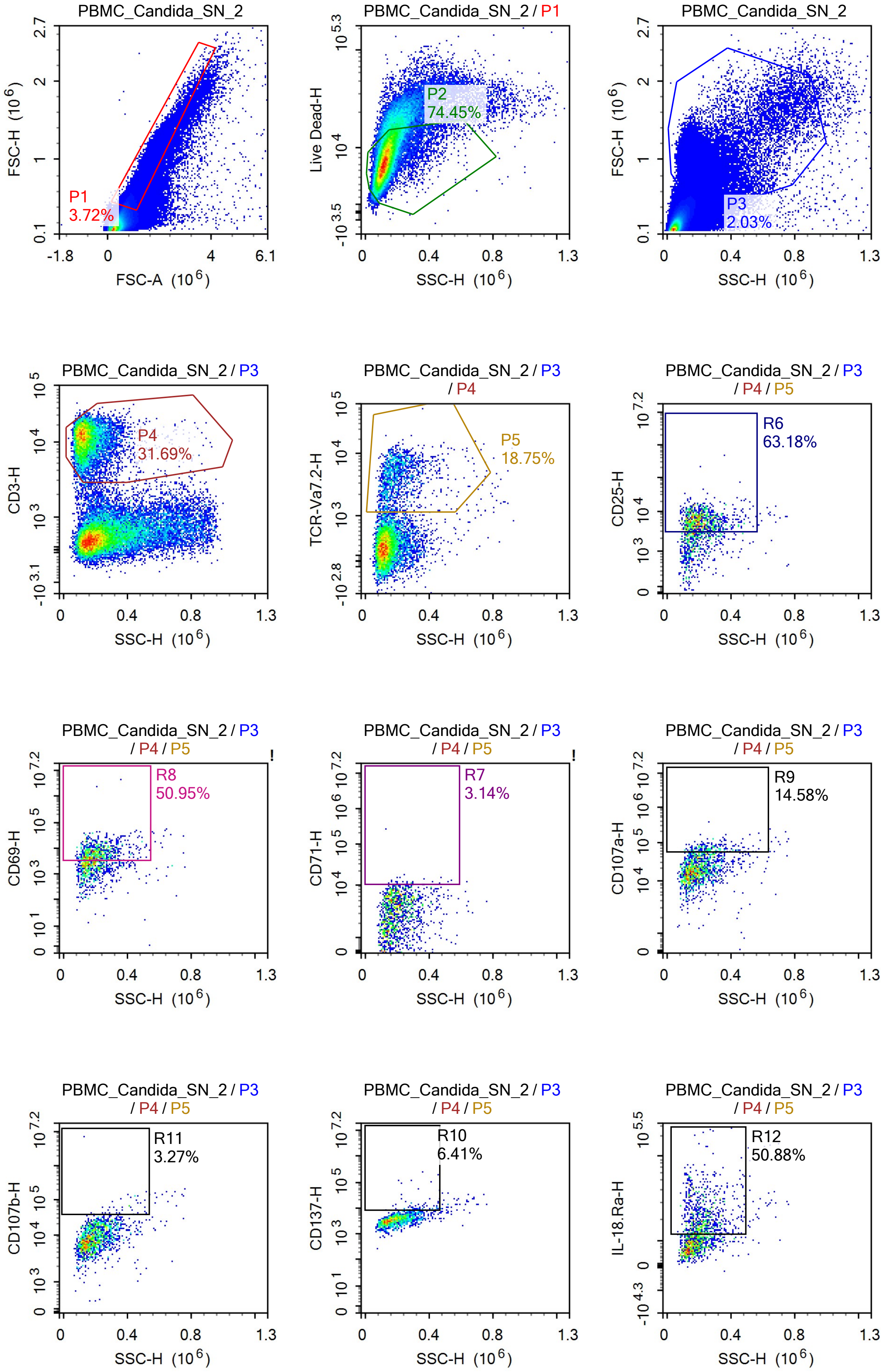
PBMC_Candida_SN_2
P1
3.72%
FSC-H (10^6)
FSC-A (10^6)
PBMC_Candida_SN_2 / P1
P2
74.45%
Live Dead-H
SSC-H (10^6)
PBMC_Candida_SN_2
P3
2.03%
FSC-H (10^6)
SSC-H (10^6)
PBMC_Candida_SN_2 / P3
P4
31.69%
CD3-H
SSC-H (10^6)
PBMC_Candida_SN_2 / P3 / P4
P5
18.75%
TCR-Va7.2-H
SSC-H (10^6)
PBMC_Candida_SN_2 / P3 / P4 / P5
R6
63.18%
CD25-H
SSC-H (10^6)
PBMC_Candida_SN_2 / P3 / P4 / P5
R8
50.95%
CD69-H
SSC-H (10^6)
PBMC_Candida_SN_2 / P3 / P4 / P5
R7
3.14%
CD71-H
SSC-H (10^6)
PBMC_Candida_SN_2 / P3 / P4 / P5
R9
14.58%
CD107a-H
SSC-H (10^6)
PBMC_Candida_SN_2 / P3 / P4 / P5
R11
3.27%
CD107b-H
SSC-H (10^6)
PBMC_Candida_SN_2 / P3 / P4 / P5
R10
6.41%
CD137-H
SSC-H (10^6)
PBMC_Candida_SN_2 / P3 / P4 / P5
R12
50.88%
IL-18.Ra-H
SSC-H (10^6)

Sample Statistics of PBMC_Candida_SN_2

| Gate | Count | % Parent | X | Y | Median X | Median Y |
|---|---|---|---|---|---|---|
| All | 1,267,702 | | | | | |
| P1 | 47,126 | 3.72% | FSC-A | FSC-H | 1,031,207 | 526,711 |
| P2 | 35,083 | 74.45% | SSC-H | Live Dead-H | 123,916 | 7,730 |
| P3 | 25,724 | 2.03% | SSC-H | FSC-H | 194,528 | 935,118 |
| P4 | 8,153 | 31.69% | SSC-H | CD3-H | 133,344 | 11,749 |
| P5 | 1,529 | 18.75% | SSC-H | TCR-Va7.2-H | 188,794 | 3,980 |
| R6 | 966 | 63.18% | SSC-H | CD25-H | 201,432 | 5,621 |
| R7 | 48 | 3.14% | SSC-H | CD71-H | 233,681 | 10,948 |
| R8 | 779 | 50.95% | SSC-H | CD69-H | 206,353 | 6,862 |
| R9 | 223 | 14.58% | SSC-H | CD107a-H | 256,055 | 67,157 |
| R10 | 98 | 6.41% | SSC-H | CD137-H | 331,762 | 10,024 |
| R11 | 50 | 3.27% | SSC-H | CD107b-H | 409,107 | 62,345 |
| R12 | 778 | 50.88% | SSC-H | IL-18.Ra-H | 216,517 | 20,160 |

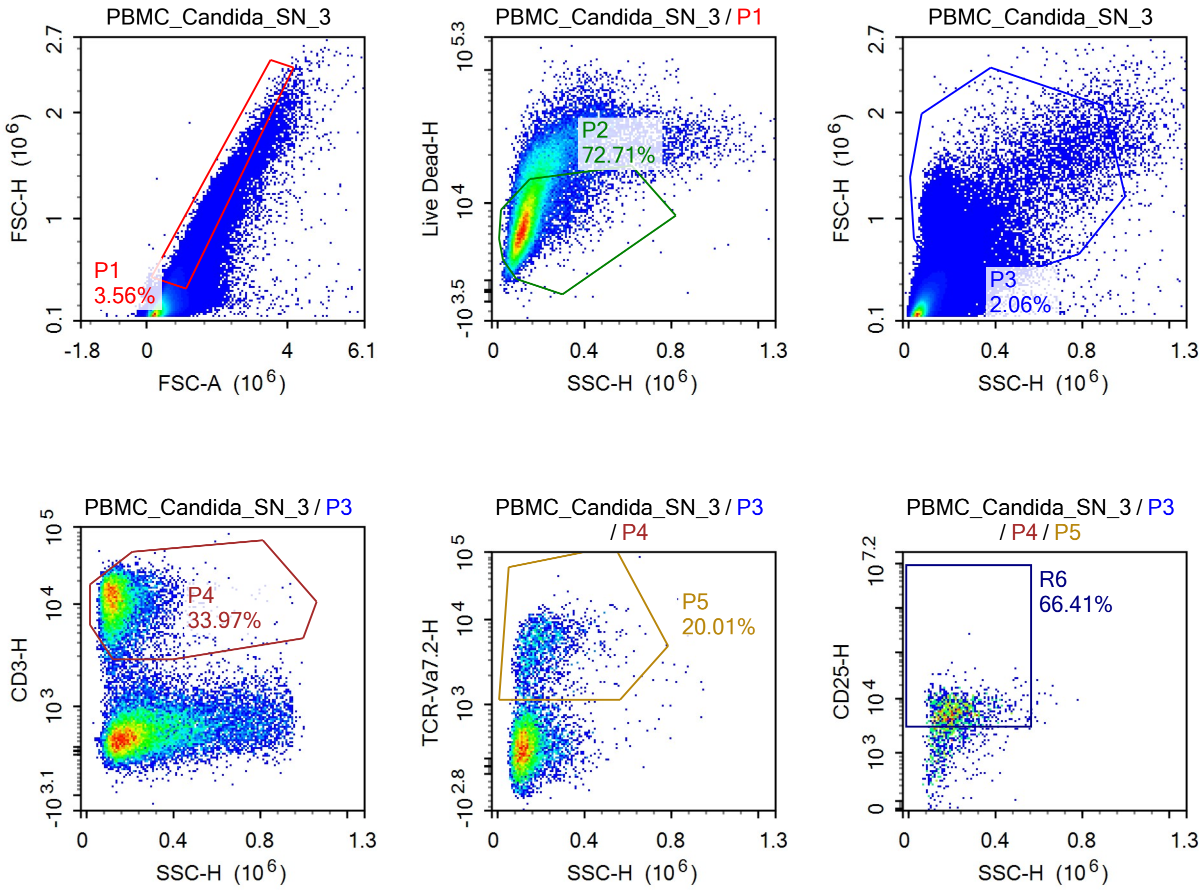

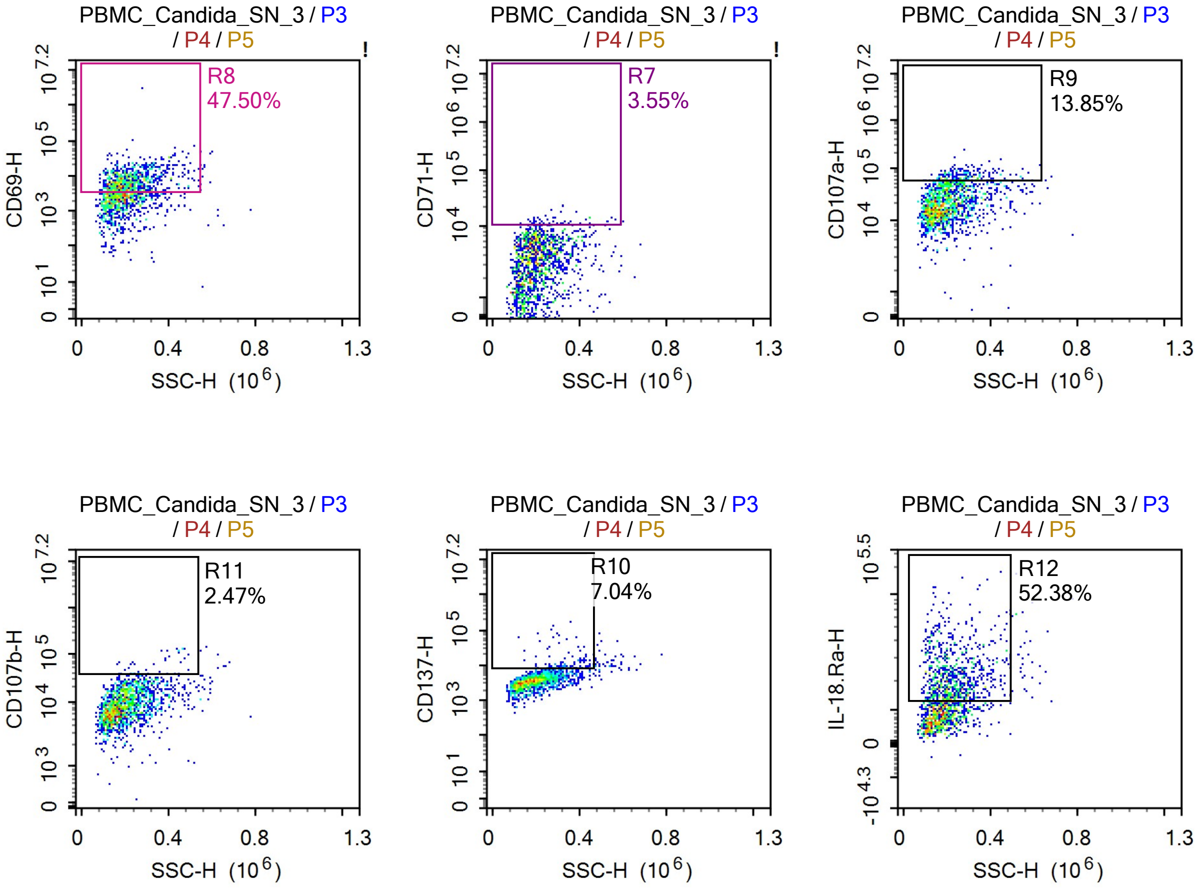


Sample Statistics of PBMC_Candida_SN_3

| Gate | Count | % Parent | X | Y | Median X | Median Y |
|---|---|---|---|---|---|---|
| All | 1,184,725 | | | | | |
| P1 | 42,211 | 3.56% | FSC-A | FSC-H | 1,065,805 | 546,428 |
| P2 | 30,692 | 72.71% | SSC-H | Live Dead-H | 123,219 | 7,449 |
| P3 | 24,436 | 2.06% | SSC-H | FSC-H | 191,455 | 959,176 |
| P4 | 8,301 | 33.97% | SSC-H | CD3-H | 132,882 | 11,220 |
| P5 | 1,661 | 20.01% | SSC-H | TCR-Va7.2-H | 191,027 | 4,272 |
| R6 | 1,103 | 66.41% | SSC-H | CD25-H | 204,556 | 5,799 |
| R7 | 59 | 3.55% | SSC-H | CD71-H | 259,801 | 11,681 |
| R8 | 789 | 47.50% | SSC-H | CD69-H | 212,422 | 7,169 |
| R9 | 230 | 13.85% | SSC-H | CD107a-H | 257,971 | 66,012 |
| R10 | 117 | 7.04% | SSC-H | CD137-H | 328,378 | 10,266 |
| R11 | 41 | 2.47% | SSC-H | CD107b-H | 332,680 | 59,129 |
| R12 | 870 | 52.38% | SSC-H | IL-18.Ra-H | 214,131 | 20,458 |

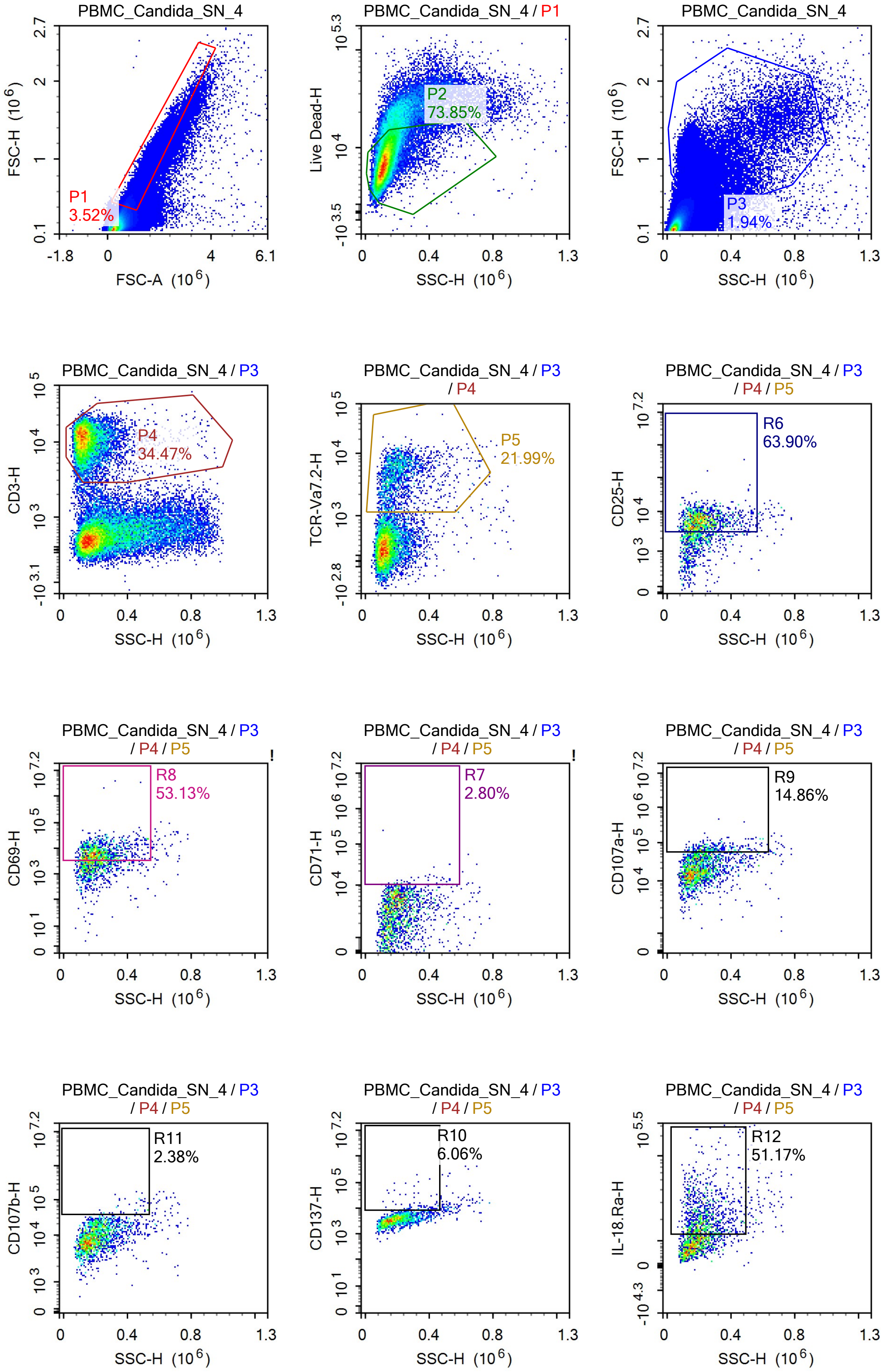
PBMC_Candida_SN_4
P1
3.52%
FSC-H (10^6)
FSC-A (10^6)
PBMC_Candida_SN_4 / P1
P2
73.85%
Live Dead-H
SSC-H (10^6)
PBMC_Candida_SN_4
P3
1.94%
FSC-H (10^6)
SSC-H (10^6)
PBMC_Candida_SN_4 / P3
P4
34.47%
CD3-H
SSC-H (10^6)
PBMC_Candida_SN_4 / P3 / P4
P5
21.99%
TCR-Va7.2-H
SSC-H (10^6)
PBMC_Candida_SN_4 / P3 / P4 / P5
R6
63.90%
CD25-H
SSC-H (10^6)
PBMC_Candida_SN_4 / P3 / P4 / P5
R8
53.13%
CD69-H
SSC-H (10^6)
PBMC_Candida_SN_4 / P3 / P4 / P5
R7
2.80%
CD71-H
SSC-H (10^6)
PBMC_Candida_SN_4 / P3 / P4 / P5
R9
14.86%
CD107a-H
SSC-H (10^6)
PBMC_Candida_SN_4 / P3 / P4 / P5
R11
2.38%
CD107b-H
SSC-H (10^6)
PBMC_Candida_SN_4 / P3 / P4 / P5
R10
6.06%
CD137-H
SSC-H (10^6)
PBMC_Candida_SN_4 / P3 / P4 / P5
R12
51.17%
IL-18.Ra-H
SSC-H (10^6)

Sample Statistics of PBMC_Candida_SN_4

| Gate | Count | % Parent | X | Y | Median X | Median Y |
|---|---|---|---|---|---|---|
| All | 1,315,223 | | | | | |
| P1 | 46,349 | 3.52% | FSC-A | FSC-H | 1,037,458 | 531,135 |
| P2 | 34,228 | 73.85% | SSC-H | Live Dead-H | 120,424 | 7,426 |
| P3 | 25,478 | 1.94% | SSC-H | FSC-H | 189,138 | 957,530 |
| P4 | 8,782 | 34.47% | SSC-H | CD3-H | 134,255 | 11,210 |
| P5 | 1,931 | 21.99% | SSC-H | TCR-Va7.2-H | 195,023 | 4,329 |
| R6 | 1,234 | 63.90% | SSC-H | CD25-H | 208,417 | 5,720 |
| R7 | 54 | 2.80% | SSC-H | CD71-H | 275,092 | 11,370 |
| R8 | 1,026 | 53.13% | SSC-H | CD69-H | 208,859 | 6,677 |
| R9 | 287 | 14.86% | SSC-H | CD107a-H | 272,521 | 67,971 |
| R10 | 117 | 6.06% | SSC-H | CD137-H | 326,729 | 9,864 |
| R11 | 46 | 2.38% | SSC-H | CD107b-H | 450,602 | 58,990 |
| R12 | 988 | 51.17% | SSC-H | IL-18.Ra-H | 219,200 | 20,393 |

de resultaten van deze readout staan op het einde !!

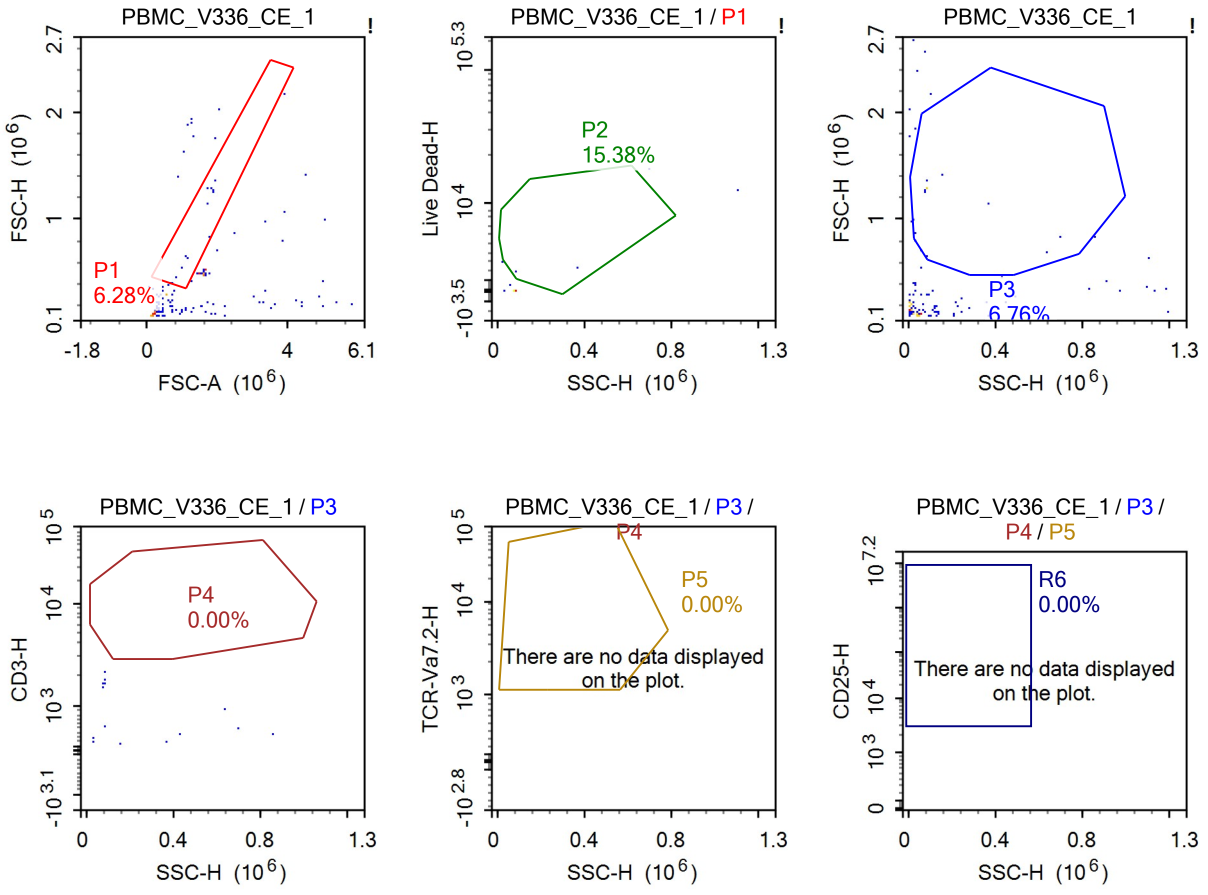

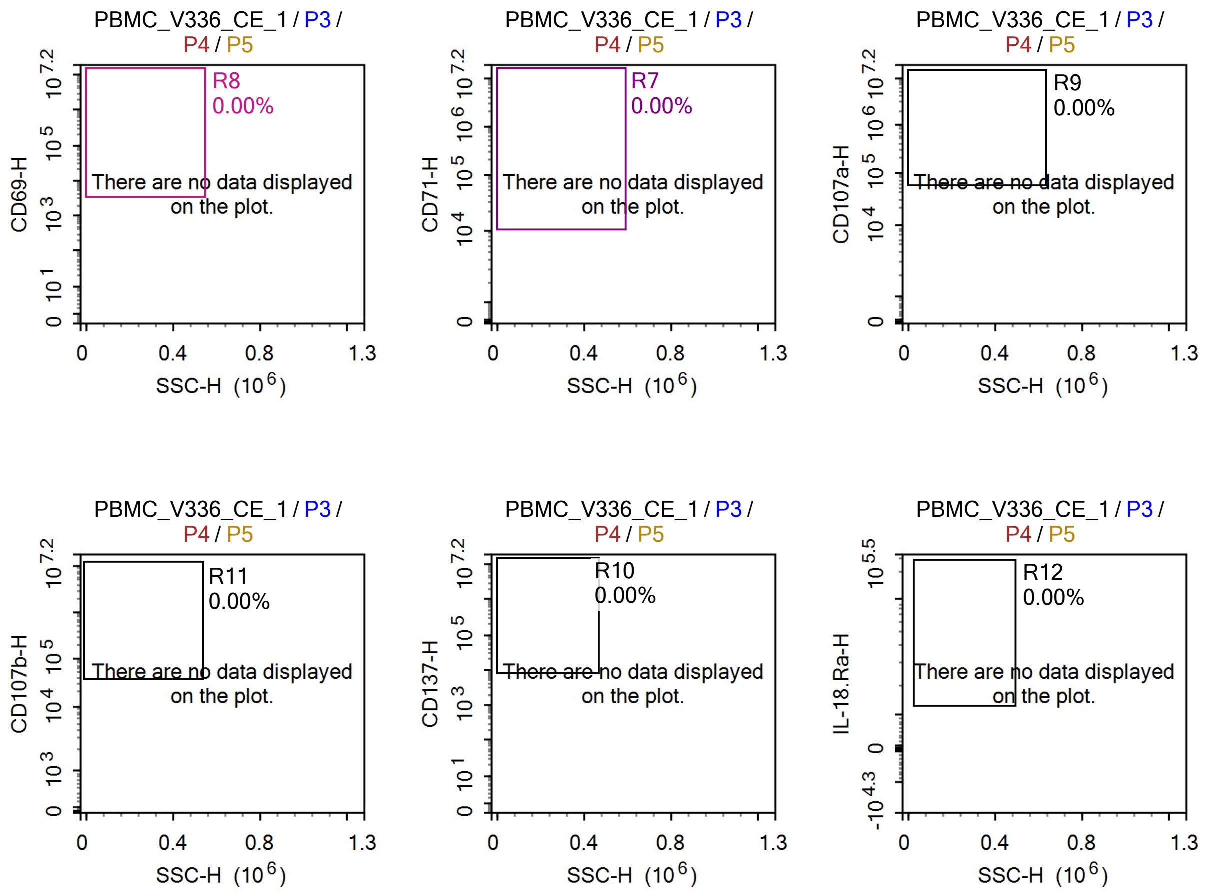


Sample Statistics of PBMC_V336_CE_1

| Gate | Count | % Parent | X | Y | Median X | Median Y |
|---|---|---|---|---|---|---|
| All | 207 | | | | | |
| P1 | 13 | 6.28% | FSC-A | FSC-H | 1,648,905 | 842,738 |
| P2 | 2 | 15.38% | SSC-H | Live Dead-H | 219,988 | 2,274 |
| P3 | 14 | 6.76% | SSC-H | FSC-H | 82,830 | 1,009,896 |
| P4 | 0 | 0.00% | SSC-H | CD3-H | 0 | 0 |
| P5 | 0 | 0.00% | SSC-H | TCR-Va7.2-H | 0 | 0 |
| R6 | 0 | 0.00% | SSC-H | CD25-H | 0 | 0 |
| R7 | 0 | 0.00% | SSC-H | CD71-H | 0 | 0 |
| R8 | 0 | 0.00% | SSC-H | CD69-H | 0 | 0 |
| R9 | 0 | 0.00% | SSC-H | CD107a-H | 0 | 0 |
| R10 | 0 | 0.00% | SSC-H | CD137-H | 0 | 0 |
| R11 | 0 | 0.00% | SSC-H | CD107b-H | 0 | 0 |
| R12 | 0 | 0.00% | SSC-H | IL-18.Ra-H | 0 | 0 |

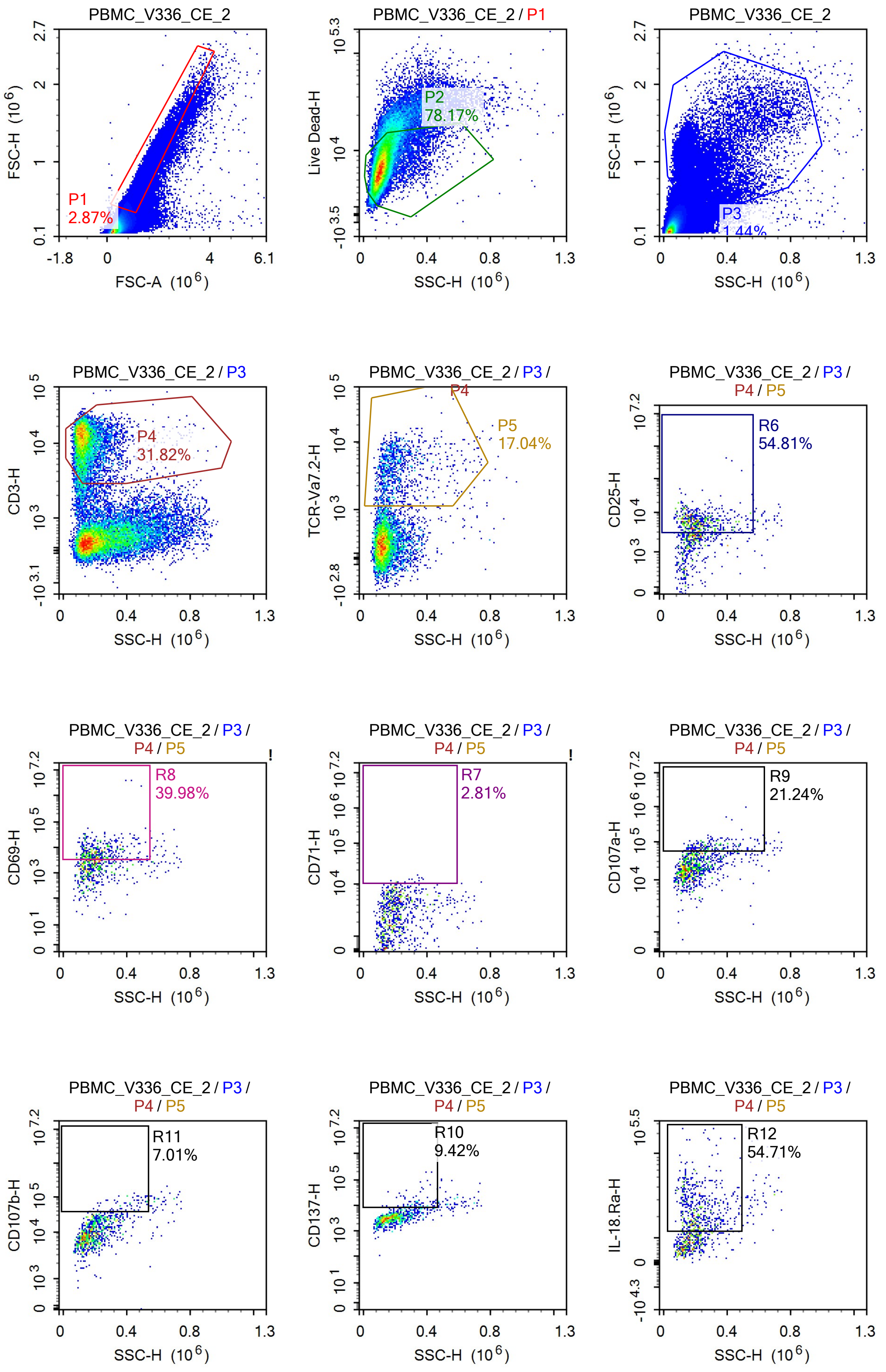
PBMC_V336_CE_2
FSC-H (10^6)
FSC-A (10^6)
P1
2.87%
PBMC_V336_CE_2 / P1
Live Dead-H
SSC-H (10^6)
P2
78.17%
PBMC_V336_CE_2
FSC-H (10^6)
SSC-H (10^6)
P3
1.44%
PBMC_V336_CE_2 / P3
CD3-H
SSC-H (10^6)
P4
31.82%
PBMC_V336_CE_2 / P3 / P4
TCR-Va7.2-H
SSC-H (10^6)
P5
17.04%
PBMC_V336_CE_2 / P3 / P4 / P5
CD25-H
SSC-H (10^6)
R6
54.81%
PBMC_V336_CE_2 / P3 / P4 / P5
CD69-H
SSC-H (10^6)
R8
39.98%
PBMC_V336_CE_2 / P3 / P4 / P5
CD71-H
SSC-H (10^6)
R7
2.81%
PBMC_V336_CE_2 / P3 / P4 / P5
CD107a-H
SSC-H (10^6)
R9
21.24%
PBMC_V336_CE_2 / P3 / P4 / P5
CD107b-H
SSC-H (10^6)
R11
7.01%
PBMC_V336_CE_2 / P3 / P4 / P5
CD137-H
SSC-H (10^6)
R10
9.42%
PBMC_V336_CE_2 / P3 / P4 / P5
IL-18.Ra-H
SSC-H (10^6)
R12
54.71%

Sample Statistics of PBMC_V336_CE_2

| Gate | Count | % Parent | X | Y | Median X | Median Y |
|---|---|---|---|---|---|---|
| All | 1,274,696 | | | | | |
| P1 | 36,590 | 2.87% | FSC-A | FSC-H | 1,010,239 | 509,907 |
| P2 | 28,604 | 78.17% | SSC-H | Live Dead-H | 113,651 | 7,168 |
| P3 | 18,407 | 1.44% | SSC-H | FSC-H | 180,205 | 993,974 |
| P4 | 5,858 | 31.82% | SSC-H | CD3-H | 131,391 | 11,207 |
| P5 | 998 | 17.04% | SSC-H | TCR-Va7.2-H | 189,132 | 3,717 |
| R6 | 547 | 54.81% | SSC-H | CD25-H | 206,240 | 5,759 |
| R7 | 28 | 2.81% | SSC-H | CD71-H | 258,538 | 11,205 |
| R8 | 399 | 39.98% | SSC-H | CD69-H | 209,723 | 6,549 |
| R9 | 212 | 21.24% | SSC-H | CD107a-H | 273,337 | 72,102 |
| R10 | 94 | 9.42% | SSC-H | CD137-H | 336,627 | 11,215 |
| R11 | 70 | 7.01% | SSC-H | CD107b-H | 401,988 | 63,974 |
| R12 | 546 | 54.71% | SSC-H | IL-18.Ra-H | 203,491 | 22,826 |

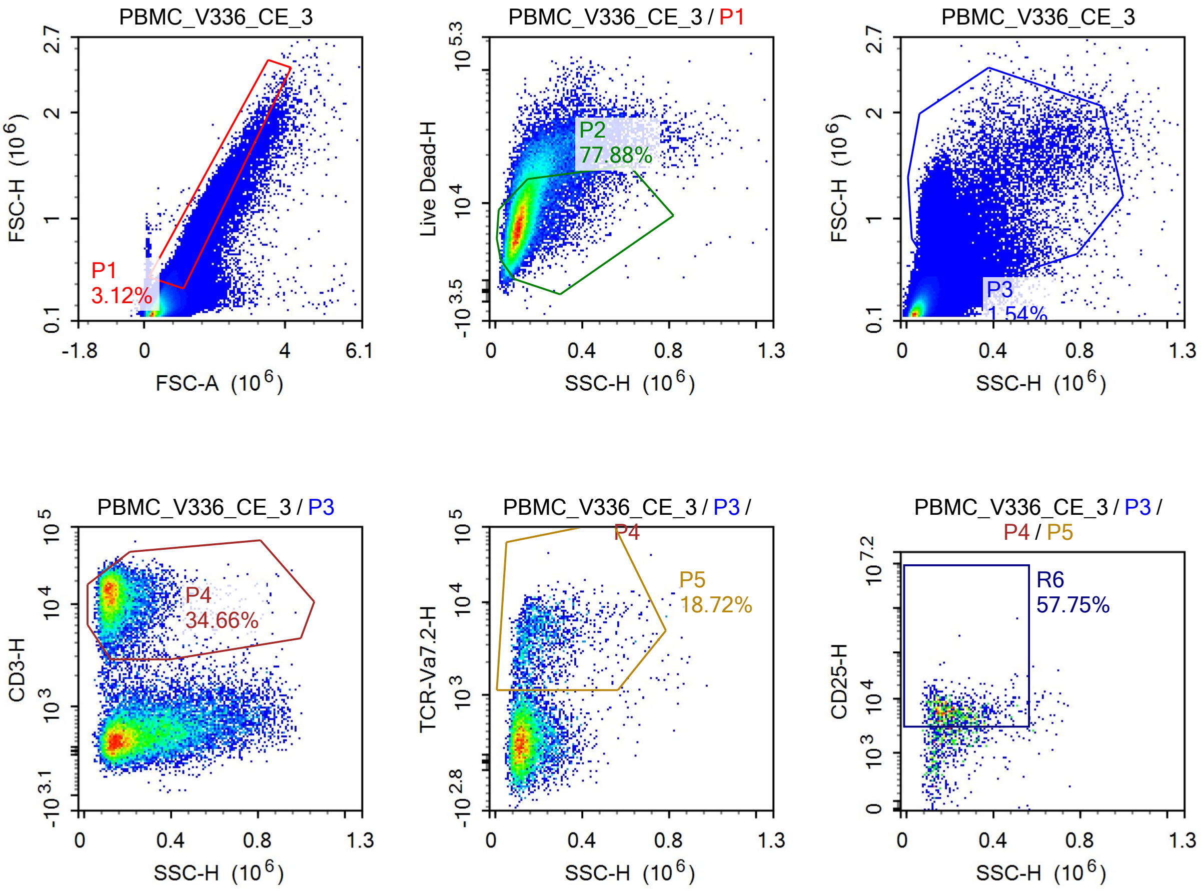

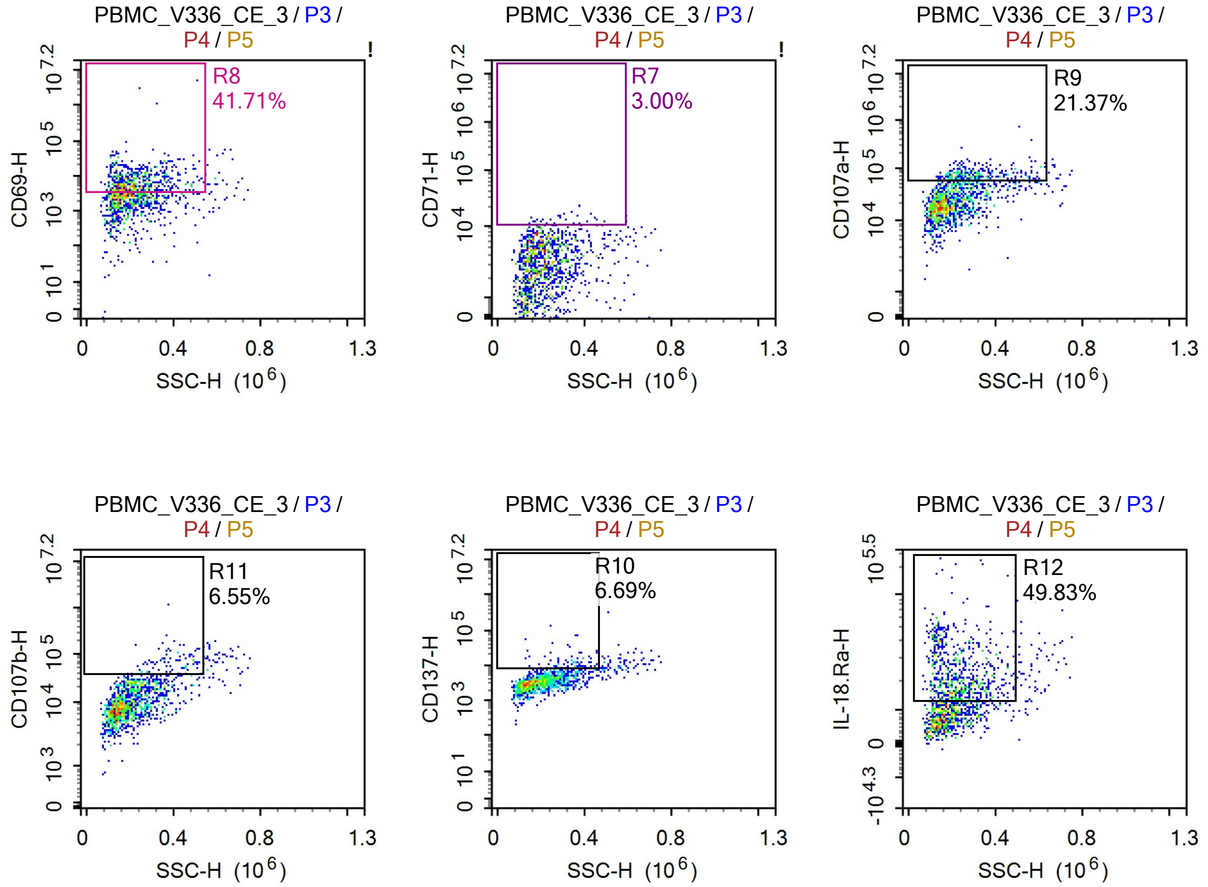


Sample Statistics of PBMC_V336_CE_3

| Gate | Count | % Parent | X | Y | Median X | Median Y |
|---|---|---|---|---|---|---|
| All | 1,467,940 | | | | | |
| P1 | 45,751 | 3.12% | FSC-A | FSC-H | 1,005,883 | 502,673 |
| P2 | 35,633 | 77.88% | SSC-H | Live Dead-H | 113,893 | 7,428 |
| P3 | 22,583 | 1.54% | SSC-H | FSC-H | 182,078 | 994,635 |
| P4 | 7,827 | 34.66% | SSC-H | CD3-H | 134,327 | 12,234 |
| P5 | 1,465 | 18.72% | SSC-H | TCR-Va7.2-H | 193,136 | 3,781 |
| R6 | 846 | 57.75% | SSC-H | CD25-H | 204,753 | 5,571 |
| R7 | 44 | 3.00% | SSC-H | CD71-H | 281,285 | 11,755 |
| R8 | 611 | 41.71% | SSC-H | CD69-H | 209,170 | 6,263 |
| R9 | 313 | 21.37% | SSC-H | CD107a-H | 281,944 | 71,353 |
| R10 | 98 | 6.69% | SSC-H | CD137-H | 347,601 | 11,225 |
| R11 | 96 | 6.55% | SSC-H | CD107b-H | 388,038 | 65,986 |
| R12 | 730 | 49.83% | SSC-H | IL-18.Ra-H | 214,808 | 20,810 |

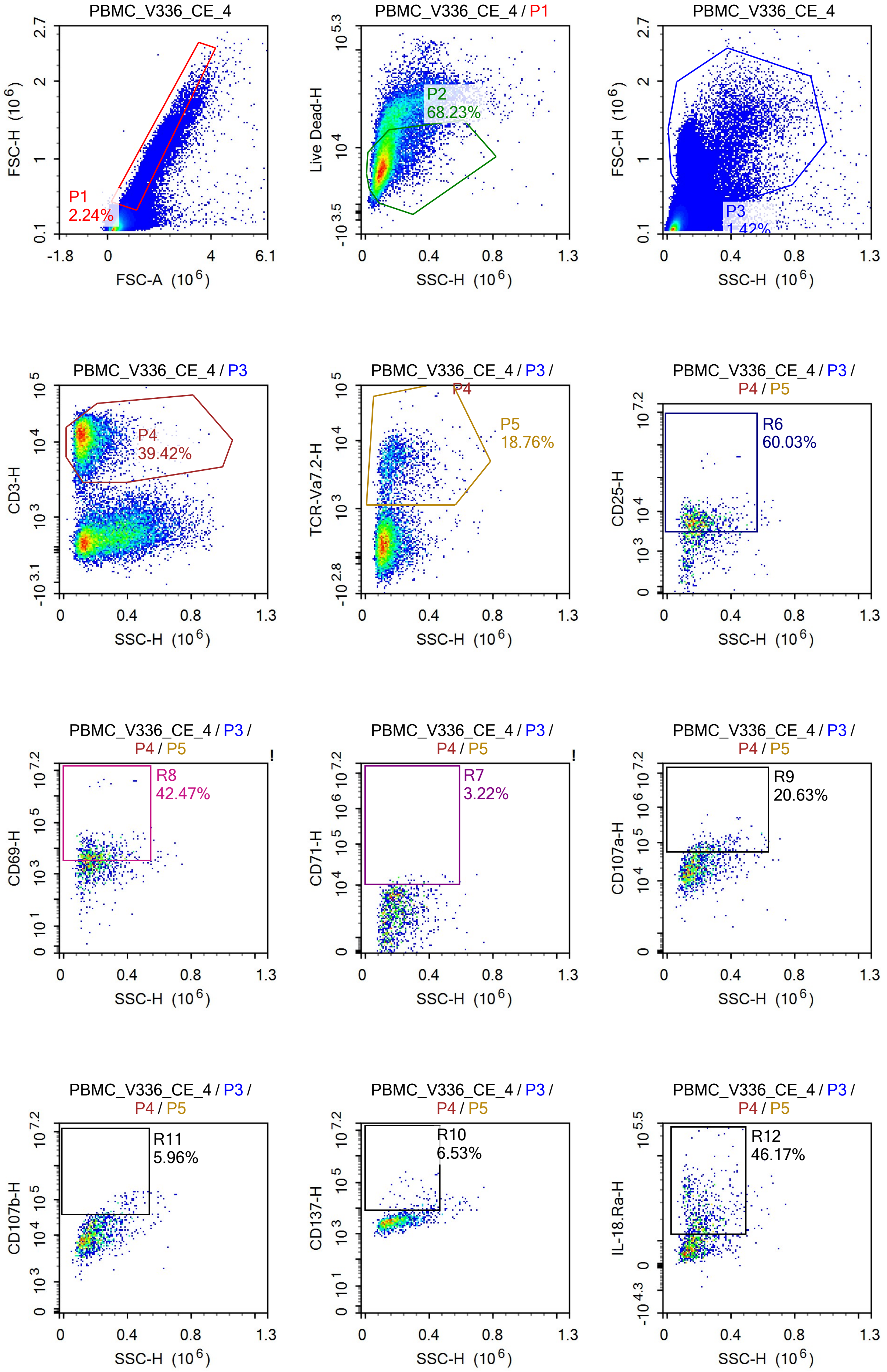
PBMC_V336_CE_4
FSC-H (10^6)
FSC-A (10^6)
P1
2.24%
PBMC_V336_CE_4 / P1
Live Dead-H
SSC-H (10^6)
P2
68.23%
P3
1.42%
PBMC_V336_CE_4 / P3
CD3-H
P4
39.42%
PBMC_V336_CE_4 / P3 / P4
TCR-Va7.2-H
P5
18.76%
PBMC_V336_CE_4 / P3 / P4 / P5
CD25-H
R6
60.03%
CD69-H
R8
42.47%
CD71-H
R7
3.22%
CD107a-H
R9
20.63%
CD107b-H
R11
5.96%
CD137-H
R10
6.53%
IL-18.Ra-H
R12
46.17%

Sample Statistics of PBMC_V336_CE_4

| Gate | Count | % Parent | X | Y | Median X | Median Y |
|---|---|---|---|---|---|---|
| All | 1,186,014 | | | | | |
| P1 | 26,509 | 2.24% | FSC-A | FSC-H | 1,120,812 | 623,271 |
| P2 | 18,088 | 68.23% | SSC-H | Live Dead-H | 110,145 | 7,016 |
| P3 | 16,784 | 1.42% | SSC-H | FSC-H | 171,986 | 1,038,096 |
| P4 | 6,616 | 39.42% | SSC-H | CD3-H | 128,488 | 11,986 |
| P5 | 1,241 | 18.76% | SSC-H | TCR-Va7.2-H | 182,442 | 4,146 |
| R6 | 745 | 60.03% | SSC-H | CD25-H | 193,905 | 5,671 |
| R7 | 40 | 3.22% | SSC-H | CD71-H | 250,486 | 11,667 |
| R8 | 527 | 42.47% | SSC-H | CD69-H | 192,873 | 5,700 |
| R9 | 256 | 20.63% | SSC-H | CD107a-H | 257,562 | 72,535 |
| R10 | 81 | 6.53% | SSC-H | CD137-H | 359,320 | 13,203 |
| R11 | 74 | 5.96% | SSC-H | CD107b-H | 374,354 | 67,947 |
| R12 | 573 | 46.17% | SSC-H | IL-18.Ra-H | 208,641 | 21,488 |

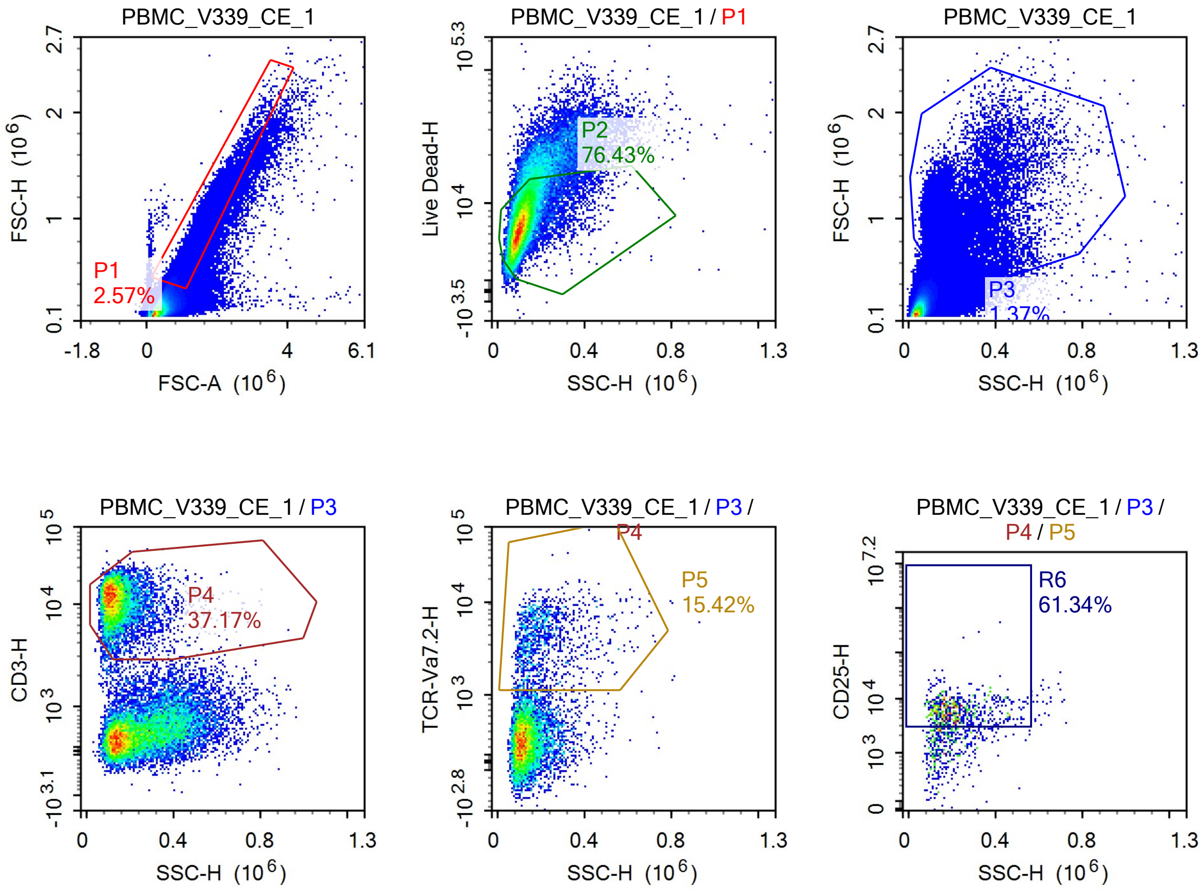

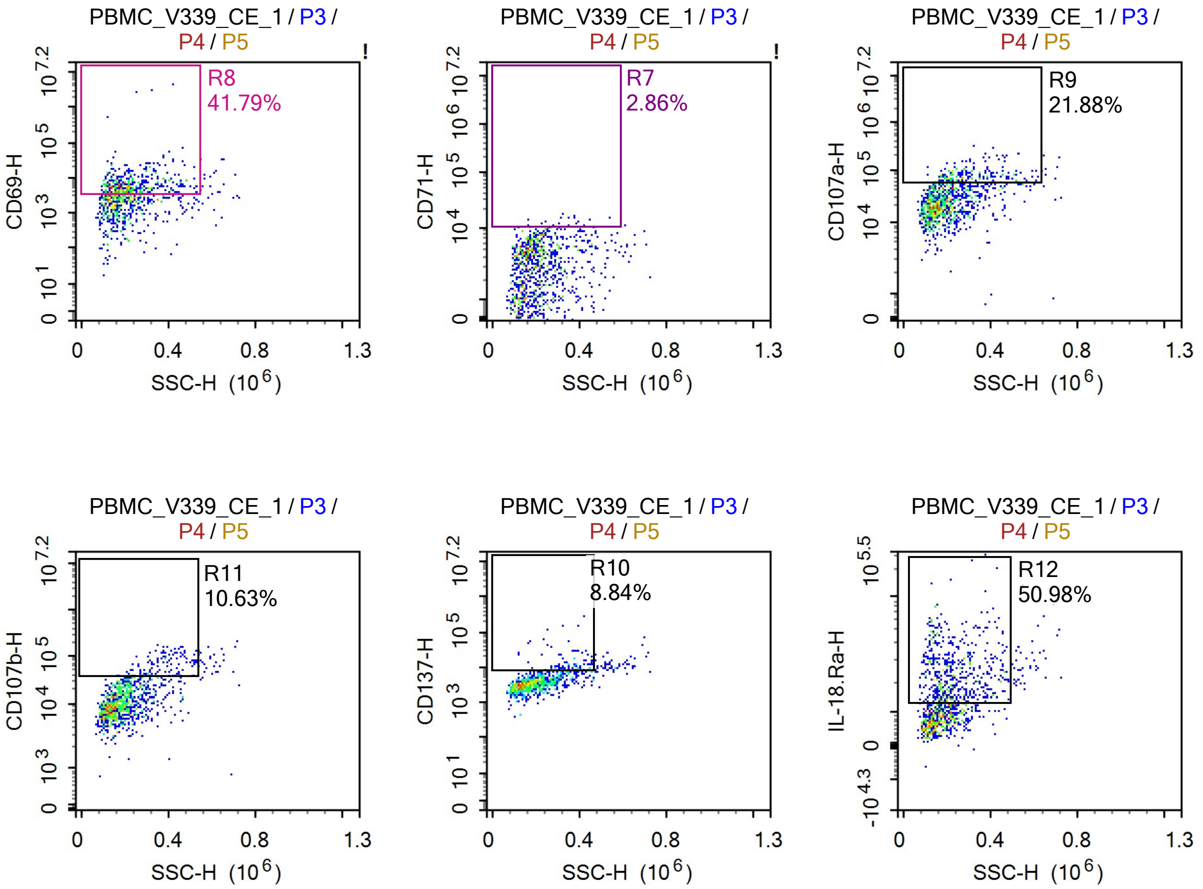


Sample Statistics of PBMC_V339_CE_1

| Gate | Count | % Parent | X | Y | Median X | Median Y |
|---|---|---|---|---|---|---|
| All | 1,425,630 | | | | | |
| P1 | 36,592 | 2.57% | FSC-A | FSC-H | 1,034,664 | 522,742 |
| P2 | 27,966 | 76.43% | SSC-H | Live Dead-H | 106,342 | 7,170 |
| P3 | 19,545 | 1.37% | SSC-H | FSC-H | 172,957 | 1,020,236 |
| P4 | 7,265 | 37.17% | SSC-H | CD3-H | 127,719 | 11,323 |
| P5 | 1,120 | 15.42% | SSC-H | TCR-Va7.2-H | 187,262 | 3,839 |
| R6 | 687 | 61.34% | SSC-H | CD25-H | 201,120 | 5,775 |
| R7 | 32 | 2.86% | SSC-H | CD71-H | 302,157 | 11,166 |
| R8 | 468 | 41.79% | SSC-H | CD69-H | 208,784 | 6,005 |
| R9 | 245 | 21.88% | SSC-H | CD107a-H | 269,150 | 73,562 |
| R10 | 99 | 8.84% | SSC-H | CD137-H | 370,361 | 11,340 |
| R11 | 119 | 10.63% | SSC-H | CD107b-H | 376,734 | 64,374 |
| R12 | 571 | 50.98% | SSC-H | IL-18.Ra-H | 210,324 | 22,598 |

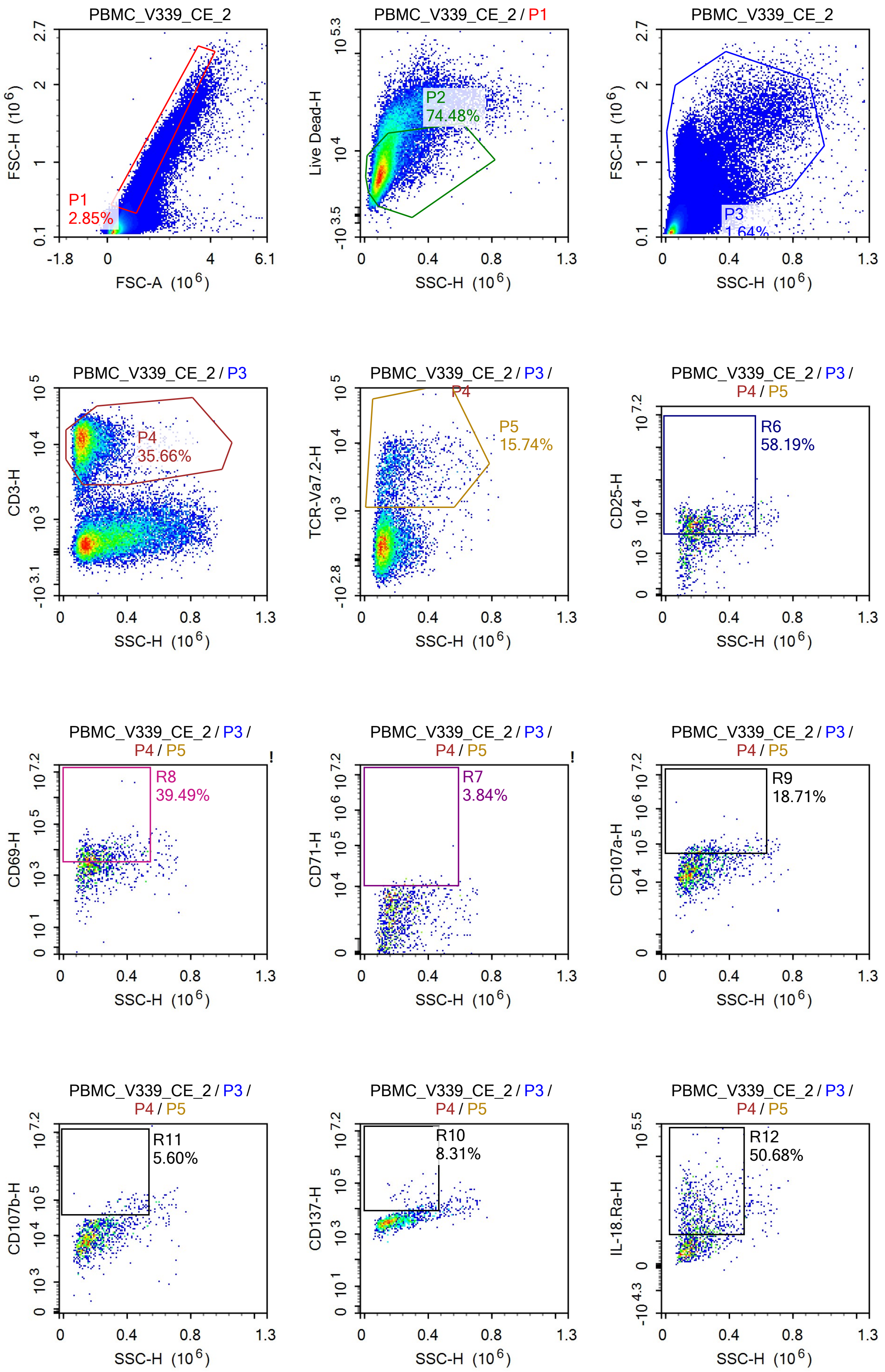
PBMC_V339_CE_2
P1
2.85%
FSC-H (10^6)
FSC-A (10^6)
PBMC_V339_CE_2 / P1
P2
74.48%
Live Dead-H
SSC-H (10^6)
PBMC_V339_CE_2
P3
1.64%
PBMC_V339_CE_2 / P3
P4
35.66%
CD3-H
PBMC_V339_CE_2 / P3 / P4
P5
15.74%
TCR-Va7.2-H
PBMC_V339_CE_2 / P3 / P4 / P5
R6
58.19%
CD25-H
R8
39.49%
CD69-H
R7
3.84%
CD71-H
R9
18.71%
CD107a-H
R11
5.60%
CD107b-H
R10
8.31%
CD137-H
R12
50.68%
IL-18.Ra-H

Sample Statistics of PBMC_V339_CE_2

| Gate | Count | % Parent | X | Y | Median X | Median Y |
|---|---|---|---|---|---|---|
| All | 1,361,102 | | | | | |
| P1 | 38,821 | 2.85% | FSC-A | FSC-H | 1,061,517 | 548,158 |
| P2 | 28,913 | 74.48% | SSC-H | Live Dead-H | 110,068 | 6,677 |
| P3 | 22,291 | 1.64% | SSC-H | FSC-H | 181,315 | 1,031,510 |
| P4 | 7,950 | 35.66% | SSC-H | CD3-H | 130,605 | 11,227 |
| P5 | 1,251 | 15.74% | SSC-H | TCR-Va7.2-H | 187,455 | 3,649 |
| R6 | 728 | 58.19% | SSC-H | CD25-H | 202,311 | 5,697 |
| R7 | 48 | 3.84% | SSC-H | CD71-H | 301,102 | 11,314 |
| R8 | 494 | 39.49% | SSC-H | CD69-H | 209,662 | 6,711 |
| R9 | 234 | 18.71% | SSC-H | CD107a-H | 271,035 | 66,714 |
| R10 | 104 | 8.31% | SSC-H | CD137-H | 356,813 | 12,413 |
| R11 | 70 | 5.60% | SSC-H | CD107b-H | 404,531 | 65,519 |
| R12 | 634 | 50.68% | SSC-H | IL-18.Ra-H | 207,801 | 22,356 |

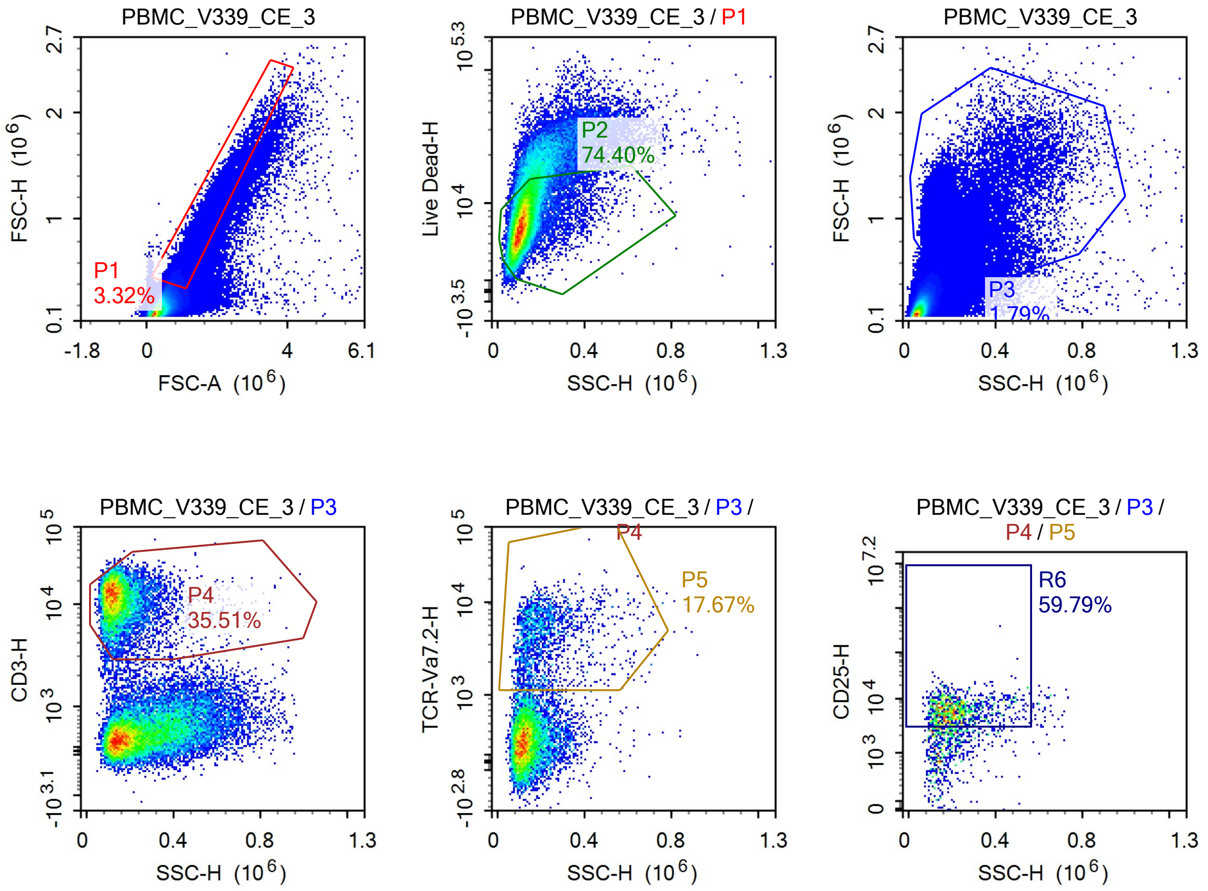

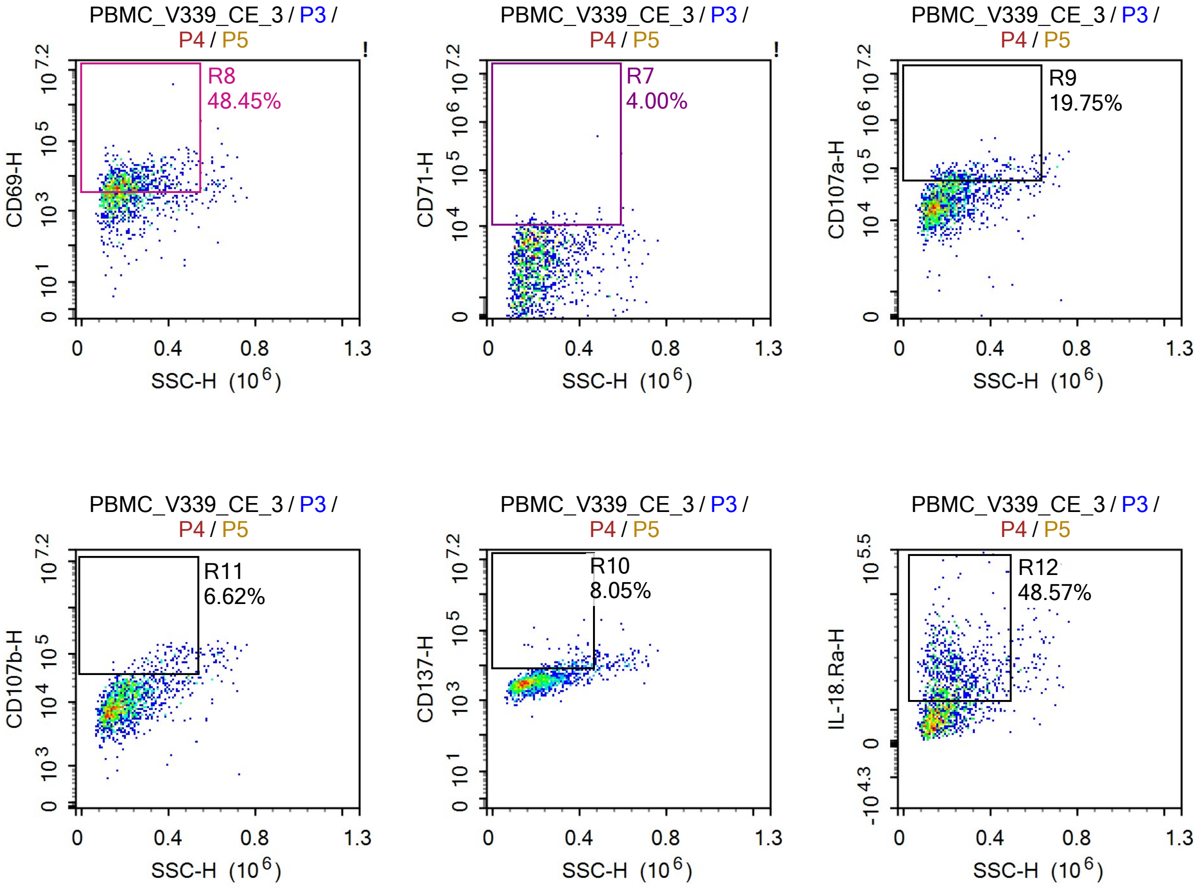


Sample Statistics of PBMC_V339_CE_3

| Gate | Count | % Parent | X | Y | Median X | Median Y |
|---|---|---|---|---|---|---|
| All | 1,491,717 | | | | | |
| P1 | 49,474 | 3.32% | FSC-A | FSC-H | 1,035,408 | 521,816 |
| P2 | 36,810 | 74.40% | SSC-H | Live Dead-H | 112,026 | 7,273 |
| P3 | 26,707 | 1.79% | SSC-H | FSC-H | 181,560 | 1,002,977 |
| P4 | 9,484 | 35.51% | SSC-H | CD3-H | 132,681 | 11,864 |
| P5 | 1,676 | 17.67% | SSC-H | TCR-Va7.2-H | 189,727 | 4,186 |
| R6 | 1,002 | 59.79% | SSC-H | CD25-H | 202,074 | 5,978 |
| R7 | 67 | 4.00% | SSC-H | CD71-H | 245,056 | 11,029 |
| R8 | 812 | 48.45% | SSC-H | CD69-H | 201,047 | 6,311 |
| R9 | 331 | 19.75% | SSC-H | CD107a-H | 281,311 | 67,815 |
| R10 | 135 | 8.05% | SSC-H | CD137-H | 363,221 | 10,408 |
| R11 | 111 | 6.62% | SSC-H | CD107b-H | 387,247 | 68,304 |
| R12 | 814 | 48.57% | SSC-H | IL-18.Ra-H | 216,513 | 21,677 |

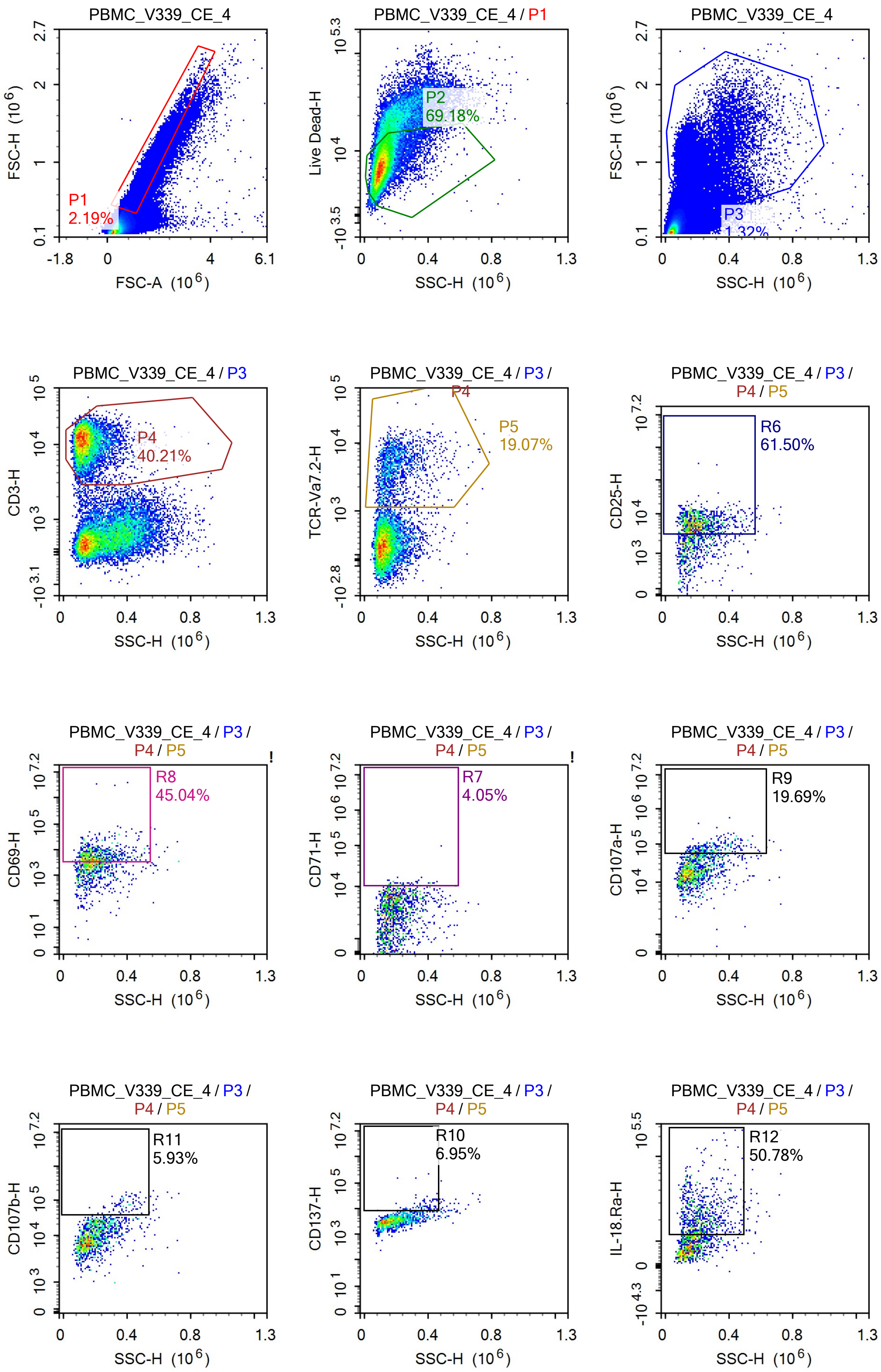
PBMC_V339_CE_4
P1
2.19%
FSC-H (10^6)
FSC-A (10^6)
PBMC_V339_CE_4 / P1
P2
69.18%
Live Dead-H
SSC-H (10^6)
PBMC_V339_CE_4
P3
1.32%
FSC-H (10^6)
SSC-H (10^6)
PBMC_V339_CE_4 / P3
P4
40.21%
CD3-H
PBMC_V339_CE_4 / P3 / P4
P5
19.07%
TCR-Va7.2-H
PBMC_V339_CE_4 / P3 / P4 / P5
R6
61.50%
CD25-H
R8
45.04%
CD69-H
R7
4.05%
CD71-H
R9
19.69%
CD107a-H
R11
5.93%
CD107b-H
R10
6.95%
CD137-H
R12
50.78%
IL-18.Ra-H

Sample Statistics of PBMC_V339_CE_4

| Gate | Count | % Parent | X | Y | Median X | Median Y |
|---|---|---|---|---|---|---|
| All | 1,462,398 | | | | | |
| P1 | 31,982 | 2.19% | FSC-A | FSC-H | 1,095,453 | 582,087 |
| P2 | 22,124 | 69.18% | SSC-H | Live Dead-H | 107,214 | 7,384 |
| P3 | 19,340 | 1.32% | SSC-H | FSC-H | 174,478 | 1,025,526 |
| P4 | 7,776 | 40.21% | SSC-H | CD3-H | 131,179 | 11,414 |
| P5 | 1,483 | 19.07% | SSC-H | TCR-Va7.2-H | 179,350 | 4,133 |
| R6 | 912 | 61.50% | SSC-H | CD25-H | 196,878 | 5,723 |
| R7 | 60 | 4.05% | SSC-H | CD71-H | 226,440 | 11,378 |
| R8 | 668 | 45.04% | SSC-H | CD69-H | 192,724 | 5,868 |
| R9 | 292 | 19.69% | SSC-H | CD107a-H | 267,325 | 70,851 |
| R10 | 103 | 6.95% | SSC-H | CD137-H | 354,039 | 10,530 |
| R11 | 88 | 5.93% | SSC-H | CD107b-H | 377,905 | 69,561 |
| R12 | 753 | 50.78% | SSC-H | IL-18.Ra-H | 213,981 | 20,805 |

### Report of PBMC 12/09/25-PBMC_Ctrl_RPMI_

Sample Name: PBMC 12/09/25-PBMC_Ctrl_RPMI_1 Run Time: 12-Sep-25 1:26 PM

Cytometer: NovoCyte Quanteon 621181110427 Software: NovoExpress 1.6.0

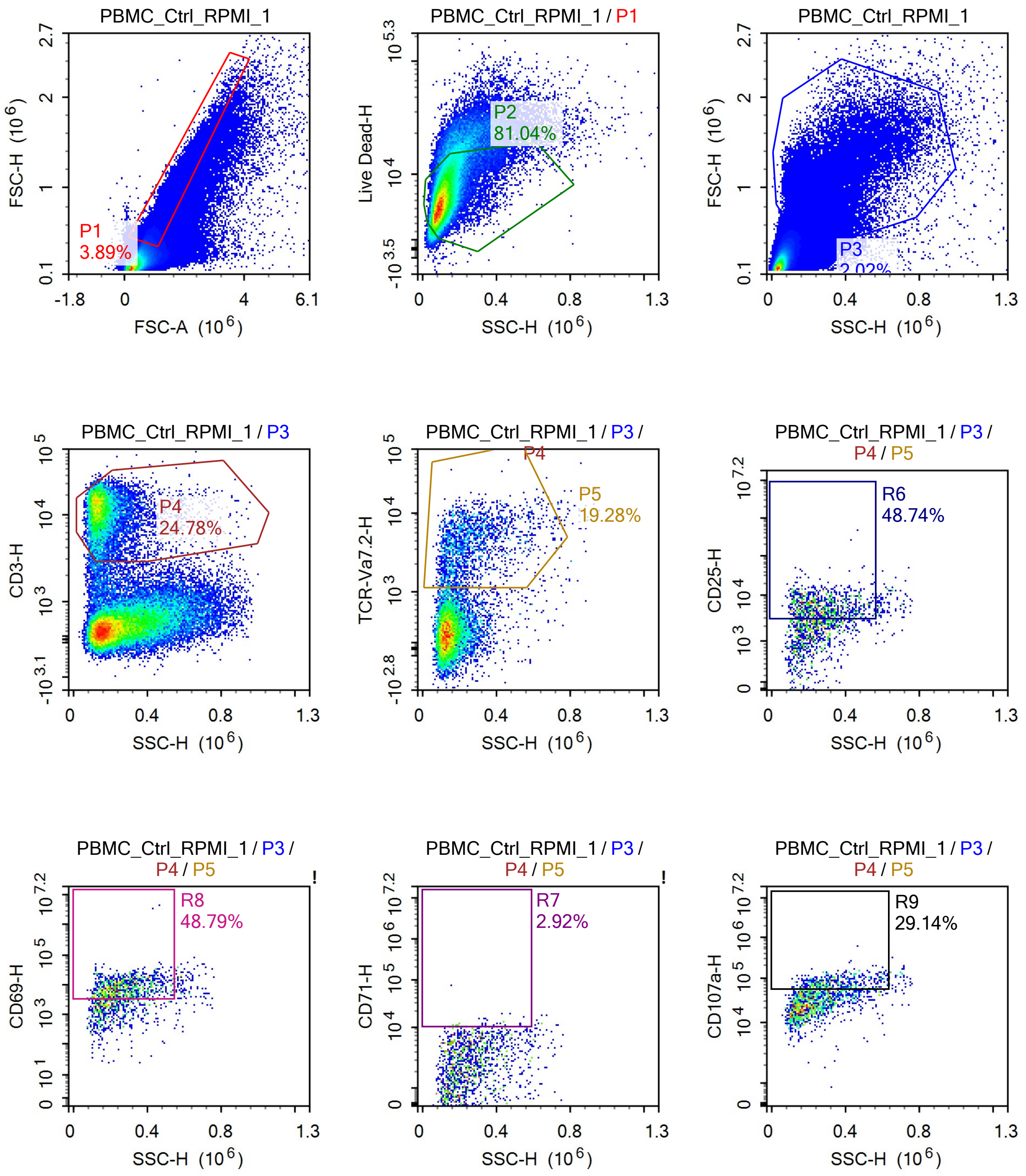

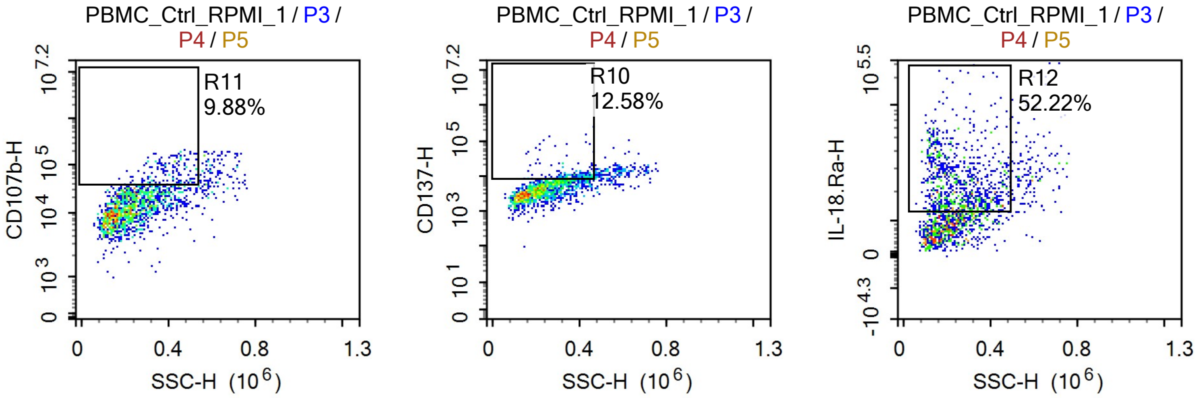


Sample Statistics

| Gate | Count | % Parent | X | Y | Median X | Median Y |
|---|---|---|---|---|---|---|
| All | 1,842,411 | | | | | |
| P1 | 71,739 | 3.89% | FSC-A | FSC-H | 1,014,041 | 489,640 |
| P2 | 58,137 | 81.04% | SSC-H | Live Dead-H | 107,806 | 6,169 |
| P3 | 37,278 | 2.02% | SSC-H | FSC-H | 204,862 | 954,967 |
| P4 | 9,237 | 24.78% | SSC-H | CD3-H | 146,861 | 11,368 |
| P5 | 1,781 | 19.28% | SSC-H | TCR-Va7.2-H | 231,900 | 3,823 |
| R6 | 868 | 48.74% | SSC-H | CD25-H | 258,153 | 5,571 |
| R7 | 52 | 2.92% | SSC-H | CD71-H | 327,375 | 11,127 |
| R8 | 869 | 48.79% | SSC-H | CD69-H | 257,849 | 8,180 |
| R9 | 519 | 29.14% | SSC-H | CD107a-H | 320,524 | 77,327 |
| R10 | 224 | 12.58% | SSC-H | CD137-H | 366,419 | 9,693 |
| R11 | 176 | 9.88% | SSC-H | CD107b-H | 400,009 | 66,444 |
| R12 | 930 | 52.22% | SSC-H | IL-18.Ra-H | 241,819 | 21,868 |

### Report of PBMC 12/09/25-PBMC_Ctrl_RPMI_

Sample Name: PBMC 12/09/25-PBMC_Ctrl_RPMI_2 Run Time: 12-Sep-25 1:28 PM

Cytometer: NovoCyte Quanteon 621181110427 Software: NovoExpress 1.6.0

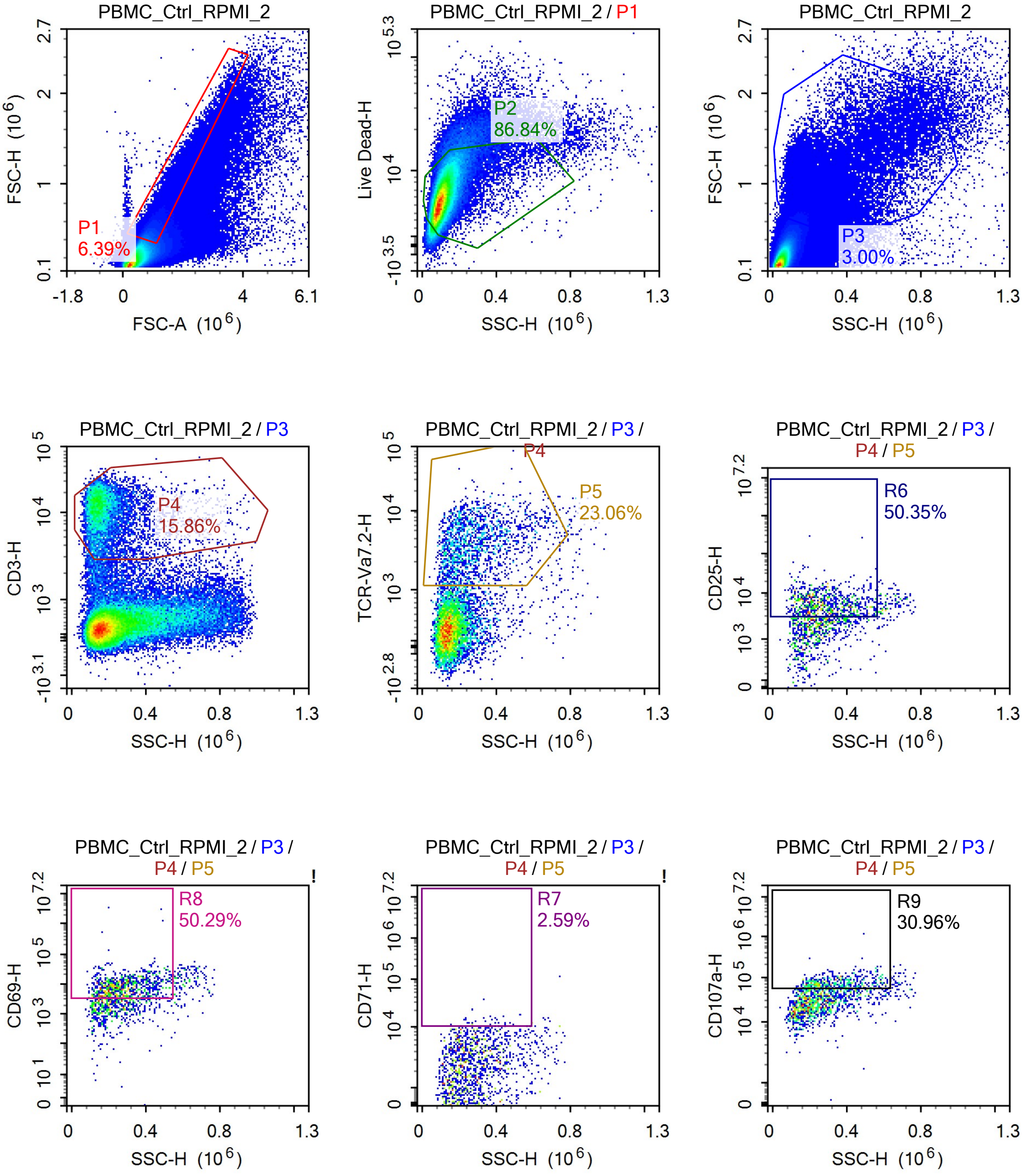

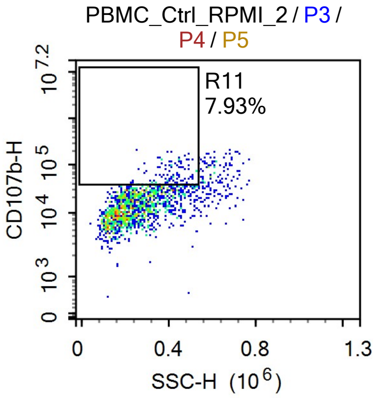


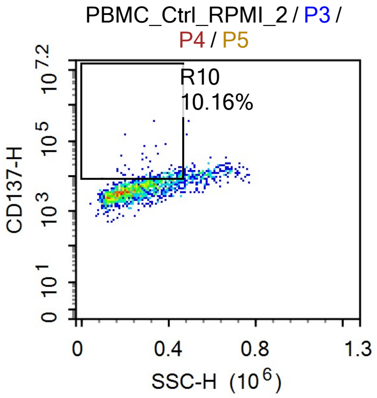


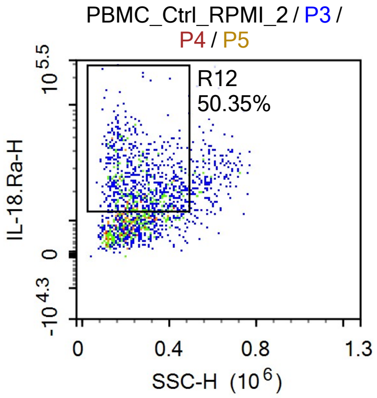


Sample Statistics

| Gate | Count | % Parent | X | Y | Median X | Median Y |
|---|---|---|---|---|---|---|
| All | 1,549,068 | | | | | |
| P1 | 99,042 | 6.39% | FSC-A | FSC-H | 990,278 | 472,055 |
| P2 | 86,010 | 86.84% | SSC-H | Live Dead-H | 105,827 | 5,831 |
| P3 | 46,549 | 3.00% | SSC-H | FSC-H | 196,318 | 825,779 |
| P4 | 7,382 | 15.86% | SSC-H | CD3-H | 161,027 | 10,793 |
| P5 | 1,702 | 23.06% | SSC-H | TCR-Va7.2-H | 252,958 | 3,481 |
| R6 | 857 | 50.35% | SSC-H | CD25-H | 275,174 | 5,458 |
| R7 | 44 | 2.59% | SSC-H | CD71-H | 347,088 | 11,927 |
| R8 | 856 | 50.29% | SSC-H | CD69-H | 274,817 | 8,325 |
| R9 | 527 | 30.96% | SSC-H | CD107a-H | 334,835 | 73,179 |
| R10 | 173 | 10.16% | SSC-H | CD137-H | 387,671 | 9,663 |
| R11 | 135 | 7.93% | SSC-H | CD107b-H | 406,445 | 53,203 |
| R12 | 857 | 50.35% | SSC-H | IL-18.Ra-H | 256,673 | 21,928 |

### Report of PBMC 12/09/25-PBMC_Ctrl_RPMI_

Sample Name: PBMC 12/09/25-PBMC_Ctrl_RPMI_3 | Run Time: 12-Sep-25 1:30 PM

Cytometer: NovoCyte Quanteon 621181110427 | Software: NovoExpress 1.6.0

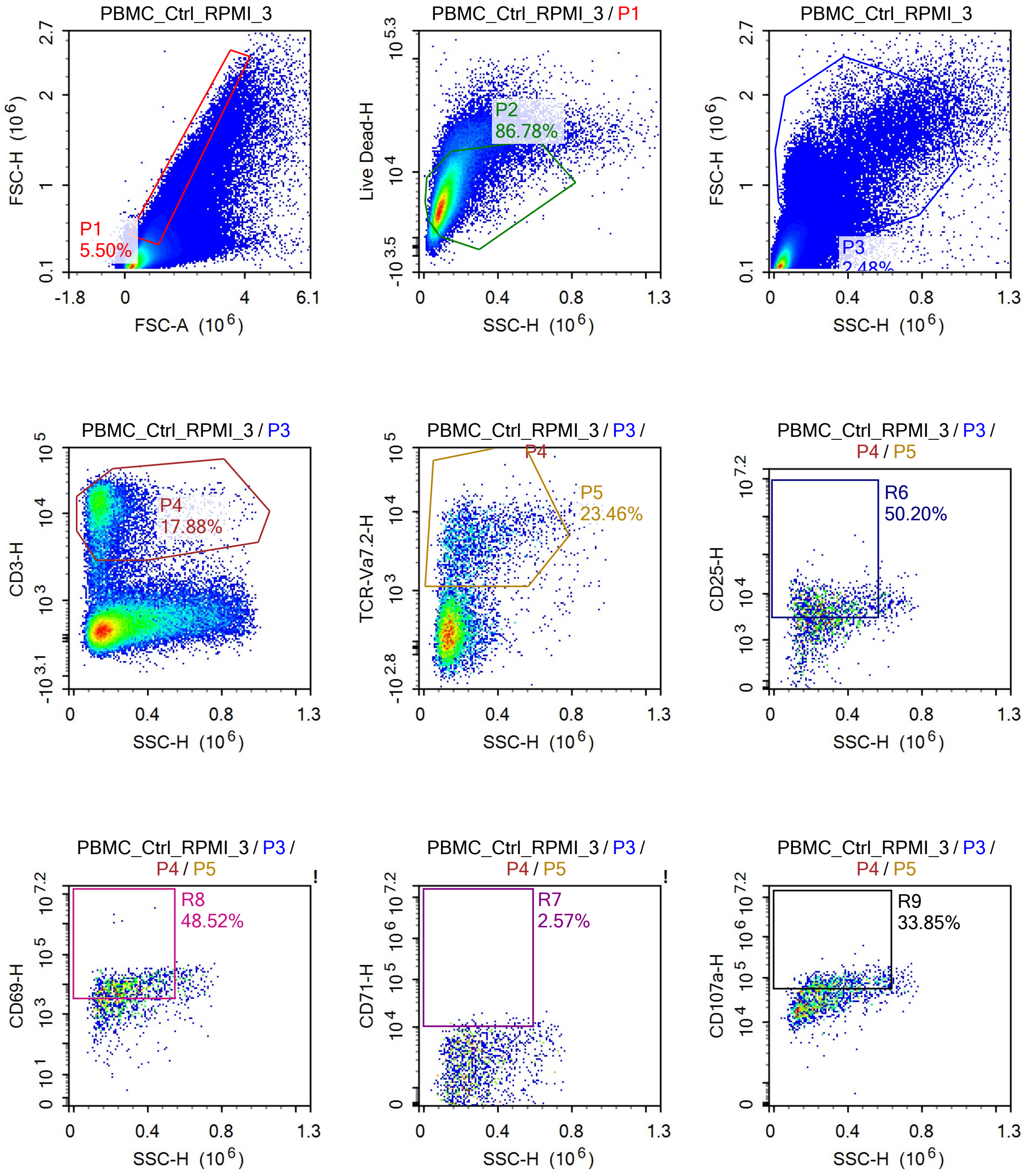

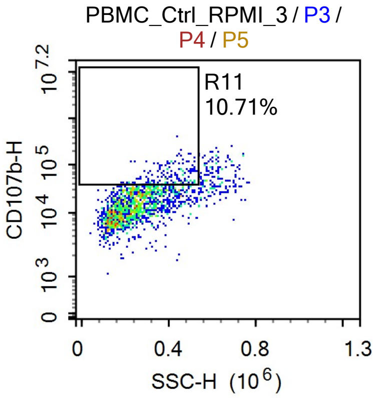

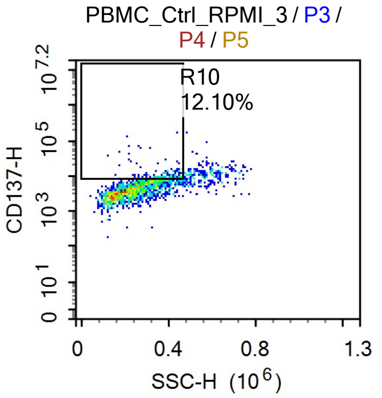

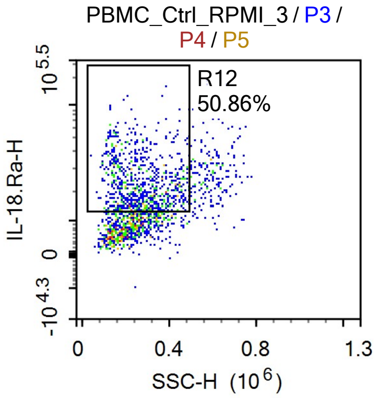


Sample Statistics

| Gate | Count | % Parent | X | Y | Median X | Median Y |
|---|---|---|---|---|---|---|
| All | 1,725,636 | | | | | |
| P1 | 94,859 | 5.50% | FSC-A | FSC-H | 986,720 | 469,540 |
| P2 | 82,321 | 86.78% | SSC-H | Live Dead-H | 104,129 | 5,650 |
| P3 | 42,742 | 2.48% | SSC-H | FSC-H | 195,647 | 840,369 |
| P4 | 7,643 | 17.88% | SSC-H | CD3-H | 157,558 | 11,567 |
| P5 | 1,793 | 23.46% | SSC-H | TCR-Va7.2-H | 253,278 | 3,799 |
| R6 | 900 | 50.20% | SSC-H | CD25-H | 261,064 | 5,612 |
| R7 | 46 | 2.57% | SSC-H | CD71-H | 363,562 | 11,789 |
| R8 | 870 | 48.52% | SSC-H | CD69-H | 274,720 | 8,280 |
| R9 | 607 | 33.85% | SSC-H | CD107a-H | 338,306 | 75,685 |
| R10 | 217 | 12.10% | SSC-H | CD137-H | 373,034 | 9,498 |
| R11 | 192 | 10.71% | SSC-H | CD107b-H | 402,920 | 48,095 |
| R12 | 912 | 50.86% | SSC-H | IL-18.Ra-H | 261,667 | 21,548 |

### Report of PBMC 12/09/25-PBMC_Ctrl_RPMI_

Sample Name: PBMC 12/09/25-PBMC_Ctrl_RPMI_4 | Run Time: 12-Sep-25 1:32 PM

Cytometer: NovoCyte Quanteon 621181110427 | Software: NovoExpress 1.6.0

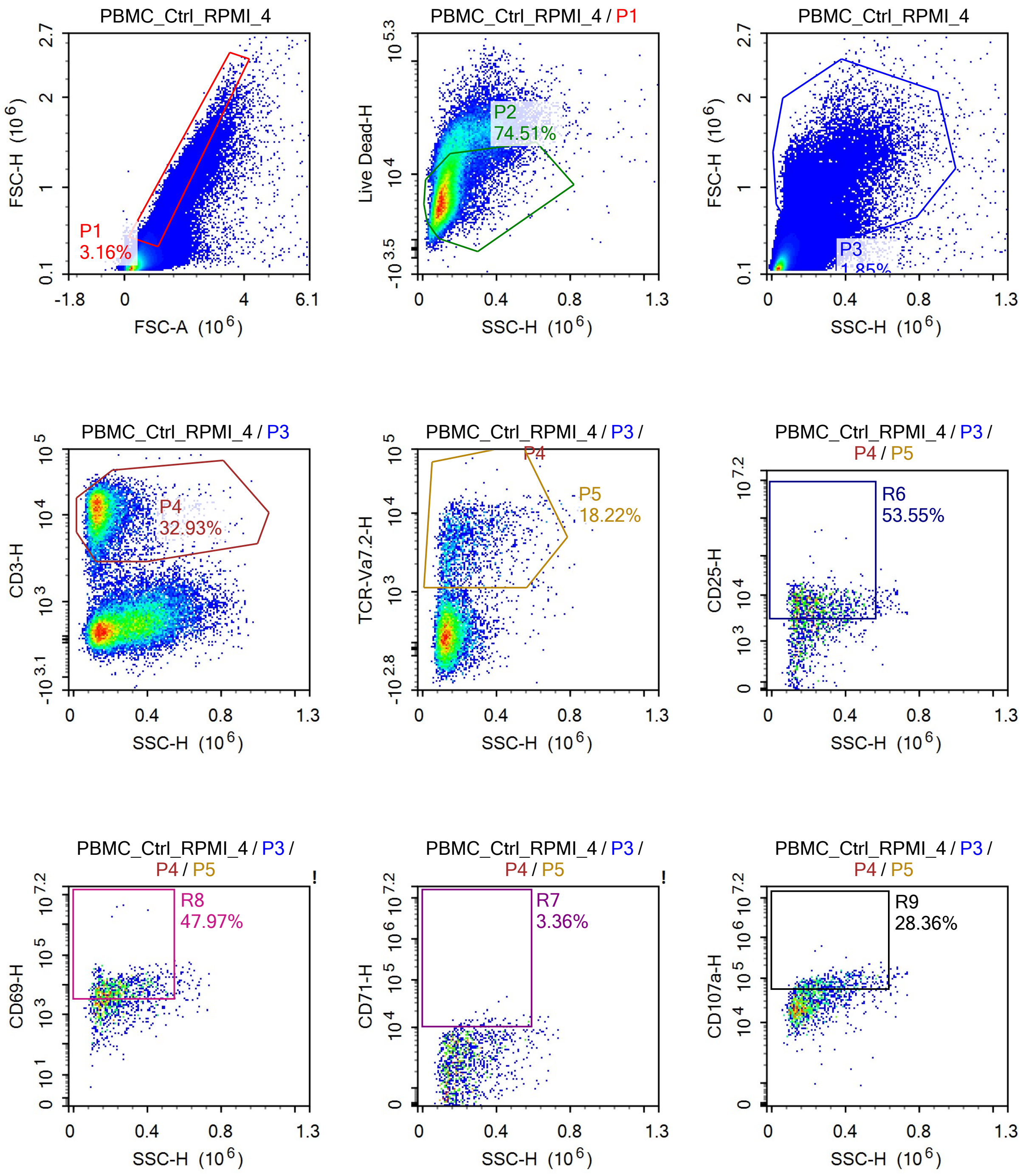

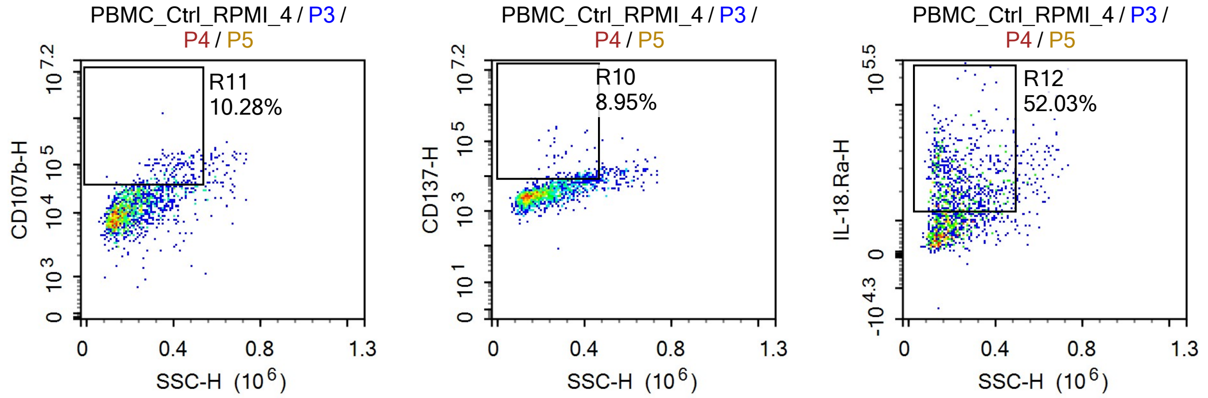


Sample Statistics

| Gate | Count | % Parent | X | Y | Median X | Median Y |
|---|---|---|---|---|---|---|
| All | 1,423,509 | | | | | |
| P1 | 45,021 | 3.16% | FSC-A | FSC-H | 1,060,403 | 531,699 |
| P2 | 33,543 | 74.51% | SSC-H | Live Dead-H | 114,934 | 6,584 |
| P3 | 26,265 | 1.85% | SSC-H | FSC-H | 195,199 | 995,281 |
| P4 | 8,648 | 32.93% | SSC-H | CD3-H | 140,461 | 11,458 |
| P5 | 1,576 | 18.22% | SSC-H | TCR-Va7.2-H | 202,106 | 3,883 |
| R6 | 844 | 53.55% | SSC-H | CD25-H | 223,597 | 6,304 |
| R7 | 53 | 3.36% | SSC-H | CD71-H | 350,403 | 12,388 |
| R8 | 756 | 47.97% | SSC-H | CD69-H | 226,637 | 7,345 |
| R9 | 447 | 28.36% | SSC-H | CD107a-H | 303,845 | 75,946 |
| R10 | 141 | 8.95% | SSC-H | CD137-H | 366,146 | 9,936 |
| R11 | 162 | 10.28% | SSC-H | CD107b-H | 393,861 | 61,424 |
| R12 | 820 | 52.03% | SSC-H | IL-18.Ra-H | 223,340 | 22,299 |

### Report of PBMC 12/09/25-PBMC_5-OP-RU-

Sample Name: PBMC 12/09/25-PBMC_5-OP-RU-8uM_1 Run Time: 12-Sep-25 1:34 PM

Cytometer: NovoCyte Quanteon 621181110427 Software: NovoExpress 1.6.0

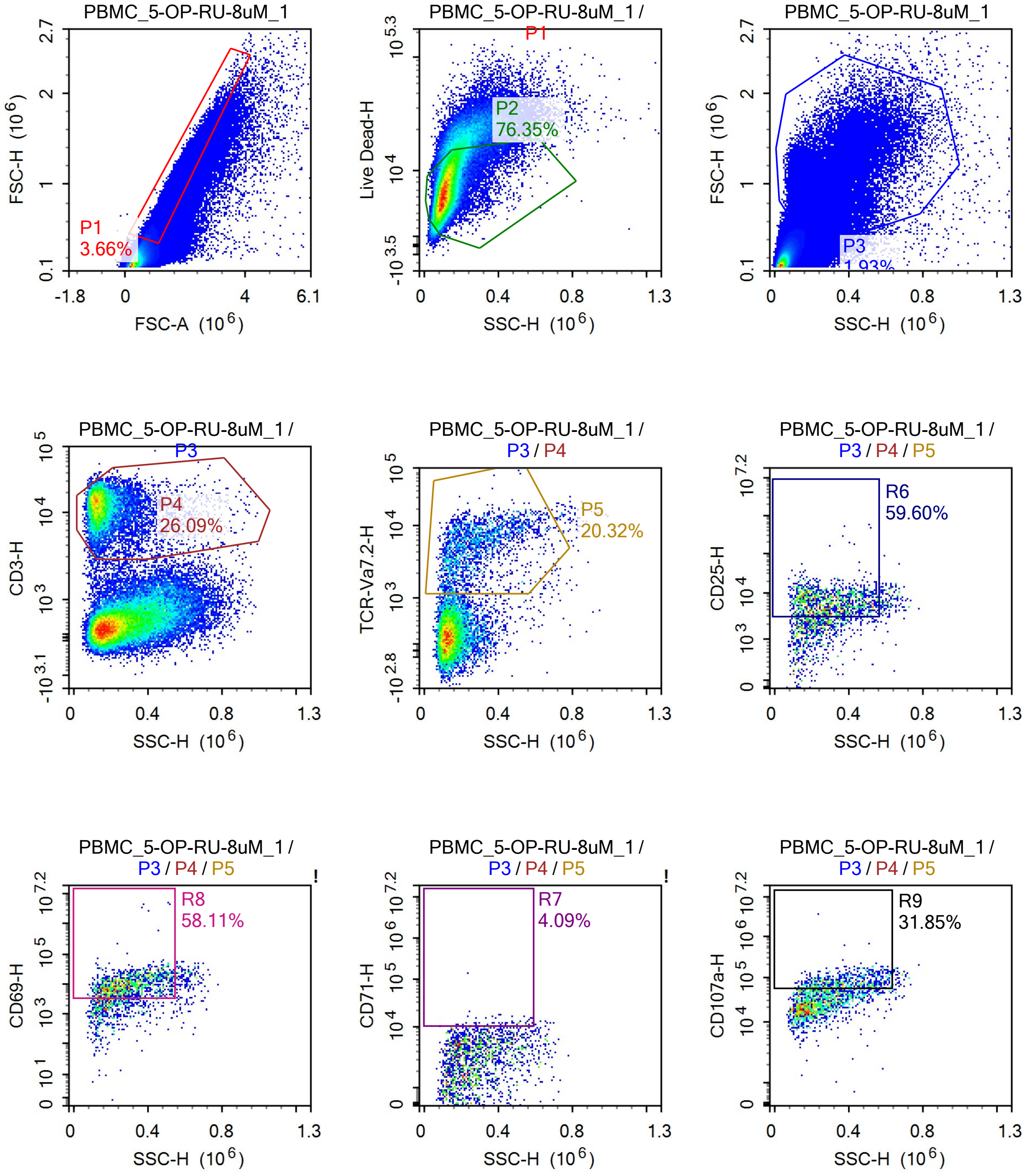

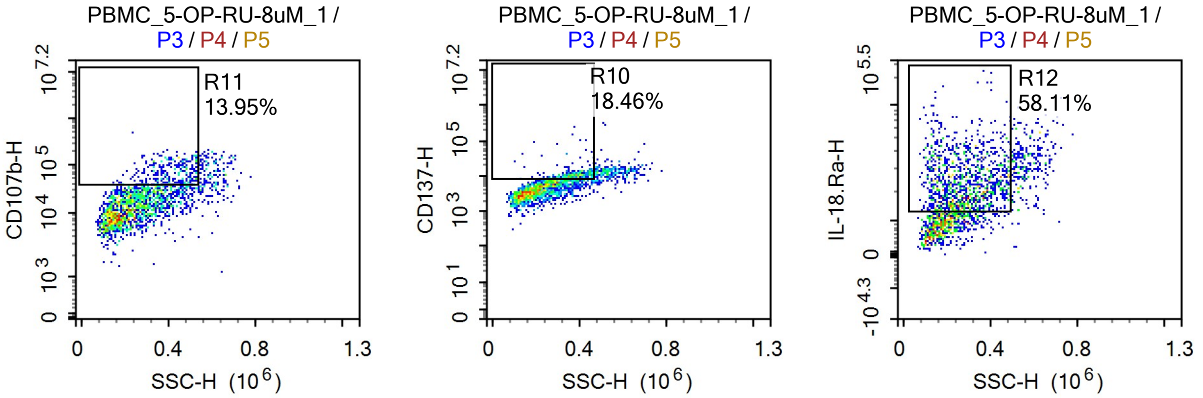


Sample Statistics

| Gate | Count | % Parent | X | Y | Median X | Median Y |
|---|---|---|---|---|---|---|
| All | 2,101,112 | | | | | |
| P1 | 76,918 | 3.66% | FSC-A | FSC-H | 1,005,279 | 501,881 |
| P2 | 58,728 | 76.35% | SSC-H | Live Dead-H | 117,049 | 7,135 |
| P3 | 40,574 | 1.93% | SSC-H | FSC-H | 211,422 | 936,969 |
| P4 | 10,587 | 26.09% | SSC-H | CD3-H | 144,061 | 12,225 |
| P5 | 2,151 | 20.32% | SSC-H | TCR-Va7.2-H | 254,947 | 5,249 |
| R6 | 1,282 | 59.60% | SSC-H | CD25-H | 287,998 | 6,204 |
| R7 | 88 | 4.09% | SSC-H | CD71-H | 381,966 | 11,129 |
| R8 | 1,250 | 58.11% | SSC-H | CD69-H | 284,115 | 9,526 |
| R9 | 685 | 31.85% | SSC-H | CD107a-H | 367,158 | 75,784 |
| R10 | 397 | 18.46% | SSC-H | CD137-H | 366,421 | 10,112 |
| R11 | 300 | 13.95% | SSC-H | CD107b-H | 408,812 | 66,626 |
| R12 | 1,250 | 58.11% | SSC-H | IL-18.Ra-H | 277,017 | 22,659 |

### Report of PBMC 12/09/25-PBMC_5-OP-RU-

Sample Name: PBMC 12/09/25-PBMC_5-OP-RU-8uM_2 | Run Time: 12-Sep-25 1:36 PM

Cytometer: NovoCyte Quanteon 621181110427 | Software: NovoExpress 1.6.0

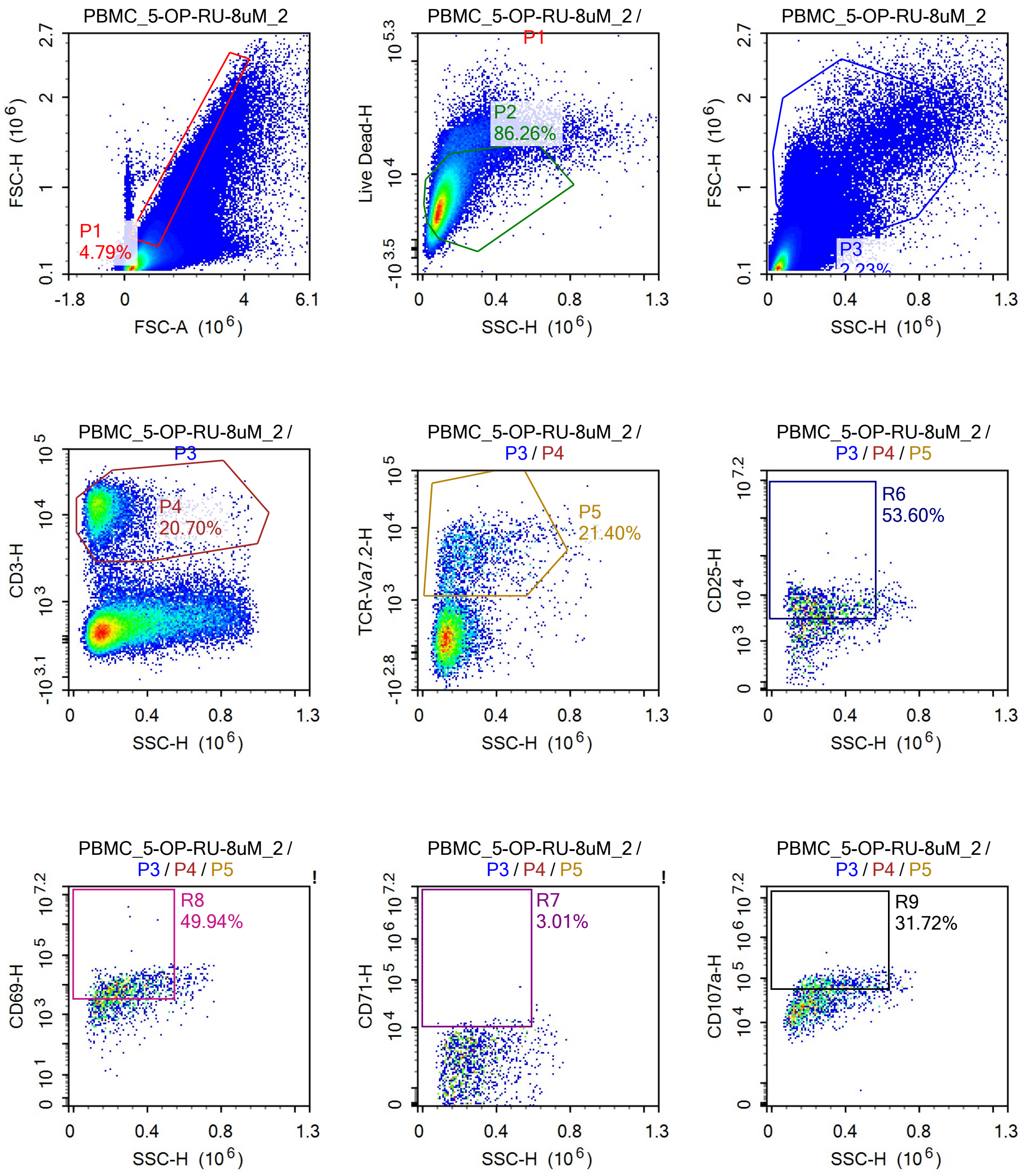

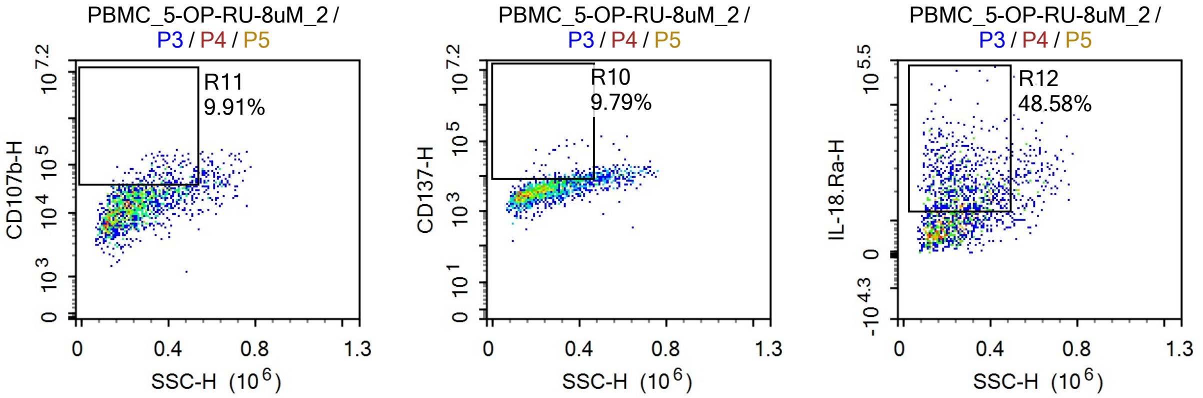


Sample Statistics

| Gate | Count | % Parent | X | Y | Median X | Median Y |
|---|---|---|---|---|---|---|
| All | 1,718,594 | | | | | |
| P1 | 82,332 | 4.79% | FSC-A | FSC-H | 992,647 | 463,118 |
| P2 | 71,021 | 86.26% | SSC-H | Live Dead-H | 100,626 | 5,494 |
| P3 | 38,291 | 2.23% | SSC-H | FSC-H | 194,743 | 862,481 |
| P4 | 7,926 | 20.70% | SSC-H | CD3-H | 150,916 | 12,251 |
| P5 | 1,696 | 21.40% | SSC-H | TCR-Va7.2-H | 239,696 | 3,798 |
| R6 | 909 | 53.60% | SSC-H | CD25-H | 249,408 | 5,735 |
| R7 | 51 | 3.01% | SSC-H | CD71-H | 383,468 | 11,153 |
| R8 | 847 | 49.94% | SSC-H | CD69-H | 269,442 | 7,874 |
| R9 | 538 | 31.72% | SSC-H | CD107a-H | 323,647 | 72,542 |
| R10 | 166 | 9.79% | SSC-H | CD137-H | 376,920 | 9,466 |
| R11 | 168 | 9.91% | SSC-H | CD107b-H | 387,057 | 58,422 |
| R12 | 824 | 48.58% | SSC-H | IL-18.Ra-H | 257,053 | 21,046 |

### Report of PBMC 12/09/25-PBMC_5-OP-RU-

Sample Name: PBMC 12/09/25-PBMC_5-OP-RU-8uM_3 | Run Time: 12-Sep-25 1:38 PM

Cytometer: NovoCyte Quanteon 621181110427 | Software: NovoExpress 1.6.0

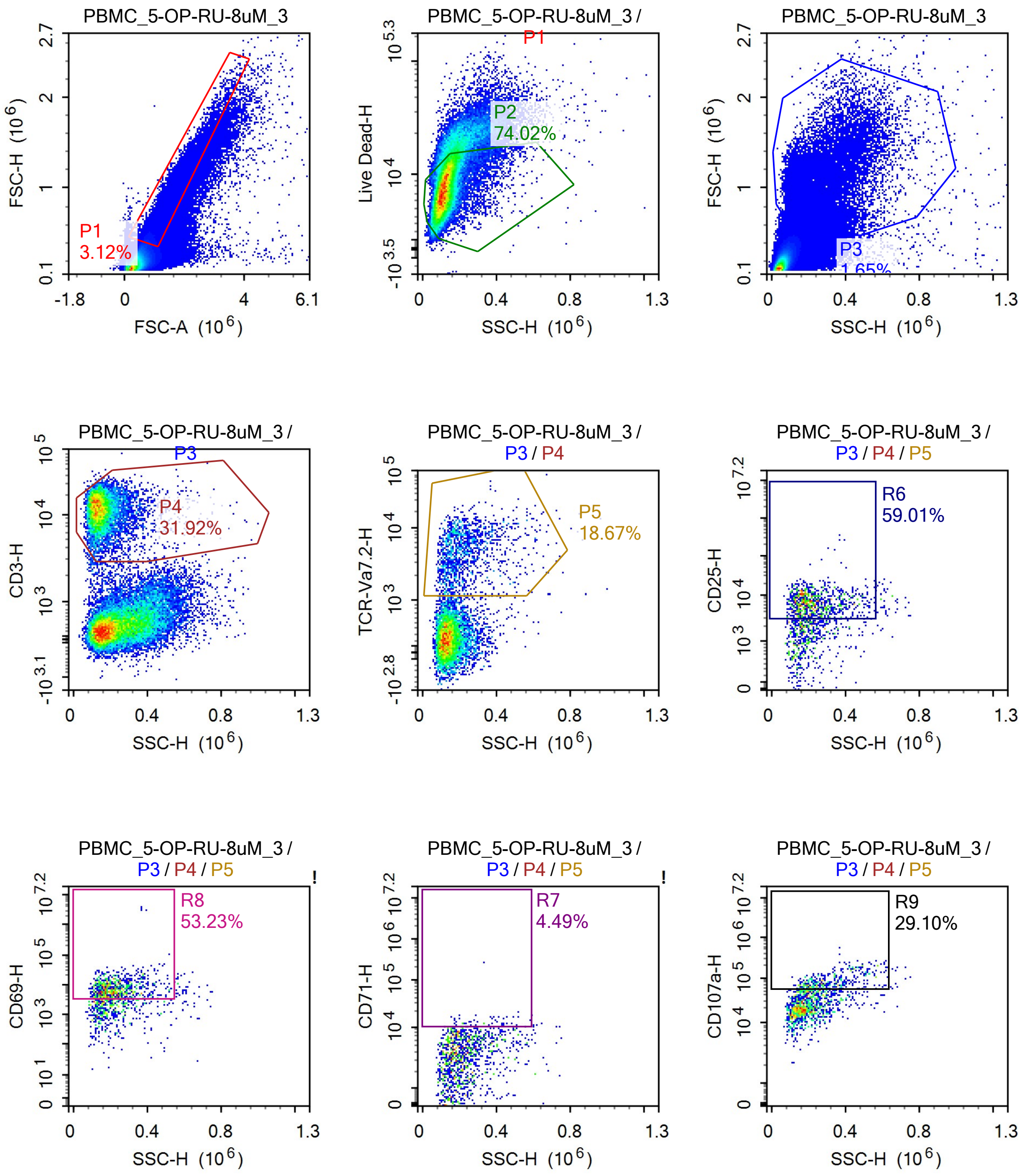

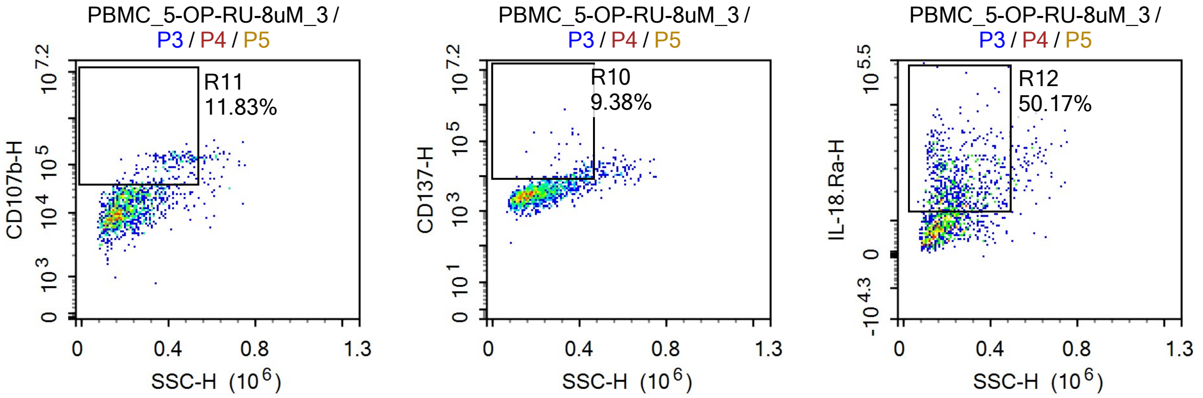


Sample Statistics

| Gate | Count | % Parent | X | Y | Median X | Median Y |
|---|---|---|---|---|---|---|
| All | 1,493,888 | | | | | |
| P1 | 46,535 | 3.12% | FSC-A | FSC-H | 1,014,305 | 508,650 |
| P2 | 34,444 | 74.02% | SSC-H | Live Dead-H | 119,118 | 7,528 |
| P3 | 24,689 | 1.65% | SSC-H | FSC-H | 193,958 | 963,657 |
| P4 | 7,880 | 31.92% | SSC-H | CD3-H | 142,962 | 12,035 |
| P5 | 1,471 | 18.67% | SSC-H | TCR-Va7.2-H | 204,296 | 4,503 |
| R6 | 868 | 59.01% | SSC-H | CD25-H | 216,002 | 6,723 |
| R7 | 66 | 4.49% | SSC-H | CD71-H | 316,834 | 12,409 |
| R8 | 783 | 53.23% | SSC-H | CD69-H | 212,605 | 7,412 |
| R9 | 428 | 29.10% | SSC-H | CD107a-H | 289,994 | 80,220 |
| R10 | 138 | 9.38% | SSC-H | CD137-H | 371,737 | 10,096 |
| R11 | 174 | 11.83% | SSC-H | CD107b-H | 402,415 | 99,604 |
| R12 | 738 | 50.17% | SSC-H | IL-18.Ra-H | 229,624 | 21,863 |

### Report of PBMC 12/09/25-PBMC_5-OP-RU-

Sample Name: PBMC 12/09/25-PBMC_5-OP-RU-8uM_4 | Run Time: 12-Sep-25 1:40 PM

Cytometer: NovoCyte Quanteon 621181110427 | Software: NovoExpress 1.6.0

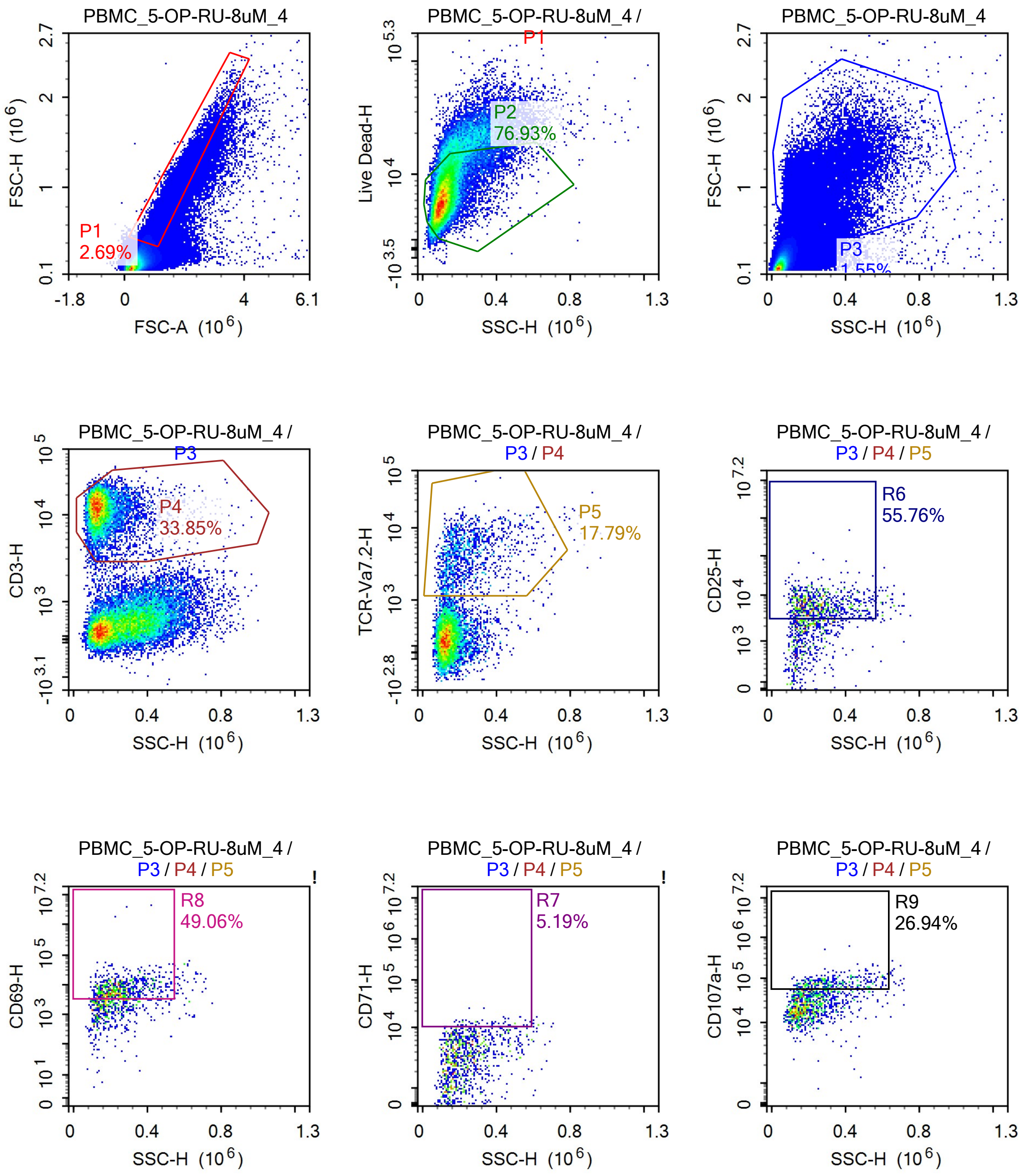

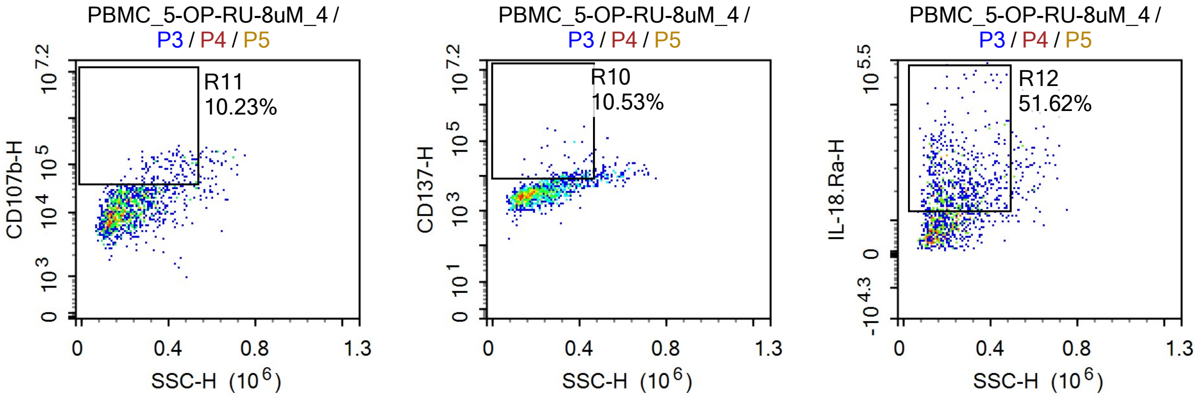


Sample Statistics

| Gate | Count | % Parent | X | Y | Median X | Median Y |
|---|---|---|---|---|---|---|
| All | 1,426,487 | | | | | |
| P1 | 38,330 | 2.69% | FSC-A | FSC-H | 1,050,687 | 534,440 |
| P2 | 29,488 | 76.93% | SSC-H | Live Dead-H | 112,987 | 6,486 |
| P3 | 22,076 | 1.55% | SSC-H | FSC-H | 194,702 | 1,013,852 |
| P4 | 7,472 | 33.85% | SSC-H | CD3-H | 138,011 | 12,366 |
| P5 | 1,329 | 17.79% | SSC-H | TCR-Va7.2-H | 209,283 | 3,987 |
| R6 | 741 | 55.76% | SSC-H | CD25-H | 232,335 | 6,008 |
| R7 | 69 | 5.19% | SSC-H | CD71-H | 365,825 | 11,885 |
| R8 | 652 | 49.06% | SSC-H | CD69-H | 239,030 | 7,261 |
| R9 | 358 | 26.94% | SSC-H | CD107a-H | 297,335 | 74,597 |
| R10 | 140 | 10.53% | SSC-H | CD137-H | 364,671 | 10,602 |
| R11 | 136 | 10.23% | SSC-H | CD107b-H | 397,808 | 80,248 |
| R12 | 686 | 51.62% | SSC-H | IL-18.Ra-H | 228,150 | 21,839 |

### Report of PBMC 12/09/25-PBMC_LGG_CE_1

Sample Name: PBMC 12/09/25-PBMC_LGG_CE_1 | Run Time: 12-Sep-25 1:42 PM

Cytometer: NovoCyte Quanteon 621181110427 | Software: NovoExpress 1.6.0

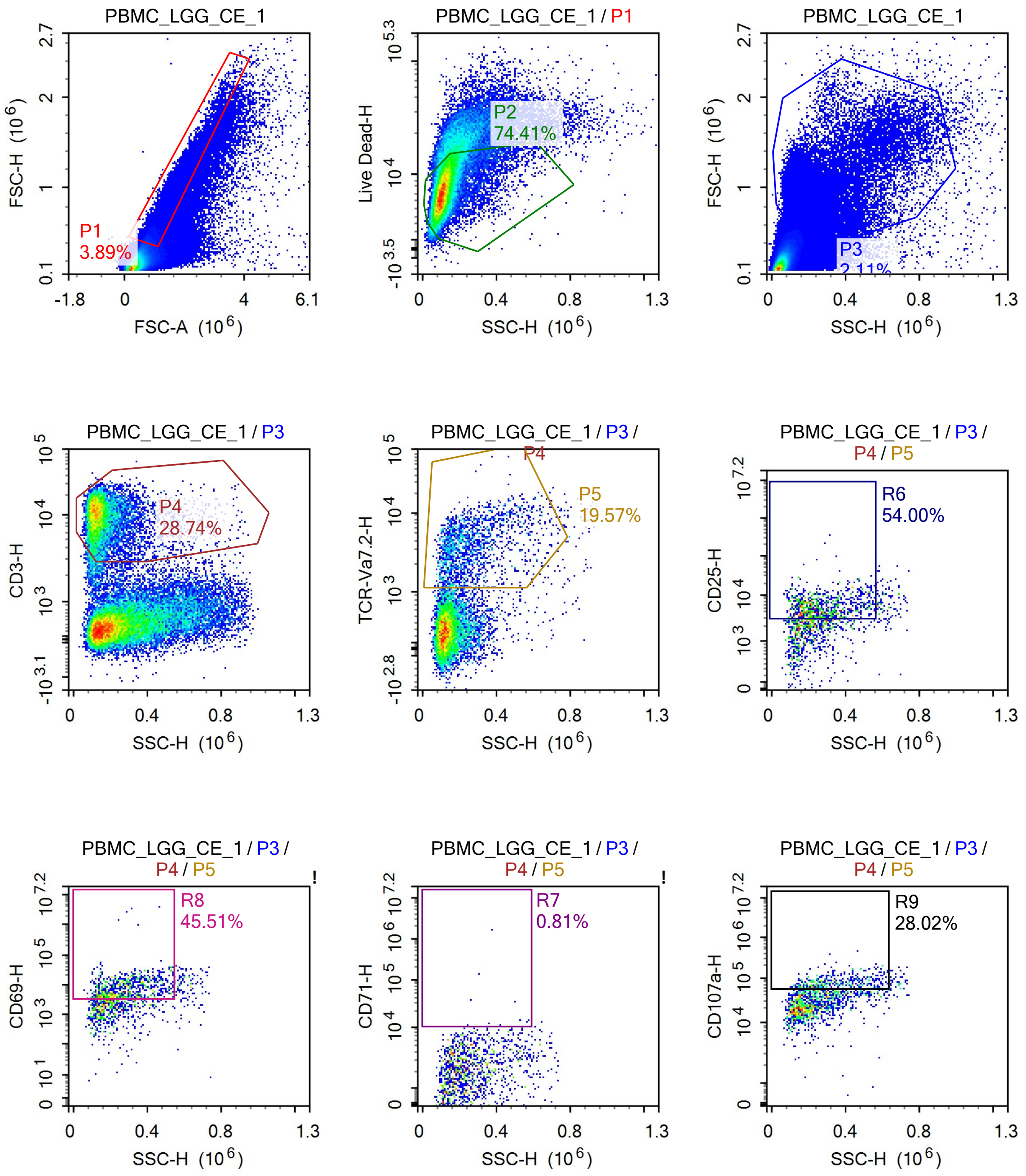

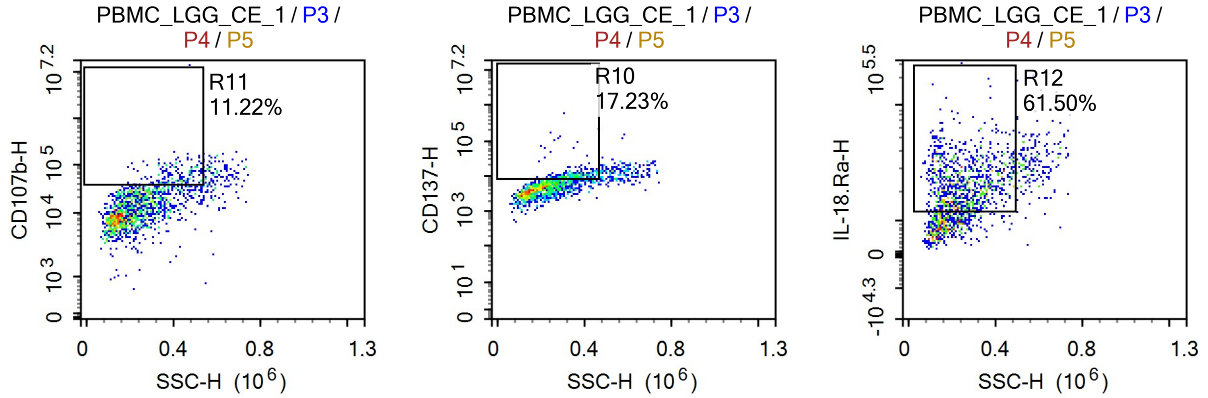


Sample Statistics

| Gate | Count | % Parent | X | Y | Median X | Median Y |
|---|---|---|---|---|---|---|
| All | 1,356,889 | | | | | |
| P1 | 52,799 | 3.89% | FSC-A | FSC-H | 1,033,187 | 505,238 |
| P2 | 39,290 | 74.41% | SSC-H | Live Dead-H | 111,967 | 7,458 |
| P3 | 28,684 | 2.11% | SSC-H | FSC-H | 201,735 | 961,711 |
| P4 | 8,243 | 28.74% | SSC-H | CD3-H | 137,616 | 9,614 |
| P5 | 1,613 | 19.57% | SSC-H | TCR-Va7.2-H | 232,277 | 3,520 |
| R6 | 871 | 54.00% | SSC-H | CD25-H | 259,447 | 5,398 |
| R7 | 13 | 0.81% | SSC-H | CD71-H | 416,473 | 13,403 |
| R8 | 734 | 45.51% | SSC-H | CD69-H | 268,451 | 8,116 |
| R9 | 452 | 28.02% | SSC-H | CD107a-H | 320,684 | 75,155 |
| R10 | 278 | 17.23% | SSC-H | CD137-H | 356,073 | 10,127 |
| R11 | 181 | 11.22% | SSC-H | CD107b-H | 416,318 | 58,835 |
| R12 | 992 | 61.50% | SSC-H | IL-18.Ra-H | 245,212 | 22,724 |

### Report of PBMC 12/09/25-PBMC_LGG_CE_2

Sample Name: PBMC 12/09/25-PBMC_LGG_CE_2 — Run Time: 12-Sep-25 1:44 PM

Cytometer: NovoCyte Quanteon 621181110427 — Software: NovoExpress 1.6.0

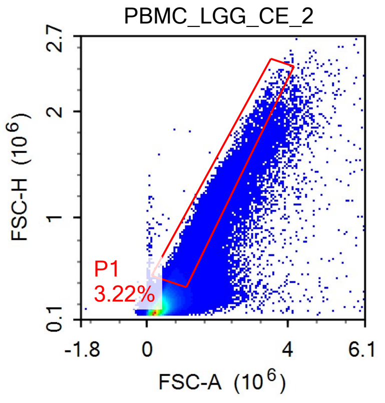


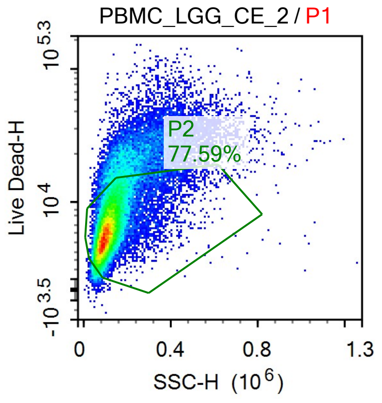


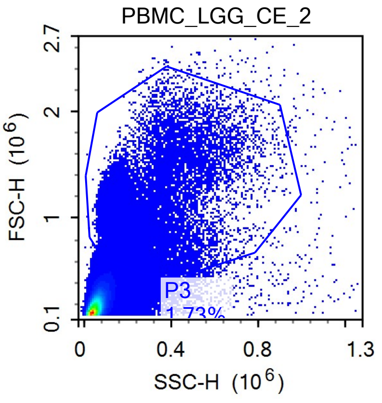


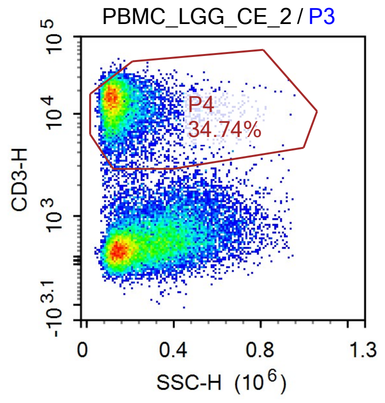


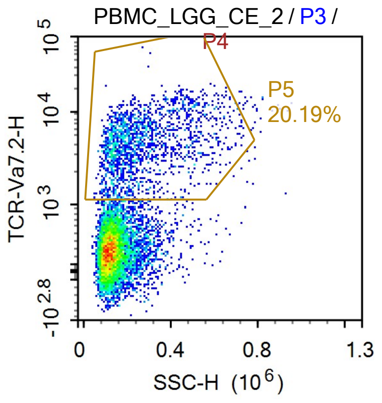


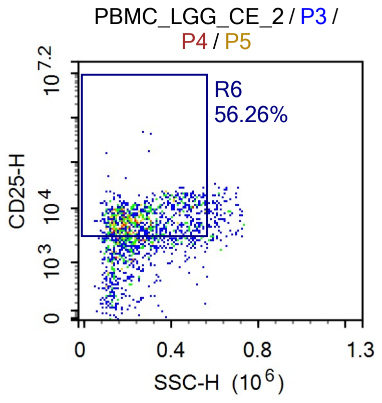


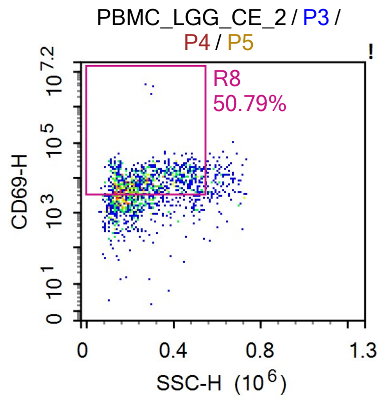


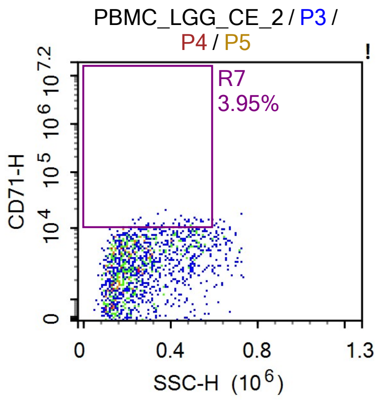


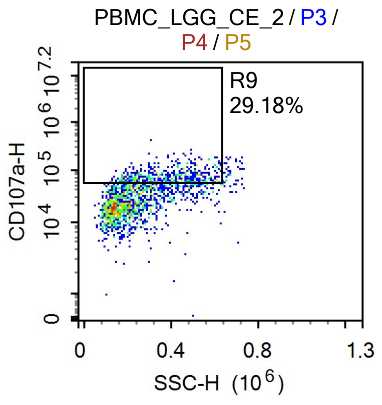

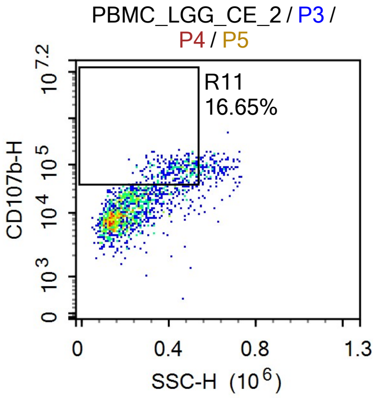


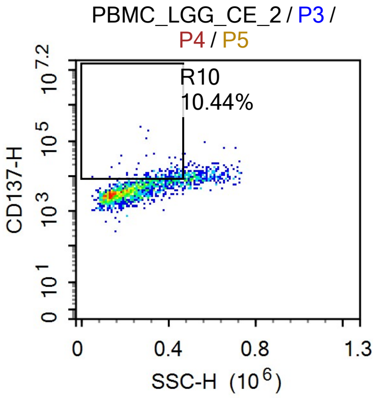


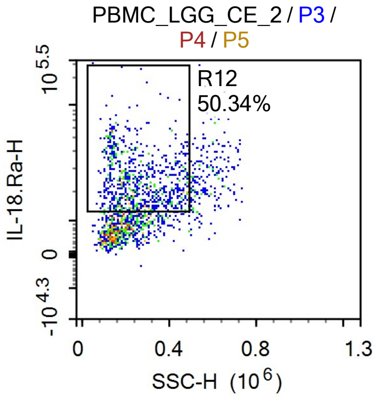


Sample Statistics

| Gate | Count | % Parent | X | Y | Median X | Median Y |
|---|---|---|---|---|---|---|
| All | 1,459,237 | | | | | |
| P1 | 47,023 | 3.22% | FSC-A | FSC-H | 1,028,974 | 510,880 |
| P2 | 36,484 | 77.59% | SSC-H | Live Dead-H | 108,284 | 6,381 |
| P3 | 25,262 | 1.73% | SSC-H | FSC-H | 189,458 | 1,024,603 |
| P4 | 8,775 | 34.74% | SSC-H | CD3-H | 135,613 | 13,359 |
| P5 | 1,772 | 20.19% | SSC-H | TCR-Va7.2-H | 228,394 | 4,174 |
| R6 | 997 | 56.26% | SSC-H | CD25-H | 248,570 | 6,101 |
| R7 | 70 | 3.95% | SSC-H | CD71-H | 385,804 | 10,949 |
| R8 | 900 | 50.79% | SSC-H | CD69-H | 257,739 | 7,951 |
| R9 | 517 | 29.18% | SSC-H | CD107a-H | 388,081 | 71,632 |
| R10 | 185 | 10.44% | SSC-H | CD137-H | 371,264 | 9,320 |
| R11 | 295 | 16.65% | SSC-H | CD107b-H | 437,240 | 78,268 |
| R12 | 892 | 50.34% | SSC-H | IL-18.Ra-H | 242,549 | 22,286 |

### Report of PBMC 12/09/25-PBMC_LGG_CE_3

Sample Name: PBMC 12/09/25-PBMC_LGG_CE_3 | Run Time: 12-Sep-25 1:46 PM

Cytometer: NovoCyte Quanteon 621181110427 | Software: NovoExpress 1.6.0

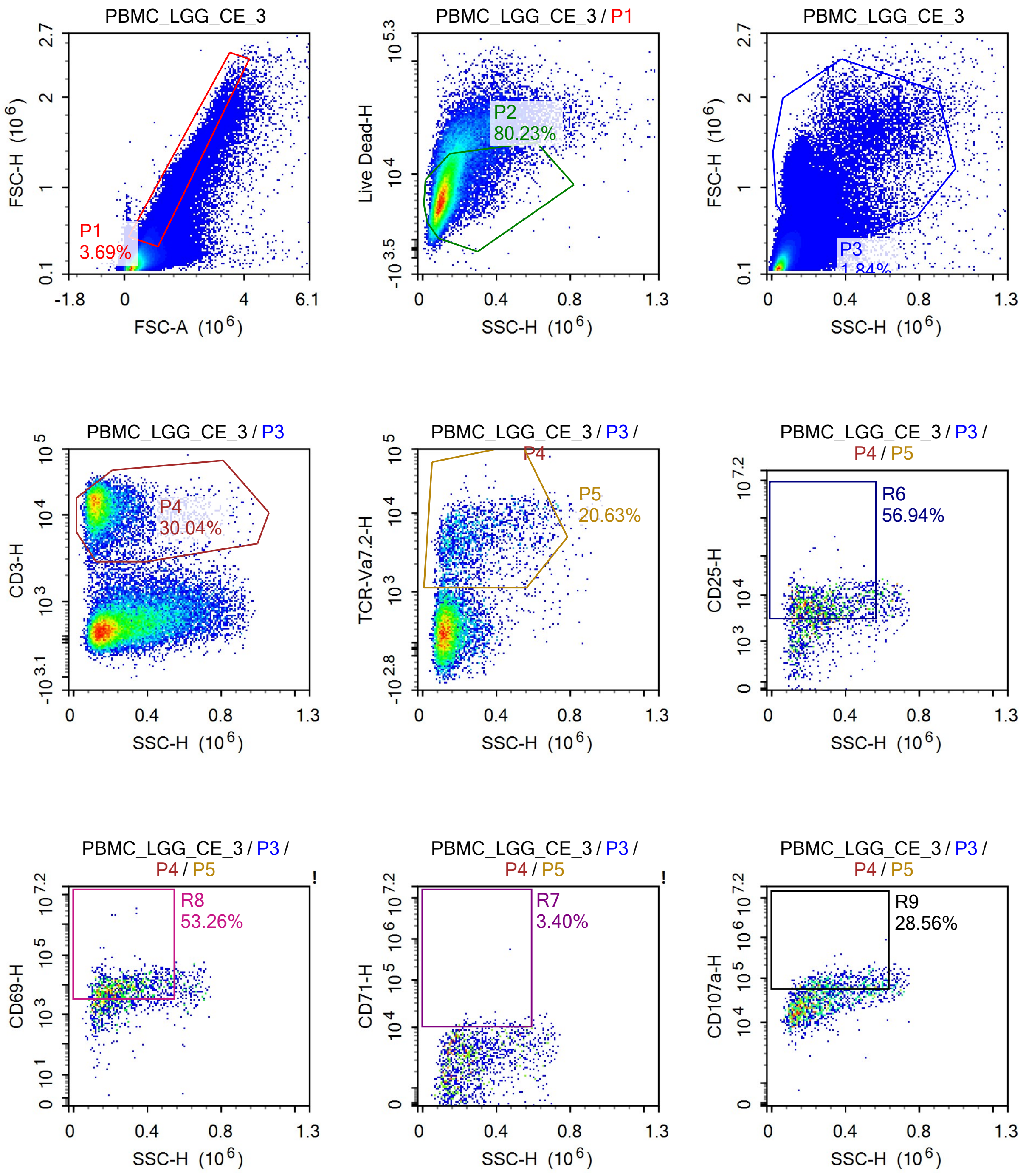

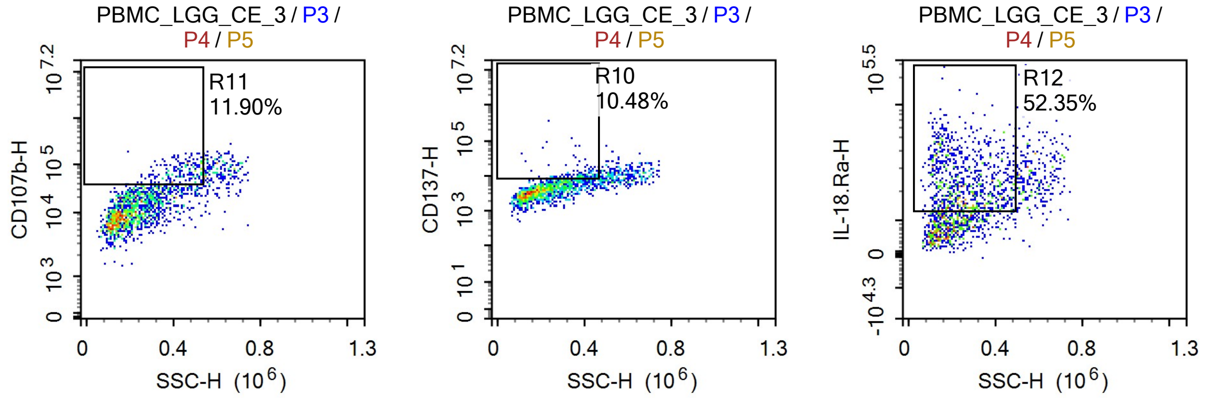


Sample Statistics

| Gate | Count | % Parent | X | Y | Median X | Median Y |
|---|---|---|---|---|---|---|
| All | 1,544,283 | | | | | |
| P1 | 56,961 | 3.69% | FSC-A | FSC-H | 1,001,051 | 493,706 |
| P2 | 45,701 | 80.23% | SSC-H | Live Dead-H | 111,200 | 6,855 |
| P3 | 28,473 | 1.84% | SSC-H | FSC-H | 192,141 | 963,794 |
| P4 | 8,554 | 30.04% | SSC-H | CD3-H | 137,900 | 12,974 |
| P5 | 1,765 | 20.63% | SSC-H | TCR-Va7.2-H | 241,230 | 4,288 |
| R6 | 1,005 | 56.94% | SSC-H | CD25-H | 261,272 | 6,265 |
| R7 | 60 | 3.40% | SSC-H | CD71-H | 387,281 | 11,803 |
| R8 | 940 | 53.26% | SSC-H | CD69-H | 258,529 | 8,679 |
| R9 | 504 | 28.56% | SSC-H | CD107a-H | 360,482 | 73,682 |
| R10 | 185 | 10.48% | SSC-H | CD137-H | 371,748 | 9,528 |
| R11 | 210 | 11.90% | SSC-H | CD107b-H | 432,002 | 67,874 |
| R12 | 924 | 52.35% | SSC-H | IL-18.Ra-H | 244,086 | 22,310 |

### Report of PBMC 12/09/25-PBMC_LGG_CE_4

Sample Name: PBMC 12/09/25-PBMC_LGG_CE_4 | Run Time: 12-Sep-25 1:48 PM

Cytometer: NovoCyte Quanteon 621181110427 | Software: NovoExpress 1.6.0

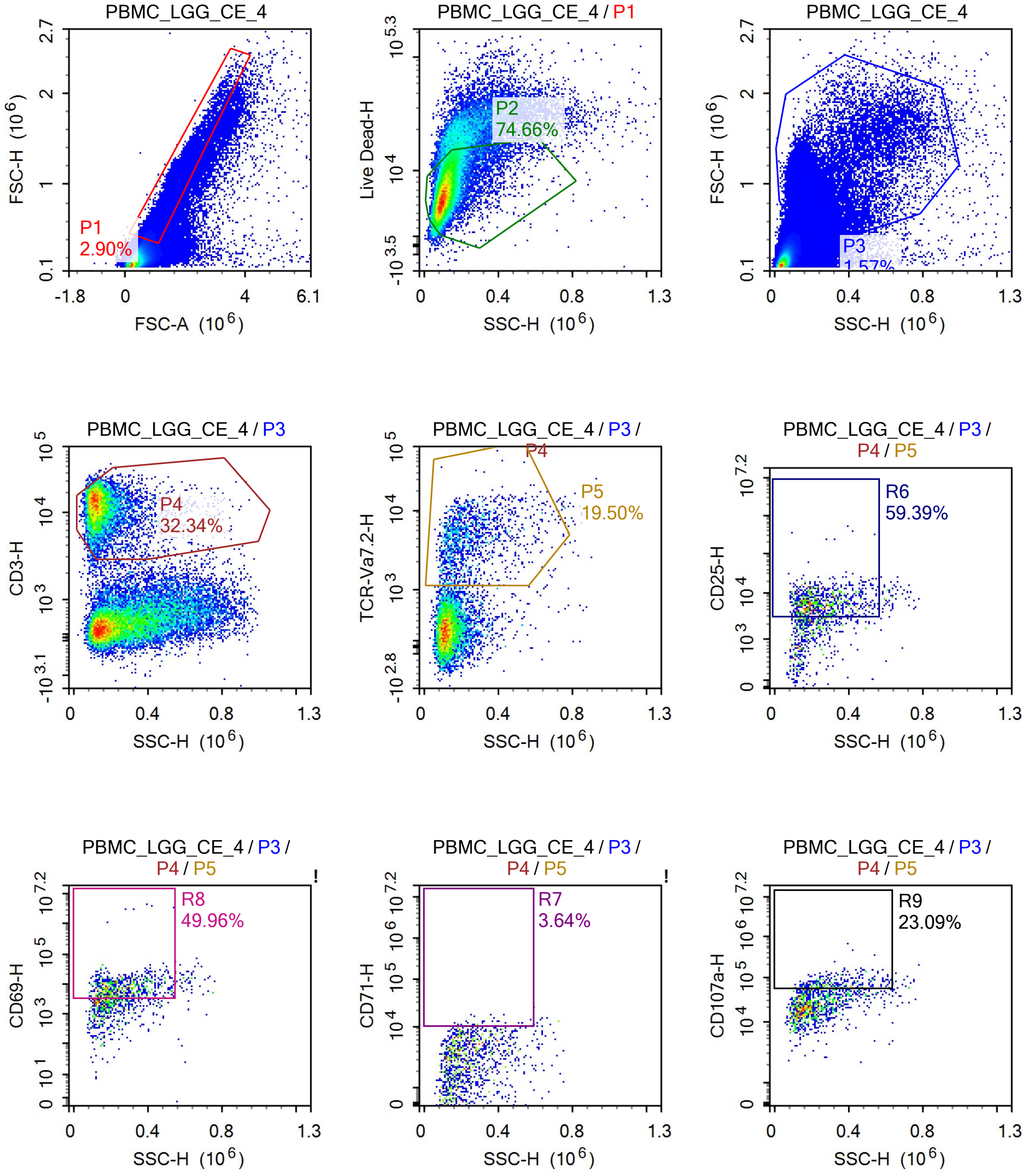

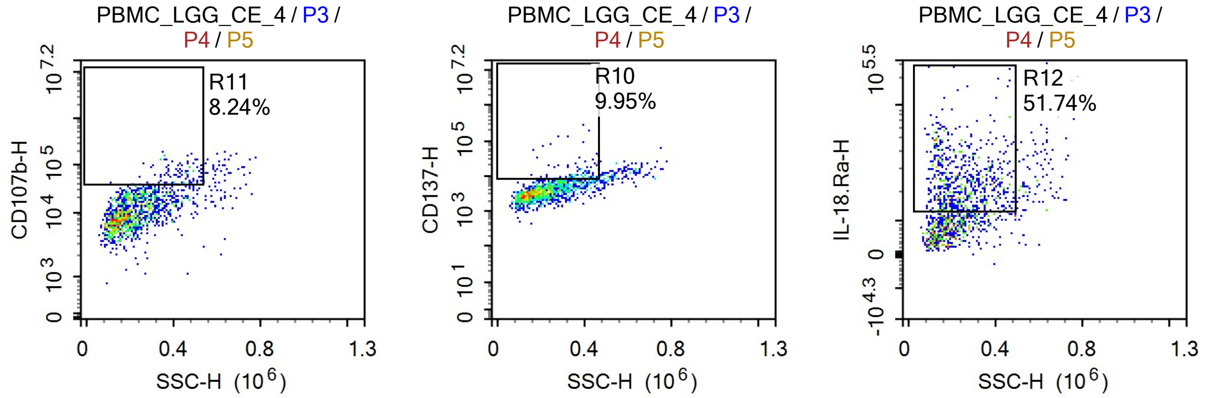


Sample Statistics

| Gate | Count | % Parent | X | Y | Median X | Median Y |
|---|---|---|---|---|---|---|
| All | 1,361,752 | | | | | |
| P1 | 39,521 | 2.90% | FSC-A | FSC-H | 1,028,901 | 512,722 |
| P2 | 29,508 | 74.66% | SSC-H | Live Dead-H | 110,543 | 6,626 |
| P3 | 21,363 | 1.57% | SSC-H | FSC-H | 194,178 | 1,030,468 |
| P4 | 6,909 | 32.34% | SSC-H | CD3-H | 131,781 | 12,193 |
| P5 | 1,347 | 19.50% | SSC-H | TCR-Va7.2-H | 212,371 | 4,156 |
| R6 | 800 | 59.39% | SSC-H | CD25-H | 233,403 | 6,106 |
| R7 | 49 | 3.64% | SSC-H | CD71-H | 312,712 | 11,123 |
| R8 | 673 | 49.96% | SSC-H | CD69-H | 243,804 | 7,974 |
| R9 | 311 | 23.09% | SSC-H | CD107a-H | 310,368 | 70,165 |
| R10 | 134 | 9.95% | SSC-H | CD137-H | 367,985 | 9,913 |
| R11 | 111 | 8.24% | SSC-H | CD107b-H | 413,259 | 60,696 |
| R12 | 697 | 51.74% | SSC-H | IL-18.Ra-H | 234,608 | 22,728 |

### Report of PBMC 12/09/25-

Sample Name: PBMC 12/09/25-PBMC_Candida_SN_1 Run Time: 12-Sep-25 1:50 PM

Cytometer: NovoCyte Quanteon 621181110427 Software: NovoExpress 1.6.0

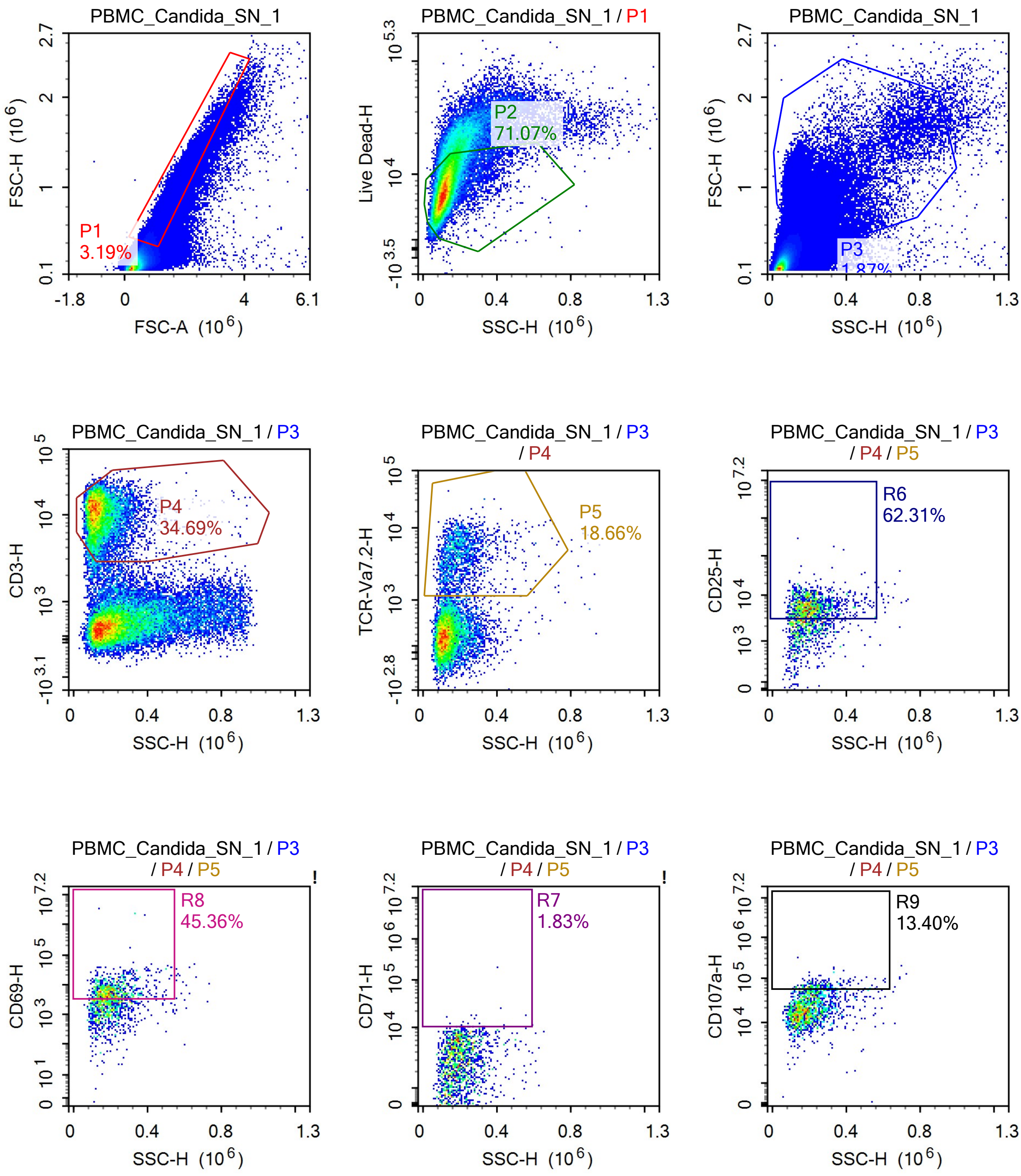

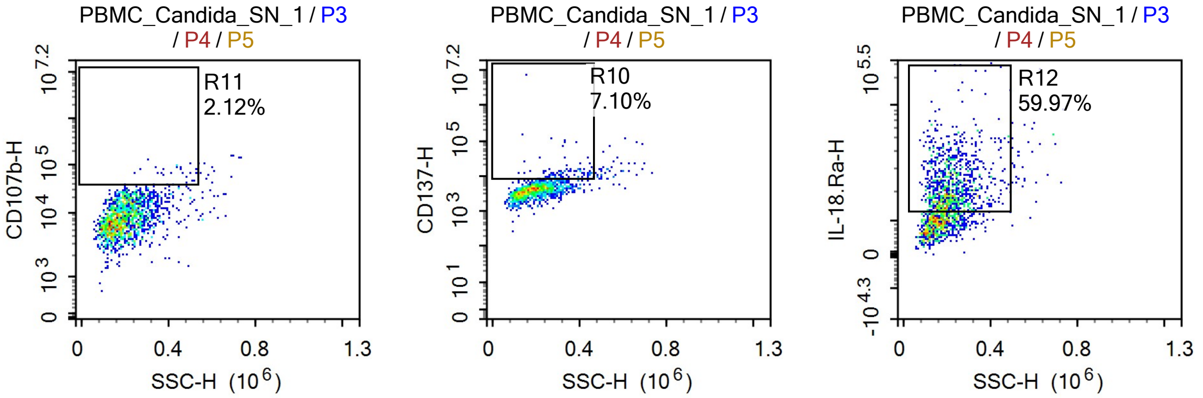


Sample Statistics

| Gate | Count | % Parent | X | Y | Median X | Median Y |
|---|---|---|---|---|---|---|
| All | 1,438,412 | | | | | |
| P1 | 45,877 | 3.19% | FSC-A | FSC-H | 1,082,465 | 556,281 |
| P2 | 32,604 | 71.07% | SSC-H | Live Dead-H | 117,114 | 7,623 |
| P3 | 26,968 | 1.87% | SSC-H | FSC-H | 186,591 | 978,540 |
| P4 | 9,355 | 34.69% | SSC-H | CD3-H | 132,041 | 10,808 |
| P5 | 1,746 | 18.66% | SSC-H | TCR-Va7.2-H | 189,047 | 3,909 |
| R6 | 1,088 | 62.31% | SSC-H | CD25-H | 200,699 | 5,480 |
| R7 | 32 | 1.83% | SSC-H | CD71-H | 270,098 | 10,844 |
| R8 | 792 | 45.36% | SSC-H | CD69-H | 202,468 | 6,411 |
| R9 | 234 | 13.40% | SSC-H | CD107a-H | 252,658 | 63,939 |
| R10 | 124 | 7.10% | SSC-H | CD137-H | 323,810 | 11,591 |
| R11 | 37 | 2.12% | SSC-H | CD107b-H | 418,688 | 67,879 |
| R12 | 1,047 | 59.97% | SSC-H | IL-18.Ra-H | 210,046 | 20,923 |

### Report of PBMC 12/09/25-

Sample Name: PBMC 12/09/25-PBMC_Candida_SN_2 | Run Time: 12-Sep-25 1:52 PM

Cytometer: NovoCyte Quanteon 621181110427 | Software: NovoExpress 1.6.0

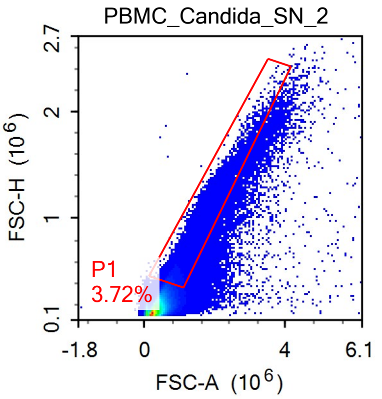


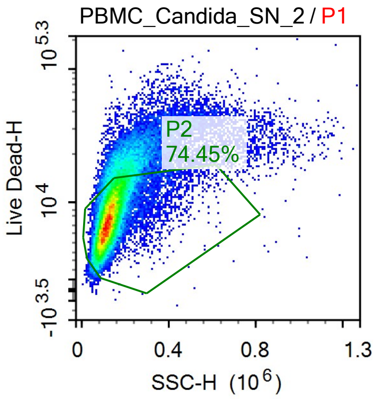


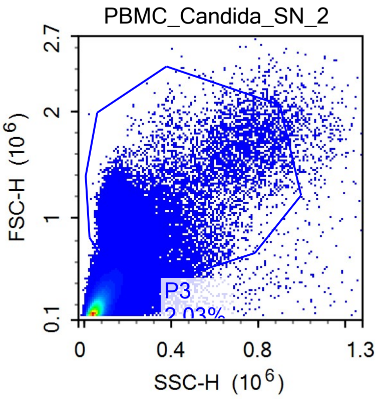


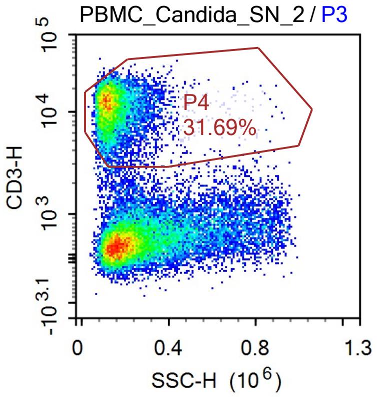


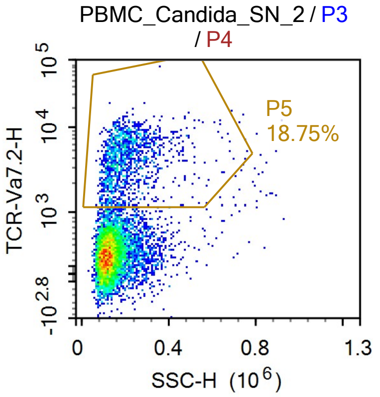


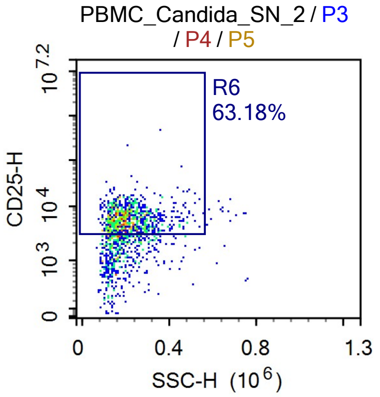


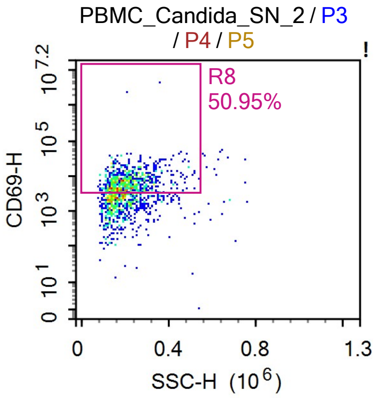


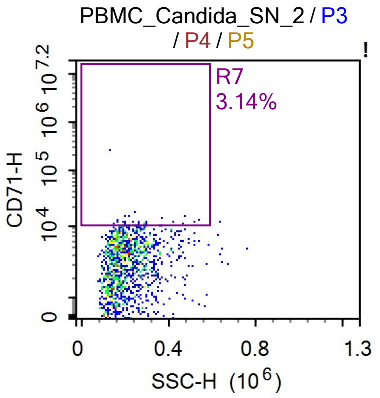


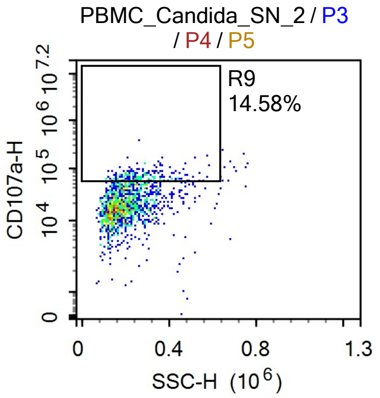

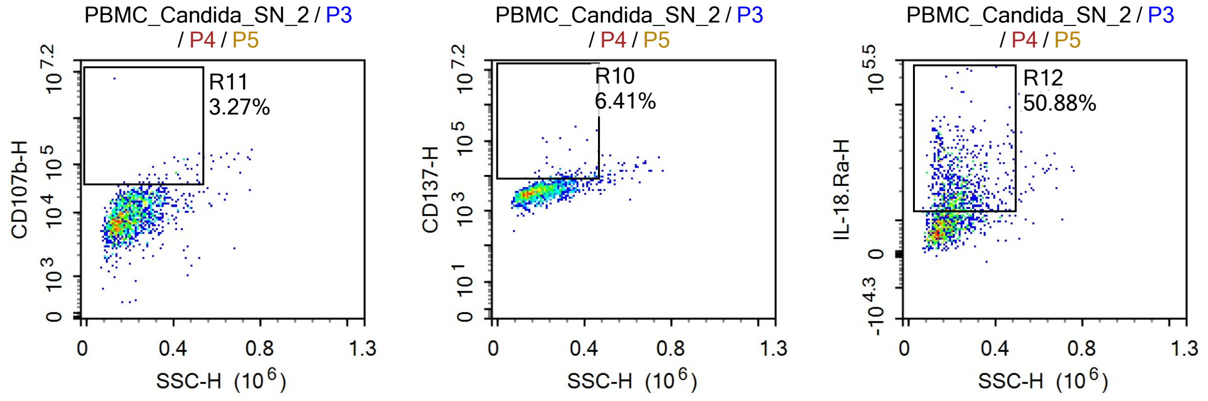


Sample Statistics

| Gate | Count | % Parent | X | Y | Median X | Median Y |
|---|---|---|---|---|---|---|
| All | 1,267,702 | | | | | |
| P1 | 47,126 | 3.72% | FSC-A | FSC-H | 1,031,207 | 526,711 |
| P2 | 35,083 | 74.45% | SSC-H | Live Dead-H | 123,916 | 7,730 |
| P3 | 25,724 | 2.03% | SSC-H | FSC-H | 194,528 | 935,118 |
| P4 | 8,153 | 31.69% | SSC-H | CD3-H | 133,344 | 11,749 |
| P5 | 1,529 | 18.75% | SSC-H | TCR-Va7.2-H | 188,794 | 3,980 |
| R6 | 966 | 63.18% | SSC-H | CD25-H | 201,432 | 5,621 |
| R7 | 48 | 3.14% | SSC-H | CD71-H | 233,681 | 10,948 |
| R8 | 779 | 50.95% | SSC-H | CD69-H | 206,353 | 6,862 |
| R9 | 223 | 14.58% | SSC-H | CD107a-H | 256,055 | 67,157 |
| R10 | 98 | 6.41% | SSC-H | CD137-H | 331,762 | 10,024 |
| R11 | 50 | 3.27% | SSC-H | CD107b-H | 409,107 | 62,345 |
| R12 | 778 | 50.88% | SSC-H | IL-18.Ra-H | 216,517 | 20,160 |

### Report of PBMC 12/09/25-

Sample Name: PBMC 12/09/25-PBMC_Candida_SN_3 | Run Time: 12-Sep-25 1:54 PM

Cytometer: NovoCyte Quanteon 621181110427 | Software: NovoExpress 1.6.0

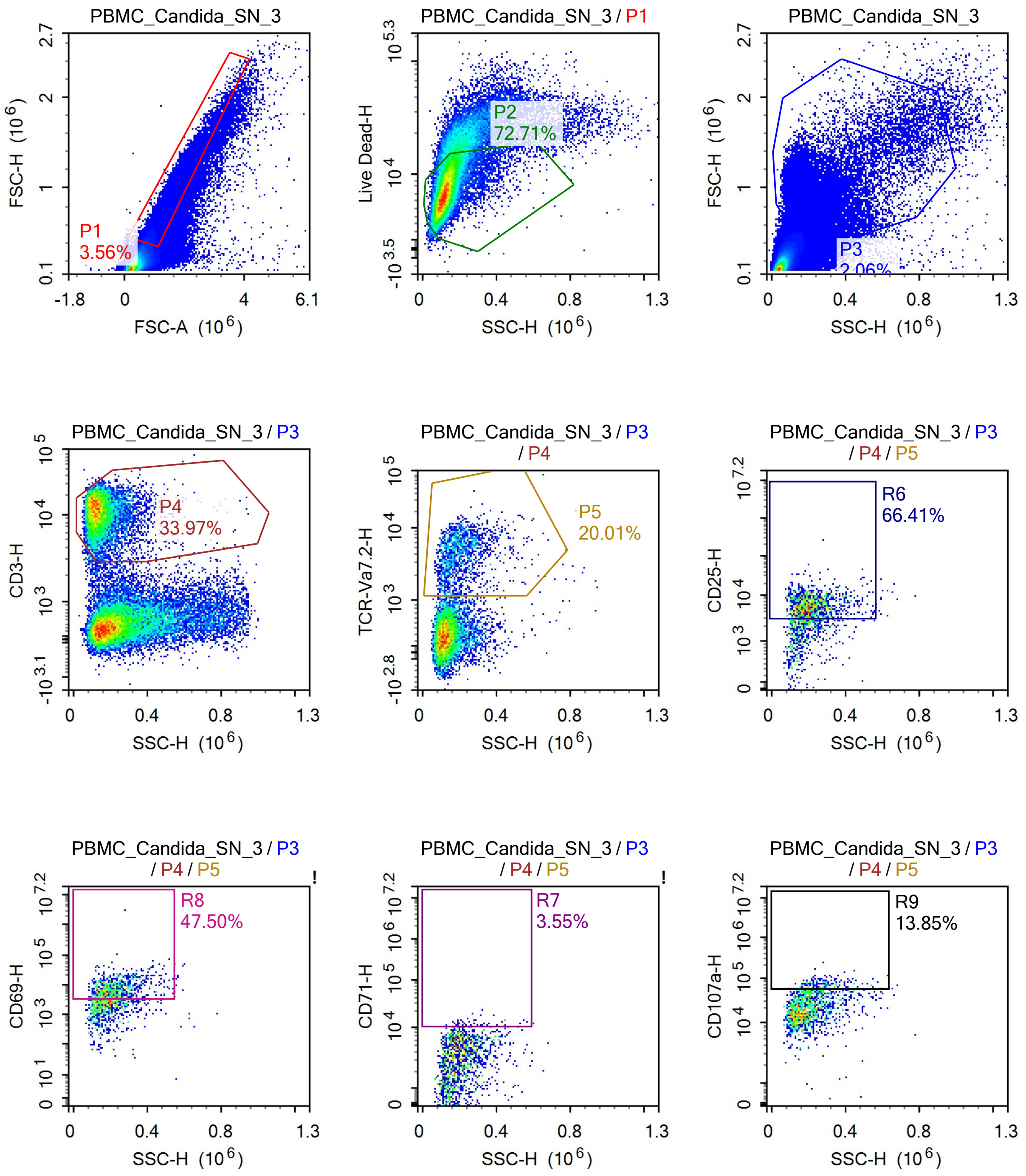

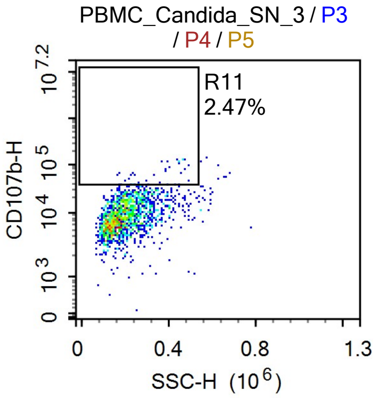


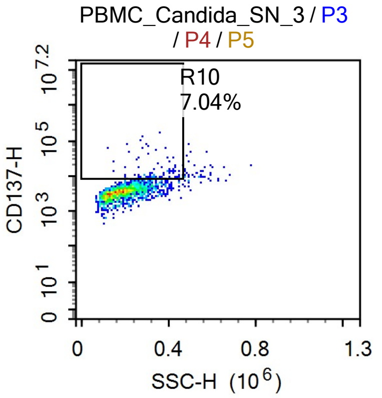


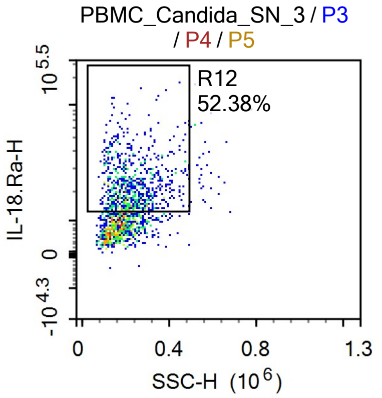


Sample Statistics

| Gate | Count | % Parent | X | Y | Median X | Median Y |
|---|---|---|---|---|---|---|
| All | 1,184,725 | | | | | |
| P1 | 42,211 | 3.56% | FSC-A | FSC-H | 1,065,805 | 546,428 |
| P2 | 30,692 | 72.71% | SSC-H | Live Dead-H | 123,219 | 7,449 |
| P3 | 24,436 | 2.06% | SSC-H | FSC-H | 191,455 | 959,176 |
| P4 | 8,301 | 33.97% | SSC-H | CD3-H | 132,882 | 11,220 |
| P5 | 1,661 | 20.01% | SSC-H | TCR-Va7.2-H | 191,027 | 4,272 |
| R6 | 1,103 | 66.41% | SSC-H | CD25-H | 204,556 | 5,799 |
| R7 | 59 | 3.55% | SSC-H | CD71-H | 259,801 | 11,681 |
| R8 | 789 | 47.50% | SSC-H | CD69-H | 212,422 | 7,169 |
| R9 | 230 | 13.85% | SSC-H | CD107a-H | 257,971 | 66,012 |
| R10 | 117 | 7.04% | SSC-H | CD137-H | 328,378 | 10,266 |
| R11 | 41 | 2.47% | SSC-H | CD107b-H | 332,680 | 59,129 |
| R12 | 870 | 52.38% | SSC-H | IL-18.Ra-H | 214,131 | 20,458 |

### Report of PBMC 12/09/25-

Sample Name: PBMC 12/09/25-PBMC_Candida_SN_4 | Run Time: 12-Sep-25 1:56 PM

Cytometer: NovoCyte Quanteon 621181110427 | Software: NovoExpress 1.6.0

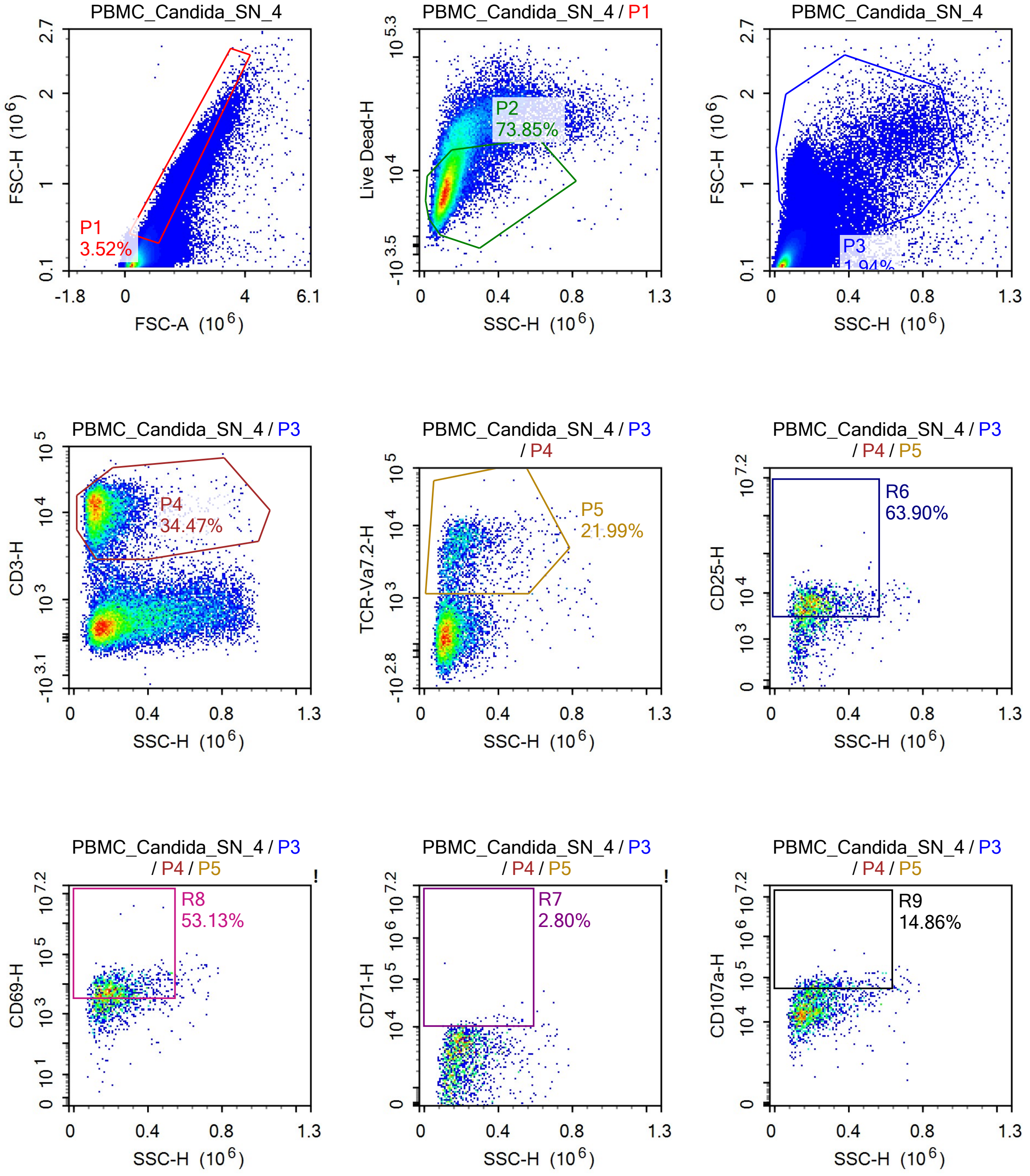

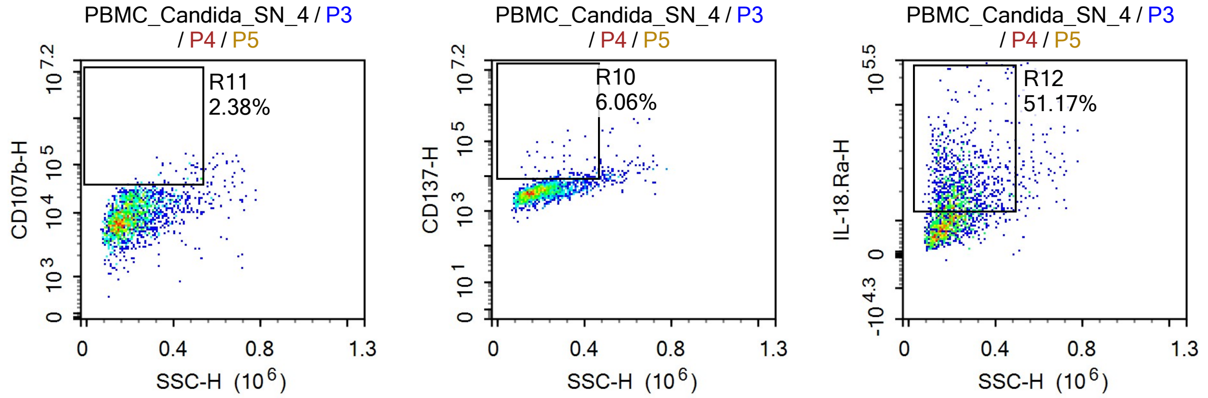


Sample Statistics

| Gate | Count | % Parent | X | Y | Median X | Median Y |
|---|---|---|---|---|---|---|
| All | 1,315,223 | | | | | |
| P1 | 46,349 | 3.52% | FSC-A | FSC-H | 1,037,458 | 531,135 |
| P2 | 34,228 | 73.85% | SSC-H | Live Dead-H | 120,424 | 7,426 |
| P3 | 25,478 | 1.94% | SSC-H | FSC-H | 189,138 | 957,530 |
| P4 | 8,782 | 34.47% | SSC-H | CD3-H | 134,255 | 11,210 |
| P5 | 1,931 | 21.99% | SSC-H | TCR-Va7.2-H | 195,023 | 4,329 |
| R6 | 1,234 | 63.90% | SSC-H | CD25-H | 208,417 | 5,720 |
| R7 | 54 | 2.80% | SSC-H | CD71-H | 275,092 | 11,370 |
| R8 | 1,026 | 53.13% | SSC-H | CD69-H | 208,859 | 6,677 |
| R9 | 287 | 14.86% | SSC-H | CD107a-H | 272,521 | 67,971 |
| R10 | 117 | 6.06% | SSC-H | CD137-H | 326,729 | 9,864 |
| R11 | 46 | 2.38% | SSC-H | CD107b-H | 450,602 | 58,990 |
| R12 | 988 | 51.17% | SSC-H | IL-18.Ra-H | 219,200 | 20,393 |

### Report of PBMC 12/09/25-PBMC_V336_CE_1

Sample Name: PBMC 12/09/25-PBMC_V336_CE_1 Run Time: 12-Sep-25 1:58 PM

Cytometer: NovoCyte Quanteon 621181110427 Software: NovoExpress 1.6.0

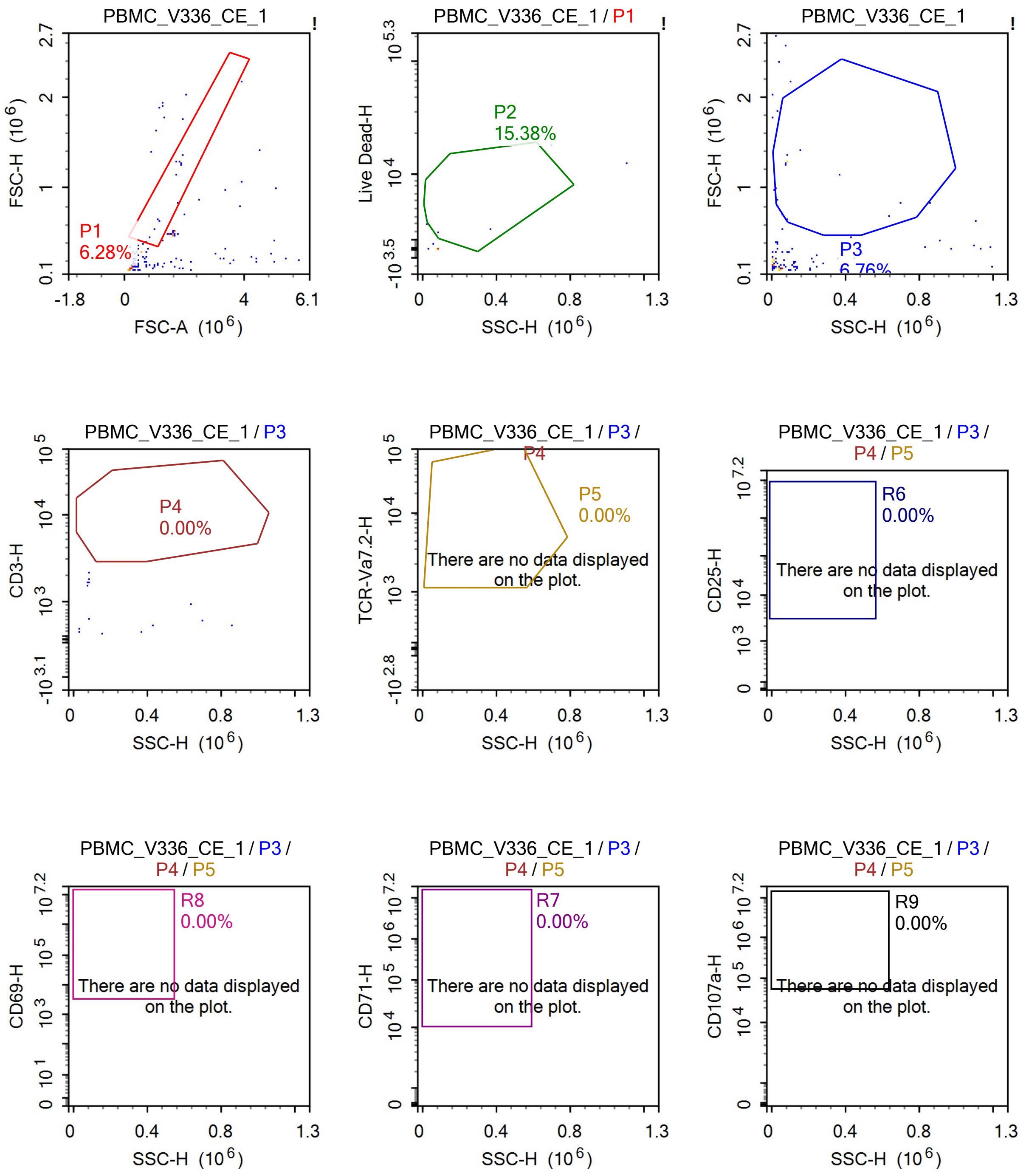

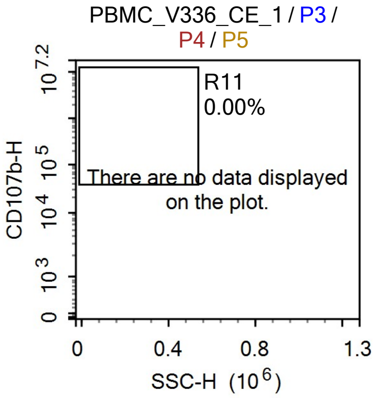


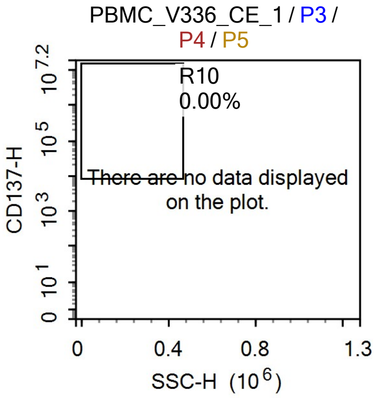


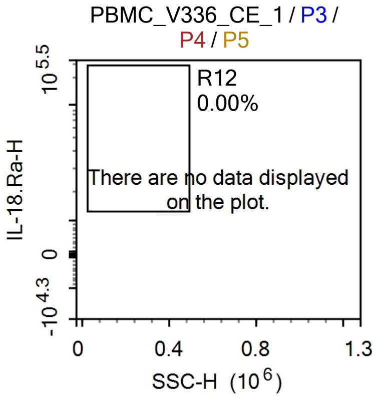


Sample Statistics

| Gate | Count | % Parent | X | Y | Median X | Median Y |
|---|---|---|---|---|---|---|
| All | 207 | | | | | |
| P1 | 13 | 6.28% | FSC-A | FSC-H | 1,648,905 | 842,738 |
| P2 | 2 | 15.38% | SSC-H | Live Dead-H | 219,988 | 2,274 |
| P3 | 14 | 6.76% | SSC-H | FSC-H | 82,830 | 1,009,896 |
| P4 | 0 | 0.00% | SSC-H | CD3-H | 0 | 0 |
| P5 | 0 | 0.00% | SSC-H | TCR-Va7.2-H | 0 | 0 |
| R6 | 0 | 0.00% | SSC-H | CD25-H | 0 | 0 |
| R7 | 0 | 0.00% | SSC-H | CD71-H | 0 | 0 |
| R8 | 0 | 0.00% | SSC-H | CD69-H | 0 | 0 |
| R9 | 0 | 0.00% | SSC-H | CD107a-H | 0 | 0 |
| R10 | 0 | 0.00% | SSC-H | CD137-H | 0 | 0 |
| R11 | 0 | 0.00% | SSC-H | CD107b-H | 0 | 0 |
| R12 | 0 | 0.00% | SSC-H | IL-18.Ra-H | 0 | 0 |

### Report of PBMC 12/09/25-PBMC_V336_CE_2

Sample Name: PBMC 12/09/25-PBMC_V336_CE_2 Run Time: 12-Sep-25 2:00 PM

Cytometer: NovoCyte Quanteon 621181110427 Software: NovoExpress 1.6.0

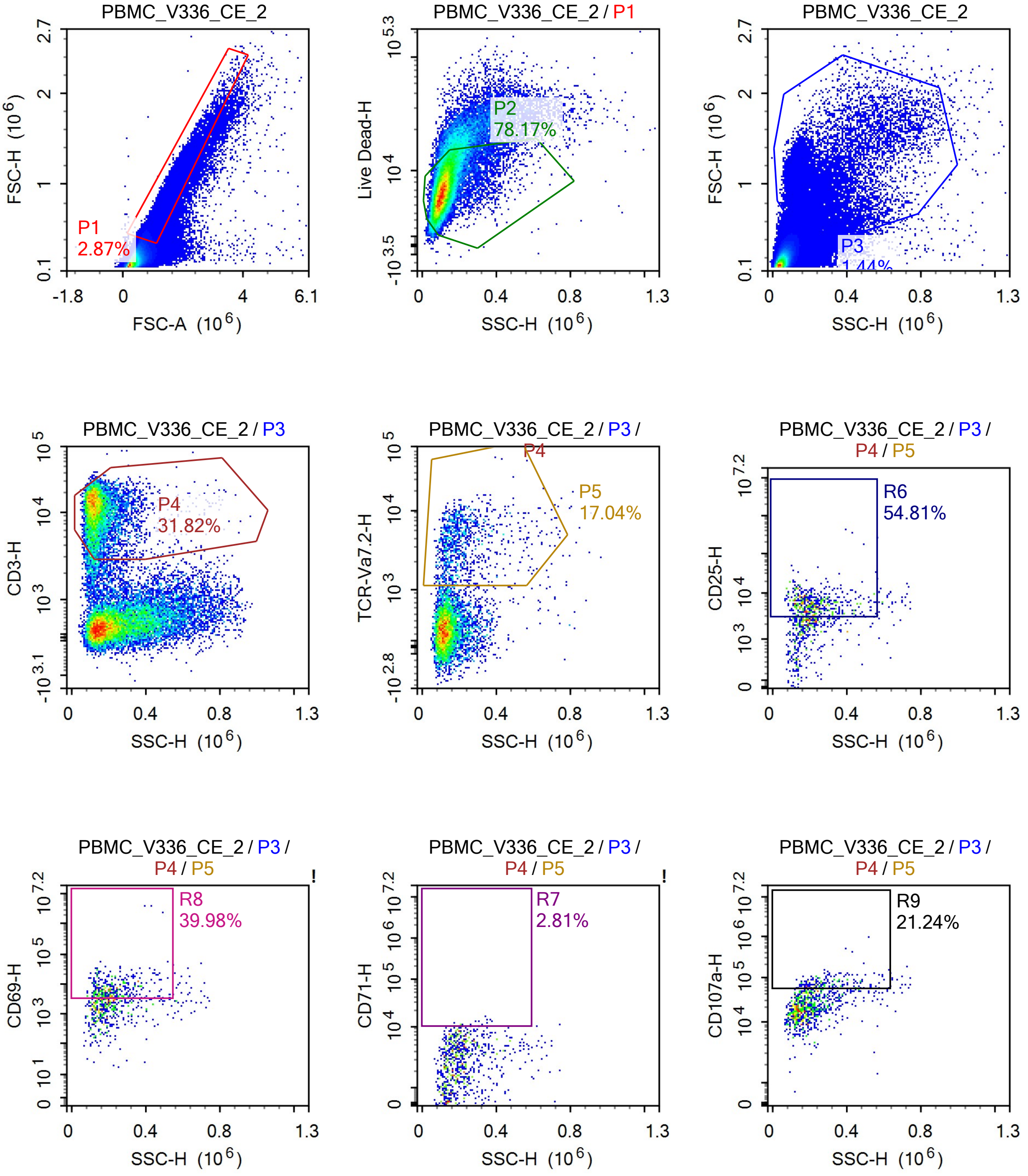

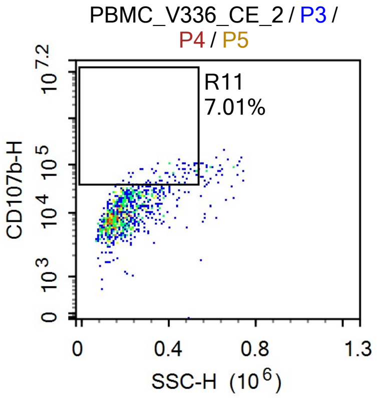


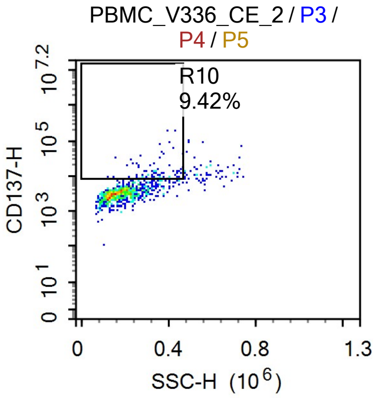


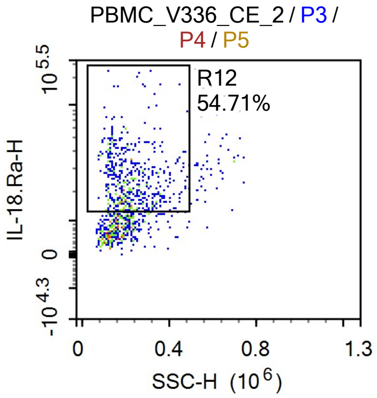


Sample Statistics

| Gate | Count | % Parent | X | Y | Median X | Median Y |
|---|---|---|---|---|---|---|
| All | 1,274,696 | | | | | |
| P1 | 36,590 | 2.87% | FSC-A | FSC-H | 1,010,239 | 509,907 |
| P2 | 28,604 | 78.17% | SSC-H | Live Dead-H | 113,651 | 7,168 |
| P3 | 18,407 | 1.44% | SSC-H | FSC-H | 180,205 | 993,974 |
| P4 | 5,858 | 31.82% | SSC-H | CD3-H | 131,391 | 11,207 |
| P5 | 998 | 17.04% | SSC-H | TCR-Va7.2-H | 189,132 | 3,717 |
| R6 | 547 | 54.81% | SSC-H | CD25-H | 206,240 | 5,759 |
| R7 | 28 | 2.81% | SSC-H | CD71-H | 258,538 | 11,205 |
| R8 | 399 | 39.98% | SSC-H | CD69-H | 209,723 | 6,549 |
| R9 | 212 | 21.24% | SSC-H | CD107a-H | 273,337 | 72,102 |
| R10 | 94 | 9.42% | SSC-H | CD137-H | 336,627 | 11,215 |
| R11 | 70 | 7.01% | SSC-H | CD107b-H | 401,988 | 63,974 |
| R12 | 546 | 54.71% | SSC-H | IL-18.Ra-H | 203,491 | 22,826 |

### Report of PBMC 12/09/25-PBMC_V336_CE_3

Sample Name: PBMC 12/09/25-PBMC_V336_CE_3 Run Time: 12-Sep-25 2:02 PM

Cytometer: NovoCyte Quanteon 621181110427 Software: NovoExpress 1.6.0

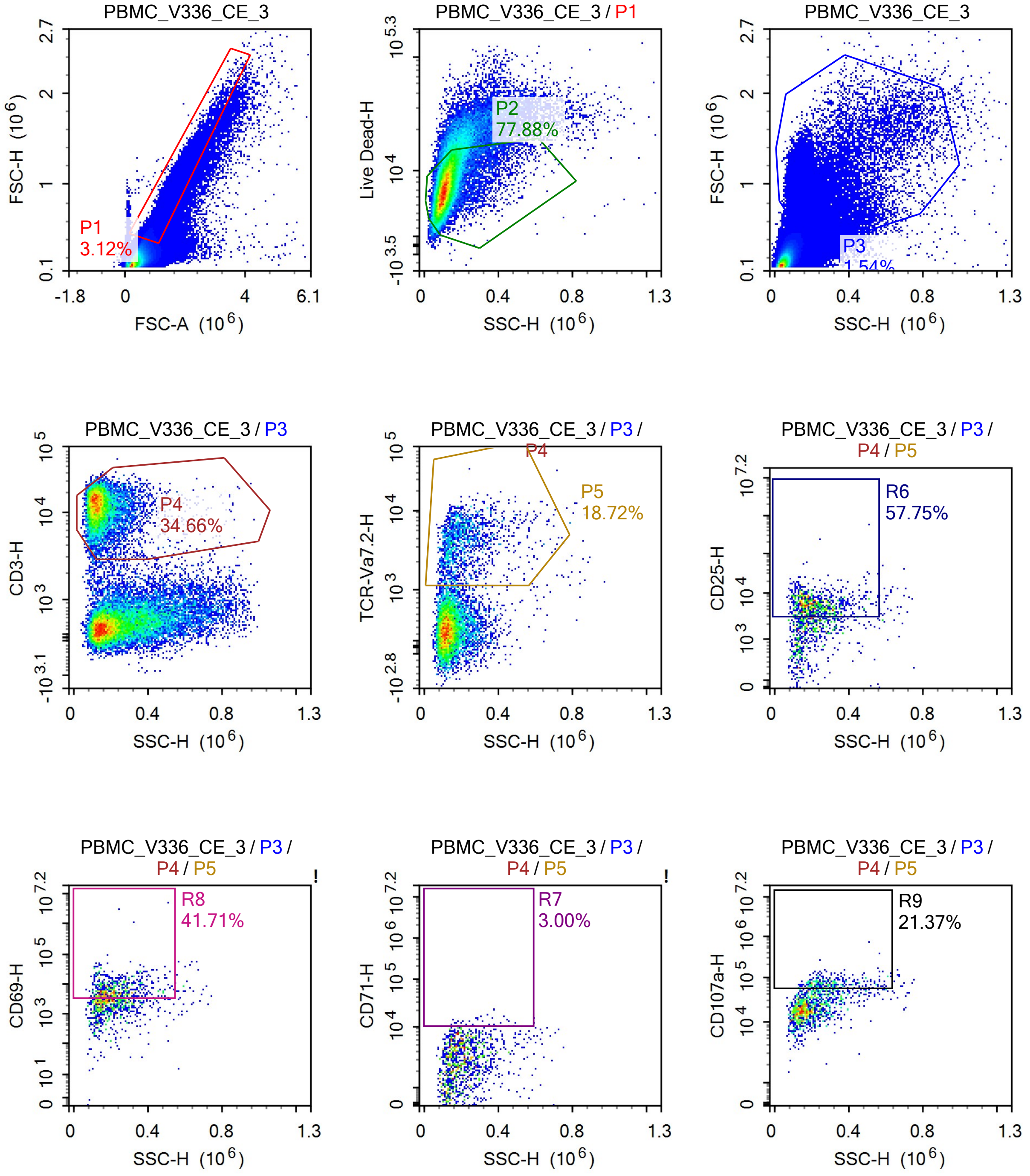

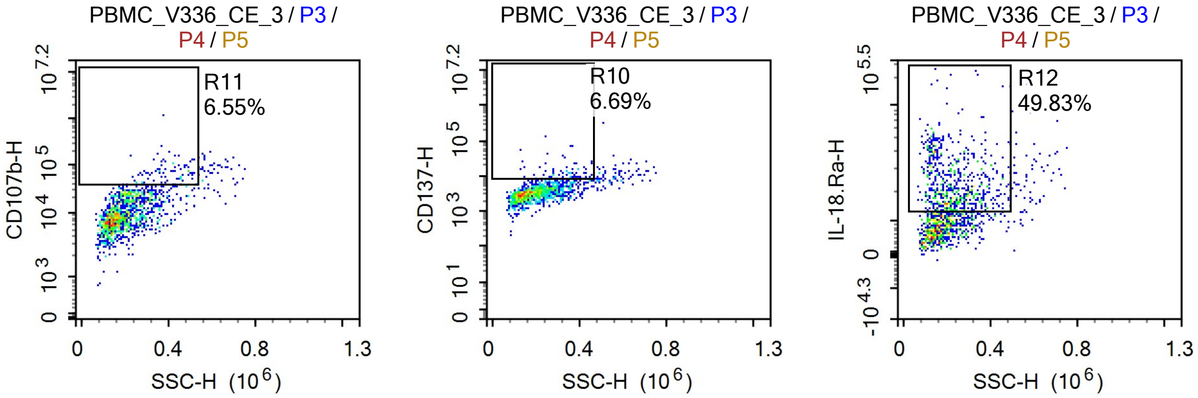


Sample Statistics

| Gate | Count | % Parent | X | Y | Median X | Median Y |
|---|---|---|---|---|---|---|
| All | 1,467,940 | | | | | |
| P1 | 45,751 | 3.12% | FSC-A | FSC-H | 1,005,883 | 502,673 |
| P2 | 35,633 | 77.88% | SSC-H | Live Dead-H | 113,893 | 7,428 |
| P3 | 22,583 | 1.54% | SSC-H | FSC-H | 182,078 | 994,635 |
| P4 | 7,827 | 34.66% | SSC-H | CD3-H | 134,327 | 12,234 |
| P5 | 1,465 | 18.72% | SSC-H | TCR-Va7.2-H | 193,136 | 3,781 |
| R6 | 846 | 57.75% | SSC-H | CD25-H | 204,753 | 5,571 |
| R7 | 44 | 3.00% | SSC-H | CD71-H | 281,285 | 11,755 |
| R8 | 611 | 41.71% | SSC-H | CD69-H | 209,170 | 6,263 |
| R9 | 313 | 21.37% | SSC-H | CD107a-H | 281,944 | 71,353 |
| R10 | 98 | 6.69% | SSC-H | CD137-H | 347,601 | 11,225 |
| R11 | 96 | 6.55% | SSC-H | CD107b-H | 388,038 | 65,986 |
| R12 | 730 | 49.83% | SSC-H | IL-18.Ra-H | 214,808 | 20,810 |

### Report of PBMC 12/09/25-PBMC_V336_CE_4

Sample Name: PBMC 12/09/25-PBMC_V336_CE_4 Run Time: 12-Sep-25 2:04 PM

Cytometer: NovoCyte Quanteon 621181110427 Software: NovoExpress 1.6.0

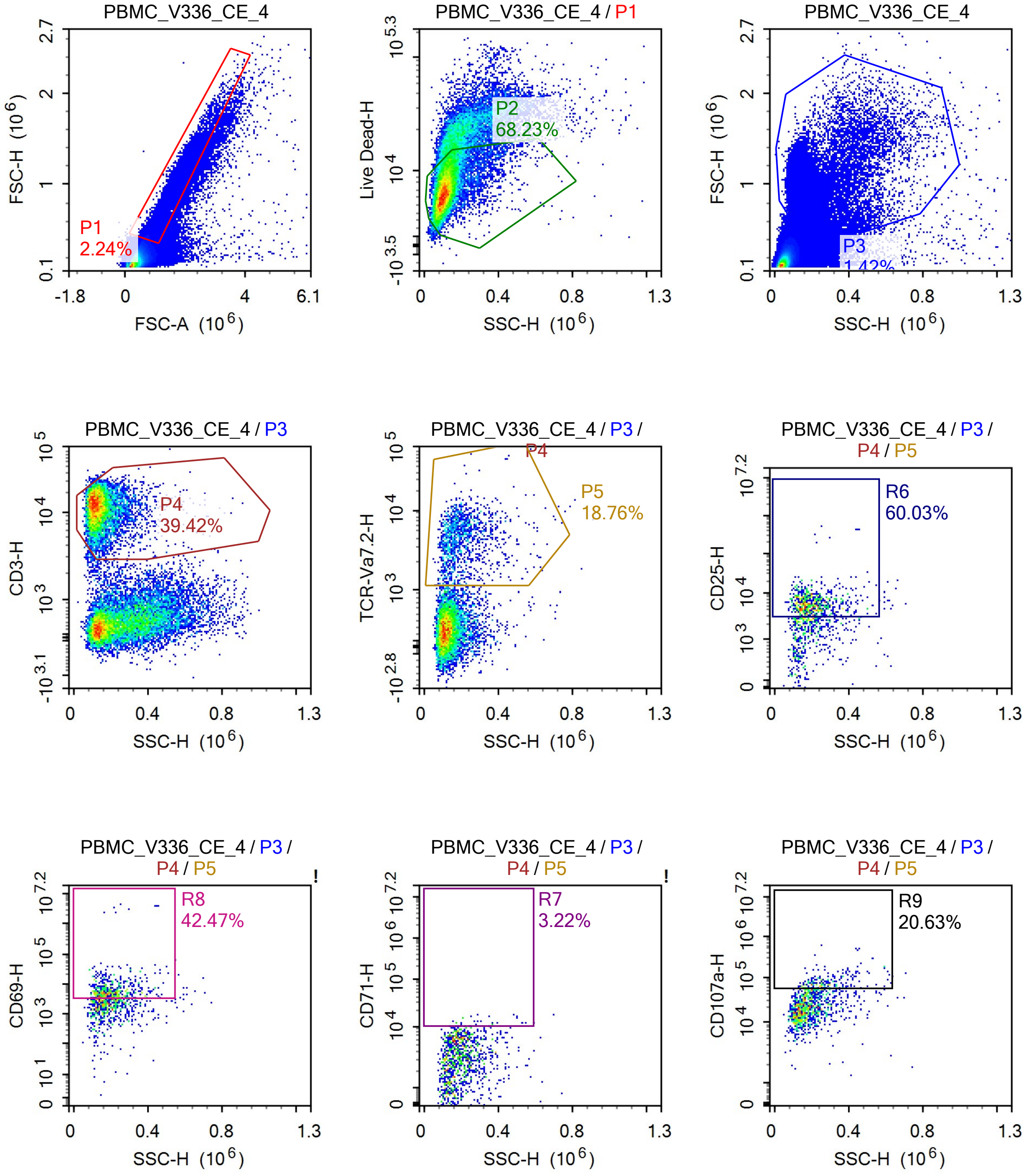

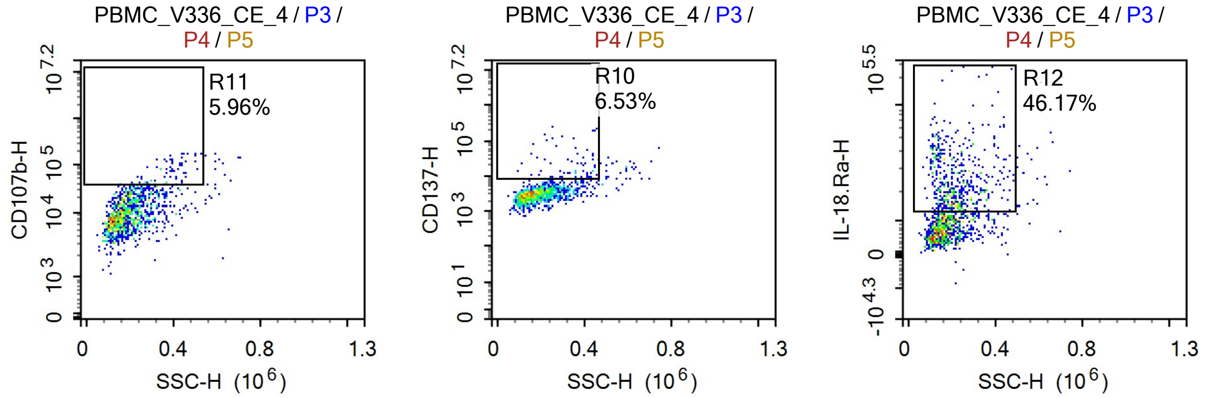


Sample Statistics

| Gate | Count | % Parent | X | Y | Median X | Median Y |
|---|---|---|---|---|---|---|
| All | 1,186,014 | | | | | |
| P1 | 26,509 | 2.24% | FSC-A | FSC-H | 1,120,812 | 623,271 |
| P2 | 18,088 | 68.23% | SSC-H | Live Dead-H | 110,145 | 7,016 |
| P3 | 16,784 | 1.42% | SSC-H | FSC-H | 171,986 | 1,038,096 |
| P4 | 6,616 | 39.42% | SSC-H | CD3-H | 128,488 | 11,986 |
| P5 | 1,241 | 18.76% | SSC-H | TCR-Va7.2-H | 182,442 | 4,146 |
| R6 | 745 | 60.03% | SSC-H | CD25-H | 193,905 | 5,671 |
| R7 | 40 | 3.22% | SSC-H | CD71-H | 250,486 | 11,667 |
| R8 | 527 | 42.47% | SSC-H | CD69-H | 192,873 | 5,700 |
| R9 | 256 | 20.63% | SSC-H | CD107a-H | 257,562 | 72,535 |
| R10 | 81 | 6.53% | SSC-H | CD137-H | 359,320 | 13,203 |
| R11 | 74 | 5.96% | SSC-H | CD107b-H | 374,354 | 67,947 |
| R12 | 573 | 46.17% | SSC-H | IL-18.Ra-H | 208,641 | 21,488 |

### Report of PBMC 12/09/25-PBMC_V339_CE_1

Sample Name: PBMC 12/09/25-PBMC_V339_CE_1 | Run Time: 12-Sep-25 2:06 PM

Cytometer: NovoCyte Quanteon 621181110427 | Software: NovoExpress 1.6.0

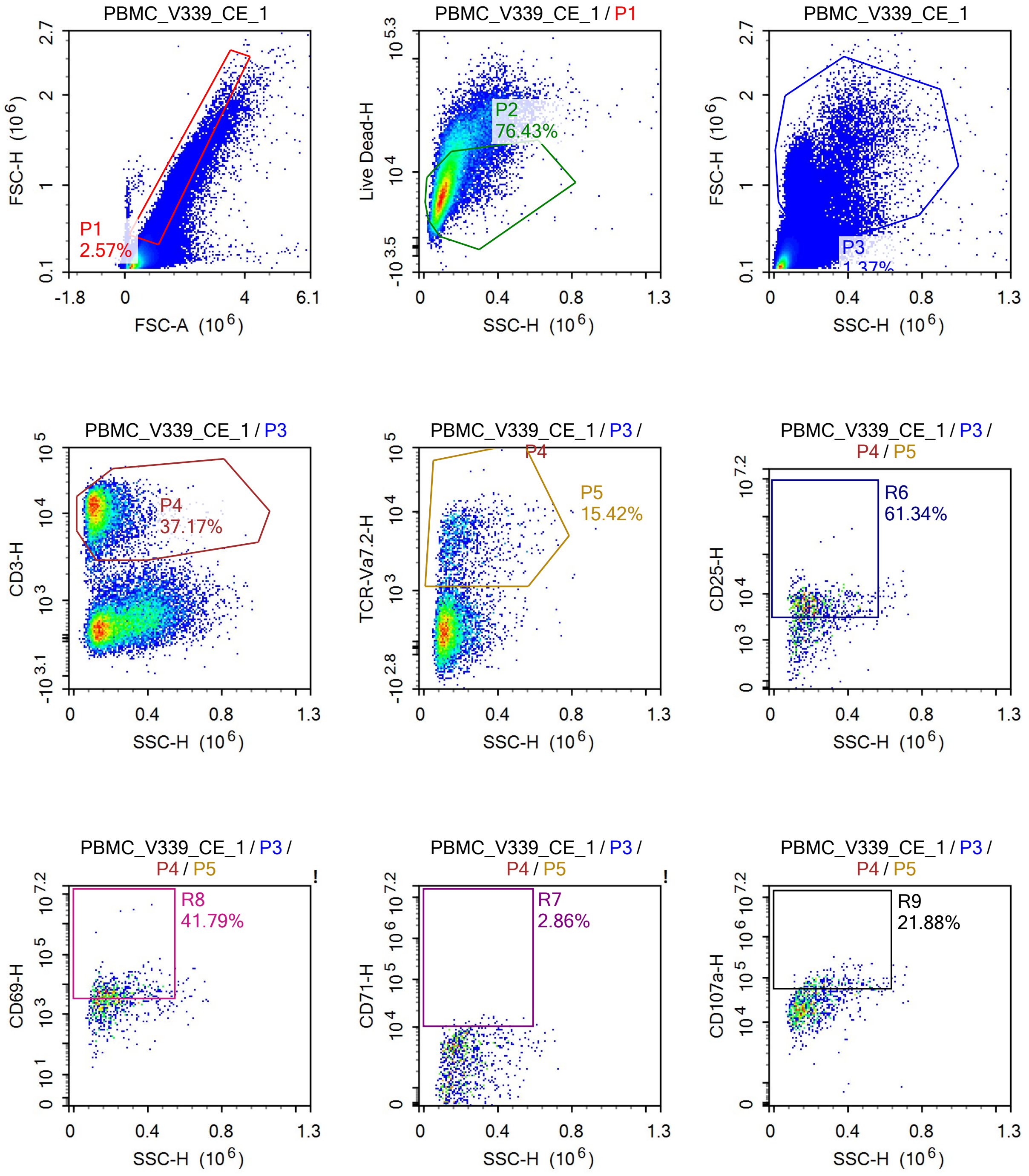

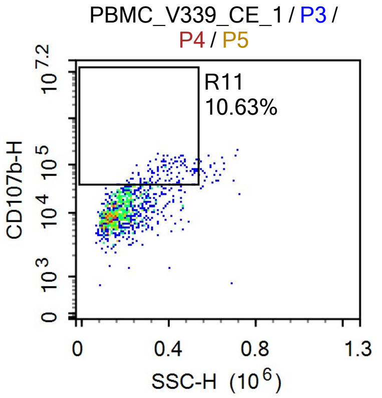


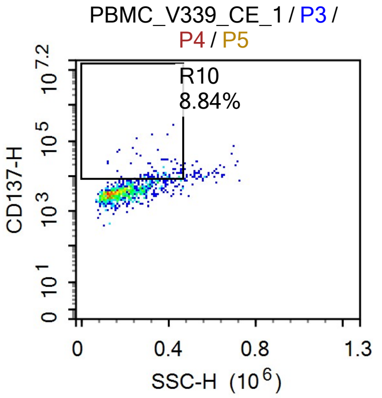


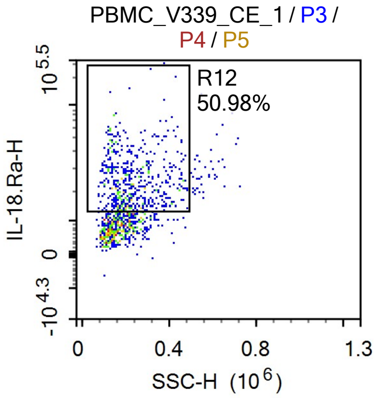


Sample Statistics

| Gate | Count | % Parent | X | Y | Median X | Median Y |
|---|---|---|---|---|---|---|
| All | 1,425,630 | | | | | |
| P1 | 36,592 | 2.57% | FSC-A | FSC-H | 1,034,664 | 522,742 |
| P2 | 27,966 | 76.43% | SSC-H | Live Dead-H | 106,342 | 7,170 |
| P3 | 19,545 | 1.37% | SSC-H | FSC-H | 172,957 | 1,020,236 |
| P4 | 7,265 | 37.17% | SSC-H | CD3-H | 127,719 | 11,323 |
| P5 | 1,120 | 15.42% | SSC-H | TCR-Va7.2-H | 187,262 | 3,839 |
| R6 | 687 | 61.34% | SSC-H | CD25-H | 201,120 | 5,775 |
| R7 | 32 | 2.86% | SSC-H | CD71-H | 302,157 | 11,166 |
| R8 | 468 | 41.79% | SSC-H | CD69-H | 208,784 | 6,005 |
| R9 | 245 | 21.88% | SSC-H | CD107a-H | 269,150 | 73,562 |
| R10 | 99 | 8.84% | SSC-H | CD137-H | 370,361 | 11,340 |
| R11 | 119 | 10.63% | SSC-H | CD107b-H | 376,734 | 64,374 |
| R12 | 571 | 50.98% | SSC-H | IL-18.Ra-H | 210,324 | 22,598 |

### Report of PBMC 12/09/25-PBMC_V339_CE_2

Sample Name: PBMC 12/09/25-PBMC_V339_CE_2 Run Time: 12-Sep-25 2:08 PM

Cytometer: NovoCyte Quanteon 621181110427 Software: NovoExpress 1.6.0

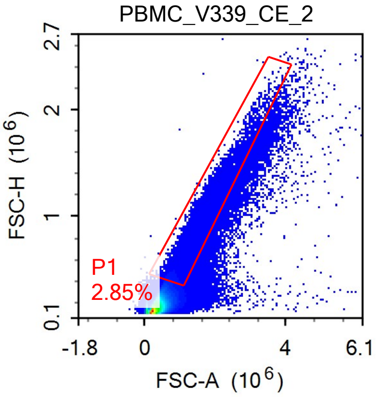


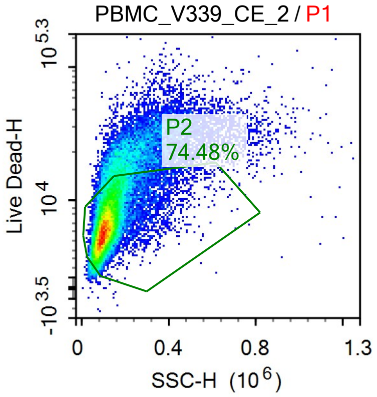


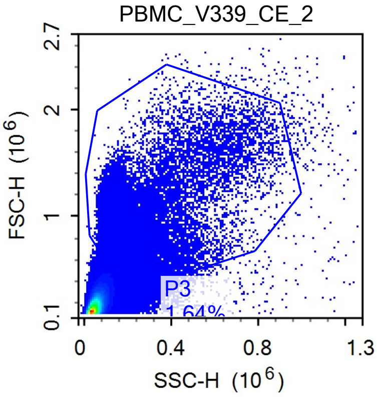


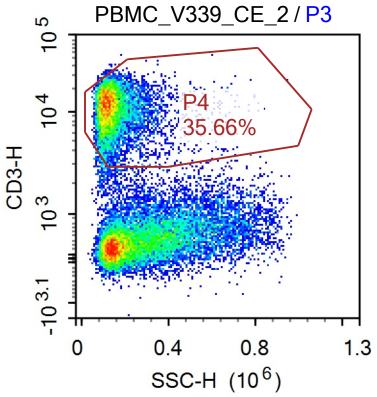


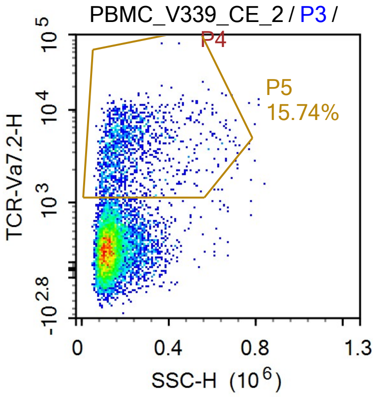


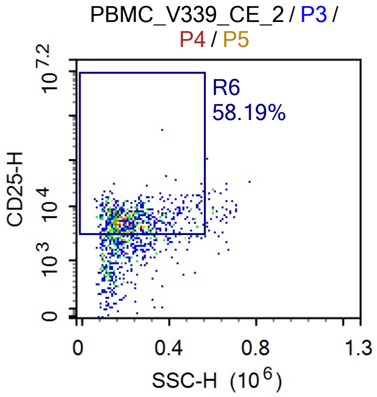


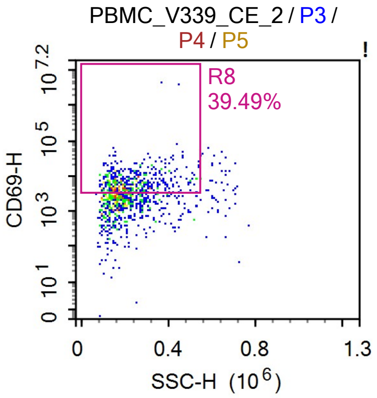


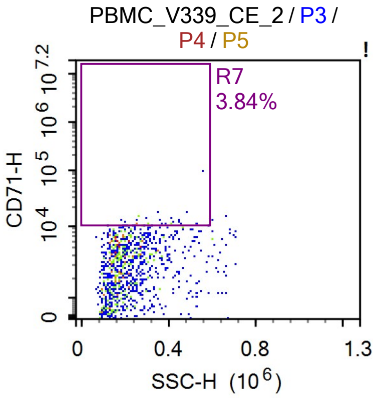


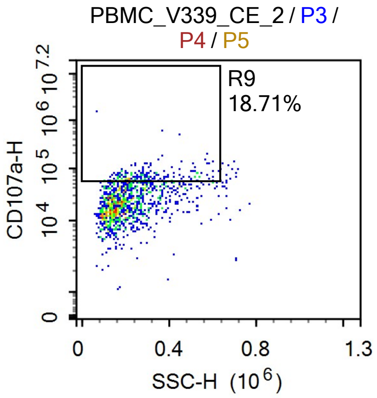

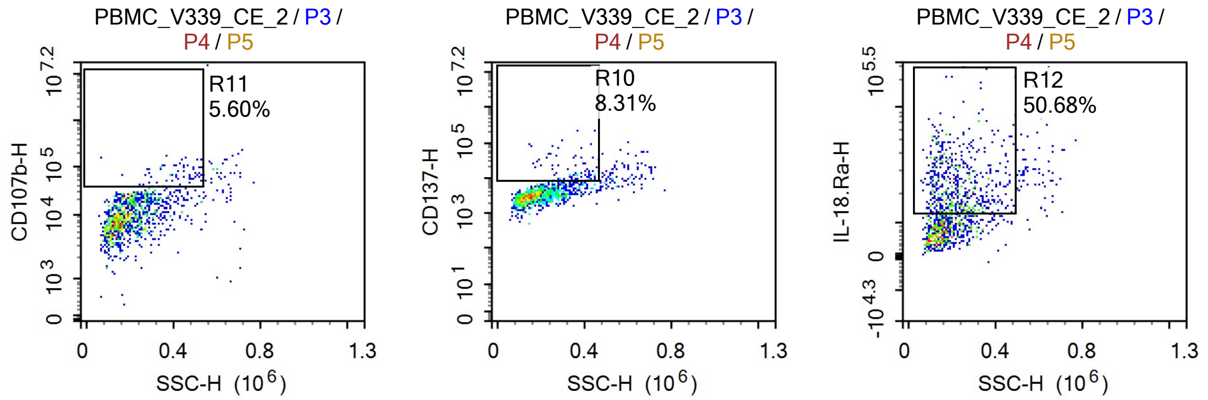


Sample Statistics

| Gate | Count | % Parent | X | Y | Median X | Median Y |
|---|---|---|---|---|---|---|
| All | 1,361,102 | | | | | |
| P1 | 38,821 | 2.85% | FSC-A | FSC-H | 1,061,517 | 548,158 |
| P2 | 28,913 | 74.48% | SSC-H | Live Dead-H | 110,068 | 6,677 |
| P3 | 22,291 | 1.64% | SSC-H | FSC-H | 181,315 | 1,031,510 |
| P4 | 7,950 | 35.66% | SSC-H | CD3-H | 130,605 | 11,227 |
| P5 | 1,251 | 15.74% | SSC-H | TCR-Va7.2-H | 187,455 | 3,649 |
| R6 | 728 | 58.19% | SSC-H | CD25-H | 202,311 | 5,697 |
| R7 | 48 | 3.84% | SSC-H | CD71-H | 301,102 | 11,314 |
| R8 | 494 | 39.49% | SSC-H | CD69-H | 209,662 | 6,711 |
| R9 | 234 | 18.71% | SSC-H | CD107a-H | 271,035 | 66,714 |
| R10 | 104 | 8.31% | SSC-H | CD137-H | 356,813 | 12,413 |
| R11 | 70 | 5.60% | SSC-H | CD107b-H | 404,531 | 65,519 |
| R12 | 634 | 50.68% | SSC-H | IL-18.Ra-H | 207,801 | 22,356 |

### Report of PBMC 12/09/25-PBMC_V339_CE_3

Sample Name: PBMC 12/09/25-PBMC_V339_CE_3 | Run Time: 12-Sep-25 2:10 PM
Cytometer: NovoCyte Quanteon 621181110427 | Software: NovoExpress 1.6.0

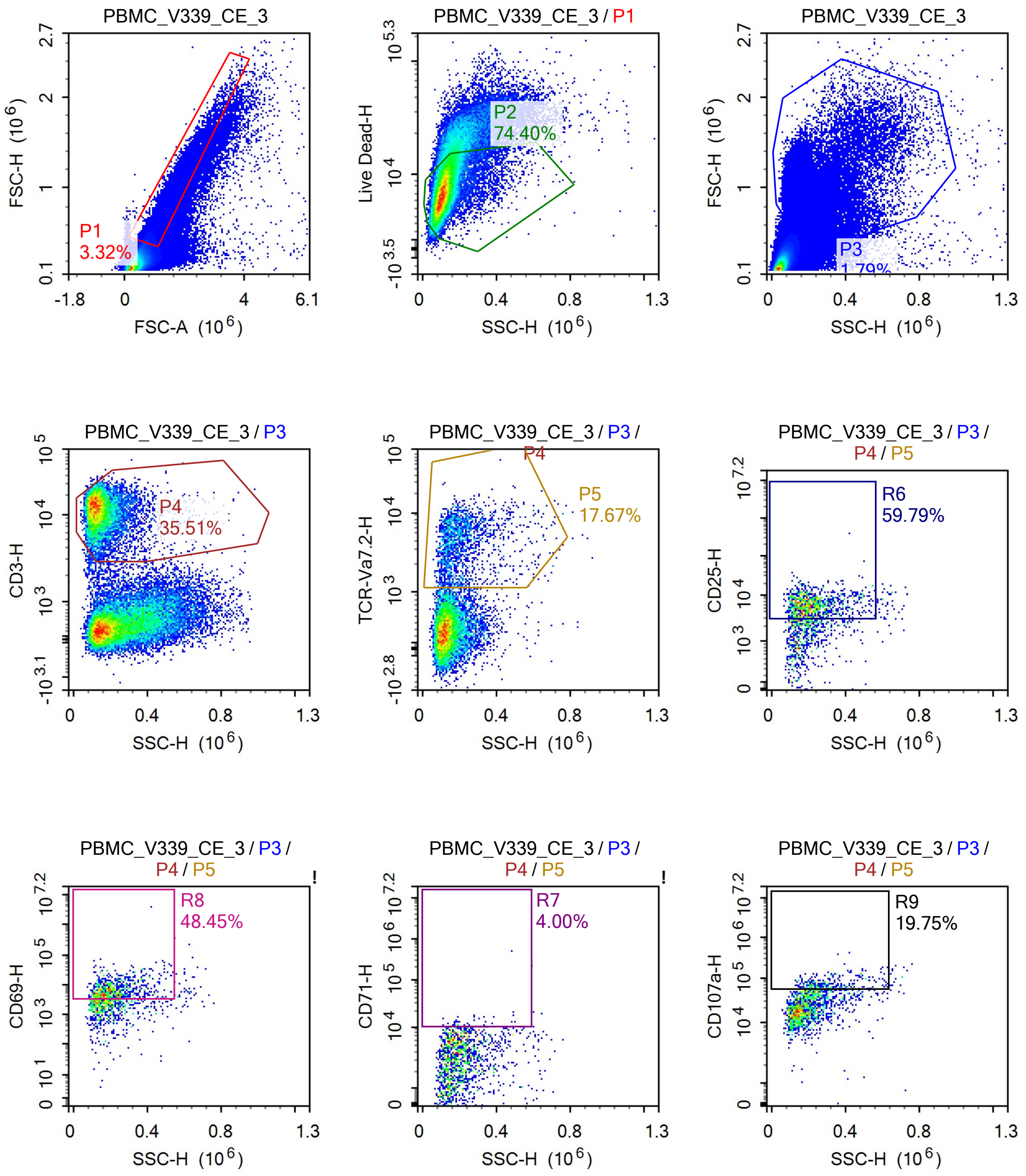

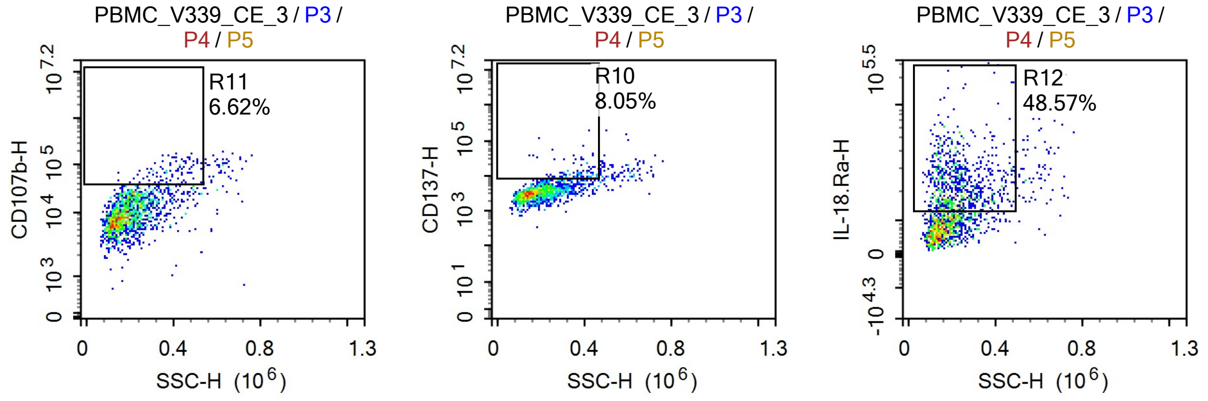


Sample Statistics

| Gate | Count | % Parent | X | Y | Median X | Median Y |
|---|---|---|---|---|---|---|
| All | 1,491,717 | | | | | |
| P1 | 49,474 | 3.32% | FSC-A | FSC-H | 1,035,408 | 521,816 |
| P2 | 36,810 | 74.40% | SSC-H | Live Dead-H | 112,026 | 7,273 |
| P3 | 26,707 | 1.79% | SSC-H | FSC-H | 181,560 | 1,002,977 |
| P4 | 9,484 | 35.51% | SSC-H | CD3-H | 132,681 | 11,864 |
| P5 | 1,676 | 17.67% | SSC-H | TCR-Va7.2-H | 189,727 | 4,186 |
| R6 | 1,002 | 59.79% | SSC-H | CD25-H | 202,074 | 5,978 |
| R7 | 67 | 4.00% | SSC-H | CD71-H | 245,056 | 11,029 |
| R8 | 812 | 48.45% | SSC-H | CD69-H | 201,047 | 6,311 |
| R9 | 331 | 19.75% | SSC-H | CD107a-H | 281,311 | 67,815 |
| R10 | 135 | 8.05% | SSC-H | CD137-H | 363,221 | 10,408 |
| R11 | 111 | 6.62% | SSC-H | CD107b-H | 387,247 | 68,304 |
| R12 | 814 | 48.57% | SSC-H | IL-18.Ra-H | 216,513 | 21,677 |

### Report of PBMC 12/09/25-PBMC_V339_CE_4

Sample Name: PBMC 12/09/25-PBMC_V339_CE_4

Run Time: 12-Sep-25 2:12 PM

Cytometer: NovoCyte Quanteon 621181110427

Software: NovoExpress 1.6.0

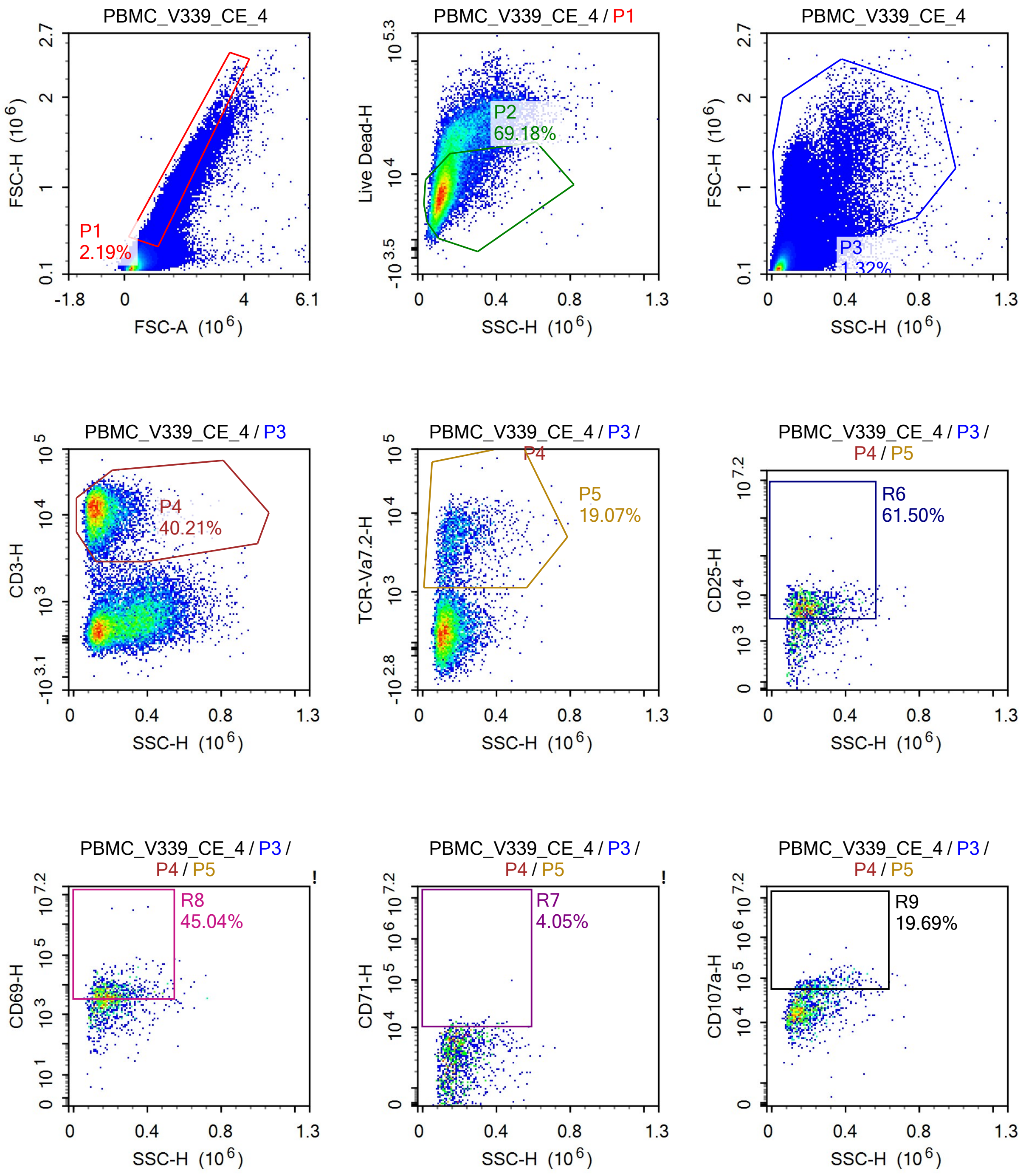

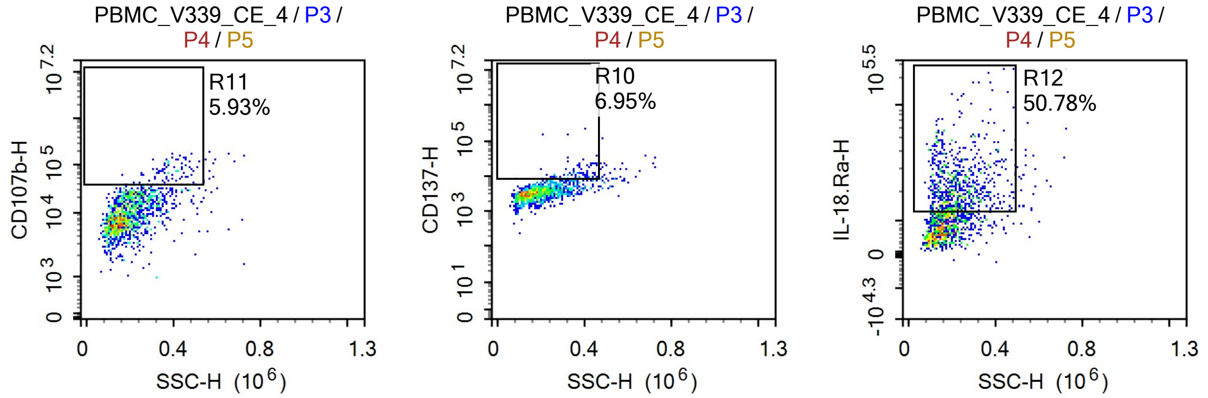


Sample Statistics

| Gate | Count | % Parent | X | Y | Median X | Median Y |
|---|---|---|---|---|---|---|
| All | 1,462,398 | | | | | |
| P1 | 31,982 | 2.19% | FSC-A | FSC-H | 1,095,453 | 582,087 |
| P2 | 22,124 | 69.18% | SSC-H | Live Dead-H | 107,214 | 7,384 |
| P3 | 19,340 | 1.32% | SSC-H | FSC-H | 174,478 | 1,025,526 |
| P4 | 7,776 | 40.21% | SSC-H | CD3-H | 131,179 | 11,414 |
| P5 | 1,483 | 19.07% | SSC-H | TCR-Va7.2-H | 179,350 | 4,133 |
| R6 | 912 | 61.50% | SSC-H | CD25-H | 196,878 | 5,723 |
| R7 | 60 | 4.05% | SSC-H | CD71-H | 226,440 | 11,378 |
| R8 | 668 | 45.04% | SSC-H | CD69-H | 192,724 | 5,868 |
| R9 | 292 | 19.69% | SSC-H | CD107a-H | 267,325 | 70,851 |
| R10 | 103 | 6.95% | SSC-H | CD137-H | 354,039 | 10,530 |
| R11 | 88 | 5.93% | SSC-H | CD107b-H | 377,905 | 69,561 |
| R12 | 753 | 50.78% | SSC-H | IL-18.Ra-H | 213,981 | 20,805 |
