## Supplementary material for "Vitamin B_2_ Production by Vaginal Lactobacilli Promotes Symbiosis": Data S10: 250919 Bacteria + CE.pdf

### Report of PBMC 19_09_25

Specimen Name: PBMC 19_09_25 | Run Time: 19-Sep-25 8:26 AM

Cytometer: NovoCyte Quanteon 621181110427 | Software: NovoExpress 1.6.0

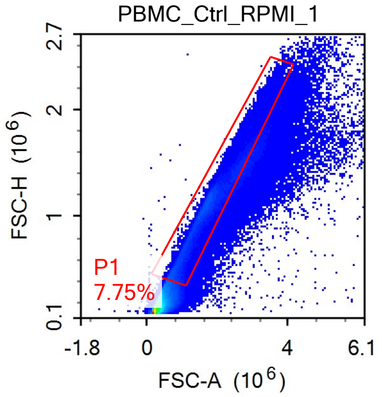


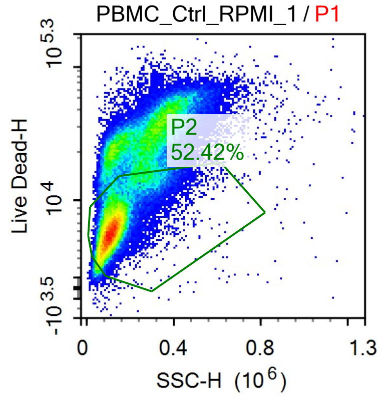


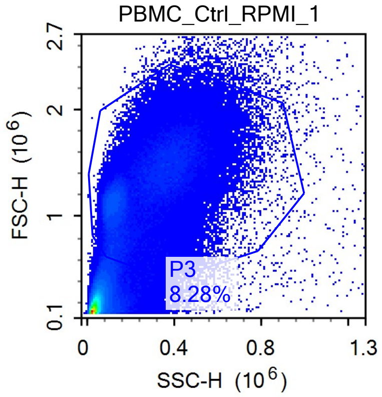


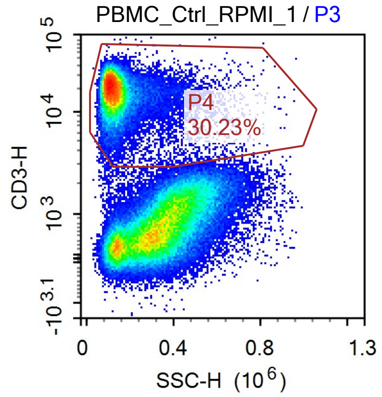


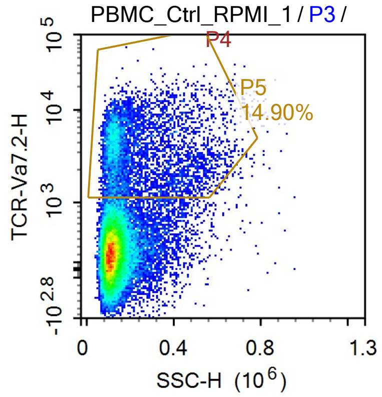


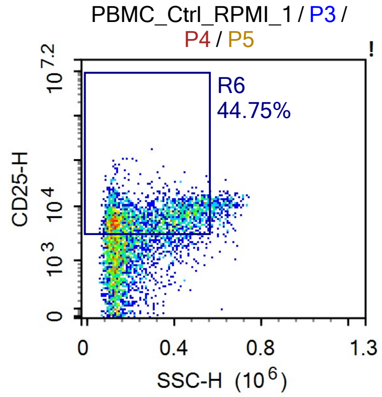


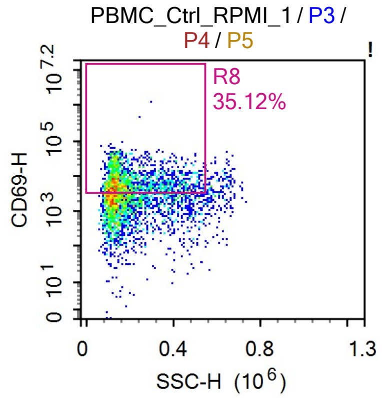


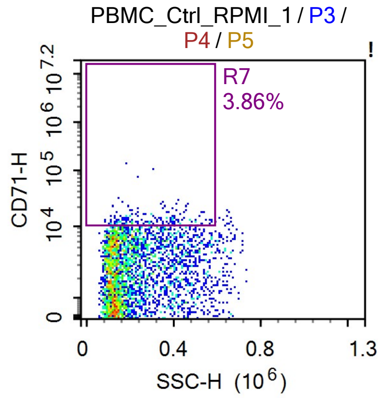


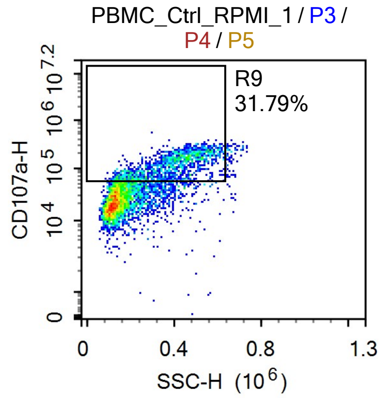

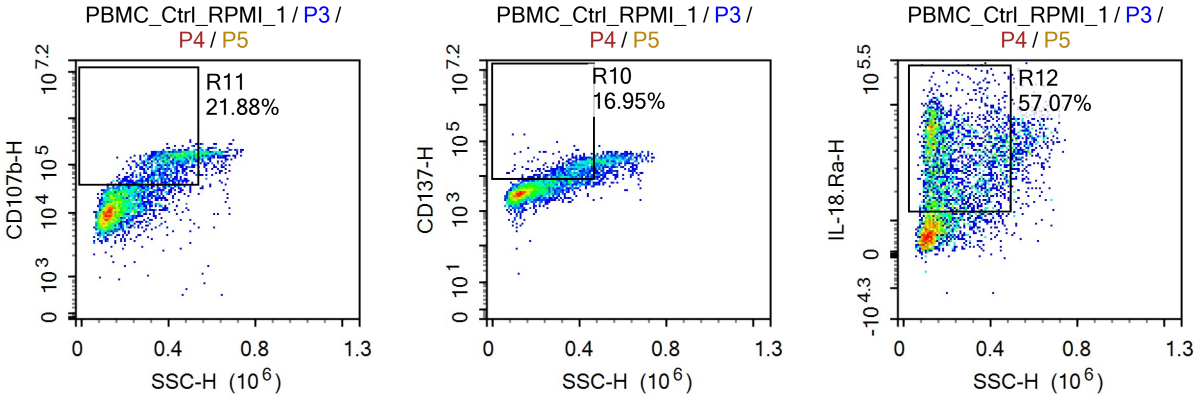


Sample Statistics of PBMC_Ctrl_RPMI_1

| Gate | Count | % Parent | X | Y | Median X | Median Y |
|---|---|---|---|---|---|---|
| All | 1,403,266 | | | | | |
| P1 | 108,728 | 7.75% | FSC-A | FSC-H | 1,650,278 | 1,069,087 |
| P2 | 56,991 | 52.42% | SSC-H | Live Dead-H | 117,528 | 6,579 |
| P3 | 116,186 | 8.28% | SSC-H | FSC-H | 273,309 | 1,222,024 |
| P4 | 35,118 | 30.23% | SSC-H | CD3-H | 121,839 | 18,197 |
| P5 | 5,234 | 14.90% | SSC-H | TCR-Va7.2-H | 170,416 | 3,299 |
| R6 | 2,342 | 44.75% | SSC-H | CD25-H | 246,614 | 6,122 |
| R7 | 202 | 3.86% | SSC-H | CD71-H | 270,516 | 12,195 |
| R8 | 1,838 | 35.12% | SSC-H | CD69-H | 163,081 | 6,881 |
| R9 | 1,664 | 31.79% | SSC-H | CD107a-H | 387,827 | 114,662 |
| R10 | 887 | 16.95% | SSC-H | CD137-H | 370,943 | 14,003 |
| R11 | 1,145 | 21.88% | SSC-H | CD107b-H | 402,053 | 117,854 |
| R12 | 2,987 | 57.07% | SSC-H | IL-18.Ra-H | 189,316 | 35,693 |

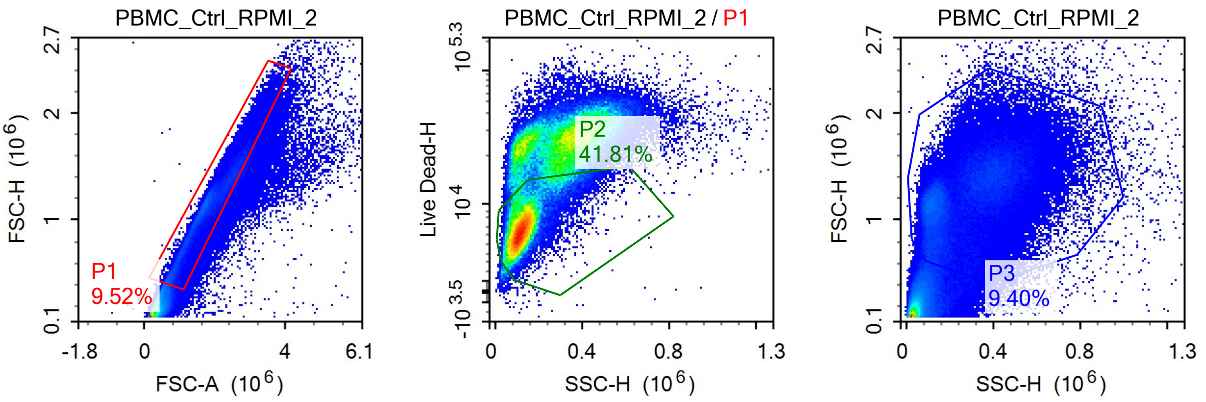

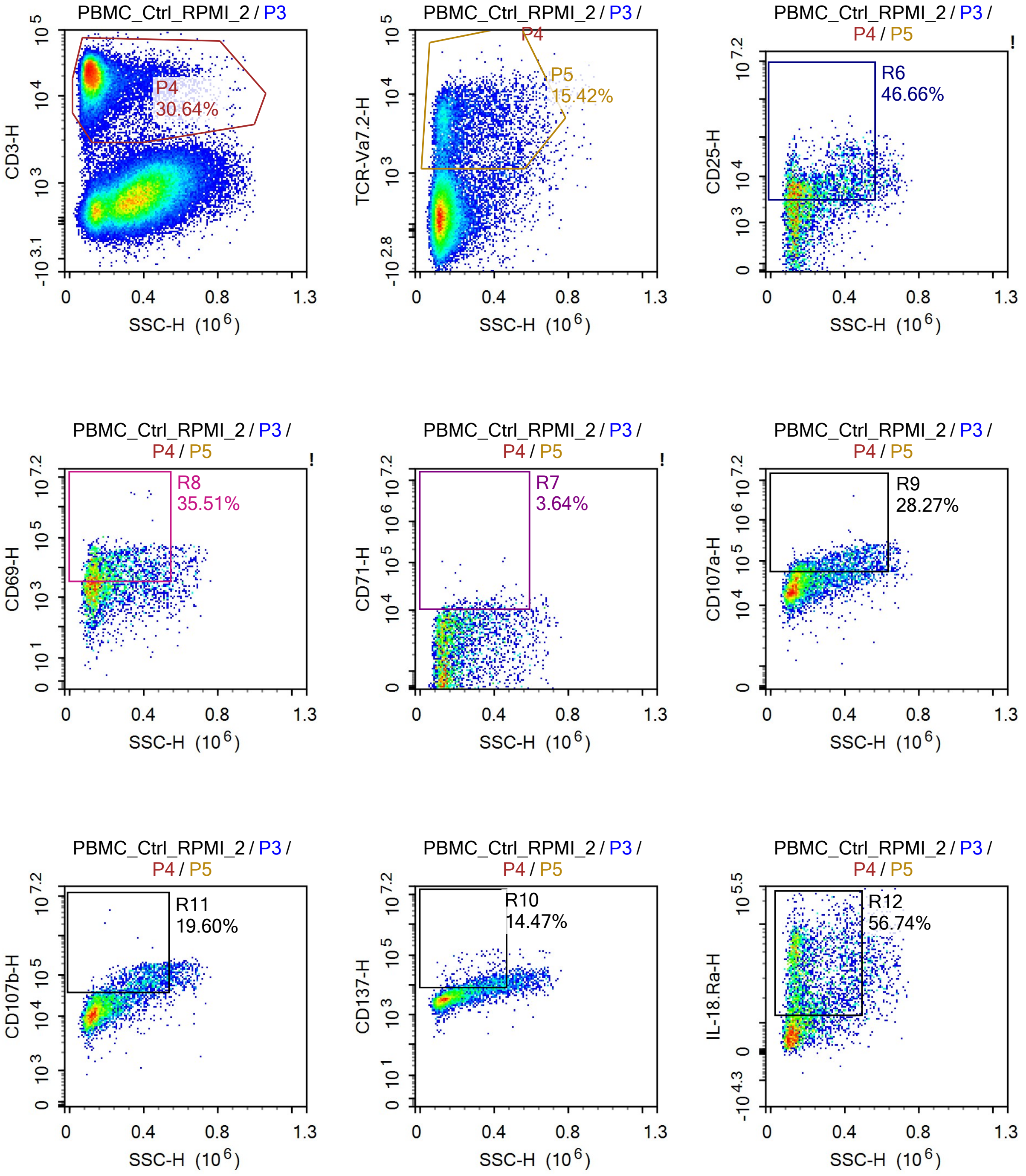
PBMC_Ctrl_RPMI_2 / P3
P4
30.64%
CD3-H
SSC-H (10^6)
PBMC_Ctrl_RPMI_2 / P3 / P4
P5
15.42%
TCR-Va7.2-H
SSC-H (10^6)
PBMC_Ctrl_RPMI_2 / P3 / P4 / P5
R6
46.66%
CD25-H
SSC-H (10^6)
PBMC_Ctrl_RPMI_2 / P3 / P4 / P5
R8
35.51%
CD69-H
SSC-H (10^6)
PBMC_Ctrl_RPMI_2 / P3 / P4 / P5
R7
3.64%
CD71-H
SSC-H (10^6)
PBMC_Ctrl_RPMI_2 / P3 / P4 / P5
R9
28.27%
CD107a-H
SSC-H (10^6)
PBMC_Ctrl_RPMI_2 / P3 / P4 / P5
R11
19.60%
CD107b-H
SSC-H (10^6)
PBMC_Ctrl_RPMI_2 / P3 / P4 / P5
R10
14.47%
CD137-H
SSC-H (10^6)
PBMC_Ctrl_RPMI_2 / P3 / P4 / P5
R12
56.74%
IL-18.Ra-H
SSC-H (10^6)

Sample Statistics of PBMC_Ctrl_RPMI_2

| Gate | Count | % Parent | X | Y | Median X | Median Y |
|---|---|---|---|---|---|---|
| All | 1,106,340 | | | | | |
| P1 | 105,377 | 9.52% | FSC-A | FSC-H | 1,727,531 | 1,119,092 |
| P2 | 44,059 | 41.81% | SSC-H | Live Dead-H | 121,134 | 6,949 |
| P3 | 103,965 | 9.40% | SSC-H | FSC-H | 281,215 | 1,202,179 |
| P4 | 31,859 | 30.64% | SSC-H | CD3-H | 122,127 | 19,313 |
| P5 | 4,914 | 15.42% | SSC-H | TCR-Va7.2-H | 169,050 | 3,633 |
| R6 | 2,293 | 46.66% | SSC-H | CD25-H | 230,180 | 6,204 |
| R7 | 179 | 3.64% | SSC-H | CD71-H | 315,046 | 12,187 |
| R8 | 1,745 | 35.51% | SSC-H | CD69-H | 188,464 | 8,794 |
| R9 | 1,389 | 28.27% | SSC-H | CD107a-H | 334,180 | 78,806 |
| R10 | 711 | 14.47% | SSC-H | CD137-H | 360,529 | 11,305 |
| R11 | 963 | 19.60% | SSC-H | CD107b-H | 401,755 | 71,328 |
| R12 | 2,788 | 56.74% | SSC-H | IL-18.Ra-H | 184,590 | 33,020 |

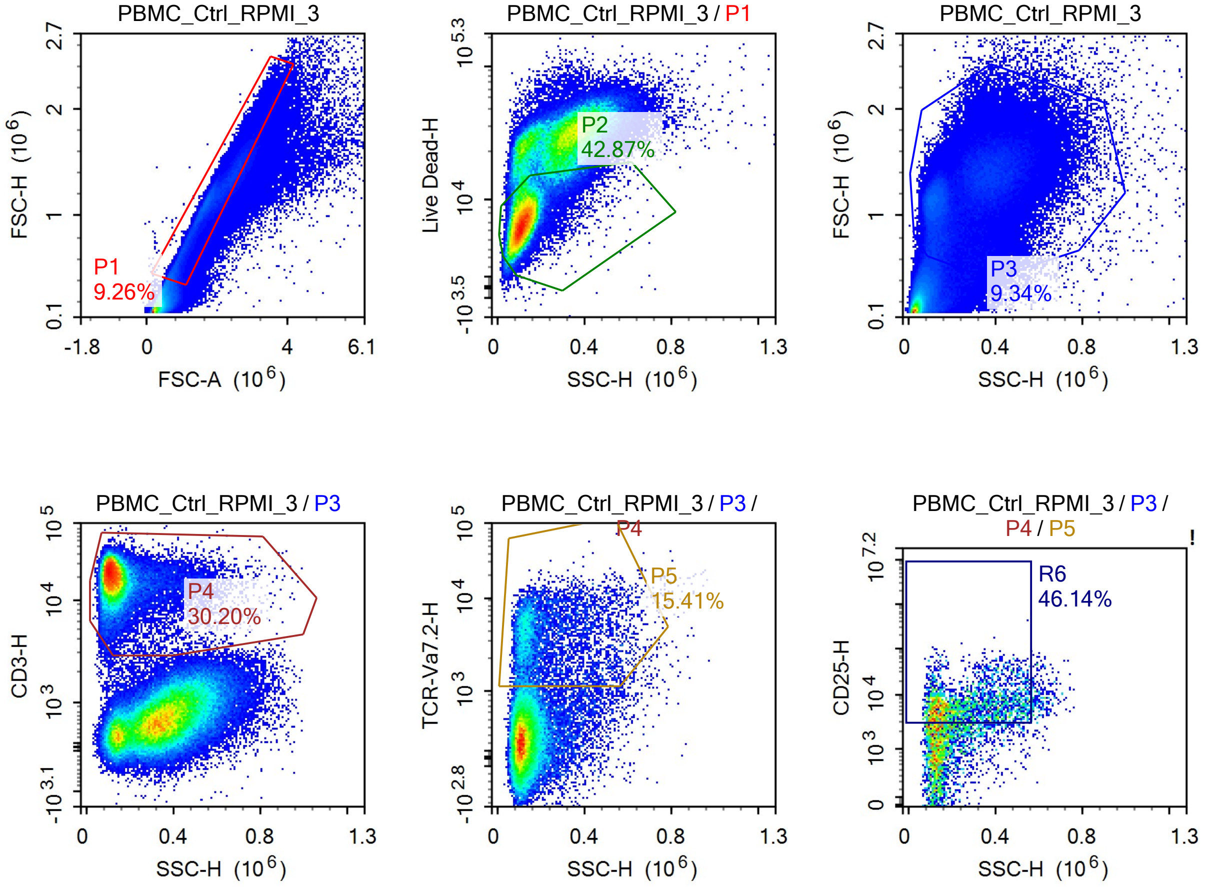

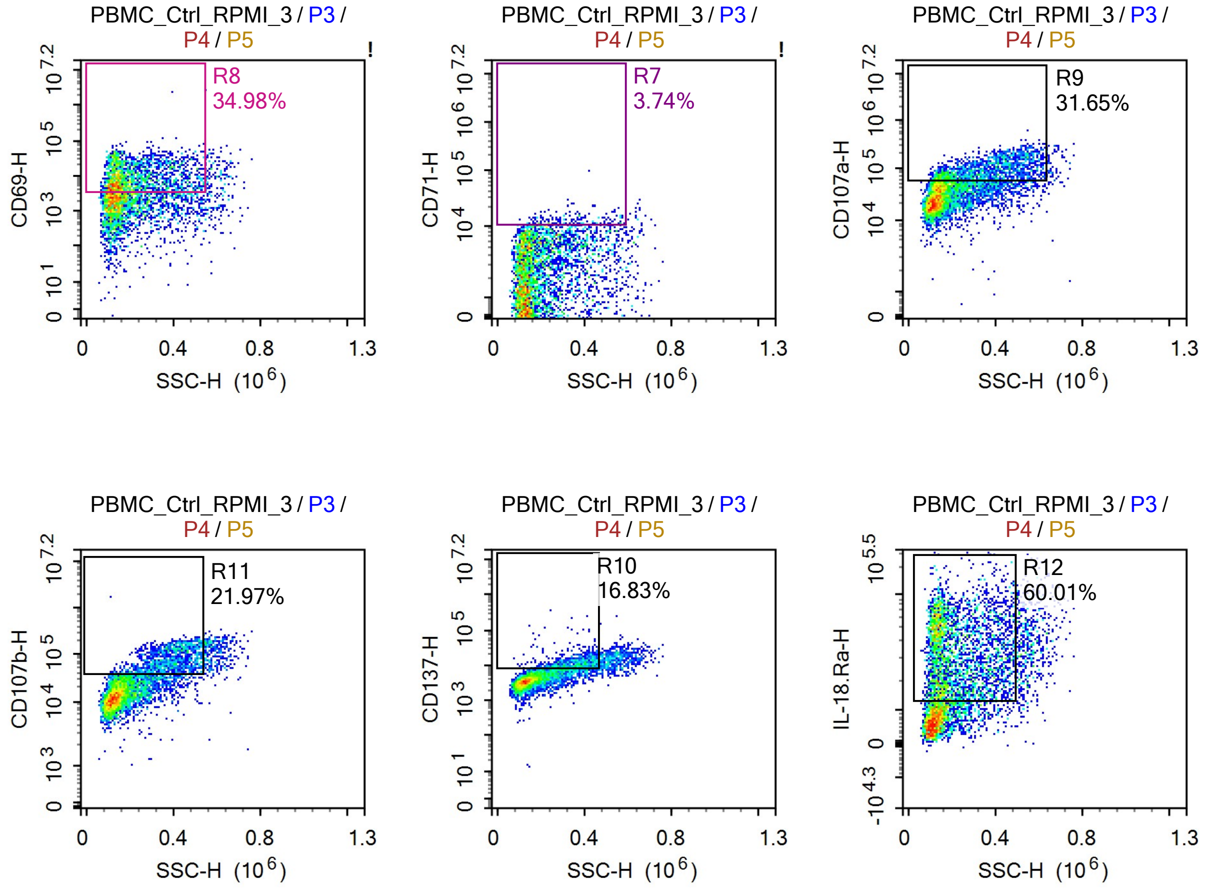


Sample Statistics of PBMC_Ctrl_RPMI_3

| Gate | Count | % Parent | X | Y | Median X | Median Y |
|---|---|---|---|---|---|---|
| All | 1,383,585 | | | | | |
| P1 | 128,187 | 9.26% | FSC-A | FSC-H | 1,720,967 | 1,106,671 |
| P2 | 54,951 | 42.87% | SSC-H | Live Dead-H | 116,786 | 7,290 |
| P3 | 129,163 | 9.34% | SSC-H | FSC-H | 283,085 | 1,211,035 |
| P4 | 39,002 | 30.20% | SSC-H | CD3-H | 121,914 | 19,052 |
| P5 | 6,012 | 15.41% | SSC-H | TCR-Va7.2-H | 169,018 | 3,528 |
| R6 | 2,774 | 46.14% | SSC-H | CD25-H | 251,223 | 6,302 |
| R7 | 225 | 3.74% | SSC-H | CD71-H | 306,514 | 11,554 |
| R8 | 2,103 | 34.98% | SSC-H | CD69-H | 178,615 | 8,431 |
| R9 | 1,903 | 31.65% | SSC-H | CD107a-H | 343,370 | 82,874 |
| R10 | 1,012 | 16.83% | SSC-H | CD137-H | 361,979 | 11,025 |
| R11 | 1,321 | 21.97% | SSC-H | CD107b-H | 388,761 | 72,931 |
| R12 | 3,608 | 60.01% | SSC-H | IL-18.Ra-H | 187,212 | 33,130 |

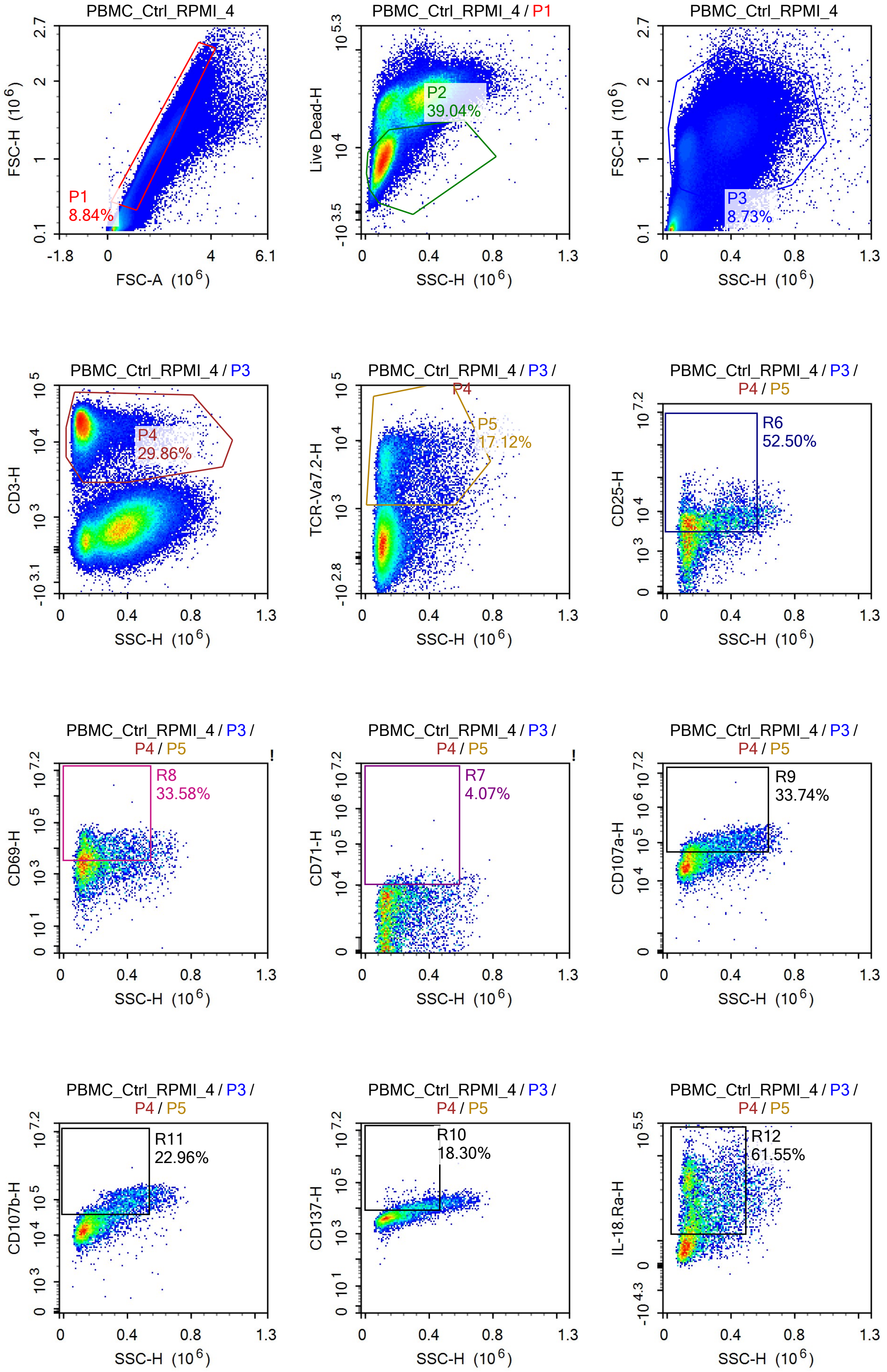
PBMC_Ctrl_RPMI_4
FSC-H (10^6)
FSC-A (10^6)
P1
8.84%
PBMC_Ctrl_RPMI_4 / P1
Live Dead-H
SSC-H (10^6)
P2
39.04%
PBMC_Ctrl_RPMI_4
FSC-H (10^6)
P3
8.73%
PBMC_Ctrl_RPMI_4 / P3
CD3-H
P4
29.86%
PBMC_Ctrl_RPMI_4 / P3 / P4
TCR-Va7.2-H
P5
17.12%
PBMC_Ctrl_RPMI_4 / P3 / P4 / P5
CD25-H
R6
52.50%
CD69-H
R8
33.58%
CD71-H
R7
4.07%
CD107a-H
R9
33.74%
CD107b-H
R11
22.96%
CD137-H
R10
18.30%
IL-18.Ra-H
R12
61.55%

Sample Statistics of PBMC_Ctrl_RPMI_4

| Gate | Count | % Parent | X | Y | Median X | Median Y |
|---|---|---|---|---|---|---|
| All | 1,641,602 | | | | | |
| P1 | 145,186 | 8.84% | FSC-A | FSC-H | 1,726,138 | 1,100,532 |
| P2 | 56,681 | 39.04% | SSC-H | Live Dead-H | 113,747 | 8,068 |
| P3 | 143,332 | 8.73% | SSC-H | FSC-H | 288,508 | 1,195,633 |
| P4 | 42,797 | 29.86% | SSC-H | CD3-H | 122,382 | 18,808 |
| P5 | 7,326 | 17.12% | SSC-H | TCR-Va7.2-H | 167,567 | 3,875 |
| R6 | 3,846 | 52.50% | SSC-H | CD25-H | 204,893 | 5,739 |
| R7 | 298 | 4.07% | SSC-H | CD71-H | 277,147 | 11,752 |
| R8 | 2,460 | 33.58% | SSC-H | CD69-H | 171,464 | 7,482 |
| R9 | 2,472 | 33.74% | SSC-H | CD107a-H | 293,602 | 79,716 |
| R10 | 1,341 | 18.30% | SSC-H | CD137-H | 342,358 | 11,506 |
| R11 | 1,682 | 22.96% | SSC-H | CD107b-H | 360,774 | 69,919 |
| R12 | 4,509 | 61.55% | SSC-H | IL-18.Ra-H | 186,643 | 31,121 |

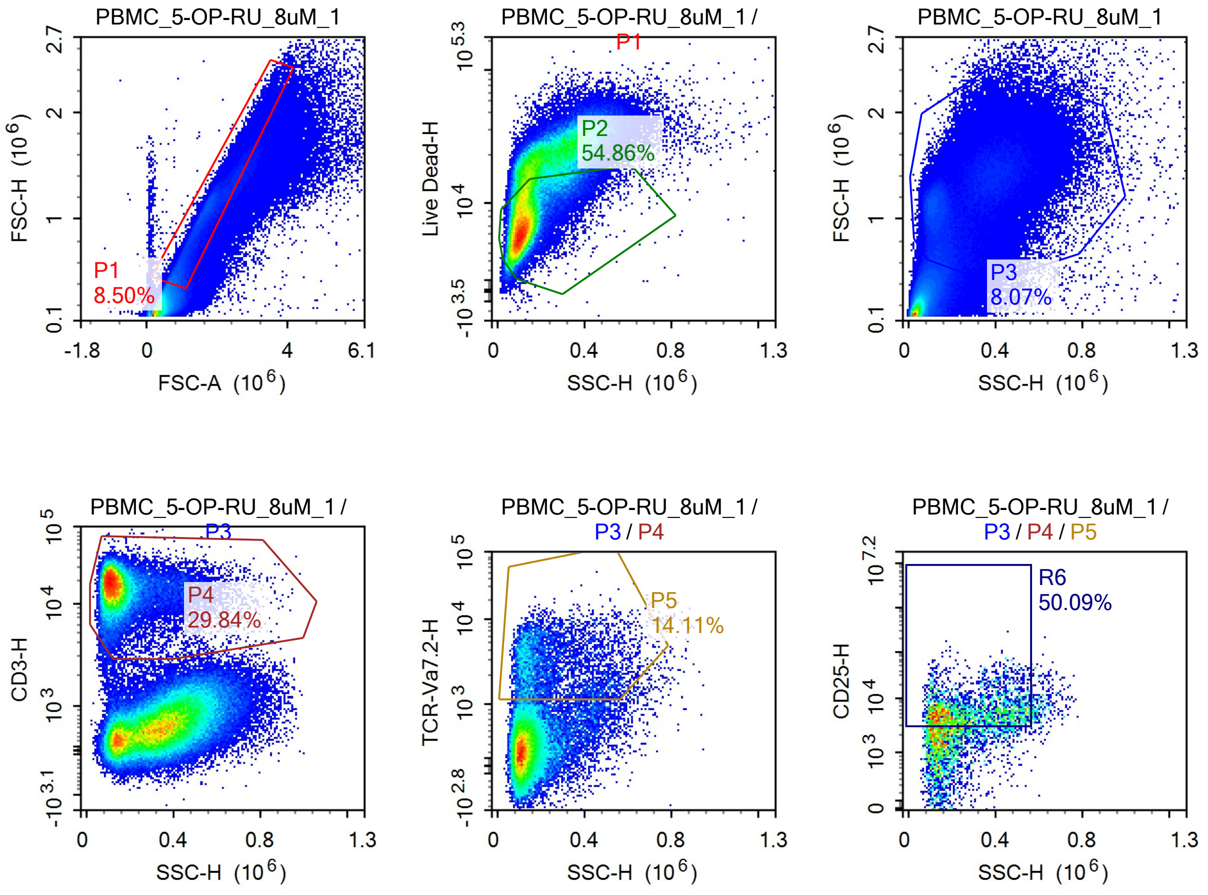

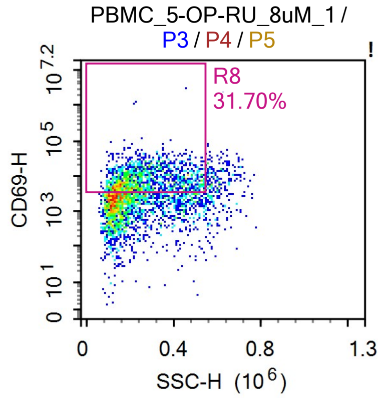


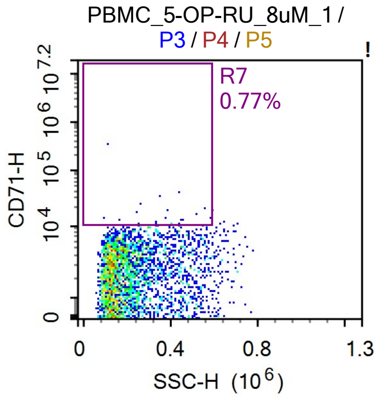


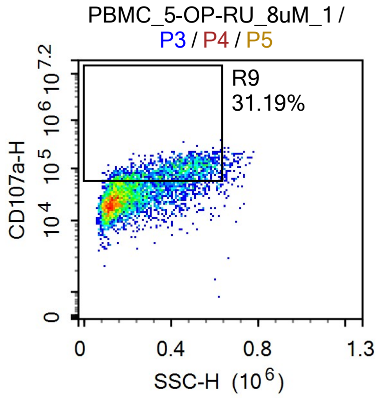


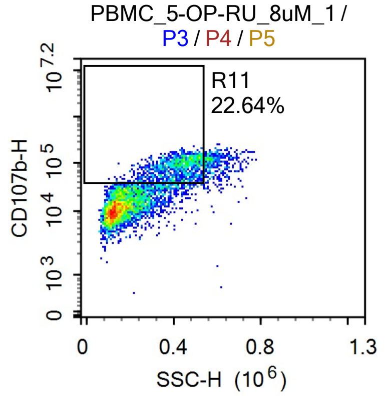


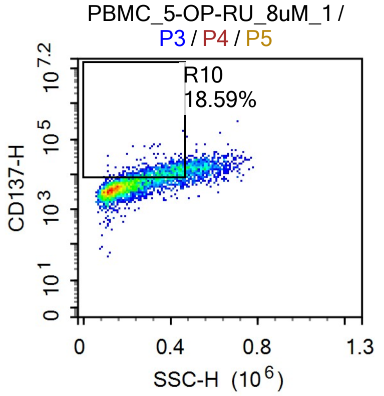


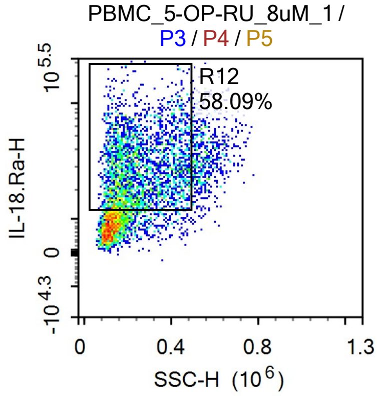


Sample Statistics of PBMC_5-OP-RU_8uM_1

| Gate | Count | % Parent | X | Y | Median X | Median Y |
|---|---|---|---|---|---|---|
| All | 1,601,334 | | | | | |
| P1 | 136,060 | 8.50% | FSC-A | FSC-H | 1,476,308 | 917,339 |
| P2 | 74,648 | 54.86% | SSC-H | Live Dead-H | 115,890 | 7,306 |
| P3 | 129,195 | 8.07% | SSC-H | FSC-H | 265,580 | 1,175,327 |
| P4 | 38,552 | 29.84% | SSC-H | CD3-H | 126,702 | 16,394 |
| P5 | 5,438 | 14.11% | SSC-H | TCR-Va7.2-H | 199,936 | 2,763 |
| R6 | 2,724 | 50.09% | SSC-H | CD25-H | 256,573 | 5,457 |
| R7 | 42 | 0.77% | SSC-H | CD71-H | 344,677 | 10,969 |
| R8 | 1,724 | 31.70% | SSC-H | CD69-H | 237,290 | 7,160 |
| R9 | 1,696 | 31.19% | SSC-H | CD107a-H | 405,039 | 83,700 |
| R10 | 1,011 | 18.59% | SSC-H | CD137-H | 369,921 | 10,969 |
| R11 | 1,231 | 22.64% | SSC-H | CD107b-H | 423,728 | 84,667 |
| R12 | 3,159 | 58.09% | SSC-H | IL-18.Ra-H | 227,711 | 27,733 |

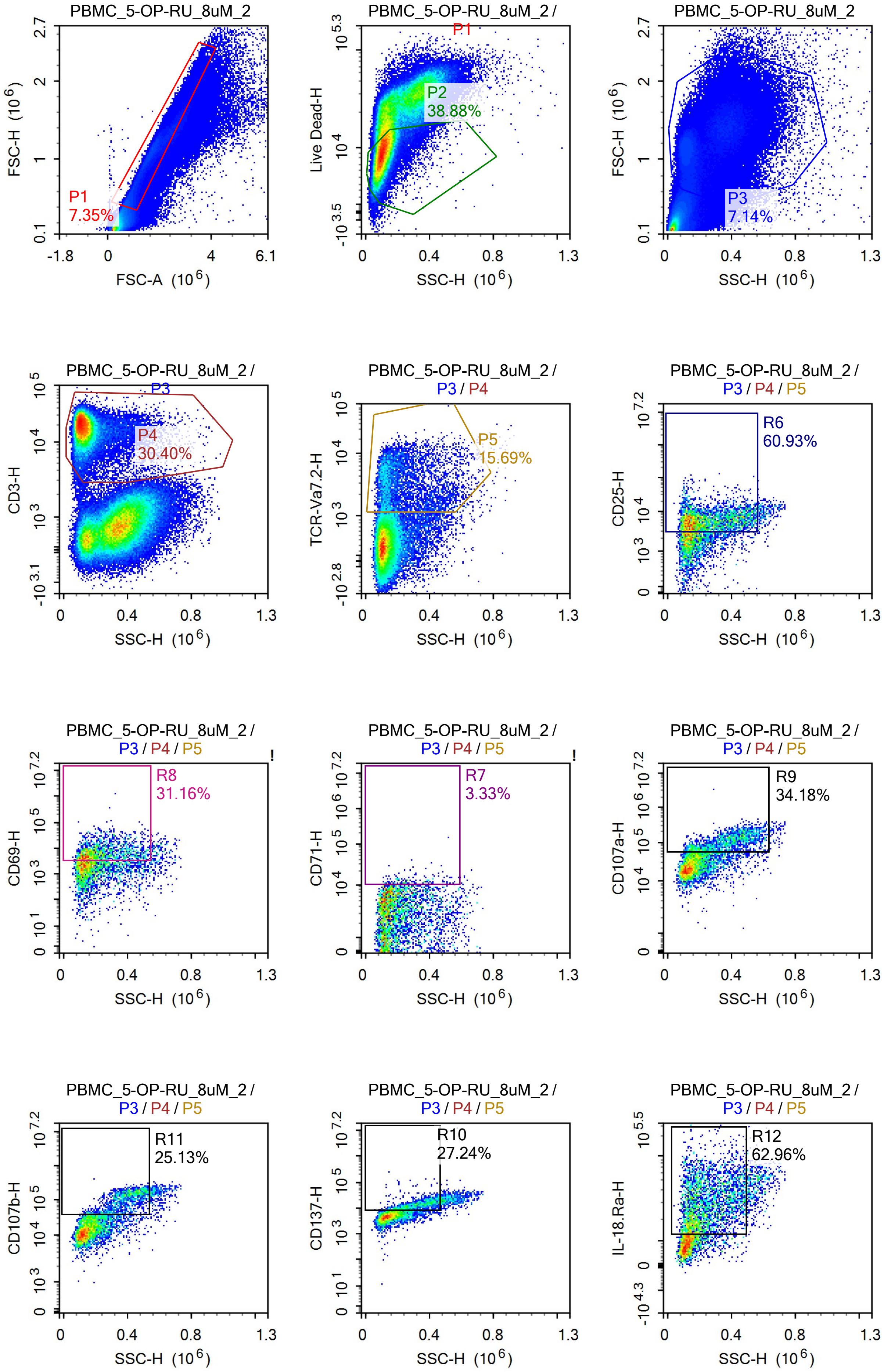
PBMC_5-OP-RU_8uM_2
P1
7.35%
FSC-H (10^6)
FSC-A (10^6)
PBMC_5-OP-RU_8uM_2 / P1
P2
38.88%
Live Dead-H
SSC-H (10^6)
PBMC_5-OP-RU_8uM_2
P3
7.14%
FSC-H (10^6)
SSC-H (10^6)
PBMC_5-OP-RU_8uM_2 / P3
P4
30.40%
CD3-H
SSC-H (10^6)
PBMC_5-OP-RU_8uM_2 / P3 / P4
P5
15.69%
TCR-Va7.2-H
SSC-H (10^6)
PBMC_5-OP-RU_8uM_2 / P3 / P4 / P5
R6
60.93%
CD25-H
SSC-H (10^6)
PBMC_5-OP-RU_8uM_2 / P3 / P4 / P5
R8
31.16%
CD69-H
SSC-H (10^6)
PBMC_5-OP-RU_8uM_2 / P3 / P4 / P5
R7
3.33%
CD71-H
SSC-H (10^6)
PBMC_5-OP-RU_8uM_2 / P3 / P4 / P5
R9
34.18%
CD107a-H
SSC-H (10^6)
PBMC_5-OP-RU_8uM_2 / P3 / P4 / P5
R11
25.13%
CD107b-H
SSC-H (10^6)
PBMC_5-OP-RU_8uM_2 / P3 / P4 / P5
R10
27.24%
CD137-H
SSC-H (10^6)
PBMC_5-OP-RU_8uM_2 / P3 / P4 / P5
R12
62.96%
IL-18.Ra-H
SSC-H (10^6)

Sample Statistics of PBMC_5-OP-RU_8uM_2

| Gate | Count | % Parent | X | Y | Median X | Median Y |
|---|---|---|---|---|---|---|
| All | 1,637,588 | | | | | |
| P1 | 120,415 | 7.35% | FSC-A | FSC-H | 1,575,460 | 995,883 |
| P2 | 46,823 | 38.88% | SSC-H | Live Dead-H | 106,628 | 8,694 |
| P3 | 116,951 | 7.14% | SSC-H | FSC-H | 270,501 | 1,200,076 |
| P4 | 35,556 | 30.40% | SSC-H | CD3-H | 123,346 | 17,376 |
| P5 | 5,580 | 15.69% | SSC-H | TCR-Va7.2-H | 188,604 | 3,092 |
| R6 | 3,400 | 60.93% | SSC-H | CD25-H | 224,709 | 5,823 |
| R7 | 186 | 3.33% | SSC-H | CD71-H | 247,035 | 11,688 |
| R8 | 1,739 | 31.16% | SSC-H | CD69-H | 209,370 | 6,385 |
| R9 | 1,907 | 34.18% | SSC-H | CD107a-H | 376,194 | 103,329 |
| R10 | 1,520 | 27.24% | SSC-H | CD137-H | 325,451 | 13,166 |
| R11 | 1,402 | 25.13% | SSC-H | CD107b-H | 389,600 | 114,484 |
| R12 | 3,513 | 62.96% | SSC-H | IL-18.Ra-H | 218,465 | 28,865 |

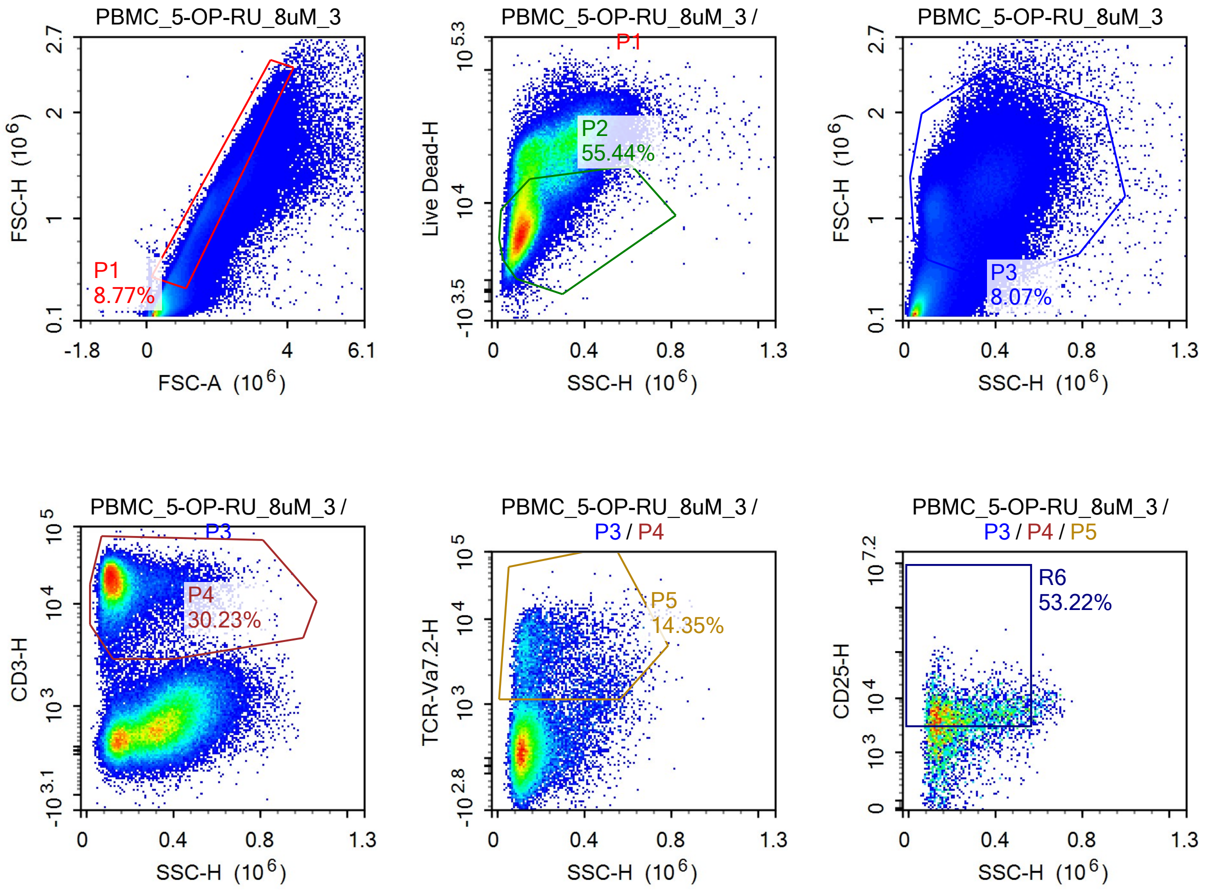

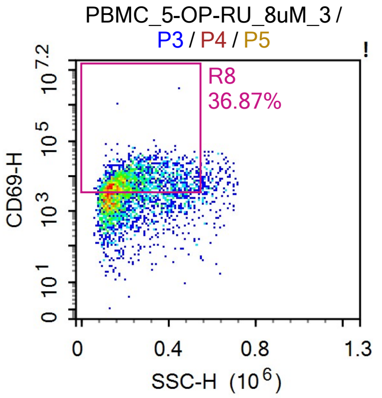


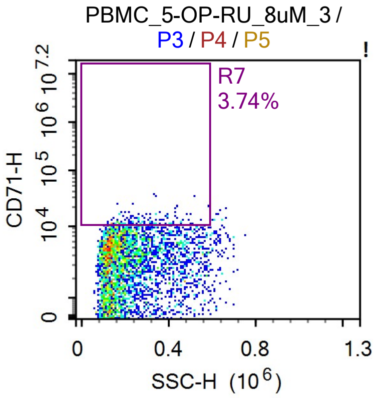


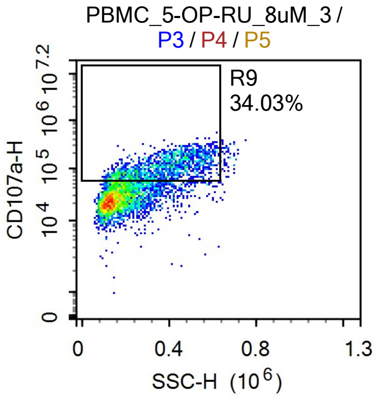


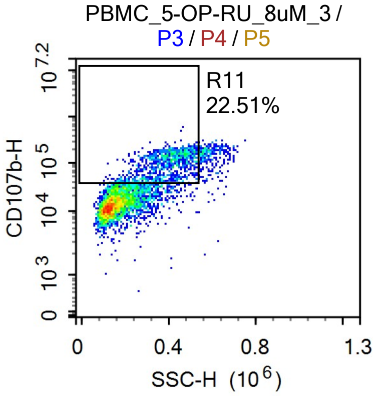


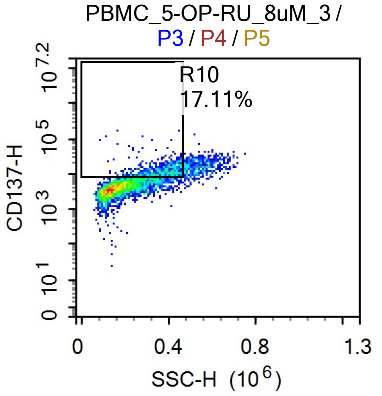


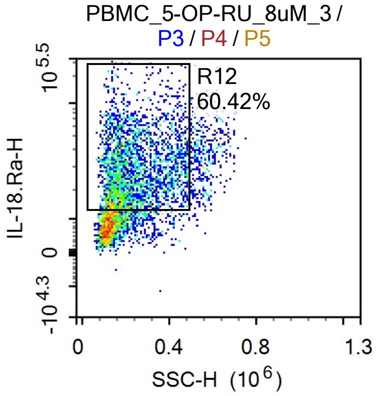


Sample Statistics of PBMC_5-OP-RU_8uM_3

| Gate | Count | % Parent | X | Y | Median X | Median Y |
|---|---|---|---|---|---|---|
| All | 1,428,070 | | | | | |
| P1 | 125,189 | 8.77% | FSC-A | FSC-H | 1,437,780 | 887,048 |
| P2 | 69,409 | 55.44% | SSC-H | Live Dead-H | 116,754 | 7,171 |
| P3 | 115,245 | 8.07% | SSC-H | FSC-H | 252,738 | 1,126,457 |
| P4 | 34,837 | 30.23% | SSC-H | CD3-H | 127,261 | 18,288 |
| P5 | 4,998 | 14.35% | SSC-H | TCR-Va7.2-H | 188,010 | 3,220 |
| R6 | 2,660 | 53.22% | SSC-H | CD25-H | 214,193 | 5,561 |
| R7 | 187 | 3.74% | SSC-H | CD71-H | 267,945 | 11,423 |
| R8 | 1,843 | 36.87% | SSC-H | CD69-H | 216,725 | 7,120 |
| R9 | 1,701 | 34.03% | SSC-H | CD107a-H | 362,841 | 98,408 |
| R10 | 855 | 17.11% | SSC-H | CD137-H | 365,692 | 11,248 |
| R11 | 1,125 | 22.51% | SSC-H | CD107b-H | 395,319 | 106,789 |
| R12 | 3,020 | 60.42% | SSC-H | IL-18.Ra-H | 214,710 | 27,079 |

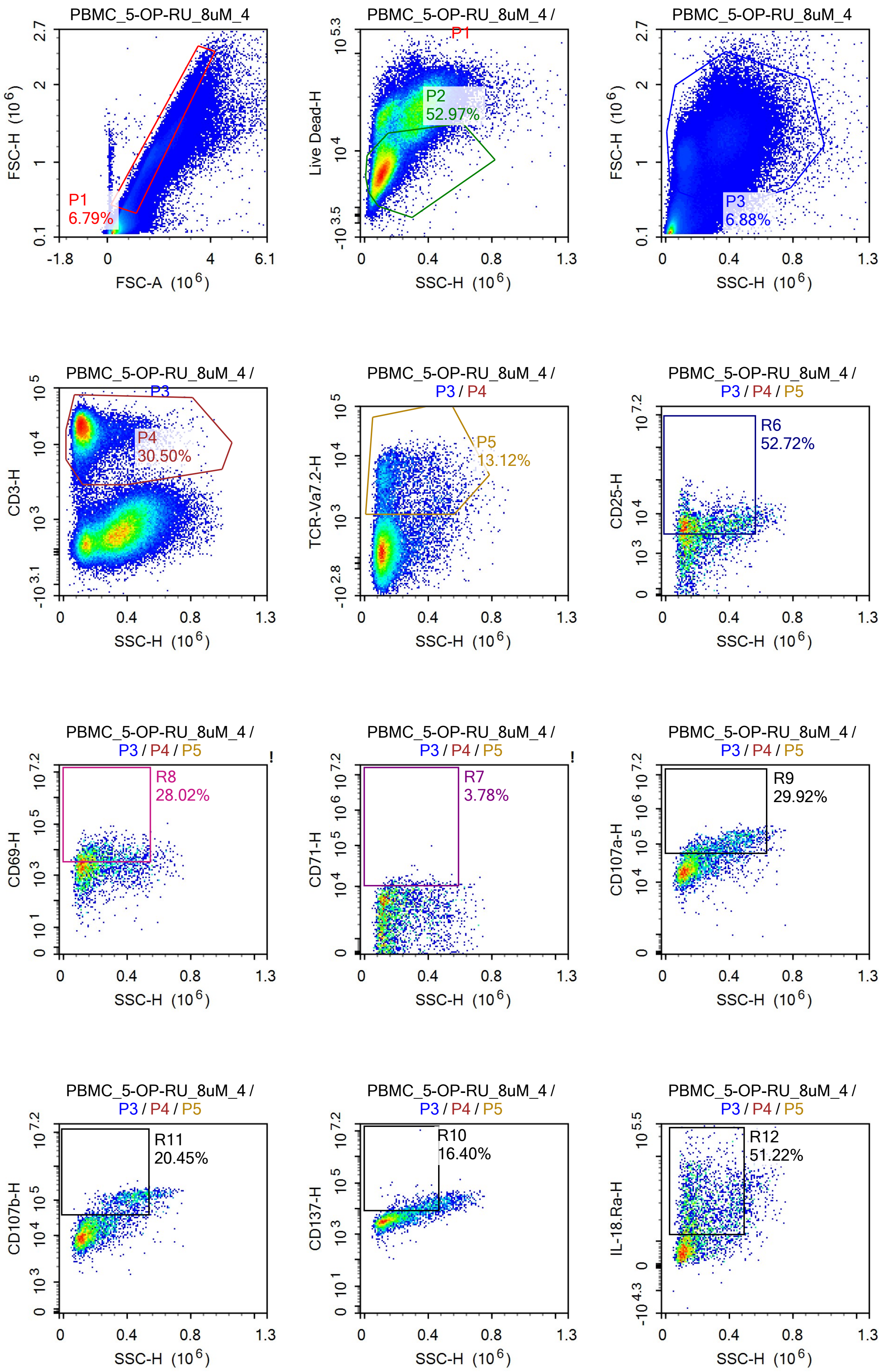
PBMC_5-OP-RU_8uM_4
P1
6.79%
FSC-H (10^6)
FSC-A (10^6)
PBMC_5-OP-RU_8uM_4 / P1
P2
52.97%
Live Dead-H
SSC-H (10^6)
PBMC_5-OP-RU_8uM_4
P3
6.88%
PBMC_5-OP-RU_8uM_4 / P3
P4
30.50%
CD3-H
PBMC_5-OP-RU_8uM_4 / P3 / P4
P5
13.12%
TCR-Va7.2-H
PBMC_5-OP-RU_8uM_4 / P3 / P4 / P5
R6
52.72%
CD25-H
R8
28.02%
CD69-H
R7
3.78%
CD71-H
R9
29.92%
CD107a-H
R11
20.45%
CD107b-H
R10
16.40%
CD137-H
R12
51.22%
IL-18.Ra-H

Sample Statistics of PBMC_5-OP-RU_8uM_4

| Gate | Count | % Parent | X | Y | Median X | Median Y |
|---|---|---|---|---|---|---|
| All | 1,373,585 | | | | | |
| P1 | 93,221 | 6.79% | FSC-A | FSC-H | 1,621,995 | 1,013,707 |
| P2 | 49,381 | 52.97% | SSC-H | Live Dead-H | 118,744 | 6,957 |
| P3 | 94,449 | 6.88% | SSC-H | FSC-H | 276,531 | 1,145,806 |
| P4 | 28,803 | 30.50% | SSC-H | CD3-H | 123,903 | 18,174 |
| P5 | 3,780 | 13.12% | SSC-H | TCR-Va7.2-H | 175,368 | 3,490 |
| R6 | 1,993 | 52.72% | SSC-H | CD25-H | 197,381 | 5,423 |
| R7 | 143 | 3.78% | SSC-H | CD71-H | 279,194 | 11,470 |
| R8 | 1,059 | 28.02% | SSC-H | CD69-H | 194,164 | 5,793 |
| R9 | 1,131 | 29.92% | SSC-H | CD107a-H | 368,305 | 92,671 |
| R10 | 620 | 16.40% | SSC-H | CD137-H | 370,789 | 11,781 |
| R11 | 773 | 20.45% | SSC-H | CD107b-H | 404,564 | 99,202 |
| R12 | 1,936 | 51.22% | SSC-H | IL-18.Ra-H | 209,057 | 26,576 |

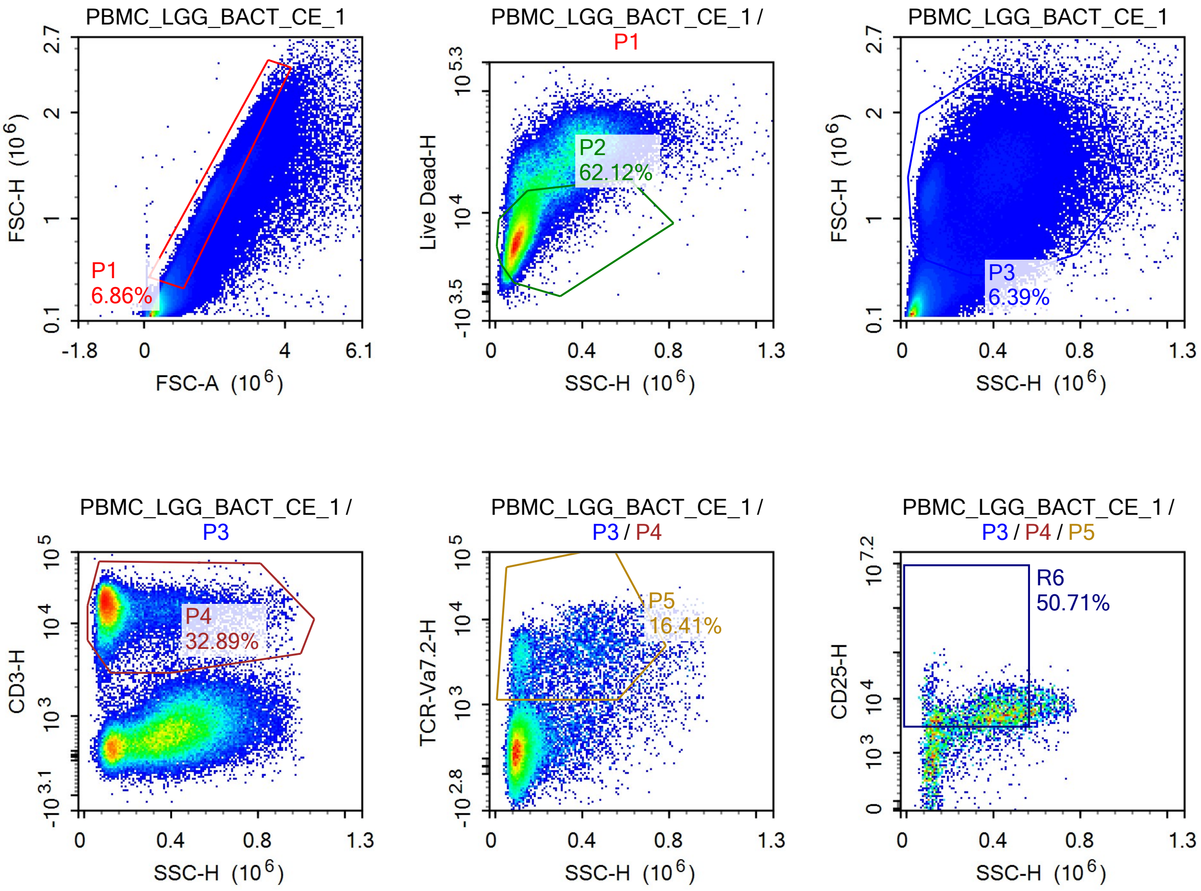

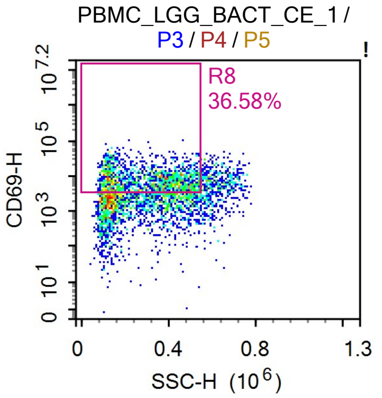


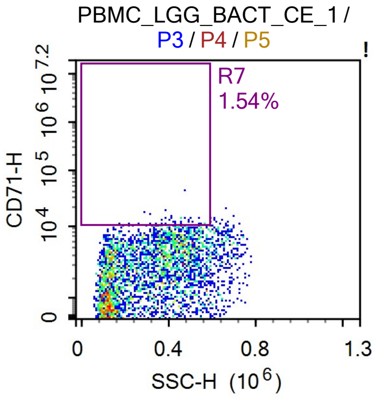


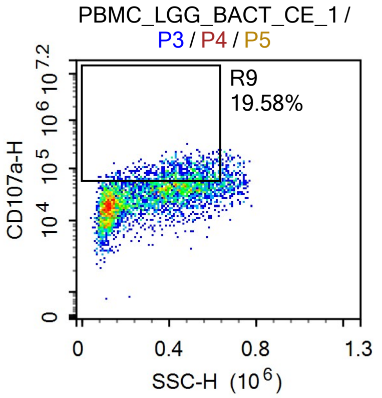


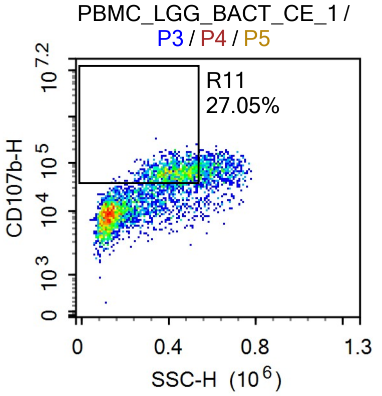


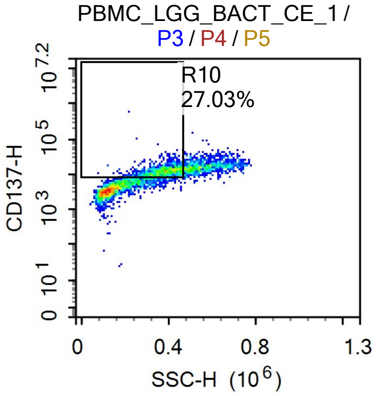


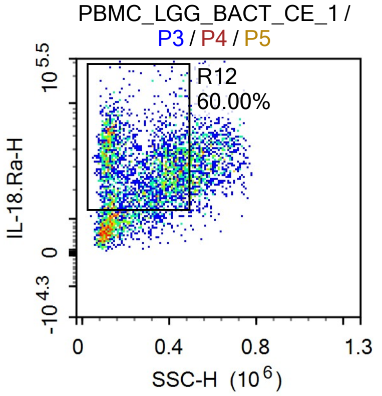


Sample Statistics of PBMC_LGG_BACT_CE_1

| Gate | Count | % Parent | X | Y | Median X | Median Y |
|---|---|---|---|---|---|---|
| All | 1,372,829 | | | | | |
| P1 | 94,214 | 6.86% | FSC-A | FSC-H | 1,466,746 | 867,619 |
| P2 | 58,529 | 62.12% | SSC-H | Live Dead-H | 104,343 | 6,763 |
| P3 | 87,743 | 6.39% | SSC-H | FSC-H | 285,451 | 1,224,059 |
| P4 | 28,858 | 32.89% | SSC-H | CD3-H | 115,733 | 16,398 |
| P5 | 4,735 | 16.41% | SSC-H | TCR-Va7.2-H | 295,384 | 3,519 |
| R6 | 2,401 | 50.71% | SSC-H | CD25-H | 375,753 | 5,291 |
| R7 | 73 | 1.54% | SSC-H | CD71-H | 431,532 | 10,936 |
| R8 | 1,732 | 36.58% | SSC-H | CD69-H | 314,707 | 7,653 |
| R9 | 927 | 19.58% | SSC-H | CD107a-H | 449,016 | 71,018 |
| R10 | 1,280 | 27.03% | SSC-H | CD137-H | 381,616 | 12,488 |
| R11 | 1,281 | 27.05% | SSC-H | CD107b-H | 427,742 | 62,414 |
| R12 | 2,841 | 60.00% | SSC-H | IL-18.Ra-H | 278,199 | 29,809 |

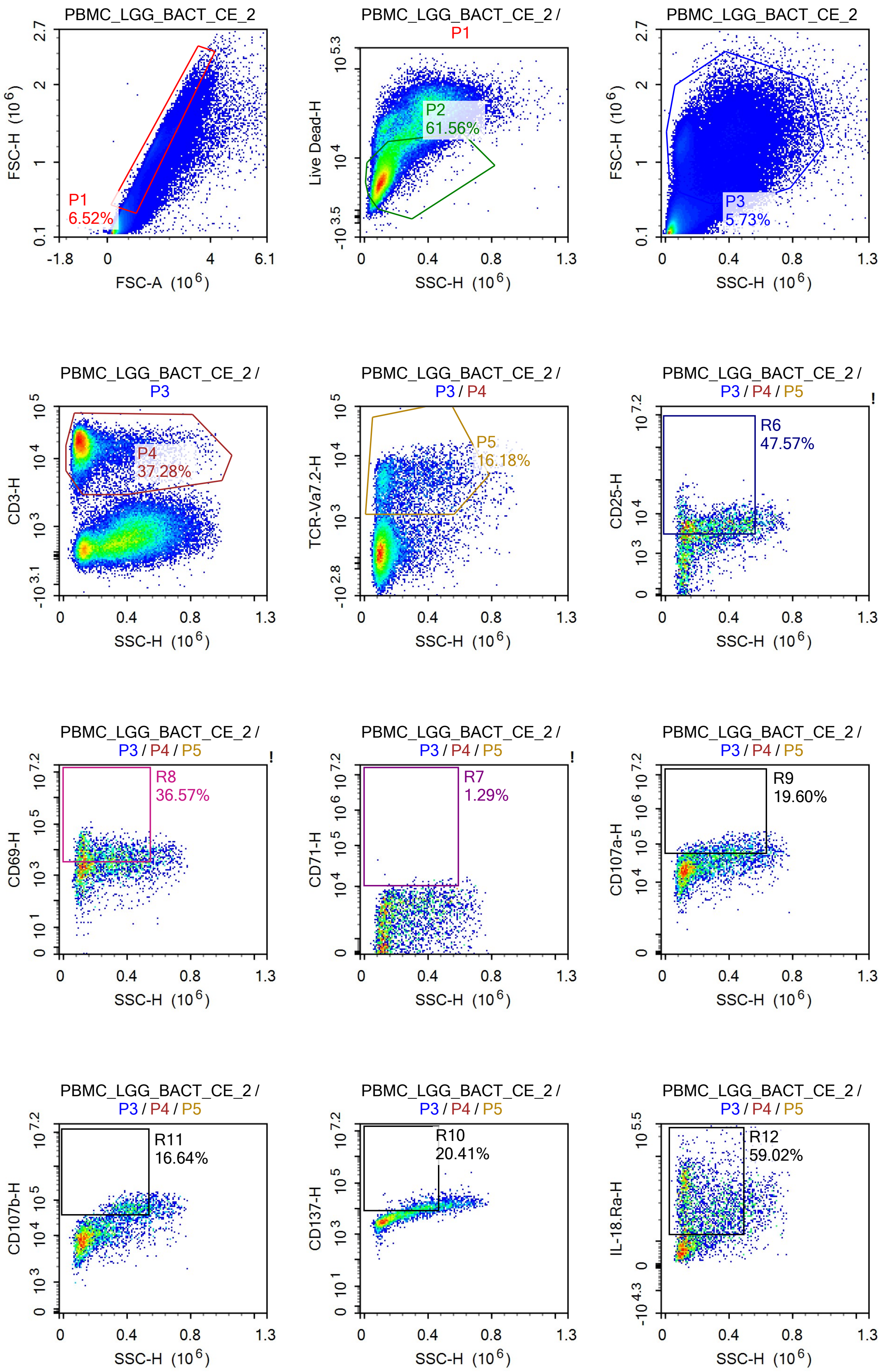
PBMC_LGG_BACT_CE_2
P1
6.52%
FSC-H (10^6)
FSC-A (10^6)
PBMC_LGG_BACT_CE_2 / P1
P2
61.56%
Live Dead-H
SSC-H (10^6)
PBMC_LGG_BACT_CE_2
P3
5.73%
FSC-H (10^6)
SSC-H (10^6)
PBMC_LGG_BACT_CE_2 / P3
P4
37.28%
CD3-H
SSC-H (10^6)
PBMC_LGG_BACT_CE_2 / P3 / P4
P5
16.18%
TCR-Va7.2-H
SSC-H (10^6)
PBMC_LGG_BACT_CE_2 / P3 / P4 / P5
R6
47.57%
CD25-H
SSC-H (10^6)
PBMC_LGG_BACT_CE_2 / P3 / P4 / P5
R8
36.57%
CD69-H
SSC-H (10^6)
PBMC_LGG_BACT_CE_2 / P3 / P4 / P5
R7
1.29%
CD71-H
SSC-H (10^6)
PBMC_LGG_BACT_CE_2 / P3 / P4 / P5
R9
19.60%
CD107a-H
SSC-H (10^6)
PBMC_LGG_BACT_CE_2 / P3 / P4 / P5
R11
16.64%
CD107b-H
SSC-H (10^6)
PBMC_LGG_BACT_CE_2 / P3 / P4 / P5
R10
20.41%
CD137-H
SSC-H (10^6)
PBMC_LGG_BACT_CE_2 / P3 / P4 / P5
R12
59.02%
IL-18.Ra-H
SSC-H (10^6)

Sample Statistics of PBMC_LGG_BACT_CE_2

| Gate | Count | % Parent | X | Y | Median X | Median Y |
|---|---|---|---|---|---|---|
| All | 1,076,879 | | | | | |
| P1 | 70,242 | 6.52% | FSC-A | FSC-H | 1,572,427 | 985,210 |
| P2 | 43,240 | 61.56% | SSC-H | Live Dead-H | 111,476 | 6,238 |
| P3 | 61,654 | 5.73% | SSC-H | FSC-H | 238,490 | 1,188,065 |
| P4 | 22,983 | 37.28% | SSC-H | CD3-H | 113,904 | 18,018 |
| P5 | 3,719 | 16.18% | SSC-H | TCR-Va7.2-H | 199,572 | 3,562 |
| R6 | 1,769 | 47.57% | SSC-H | CD25-H | 277,048 | 5,185 |
| R7 | 48 | 1.29% | SSC-H | CD71-H | 404,128 | 11,318 |
| R8 | 1,360 | 36.57% | SSC-H | CD69-H | 224,234 | 7,281 |
| R9 | 729 | 19.60% | SSC-H | CD107a-H | 395,304 | 69,922 |
| R10 | 759 | 20.41% | SSC-H | CD137-H | 366,496 | 10,817 |
| R11 | 619 | 16.64% | SSC-H | CD107b-H | 418,767 | 60,027 |
| R12 | 2,195 | 59.02% | SSC-H | IL-18.Ra-H | 206,894 | 29,297 |

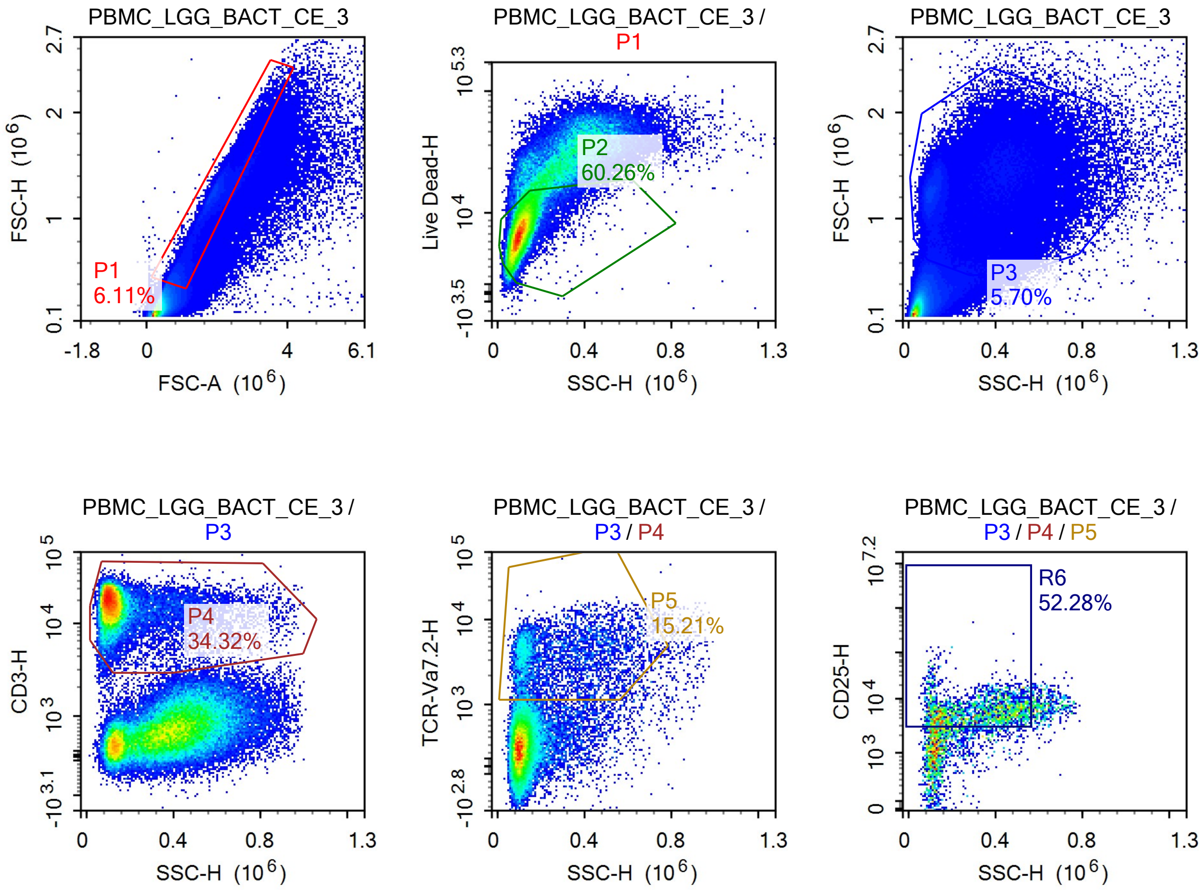

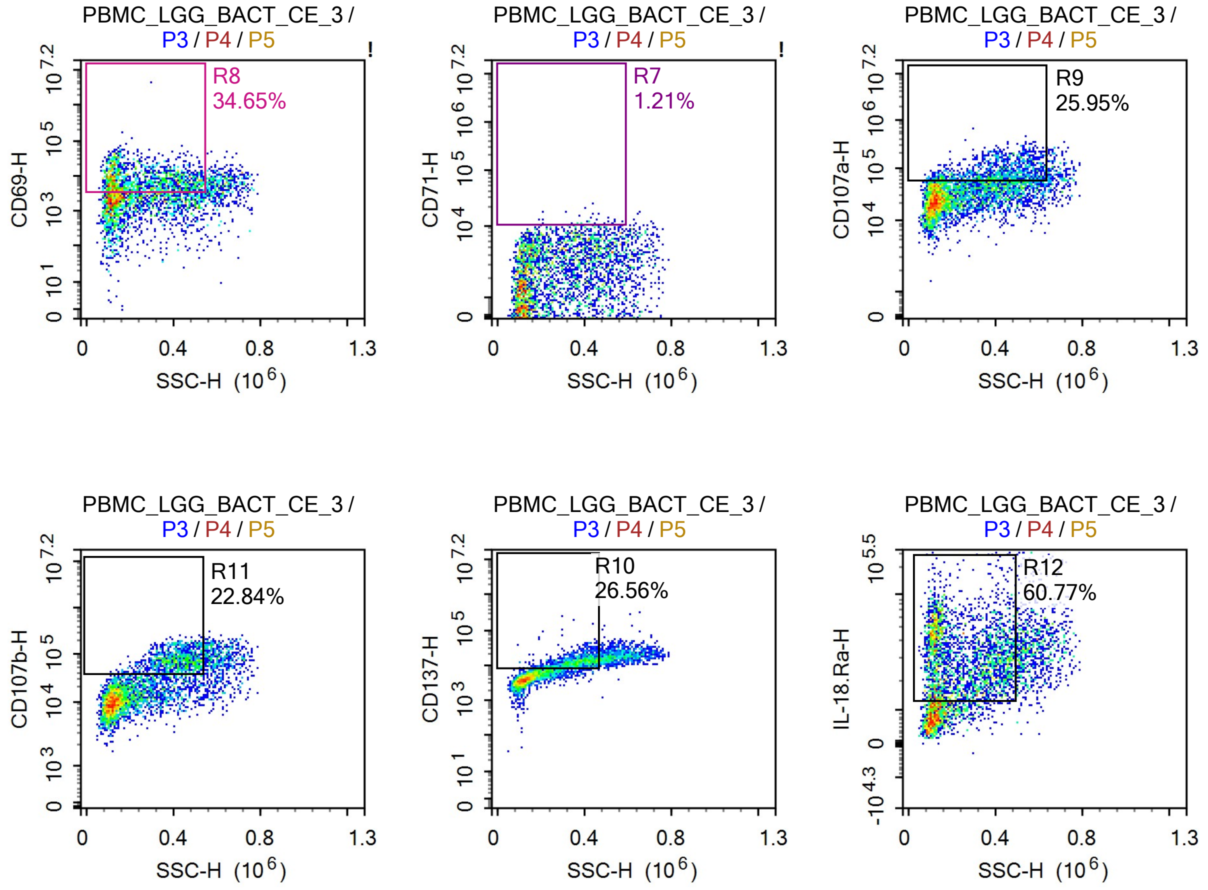


Sample Statistics of PBMC_LGG_BACT_CE_3

| Gate | Count | % Parent | X | Y | Median X | Median Y |
|---|---|---|---|---|---|---|
| All | 1,607,669 | | | | | |
| P1 | 98,233 | 6.11% | FSC-A | FSC-H | 1,606,912 | 997,133 |
| P2 | 59,196 | 60.26% | SSC-H | Live Dead-H | 101,716 | 7,354 |
| P3 | 91,622 | 5.70% | SSC-H | FSC-H | 274,493 | 1,246,475 |
| P4 | 31,445 | 34.32% | SSC-H | CD3-H | 113,586 | 17,844 |
| P5 | 4,782 | 15.21% | SSC-H | TCR-Va7.2-H | 226,256 | 3,473 |
| R6 | 2,500 | 52.28% | SSC-H | CD25-H | 325,818 | 5,584 |
| R7 | 58 | 1.21% | SSC-H | CD71-H | 422,921 | 10,947 |
| R8 | 1,657 | 34.65% | SSC-H | CD69-H | 237,305 | 7,317 |
| R9 | 1,241 | 25.95% | SSC-H | CD107a-H | 420,757 | 74,579 |
| R10 | 1,270 | 26.56% | SSC-H | CD137-H | 365,275 | 12,588 |
| R11 | 1,092 | 22.84% | SSC-H | CD107b-H | 417,237 | 72,257 |
| R12 | 2,906 | 60.77% | SSC-H | IL-18.Ra-H | 222,361 | 29,837 |

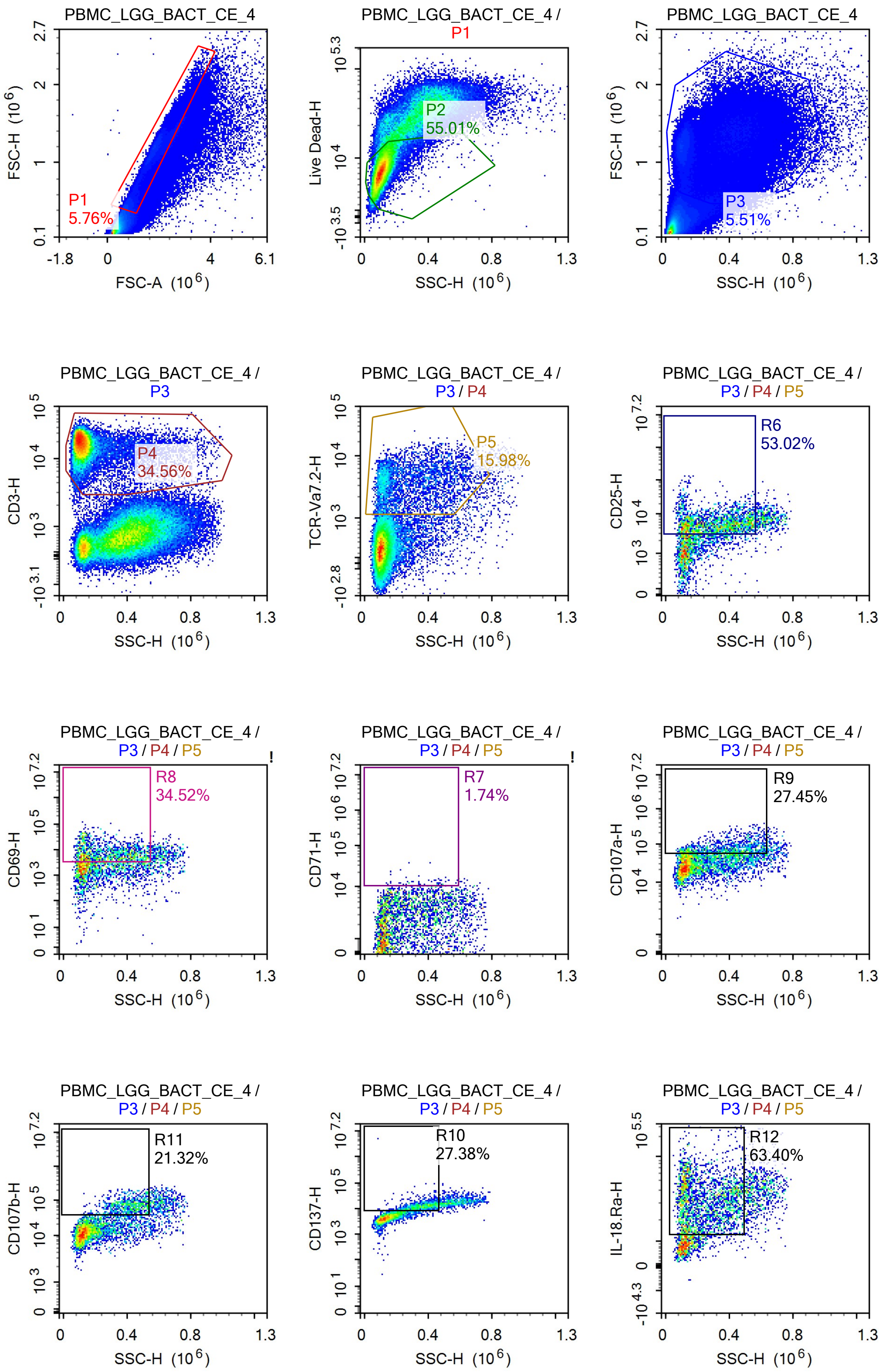
PBMC_LGG_BACT_CE_4
P1
5.76%
FSC-H (10^6)
FSC-A (10^6)
PBMC_LGG_BACT_CE_4 / P1
P2
55.01%
Live Dead-H
SSC-H (10^6)
PBMC_LGG_BACT_CE_4
P3
5.51%
FSC-H (10^6)
SSC-H (10^6)
PBMC_LGG_BACT_CE_4 / P3
P4
34.56%
CD3-H
SSC-H (10^6)
PBMC_LGG_BACT_CE_4 / P3 / P4
P5
15.98%
TCR-Va7.2-H
SSC-H (10^6)
PBMC_LGG_BACT_CE_4 / P3 / P4 / P5
R6
53.02%
CD25-H
SSC-H (10^6)
PBMC_LGG_BACT_CE_4 / P3 / P4 / P5
R8
34.52%
CD69-H
SSC-H (10^6)
PBMC_LGG_BACT_CE_4 / P3 / P4 / P5
R7
1.74%
CD71-H
SSC-H (10^6)
PBMC_LGG_BACT_CE_4 / P3 / P4 / P5
R9
27.45%
CD107a-H
SSC-H (10^6)
PBMC_LGG_BACT_CE_4 / P3 / P4 / P5
R11
21.32%
CD107b-H
SSC-H (10^6)
PBMC_LGG_BACT_CE_4 / P3 / P4 / P5
R10
27.38%
CD137-H
SSC-H (10^6)
PBMC_LGG_BACT_CE_4 / P3 / P4 / P5
R12
63.40%
IL-18.Ra-H
SSC-H (10^6)

Sample Statistics of PBMC_LGG_BACT_CE_4

| Gate | Count | % Parent | X | Y | Median X | Median Y |
|---|---|---|---|---|---|---|
| All | 1,452,583 | | | | | |
| P1 | 83,703 | 5.76% | FSC-A | FSC-H | 1,644,877 | 1,030,390 |
| P2 | 46,044 | 55.01% | SSC-H | Live Dead-H | 103,956 | 7,578 |
| P3 | 80,086 | 5.51% | SSC-H | FSC-H | 288,590 | 1,200,731 |
| P4 | 27,674 | 34.56% | SSC-H | CD3-H | 114,154 | 18,326 |
| P5 | 4,423 | 15.98% | SSC-H | TCR-Va7.2-H | 227,117 | 3,589 |
| R6 | 2,345 | 53.02% | SSC-H | CD25-H | 311,443 | 5,492 |
| R7 | 77 | 1.74% | SSC-H | CD71-H | 402,798 | 11,720 |
| R8 | 1,527 | 34.52% | SSC-H | CD69-H | 253,089 | 7,027 |
| R9 | 1,214 | 27.45% | SSC-H | CD107a-H | 392,981 | 69,835 |
| R10 | 1,211 | 27.38% | SSC-H | CD137-H | 361,372 | 12,308 |
| R11 | 943 | 21.32% | SSC-H | CD107b-H | 411,011 | 70,724 |
| R12 | 2,804 | 63.40% | SSC-H | IL-18.Ra-H | 217,623 | 31,916 |

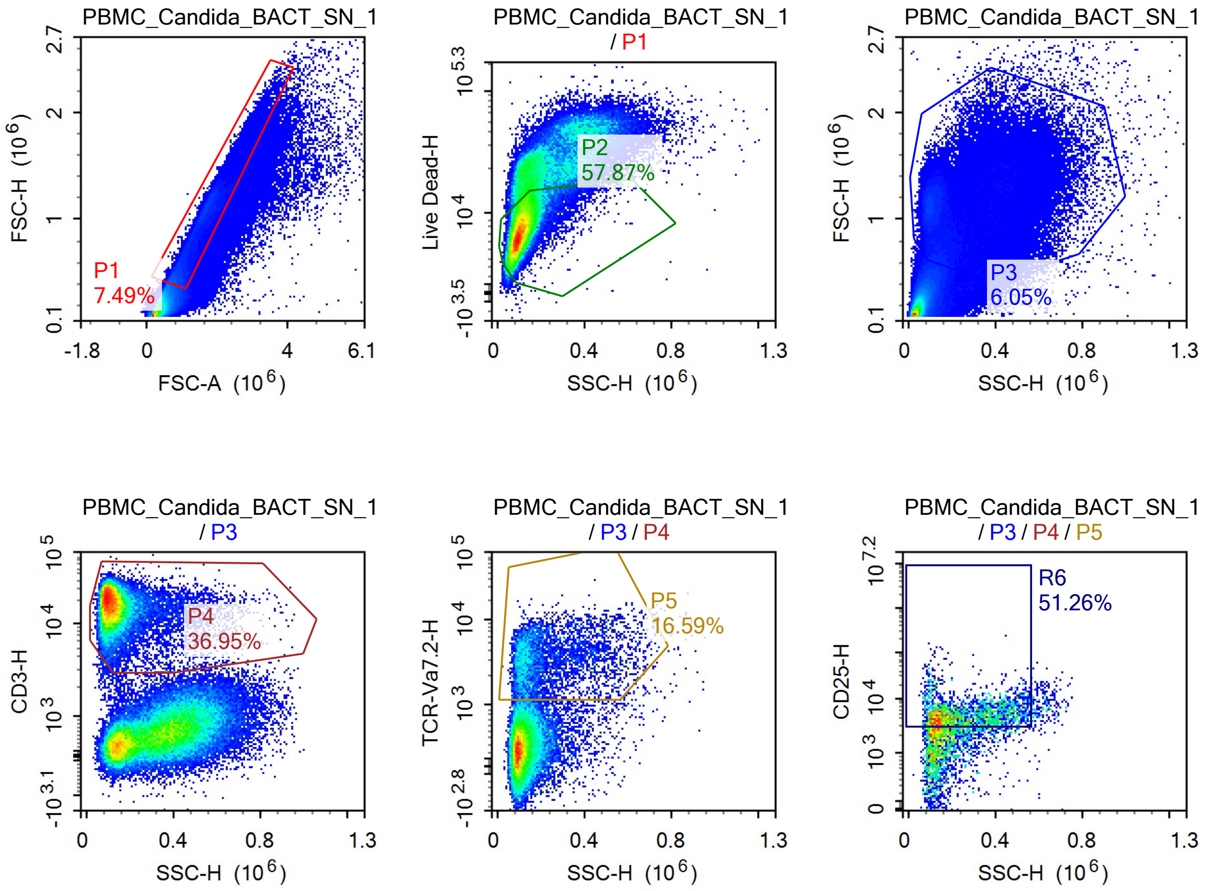

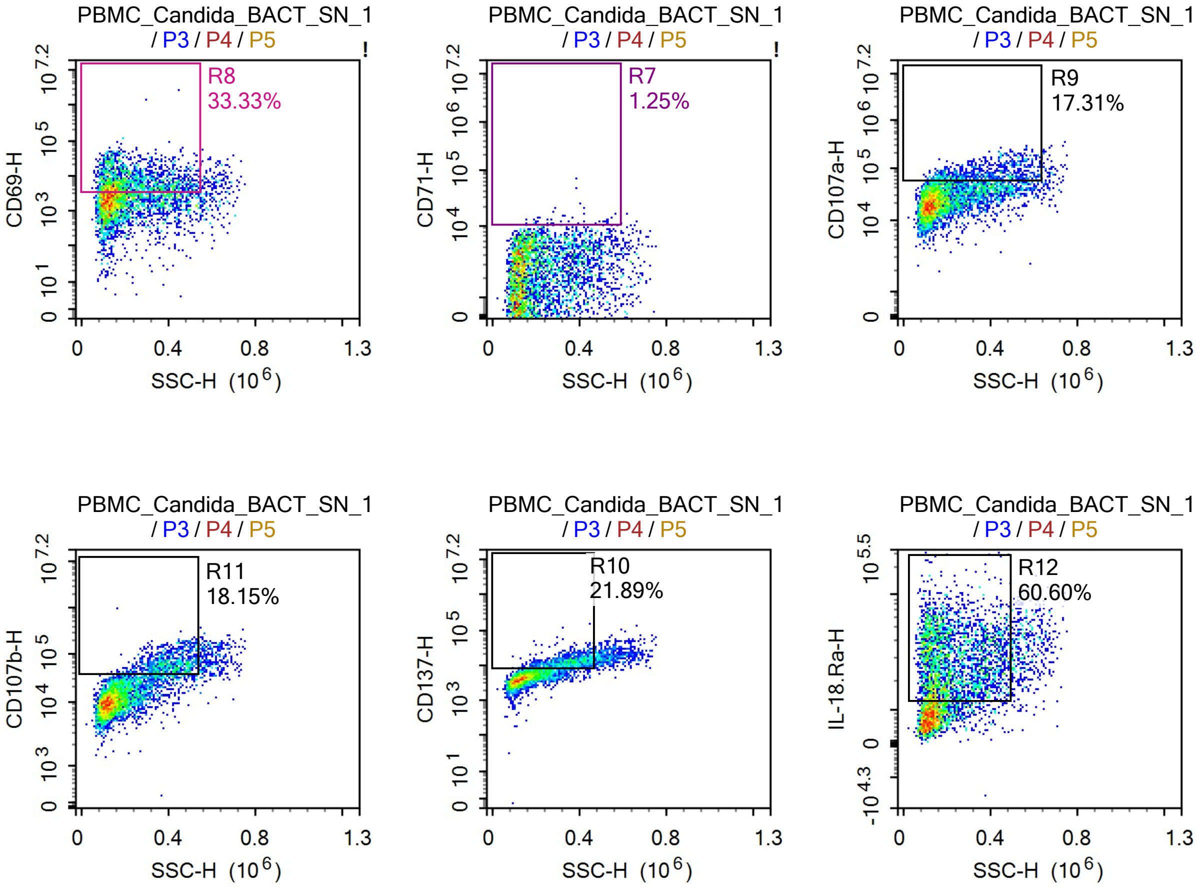


Sample Statistics of PBMC_Candida_BACT_SN_1

| Gate | Count | % Parent | X | Y | Median X | Median Y |
|---|---|---|---|---|---|---|
| All | 1,376,705 | | | | | |
| P1 | 103,080 | 7.49% | FSC-A | FSC-H | 1,322,647 | 777,457 |
| P2 | 59,649 | 57.87% | SSC-H | Live Dead-H | 102,081 | 7,465 |
| P3 | 83,338 | 6.05% | SSC-H | FSC-H | 204,417 | 1,094,415 |
| P4 | 30,790 | 36.95% | SSC-H | CD3-H | 112,084 | 17,615 |
| P5 | 5,107 | 16.59% | SSC-H | TCR-Va7.2-H | 171,966 | 3,164 |
| R6 | 2,618 | 51.26% | SSC-H | CD25-H | 217,050 | 4,837 |
| R7 | 64 | 1.25% | SSC-H | CD71-H | 318,158 | 11,599 |
| R8 | 1,702 | 33.33% | SSC-H | CD69-H | 192,779 | 6,699 |
| R9 | 884 | 17.31% | SSC-H | CD107a-H | 367,153 | 70,459 |
| R10 | 1,118 | 21.89% | SSC-H | CD137-H | 350,538 | 12,249 |
| R11 | 927 | 18.15% | SSC-H | CD107b-H | 401,747 | 60,204 |
| R12 | 3,095 | 60.60% | SSC-H | IL-18.Ra-H | 194,715 | 29,049 |

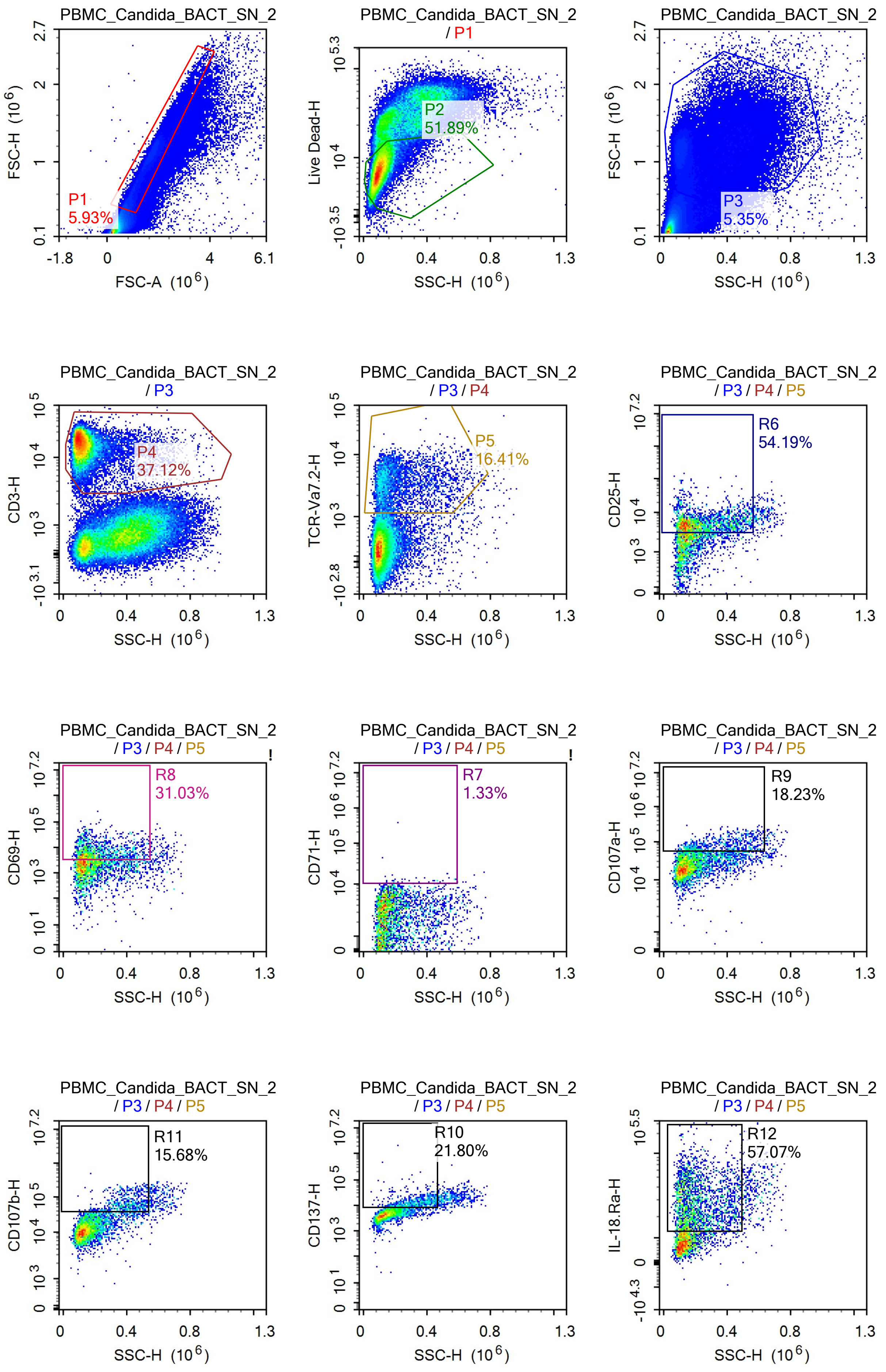

PBMC_Candida_BACT_SN_2
P1
5.93%
FSC-H (10^6)
FSC-A (10^6)
PBMC_Candida_BACT_SN_2
/ P1
P2
51.89%
Live Dead-H
SSC-H (10^6)
PBMC_Candida_BACT_SN_2
P3
5.35%
FSC-H (10^6)
SSC-H (10^6)
PBMC_Candida_BACT_SN_2
/ P3
P4
37.12%
CD3-H
SSC-H (10^6)
PBMC_Candida_BACT_SN_2
/ P3 / P4
P5
16.41%
TCR-Va7.2-H
SSC-H (10^6)
PBMC_Candida_BACT_SN_2
/ P3 / P4 / P5
R6
54.19%
CD25-H
SSC-H (10^6)
PBMC_Candida_BACT_SN_2
/ P3 / P4 / P5
R8
31.03%
CD69-H
SSC-H (10^6)
PBMC_Candida_BACT_SN_2
/ P3 / P4 / P5
R7
1.33%
CD71-H
SSC-H (10^6)
PBMC_Candida_BACT_SN_2
/ P3 / P4 / P5
R9
18.23%
CD107a-H
SSC-H (10^6)
PBMC_Candida_BACT_SN_2
/ P3 / P4 / P5
R11
15.68%
CD107b-H
SSC-H (10^6)
PBMC_Candida_BACT_SN_2
/ P3 / P4 / P5
R10
21.80%
CD137-H
SSC-H (10^6)
PBMC_Candida_BACT_SN_2
/ P3 / P4 / P5
R12
57.07%
IL-18.Ra-H
SSC-H (10^6)

Sample Statistics of PBMC_Candida_BACT_SN_2

| Gate | Count | % Parent | X | Y | Median X | Median Y |
|---|---|---|---|---|---|---|
| All | 1,288,482 | | | | | |
| P1 | 76,353 | 5.93% | FSC-A | FSC-H | 1,490,615 | 916,206 |
| P2 | 39,621 | 51.89% | SSC-H | Live Dead-H | 98,265 | 7,606 |
| P3 | 68,973 | 5.35% | SSC-H | FSC-H | 231,843 | 1,119,135 |
| P4 | 25,606 | 37.12% | SSC-H | CD3-H | 111,452 | 17,446 |
| P5 | 4,202 | 16.41% | SSC-H | TCR-Va7.2-H | 167,157 | 3,264 |
| R6 | 2,277 | 54.19% | SSC-H | CD25-H | 193,722 | 4,921 |
| R7 | 56 | 1.33% | SSC-H | CD71-H | 235,834 | 11,441 |
| R8 | 1,304 | 31.03% | SSC-H | CD69-H | 177,887 | 6,059 |
| R9 | 766 | 18.23% | SSC-H | CD107a-H | 355,626 | 72,339 |
| R10 | 916 | 21.80% | SSC-H | CD137-H | 330,539 | 12,716 |
| R11 | 659 | 15.68% | SSC-H | CD107b-H | 398,872 | 61,219 |
| R12 | 2,398 | 57.07% | SSC-H | IL-18.Ra-H | 187,654 | 27,795 |

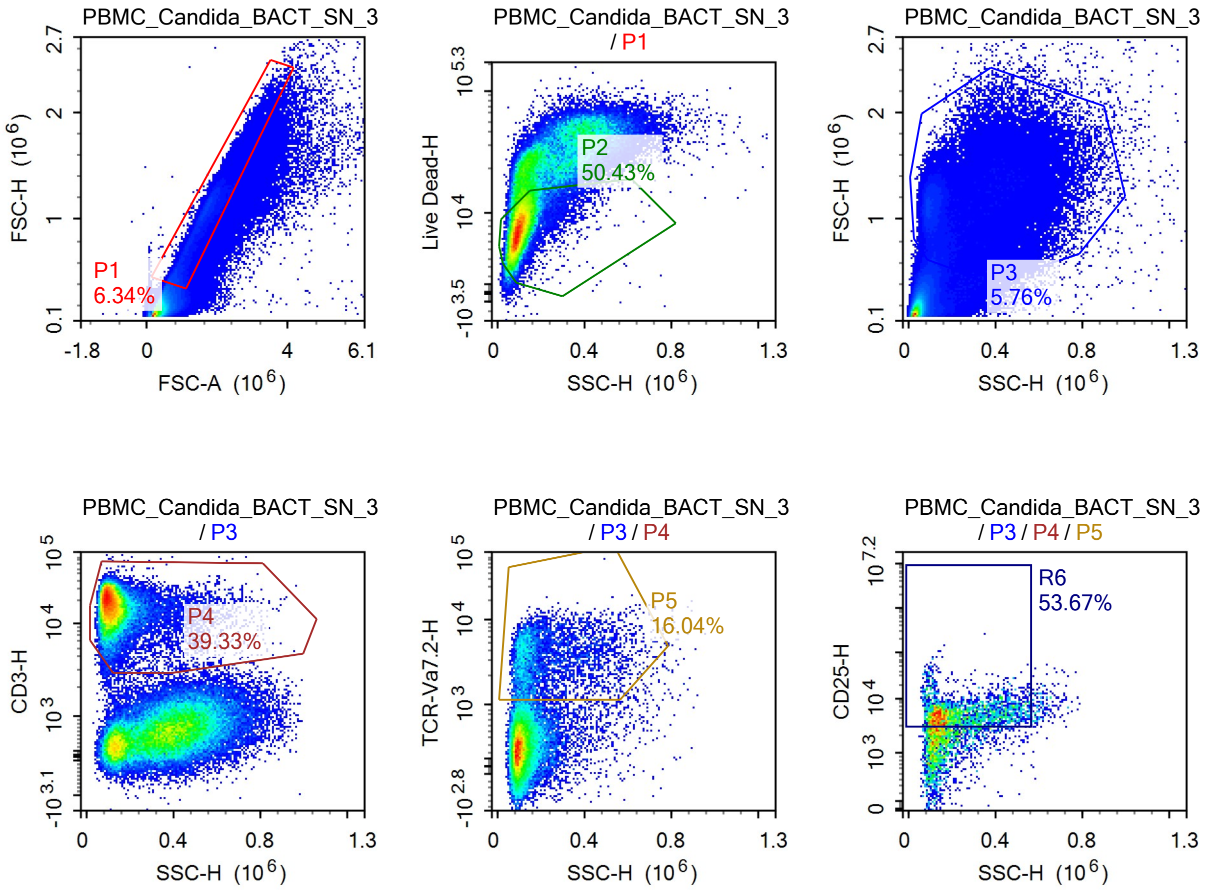

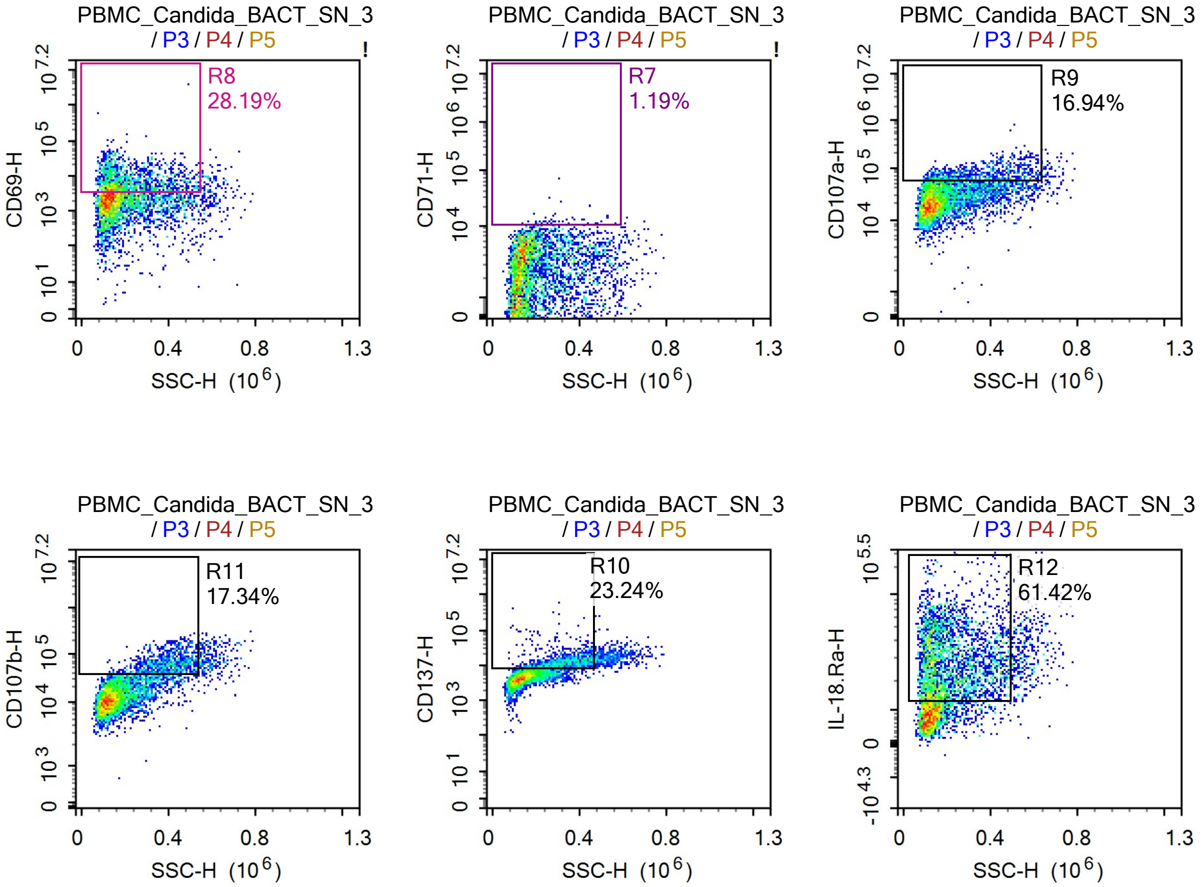


Sample Statistics of PBMC_Candida_BACT_SN_3

| Gate | Count | % Parent | X | Y | Median X | Median Y |
|---|---|---|---|---|---|---|
| All | 1,415,403 | | | | | |
| P1 | 89,758 | 6.34% | FSC-A | FSC-H | 1,510,094 | 934,596 |
| P2 | 45,261 | 50.43% | SSC-H | Live Dead-H | 96,571 | 7,722 |
| P3 | 81,529 | 5.76% | SSC-H | FSC-H | 208,046 | 1,117,747 |
| P4 | 32,062 | 39.33% | SSC-H | CD3-H | 109,365 | 17,273 |
| P5 | 5,143 | 16.04% | SSC-H | TCR-Va7.2-H | 163,296 | 3,234 |
| R6 | 2,760 | 53.67% | SSC-H | CD25-H | 198,951 | 5,131 |
| R7 | 61 | 1.19% | SSC-H | CD71-H | 277,256 | 11,657 |
| R8 | 1,450 | 28.19% | SSC-H | CD69-H | 180,822 | 6,516 |
| R9 | 871 | 16.94% | SSC-H | CD107a-H | 336,643 | 69,356 |
| R10 | 1,195 | 23.24% | SSC-H | CD137-H | 325,604 | 12,271 |
| R11 | 892 | 17.34% | SSC-H | CD107b-H | 396,836 | 61,034 |
| R12 | 3,159 | 61.42% | SSC-H | IL-18.Ra-H | 183,229 | 28,037 |

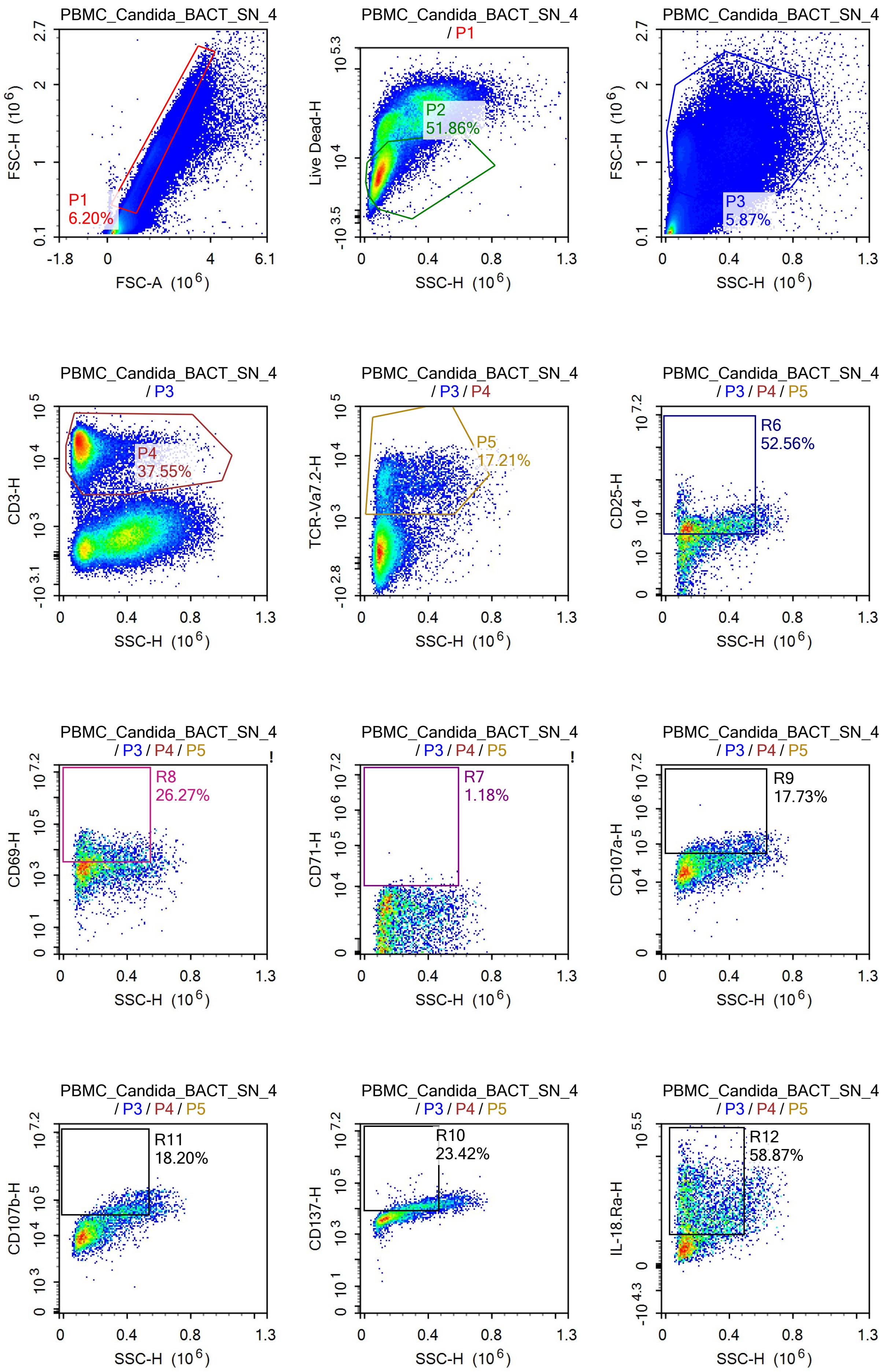

PBMC_Candida_BACT_SN_4
P1
6.20%
FSC-H (10^6)
FSC-A (10^6)
PBMC_Candida_BACT_SN_4
/ P1
P2
51.86%
Live Dead-H
SSC-H (10^6)
PBMC_Candida_BACT_SN_4
P3
5.87%
FSC-H (10^6)
SSC-H (10^6)
PBMC_Candida_BACT_SN_4
/ P3
P4
37.55%
CD3-H
SSC-H (10^6)
PBMC_Candida_BACT_SN_4
/ P3 / P4
P5
17.21%
TCR-Va7.2-H
SSC-H (10^6)
PBMC_Candida_BACT_SN_4
/ P3 / P4 / P5
R6
52.56%
CD25-H
SSC-H (10^6)
PBMC_Candida_BACT_SN_4
/ P3 / P4 / P5
R8
26.27%
CD69-H
SSC-H (10^6)
PBMC_Candida_BACT_SN_4
/ P3 / P4 / P5
R7
1.18%
CD71-H
SSC-H (10^6)
PBMC_Candida_BACT_SN_4
/ P3 / P4 / P5
R9
17.73%
CD107a-H
SSC-H (10^6)
PBMC_Candida_BACT_SN_4
/ P3 / P4 / P5
R11
18.20%
CD107b-H
SSC-H (10^6)
PBMC_Candida_BACT_SN_4
/ P3 / P4 / P5
R10
23.42%
CD137-H
SSC-H (10^6)
PBMC_Candida_BACT_SN_4
/ P3 / P4 / P5
R12
58.87%
IL-18.Ra-H
SSC-H (10^6)

Sample Statistics of PBMC_Candida_BACT_SN_4

| Gate | Count | % Parent | X | Y | Median X | Median Y |
|---|---|---|---|---|---|---|
| All | 1,451,807 | | | | | |
| P1 | 90,068 | 6.20% | FSC-A | FSC-H | 1,538,433 | 947,883 |
| P2 | 46,711 | 51.86% | SSC-H | Live Dead-H | 99,963 | 7,419 |
| P3 | 85,171 | 5.87% | SSC-H | FSC-H | 238,868 | 1,105,593 |
| P4 | 31,981 | 37.55% | SSC-H | CD3-H | 112,692 | 16,503 |
| P5 | 5,504 | 17.21% | SSC-H | TCR-Va7.2-H | 179,878 | 3,213 |
| R6 | 2,893 | 52.56% | SSC-H | CD25-H | 221,448 | 4,696 |
| R7 | 65 | 1.18% | SSC-H | CD71-H | 291,297 | 10,825 |
| R8 | 1,446 | 26.27% | SSC-H | CD69-H | 193,409 | 6,722 |
| R9 | 976 | 17.73% | SSC-H | CD107a-H | 359,257 | 71,653 |
| R10 | 1,289 | 23.42% | SSC-H | CD137-H | 338,357 | 11,882 |
| R11 | 1,002 | 18.20% | SSC-H | CD107b-H | 399,326 | 55,488 |
| R12 | 3,240 | 58.87% | SSC-H | IL-18.Ra-H | 203,579 | 26,121 |

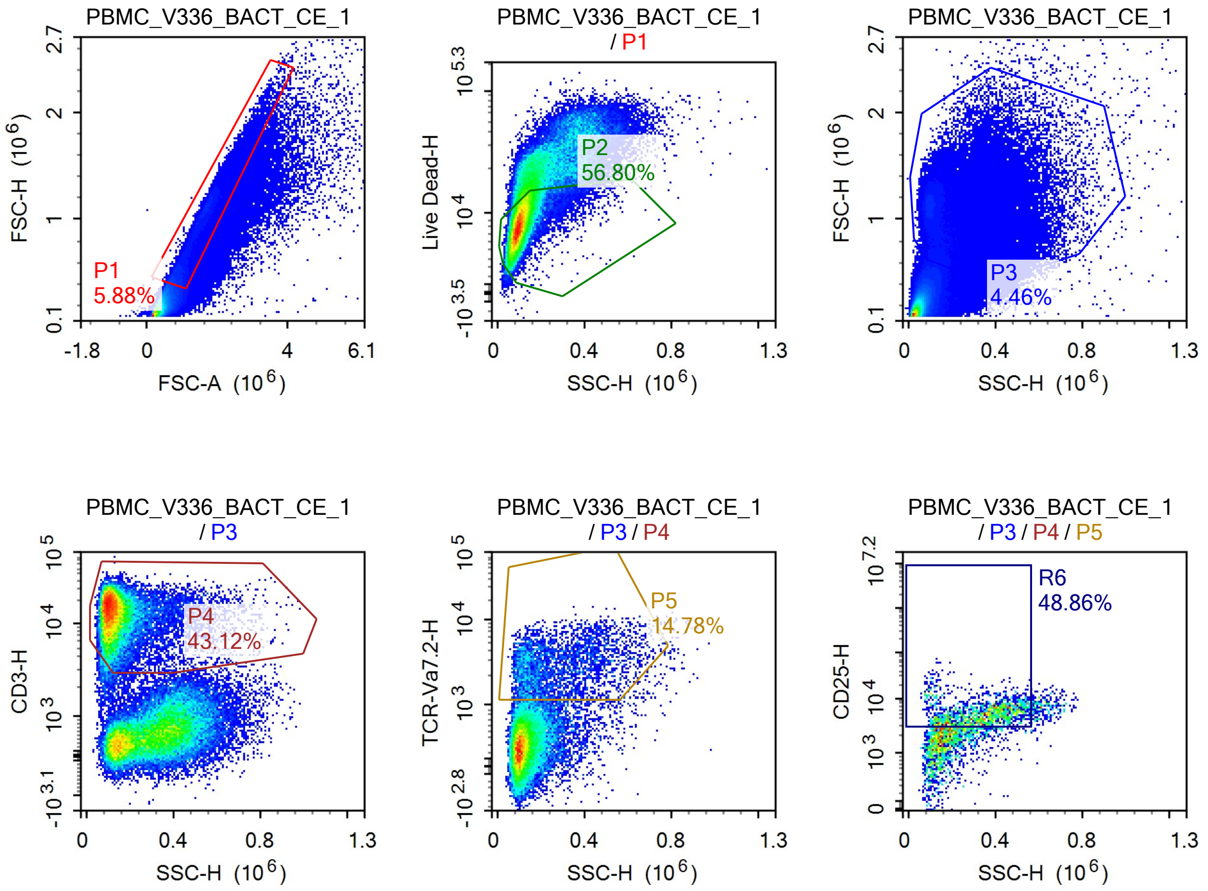

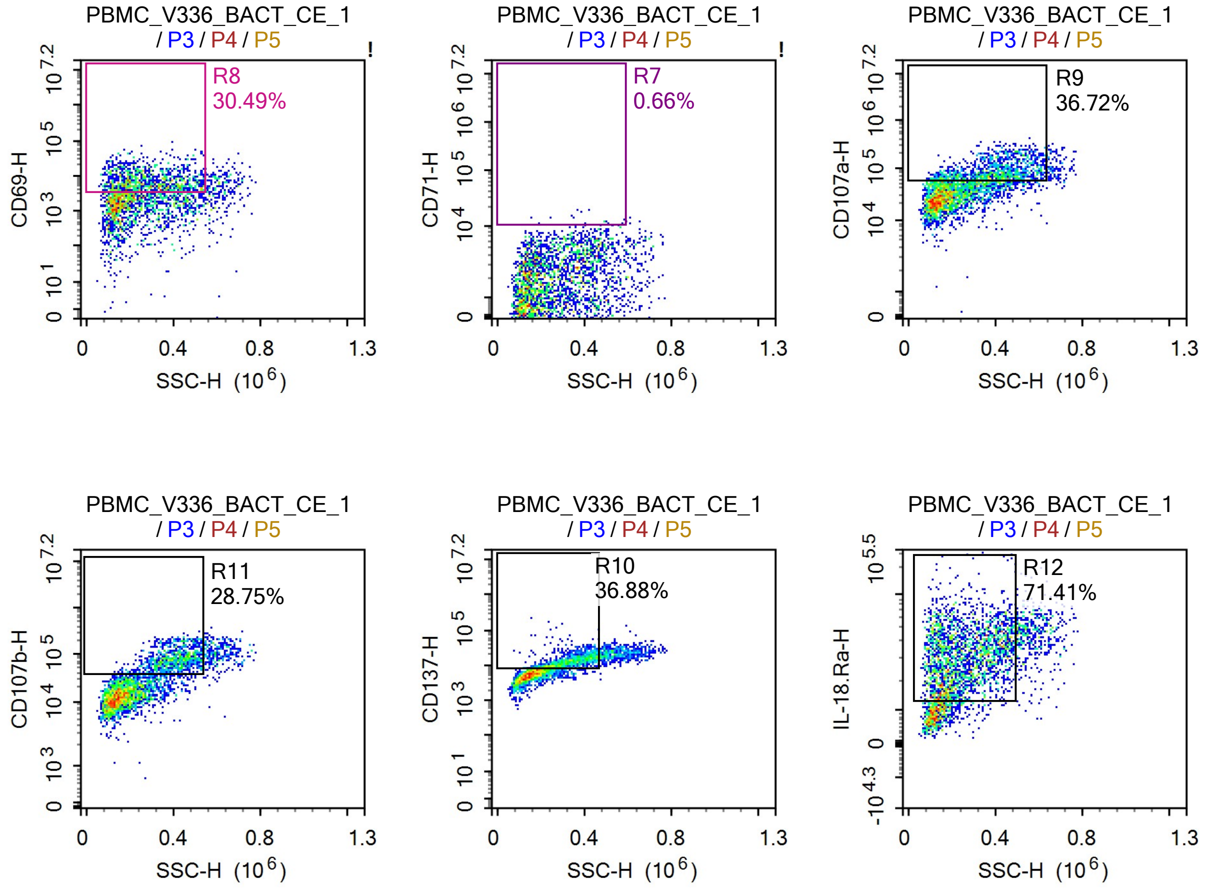


Sample Statistics of PBMC_V336_BACT_CE_1

| Gate | Count | % Parent | X | Y | Median X | Median Y |
|---|---|---|---|---|---|---|
| All | 1,331,564 | | | | | |
| P1 | 78,245 | 5.88% | FSC-A | FSC-H | 1,264,229 | 730,875 |
| P2 | 44,447 | 56.80% | SSC-H | Live Dead-H | 98,429 | 8,253 |
| P3 | 59,432 | 4.46% | SSC-H | FSC-H | 187,582 | 1,041,932 |
| P4 | 25,629 | 43.12% | SSC-H | CD3-H | 119,392 | 13,542 |
| P5 | 3,788 | 14.78% | SSC-H | TCR-Va7.2-H | 227,806 | 2,671 |
| R6 | 1,851 | 48.86% | SSC-H | CD25-H | 336,425 | 5,072 |
| R7 | 25 | 0.66% | SSC-H | CD71-H | 422,109 | 11,388 |
| R8 | 1,155 | 30.49% | SSC-H | CD69-H | 262,966 | 6,564 |
| R9 | 1,391 | 36.72% | SSC-H | CD107a-H | 397,673 | 76,288 |
| R10 | 1,397 | 36.88% | SSC-H | CD137-H | 334,917 | 12,924 |
| R11 | 1,089 | 28.75% | SSC-H | CD107b-H | 404,725 | 76,516 |
| R12 | 2,705 | 71.41% | SSC-H | IL-18.Ra-H | 237,360 | 31,400 |

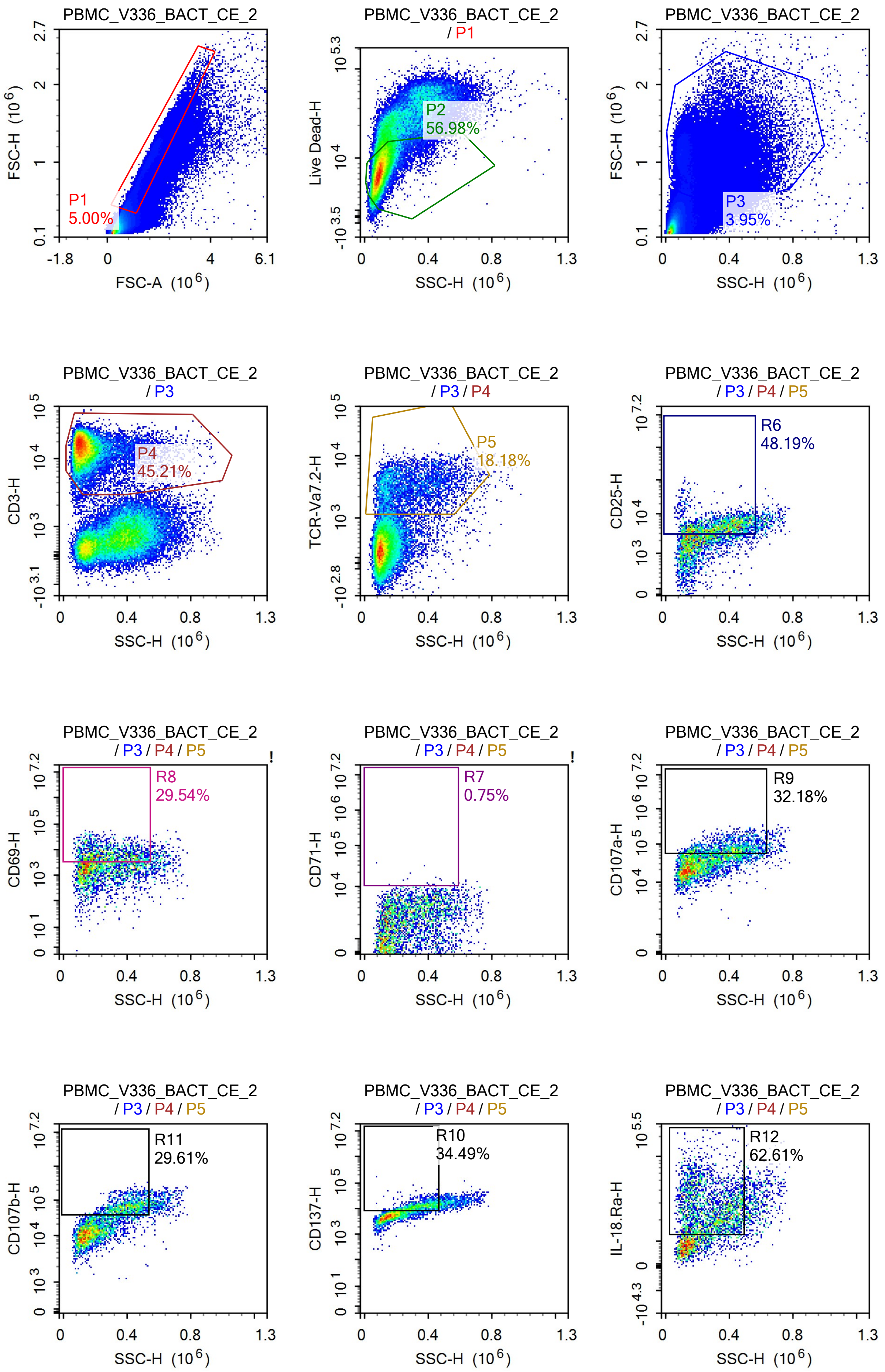
PBMC_V336_BACT_CE_2
FSC-H (10^6)
FSC-A (10^6)
P1
5.00%
PBMC_V336_BACT_CE_2
/ P1
Live Dead-H
SSC-H (10^6)
P2
56.98%
PBMC_V336_BACT_CE_2
FSC-H (10^6)
SSC-H (10^6)
P3
3.95%
PBMC_V336_BACT_CE_2
/ P3
CD3-H
SSC-H (10^6)
P4
45.21%
PBMC_V336_BACT_CE_2
/ P3 / P4
TCR-Va7.2-H
SSC-H (10^6)
P5
18.18%
PBMC_V336_BACT_CE_2
/ P3 / P4 / P5
CD25-H
SSC-H (10^6)
R6
48.19%
PBMC_V336_BACT_CE_2
/ P3 / P4 / P5
CD69-H
SSC-H (10^6)
R8
29.54%
PBMC_V336_BACT_CE_2
/ P3 / P4 / P5
CD71-H
SSC-H (10^6)
R7
0.75%
PBMC_V336_BACT_CE_2
/ P3 / P4 / P5
CD107a-H
SSC-H (10^6)
R9
32.18%
PBMC_V336_BACT_CE_2
/ P3 / P4 / P5
CD107b-H
SSC-H (10^6)
R11
29.61%
PBMC_V336_BACT_CE_2
/ P3 / P4 / P5
CD137-H
SSC-H (10^6)
R10
34.49%
PBMC_V336_BACT_CE_2
/ P3 / P4 / P5
IL-18.Ra-H
SSC-H (10^6)
R12
62.61%

Sample Statistics of PBMC_V336_BACT_CE_2

| Gate | Count | % Parent | X | Y | Median X | Median Y |
|---|---|---|---|---|---|---|
| All | 1,318,516 | | | | | |
| P1 | 65,924 | 5.00% | FSC-A | FSC-H | 1,286,714 | 739,922 |
| P2 | 37,565 | 56.98% | SSC-H | Live Dead-H | 96,492 | 7,595 |
| P3 | 52,130 | 3.95% | SSC-H | FSC-H | 194,715 | 1,040,206 |
| P4 | 23,568 | 45.21% | SSC-H | CD3-H | 120,678 | 15,525 |
| P5 | 4,285 | 18.18% | SSC-H | TCR-Va7.2-H | 247,041 | 2,938 |
| R6 | 2,065 | 48.19% | SSC-H | CD25-H | 339,247 | 4,596 |
| R7 | 32 | 0.75% | SSC-H | CD71-H | 383,359 | 10,920 |
| R8 | 1,266 | 29.54% | SSC-H | CD69-H | 263,637 | 6,527 |
| R9 | 1,379 | 32.18% | SSC-H | CD107a-H | 411,765 | 74,097 |
| R10 | 1,478 | 34.49% | SSC-H | CD137-H | 344,946 | 11,933 |
| R11 | 1,269 | 29.61% | SSC-H | CD107b-H | 408,578 | 65,665 |
| R12 | 2,683 | 62.61% | SSC-H | IL-18.Ra-H | 262,245 | 26,283 |

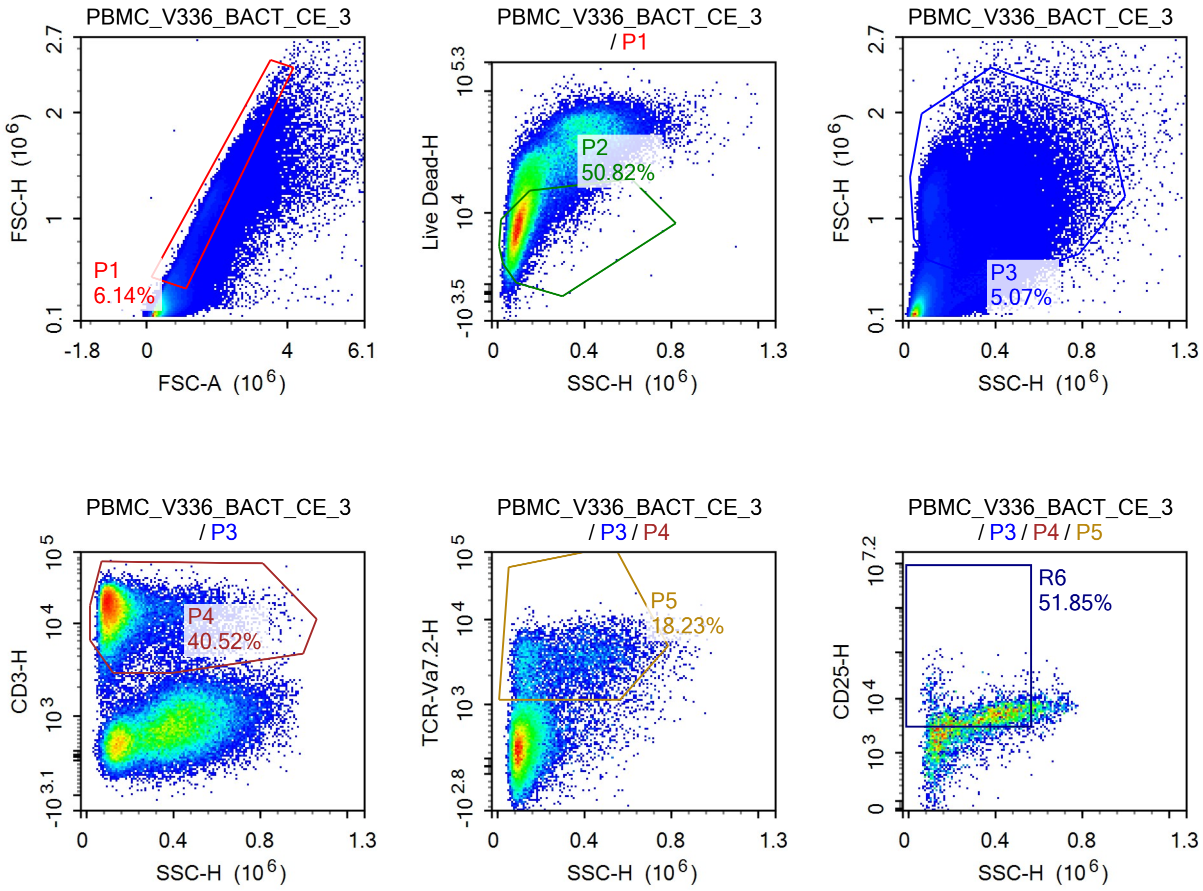

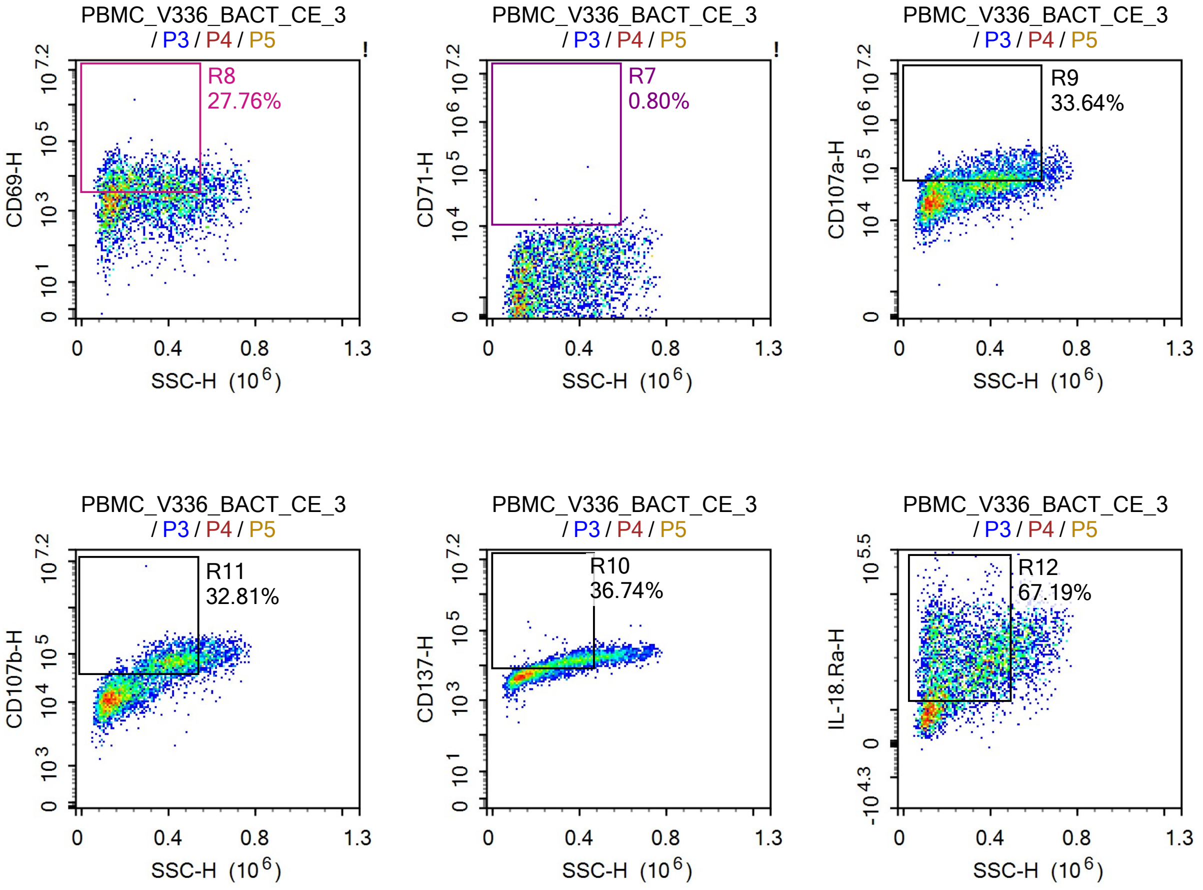


Sample Statistics of PBMC_V336_BACT_CE_3

| Gate | Count | % Parent | X | Y | Median X | Median Y |
|---|---|---|---|---|---|---|
| All | 1,438,199 | | | | | |
| P1 | 88,255 | 6.14% | FSC-A | FSC-H | 1,293,240 | 739,046 |
| P2 | 44,850 | 50.82% | SSC-H | Live Dead-H | 96,545 | 8,247 |
| P3 | 72,950 | 5.07% | SSC-H | FSC-H | 216,194 | 1,076,175 |
| P4 | 29,557 | 40.52% | SSC-H | CD3-H | 118,782 | 15,044 |
| P5 | 5,389 | 18.23% | SSC-H | TCR-Va7.2-H | 264,444 | 2,910 |
| R6 | 2,794 | 51.85% | SSC-H | CD25-H | 363,455 | 4,697 |
| R7 | 43 | 0.80% | SSC-H | CD71-H | 424,672 | 10,869 |
| R8 | 1,496 | 27.76% | SSC-H | CD69-H | 254,885 | 7,059 |
| R9 | 1,813 | 33.64% | SSC-H | CD107a-H | 417,315 | 71,324 |
| R10 | 1,980 | 36.74% | SSC-H | CD137-H | 347,178 | 12,395 |
| R11 | 1,768 | 32.81% | SSC-H | CD107b-H | 410,200 | 68,865 |
| R12 | 3,621 | 67.19% | SSC-H | IL-18.Ra-H | 261,289 | 28,438 |

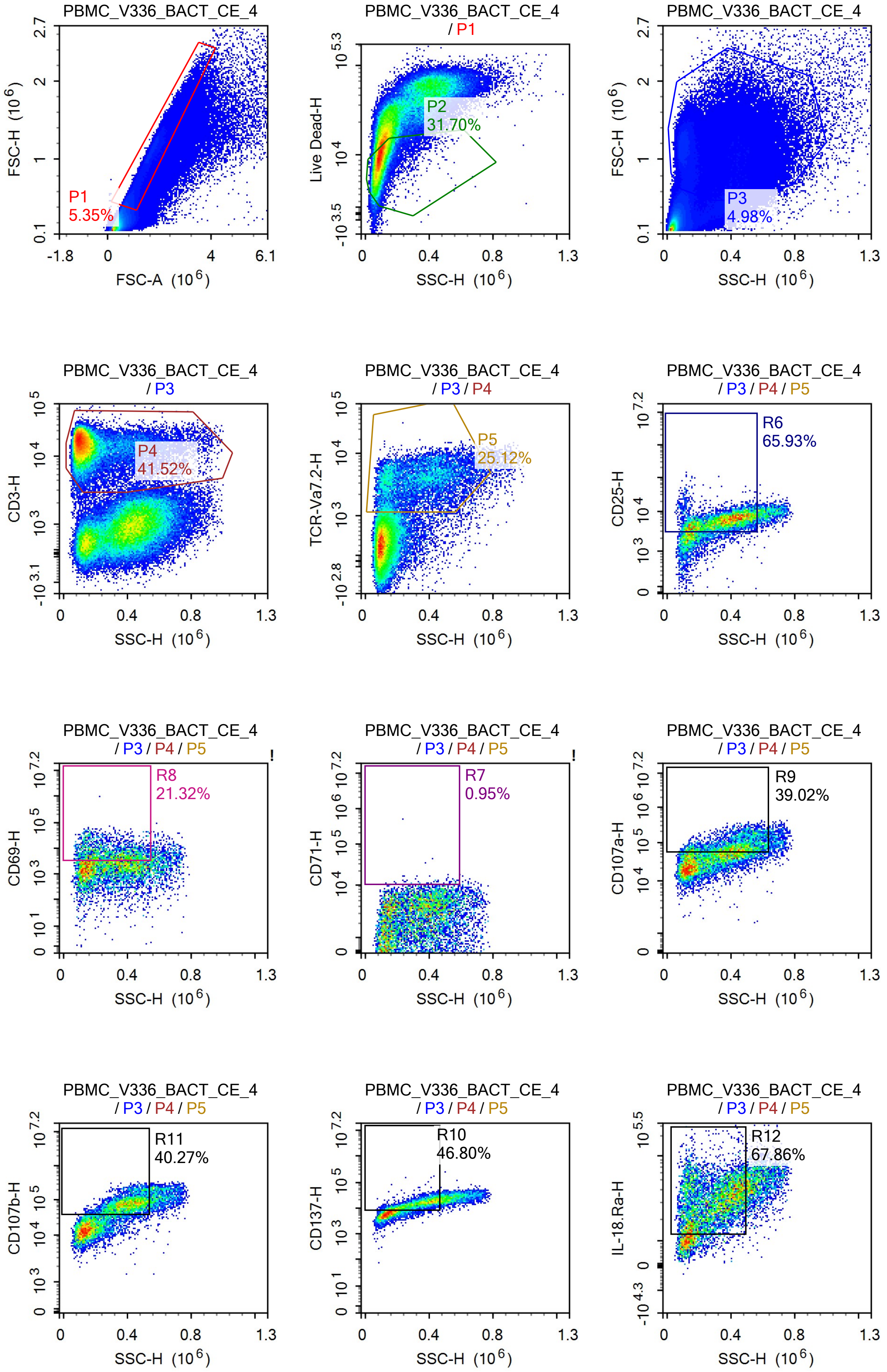

PBMC_V336_BACT_CE_4
P1
5.35%
FSC-H (10^6)
FSC-A (10^6)
PBMC_V336_BACT_CE_4
/ P1
P2
31.70%
Live Dead-H
SSC-H (10^6)
PBMC_V336_BACT_CE_4
P3
4.98%
FSC-H (10^6)
SSC-H (10^6)
PBMC_V336_BACT_CE_4
/ P3
P4
41.52%
CD3-H
SSC-H (10^6)
PBMC_V336_BACT_CE_4
/ P3 / P4
P5
25.12%
TCR-Va7.2-H
SSC-H (10^6)
PBMC_V336_BACT_CE_4
/ P3 / P4 / P5
R6
65.93%
CD25-H
SSC-H (10^6)
PBMC_V336_BACT_CE_4
/ P3 / P4 / P5
R8
21.32%
CD69-H
SSC-H (10^6)
PBMC_V336_BACT_CE_4
/ P3 / P4 / P5
R7
0.95%
CD71-H
SSC-H (10^6)
PBMC_V336_BACT_CE_4
/ P3 / P4 / P5
R9
39.02%
CD107a-H
SSC-H (10^6)
PBMC_V336_BACT_CE_4
/ P3 / P4 / P5
R11
40.27%
CD107b-H
SSC-H (10^6)
PBMC_V336_BACT_CE_4
/ P3 / P4 / P5
R10
46.80%
CD137-H
SSC-H (10^6)
PBMC_V336_BACT_CE_4
/ P3 / P4 / P5
R12
67.86%
IL-18.Ra-H
SSC-H (10^6)

Sample Statistics of PBMC_V336_BACT_CE_4

| Gate | Count | % Parent | X | Y | Median X | Median Y |
|---|---|---|---|---|---|---|
| All | 1,812,759 | | | | | |
| P1 | 97,031 | 5.35% | FSC-A | FSC-H | 1,417,979 | 839,634 |
| P2 | 30,760 | 31.70% | SSC-H | Live Dead-H | 88,928 | 8,598 |
| P3 | 90,211 | 4.98% | SSC-H | FSC-H | 272,864 | 1,071,255 |
| P4 | 37,460 | 41.52% | SSC-H | CD3-H | 125,191 | 15,960 |
| P5 | 9,410 | 25.12% | SSC-H | TCR-Va7.2-H | 323,508 | 3,450 |
| R6 | 6,204 | 65.93% | SSC-H | CD25-H | 354,459 | 5,748 |
| R7 | 89 | 0.95% | SSC-H | CD71-H | 418,260 | 11,207 |
| R8 | 2,006 | 21.32% | SSC-H | CD69-H | 292,854 | 6,273 |
| R9 | 3,672 | 39.02% | SSC-H | CD107a-H | 441,302 | 78,899 |
| R10 | 4,404 | 46.80% | SSC-H | CD137-H | 330,253 | 14,595 |
| R11 | 3,789 | 40.27% | SSC-H | CD107b-H | 406,901 | 75,313 |
| R12 | 6,386 | 67.86% | SSC-H | IL-18.Ra-H | 290,078 | 31,550 |

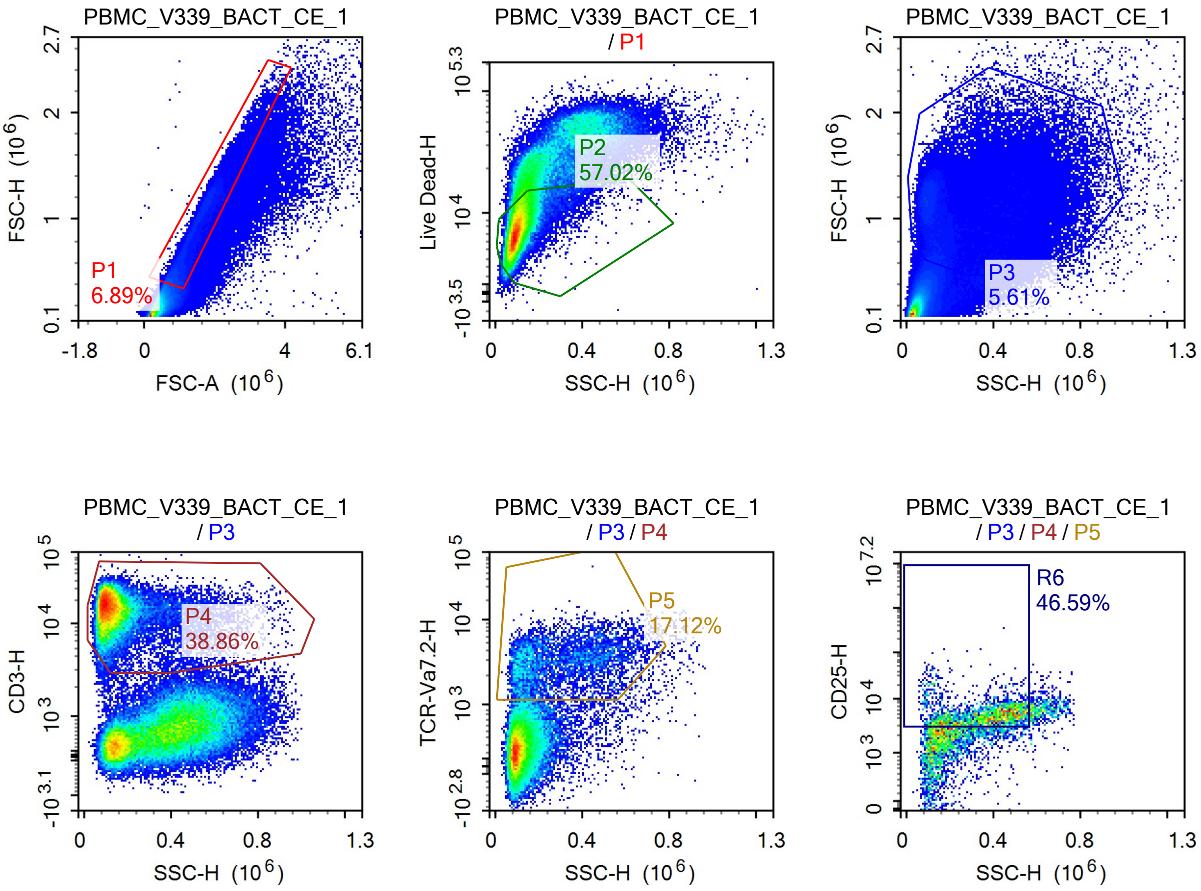

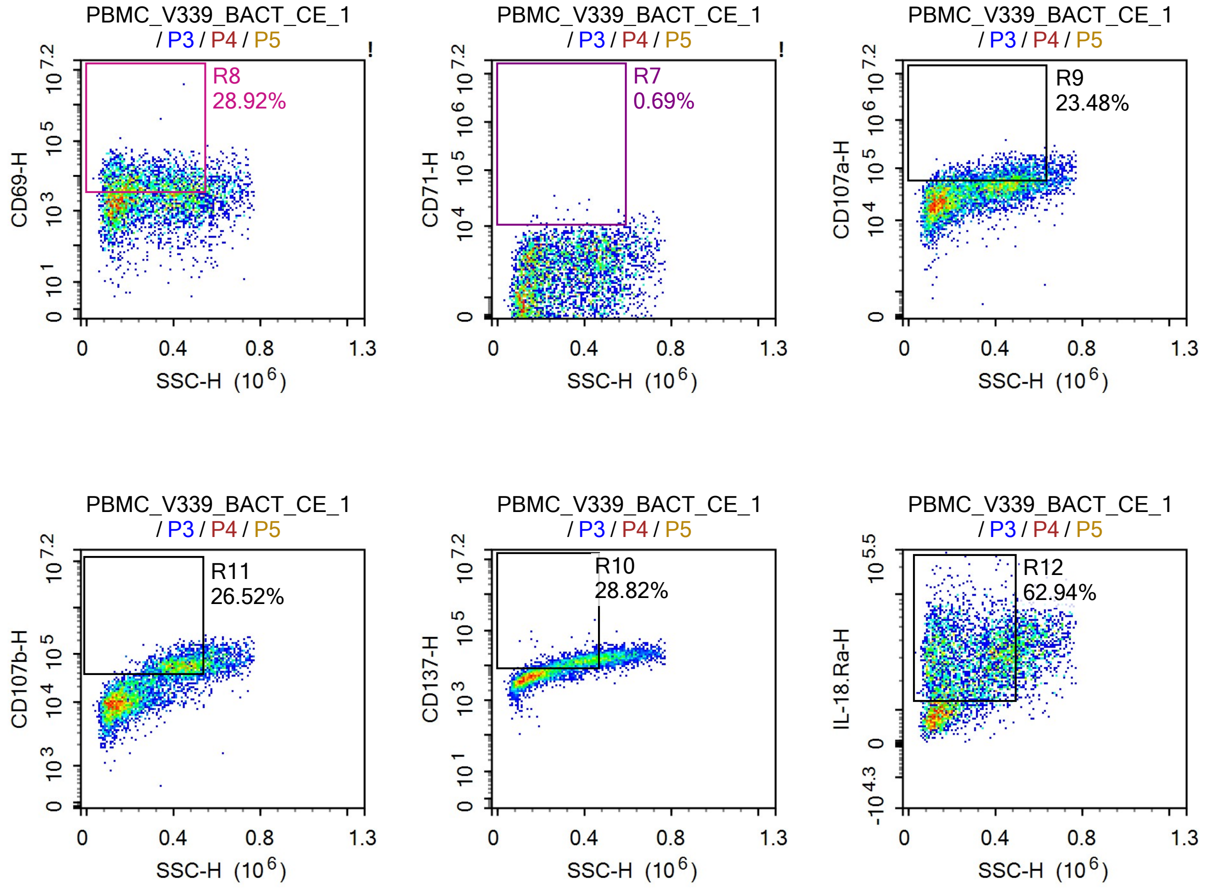


Sample Statistics of PBMC_V339_BACT_CE_1

| Gate | Count | % Parent | X | Y | Median X | Median Y |
|---|---|---|---|---|---|---|
| All | 1,486,064 | | | | | |
| P1 | 102,367 | 6.89% | FSC-A | FSC-H | 1,257,556 | 715,116 |
| P2 | 58,366 | 57.02% | SSC-H | Live Dead-H | 98,705 | 7,701 |
| P3 | 83,303 | 5.61% | SSC-H | FSC-H | 226,871 | 1,058,799 |
| P4 | 32,375 | 38.86% | SSC-H | CD3-H | 117,090 | 15,384 |
| P5 | 5,542 | 17.12% | SSC-H | TCR-Va7.2-H | 268,258 | 2,962 |
| R6 | 2,582 | 46.59% | SSC-H | CD25-H | 376,873 | 4,673 |
| R7 | 38 | 0.69% | SSC-H | CD71-H | 401,836 | 11,345 |
| R8 | 1,603 | 28.92% | SSC-H | CD69-H | 253,397 | 7,155 |
| R9 | 1,301 | 23.48% | SSC-H | CD107a-H | 446,661 | 69,150 |
| R10 | 1,597 | 28.82% | SSC-H | CD137-H | 369,521 | 12,387 |
| R11 | 1,470 | 26.52% | SSC-H | CD107b-H | 431,100 | 57,616 |
| R12 | 3,488 | 62.94% | SSC-H | IL-18.Ra-H | 263,455 | 29,045 |

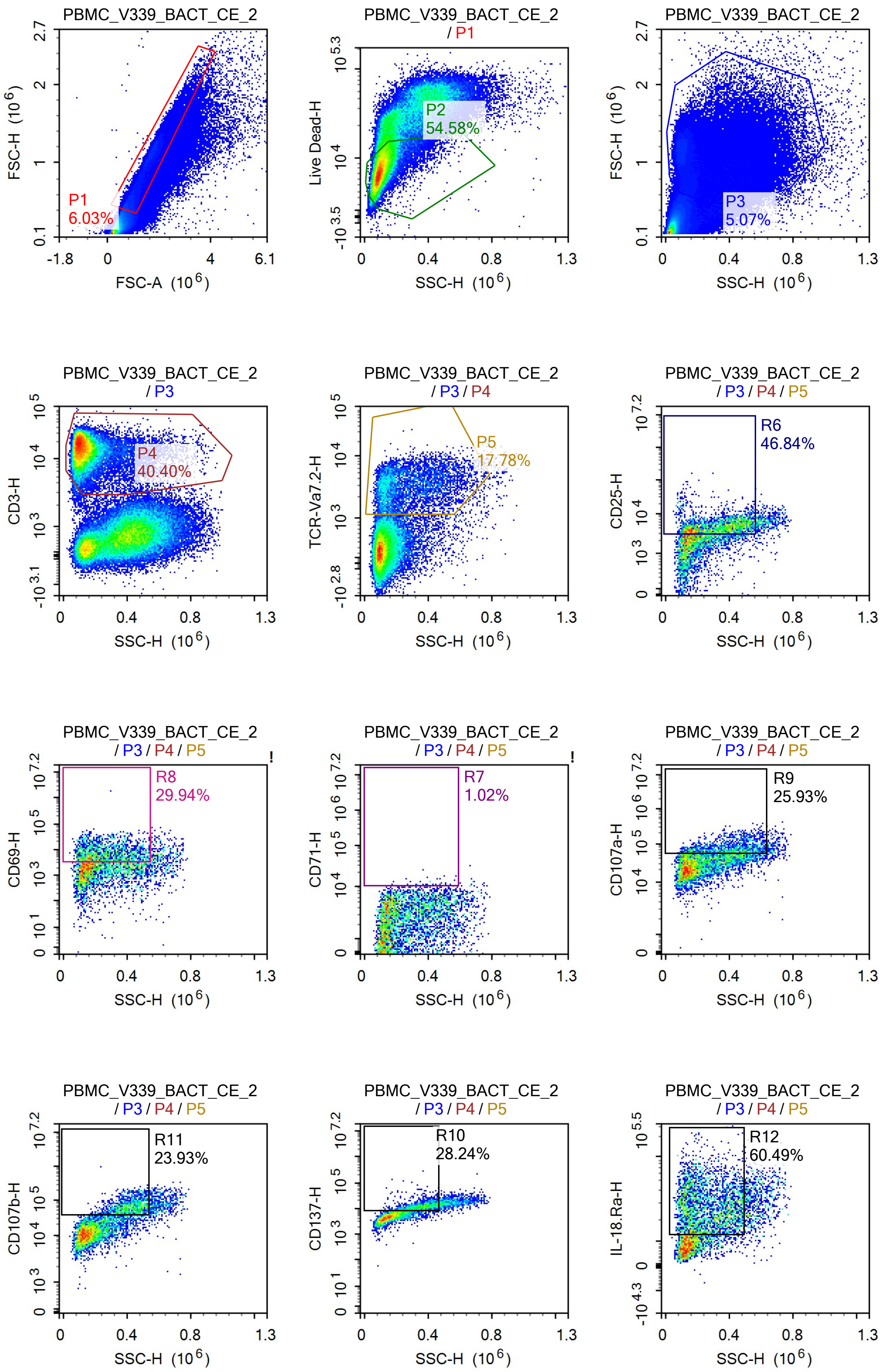
PBMC_V339_BACT_CE_2
P1
6.03%
FSC-H (10^6)
FSC-A (10^6)
PBMC_V339_BACT_CE_2
/ P1
P2
54.58%
Live Dead-H
SSC-H (10^6)
PBMC_V339_BACT_CE_2
P3
5.07%
FSC-H (10^6)
SSC-H (10^6)
PBMC_V339_BACT_CE_2
/ P3
P4
40.40%
CD3-H
SSC-H (10^6)
PBMC_V339_BACT_CE_2
/ P3 / P4
P5
17.78%
TCR-Va7.2-H
SSC-H (10^6)
PBMC_V339_BACT_CE_2
/ P3 / P4 / P5
R6
46.84%
CD25-H
SSC-H (10^6)
PBMC_V339_BACT_CE_2
/ P3 / P4 / P5
R8
29.94%
CD69-H
SSC-H (10^6)
PBMC_V339_BACT_CE_2
/ P3 / P4 / P5
R7
1.02%
CD71-H
SSC-H (10^6)
PBMC_V339_BACT_CE_2
/ P3 / P4 / P5
R9
25.93%
CD107a-H
SSC-H (10^6)
PBMC_V339_BACT_CE_2
/ P3 / P4 / P5
R11
23.93%
CD107b-H
SSC-H (10^6)
PBMC_V339_BACT_CE_2
/ P3 / P4 / P5
R10
28.24%
CD137-H
SSC-H (10^6)
PBMC_V339_BACT_CE_2
/ P3 / P4 / P5
R12
60.49%
IL-18.Ra-H
SSC-H (10^6)

Sample Statistics of PBMC_V339_BACT_CE_2

| Gate | Count | % Parent | X | Y | Median X | Median Y |
|---|---|---|---|---|---|---|
| All | 1,533,753 | | | | | |
| P1 | 92,533 | 6.03% | FSC-A | FSC-H | 1,351,387 | 796,287 |
| P2 | 50,506 | 54.58% | SSC-H | Live Dead-H | 98,167 | 7,696 |
| P3 | 77,832 | 5.07% | SSC-H | FSC-H | 231,777 | 1,045,832 |
| P4 | 31,442 | 40.40% | SSC-H | CD3-H | 117,525 | 15,040 |
| P5 | 5,591 | 17.78% | SSC-H | TCR-Va7.2-H | 225,575 | 3,082 |
| R6 | 2,619 | 46.84% | SSC-H | CD25-H | 332,098 | 4,460 |
| R7 | 57 | 1.02% | SSC-H | CD71-H | 376,333 | 11,235 |
| R8 | 1,674 | 29.94% | SSC-H | CD69-H | 246,685 | 7,130 |
| R9 | 1,450 | 25.93% | SSC-H | CD107a-H | 409,305 | 71,663 |
| R10 | 1,579 | 28.24% | SSC-H | CD137-H | 354,273 | 12,013 |
| R11 | 1,338 | 23.93% | SSC-H | CD107b-H | 418,107 | 63,503 |
| R12 | 3,382 | 60.49% | SSC-H | IL-18.Ra-H | 244,154 | 26,637 |

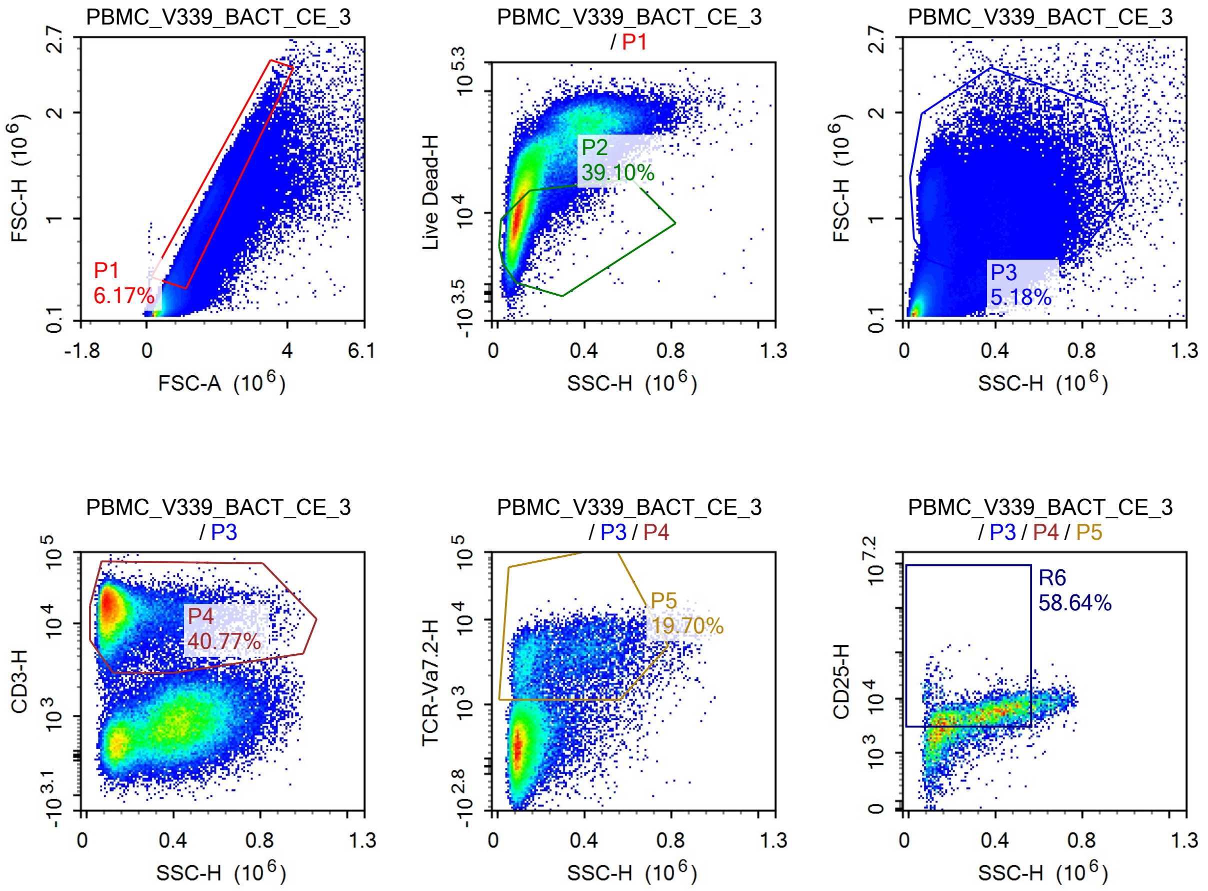

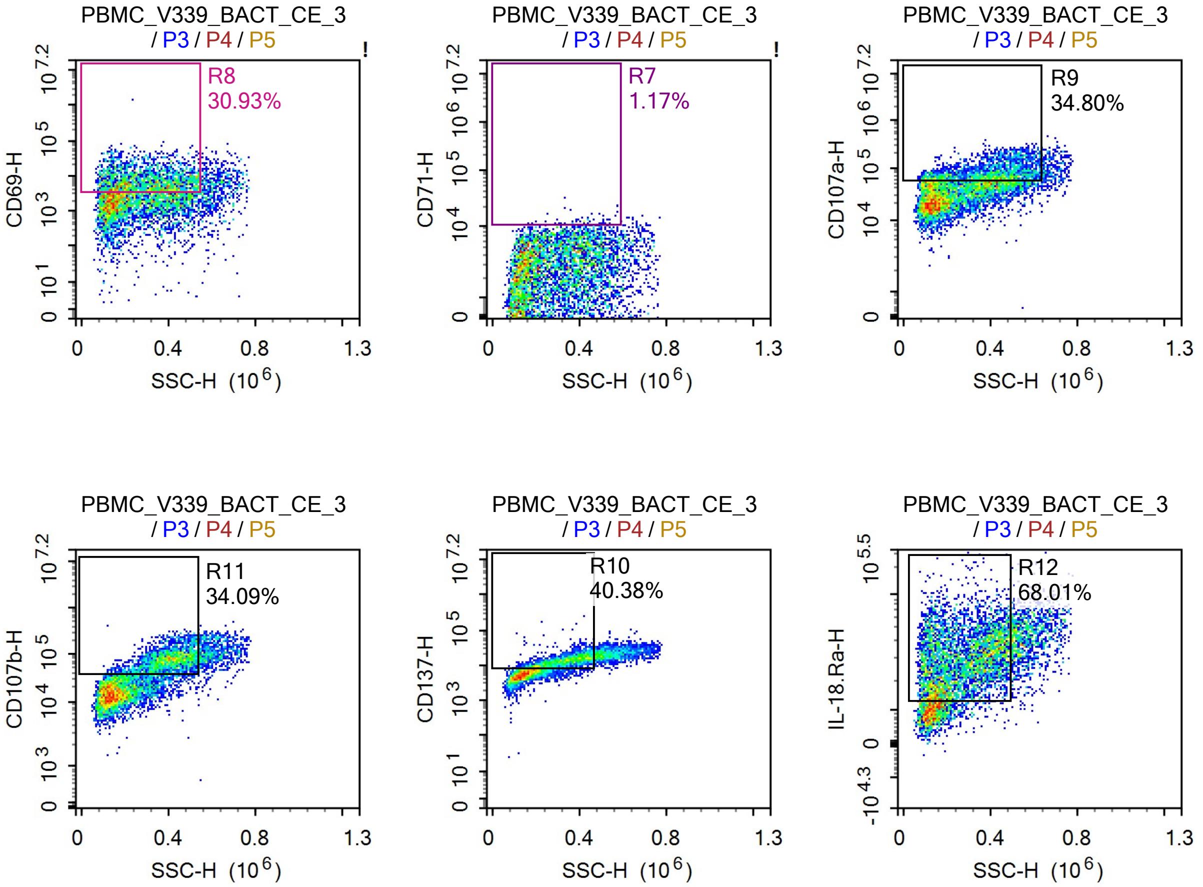


Sample Statistics of PBMC_V339_BACT_CE_3

| Gate | Count | % Parent | X | Y | Median X | Median Y |
|---|---|---|---|---|---|---|
| All | 1,801,968 | | | | | |
| P1 | 111,162 | 6.17% | FSC-A | FSC-H | 1,326,698 | 776,070 |
| P2 | 43,466 | 39.10% | SSC-H | Live Dead-H | 89,122 | 8,572 |
| P3 | 93,294 | 5.18% | SSC-H | FSC-H | 233,273 | 1,064,216 |
| P4 | 38,032 | 40.77% | SSC-H | CD3-H | 114,882 | 15,131 |
| P5 | 7,491 | 19.70% | SSC-H | TCR-Va7.2-H | 270,078 | 3,200 |
| R6 | 4,393 | 58.64% | SSC-H | CD25-H | 344,065 | 5,204 |
| R7 | 88 | 1.17% | SSC-H | CD71-H | 415,938 | 11,190 |
| R8 | 2,317 | 30.93% | SSC-H | CD69-H | 295,434 | 6,945 |
| R9 | 2,607 | 34.80% | SSC-H | CD107a-H | 412,609 | 74,024 |
| R10 | 3,025 | 40.38% | SSC-H | CD137-H | 333,471 | 13,391 |
| R11 | 2,554 | 34.09% | SSC-H | CD107b-H | 405,510 | 75,819 |
| R12 | 5,095 | 68.01% | SSC-H | IL-18.Ra-H | 260,383 | 29,634 |

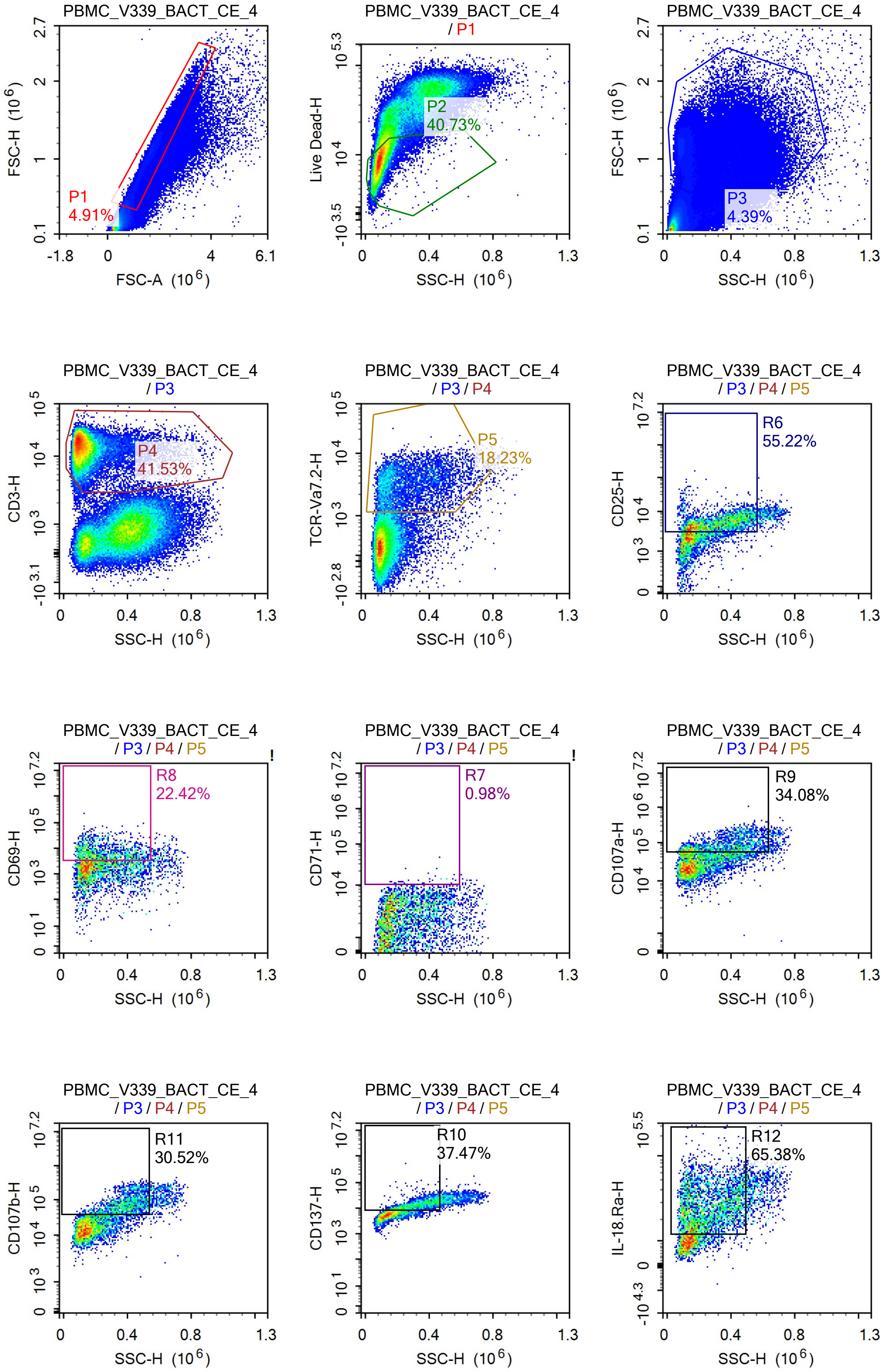
PBMC_V339_BACT_CE_4
FSC-H (10^6)
FSC-A (10^6)
P1
4.91%
PBMC_V339_BACT_CE_4
/ P1
Live Dead-H
SSC-H (10^6)
P2
40.73%
PBMC_V339_BACT_CE_4
FSC-H (10^6)
SSC-H (10^6)
P3
4.39%
PBMC_V339_BACT_CE_4
/ P3
CD3-H
SSC-H (10^6)
P4
41.53%
PBMC_V339_BACT_CE_4
/ P3 / P4
TCR-Va7.2-H
SSC-H (10^6)
P5
18.23%
PBMC_V339_BACT_CE_4
/ P3 / P4 / P5
CD25-H
SSC-H (10^6)
R6
55.22%
PBMC_V339_BACT_CE_4
/ P3 / P4 / P5
CD69-H
SSC-H (10^6)
R8
22.42%
PBMC_V339_BACT_CE_4
/ P3 / P4 / P5
CD71-H
SSC-H (10^6)
R7
0.98%
PBMC_V339_BACT_CE_4
/ P3 / P4 / P5
CD107a-H
SSC-H (10^6)
R9
34.08%
PBMC_V339_BACT_CE_4
/ P3 / P4 / P5
CD107b-H
SSC-H (10^6)
R11
30.52%
PBMC_V339_BACT_CE_4
/ P3 / P4 / P5
CD137-H
SSC-H (10^6)
R10
37.47%
PBMC_V339_BACT_CE_4
/ P3 / P4 / P5
IL-18.Ra-H
SSC-H (10^6)
R12
65.38%

Sample Statistics of PBMC_V339_BACT_CE_4

| Gate | Count | % Parent | X | Y | Median X | Median Y |
|---|---|---|---|---|---|---|
| All | 1,724,278 | | | | | |
| P1 | 84,610 | 4.91% | FSC-A | FSC-H | 1,473,168 | 888,786 |
| P2 | 34,465 | 40.73% | SSC-H | Live Dead-H | 88,464 | 8,743 |
| P3 | 75,669 | 4.39% | SSC-H | FSC-H | 232,694 | 1,047,606 |
| P4 | 31,424 | 41.53% | SSC-H | CD3-H | 111,195 | 14,804 |
| P5 | 5,728 | 18.23% | SSC-H | TCR-Va7.2-H | 222,247 | 3,096 |
| R6 | 3,163 | 55.22% | SSC-H | CD25-H | 308,761 | 4,972 |
| R7 | 56 | 0.98% | SSC-H | CD71-H | 358,526 | 11,178 |
| R8 | 1,284 | 22.42% | SSC-H | CD69-H | 232,303 | 6,422 |
| R9 | 1,952 | 34.08% | SSC-H | CD107a-H | 384,904 | 77,294 |
| R10 | 2,146 | 37.47% | SSC-H | CD137-H | 320,177 | 13,044 |
| R11 | 1,748 | 30.52% | SSC-H | CD107b-H | 398,937 | 74,273 |
| R12 | 3,745 | 65.38% | SSC-H | IL-18.Ra-H | 236,038 | 26,799 |
