## Supplementary material for "Vitamin B_2_ Production by Vaginal Lactobacilli Promotes Symbiosis": Data S10: MR1 expression on VK2 E6E7 (attempt 1, one gating strategy).pdf

### Report of Specimen8

Specimen Name: Specimen8 | Run Time: 28-Feb-25 1:37 PM

Cytometer: NovoCyte Quanteon 621181110427 | Software: NovoExpress 1.6.0

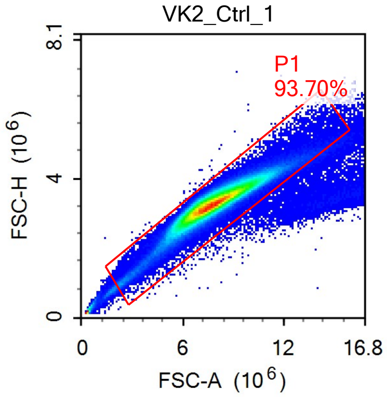


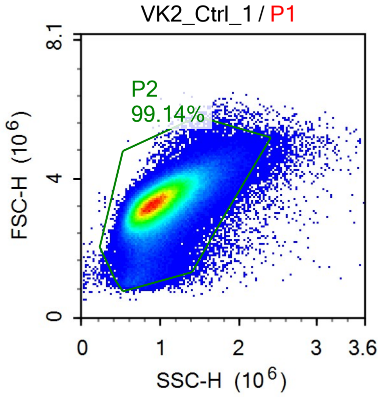


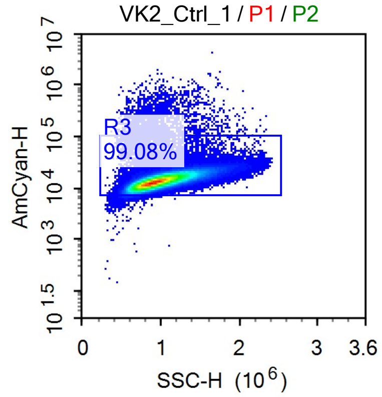


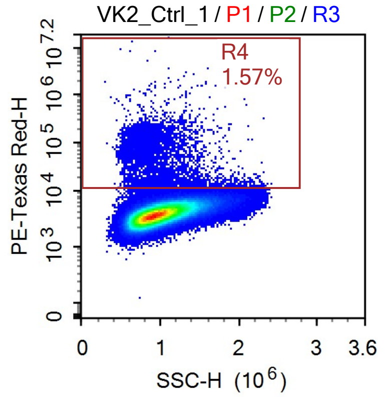

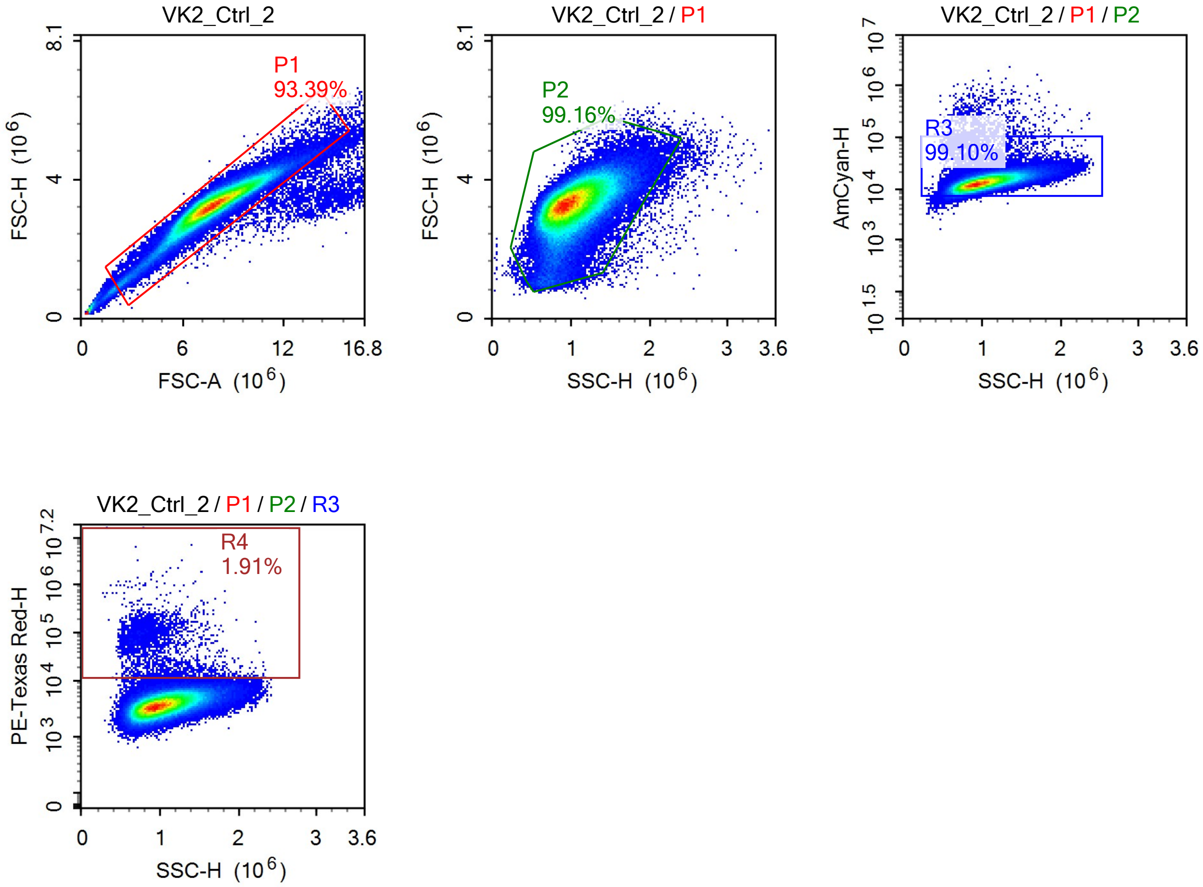
VK2_Ctrl_2
P1
93.39%
FSC-H (10^6)
FSC-A (10^6)
VK2_Ctrl_2 / P1
P2
99.16%
FSC-H (10^6)
SSC-H (10^6)
VK2_Ctrl_2 / P1 / P2
R3
99.10%
AmCyan-H
SSC-H (10^6)
VK2_Ctrl_2 / P1 / P2 / R3
R4
1.91%
PE-Texas Red-H
SSC-H (10^6)

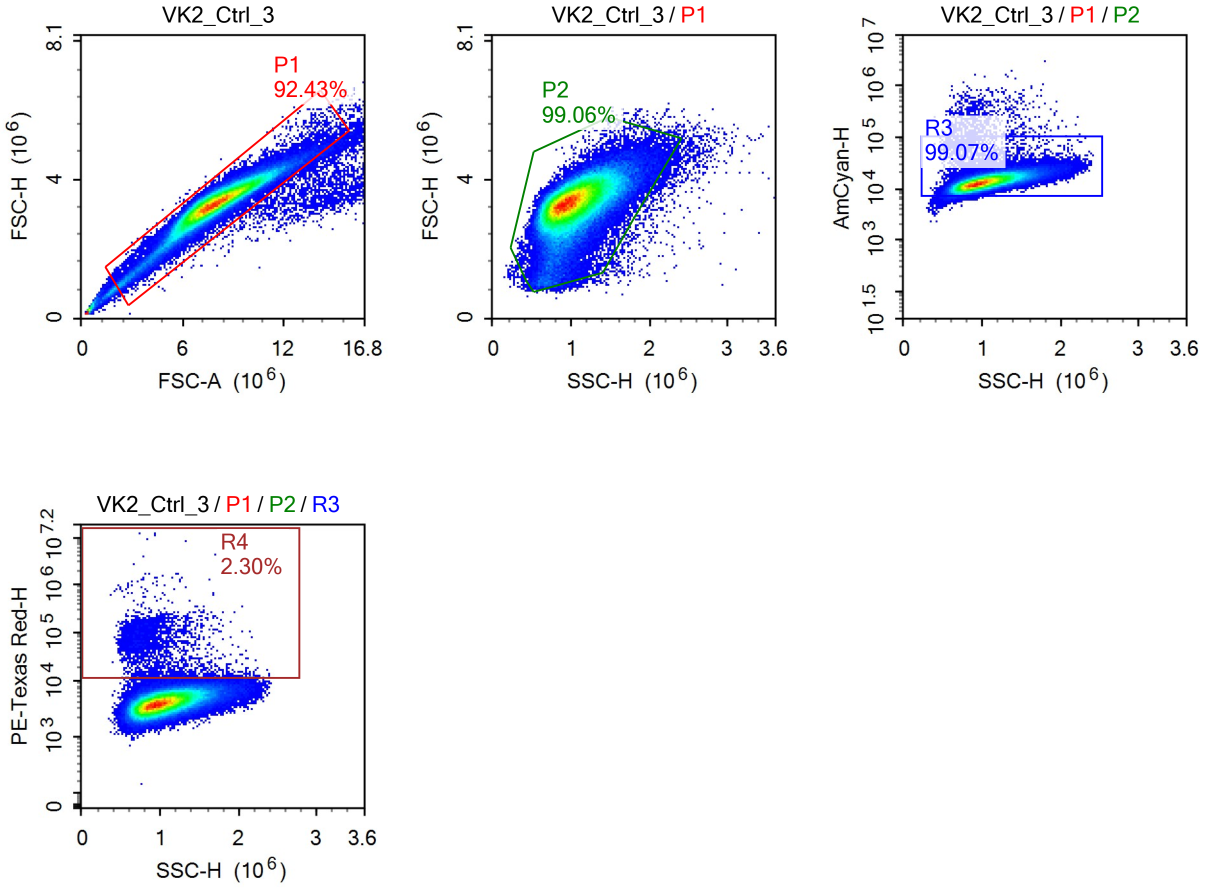
VK2_Ctrl_3
P1
92.43%
FSC-H (10^6)
FSC-A (10^6)
VK2_Ctrl_3 / P1
P2
99.06%
FSC-H (10^6)
SSC-H (10^6)
VK2_Ctrl_3 / P1 / P2
R3
99.07%
AmCyan-H
SSC-H (10^6)
VK2_Ctrl_3 / P1 / P2 / R3
R4
2.30%
PE-Texas Red-H
SSC-H (10^6)

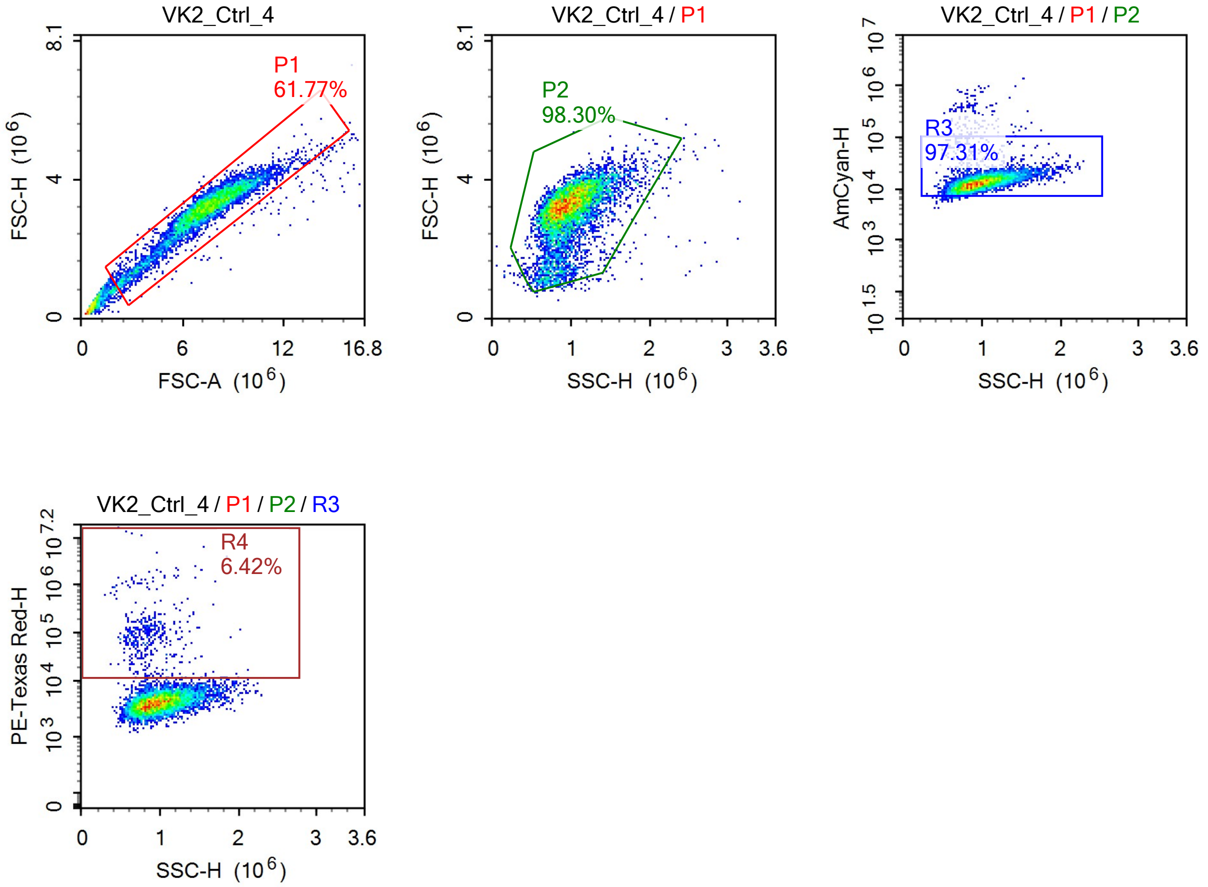
VK2_Ctrl_4
P1
61.77%
FSC-H (10^6)
FSC-A (10^6)
VK2_Ctrl_4 / P1
P2
98.30%
SSC-H (10^6)
VK2_Ctrl_4 / P1 / P2
R3
97.31%
AmCyan-H
VK2_Ctrl_4 / P1 / P2 / R3
R4
6.42%
PE-Texas Red-H

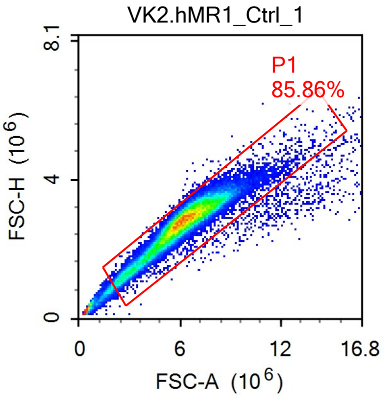
VK2.hMR1_Ctrl_1
P1
85.86%
FSC-H (10^6)
FSC-A (10^6)
0
4
8.1
6
12
16.8

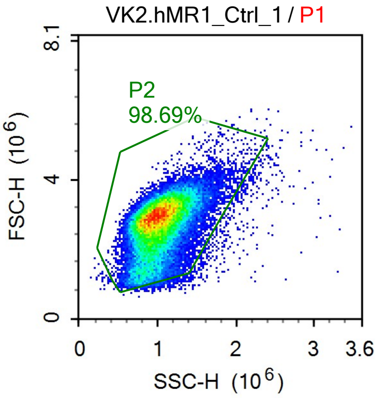
VK2.hMR1_Ctrl_1 / P1
P2
98.69%
FSC-H (10^6)
SSC-H (10^6)
0
4
8.1
1
2
3
3.6

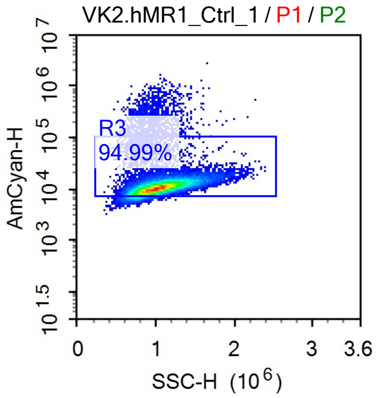
VK2.hMR1_Ctrl_1 / P1 / P2
R3
94.99%
AmCyan-H
SSC-H (10^6)
10^1.5
10^3
10^4
10^5
10^6
10^7
0
1
2
3
3.6

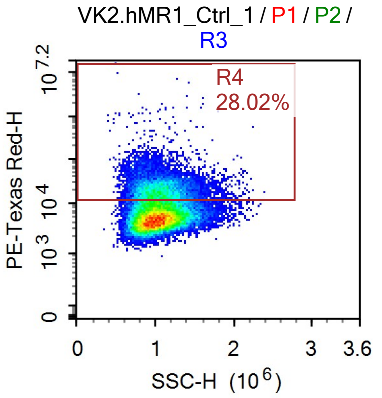
VK2.hMR1_Ctrl_1 / P1 / P2 / R3
R4
28.02%
PE-Texas Red-H
SSC-H (10^6)
0
10^3
10^4
10^7.2
1
2
3
3.6

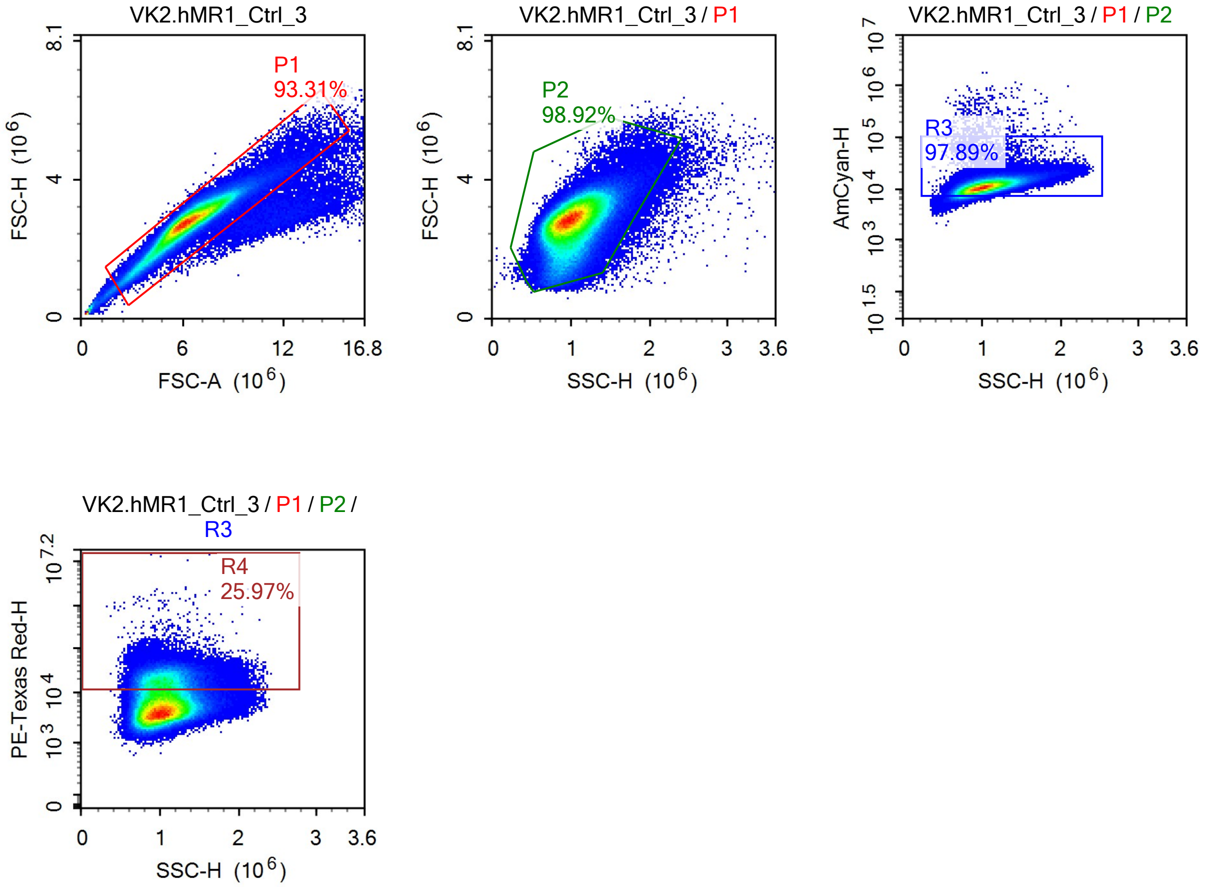
VK2.hMR1_Ctrl_3
P1
93.31%
FSC-H (10^6)
FSC-A (10^6)
VK2.hMR1_Ctrl_3 / P1
P2
98.92%
FSC-H (10^6)
SSC-H (10^6)
VK2.hMR1_Ctrl_3 / P1 / P2
R3
97.89%
AmCyan-H
SSC-H (10^6)
VK2.hMR1_Ctrl_3 / P1 / P2 / R3
R4
25.97%
PE-Texas Red-H
SSC-H (10^6)

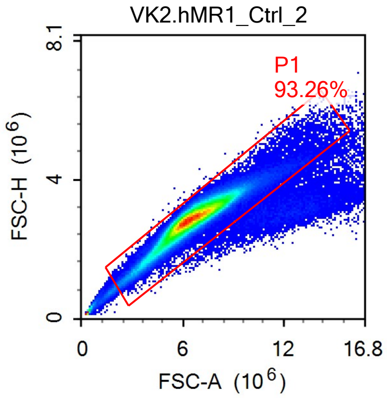
VK2.hMR1_Ctrl_2
P1
93.26%
FSC-H (10^6)
FSC-A (10^6)
0
4
8.1
0
6
12
16.8

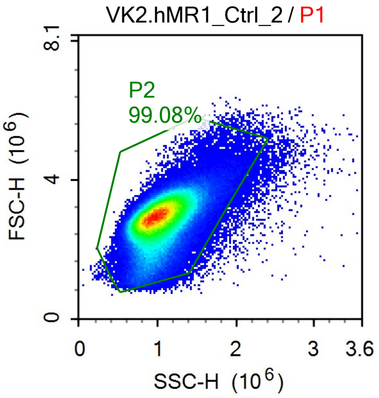
VK2.hMR1_Ctrl_2 / P1
P2
99.08%
FSC-H (10^6)
SSC-H (10^6)
0
4
8.1
0
1
2
3
3.6

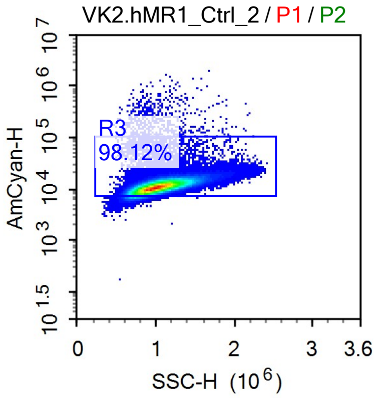
VK2.hMR1_Ctrl_2 / P1 / P2
R3
98.12%
AmCyan-H
10^1.5
10^3
10^4
10^5
10^6
10^7
SSC-H (10^6)
0
1
2
3
3.6

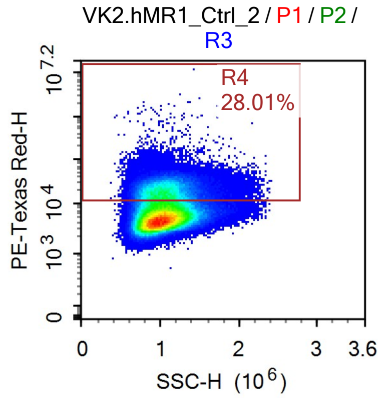
VK2.hMR1_Ctrl_2 / P1 / P2 / R3
R4
28.01%
PE-Texas Red-H
0
10^3
10^4
10^7.2
SSC-H (10^6)
0
1
2
3
3.6

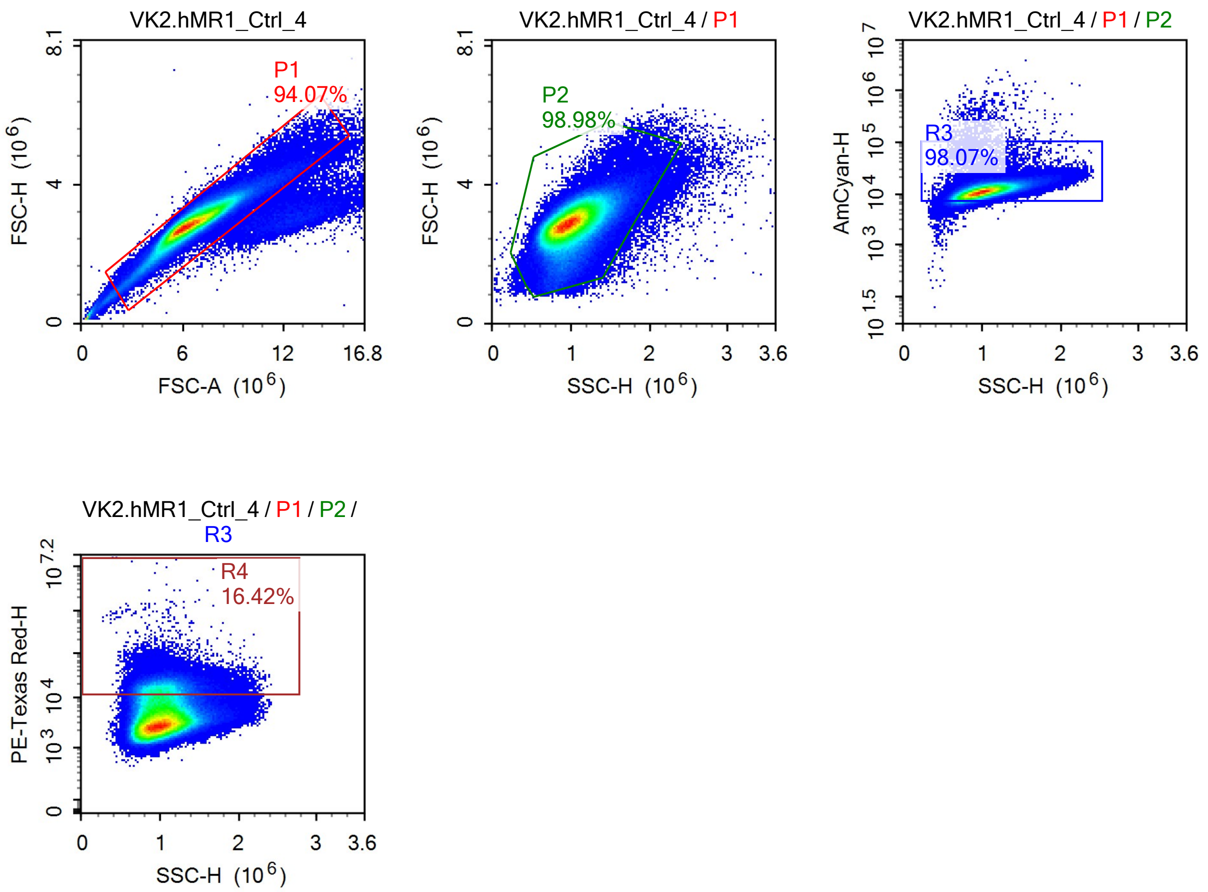
VK2.hMR1_Ctrl_4
P1
94.07%
FSC-H (10^6)
FSC-A (10^6)
VK2.hMR1_Ctrl_4 / P1
P2
98.98%
FSC-H (10^6)
SSC-H (10^6)
VK2.hMR1_Ctrl_4 / P1 / P2
R3
98.07%
AmCyan-H
SSC-H (10^6)
VK2.hMR1_Ctrl_4 / P1 / P2 / R3
R4
16.42%
PE-Texas Red-H
SSC-H (10^6)

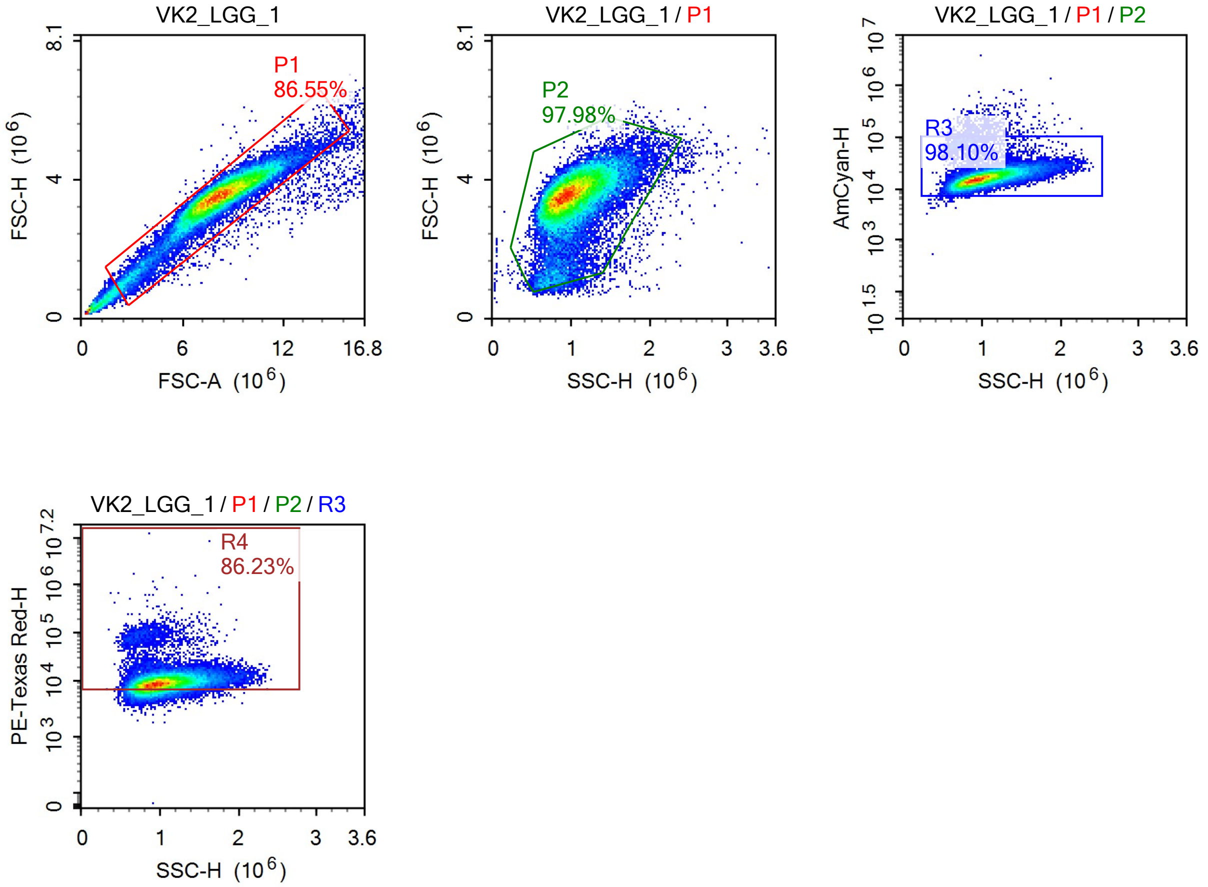
VK2_LGG_1
P1
86.55%
FSC-H (10^6)
FSC-A (10^6)
VK2_LGG_1 / P1
P2
97.98%
SSC-H (10^6)
VK2_LGG_1 / P1 / P2
R3
98.10%
AmCyan-H
VK2_LGG_1 / P1 / P2 / R3
R4
86.23%
PE-Texas Red-H

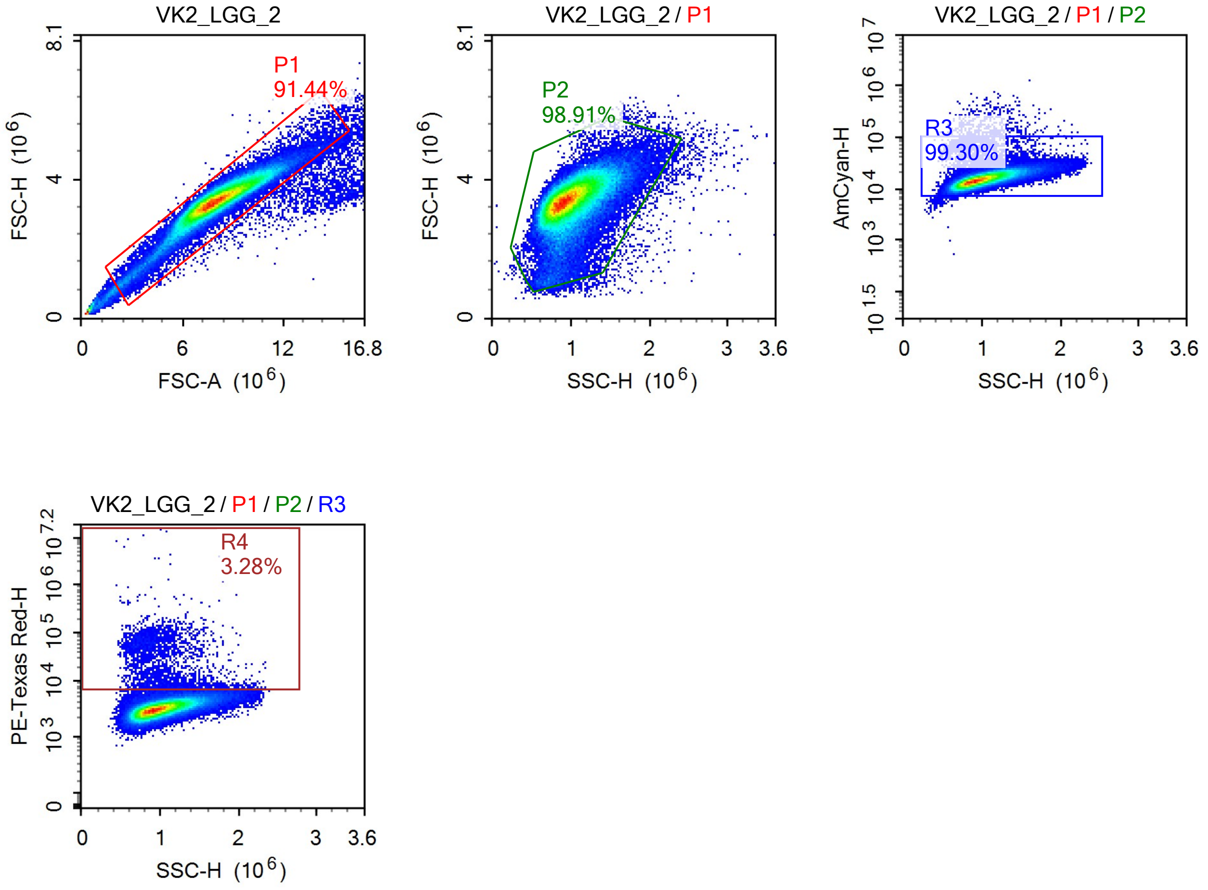

VK2_LGG_2
P1
91.44%
FSC-H (10^6)
FSC-A (10^6)
VK2_LGG_2 / P1
P2
98.91%
FSC-H (10^6)
SSC-H (10^6)
VK2_LGG_2 / P1 / P2
R3
99.30%
AmCyan-H
SSC-H (10^6)
VK2_LGG_2 / P1 / P2 / R3
R4
3.28%
PE-Texas Red-H
SSC-H (10^6)

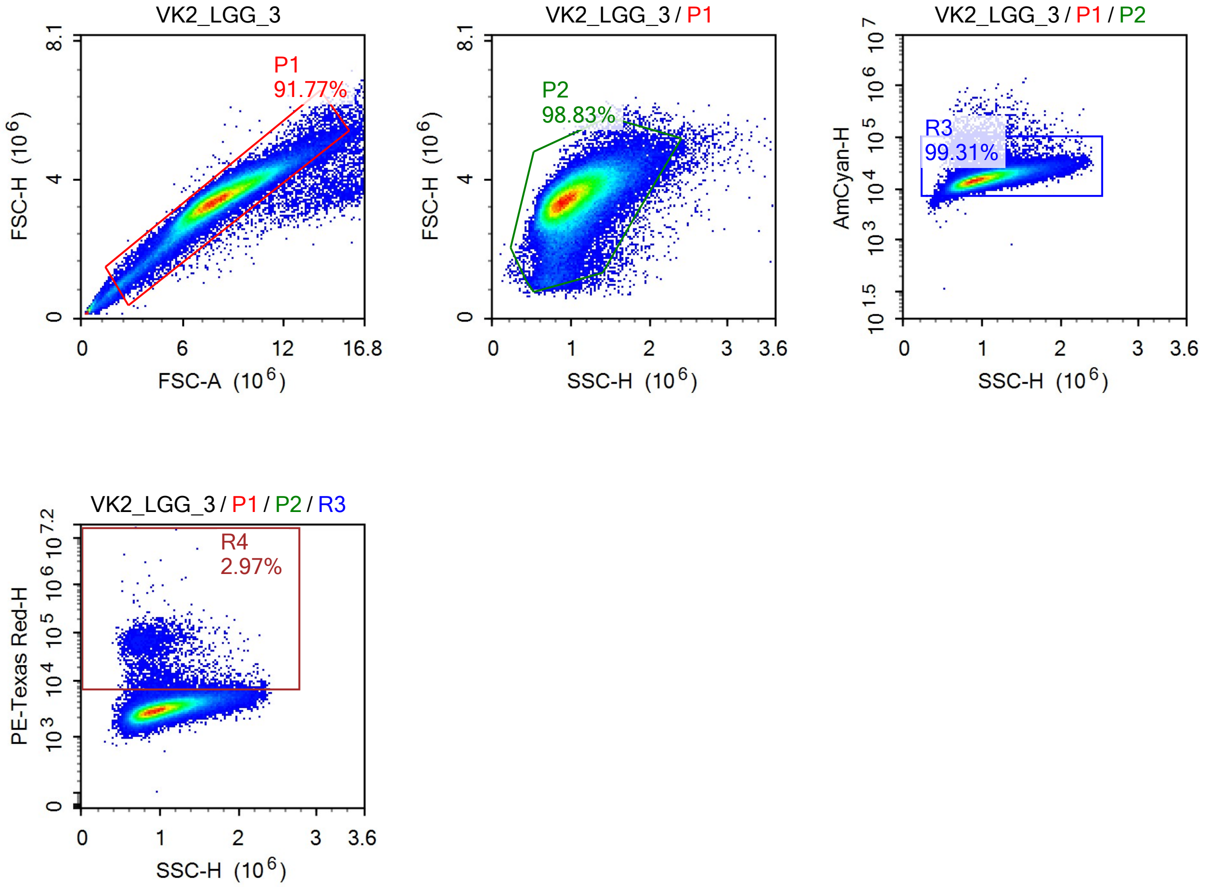
VK2_LGG_3
P1
91.77%
FSC-H (10^6)
FSC-A (10^6)
VK2_LGG_3 / P1
P2
98.83%
FSC-H (10^6)
SSC-H (10^6)
VK2_LGG_3 / P1 / P2
R3
99.31%
AmCyan-H
SSC-H (10^6)
VK2_LGG_3 / P1 / P2 / R3
R4
2.97%
PE-Texas Red-H
SSC-H (10^6)

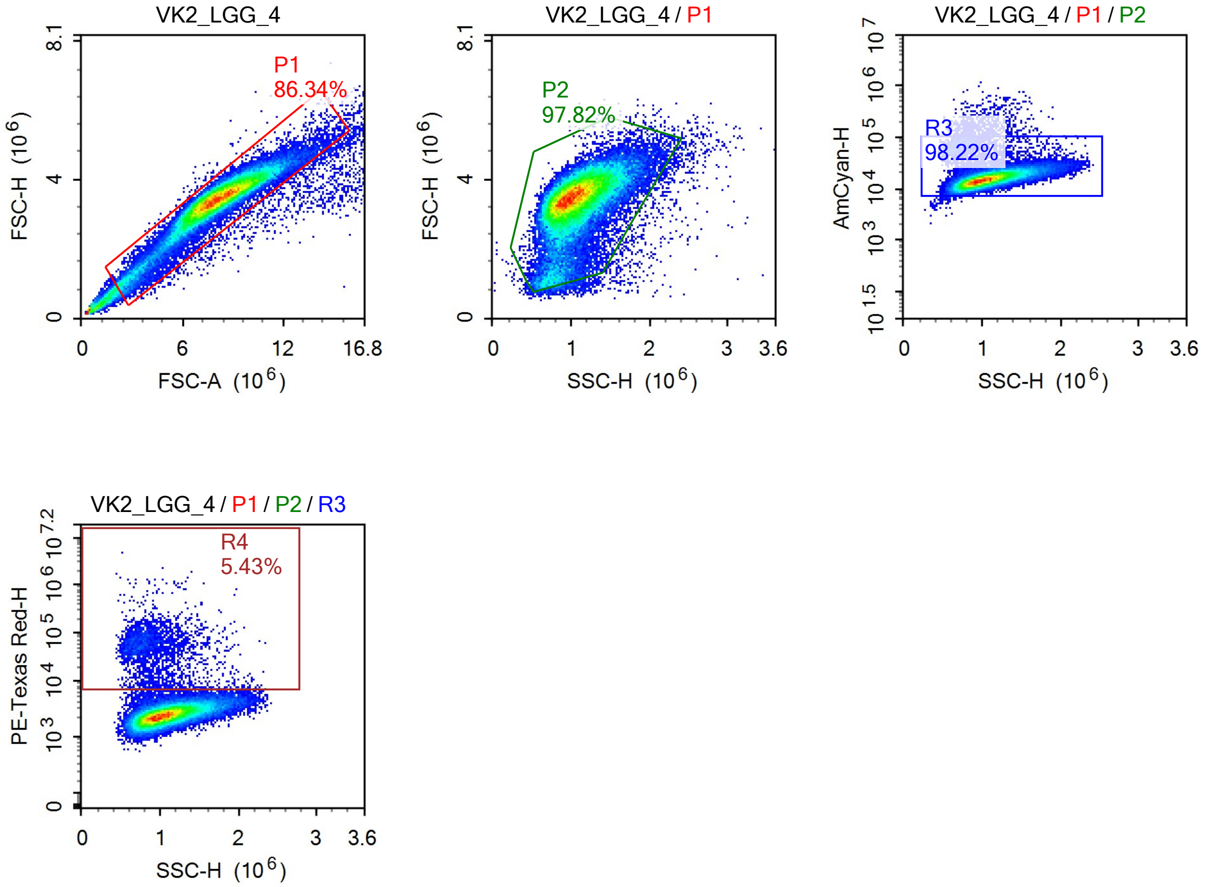
VK2_LGG_4
P1
86.34%
FSC-H (10^6)
FSC-A (10^6)
VK2_LGG_4 / P1
P2
97.82%
FSC-H (10^6)
SSC-H (10^6)
VK2_LGG_4 / P1 / P2
R3
98.22%
AmCyan-H
SSC-H (10^6)
VK2_LGG_4 / P1 / P2 / R3
R4
5.43%
PE-Texas Red-H
SSC-H (10^6)

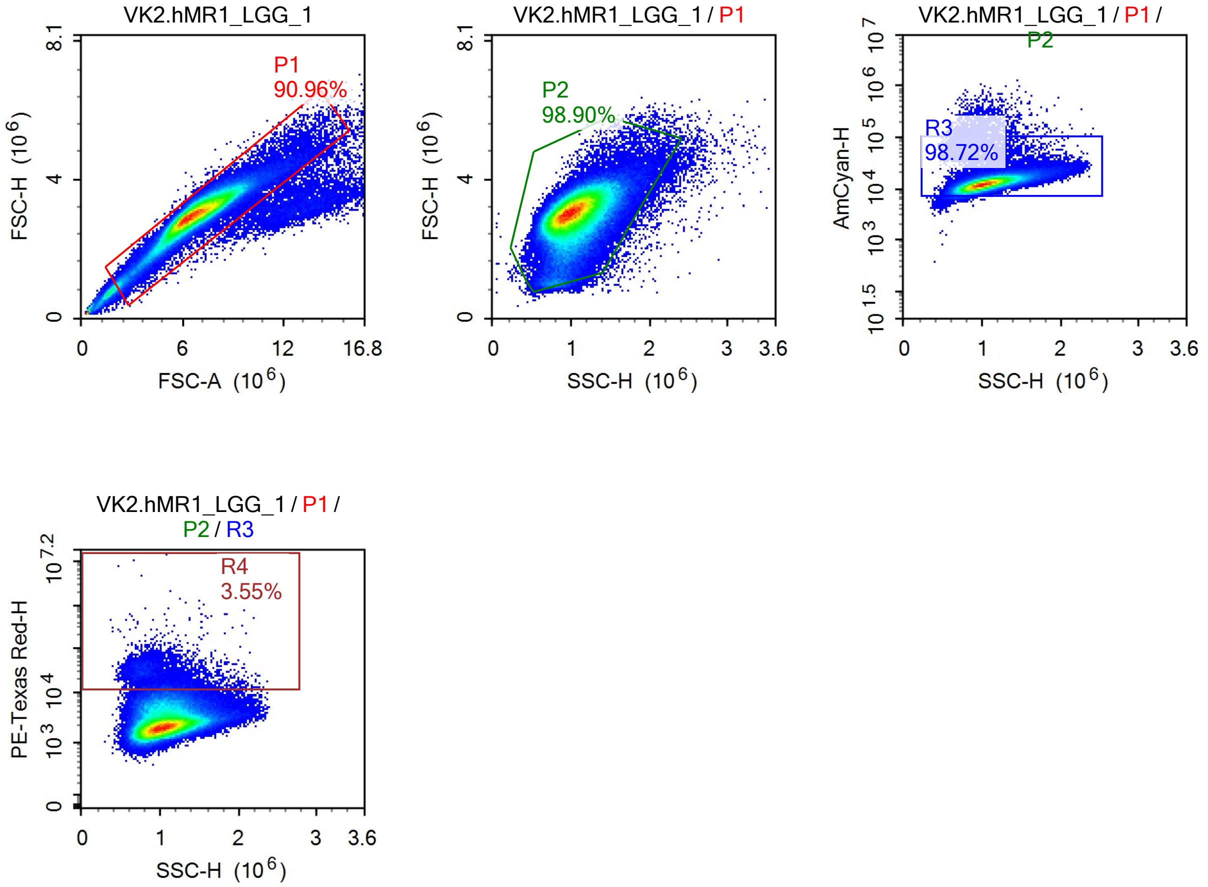
VK2.hMR1_LGG_1
P1
90.96%
FSC-H (10^6)
FSC-A (10^6)
0
4
8.1
0
6
12
16.8
VK2.hMR1_LGG_1 / P1
P2
98.90%
FSC-H (10^6)
SSC-H (10^6)
0
1
2
3
3.6
VK2.hMR1_LGG_1 / P1 / P2
R3
98.72%
AmCyan-H
10^1.5
10^3
10^4
10^5
10^6
10^7
SSC-H (10^6)
VK2.hMR1_LGG_1 / P1 / P2 / R3
R4
3.55%
PE-Texas Red-H
10^3
10^4
10^7.2
SSC-H (10^6)

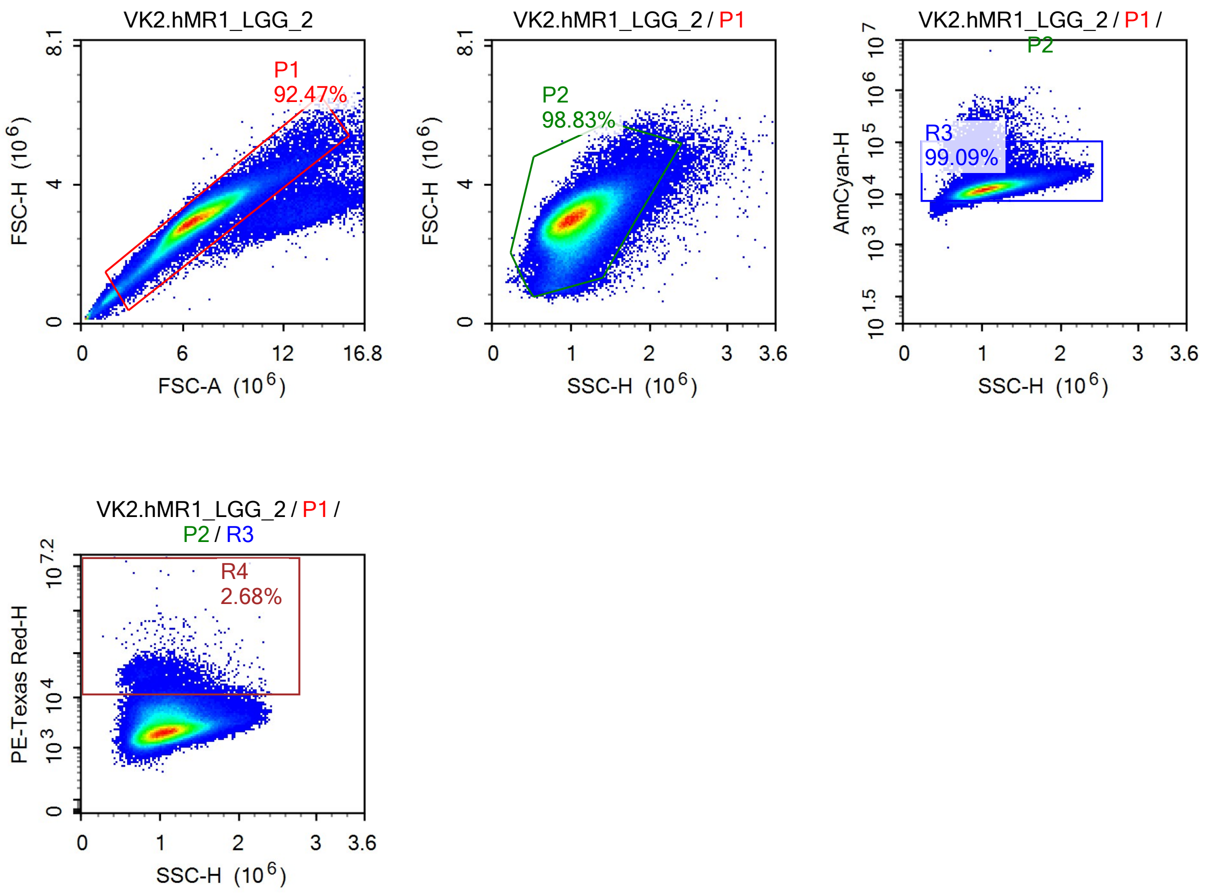
VK2.hMR1_LGG_2
P1
92.47%
FSC-H (10^6)
FSC-A (10^6)
VK2.hMR1_LGG_2 / P1
P2
98.83%
FSC-H (10^6)
SSC-H (10^6)
VK2.hMR1_LGG_2 / P1 / P2
R3
99.09%
AmCyan-H
SSC-H (10^6)
VK2.hMR1_LGG_2 / P1 / P2 / R3
R4
2.68%
PE-Texas Red-H
SSC-H (10^6)

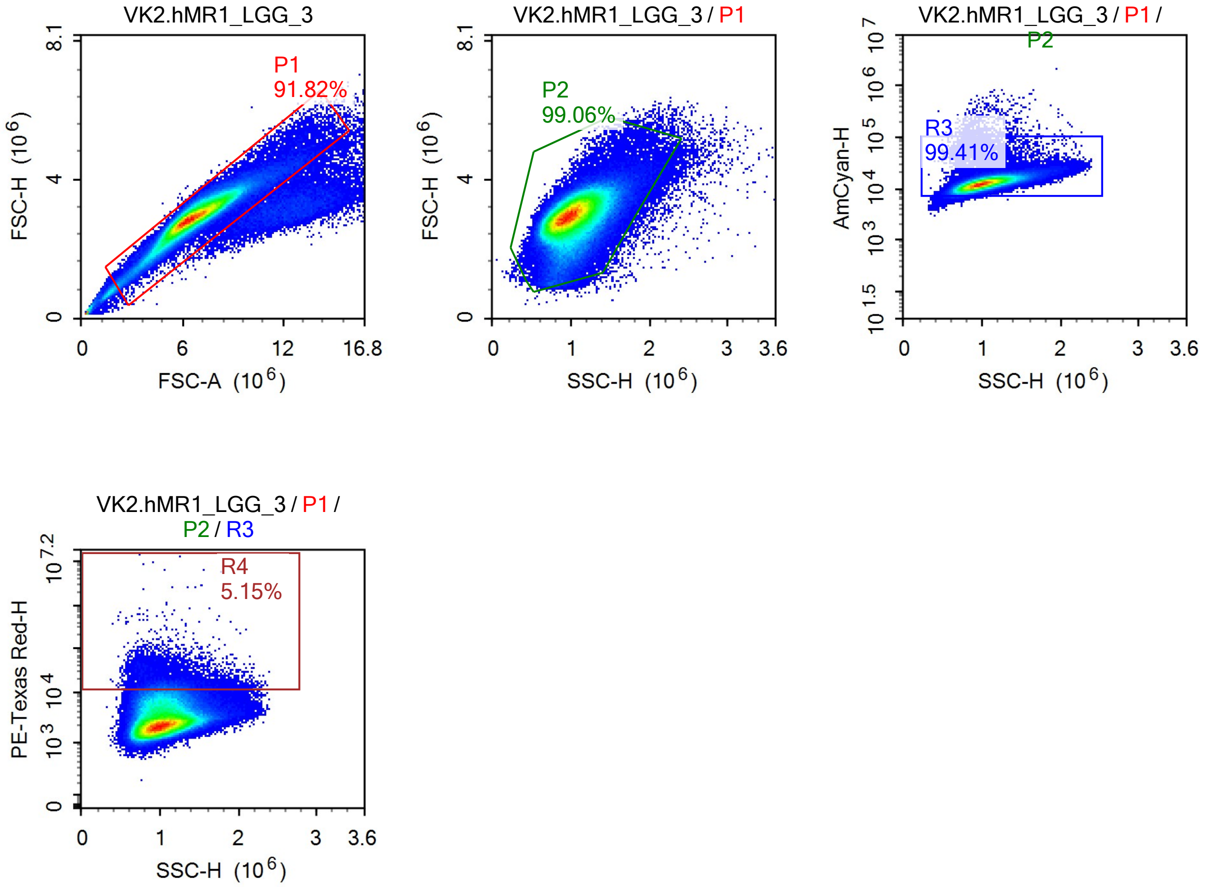
VK2.hMR1_LGG_3
P1
91.82%
FSC-H (10^6)
FSC-A (10^6)
VK2.hMR1_LGG_3 / P1
P2
99.06%
FSC-H (10^6)
SSC-H (10^6)
VK2.hMR1_LGG_3 / P1 / P2
R3
99.41%
AmCyan-H
SSC-H (10^6)
VK2.hMR1_LGG_3 / P1 / P2 / R3
R4
5.15%
PE-Texas Red-H
SSC-H (10^6)

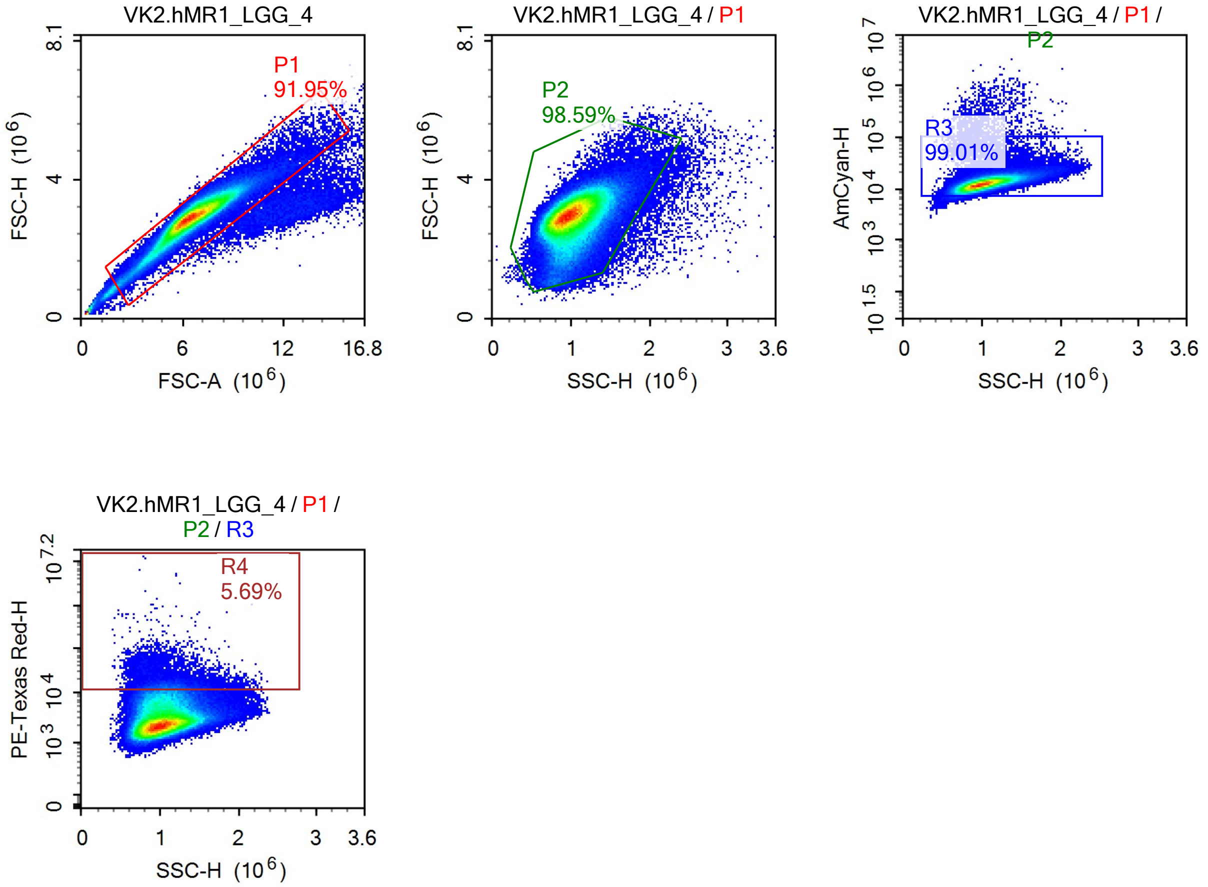
VK2.hMR1_LGG_4
P1
91.95%
FSC-H (10^6)
FSC-A (10^6)
VK2.hMR1_LGG_4 / P1
P2
98.59%
FSC-H (10^6)
SSC-H (10^6)
VK2.hMR1_LGG_4 / P1 / P2
R3
99.01%
AmCyan-H
SSC-H (10^6)
VK2.hMR1_LGG_4 / P1 / P2 / R3
R4
5.69%
PE-Texas Red-H
SSC-H (10^6)

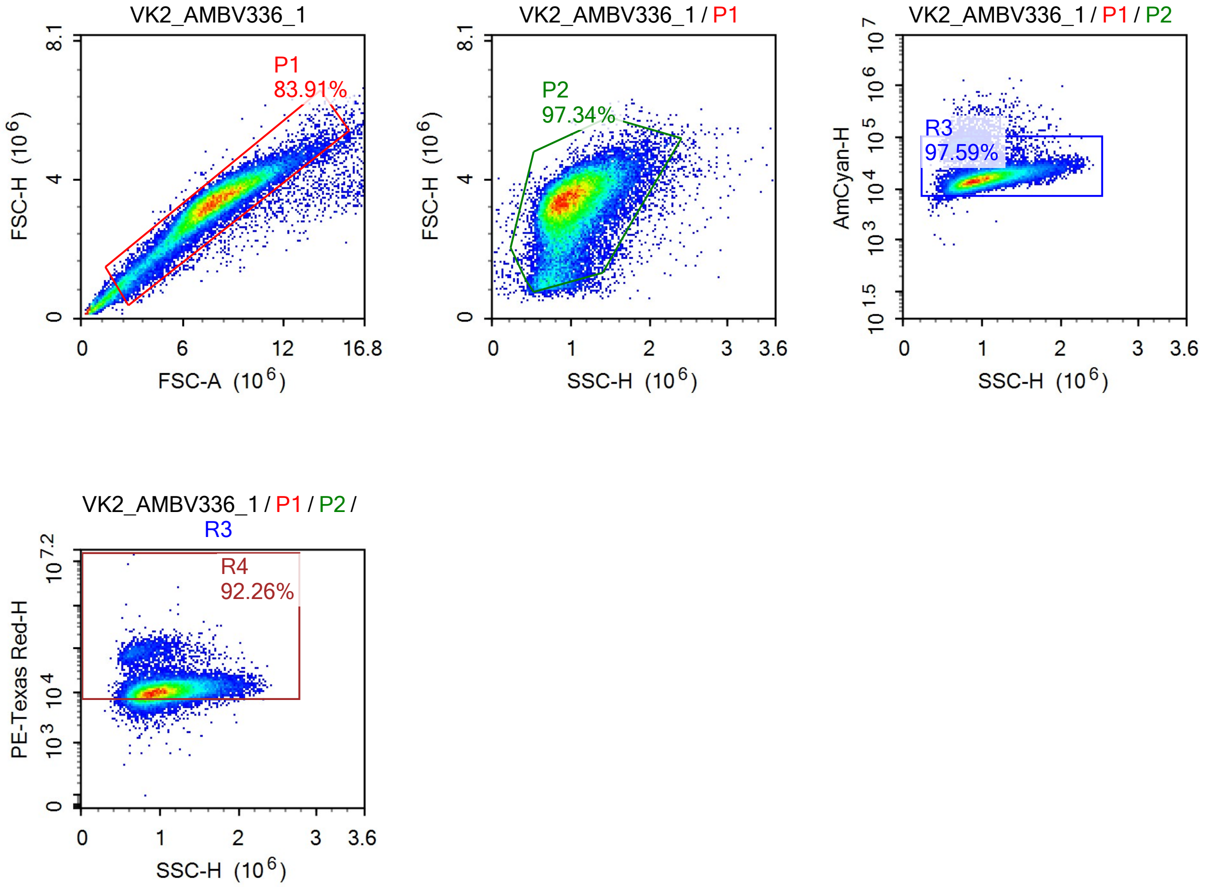
VK2_AMBV336_1
P1
83.91%
FSC-H (10^6)
FSC-A (10^6)
VK2_AMBV336_1 / P1
P2
97.34%
FSC-H (10^6)
SSC-H (10^6)
VK2_AMBV336_1 / P1 / P2
R3
97.59%
AmCyan-H
SSC-H (10^6)
VK2_AMBV336_1 / P1 / P2 / R3
R4
92.26%
PE-Texas Red-H
SSC-H (10^6)

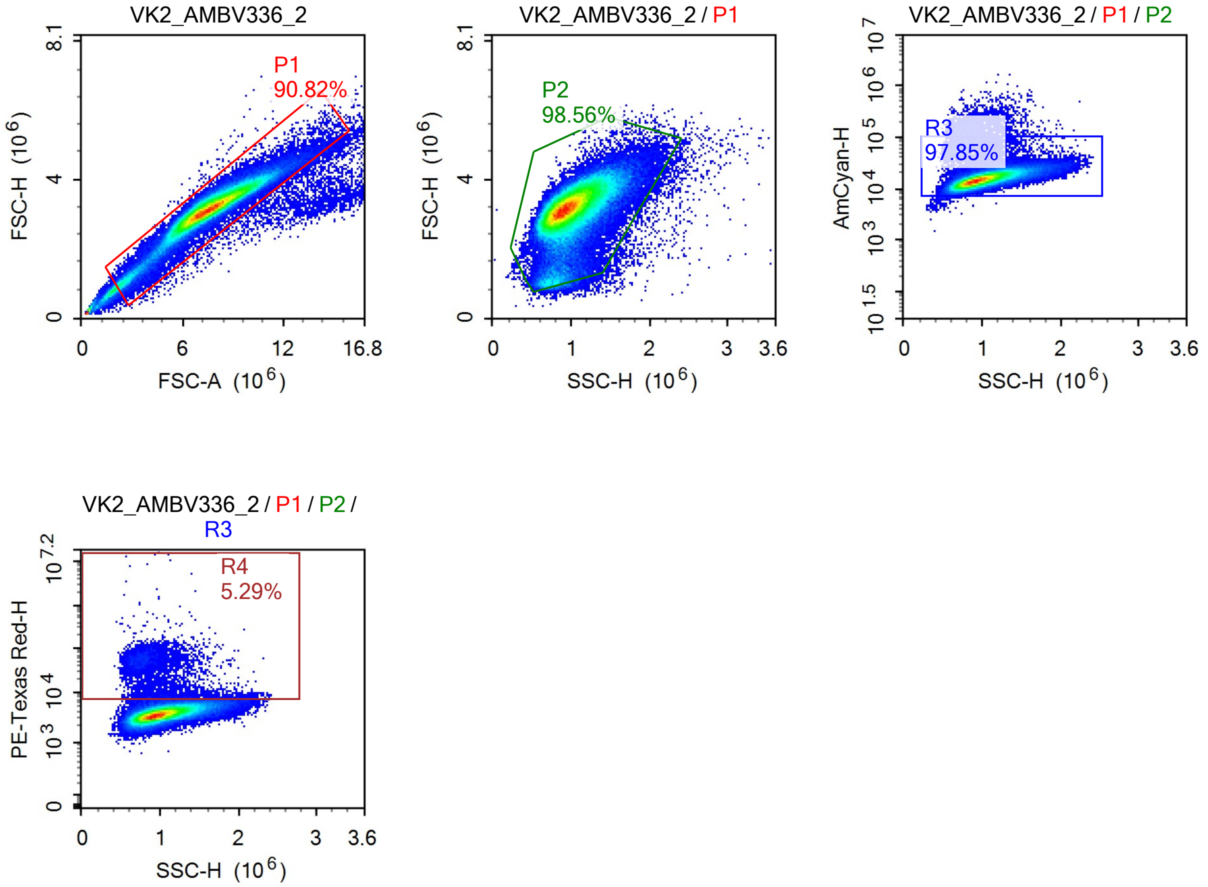
VK2_AMBV336_2
P1
90.82%
FSC-H (10^6)
FSC-A (10^6)
VK2_AMBV336_2 / P1
P2
98.56%
FSC-H (10^6)
SSC-H (10^6)
VK2_AMBV336_2 / P1 / P2
R3
97.85%
AmCyan-H
SSC-H (10^6)
VK2_AMBV336_2 / P1 / P2 / R3
R4
5.29%
PE-Texas Red-H
SSC-H (10^6)

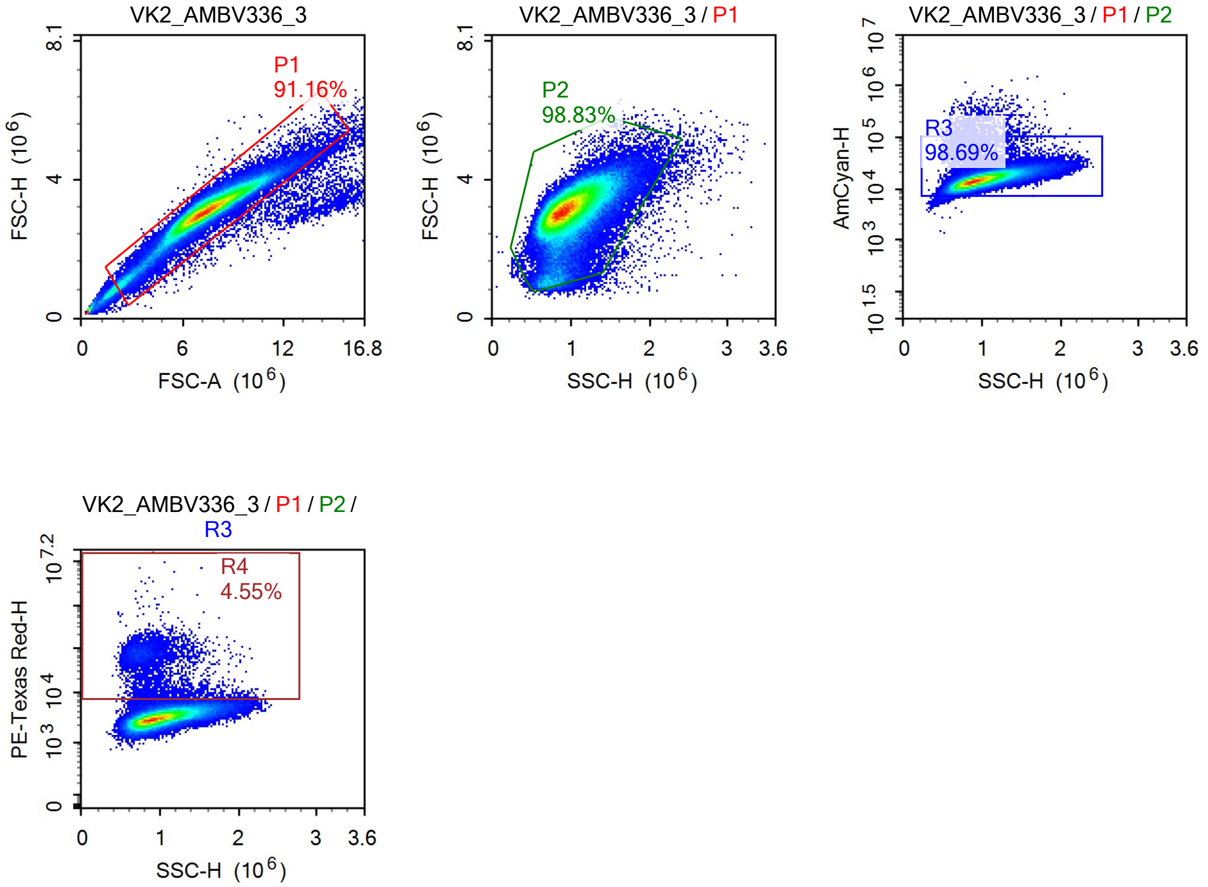
VK2_AMBV336_3
P1
91.16%
FSC-H (10^6)
FSC-A (10^6)
VK2_AMBV336_3 / P1
P2
98.83%
FSC-H (10^6)
SSC-H (10^6)
VK2_AMBV336_3 / P1 / P2
R3
98.69%
AmCyan-H
SSC-H (10^6)
VK2_AMBV336_3 / P1 / P2 / R3
R4
4.55%
PE-Texas Red-H
SSC-H (10^6)

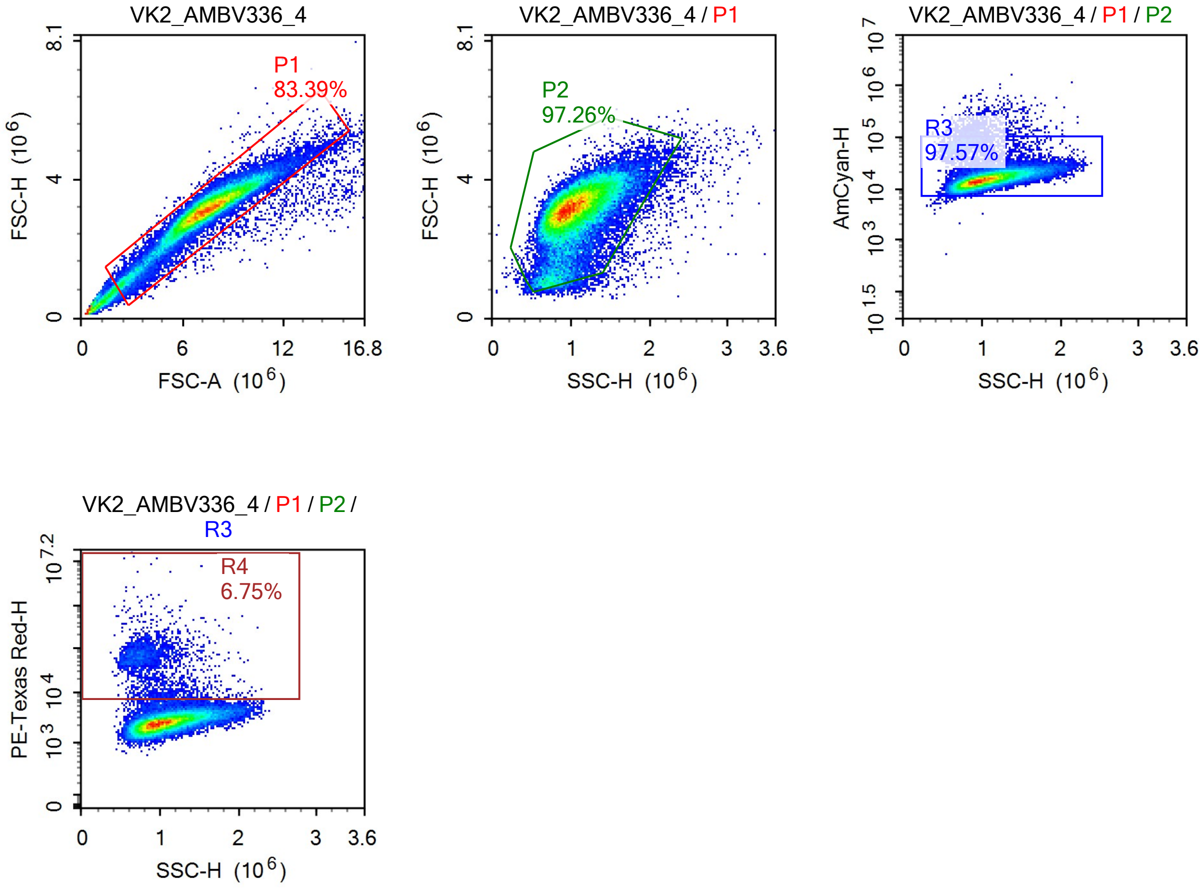
VK2_AMBV336_4
P1
83.39%
FSC-H (10^6)
FSC-A (10^6)
VK2_AMBV336_4 / P1
P2
97.26%
FSC-H (10^6)
SSC-H (10^6)
VK2_AMBV336_4 / P1 / P2
R3
97.57%
AmCyan-H
SSC-H (10^6)
VK2_AMBV336_4 / P1 / P2 / R3
R4
6.75%
PE-Texas Red-H
SSC-H (10^6)

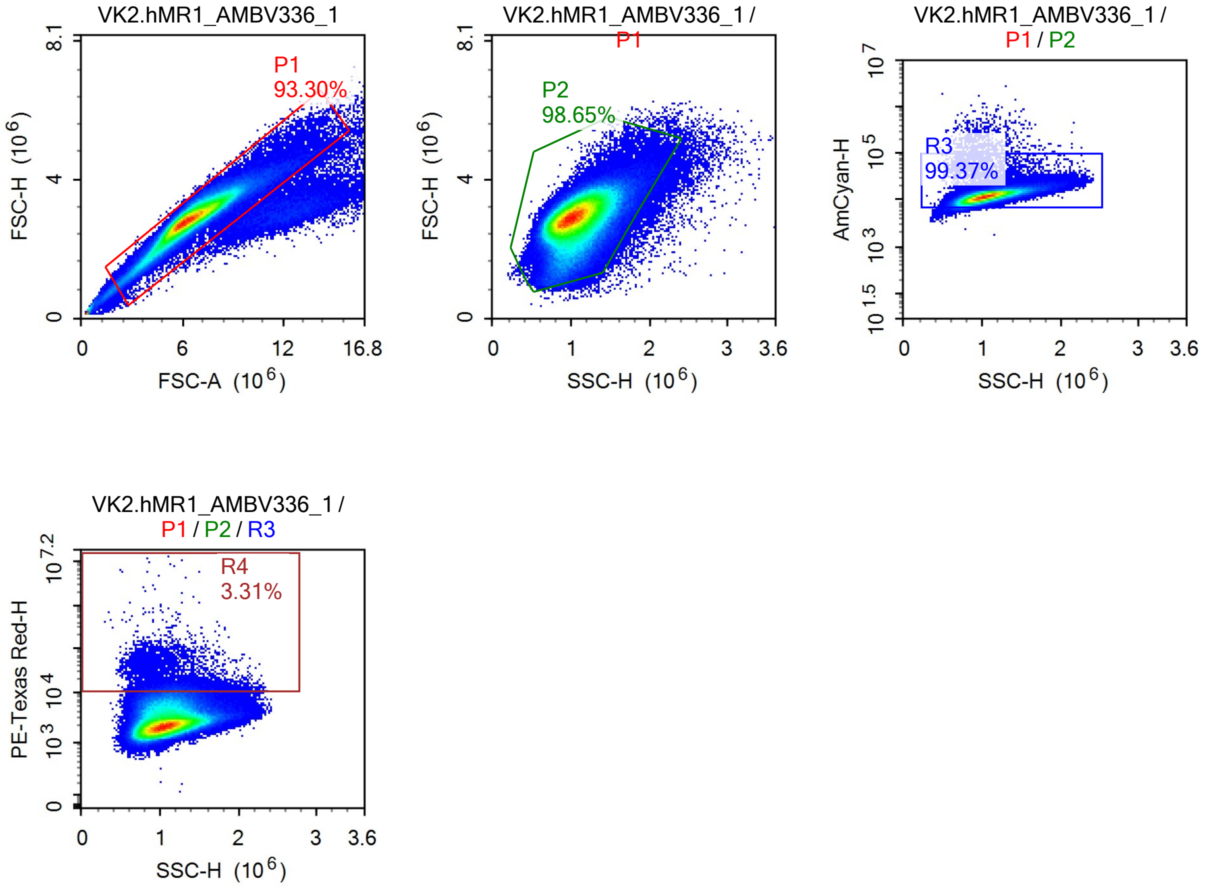
VK2.hMR1_AMBV336_1
P1
93.30%
FSC-H (10^6)
FSC-A (10^6)
VK2.hMR1_AMBV336_1 / P1
P2
98.65%
SSC-H (10^6)
VK2.hMR1_AMBV336_1 / P1 / P2
R3
99.37%
AmCyan-H
VK2.hMR1_AMBV336_1 / P1 / P2 / R3
R4
3.31%
PE-Texas Red-H

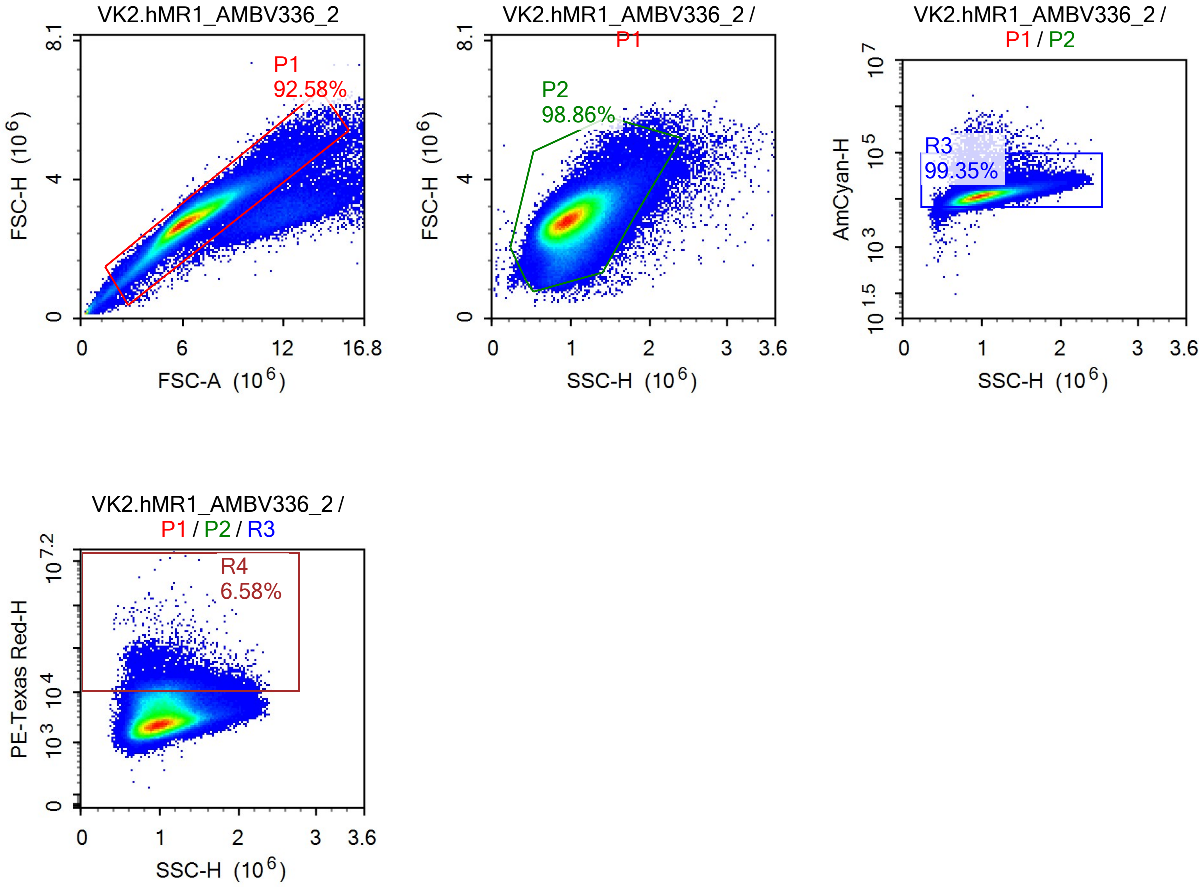
VK2.hMR1_AMBV336_2
P1
92.58%
FSC-H (10^6)
FSC-A (10^6)
VK2.hMR1_AMBV336_2 / P1
P2
98.86%
FSC-H (10^6)
SSC-H (10^6)
VK2.hMR1_AMBV336_2 / P1 / P2
R3
99.35%
AmCyan-H
SSC-H (10^6)
VK2.hMR1_AMBV336_2 / P1 / P2 / R3
R4
6.58%
PE-Texas Red-H
SSC-H (10^6)

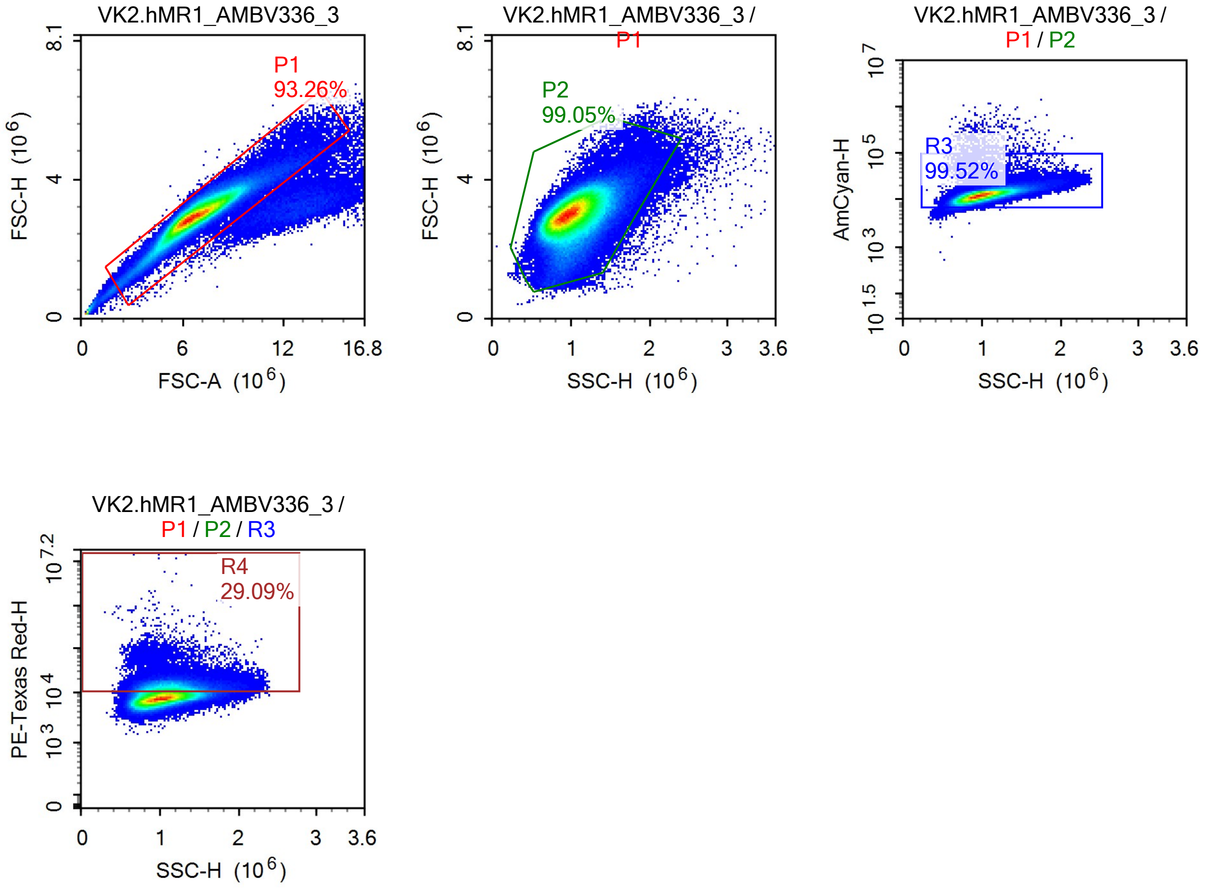

VK2.hMR1_AMBV336_3
P1
93.26%
FSC-H (10^6)
FSC-A (10^6)
VK2.hMR1_AMBV336_3 / P1
P2
99.05%
SSC-H (10^6)
VK2.hMR1_AMBV336_3 / P1 / P2
R3
99.52%
AmCyan-H
VK2.hMR1_AMBV336_3 / P1 / P2 / R3
R4
29.09%
PE-Texas Red-H

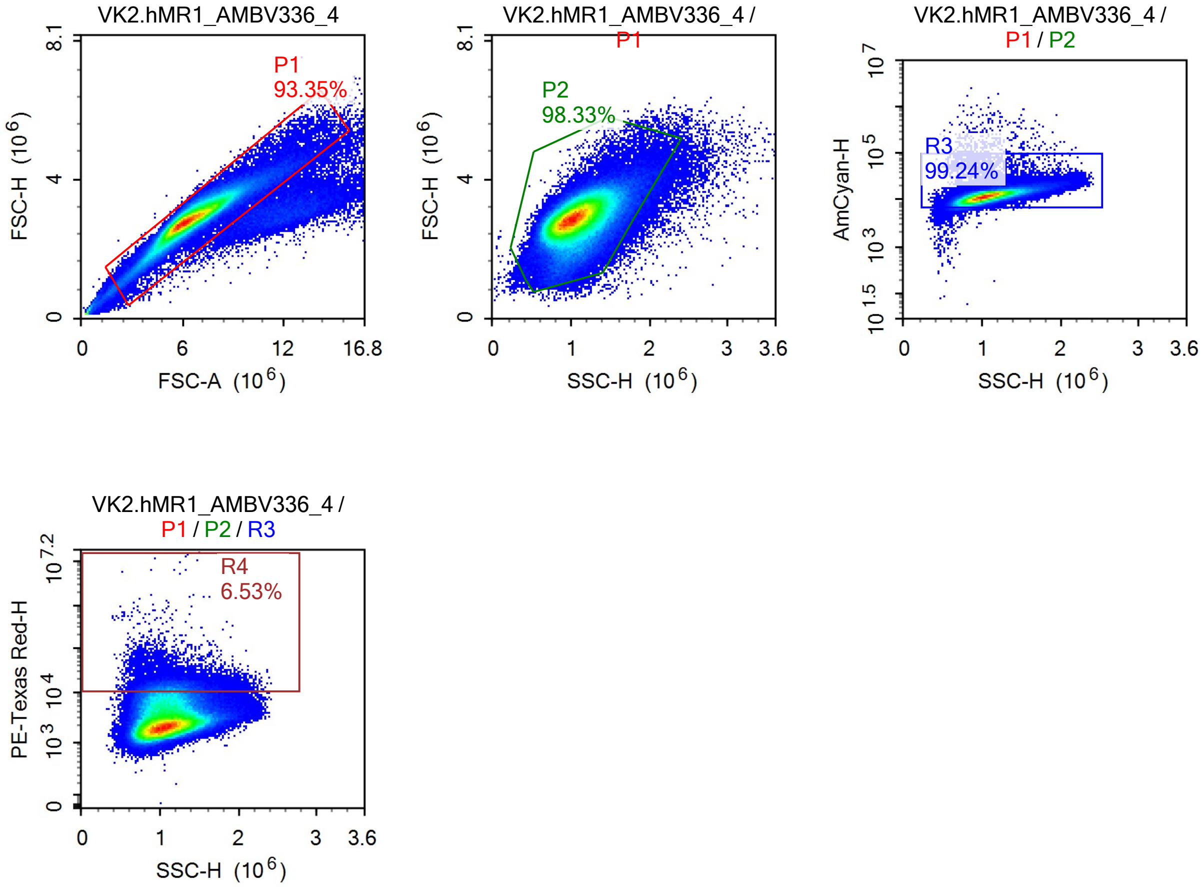
VK2.hMR1_AMBV336_4
P1
93.35%
FSC-H (10^6)
FSC-A (10^6)
VK2.hMR1_AMBV336_4 / P1
P2
98.33%
FSC-H (10^6)
SSC-H (10^6)
VK2.hMR1_AMBV336_4 / P1 / P2
R3
99.24%
AmCyan-H
SSC-H (10^6)
VK2.hMR1_AMBV336_4 / P1 / P2 / R3
R4
6.53%
PE-Texas Red-H
SSC-H (10^6)

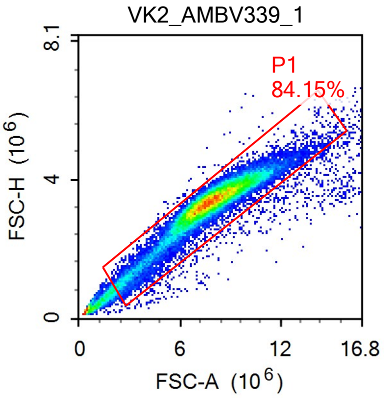
VK2_AMBV339_1
P1
84.15%
FSC-H (10^6)
8.1
4
0
0
6
12
16.8
FSC-A (10^6)

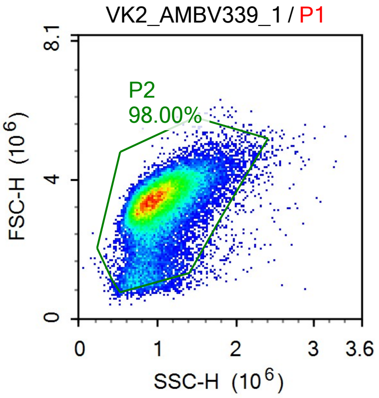
VK2_AMBV339_1 / P1
P2
98.00%
FSC-H (10^6)
8.1
4
0
0
1
2
3
3.6
SSC-H (10^6)

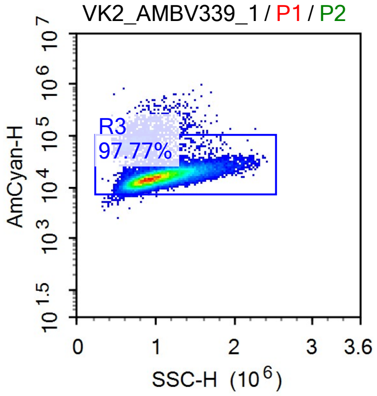
VK2_AMBV339_1 / P1 / P2
R3
97.77%
AmCyan-H
10^7
10^6
10^5
10^4
10^3
10^1.5
0
1
2
3
3.6
SSC-H (10^6)

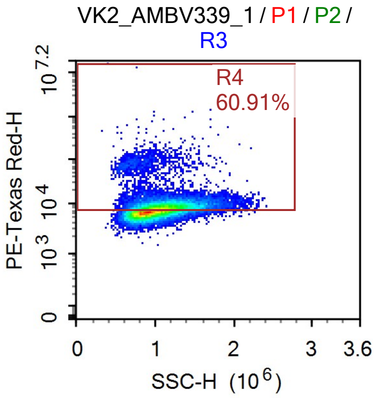
VK2_AMBV339_1 / P1 / P2 / R3
R4
60.91%
PE-Texas Red-H
10^7.2
10^4
10^3
0
0
1
2
3
3.6
SSC-H (10^6)

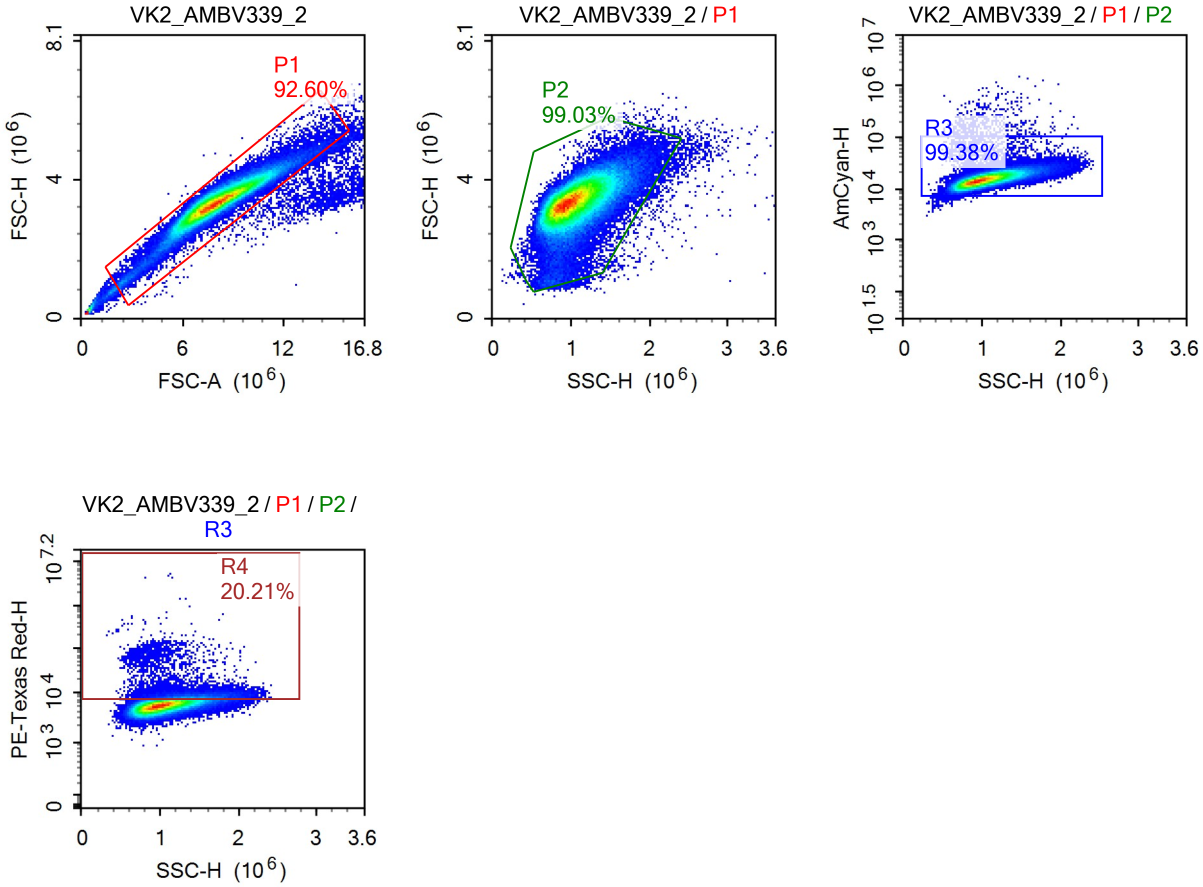
VK2_AMBV339_2
P1
92.60%
FSC-H (10^6)
FSC-A (10^6)
VK2_AMBV339_2 / P1
P2
99.03%
FSC-H (10^6)
SSC-H (10^6)
VK2_AMBV339_2 / P1 / P2
R3
99.38%
AmCyan-H
SSC-H (10^6)
VK2_AMBV339_2 / P1 / P2 / R3
R4
20.21%
PE-Texas Red-H
SSC-H (10^6)

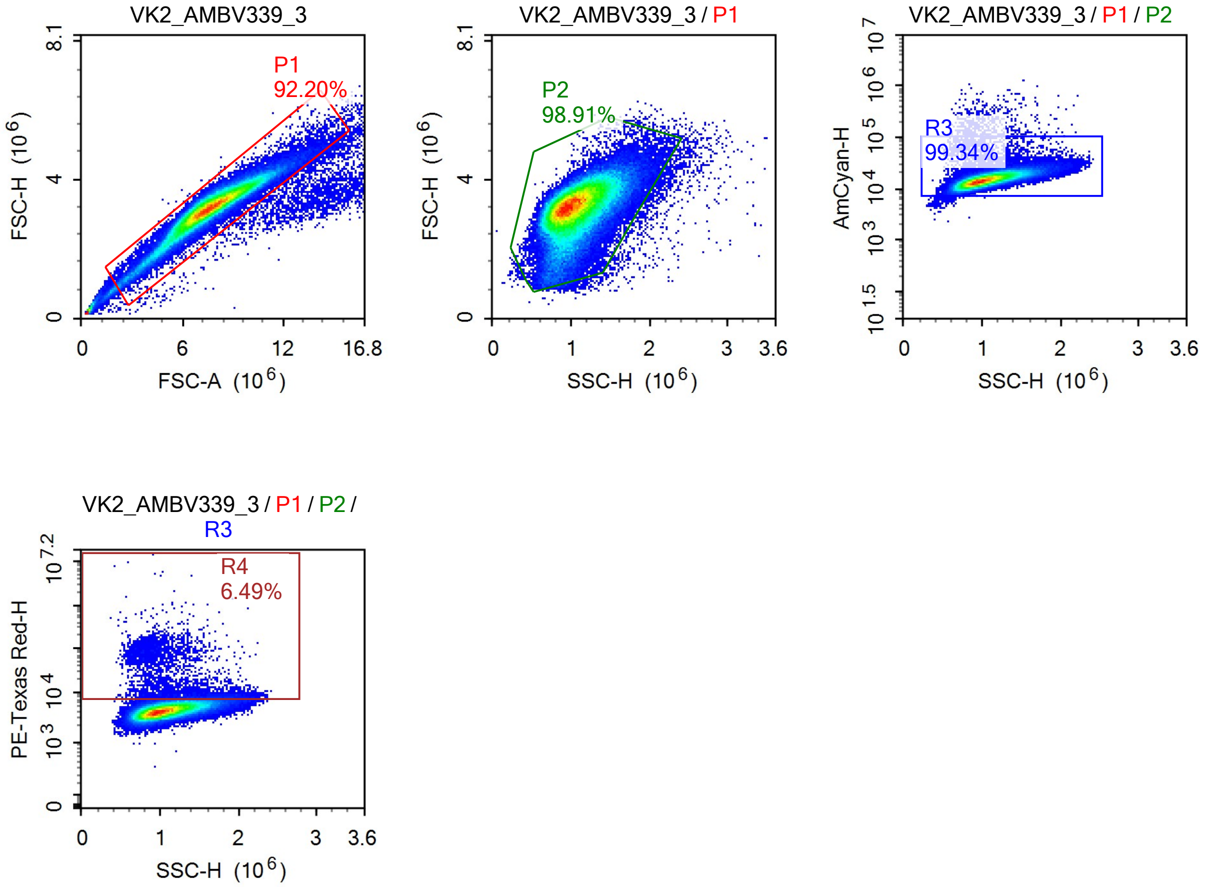

VK2_AMBV339_3
P1
92.20%
FSC-H (10^6)
FSC-A (10^6)
VK2_AMBV339_3 / P1
P2
98.91%
FSC-H (10^6)
SSC-H (10^6)
VK2_AMBV339_3 / P1 / P2
R3
99.34%
AmCyan-H
SSC-H (10^6)
VK2_AMBV339_3 / P1 / P2 / R3
R4
6.49%
PE-Texas Red-H
SSC-H (10^6)

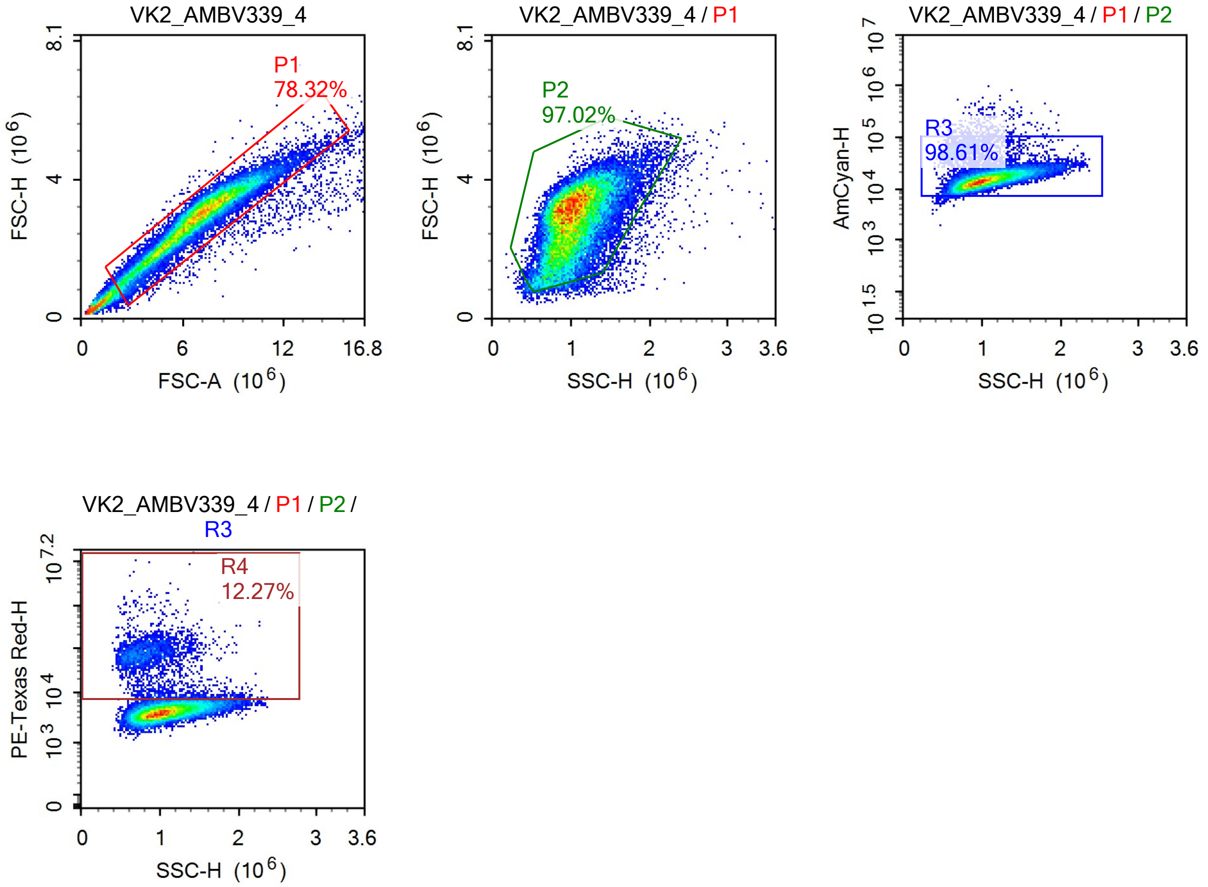
VK2_AMBV339_4
P1
78.32%
FSC-H (10^6)
FSC-A (10^6)
VK2_AMBV339_4 / P1
P2
97.02%
FSC-H (10^6)
SSC-H (10^6)
VK2_AMBV339_4 / P1 / P2
R3
98.61%
AmCyan-H
SSC-H (10^6)
VK2_AMBV339_4 / P1 / P2 / R3
R4
12.27%
PE-Texas Red-H
SSC-H (10^6)

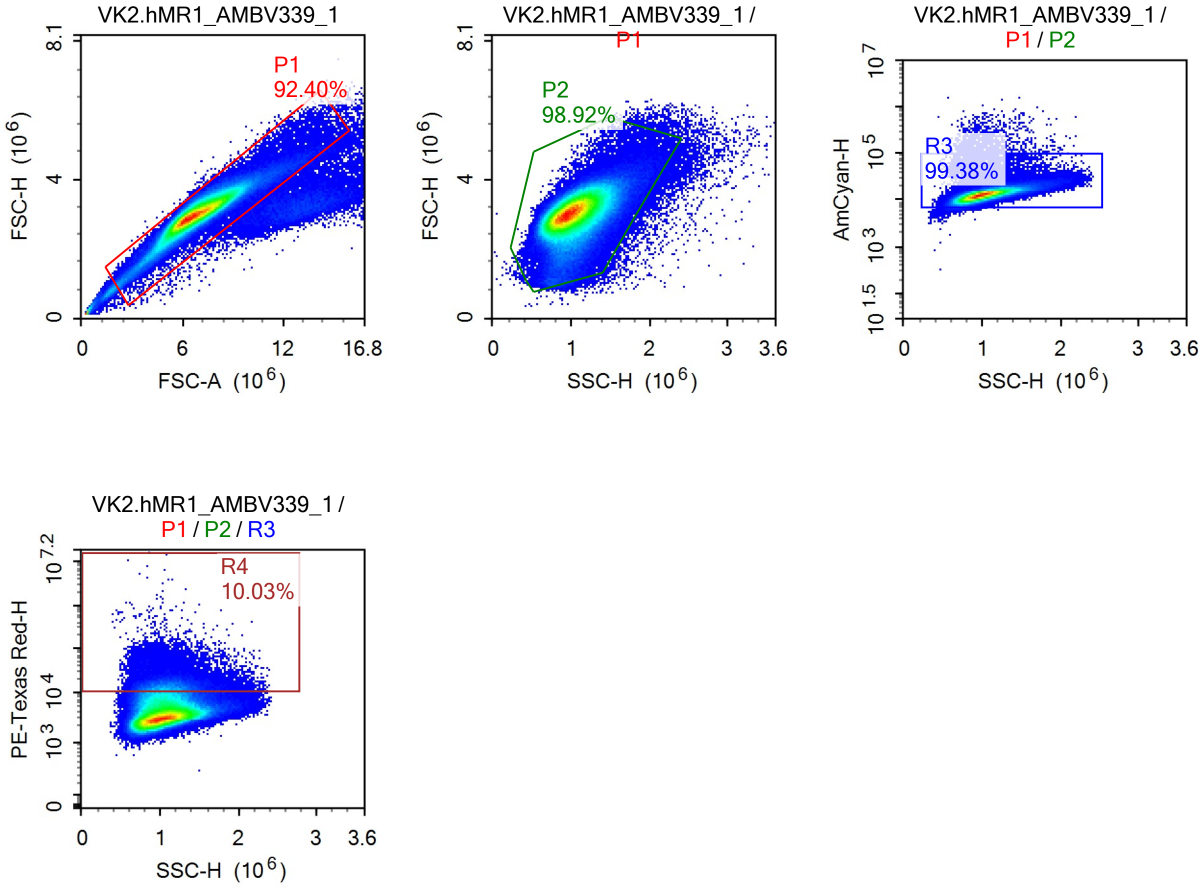
VK2.hMR1_AMBV339_1
P1
92.40%
FSC-H (10^6)
FSC-A (10^6)
VK2.hMR1_AMBV339_1 / P1
P2
98.92%
FSC-H (10^6)
SSC-H (10^6)
VK2.hMR1_AMBV339_1 / P1 / P2
R3
99.38%
AmCyan-H
SSC-H (10^6)
VK2.hMR1_AMBV339_1 / P1 / P2 / R3
R4
10.03%
PE-Texas Red-H
SSC-H (10^6)

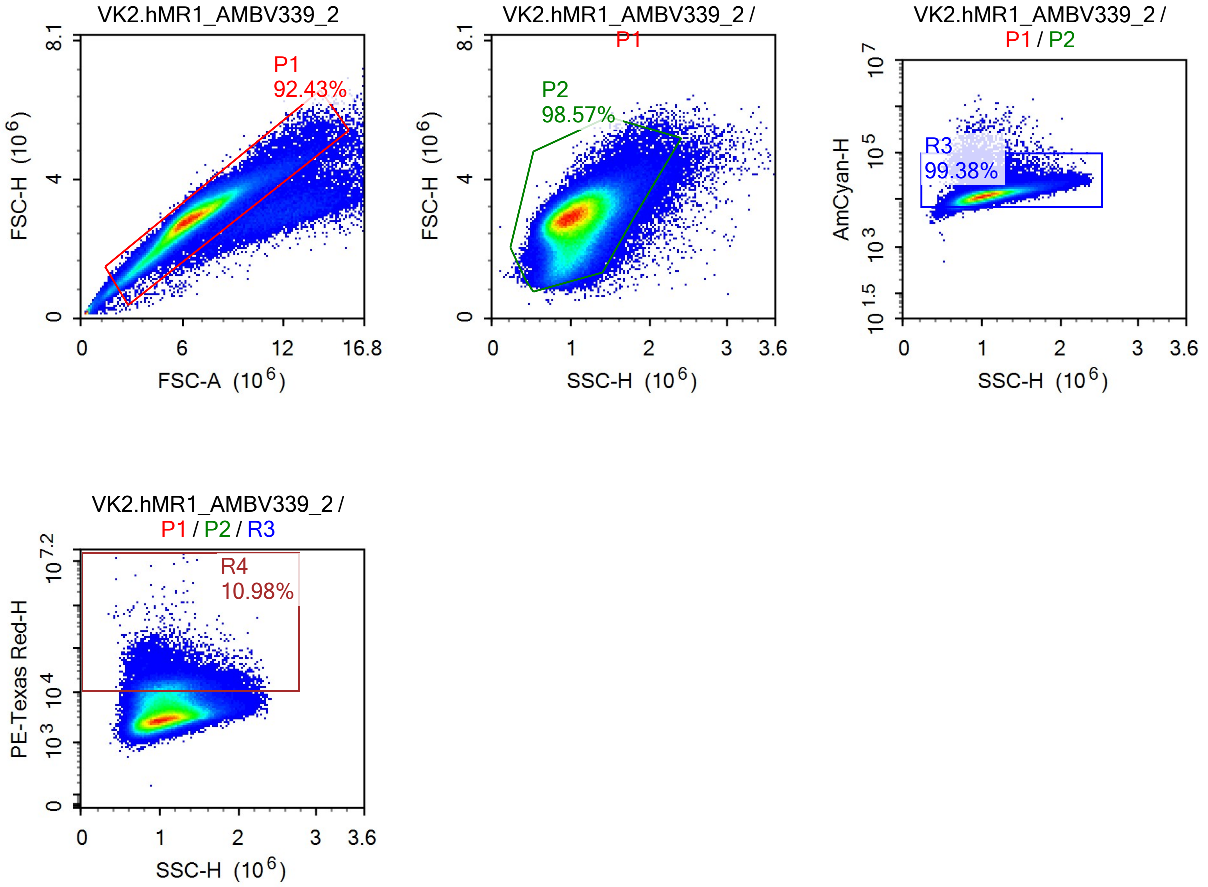
VK2.hMR1_AMBV339_2
P1
92.43%
FSC-H (10^6)
FSC-A (10^6)
0
6
12
16.8
4
8.1
VK2.hMR1_AMBV339_2 /
P1
P2
98.57%
FSC-H (10^6)
SSC-H (10^6)
1
2
3
3.6
VK2.hMR1_AMBV339_2 /
P1 / P2
R3
99.38%
AmCyan-H
10^7
10^5
10^3
10^1.5
SSC-H (10^6)
VK2.hMR1_AMBV339_2 /
P1 / P2 / R3
R4
10.98%
PE-Texas Red-H
10^7.2
10^4
10^3
SSC-H (10^6)

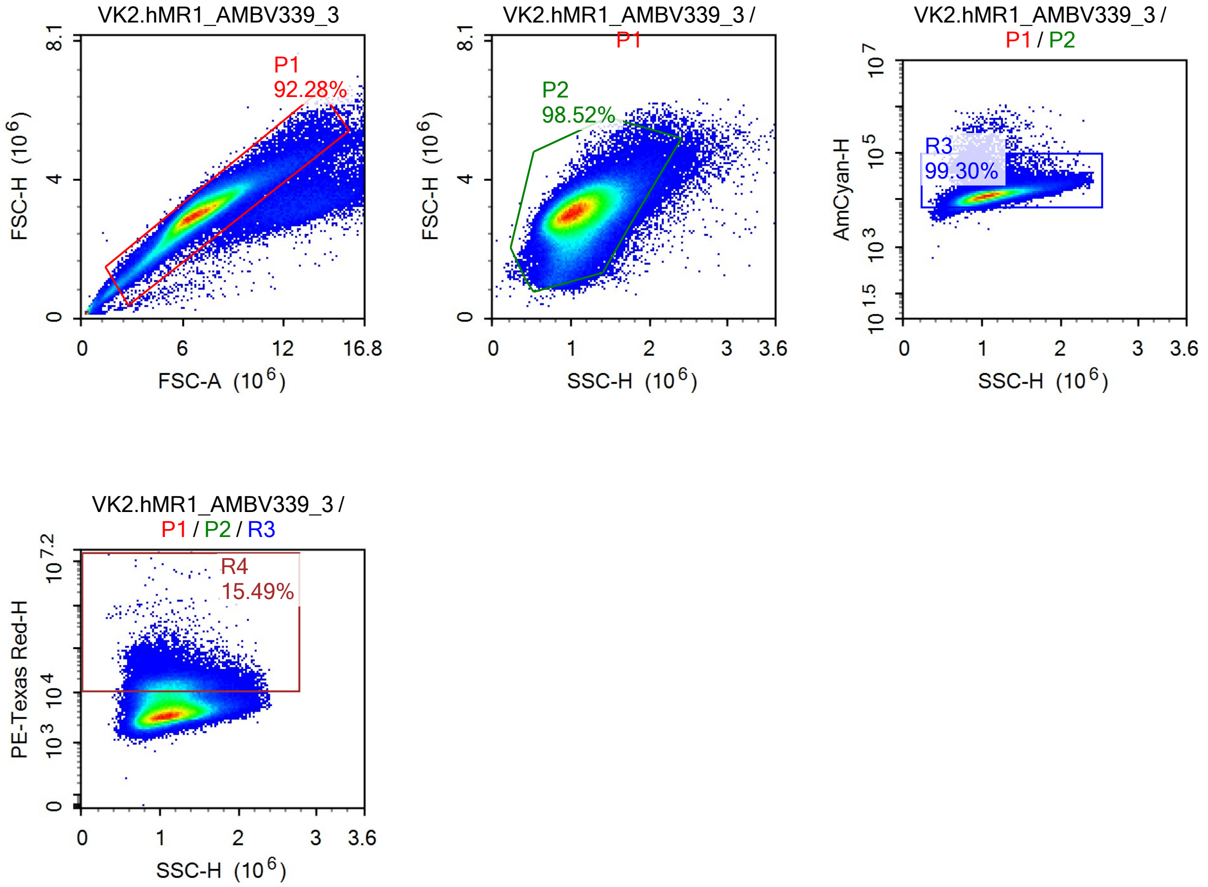
VK2.hMR1_AMBV339_3
P1
92.28%
FSC-H (10^6)
FSC-A (10^6)
0
6
12
16.8
4
8.1
VK2.hMR1_AMBV339_3 / P1
P2
98.52%
SSC-H (10^6)
1
2
3
3.6
VK2.hMR1_AMBV339_3 / P1 / P2
R3
99.30%
AmCyan-H
10^7
10^5
10^3
10^1.5
VK2.hMR1_AMBV339_3 / P1 / P2 / R3
R4
15.49%
PE-Texas Red-H
10^7.2
10^4

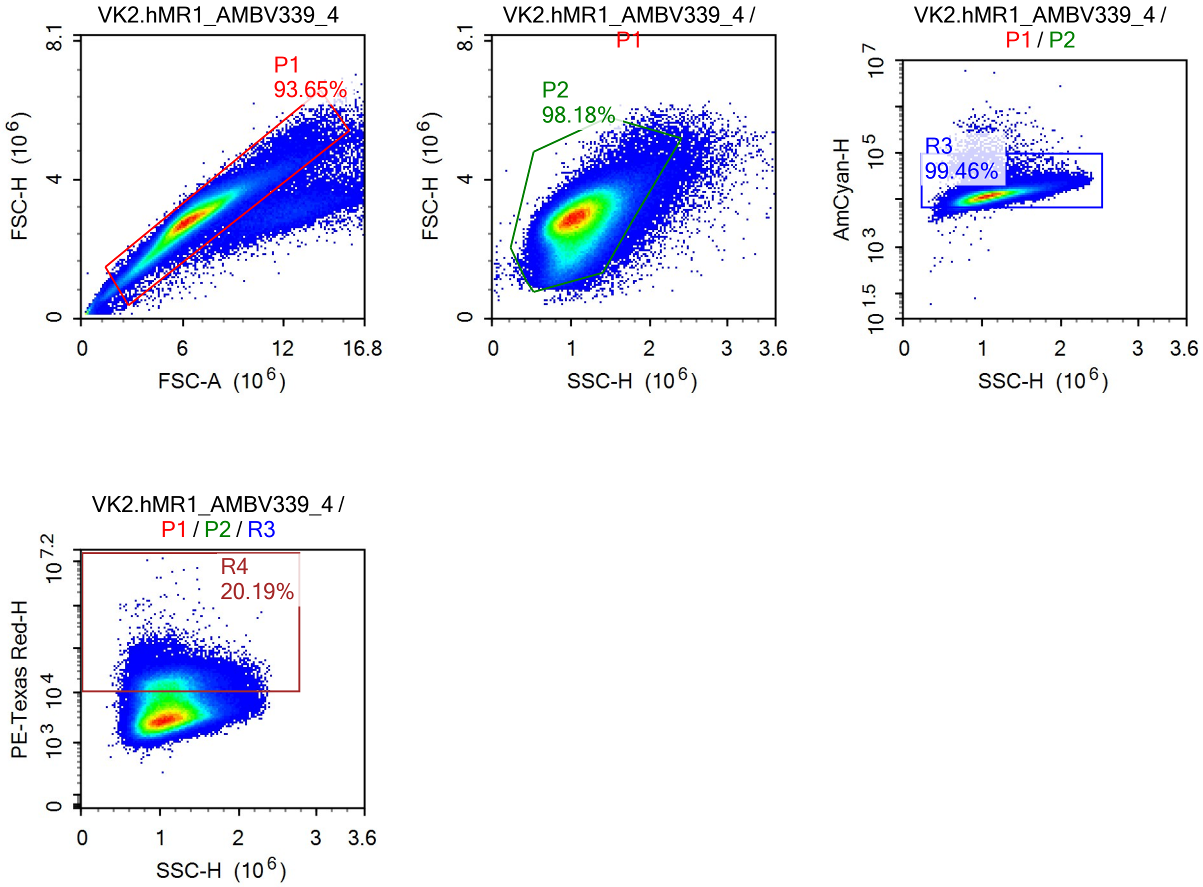
VK2.hMR1_AMBV339_4
P1
93.65%
FSC-H (10^6)
FSC-A (10^6)
VK2.hMR1_AMBV339_4 / P1
P2
98.18%
FSC-H (10^6)
SSC-H (10^6)
VK2.hMR1_AMBV339_4 / P1 / P2
R3
99.46%
AmCyan-H
SSC-H (10^6)
VK2.hMR1_AMBV339_4 / P1 / P2 / R3
R4
20.19%
PE-Texas Red-H
SSC-H (10^6)

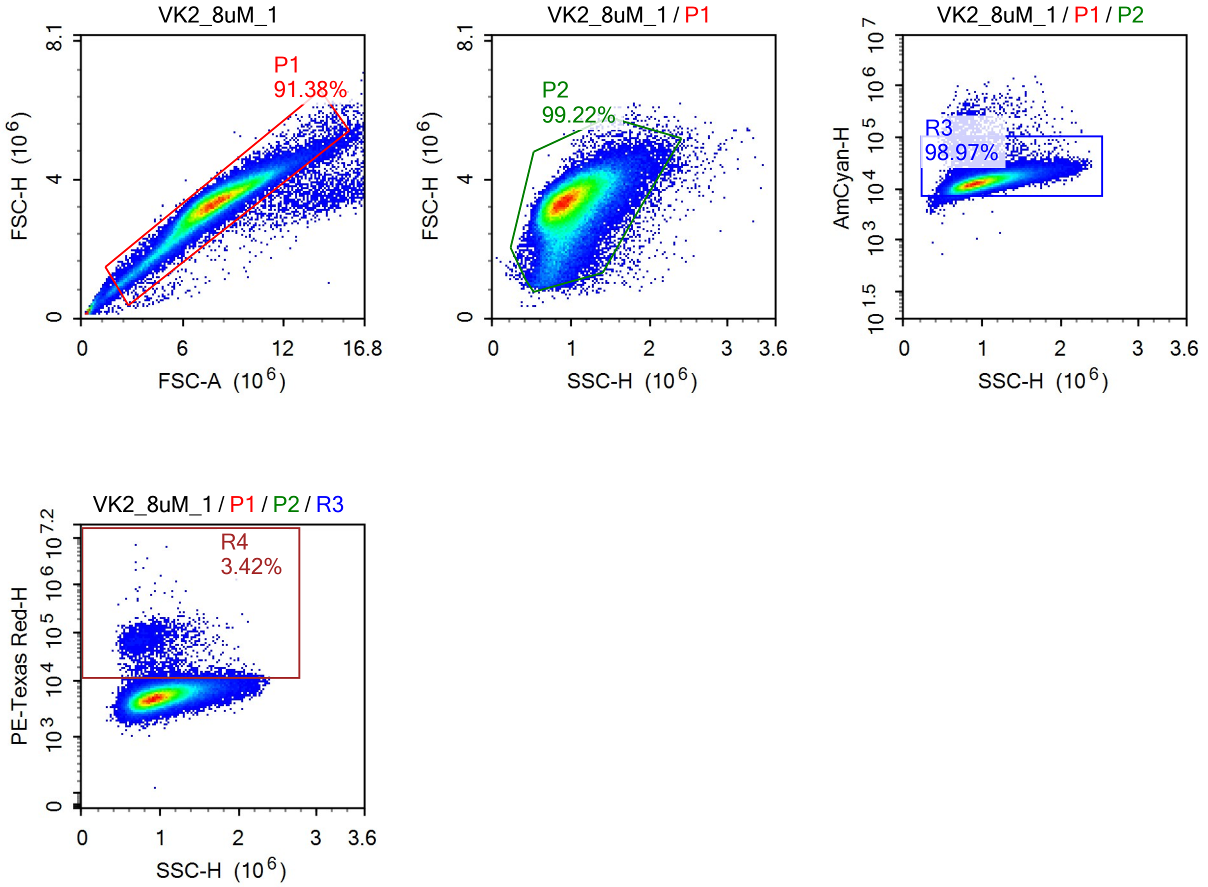
VK2_8uM_1
P1
91.38%
FSC-H (10^6)
FSC-A (10^6)
VK2_8uM_1 / P1
P2
99.22%
FSC-H (10^6)
SSC-H (10^6)
VK2_8uM_1 / P1 / P2
R3
98.97%
AmCyan-H
SSC-H (10^6)
VK2_8uM_1 / P1 / P2 / R3
R4
3.42%
PE-Texas Red-H
SSC-H (10^6)

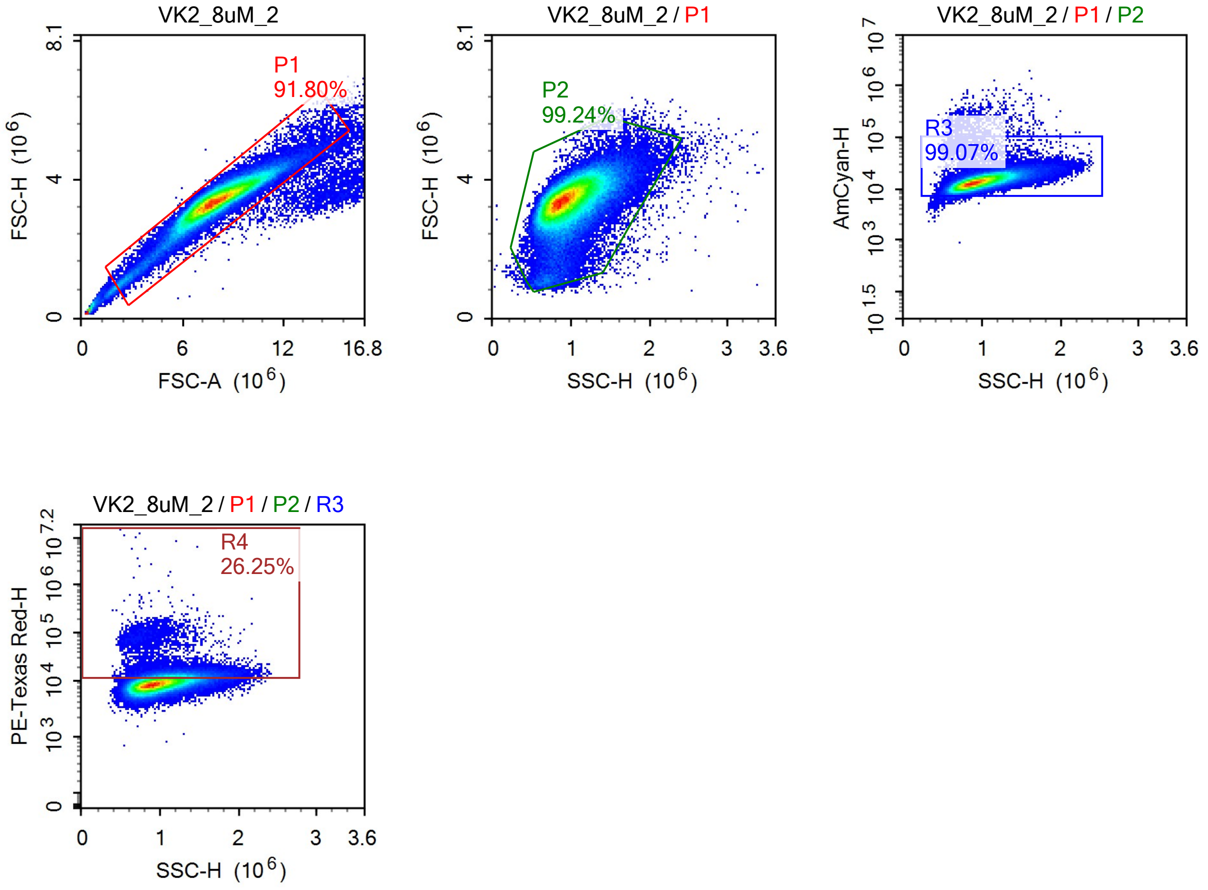
VK2_8uM_2
P1
91.80%
FSC-H (10^6)
FSC-A (10^6)
VK2_8uM_2 / P1
P2
99.24%
FSC-H (10^6)
SSC-H (10^6)
VK2_8uM_2 / P1 / P2
R3
99.07%
AmCyan-H
SSC-H (10^6)
VK2_8uM_2 / P1 / P2 / R3
R4
26.25%
PE-Texas Red-H
SSC-H (10^6)

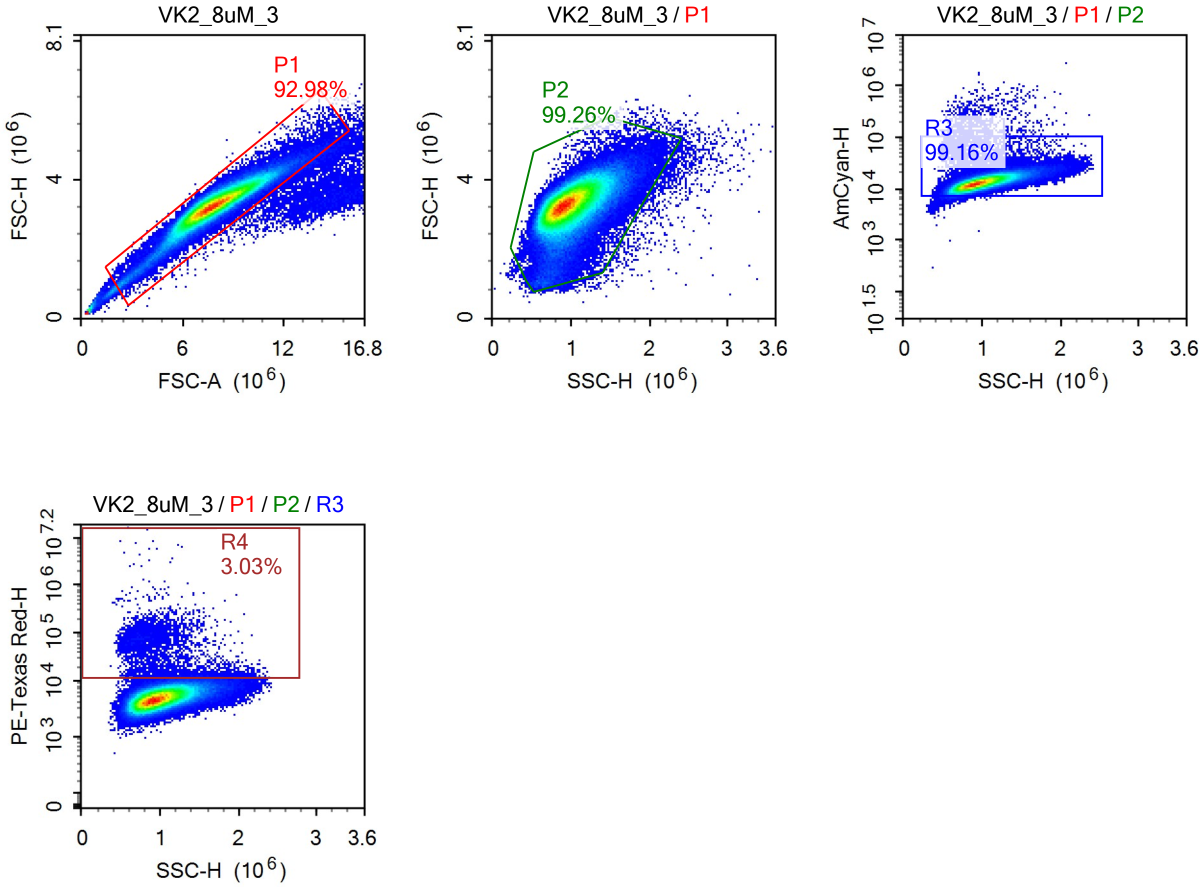
VK2_8uM_3
P1
92.98%
FSC-H (10^6)
FSC-A (10^6)
VK2_8uM_3 / P1
P2
99.26%
FSC-H (10^6)
SSC-H (10^6)
VK2_8uM_3 / P1 / P2
R3
99.16%
AmCyan-H
SSC-H (10^6)
VK2_8uM_3 / P1 / P2 / R3
R4
3.03%
PE-Texas Red-H
SSC-H (10^6)

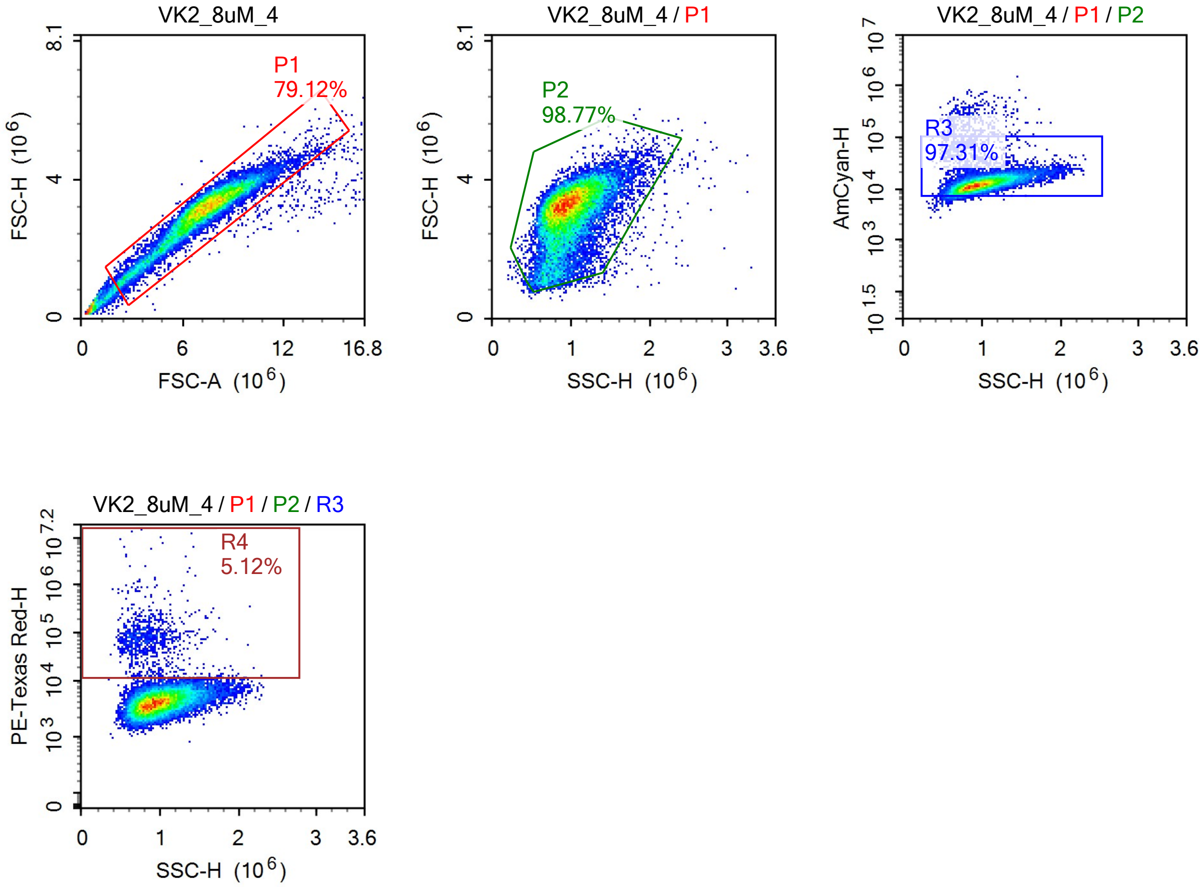
VK2_8uM_4
P1
79.12%
FSC-H (10^6)
FSC-A (10^6)
VK2_8uM_4 / P1
P2
98.77%
FSC-H (10^6)
SSC-H (10^6)
VK2_8uM_4 / P1 / P2
R3
97.31%
AmCyan-H
SSC-H (10^6)
VK2_8uM_4 / P1 / P2 / R3
R4
5.12%
PE-Texas Red-H
SSC-H (10^6)

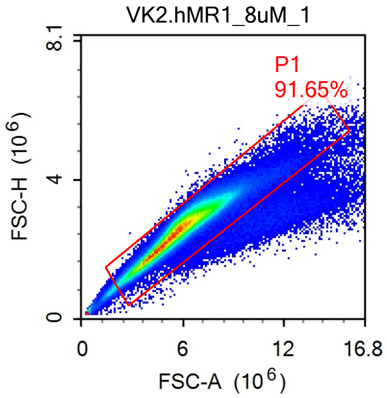
VK2.hMR1_8uM_1
P1
91.65%
FSC-H (10^6)
8.1
4
0
0
6
12
16.8
FSC-A (10^6)

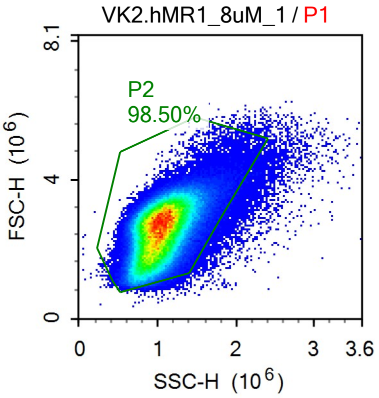
VK2.hMR1_8uM_1 / P1
P2
98.50%
FSC-H (10^6)
8.1
4
0
0
1
2
3
3.6
SSC-H (10^6)

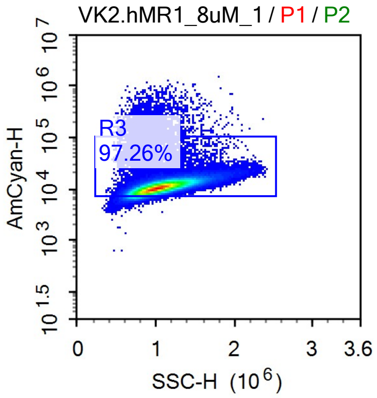
VK2.hMR1_8uM_1 / P1 / P2
R3
97.26%
AmCyan-H
10^7
10^6
10^5
10^4
10^3
10^1.5
0
1
2
3
3.6
SSC-H (10^6)

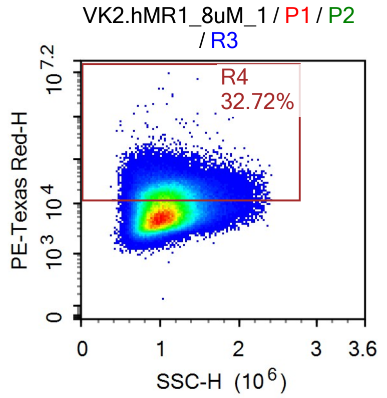
VK2.hMR1_8uM_1 / P1 / P2
/ R3
R4
32.72%
PE-Texas Red-H
10^7.2
10^4
10^3
0
0
1
2
3
3.6
SSC-H (10^6)

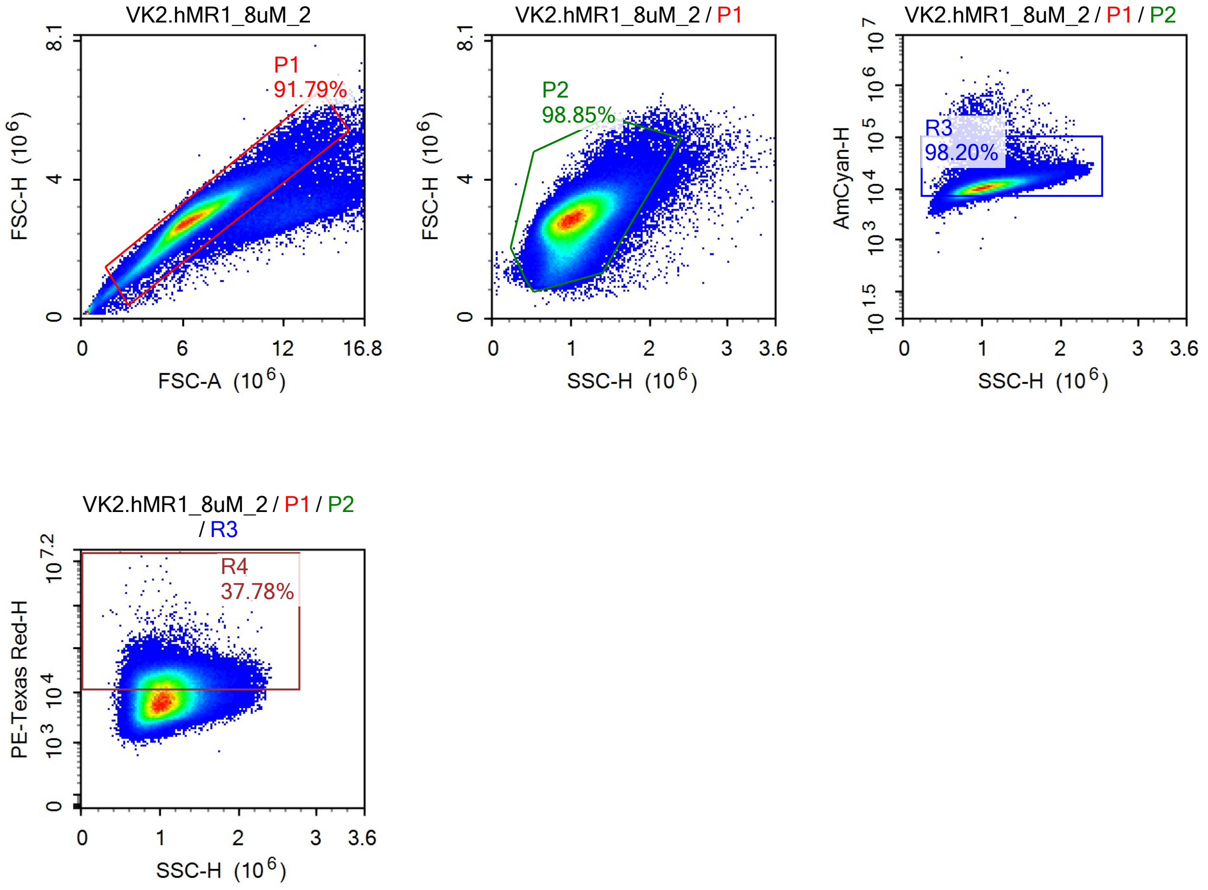
VK2.hMR1_8uM_2
P1
91.79%
FSC-H (10^6)
FSC-A (10^6)
VK2.hMR1_8uM_2 / P1
P2
98.85%
FSC-H (10^6)
SSC-H (10^6)
VK2.hMR1_8uM_2 / P1 / P2
R3
98.20%
AmCyan-H
SSC-H (10^6)
VK2.hMR1_8uM_2 / P1 / P2 / R3
R4
37.78%
PE-Texas Red-H
SSC-H (10^6)

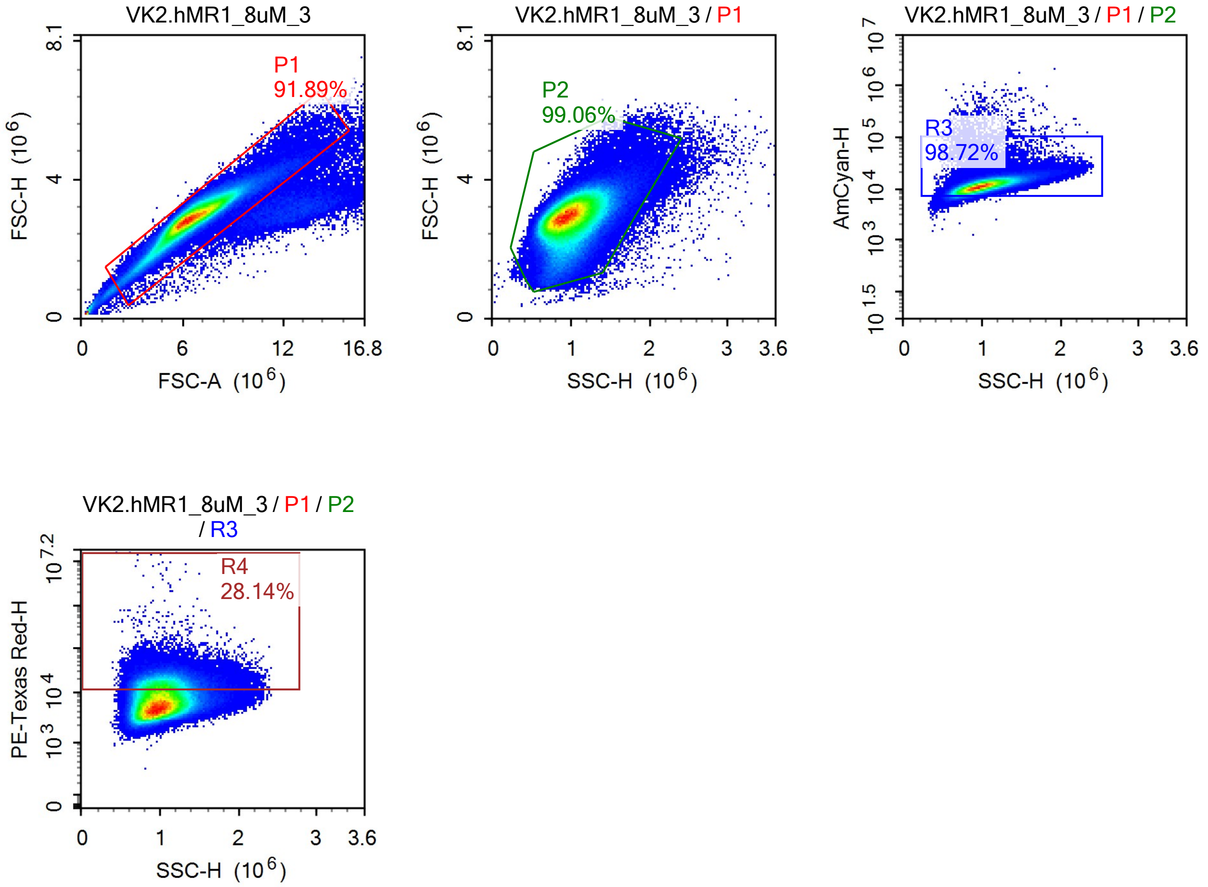
VK2.hMR1_8uM_3
P1
91.89%
FSC-H (10^6)
FSC-A (10^6)
VK2.hMR1_8uM_3 / P1
P2
99.06%
FSC-H (10^6)
SSC-H (10^6)
VK2.hMR1_8uM_3 / P1 / P2
R3
98.72%
AmCyan-H
SSC-H (10^6)
VK2.hMR1_8uM_3 / P1 / P2 / R3
R4
28.14%
PE-Texas Red-H
SSC-H (10^6)

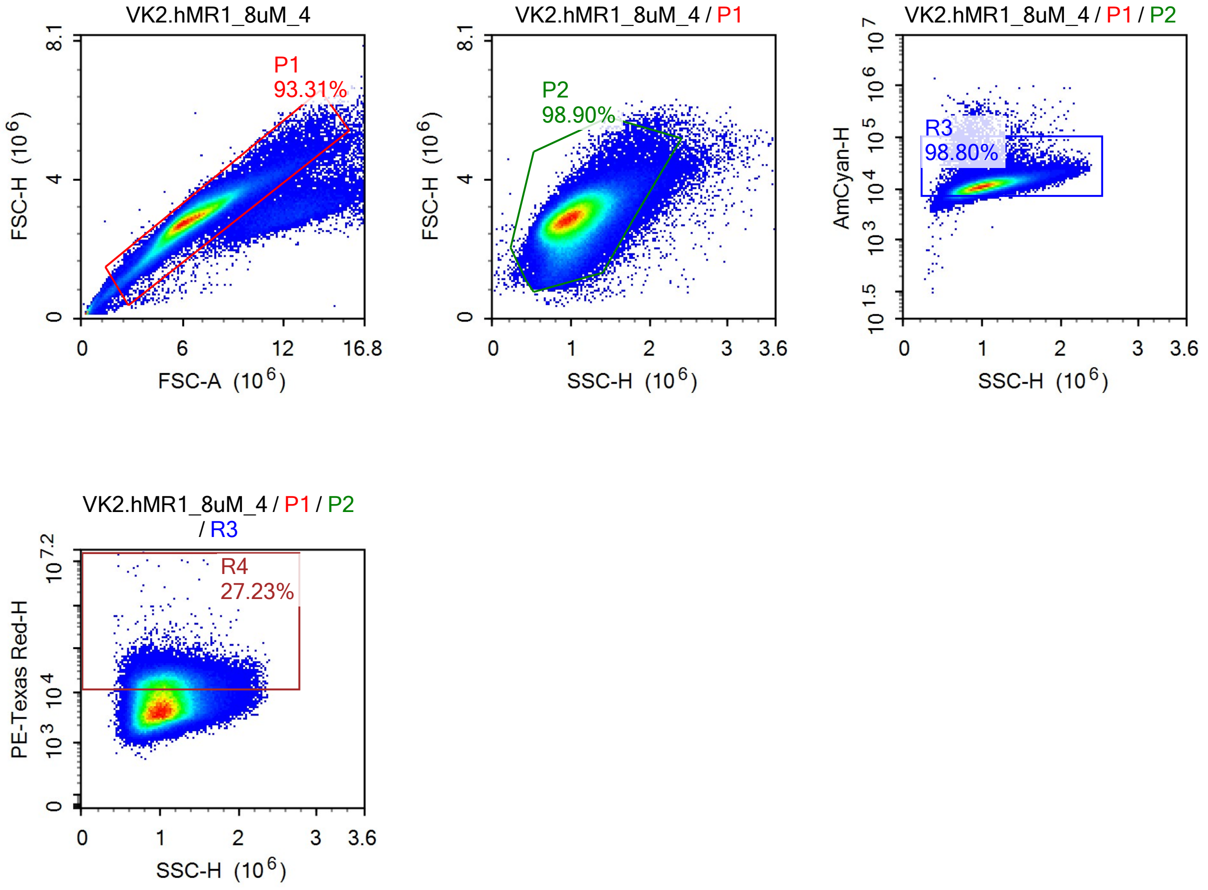
VK2.hMR1_8uM_4
P1
93.31%
FSC-H (10^6)
FSC-A (10^6)
VK2.hMR1_8uM_4 / P1
P2
98.90%
FSC-H (10^6)
SSC-H (10^6)
VK2.hMR1_8uM_4 / P1 / P2
R3
98.80%
AmCyan-H
SSC-H (10^6)
VK2.hMR1_8uM_4 / P1 / P2 / R3
R4
27.23%
PE-Texas Red-H
SSC-H (10^6)

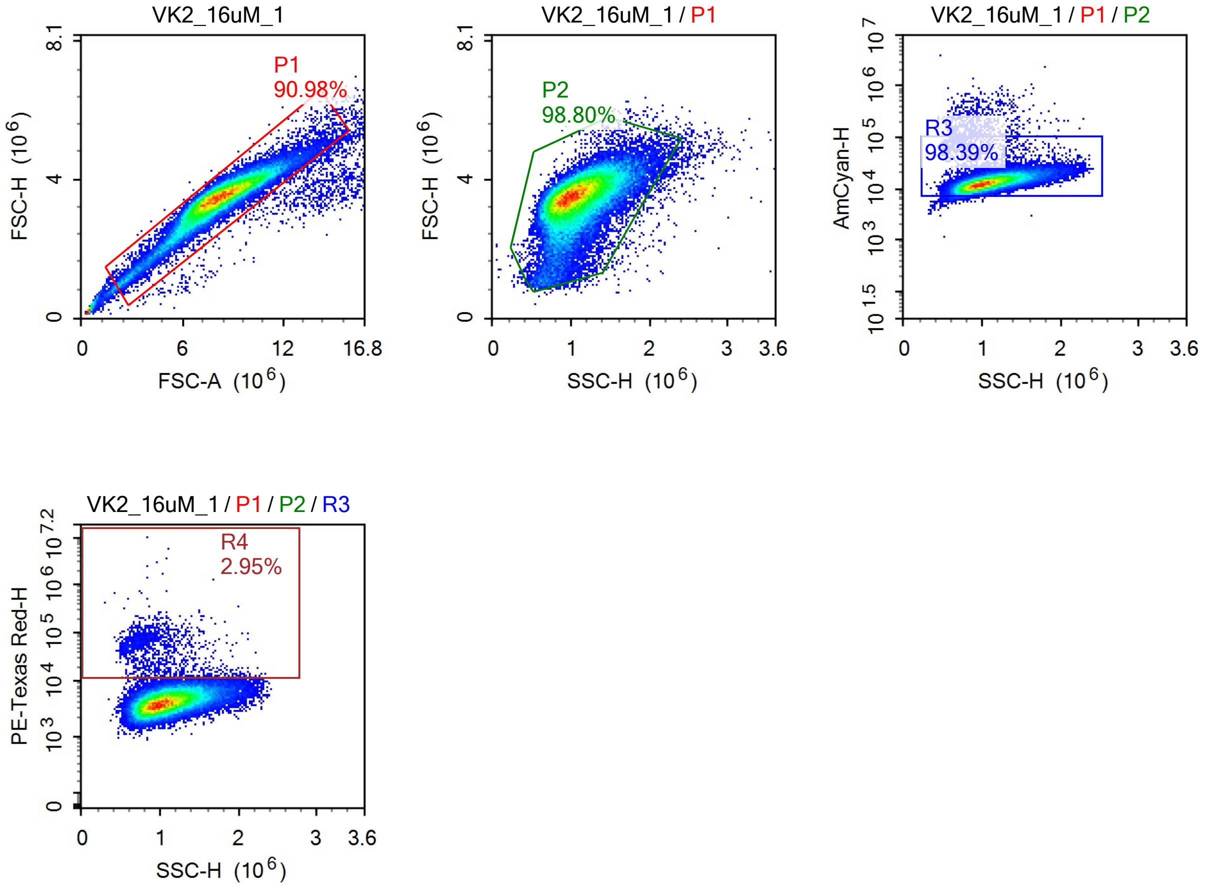
VK2_16uM_1
P1
90.98%
FSC-H (10^6)
FSC-A (10^6)
VK2_16uM_1 / P1
P2
98.80%
SSC-H (10^6)
VK2_16uM_1 / P1 / P2
R3
98.39%
AmCyan-H
VK2_16uM_1 / P1 / P2 / R3
R4
2.95%
PE-Texas Red-H

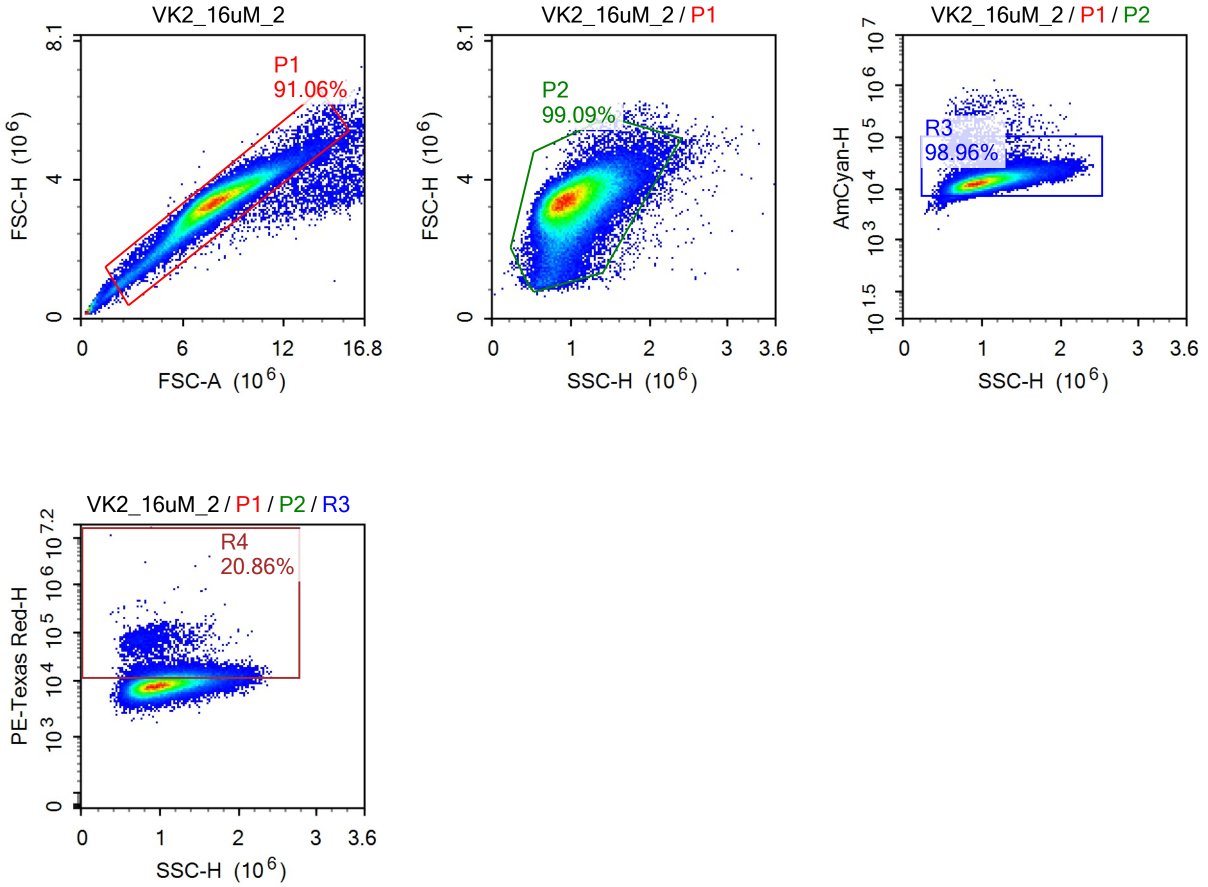
VK2_16uM_2
P1
91.06%
FSC-H (10^6)
FSC-A (10^6)
VK2_16uM_2 / P1
P2
99.09%
FSC-H (10^6)
SSC-H (10^6)
VK2_16uM_2 / P1 / P2
R3
98.96%
AmCyan-H
SSC-H (10^6)
VK2_16uM_2 / P1 / P2 / R3
R4
20.86%
PE-Texas Red-H
SSC-H (10^6)

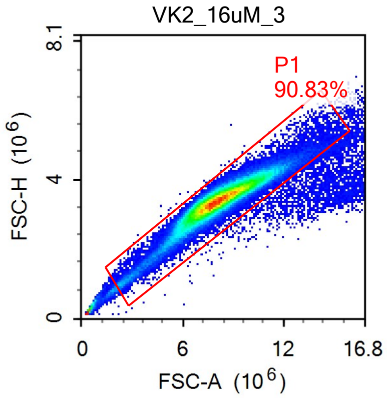
VK2_16uM_3
P1
90.83%
FSC-H (10^6)
FSC-A (10^6)
0
4
8.1
0
6
12
16.8

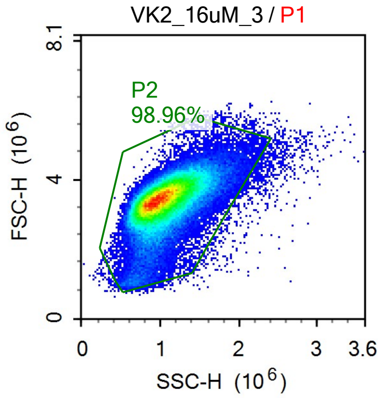
VK2_16uM_3 / P1
P2
98.96%
FSC-H (10^6)
SSC-H (10^6)
0
4
8.1
0
1
2
3
3.6

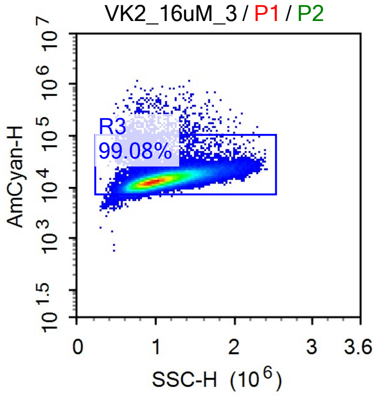
VK2_16uM_3 / P1 / P2
R3
99.08%
AmCyan-H
SSC-H (10^6)
10^1.5
10^3
10^4
10^5
10^6
10^7
0
1
2
3
3.6

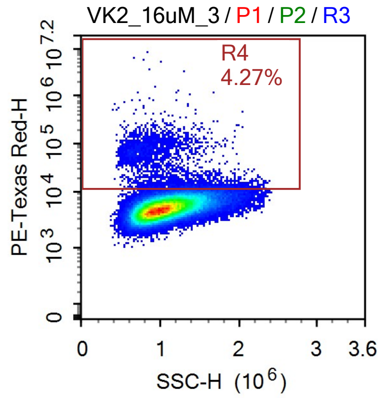
VK2_16uM_3 / P1 / P2 / R3
R4
4.27%
PE-Texas Red-H
SSC-H (10^6)
0
10^3
10^4
10^5
10^6
10^7.2
0
1
2
3
3.6

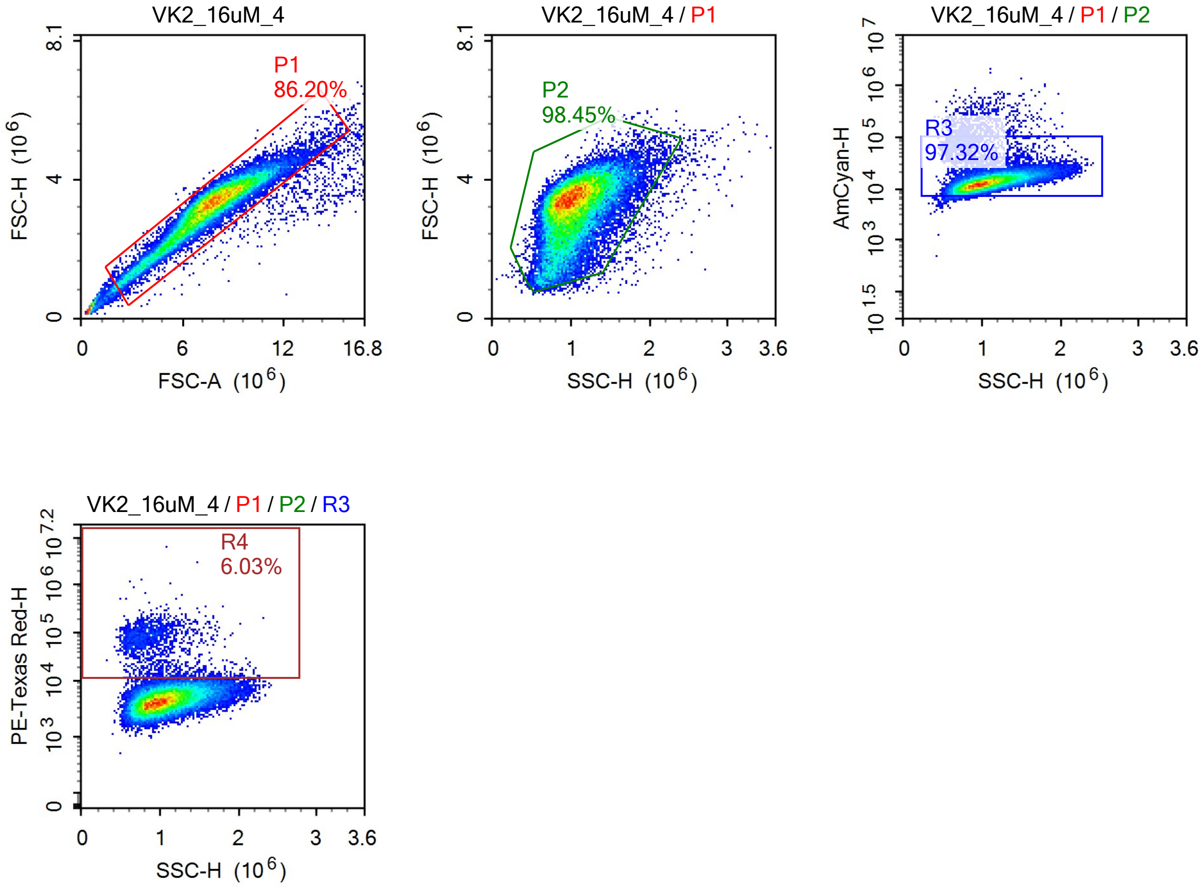
VK2_16uM_4
P1
86.20%
FSC-H (10^6)
FSC-A (10^6)
VK2_16uM_4 / P1
P2
98.45%
FSC-H (10^6)
SSC-H (10^6)
VK2_16uM_4 / P1 / P2
R3
97.32%
AmCyan-H
SSC-H (10^6)
VK2_16uM_4 / P1 / P2 / R3
R4
6.03%
PE-Texas Red-H
SSC-H (10^6)

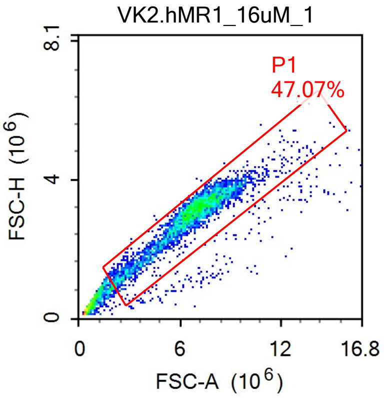
VK2.hMR1_16uM_1
P1
47.07%
FSC-H (10^6)
FSC-A (10^6)
0
6
12
16.8
4
8.1

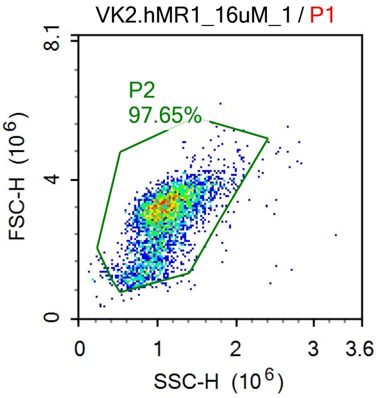
VK2.hMR1_16uM_1 / P1
P2
97.65%
FSC-H (10^6)
SSC-H (10^6)
0
1
2
3
3.6
4
8.1

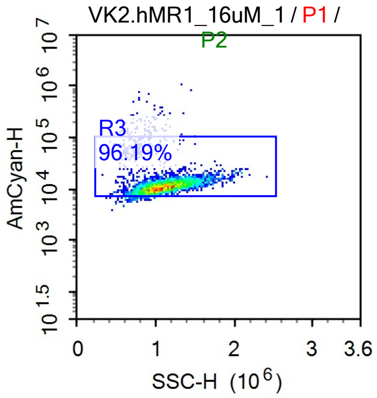
VK2.hMR1_16uM_1 / P1 / P2
R3
96.19%
AmCyan-H
SSC-H (10^6)
0
1
2
3
3.6
10^1.5
10^3
10^4
10^5
10^6
10^7

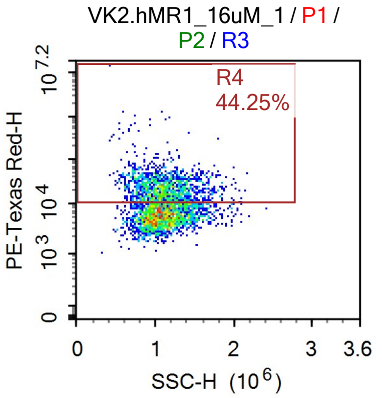
VK2.hMR1_16uM_1 / P1 / P2 / R3
R4
44.25%
PE-Texas Red-H
SSC-H (10^6)
0
1
2
3
3.6
10^3
10^4
10^7.2

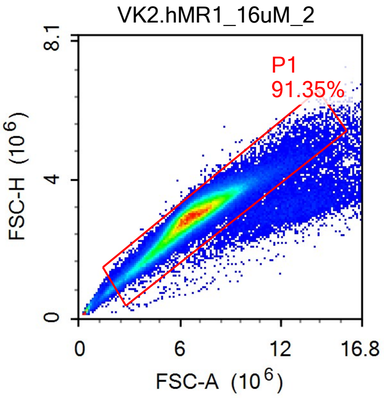
VK2.hMR1_16uM_2
P1
91.35%
FSC-H (10^6)
FSC-A (10^6)
0
4
8.1
0
6
12
16.8

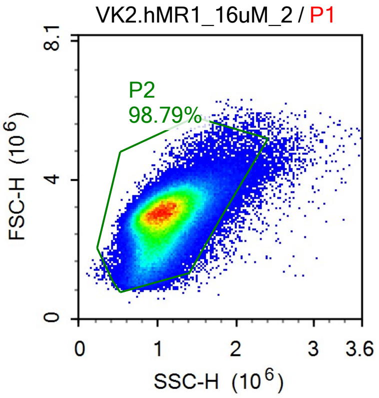
VK2.hMR1_16uM_2 / P1
P2
98.79%
FSC-H (10^6)
SSC-H (10^6)
0
4
8.1
0
1
2
3
3.6

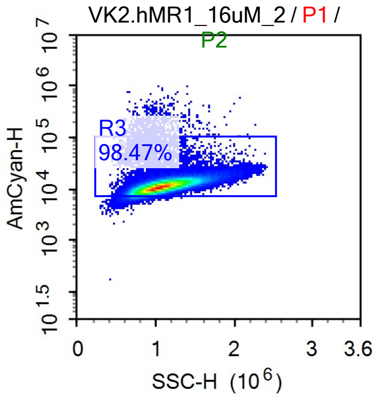
VK2.hMR1_16uM_2 / P1 / P2
R3
98.47%
AmCyan-H
SSC-H (10^6)
10^1.5
10^3
10^4
10^5
10^6
10^7
0
1
2
3
3.6

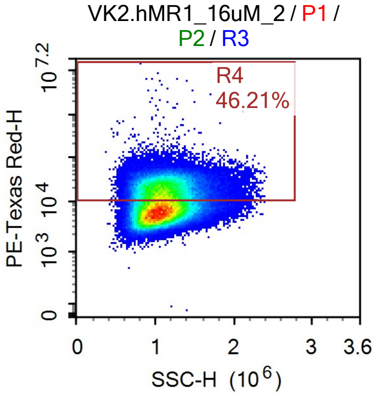
VK2.hMR1_16uM_2 / P1 / P2 / R3
R4
46.21%
PE-Texas Red-H
SSC-H (10^6)
0
10^3
10^4
10^7.2
0
1
2
3
3.6

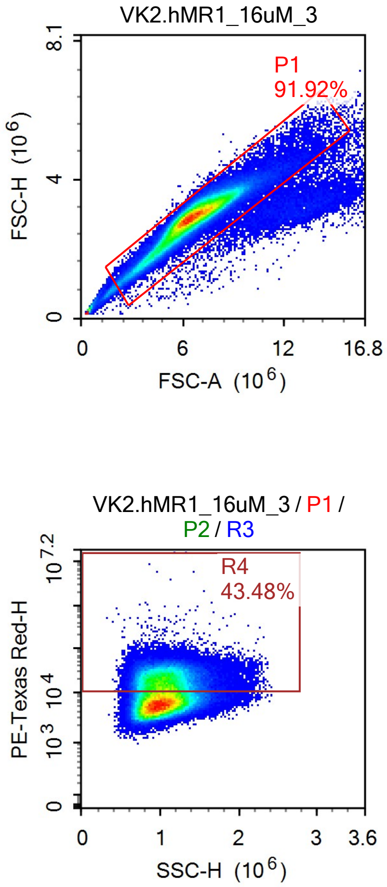

VK2.hMR1_16uM_3
P1
91.92%
FSC-H (10^6)
FSC-A (10^6)
VK2.hMR1_16uM_3 / P1
P2
99.19%
SSC-H (10^6)
VK2.hMR1_16uM_3 / P1 / P2
R3
98.76%
AmCyan-H
VK2.hMR1_16uM_3 / P1 / P2 / R3
R4
43.48%
PE-Texas Red-H

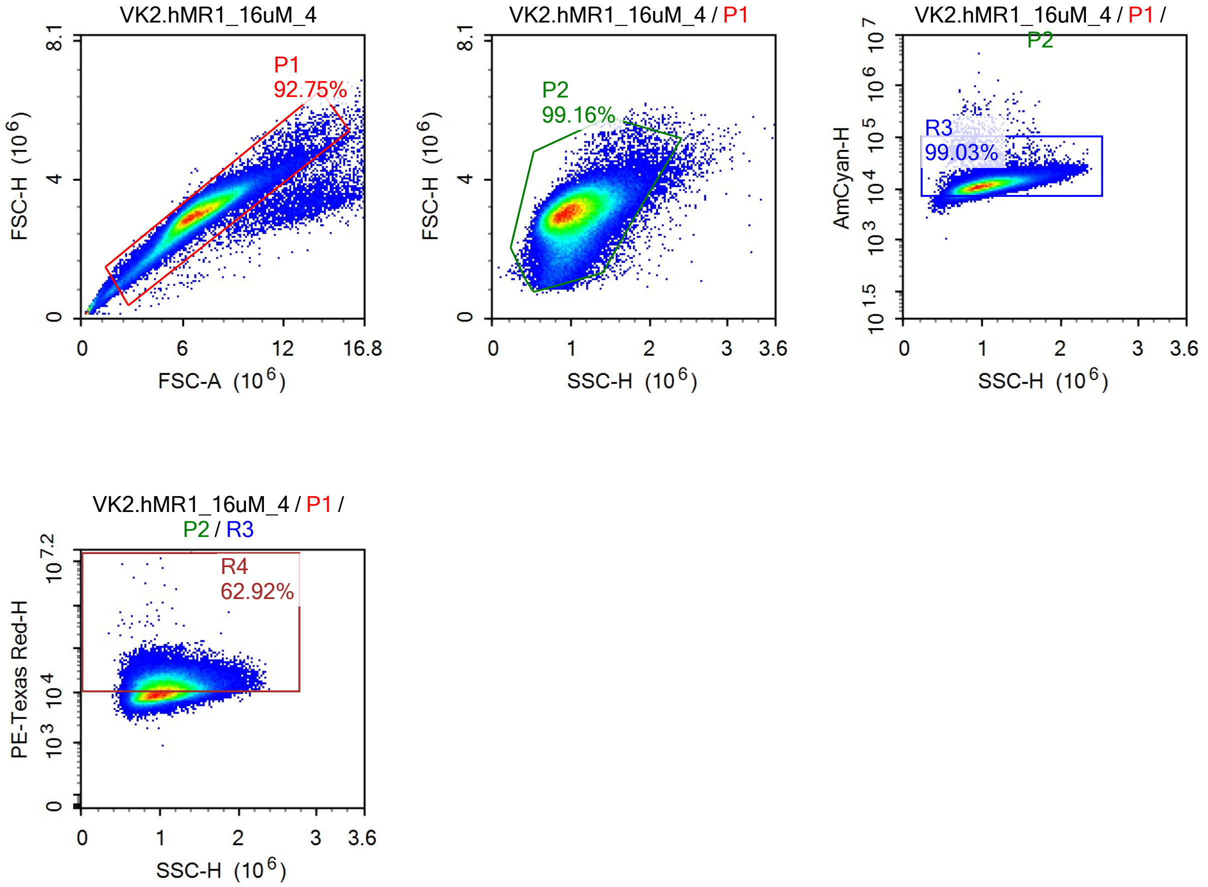
VK2.hMR1_16uM_4
P1
92.75%
FSC-H (10^6)
FSC-A (10^6)
0
4
8.1
0
6
12
16.8
VK2.hMR1_16uM_4 / P1
P2
99.16%
FSC-H (10^6)
SSC-H (10^6)
0
1
2
3
3.6
VK2.hMR1_16uM_4 / P1 / P2
R3
99.03%
AmCyan-H
10^1.5
10^3
10^4
10^5
10^6
10^7
SSC-H (10^6)
VK2.hMR1_16uM_4 / P1 / P2 / R3
R4
62.92%
PE-Texas Red-H
0
10^3
10^4
10^7.2
SSC-H (10^6)

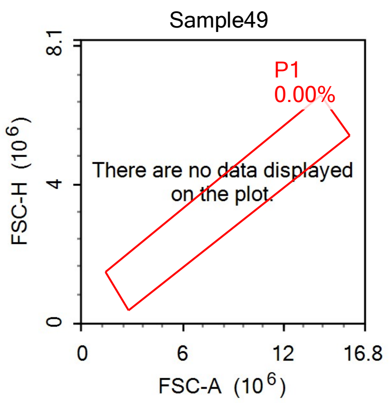
Sample49
P1
0.00%
There are no data displayed on the plot.
FSC-H (10^6)
FSC-A (10^6)

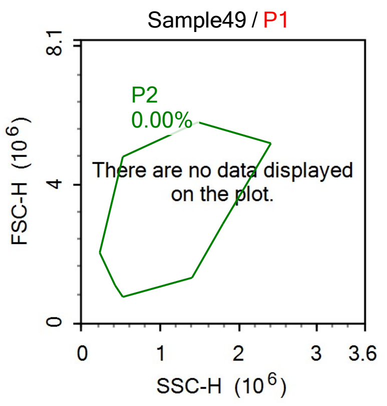
Sample49 / P1
P2
0.00%
There are no data displayed on the plot.
FSC-H (10^6)
SSC-H (10^6)

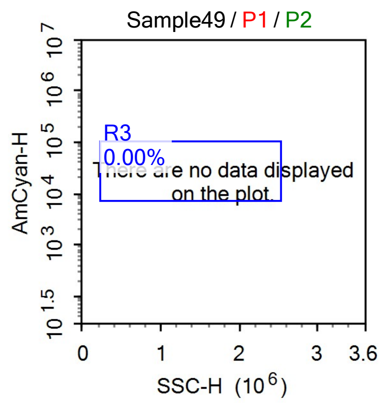
Sample49 / P1 / P2
R3
0.00%
There are no data displayed on the plot.
AmCyan-H
SSC-H (10^6)

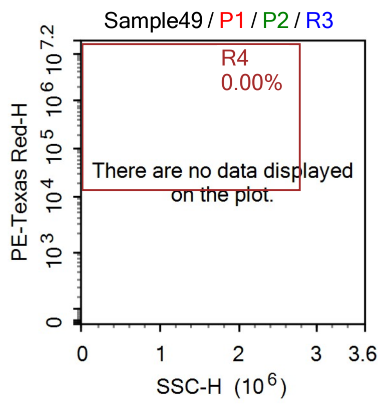
Sample49 / P1 / P2 / R3
R4
0.00%
There are no data displayed on the plot.
PE-Texas Red-H
SSC-H (10^6)

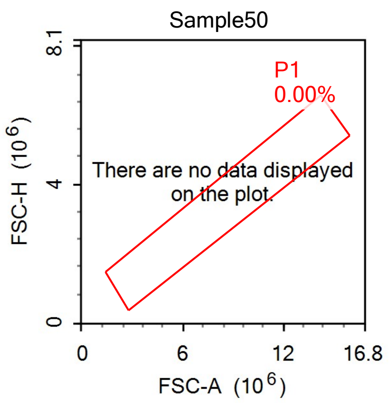
Sample50
P1
0.00%
There are no data displayed on the plot.
FSC-H (10^6)
FSC-A (10^6)

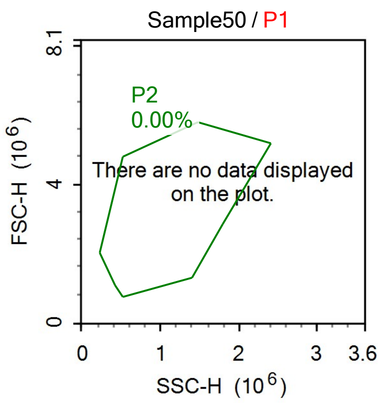
Sample50 / P1
P2
0.00%
There are no data displayed on the plot.
FSC-H (10^6)
SSC-H (10^6)

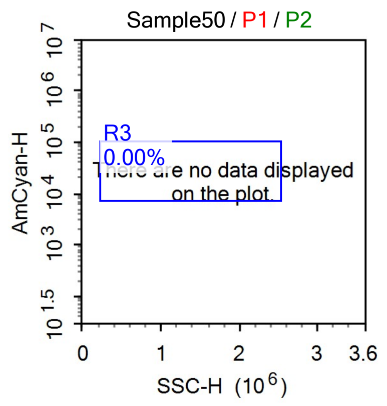
Sample50 / P1 / P2
R3
0.00%
There are no data displayed on the plot.
AmCyan-H
SSC-H (10^6)

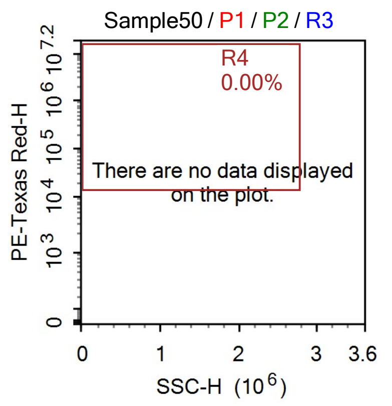
Sample50 / P1 / P2 / R3
R4
0.00%
There are no data displayed on the plot.
PE-Texas Red-H
SSC-H (10^6)

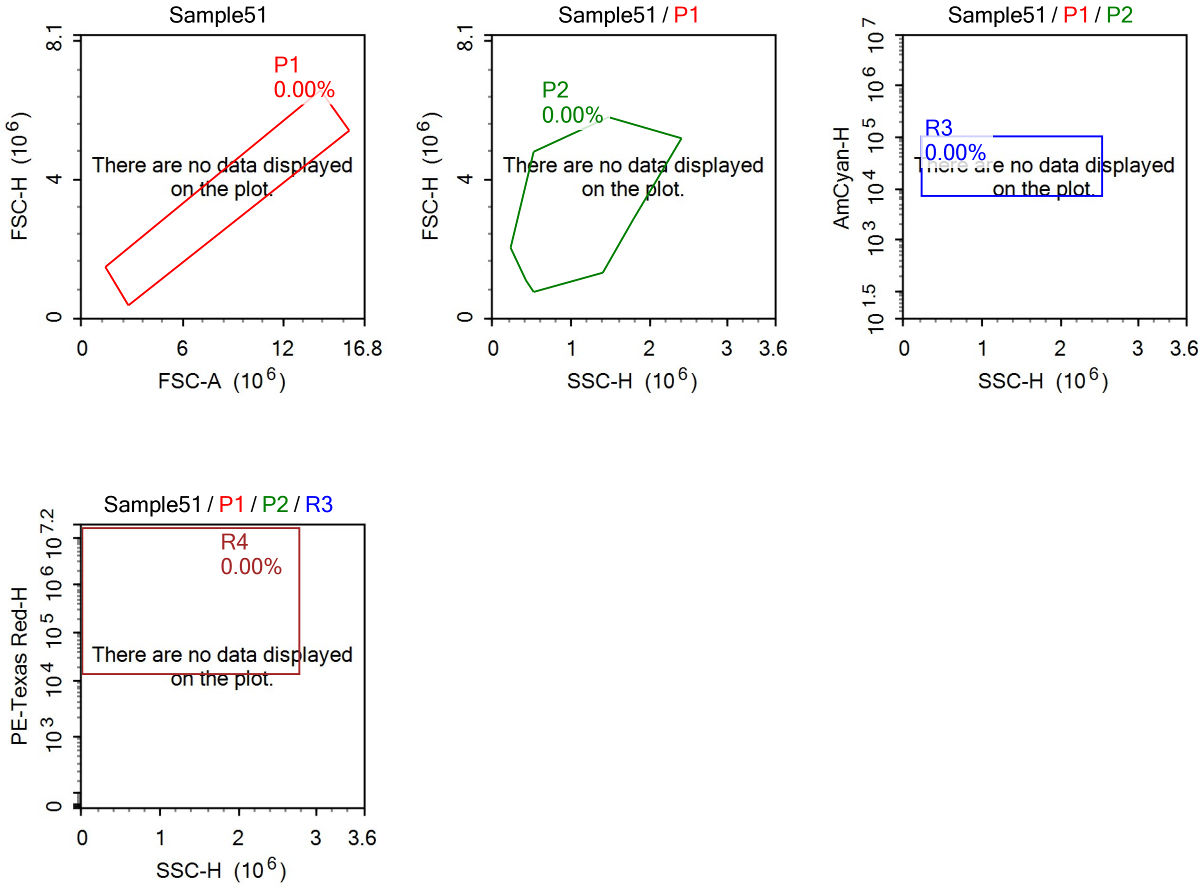

Sample51
P1
0.00%
There are no data displayed on the plot.
FSC-H (10^6)
FSC-A (10^6)
Sample51 / P1
P2
0.00%
There are no data displayed on the plot.
FSC-H (10^6)
SSC-H (10^6)
Sample51 / P1 / P2
R3
0.00%
There are no data displayed on the plot.
AmCyan-H
SSC-H (10^6)
Sample51 / P1 / P2 / R3
R4
0.00%
There are no data displayed on the plot.
PE-Texas Red-H
SSC-H (10^6)

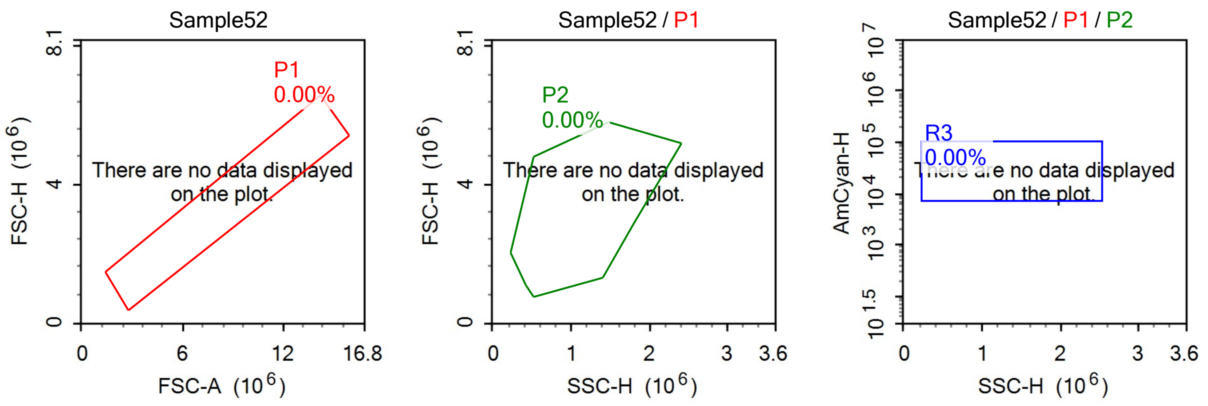

Sample52
P1
0.00%
There are no data displayed on the plot.
FSC-H (10^6)
FSC-A (10^6)
Sample52 / P1
P2
0.00%
There are no data displayed on the plot.
FSC-H (10^6)
SSC-H (10^6)
Sample52 / P1 / P2
R3
0.00%
There are no data displayed on the plot.
AmCyan-H
SSC-H (10^6)


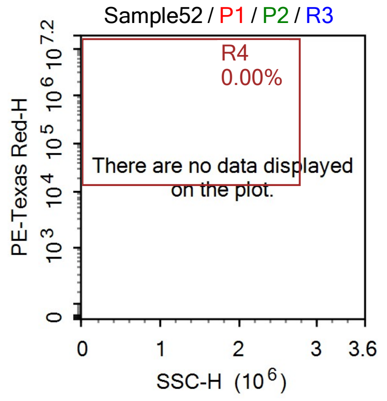

Sample52 / P1 / P2 / R3
R4
0.00%
There are no data displayed on the plot.
PE-Texas Red-H
SSC-H (10^6)

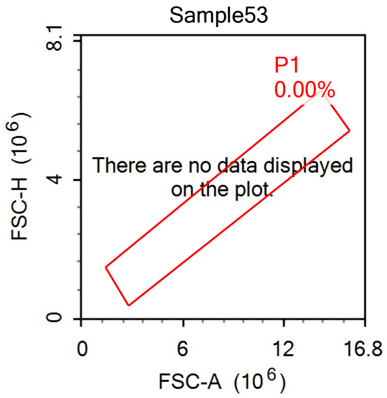
Sample53
P1
0.00%
There are no data displayed on the plot.
FSC-H (10^6)
FSC-A (10^6)
0
4
8.1
0
6
12
16.8

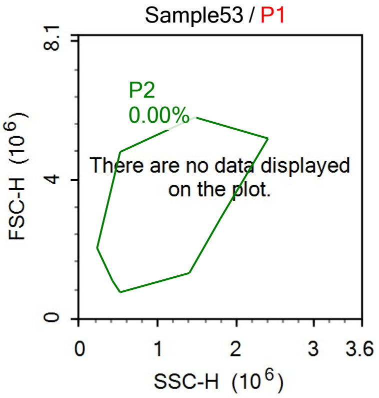
Sample53 / P1
P2
0.00%
There are no data displayed on the plot.
FSC-H (10^6)
SSC-H (10^6)
0
4
8.1
0
1
2
3
3.6

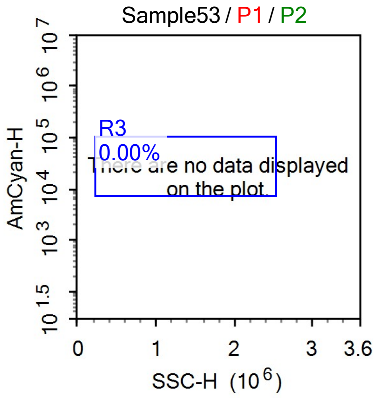
Sample53 / P1 / P2
R3
0.00%
There are no data displayed on the plot.
AmCyan-H
SSC-H (10^6)
0
1
2
3
3.6

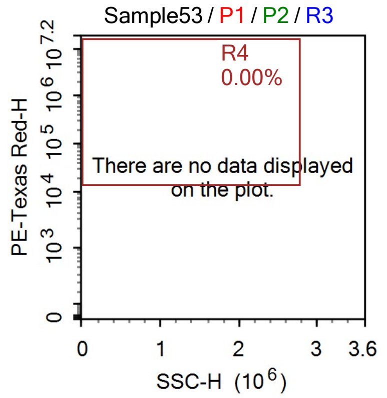
Sample53 / P1 / P2 / R3
R4
0.00%
There are no data displayed on the plot.
PE-Texas Red-H
SSC-H (10^6)
0
1
2
3
3.6

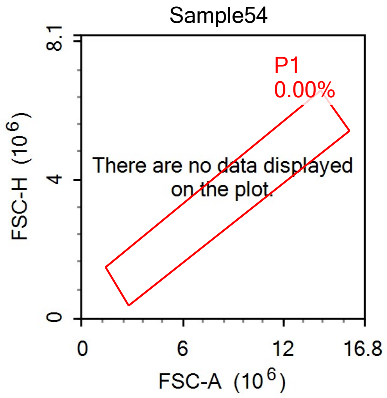
Sample54
P1
0.00%
There are no data displayed on the plot.
FSC-H (10^6)
FSC-A (10^6)
0
4
8.1
0
6
12
16.8

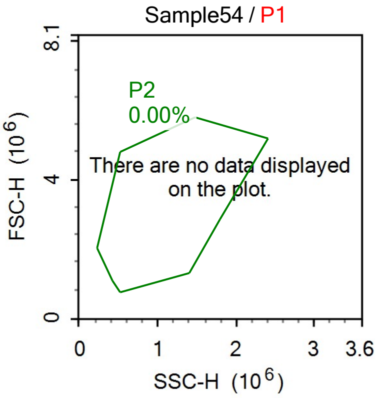
Sample54 / P1
P2
0.00%
There are no data displayed on the plot.
FSC-H (10^6)
SSC-H (10^6)
0
4
8.1
0
1
2
3
3.6

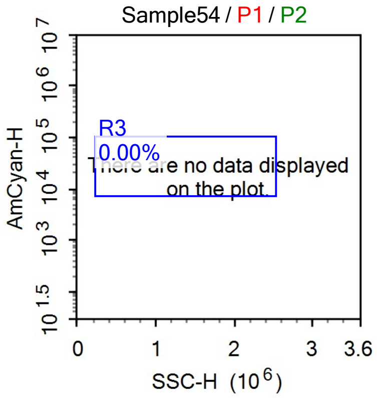
Sample54 / P1 / P2
R3
0.00%
There are no data displayed on the plot.
AmCyan-H
SSC-H (10^6)
10^1.5
10^3
10^4
10^5
10^6
10^7
0
1
2
3
3.6

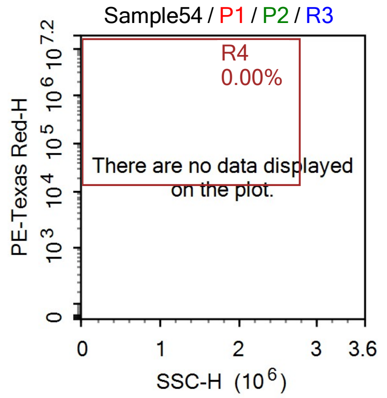
Sample54 / P1 / P2 / R3
R4
0.00%
There are no data displayed on the plot.
PE-Texas Red-H
SSC-H (10^6)
0
10^3
10^4
10^5
10^6
10^7.2
0
1
2
3
3.6

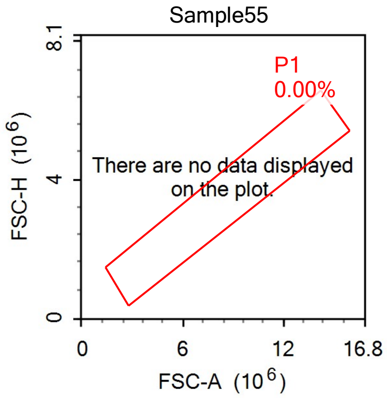
Sample55
P1
0.00%
There are no data displayed on the plot.
FSC-H (10^6)
FSC-A (10^6)
0
4
8.1
0
6
12
16.8

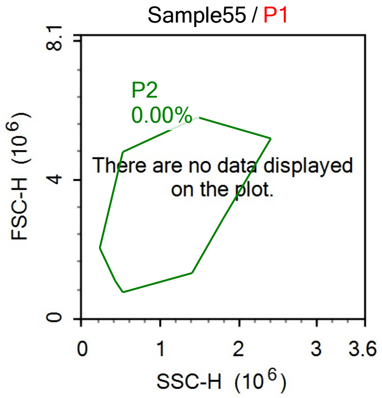
Sample55 / P1
P2
0.00%
There are no data displayed on the plot.
FSC-H (10^6)
SSC-H (10^6)
0
4
8.1
0
1
2
3
3.6

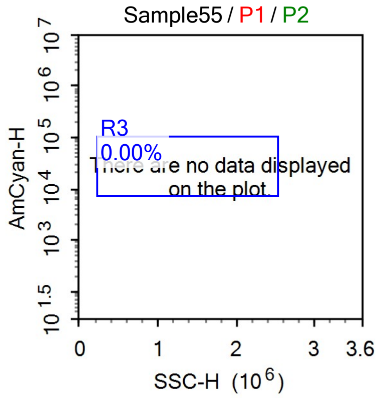
Sample55 / P1 / P2
R3
0.00%
There are no data displayed on the plot.
AmCyan-H
SSC-H (10^6)
10^1.5
10^3
10^4
10^5
10^6
10^7
0
1
2
3
3.6

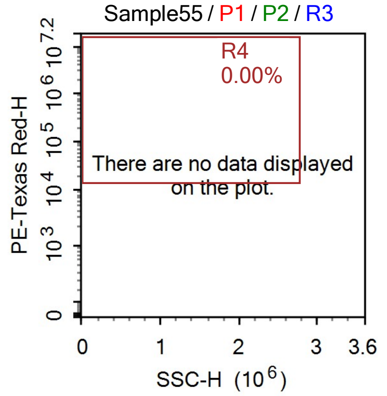
Sample55 / P1 / P2 / R3
R4
0.00%
There are no data displayed on the plot.
PE-Texas Red-H
SSC-H (10^6)
0
10^3
10^4
10^5
10^6
10^7.2
0
1
2
3
3.6

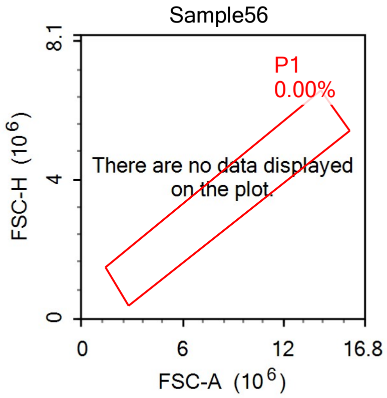
Sample56
P1
0.00%
There are no data displayed on the plot.
FSC-H (10^6)
FSC-A (10^6)
0
4
8.1
0
6
12
16.8

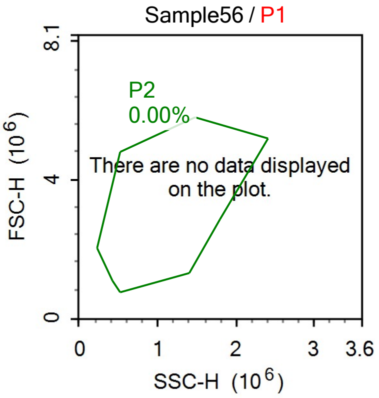
Sample56 / P1
P2
0.00%
There are no data displayed on the plot.
FSC-H (10^6)
SSC-H (10^6)
0
4
8.1
0
1
2
3
3.6

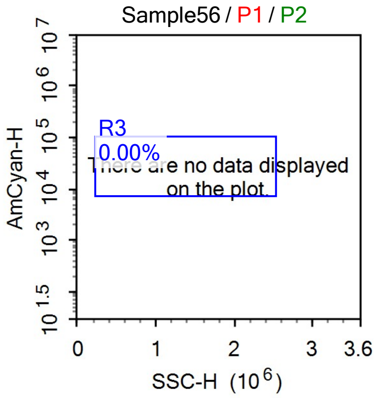
Sample56 / P1 / P2
R3
0.00%
There are no data displayed on the plot.
AmCyan-H
SSC-H (10^6)
10^1.5
10^3
10^4
10^5
10^6
10^7
0
1
2
3
3.6

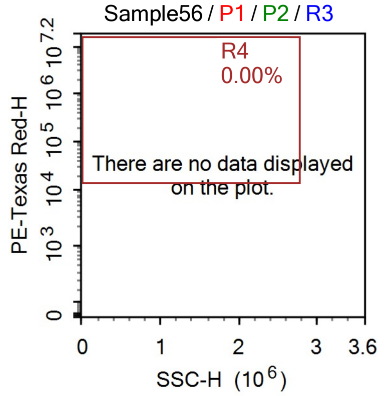
Sample56 / P1 / P2 / R3
R4
0.00%
There are no data displayed on the plot.
PE-Texas Red-H
SSC-H (10^6)
0
10^3
10^4
10^5
10^6
10^7.2
0
1
2
3
3.6

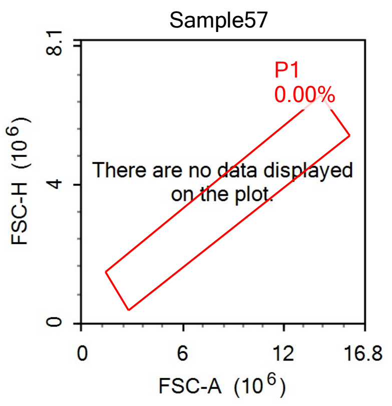
Sample57
P1
0.00%
There are no data displayed on the plot.
FSC-H (10^6)
FSC-A (10^6)
0
4
8.1
6
12
16.8

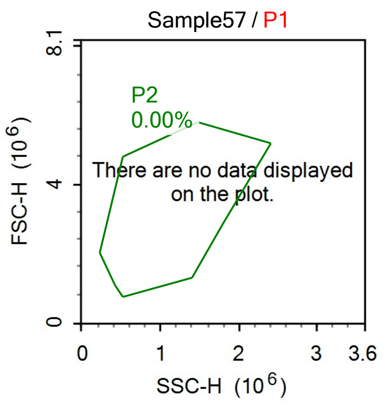
Sample57 / P1
P2
0.00%
There are no data displayed on the plot.
FSC-H (10^6)
SSC-H (10^6)
0
4
8.1
1
2
3
3.6

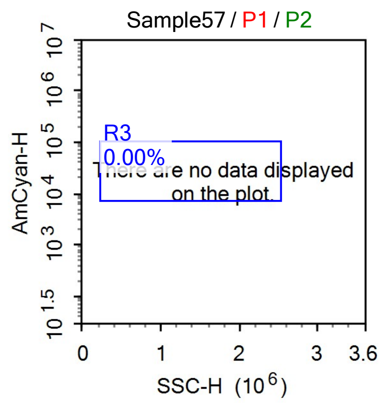
Sample57 / P1 / P2
R3
0.00%
There are no data displayed on the plot.
AmCyan-H
SSC-H (10^6)
0
1
2
3
3.6

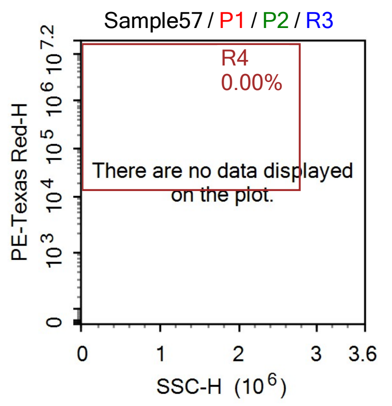
Sample57 / P1 / P2 / R3
R4
0.00%
There are no data displayed on the plot.
PE-Texas Red-H
SSC-H (10^6)
0
1
2
3
3.6

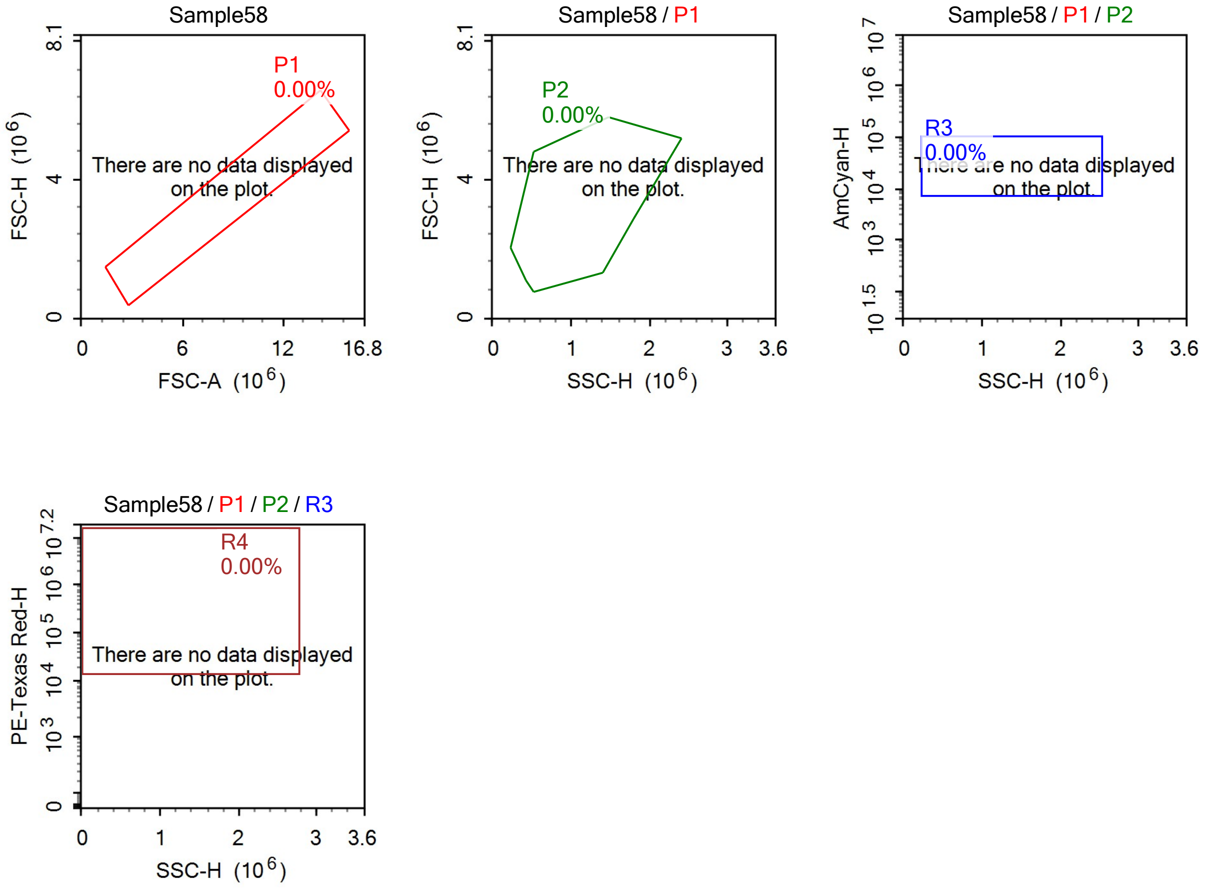

Sample58
P1
0.00%
There are no data displayed on the plot.
FSC-H (10^6)
FSC-A (10^6)
Sample58 / P1
P2
0.00%
There are no data displayed on the plot.
FSC-H (10^6)
SSC-H (10^6)
Sample58 / P1 / P2
R3
0.00%
AmCyan-H
SSC-H (10^6)
Sample58 / P1 / P2 / R3
R4
0.00%
There are no data displayed on the plot.
PE-Texas Red-H
SSC-H (10^6)

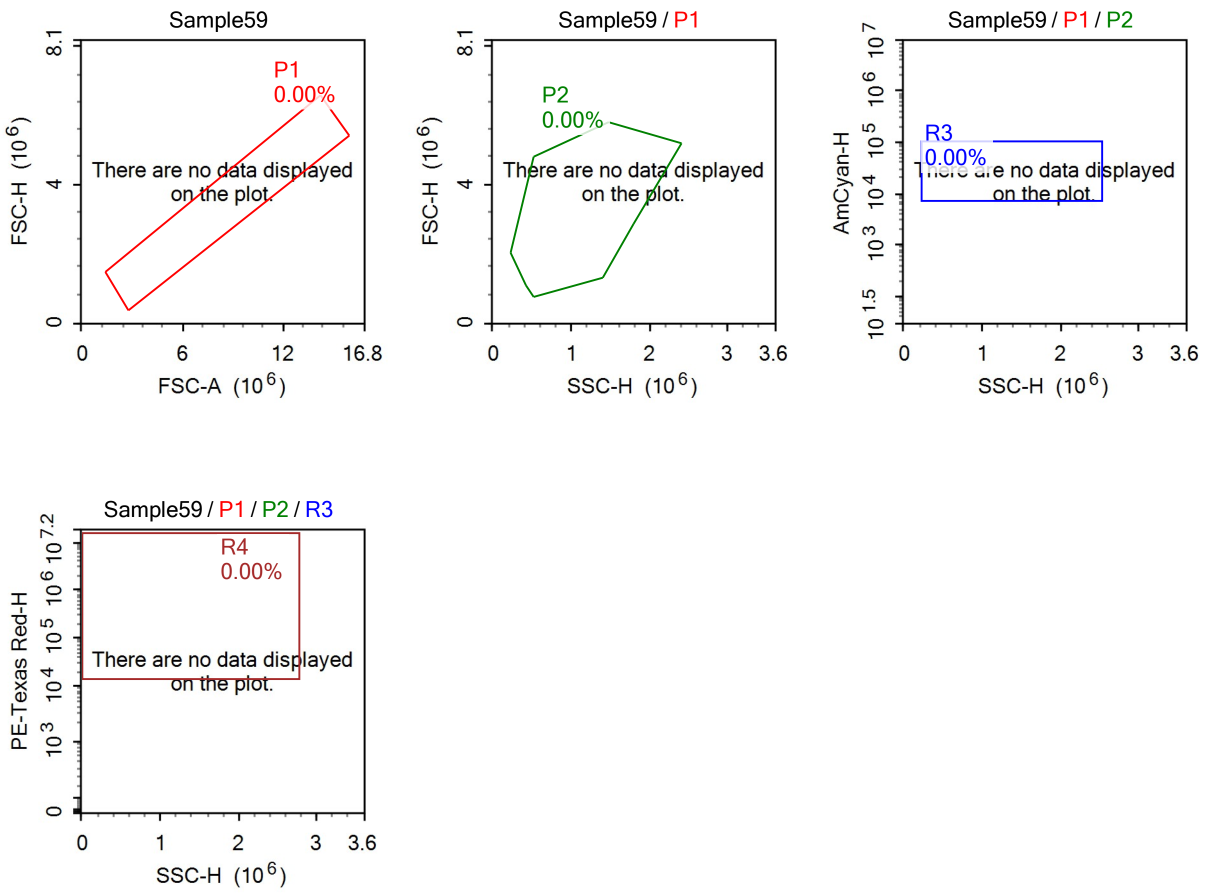
Sample59
P1
0.00%
There are no data displayed on the plot.
FSC-H (10^6)
FSC-A (10^6)
Sample59 / P1
P2
0.00%
There are no data displayed on the plot.
FSC-H (10^6)
SSC-H (10^6)
Sample59 / P1 / P2
R3
0.00%
There are no data displayed on the plot.
AmCyan-H
SSC-H (10^6)
Sample59 / P1 / P2 / R3
R4
0.00%
There are no data displayed on the plot.
PE-Texas Red-H
SSC-H (10^6)

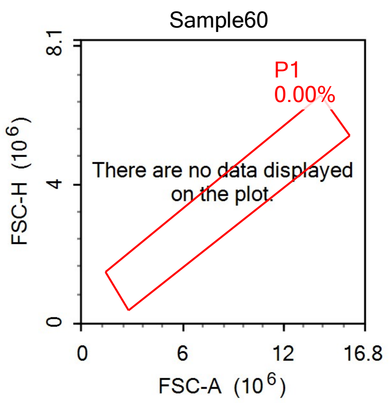
Sample60
P1
0.00%
There are no data displayed on the plot.
FSC-H (10^6)
FSC-A (10^6)
0
4
8.1
0
6
12
16.8

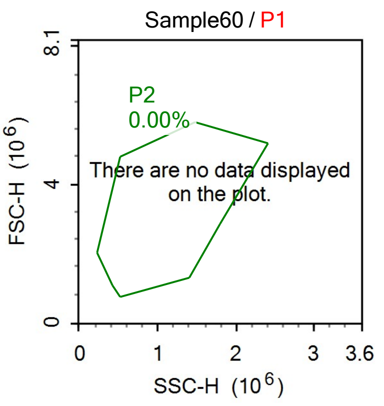
Sample60 / P1
P2
0.00%
There are no data displayed on the plot.
FSC-H (10^6)
SSC-H (10^6)
0
4
8.1
0
1
2
3
3.6

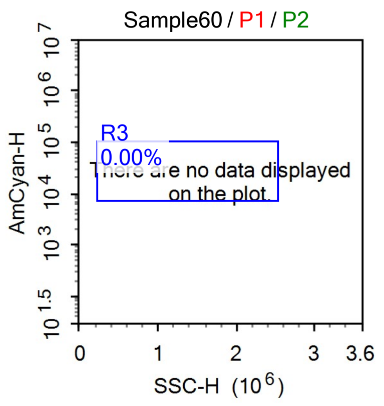
Sample60 / P1 / P2
R3
0.00%
There are no data displayed on the plot.
AmCyan-H
SSC-H (10^6)
0
1
2
3
3.6

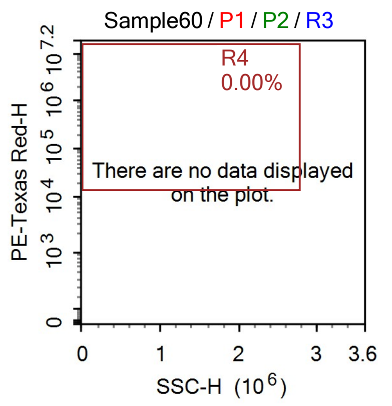
Sample60 / P1 / P2 / R3
R4
0.00%
There are no data displayed on the plot.
PE-Texas Red-H
SSC-H (10^6)
0
0
1
2
3
3.6

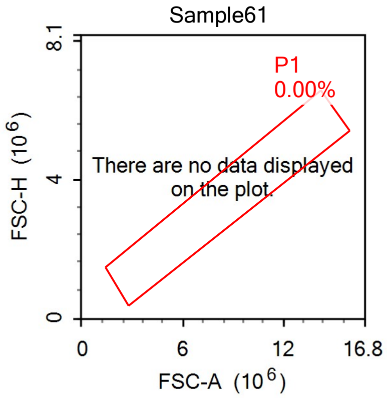
Sample61
P1
0.00%
There are no data displayed on the plot.
FSC-H ($10^6$)
FSC-A ($10^6$)
0
4
8.1
0
6
12
16.8

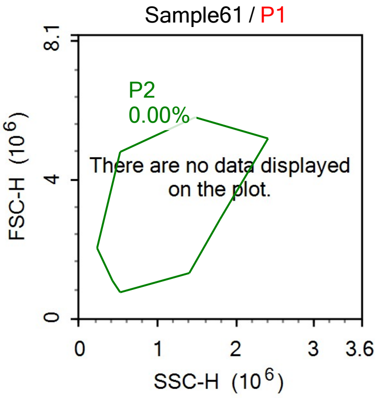
Sample61 / P1
P2
0.00%
There are no data displayed on the plot.
FSC-H ($10^6$)
SSC-H ($10^6$)
0
4
8.1
0
1
2
3
3.6

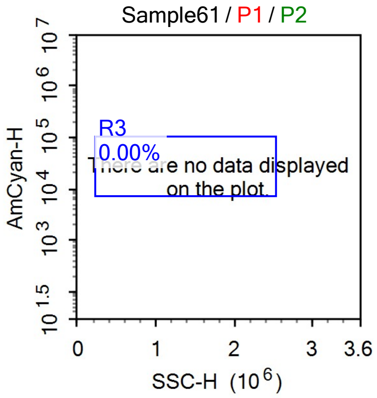
Sample61 / P1 / P2
R3
0.00%
There are no data displayed on the plot.
AmCyan-H
SSC-H ($10^6$)
$10^{1.5}$
$10^3$
$10^4$
$10^5$
$10^6$
$10^7$
0
1
2
3
3.6

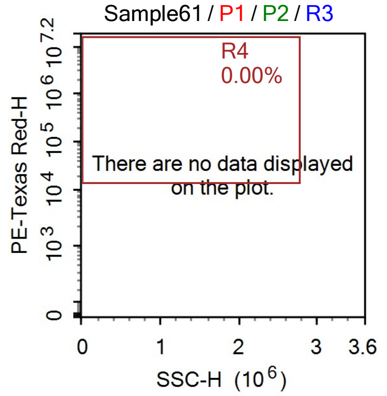
Sample61 / P1 / P2 / R3
R4
0.00%
There are no data displayed on the plot.
PE-Texas Red-H
SSC-H ($10^6$)
0
$10^3$
$10^4$
$10^5$
$10^6$
$10^{7.2}$
0
1
2
3
3.6

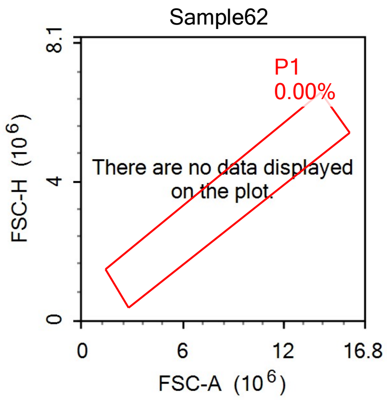
Sample62
P1
0.00%
There are no data displayed on the plot.
FSC-H (10^6)
FSC-A (10^6)

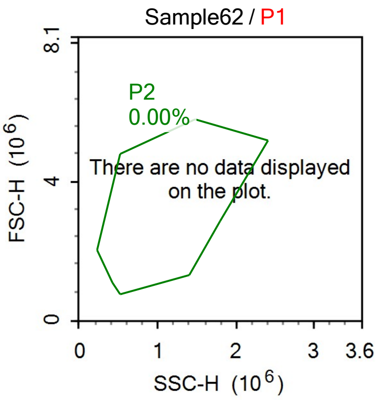
Sample62 / P1
P2
0.00%
There are no data displayed on the plot.
FSC-H (10^6)
SSC-H (10^6)

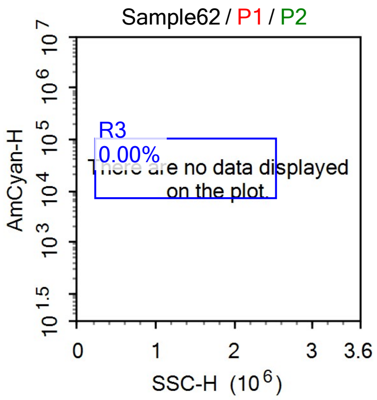
Sample62 / P1 / P2
R3
0.00%
There are no data displayed on the plot.
AmCyan-H
SSC-H (10^6)

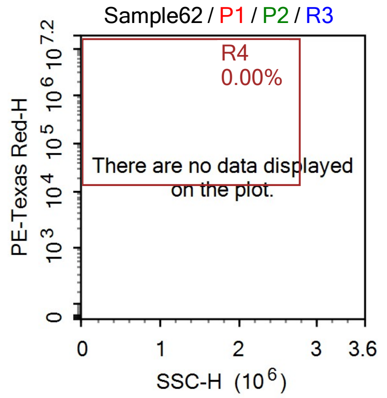
Sample62 / P1 / P2 / R3
R4
0.00%
There are no data displayed on the plot.
PE-Texas Red-H
SSC-H (10^6)

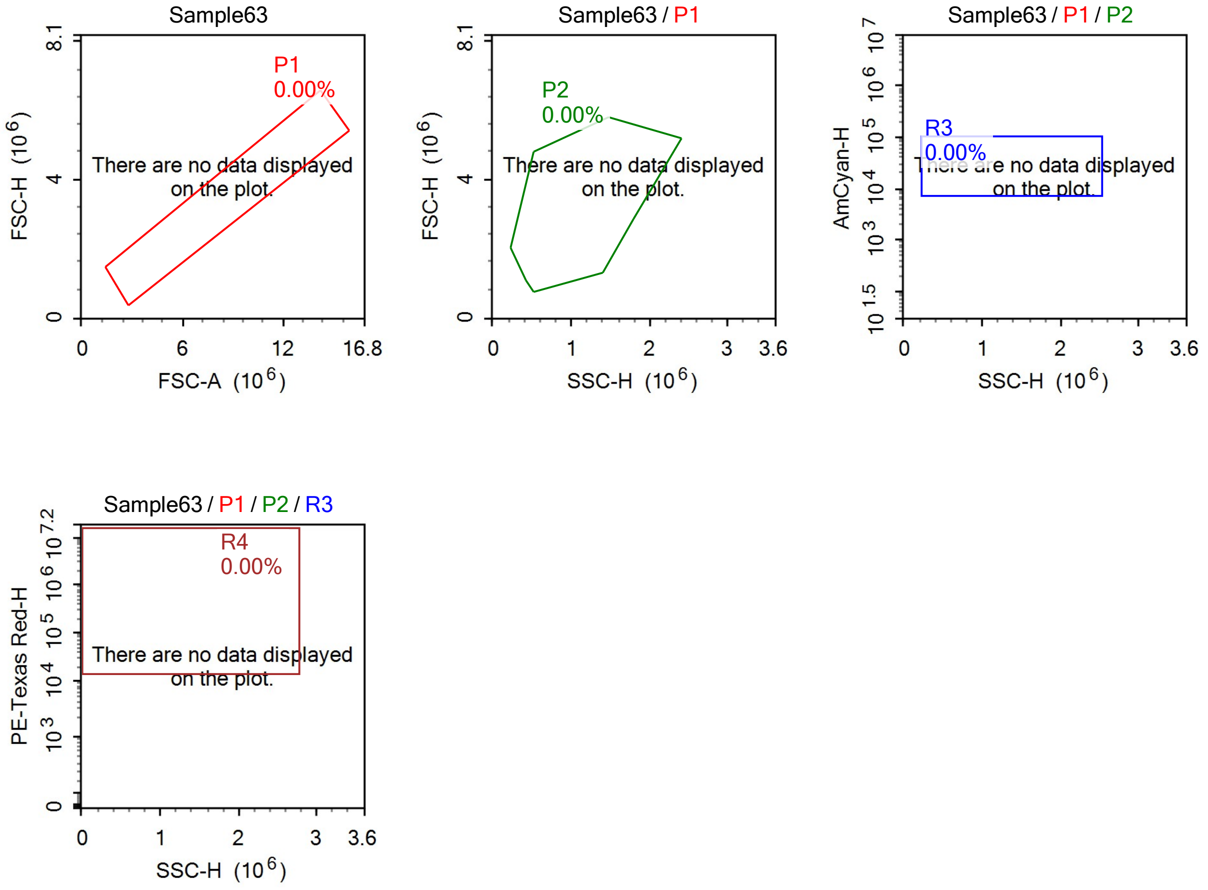
Sample63
P1
0.00%
There are no data displayed on the plot.
FSC-H (10^6)
FSC-A (10^6)
Sample63 / P1
P2
0.00%
There are no data displayed on the plot.
FSC-H (10^6)
SSC-H (10^6)
Sample63 / P1 / P2
R3
0.00%
There are no data displayed on the plot.
AmCyan-H
SSC-H (10^6)
Sample63 / P1 / P2 / R3
R4
0.00%
There are no data displayed on the plot.
PE-Texas Red-H
SSC-H (10^6)

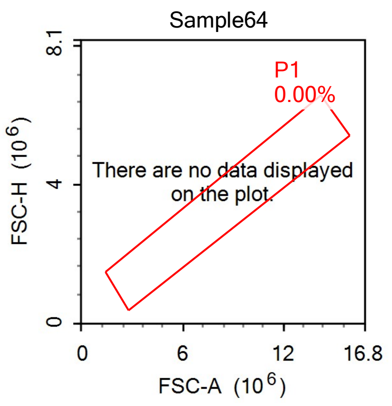
Sample64
P1
0.00%
There are no data displayed on the plot.
FSC-H (10^6)
FSC-A (10^6)
0
6
12
16.8
4
8.1

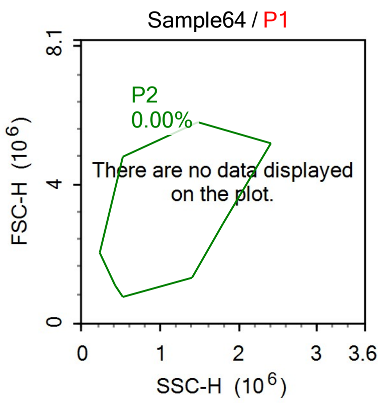
Sample64 / P1
P2
0.00%
There are no data displayed on the plot.
FSC-H (10^6)
SSC-H (10^6)
0
1
2
3
3.6
4
8.1

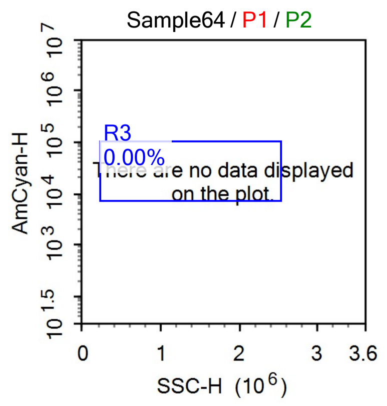
Sample64 / P1 / P2
R3
0.00%
There are no data displayed on the plot.
AmCyan-H
SSC-H (10^6)
0
1
2
3
3.6

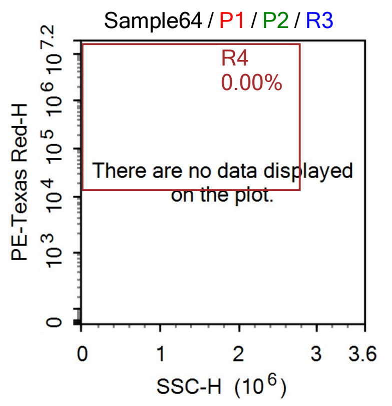
Sample64 / P1 / P2 / R3
R4
0.00%
There are no data displayed on the plot.
PE-Texas Red-H
SSC-H (10^6)
0
1
2
3
3.6
