## Supplementary material for "Vitamin B_2_ Production by Vaginal Lactobacilli Promotes Symbiosis": Data S10: MR1 expression on VK2E6E7(.hMR1) (attempt 2, gating per cell line).pdf

### Report of Specimen9

Specimen Name: Specimen9 | Run Time: 19-Mar-25 3:49 PM

Cytometer: NovoCyte Quanteon 621181110427 | Software: NovoExpress 1.6.0

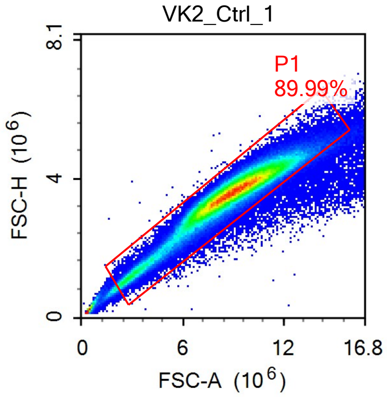


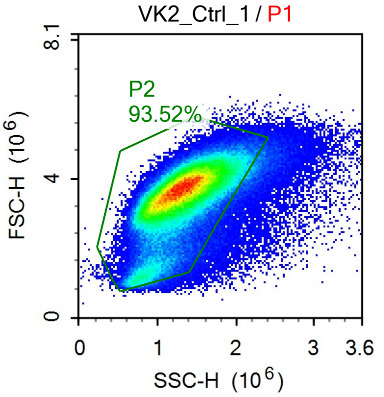


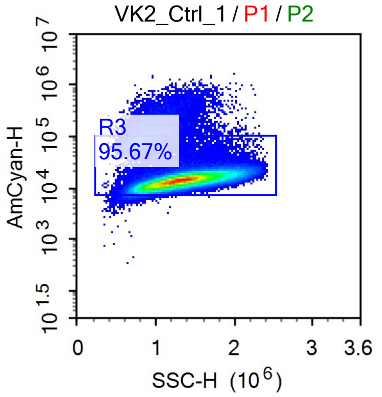


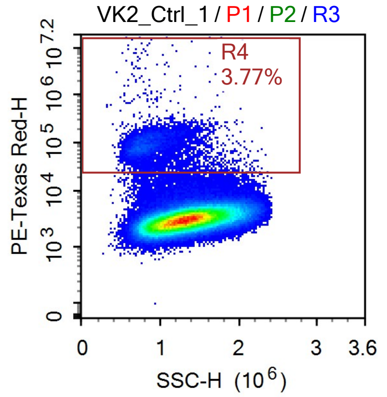


Sample Statistics of VK2_Ctrl_1

| Gate | Count | % Parent | X | Y | Median X | Median Y |
|---|---|---|---|---|---|---|
| All | 236,430 | | | | | |
| P1 | 212,752 | 89.99% | FSC-A | FSC-H | 8,879,873 | 3,587,677 |
| P2 | 198,959 | 93.52% | SSC-H | FSC-H | 1,302,124 | 3,575,605 |
| R3 | 190,352 | 95.67% | SSC-H | AmCyan-H | 1,314,958 | 14,195 |
| R4 | 7,181 | 3.77% | SSC-H | PE-Texas Red-H | 854,772 | 88,679 |

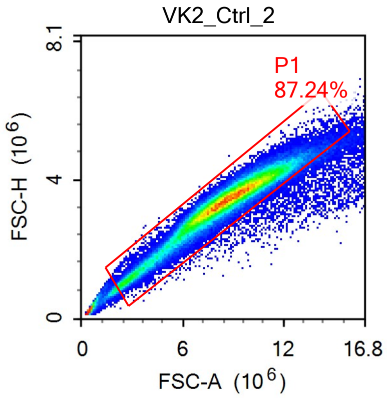


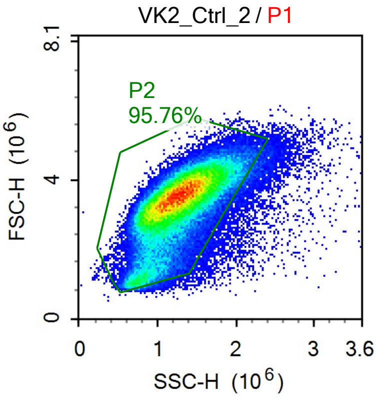


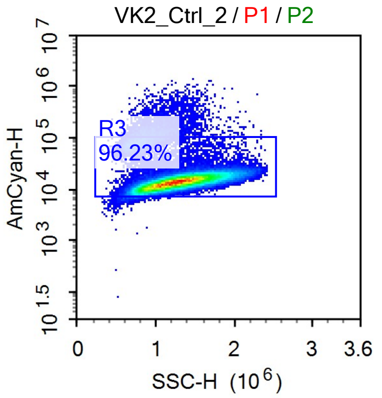

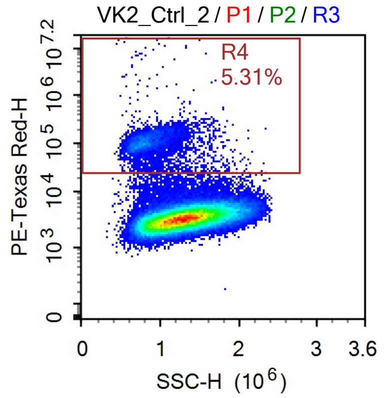


Sample Statistics of VK2_Ctrl_2

| Gate | Count | % Parent | X | Y | Median X | Median Y |
|---|---|---|---|---|---|---|
| All | 114,302 | | | | | |
| └ P1 | 99,716 | 87.24% | FSC-A | FSC-H | 8,551,798 | 3,407,710 |
| └ P2 | 95,490 | 95.76% | SSC-H | FSC-H | 1,256,015 | 3,400,918 |
| └ R3 | 91,893 | 96.23% | SSC-H | AmCyan-H | 1,267,227 | 14,111 |
| └ R4 | 4,884 | 5.31% | SSC-H | PE-Texas Red-H | 835,463 | 104,330 |

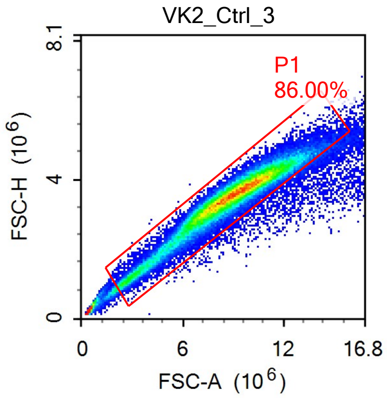


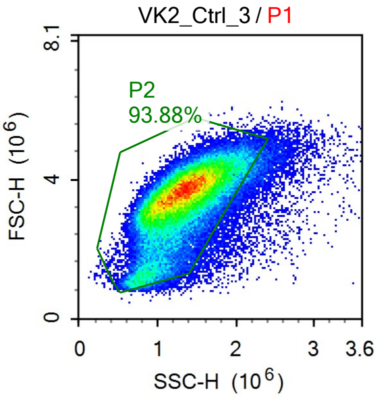


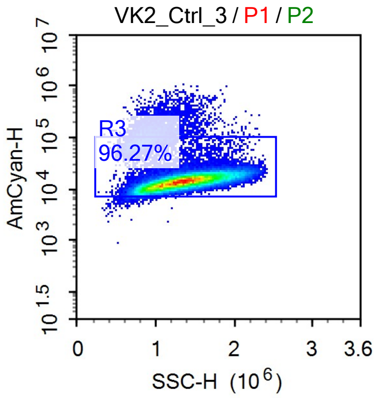


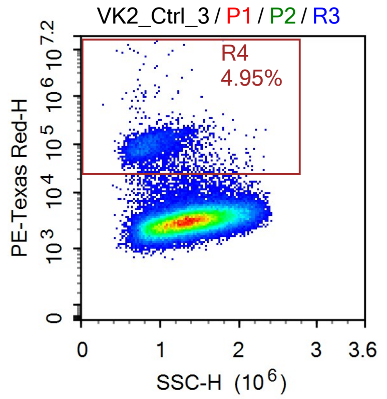

Sample Statistics of VK2_Ctrl_3

| Gate | Count | % Parent | X | Y | Median X | Median Y |
|---|---|---|---|---|---|---|
| All | 88,387 | | | | | |
| P1 | 76,013 | 86.00% | FSC-A | FSC-H | 8,865,887 | 3,522,486 |
| P2 | 71,362 | 93.88% | SSC-H | FSC-H | 1,314,128 | 3,511,782 |
| R3 | 68,701 | 96.27% | SSC-H | AmCyan-H | 1,325,149 | 14,125 |
| R4 | 3,400 | 4.95% | SSC-H | PE-Texas Red-H | 875,729 | 93,993 |

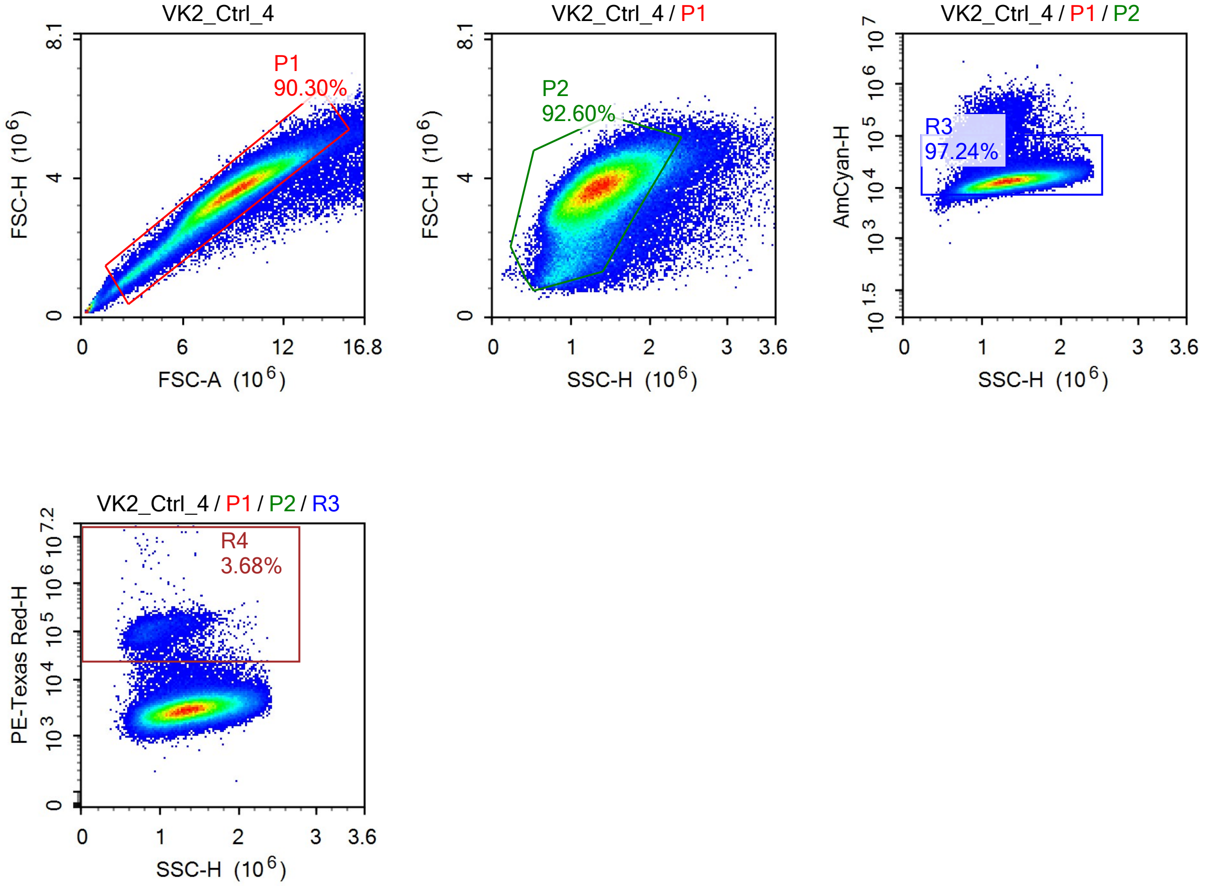


Sample Statistics of VK2_Ctrl_4

| Gate | Count | % Parent | X | Y | Median X | Median Y |
|---|---|---|---|---|---|---|
| All | 166,371 | | | | | |
| P1 | 150,229 | 90.30% | FSC-A | FSC-H | 8,951,205 | 3,596,502 |
| P2 | 139,117 | 92.60% | SSC-H | FSC-H | 1,334,975 | 3,585,551 |
| R3 | 135,273 | 97.24% | SSC-H | AmCyan-H | 1,342,082 | 13,339 |
| R4 | 4,974 | 3.68% | SSC-H | PE-Texas Red-H | 950,596 | 103,154 |

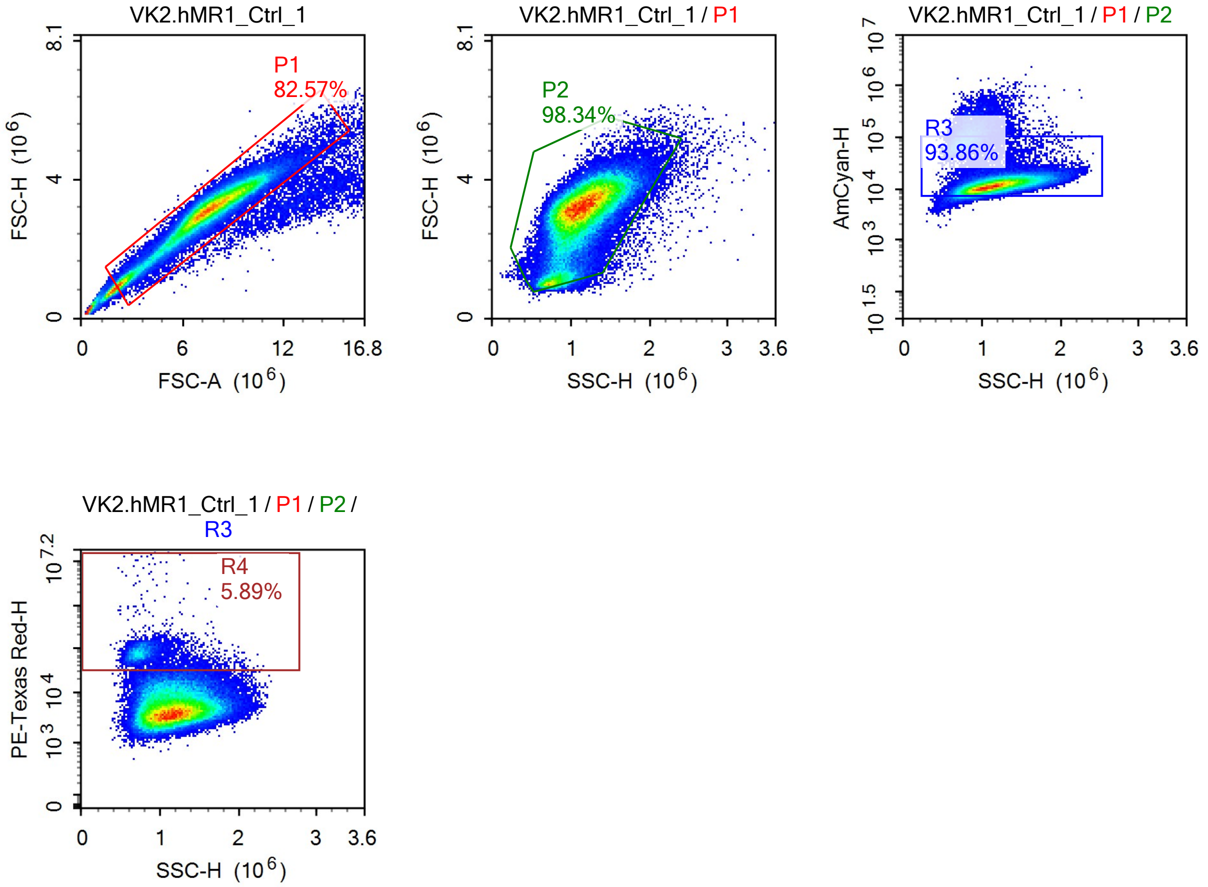


Sample Statistics of VK2.hMR1_Ctrl_1

| Gate | Count | % Parent | X | Y | Median X | Median Y |
|---|---|---|---|---|---|---|
| All | 110,820 | | | | | |
| P1 | 91,504 | 82.57% | FSC-A | FSC-H | 7,464,061 | 3,110,558 |
| P2 | 89,983 | 98.34% | SSC-H | FSC-H | 1,118,753 | 3,112,641 |
| R3 | 84,460 | 93.86% | SSC-H | AmCyan-H | 1,136,985 | 11,837 |
| R4 | 4,974 | 5.89% | SSC-H | PE-Texas Red-H | 788,171 | 74,291 |

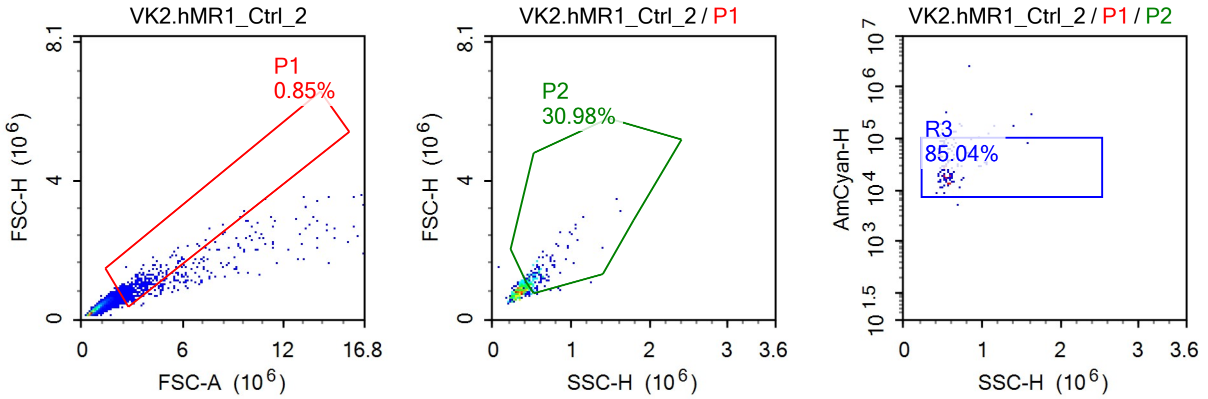

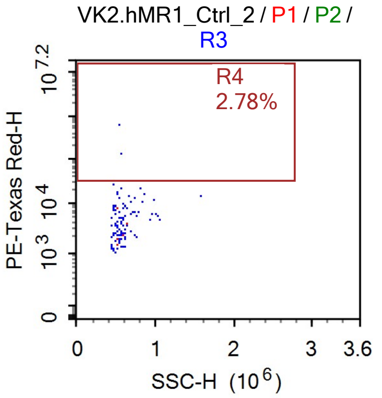


Sample Statistics of VK2.hMR1_Ctrl_2

| Gate | Count | % Parent | X | Y | Median X | Median Y |
|---|---|---|---|---|---|---|
| All | 48,470 | | | | | |
| P1 | 410 | 0.85% | FSC-A | FSC-H | 2,996,505 | 871,143 |
| P2 | 127 | 30.98% | SSC-H | FSC-H | 552,866 | 1,258,282 |
| R3 | 108 | 85.04% | SSC-H | AmCyan-H | 540,164 | 29,515 |
| R4 | 3 | 2.78% | SSC-H | PE-Texas Red-H | 541,510 | 452,676 |

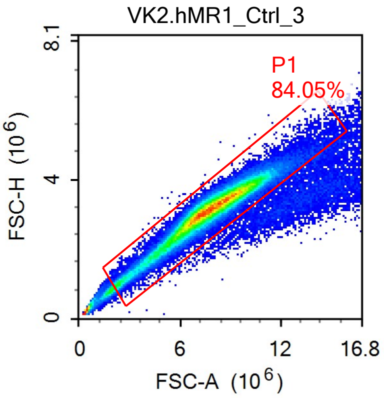


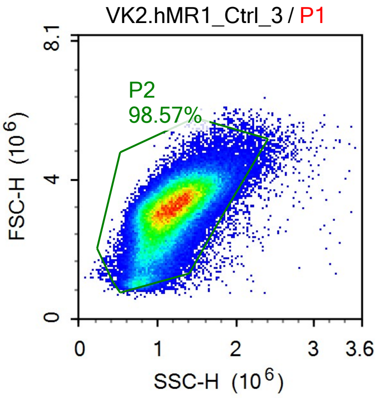


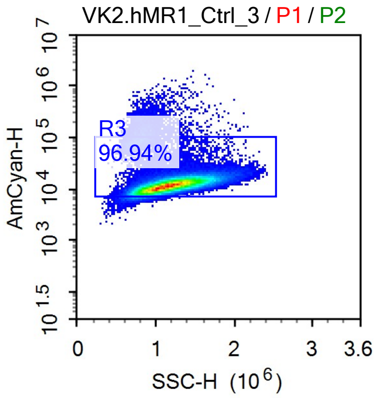


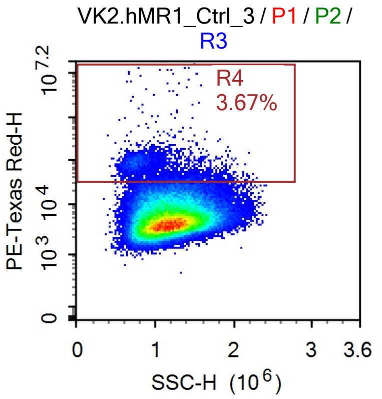

Sample Statistics of VK2.hMR1_Ctrl_3

| Gate | Count | % Parent | X | Y | Median X | Median Y |
|---|---|---|---|---|---|---|
| All | 120,517 | | | | | |
| P1 | 101,294 | 84.05% | FSC-A | FSC-H | 7,684,813 | 3,129,962 |
| P2 | 99,841 | 98.57% | SSC-H | FSC-H | 1,157,231 | 3,128,508 |
| R3 | 96,789 | 96.94% | SSC-H | AmCyan-H | 1,167,389 | 11,914 |
| R4 | 3,549 | 3.67% | SSC-H | PE-Texas Red-H | 942,784 | 69,466 |

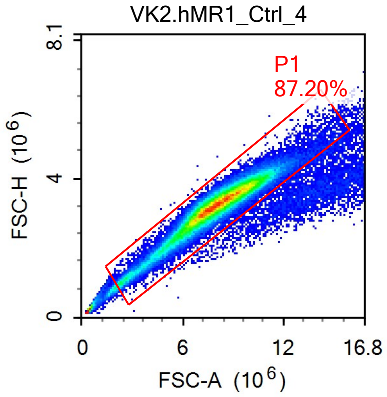


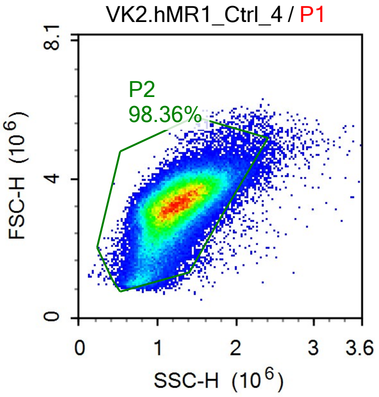


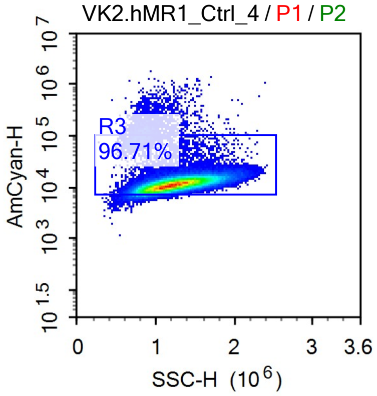


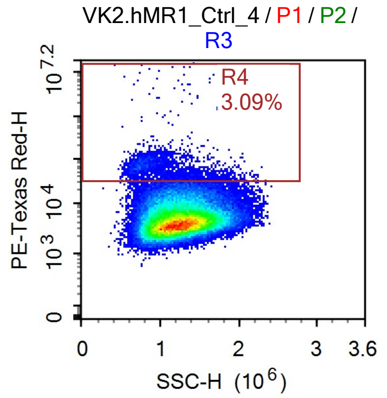


Sample Statistics of VK2.hMR1_Ctrl_4

| Gate | Count | % Parent | X | Y | Median X | Median Y |
|---|---|---|---|---|---|---|
| All | 95,119 | | | | | |
| P1 | 82,943 | 87.20% | FSC-A | FSC-H | 7,940,226 | 3,239,076 |
| P2 | 81,581 | 98.36% | SSC-H | FSC-H | 1,213,323 | 3,234,310 |
| R3 | 78,898 | 96.71% | SSC-H | AmCyan-H | 1,224,096 | 11,485 |
| R4 | 2,436 | 3.09% | SSC-H | PE-Texas Red-H | 960,324 | 62,378 |

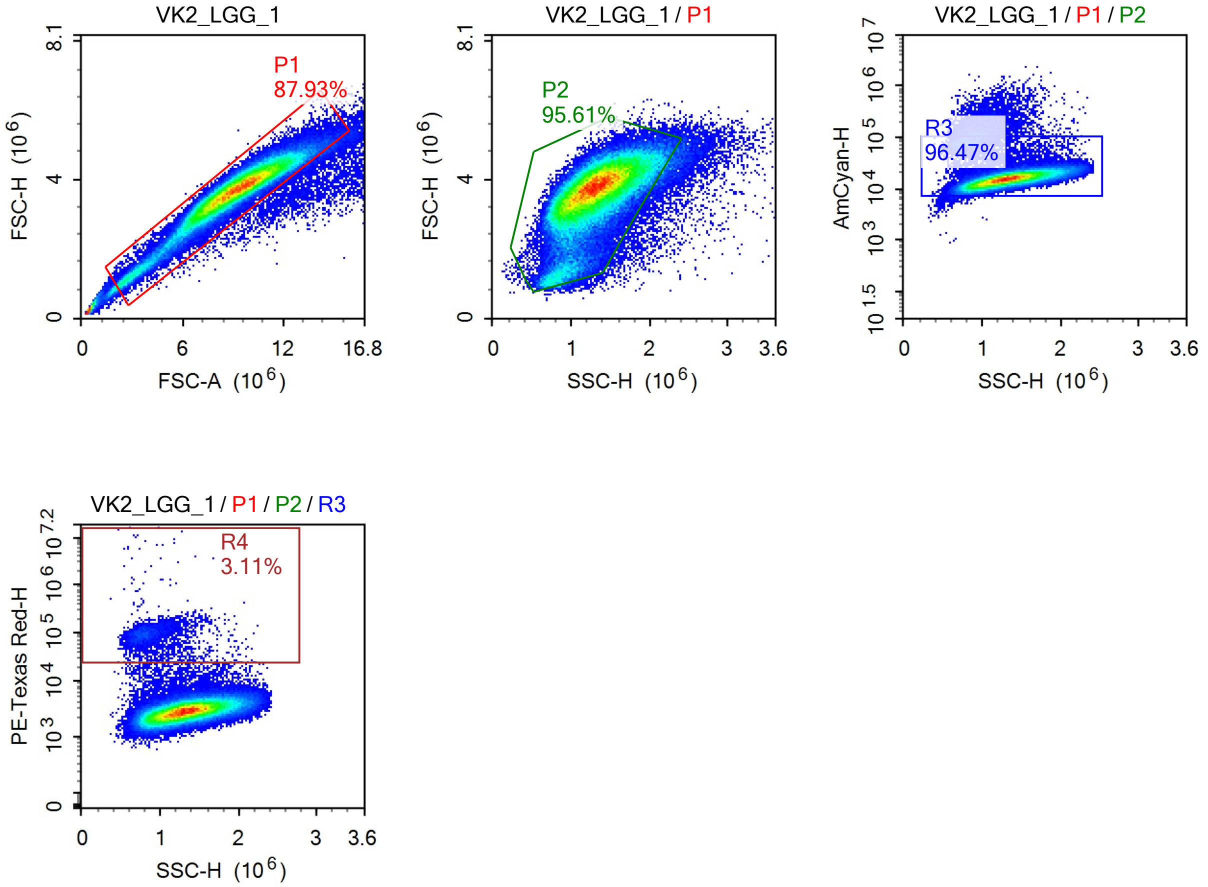


Sample Statistics of VK2_LGG_1

| Gate | Count | % Parent | X | Y | Median X | Median Y |
|---|---|---|---|---|---|---|
| All | 122,147 | | | | | |
| P1 | 107,409 | 87.93% | FSC-A | FSC-H | 9,145,303 | 3,703,138 |
| P2 | 102,699 | 95.61% | SSC-H | FSC-H | 1,298,473 | 3,688,976 |
| R3 | 99,077 | 96.47% | SSC-H | AmCyan-H | 1,309,038 | 15,651 |
| R4 | 3,080 | 3.11% | SSC-H | PE-Texas Red-H | 845,145 | 91,949 |

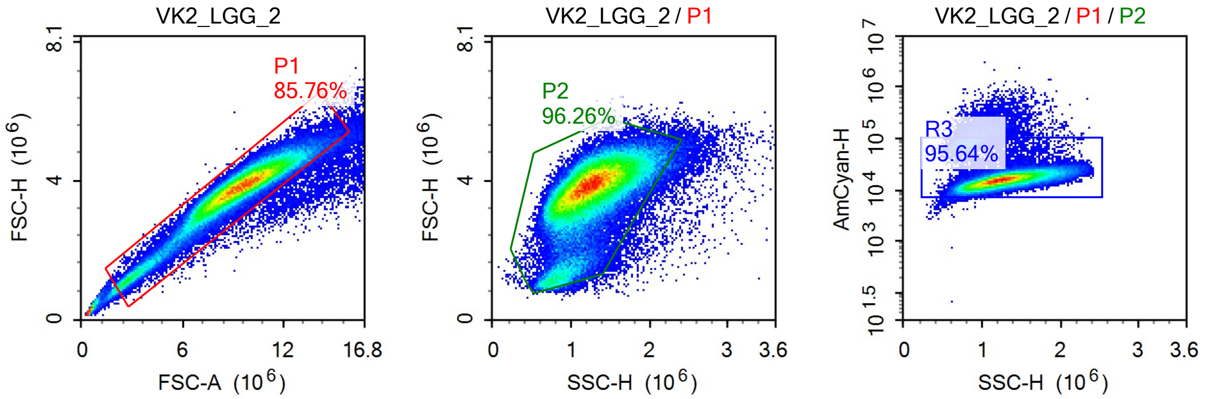

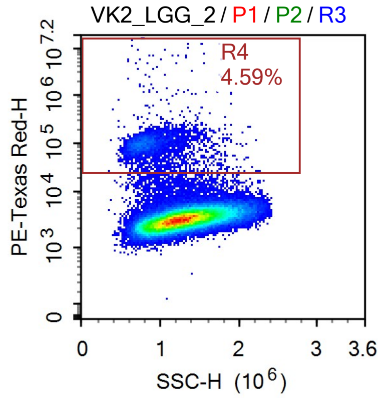


Sample Statistics of VK2_LGG_2

| Gate | Count | % Parent | X | Y | Median X | Median Y |
|---|---|---|---|---|---|---|
| All | 113,178 | | | | | |
| P1 | 97,060 | 85.76% | FSC-A | FSC-H | 9,320,344 | 3,790,748 |
| P2 | 93,432 | 96.26% | SSC-H | FSC-H | 1,242,899 | 3,782,672 |
| R3 | 89,359 | 95.64% | SSC-H | AmCyan-H | 1,253,649 | 15,911 |
| R4 | 4,104 | 4.59% | SSC-H | PE-Texas Red-H | 836,983 | 96,529 |

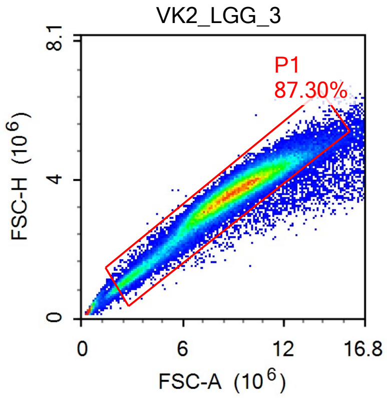


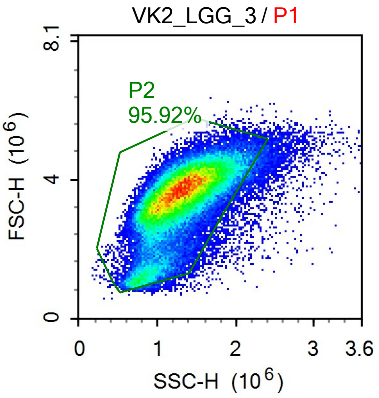


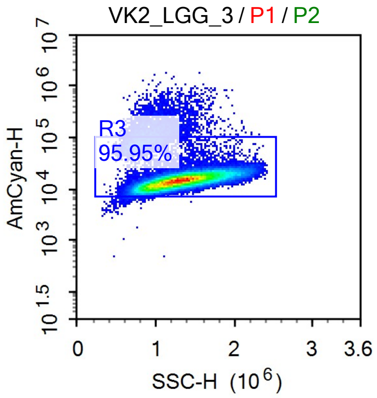


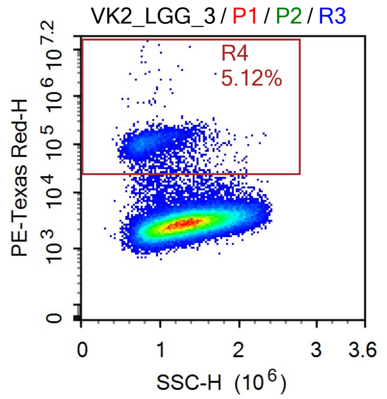

Sample Statistics of VK2_LGG_3

| Gate | Count | % Parent | X | Y | Median X | Median Y |
|---|---|---|---|---|---|---|
| All | 83,288 | | | | | |
| P1 | 72,712 | 87.30% | FSC-A | FSC-H | 8,859,694 | 3,593,505 |
| P2 | 69,744 | 95.92% | SSC-H | FSC-H | 1,279,219 | 3,581,043 |
| R3 | 66,916 | 95.95% | SSC-H | AmCyan-H | 1,291,268 | 14,799 |
| R4 | 3,427 | 5.12% | SSC-H | PE-Texas Red-H | 856,218 | 108,660 |

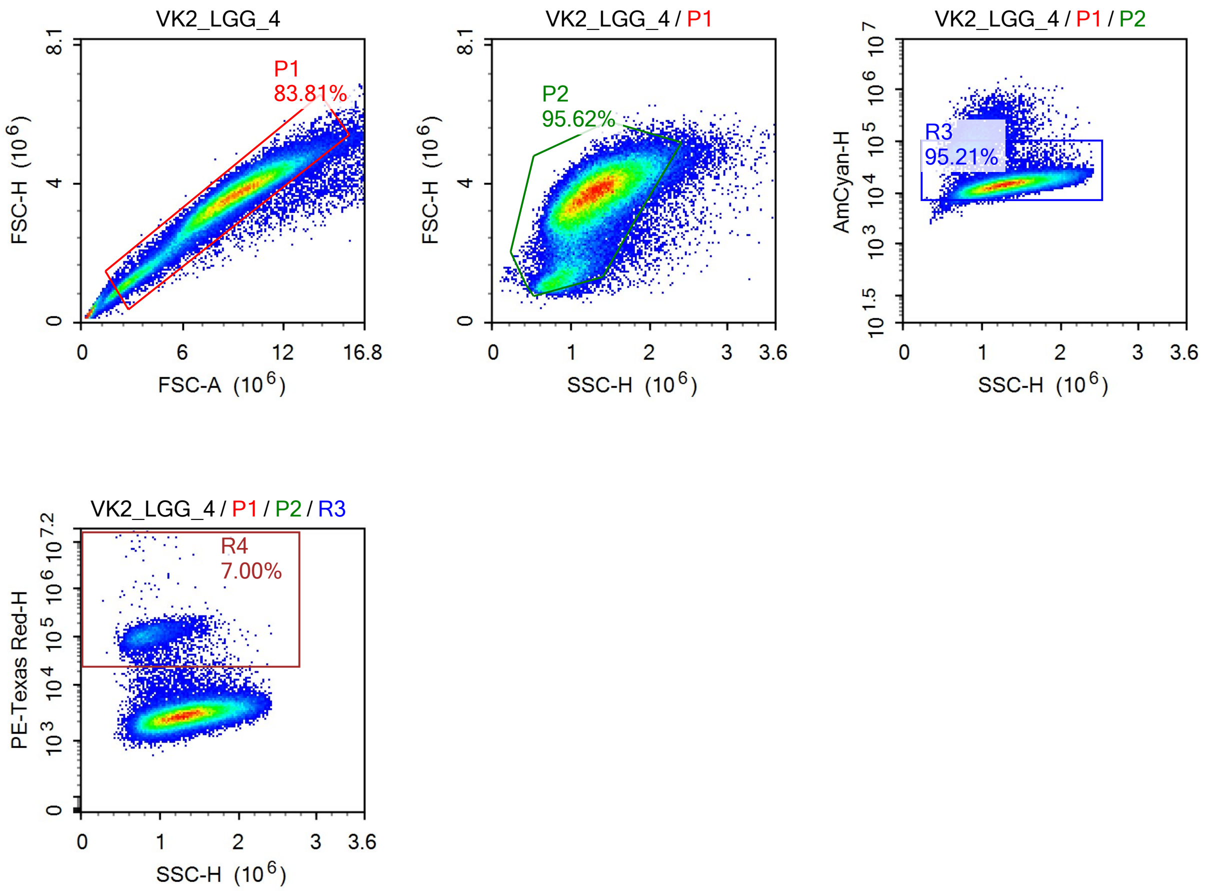


Sample Statistics of VK2_LGG_4

| Gate | Count | % Parent | X | Y | Median X | Median Y |
|---|---|---|---|---|---|---|
| All | 83,338 | | | | | |
| P1 | 69,846 | 83.81% | FSC-A | FSC-H | 8,842,944 | 3,600,152 |
| P2 | 66,789 | 95.62% | SSC-H | FSC-H | 1,262,145 | 3,591,099 |
| R3 | 63,587 | 95.21% | SSC-H | AmCyan-H | 1,277,723 | 14,730 |
| R4 | 4,448 | 7.00% | SSC-H | PE-Texas Red-H | 855,166 | 106,400 |

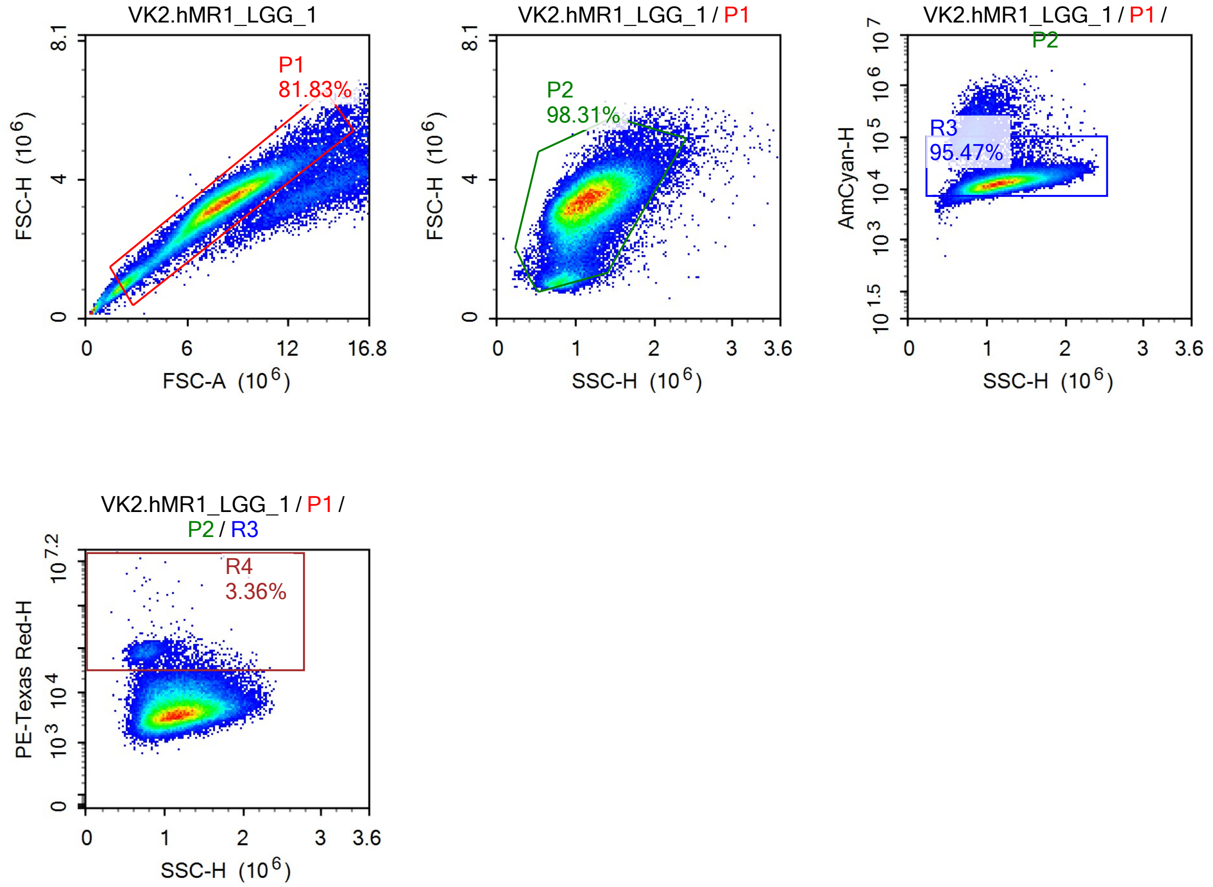


Sample Statistics of VK2.hMR1_LGG_1

| Gate | Count | % Parent | X | Y | Median X | Median Y |
|---|---|---|---|---|---|---|
| All | 86,842 | | | | | |
| P1 | 71,060 | 81.83% | FSC-A | FSC-H | 8,054,100 | 3,327,143 |
| P2 | 69,861 | 98.31% | SSC-H | FSC-H | 1,144,386 | 3,325,115 |
| R3 | 66,693 | 95.47% | SSC-H | AmCyan-H | 1,157,133 | 13,203 |
| R4 | 2,240 | 3.36% | SSC-H | PE-Texas Red-H | 821,768 | 77,958 |

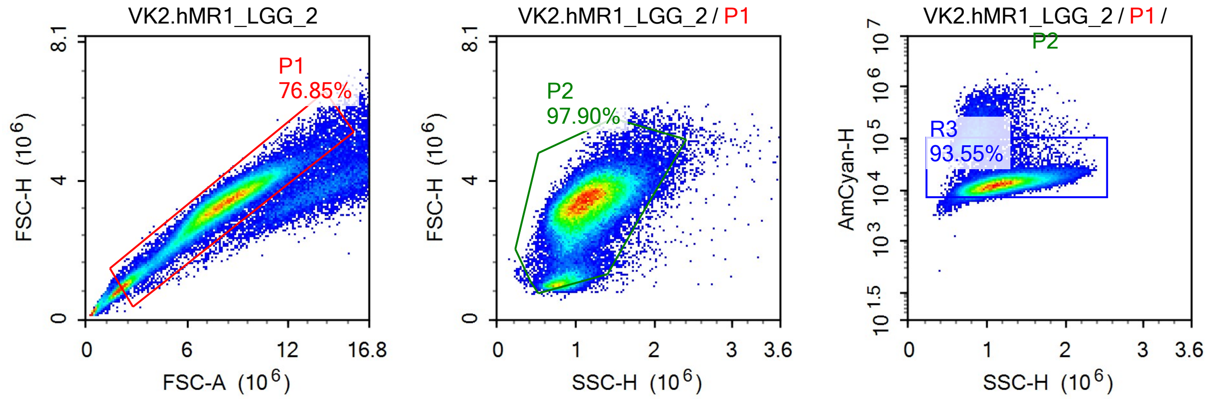

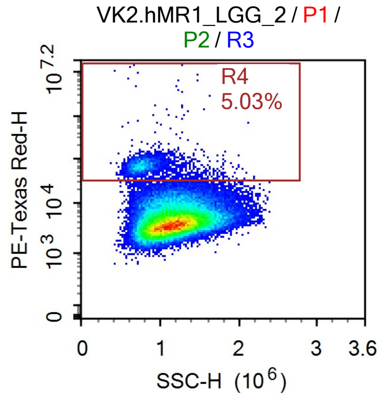


Sample Statistics of VK2.hMR1_LGG_2

| Gate | Count | % Parent | X | Y | Median X | Median Y |
|---|---|---|---|---|---|---|
| All | 80,699 | | | | | |
| P1 | 62,016 | 76.85% | FSC-A | FSC-H | 8,005,536 | 3,302,596 |
| P2 | 60,716 | 97.90% | SSC-H | FSC-H | 1,118,008 | 3,302,219 |
| R3 | 56,799 | 93.55% | SSC-H | AmCyan-H | 1,135,033 | 13,244 |
| R4 | 2,859 | 5.03% | SSC-H | PE-Texas Red-H | 779,920 | 67,902 |

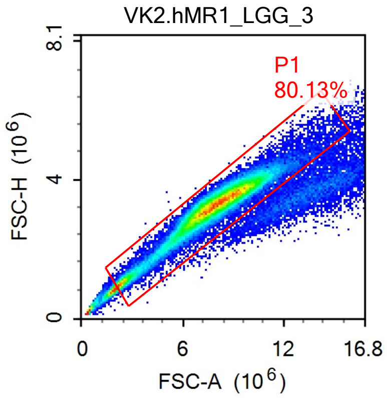


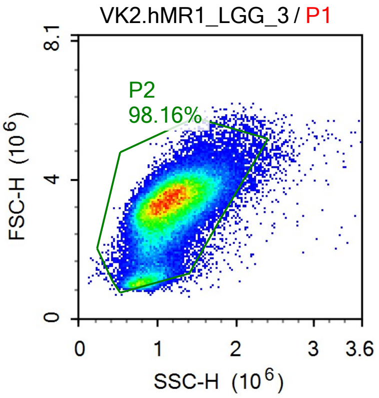


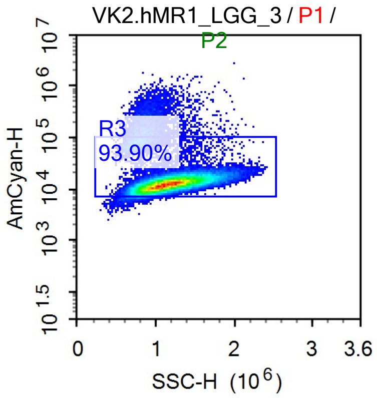


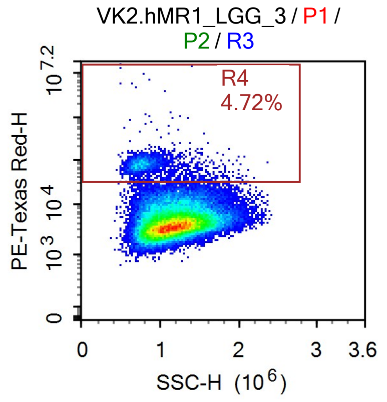

Sample Statistics of VK2.hMR1_LGG_3

| Gate | Count | % Parent | X | Y | Median X | Median Y |
|---|---|---|---|---|---|---|
| All | 76,312 | | | | | |
| P1 | 61,151 | 80.13% | FSC-A | FSC-H | 7,862,102 | 3,266,130 |
| P2 | 60,027 | 98.16% | SSC-H | FSC-H | 1,127,765 | 3,263,521 |
| R3 | 56,367 | 93.90% | SSC-H | AmCyan-H | 1,145,658 | 12,826 |
| R4 | 2,662 | 4.72% | SSC-H | PE-Texas Red-H | 796,813 | 78,297 |

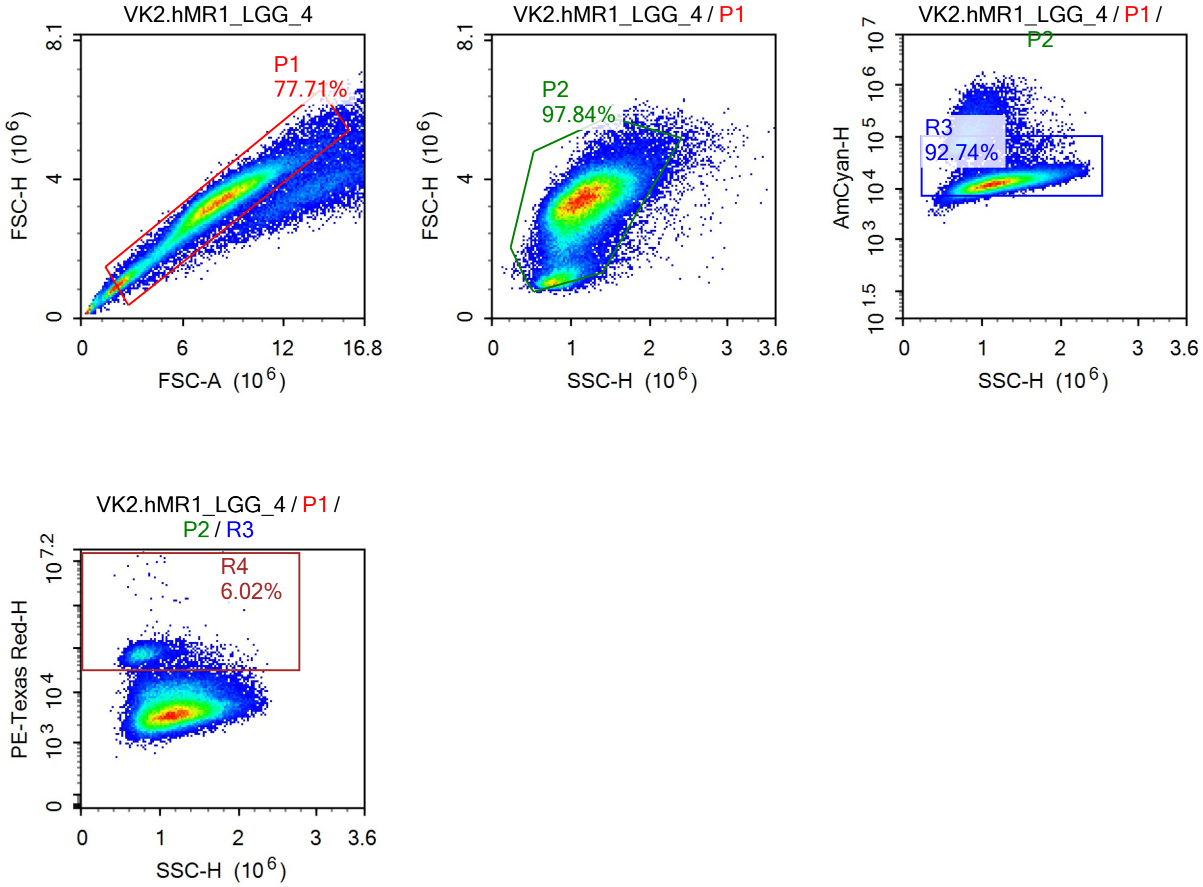


Sample Statistics of VK2.hMR1_LGG_4

| Gate | Count | % Parent | X | Y | Median X | Median Y |
|---|---|---|---|---|---|---|
| All | 88,322 | | | | | |
| P1 | 68,633 | 77.71% | FSC-A | FSC-H | 7,834,561 | 3,274,430 |
| P2 | 67,148 | 97.84% | SSC-H | FSC-H | 1,132,345 | 3,272,088 |
| R3 | 62,274 | 92.74% | SSC-H | AmCyan-H | 1,152,421 | 12,938 |
| R4 | 3,752 | 6.02% | SSC-H | PE-Texas Red-H | 797,192 | 71,607 |

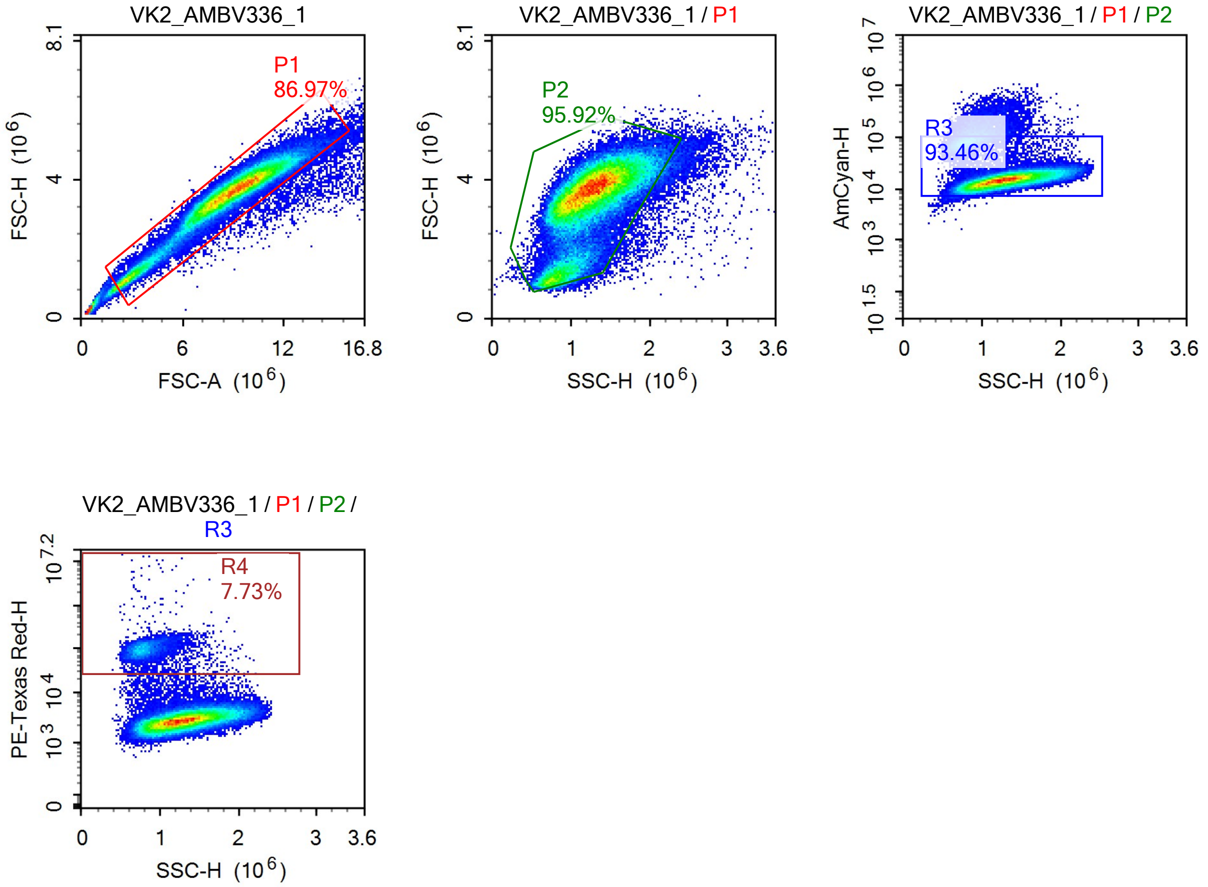


Sample Statistics of VK2_AMBV336_1

| Gate | Count | % Parent | X | Y | Median X | Median Y |
|---|---|---|---|---|---|---|
| All | 92,640 | | | | | |
| P1 | 80,570 | 86.97% | FSC-A | FSC-H | 8,809,750 | 3,612,630 |
| P2 | 77,282 | 95.92% | SSC-H | FSC-H | 1,237,389 | 3,600,111 |
| R3 | 72,224 | 93.46% | SSC-H | AmCyan-H | 1,258,022 | 15,047 |
| R4 | 5,584 | 7.73% | SSC-H | PE-Texas Red-H | 802,787 | 95,075 |

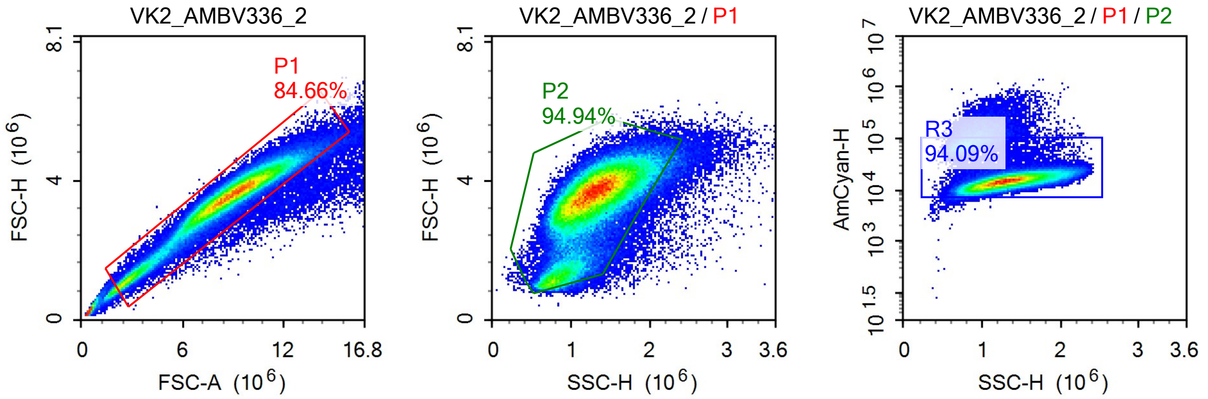

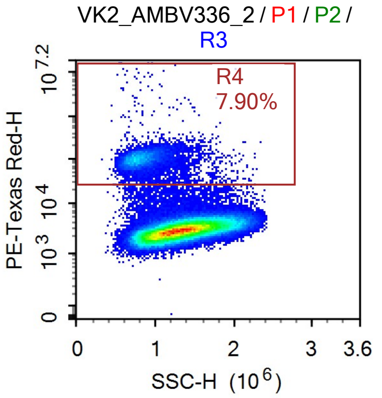


Sample Statistics of VK2_AMBV336_2

| Gate | Count | % Parent | X | Y | Median X | Median Y |
|---|---|---|---|---|---|---|
| All | 148,482 | | | | | |
| P1 | 125,709 | 84.66% | FSC-A | FSC-H | 8,734,182 | 3,539,809 |
| P2 | 119,350 | 94.94% | SSC-H | FSC-H | 1,267,718 | 3,525,008 |
| R3 | 112,295 | 94.09% | SSC-H | AmCyan-H | 1,285,940 | 15,187 |
| R4 | 8,866 | 7.90% | SSC-H | PE-Texas Red-H | 824,938 | 100,256 |

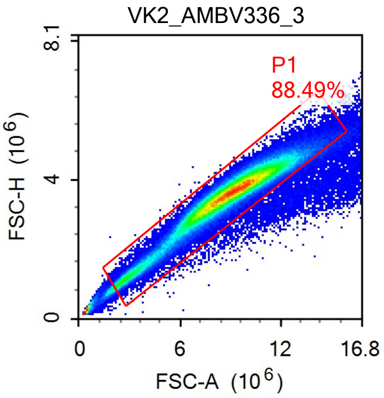


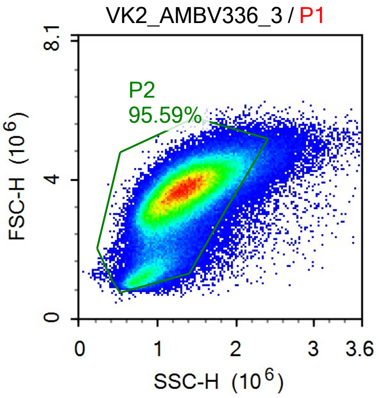


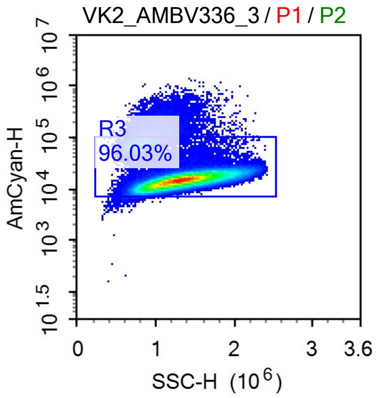


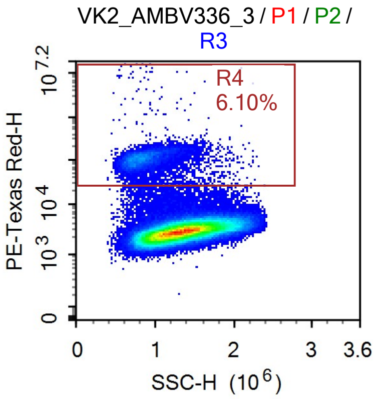

Sample Statistics of VK2_AMBV336_3

| Gate | Count | % Parent | X | Y | Median X | Median Y |
|---|---|---|---|---|---|---|
| All | 199,300 | | | | | |
| P1 | 176,363 | 88.49% | FSC-A | FSC-H | 8,922,351 | 3,604,883 |
| P2 | 168,578 | 95.59% | SSC-H | FSC-H | 1,295,085 | 3,587,765 |
| R3 | 161,878 | 96.03% | SSC-H | AmCyan-H | 1,307,231 | 15,026 |
| R4 | 9,871 | 6.10% | SSC-H | PE-Texas Red-H | 849,921 | 107,038 |

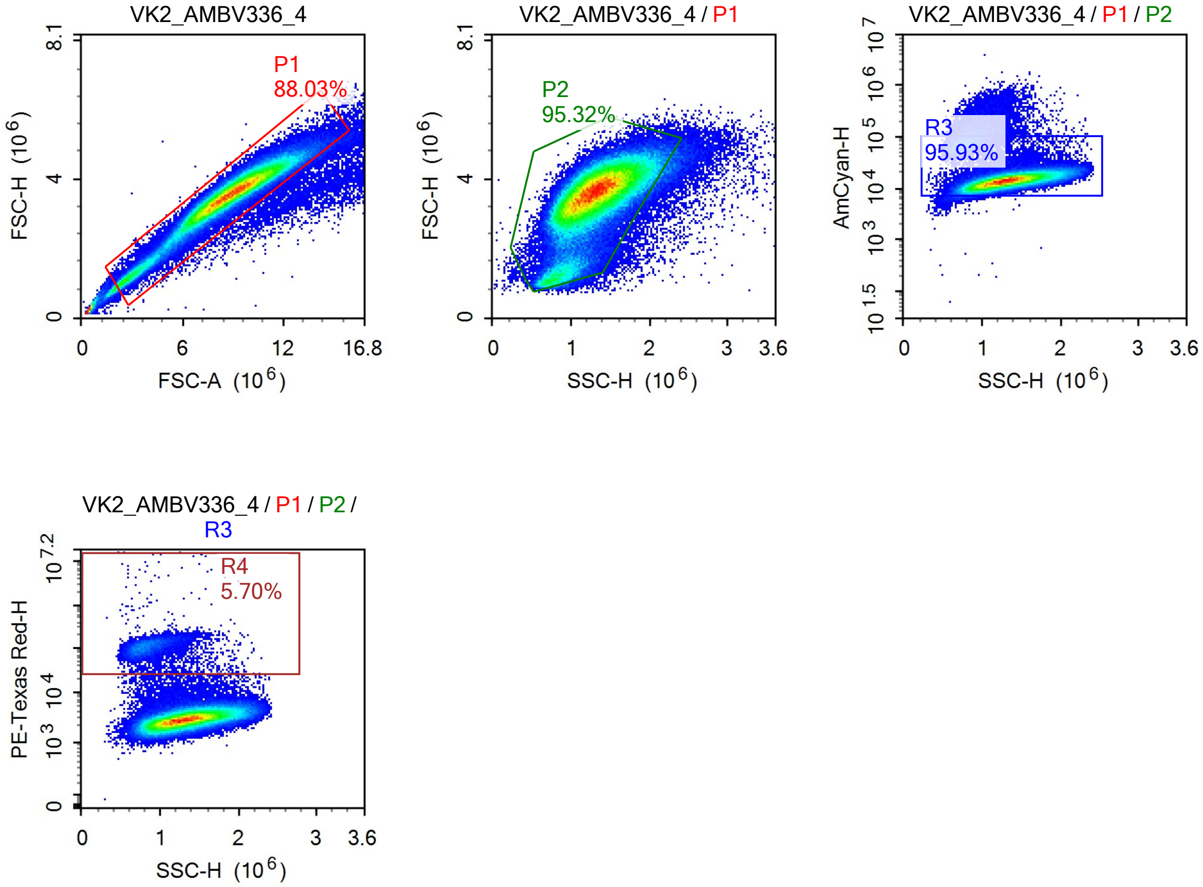


Sample Statistics of VK2_AMBV336_4

| Gate | Count | % Parent | X | Y | Median X | Median Y |
|---|---|---|---|---|---|---|
| All | 155,564 | | | | | |
| P1 | 136,943 | 88.03% | FSC-A | FSC-H | 8,802,005 | 3,565,066 |
| P2 | 130,528 | 95.32% | SSC-H | FSC-H | 1,288,307 | 3,547,654 |
| R3 | 125,210 | 95.93% | SSC-H | AmCyan-H | 1,302,226 | 14,259 |
| R4 | 7,143 | 5.70% | SSC-H | PE-Texas Red-H | 854,828 | 108,467 |

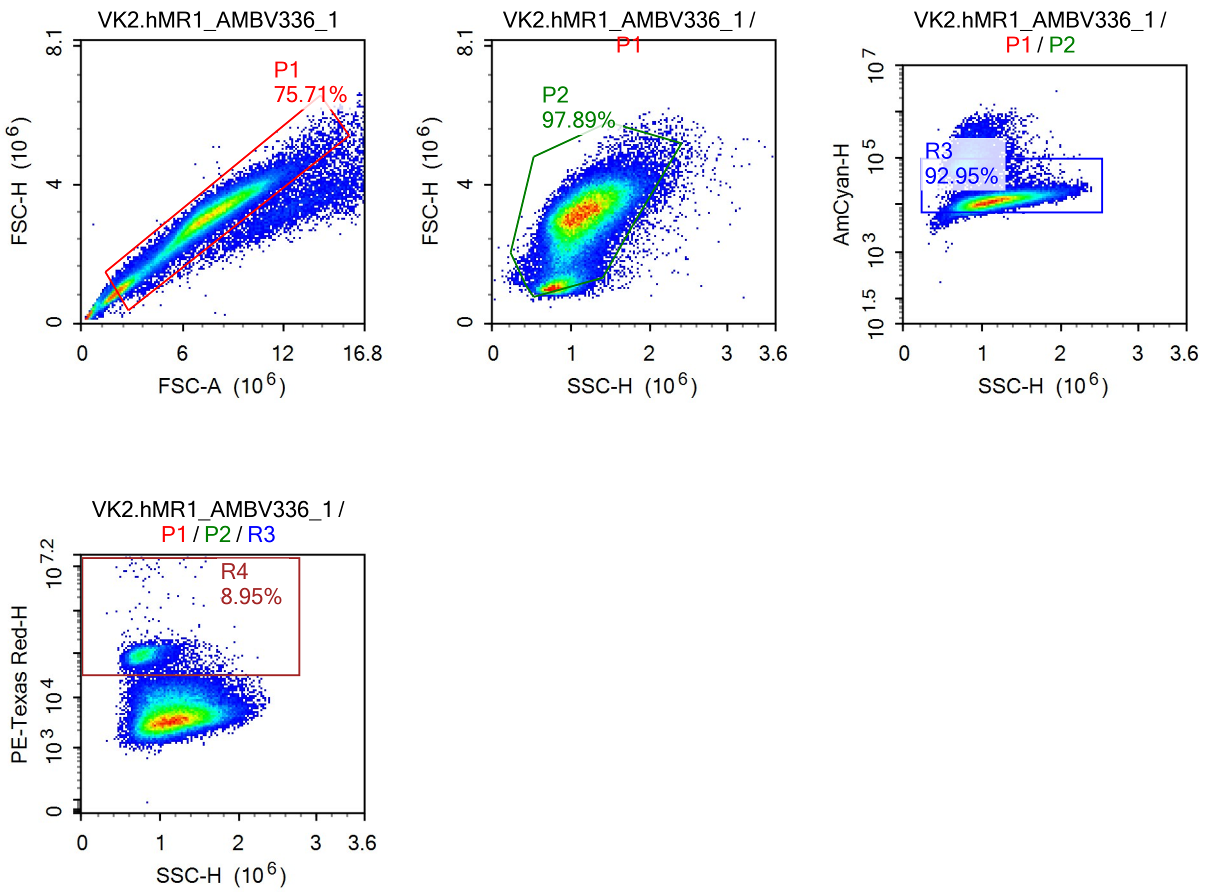


Sample Statistics of VK2.hMR1_AMBV336_1

| Gate | Count | % Parent | X | Y | Median X | Median Y |
|---|---|---|---|---|---|---|
| All | 86,856 | | | | | |
| P1 | 65,757 | 75.71% | FSC-A | FSC-H | 7,267,949 | 3,012,623 |
| P2 | 64,370 | 97.89% | SSC-H | FSC-H | 1,101,514 | 3,019,189 |
| R3 | 59,831 | 92.95% | SSC-H | AmCyan-H | 1,123,232 | 12,826 |
| R4 | 5,355 | 8.95% | SSC-H | PE-Texas Red-H | 786,167 | 91,461 |

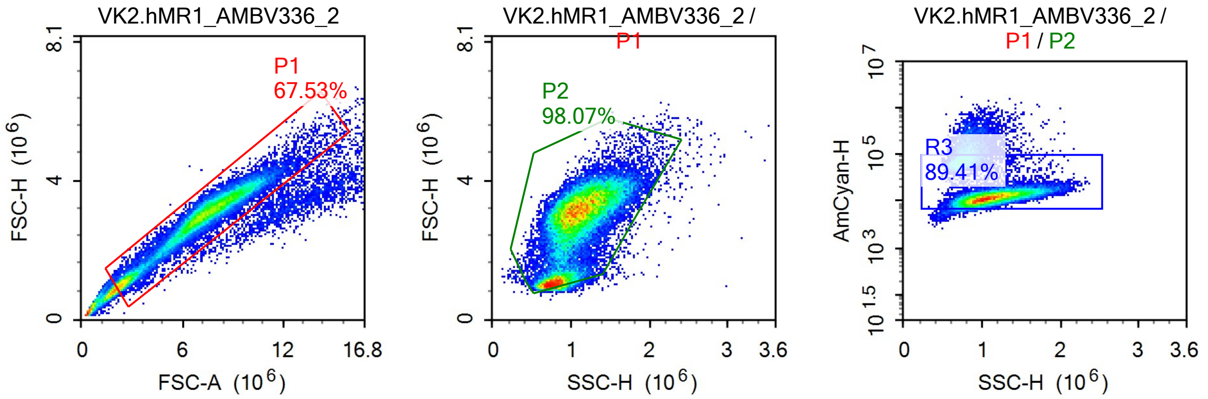

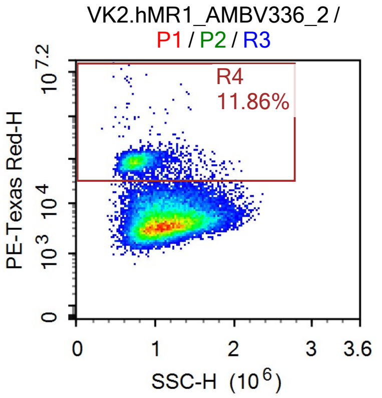


Sample Statistics of VK2.hMR1_AMBV336_2

| Gate | Count | % Parent | X | Y | Median X | Median Y |
|---|---|---|---|---|---|---|
| All | 52,472 | | | | | |
| P1 | 35,435 | 67.53% | FSC-A | FSC-H | 7,037,553 | 2,953,829 |
| P2 | 34,751 | 98.07% | SSC-H | FSC-H | 1,045,420 | 2,964,692 |
| R3 | 31,071 | 89.41% | SSC-H | AmCyan-H | 1,075,594 | 12,806 |
| R4 | 3,684 | 11.86% | SSC-H | PE-Texas Red-H | 751,749 | 87,563 |

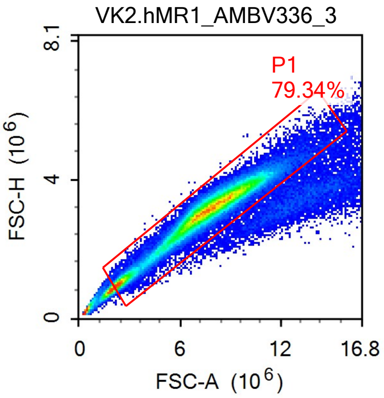


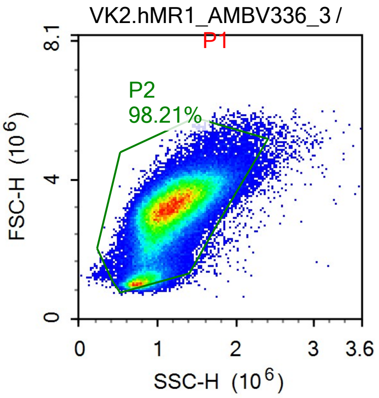


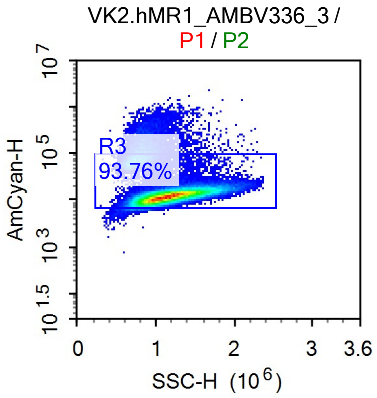


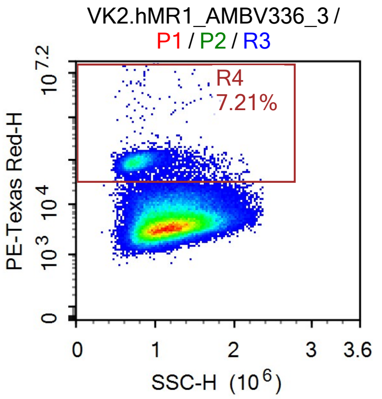

Sample Statistics of VK2.hMR1_AMBV336_3

| Gate | Count | % Parent | X | Y | Median X | Median Y |
|---|---|---|---|---|---|---|
| All | 126,508 | | | | | |
| P1 | 100,372 | 79.34% | FSC-A | FSC-H | 7,608,183 | 3,129,788 |
| P2 | 98,573 | 98.21% | SSC-H | FSC-H | 1,135,944 | 3,128,204 |
| R3 | 92,426 | 93.76% | SSC-H | AmCyan-H | 1,156,599 | 12,909 |
| R4 | 6,668 | 7.21% | SSC-H | PE-Texas Red-H | 775,193 | 90,068 |

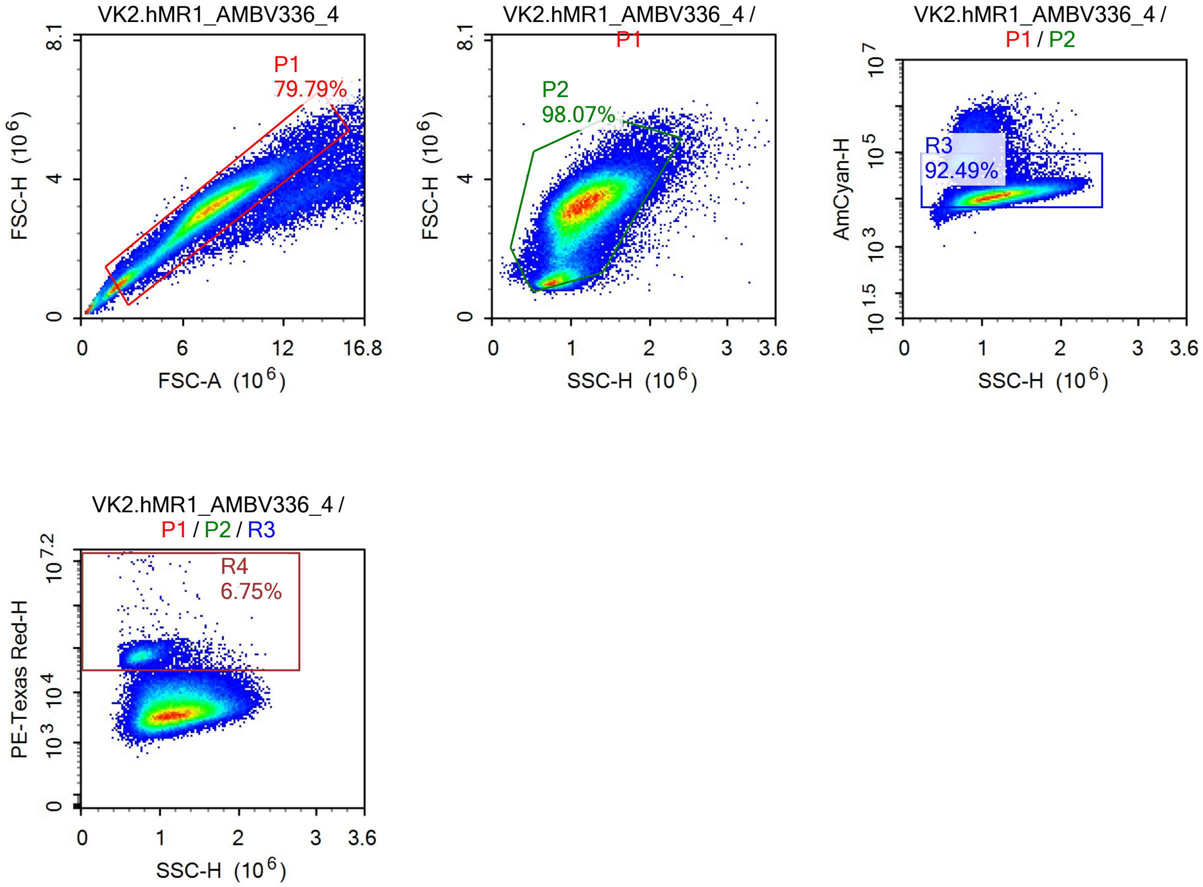


Sample Statistics of VK2.hMR1_AMBV336_4

| Gate | Count | % Parent | X | Y | Median X | Median Y |
|---|---|---|---|---|---|---|
| All | 115,767 | | | | | |
| P1 | 92,367 | 79.79% | FSC-A | FSC-H | 7,483,276 | 3,127,722 |
| P2 | 90,585 | 98.07% | SSC-H | FSC-H | 1,131,298 | 3,127,765 |
| R3 | 83,780 | 92.49% | SSC-H | AmCyan-H | 1,156,123 | 12,483 |
| R4 | 5,654 | 6.75% | SSC-H | PE-Texas Red-H | 780,858 | 69,052 |

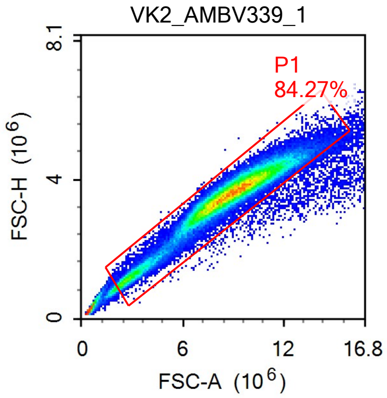


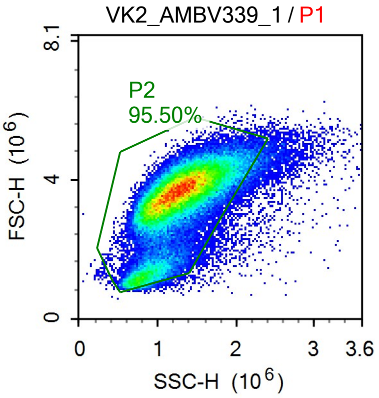


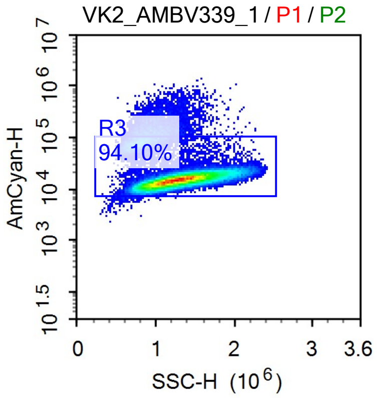


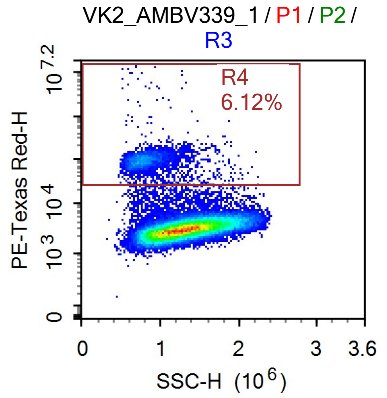


Sample Statistics of VK2_AMBV339_1

| Gate | Count | % Parent | X | Y | Median X | Median Y |
|---|---|---|---|---|---|---|
| All | 84,837 | | | | | |
| P1 | 71,488 | 84.27% | FSC-A | FSC-H | 8,807,462 | 3,596,674 |
| P2 | 68,269 | 95.50% | SSC-H | FSC-H | 1,257,905 | 3,582,933 |
| R3 | 64,240 | 94.10% | SSC-H | AmCyan-H | 1,275,515 | 15,260 |
| R4 | 3,933 | 6.12% | SSC-H | PE-Texas Red-H | 800,026 | 94,913 |

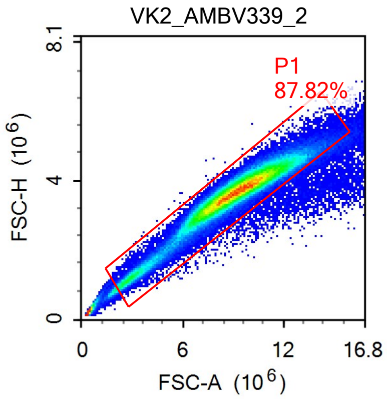


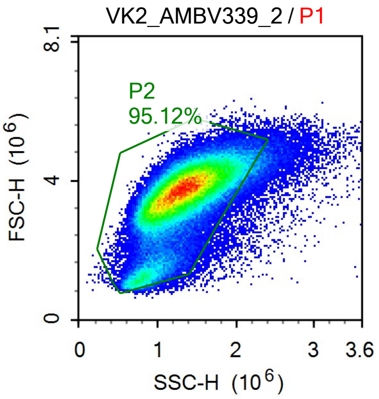


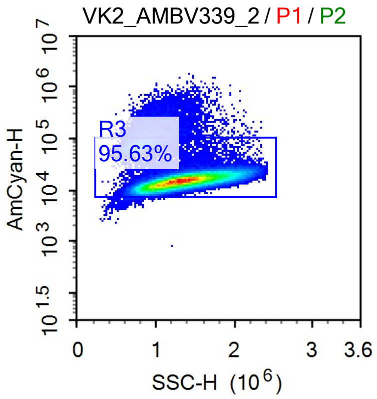

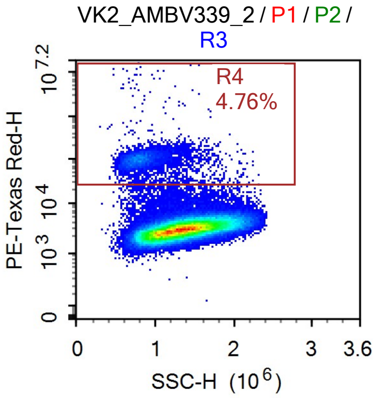


Sample Statistics of VK2_AMBV339_2

| Gate | Count | % Parent | X | Y | Median X | Median Y |
|---|---|---|---|---|---|---|
| All | 157,565 | | | | | |
| P1 | 138,367 | 87.82% | FSC-A | FSC-H | 9,128,234 | 3,678,816 |
| P2 | 131,616 | 95.12% | SSC-H | FSC-H | 1,308,204 | 3,657,370 |
| R3 | 125,870 | 95.63% | SSC-H | AmCyan-H | 1,321,666 | 15,199 |
| R4 | 5,994 | 4.76% | SSC-H | PE-Texas Red-H | 817,091 | 98,017 |

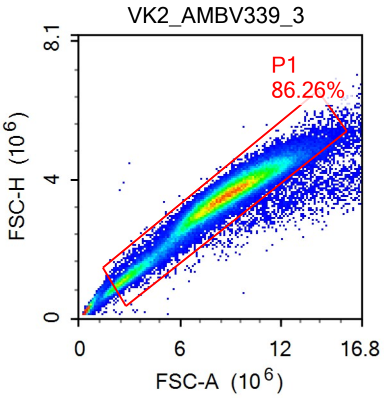


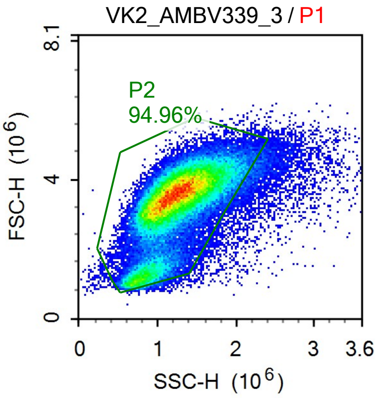


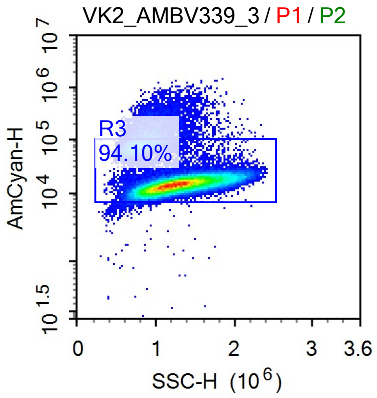


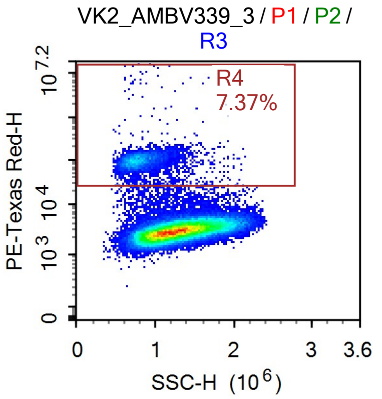

Sample Statistics of VK2_AMBV339_3

| Gate | Count | % Parent | X | Y | Median X | Median Y |
|---|---|---|---|---|---|---|
| All | 96,722 | | | | | |
| P1 | 83,432 | 86.26% | FSC-A | FSC-H | 8,598,250 | 3,508,558 |
| P2 | 79,228 | 94.96% | SSC-H | FSC-H | 1,225,679 | 3,494,444 |
| R3 | 74,553 | 94.10% | SSC-H | AmCyan-H | 1,243,158 | 14,864 |
| R4 | 5,492 | 7.37% | SSC-H | PE-Texas Red-H | 790,538 | 97,550 |

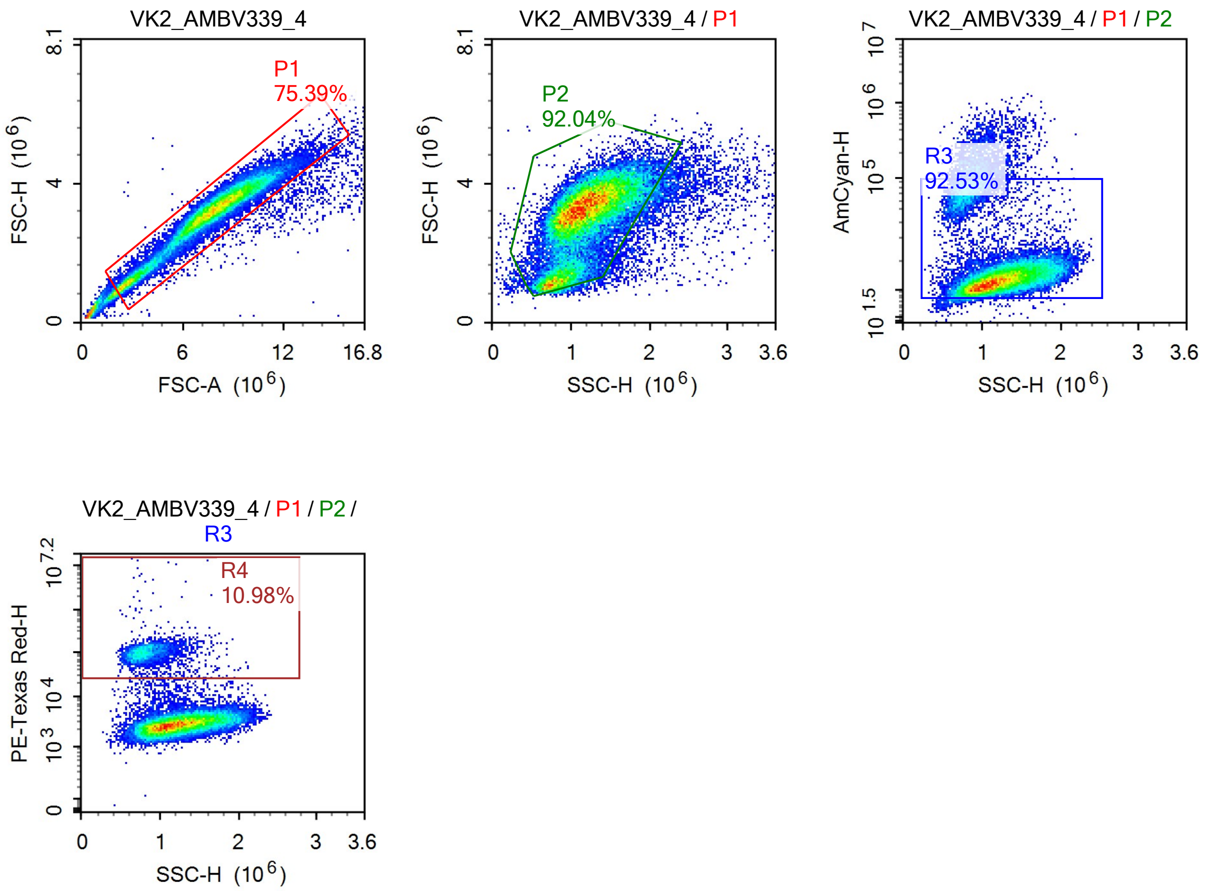


Sample Statistics of VK2_AMBV339_4

| Gate | Count | % Parent | X | Y | Median X | Median Y |
|---|---|---|---|---|---|---|
| All | 45,141 | | | | | |
| P1 | 34,033 | 75.39% | FSC-A | FSC-H | 7,715,882 | 3,200,911 |
| P2 | 31,325 | 92.04% | SSC-H | FSC-H | 1,154,114 | 3,188,625 |
| R3 | 28,985 | 92.53% | SSC-H | AmCyan-H | 1,174,134 | 13,331 |
| R4 | 3,183 | 10.98% | SSC-H | PE-Texas Red-H | 808,293 | 99,011 |

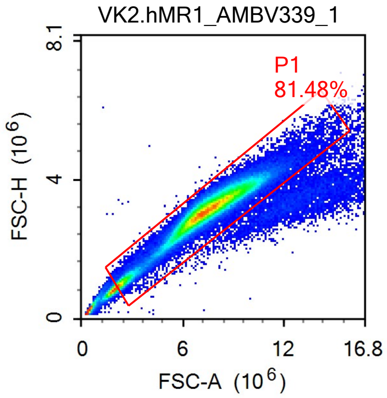


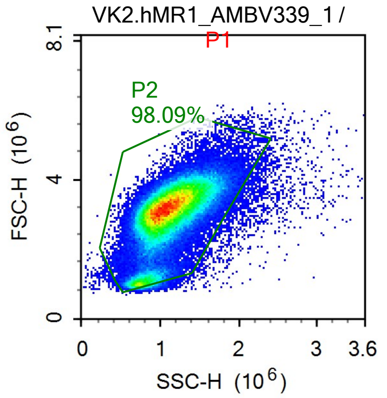


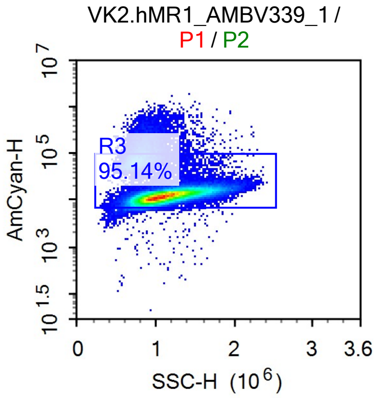


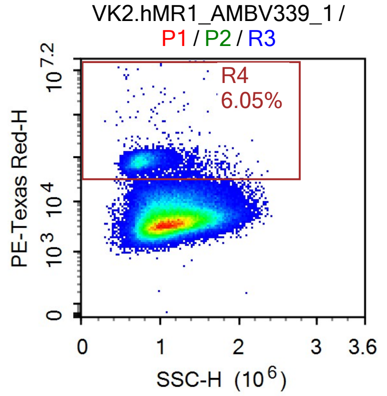


Sample Statistics of VK2.hMR1_AMBV339_1

| Gate | Count | % Parent | X | Y | Median X | Median Y |
|---|---|---|---|---|---|---|
| All | 134,881 | | | | | |
| P1 | 109,906 | 81.48% | FSC-A | FSC-H | 7,394,257 | 3,128,506 |
| P2 | 107,806 | 98.09% | SSC-H | FSC-H | 1,091,235 | 3,129,387 |
| R3 | 102,571 | 95.14% | SSC-H | AmCyan-H | 1,104,107 | 12,823 |
| R4 | 6,205 | 6.05% | SSC-H | PE-Texas Red-H | 768,713 | 80,883 |

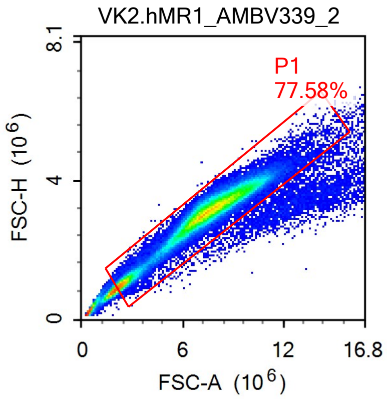


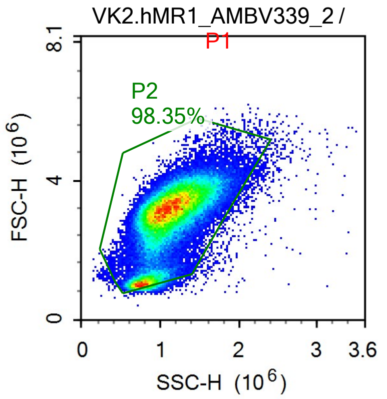


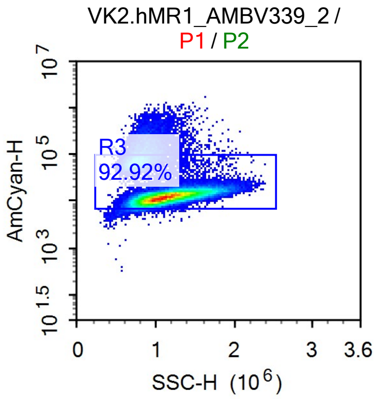

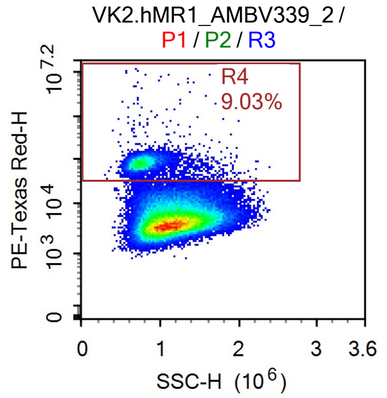


Sample Statistics of VK2.hMR1_AMBV339_2

| Gate | Count | % Parent | X | Y | Median X | Median Y |
|---|---|---|---|---|---|---|
| All | 103,721 | | | | | |
| P1 | 80,464 | 77.58% | FSC-A | FSC-H | 7,365,231 | 3,088,601 |
| P2 | 79,137 | 98.35% | SSC-H | FSC-H | 1,093,111 | 3,093,364 |
| R3 | 73,532 | 92.92% | SSC-H | AmCyan-H | 1,112,821 | 13,074 |
| R4 | 6,637 | 9.03% | SSC-H | PE-Texas Red-H | 769,154 | 80,695 |

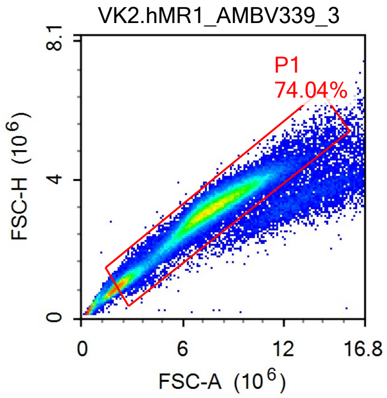


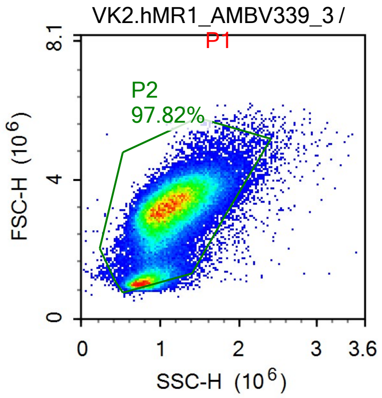


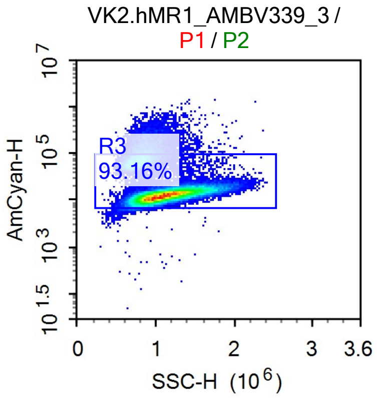


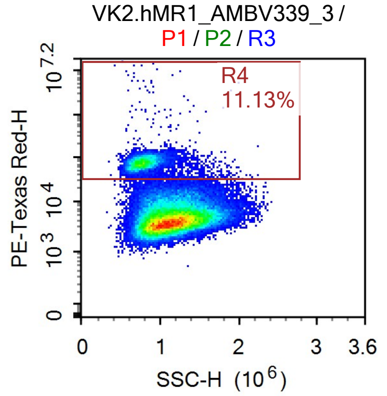

Sample Statistics of VK2.hMR1_AMBV339_3

| Gate | Count | % Parent | X | Y | Median X | Median Y |
|---|---|---|---|---|---|---|
| All | 95,551 | | | | | |
| └ P1 | 70,749 | 74.04% | FSC-A | FSC-H | 7,374,406 | 3,081,183 |
| └ P2 | 69,205 | 97.82% | SSC-H | FSC-H | 1,077,944 | 3,087,776 |
| └ R3 | 64,472 | 93.16% | SSC-H | AmCyan-H | 1,094,372 | 13,670 |
| └ R4 | 7,177 | 11.13% | SSC-H | PE-Texas Red-H | 773,630 | 74,777 |

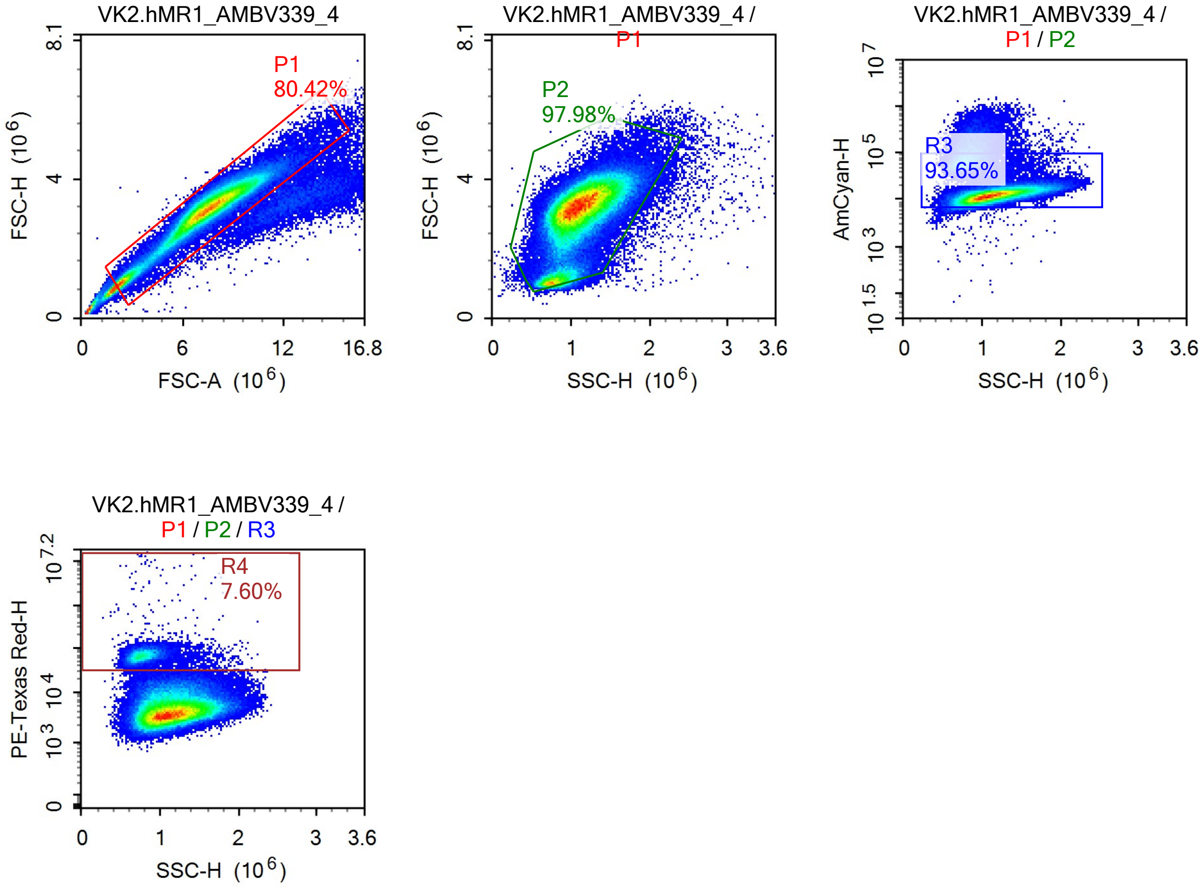


Sample Statistics of VK2.hMR1_AMBV339_4

| Gate | Count | % Parent | X | Y | Median X | Median Y |
|---|---|---|---|---|---|---|
| All | 158,044 | | | | | |
| └ P1 | 127,095 | 80.42% | FSC-A | FSC-H | 7,384,728 | 3,123,634 |
| └ P2 | 124,522 | 97.98% | SSC-H | FSC-H | 1,090,682 | 3,123,951 |
| └ R3 | 116,621 | 93.65% | SSC-H | AmCyan-H | 1,107,537 | 12,918 |
| └ R4 | 8,867 | 7.60% | SSC-H | PE-Texas Red-H | 781,698 | 67,999 |

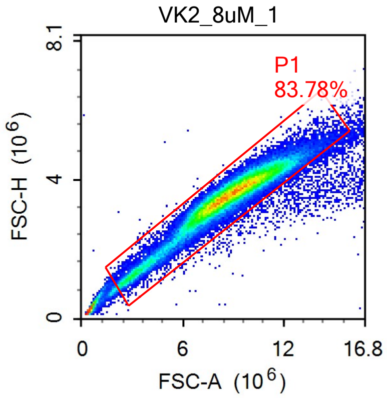


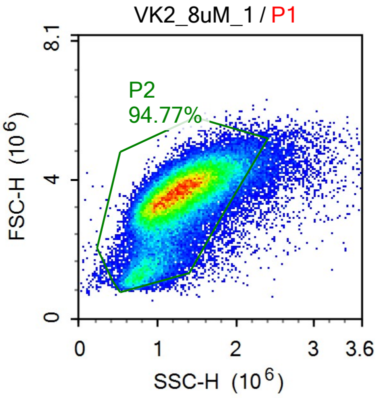


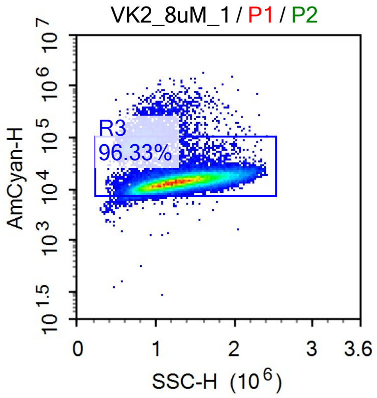


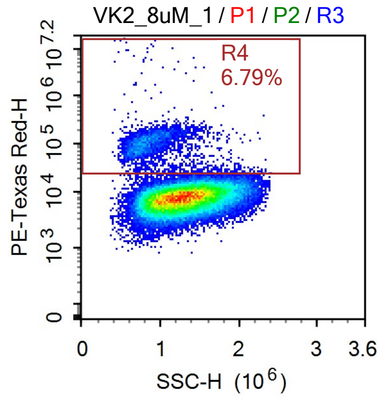


Sample Statistics of VK2_8uM_1

| Gate | Count | % Parent | X | Y | Median X | Median Y |
|---|---|---|---|---|---|---|
| All | 72,541 | | | | | |
| P1 | 60,778 | 83.78% | FSC-A | FSC-H | 8,705,908 | 3,544,727 |
| P2 | 57,600 | 94.77% | SSC-H | FSC-H | 1,251,396 | 3,529,520 |
| R3 | 55,486 | 96.33% | SSC-H | AmCyan-H | 1,263,665 | 13,906 |
| R4 | 3,765 | 6.79% | SSC-H | PE-Texas Red-H | 851,594 | 100,101 |

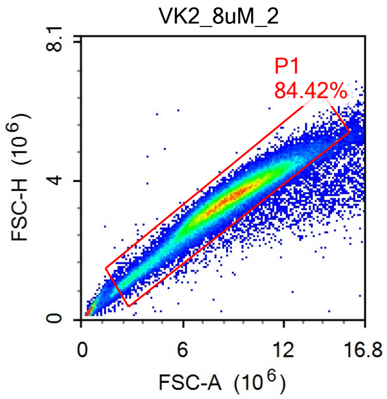


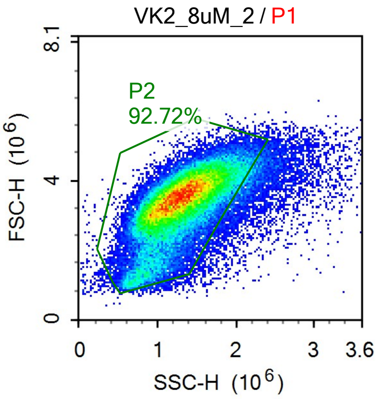


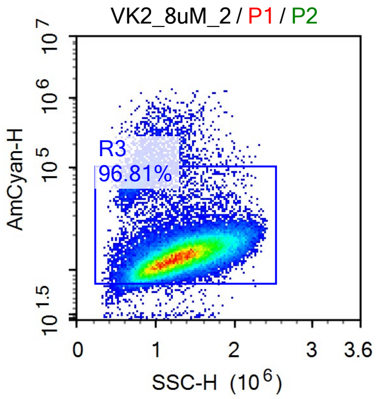

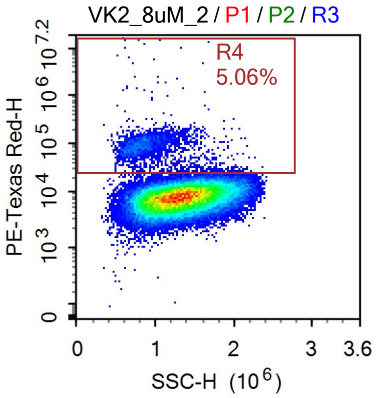


Sample Statistics of VK2_8uM_2

| Gate | Count | % Parent | X | Y | Median X | Median Y |
|---|---|---|---|---|---|---|
| All | 76,801 | | | | | |
| P1 | 64,834 | 84.42% | FSC-A | FSC-H | 8,668,046 | 3,455,183 |
| P2 | 60,115 | 92.72% | SSC-H | FSC-H | 1,283,807 | 3,433,937 |
| R3 | 58,196 | 96.81% | SSC-H | AmCyan-H | 1,293,369 | 13,833 |
| R4 | 2,942 | 5.06% | SSC-H | PE-Texas Red-H | 874,083 | 86,949 |

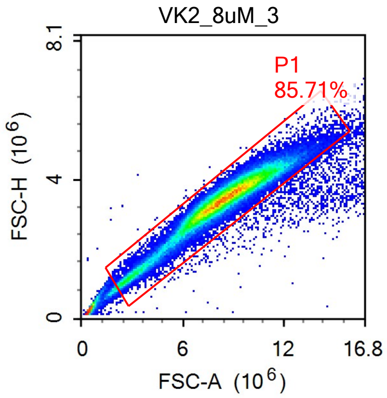


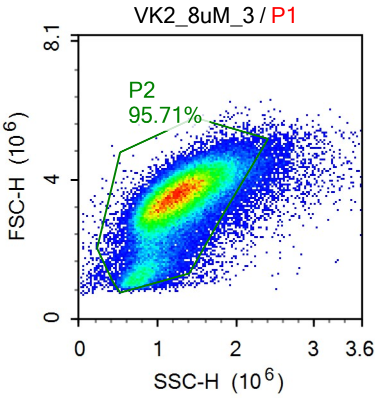


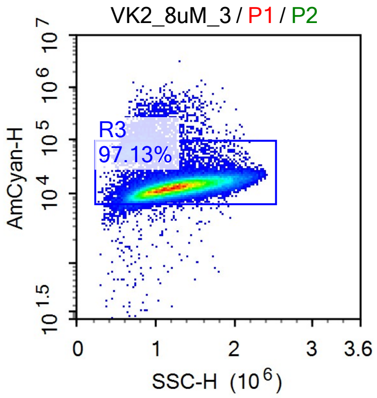


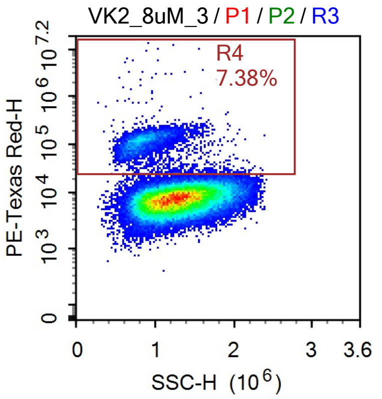

Sample Statistics of VK2_8uM_3

| Gate | Count | % Parent | X | Y | Median X | Median Y |
|---|---|---|---|---|---|---|
| All | 82,173 | | | | | |
| P1 | 70,428 | 85.71% | FSC-A | FSC-H | 8,405,141 | 3,447,312 |
| P2 | 67,407 | 95.71% | SSC-H | FSC-H | 1,232,131 | 3,435,778 |
| R3 | 65,472 | 97.13% | SSC-H | AmCyan-H | 1,241,265 | 13,449 |
| R4 | 4,835 | 7.38% | SSC-H | PE-Texas Red-H | 849,898 | 114,905 |

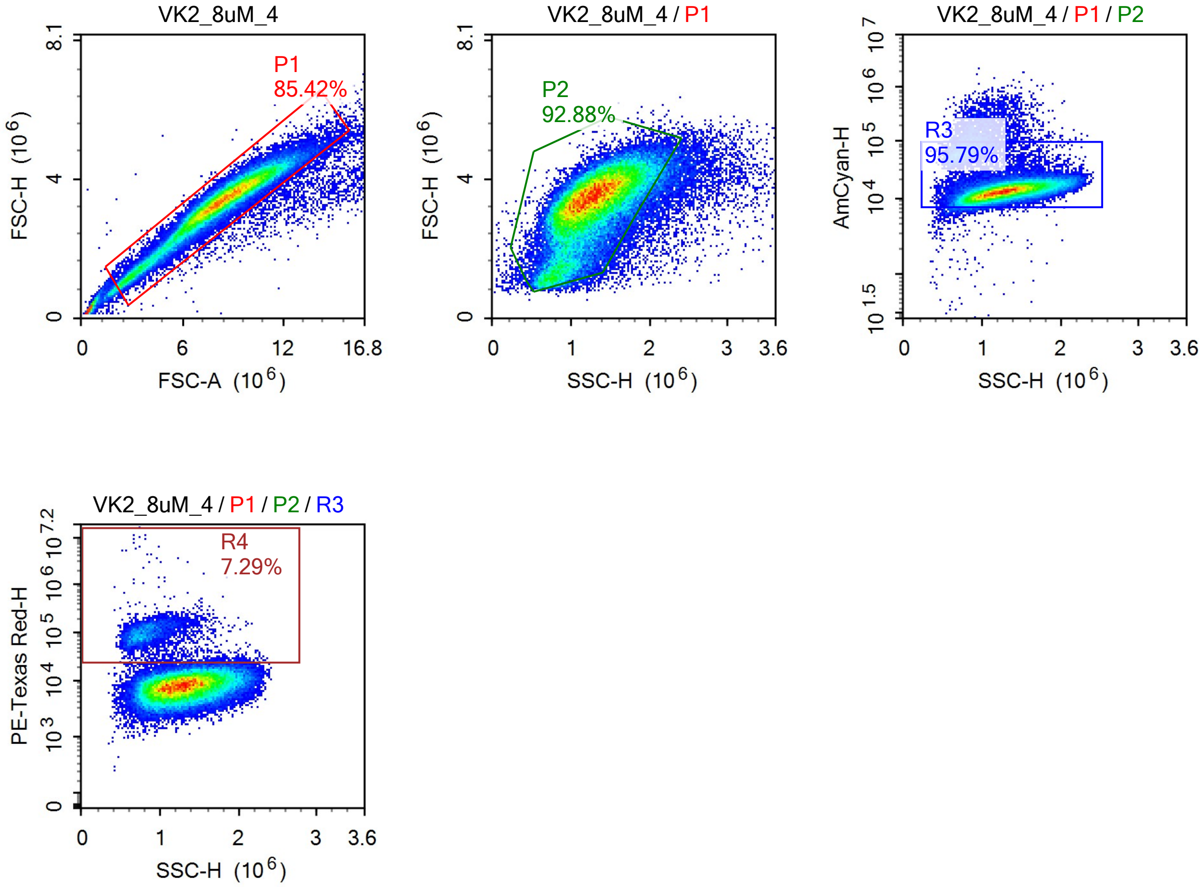


Sample Statistics of VK2_8uM_4

| Gate | Count | % Parent | X | Y | Median X | Median Y |
|---|---|---|---|---|---|---|
| All | 74,521 | | | | | |
| P1 | 63,659 | 85.42% | FSC-A | FSC-H | 8,283,757 | 3,368,992 |
| P2 | 59,126 | 92.88% | SSC-H | FSC-H | 1,250,567 | 3,351,860 |
| R3 | 56,638 | 95.79% | SSC-H | AmCyan-H | 1,263,226 | 13,412 |
| R4 | 4,131 | 7.29% | SSC-H | PE-Texas Red-H | 850,899 | 93,216 |

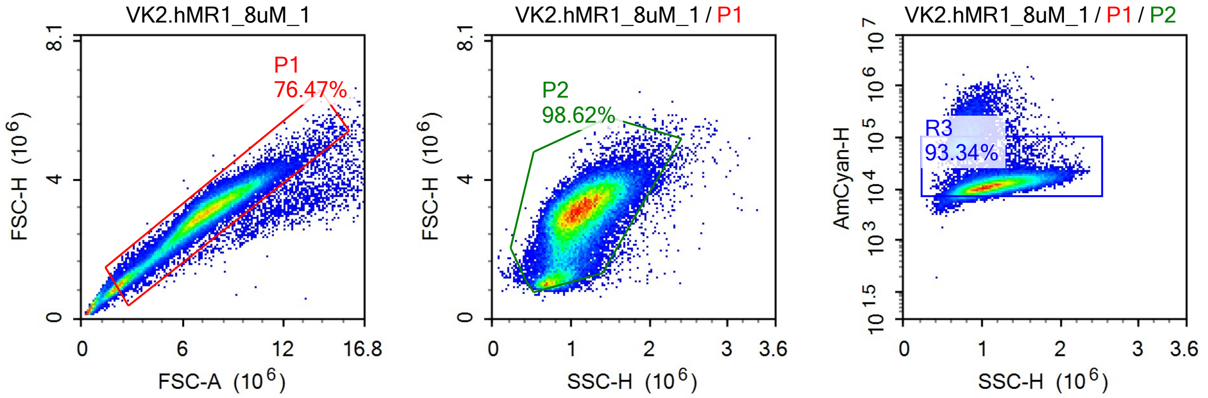


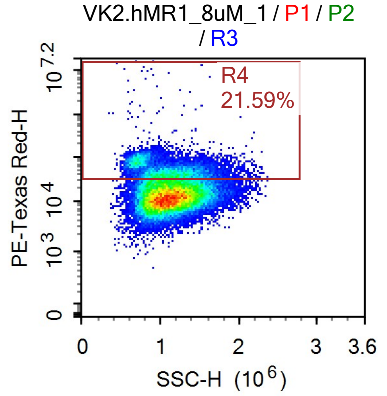


Sample Statistics of VK2.hMR1_8uM_1

| Gate | Count | % Parent | X | Y | Median X | Median Y |
|---|---|---|---|---|---|---|
| All | 69,052 | | | | | |
| P1 | 52,804 | 76.47% | FSC-A | FSC-H | 7,155,727 | 2,982,166 |
| P2 | 52,077 | 98.62% | SSC-H | FSC-H | 1,079,640 | 2,984,785 |
| R3 | 48,609 | 93.34% | SSC-H | AmCyan-H | 1,099,276 | 11,962 |
| R4 | 10,495 | 21.59% | SSC-H | PE-Texas Red-H | 1,059,897 | 49,113 |

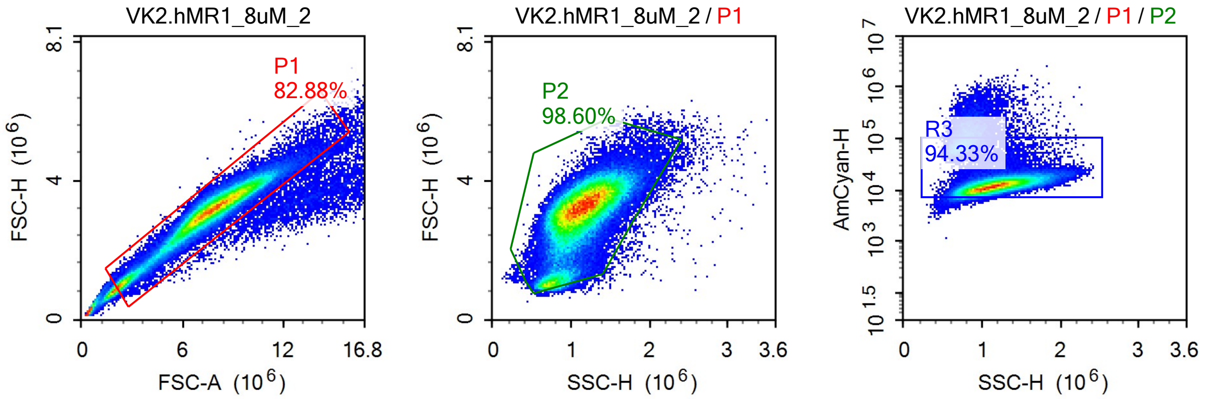

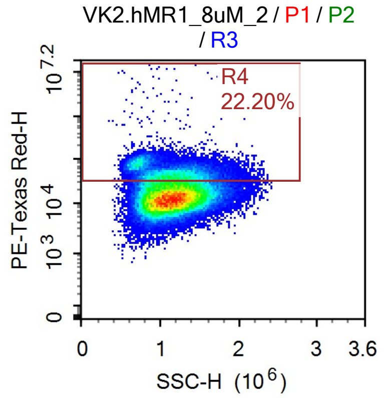


Sample Statistics of VK2.hMR1_8uM_2

| Gate | Count | % Parent | X | Y | Median X | Median Y |
|---|---|---|---|---|---|---|
| All | 114,931 | | | | | |
| P1 | 95,259 | 82.88% | FSC-A | FSC-H | 7,661,931 | 3,158,153 |
| P2 | 93,923 | 98.60% | SSC-H | FSC-H | 1,117,269 | 3,158,415 |
| R3 | 88,600 | 94.33% | SSC-H | AmCyan-H | 1,134,857 | 12,376 |
| R4 | 19,666 | 22.20% | SSC-H | PE-Texas Red-H | 1,121,696 | 48,436 |

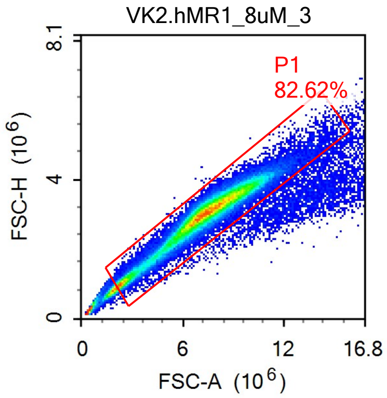


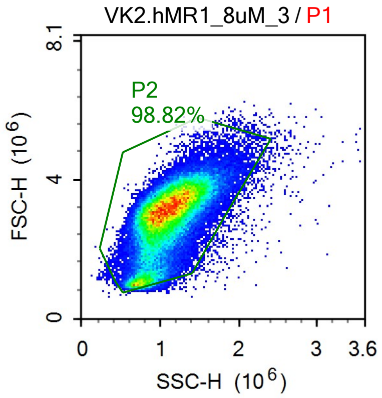


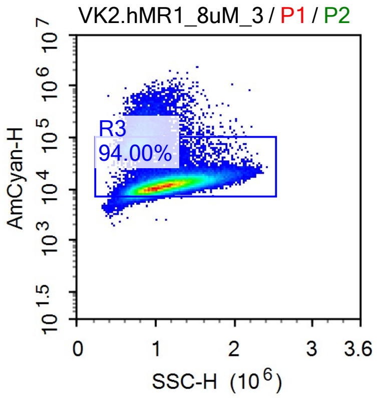


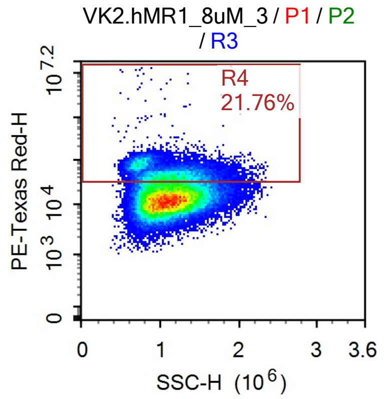

Sample Statistics of VK2.hMR1_8uM_3

| Gate | Count | % Parent | X | Y | Median X | Median Y |
|---|---|---|---|---|---|---|
| All | 92,933 | | | | | |
| P1 | 76,782 | 82.62% | FSC-A | FSC-H | 7,343,198 | 3,047,619 |
| P2 | 75,879 | 98.82% | SSC-H | FSC-H | 1,076,430 | 3,049,286 |
| R3 | 71,325 | 94.00% | SSC-H | AmCyan-H | 1,093,983 | 11,992 |
| R4 | 15,523 | 21.76% | SSC-H | PE-Texas Red-H | 1,079,416 | 48,338 |

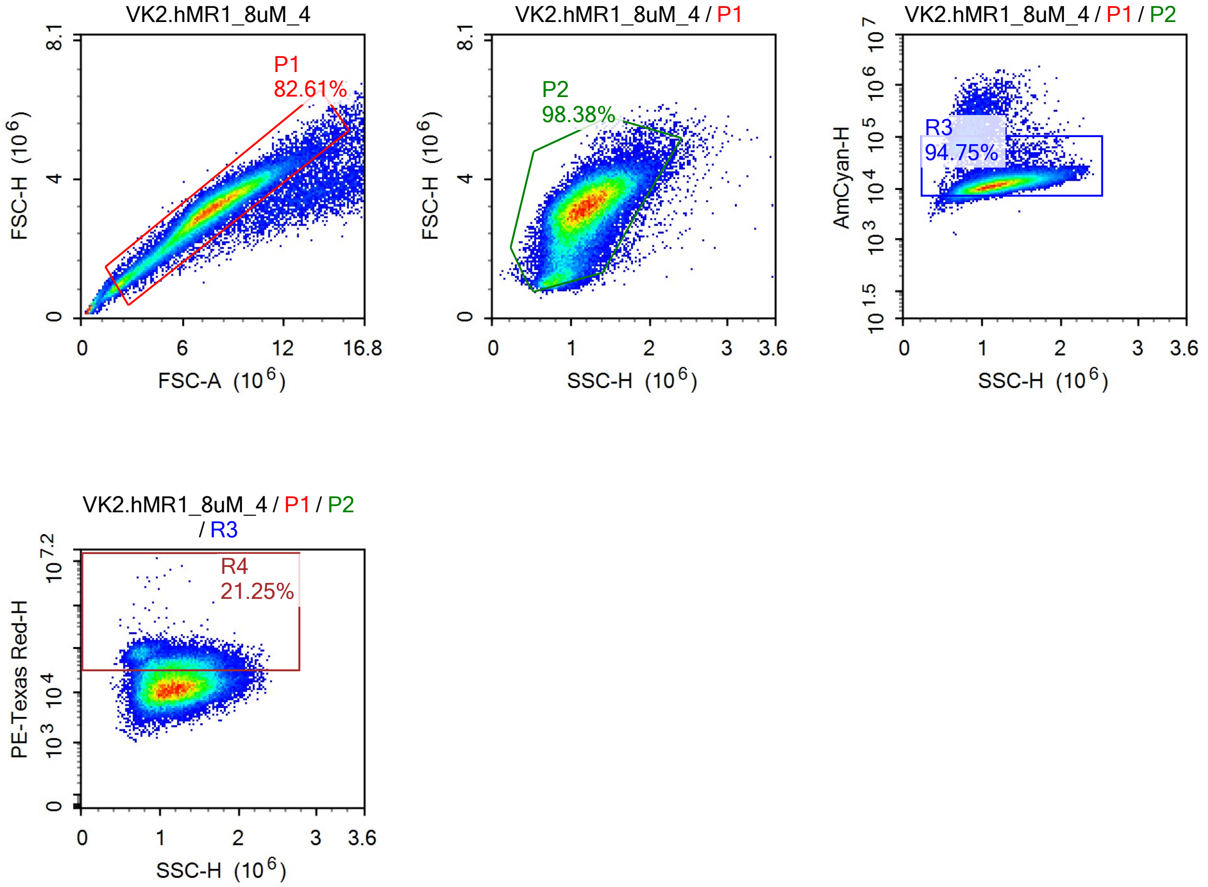


Sample Statistics of VK2.hMR1_8uM_4

| Gate | Count | % Parent | X | Y | Median X | Median Y |
|---|---|---|---|---|---|---|
| All | 65,257 | | | | | |
| P1 | 53,912 | 82.61% | FSC-A | FSC-H | 7,572,097 | 3,103,816 |
| P2 | 53,040 | 98.38% | SSC-H | FSC-H | 1,152,443 | 3,102,489 |
| R3 | 50,257 | 94.75% | SSC-H | AmCyan-H | 1,167,760 | 12,007 |
| R4 | 10,680 | 21.25% | SSC-H | PE-Texas Red-H | 1,182,703 | 46,933 |

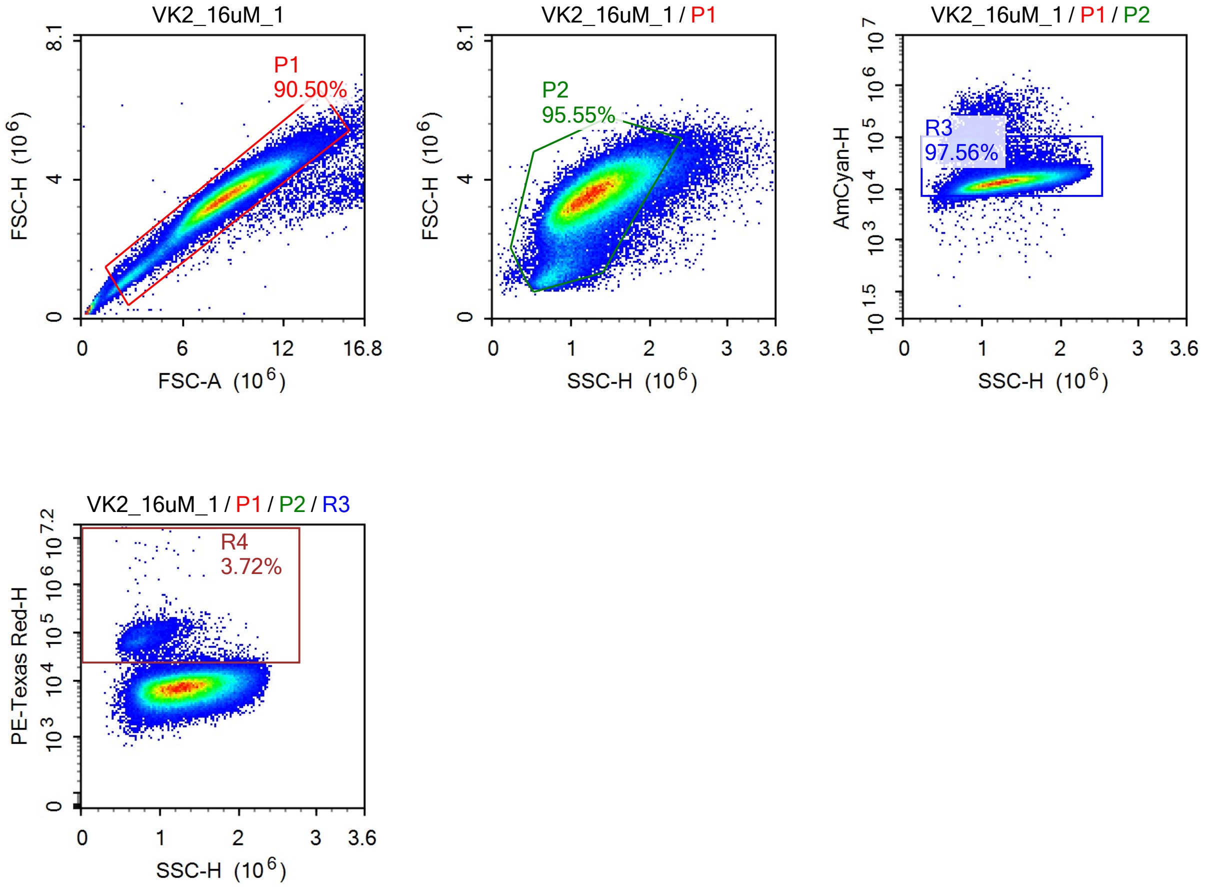


Sample Statistics of VK2_16uM_1

| Gate | Count | % Parent | X | Y | Median X | Median Y |
|---|---|---|---|---|---|---|
| All | 121,297 | | | | | |
| P1 | 109,776 | 90.50% | FSC-A | FSC-H | 8,608,862 | 3,531,274 |
| P2 | 104,892 | 95.55% | SSC-H | FSC-H | 1,273,352 | 3,514,047 |
| R3 | 102,335 | 97.56% | SSC-H | AmCyan-H | 1,280,717 | 13,761 |
| R4 | 3,804 | 3.72% | SSC-H | PE-Texas Red-H | 805,685 | 73,341 |

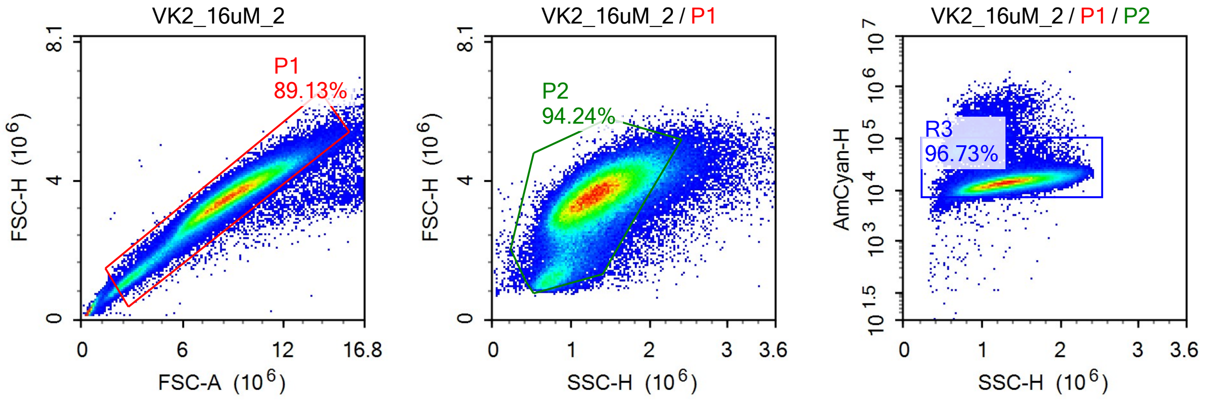

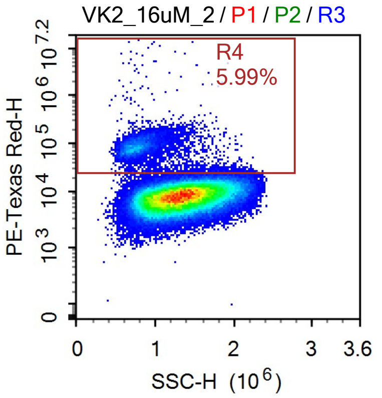


Sample Statistics of VK2_16uM_2

| Gate | Count | % Parent | X | Y | Median X | Median Y |
|---|---|---|---|---|---|---|
| All | 138,788 | | | | | |
| P1 | 123,699 | 89.13% | FSC-A | FSC-H | 8,679,775 | 3,512,208 |
| P2 | 116,575 | 94.24% | SSC-H | FSC-H | 1,292,612 | 3,488,345 |
| R3 | 112,761 | 96.73% | SSC-H | AmCyan-H | 1,302,101 | 14,239 |
| R4 | 6,756 | 5.99% | SSC-H | PE-Texas Red-H | 806,337 | 84,319 |

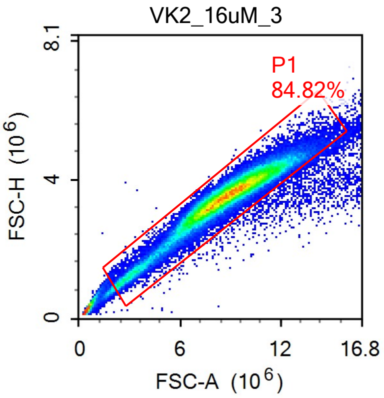


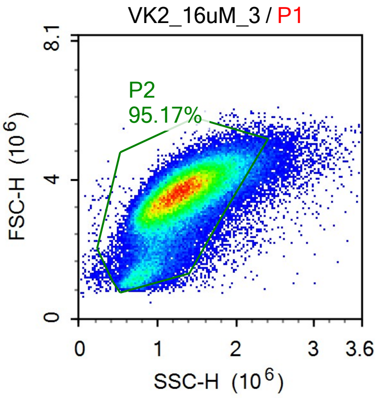


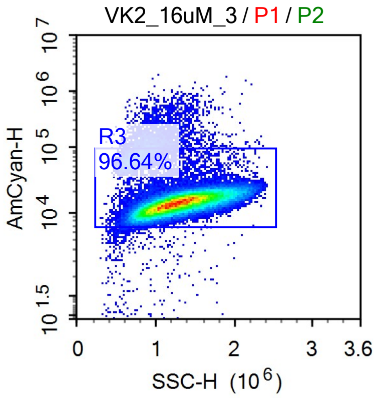


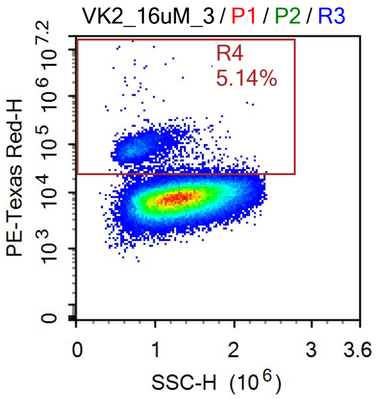

Sample Statistics of VK2_16uM_3

| Gate | Count | % Parent | X | Y | Median X | Median Y |
|---|---|---|---|---|---|---|
| All | 89,661 | | | | | |
| P1 | 76,050 | 84.82% | FSC-A | FSC-H | 8,768,270 | 3,541,852 |
| P2 | 72,379 | 95.17% | SSC-H | FSC-H | 1,272,147 | 3,525,819 |
| R3 | 69,948 | 96.64% | SSC-H | AmCyan-H | 1,282,324 | 14,277 |
| R4 | 3,593 | 5.14% | SSC-H | PE-Texas Red-H | 814,662 | 80,424 |

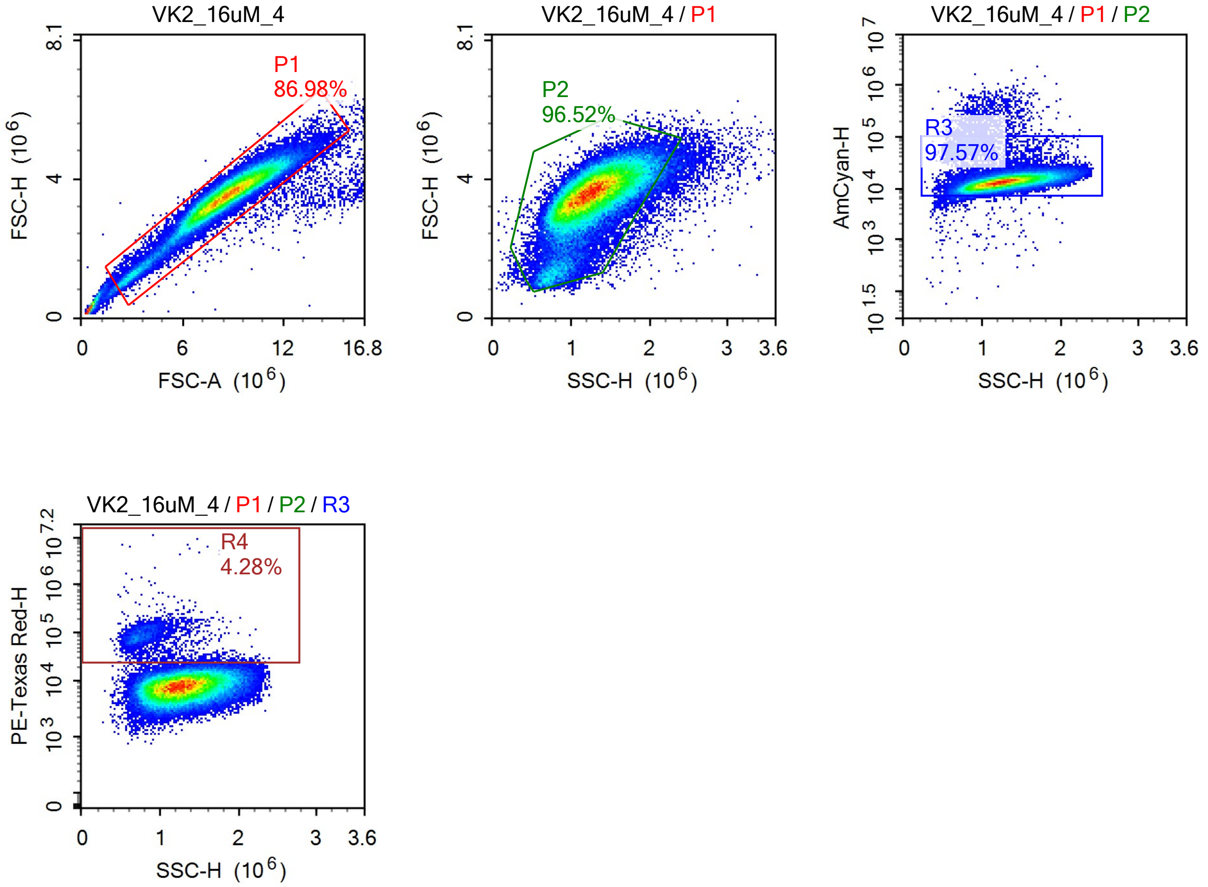


Sample Statistics of VK2_16uM_4

| Gate | Count | % Parent | X | Y | Median X | Median Y |
|---|---|---|---|---|---|---|
| All | 83,563 | | | | | |
| P1 | 72,684 | 86.98% | FSC-A | FSC-H | 8,672,274 | 3,564,755 |
| P2 | 70,156 | 96.52% | SSC-H | FSC-H | 1,256,278 | 3,552,946 |
| R3 | 68,452 | 97.57% | SSC-H | AmCyan-H | 1,263,443 | 13,518 |
| R4 | 2,933 | 4.28% | SSC-H | PE-Texas Red-H | 823,037 | 84,104 |

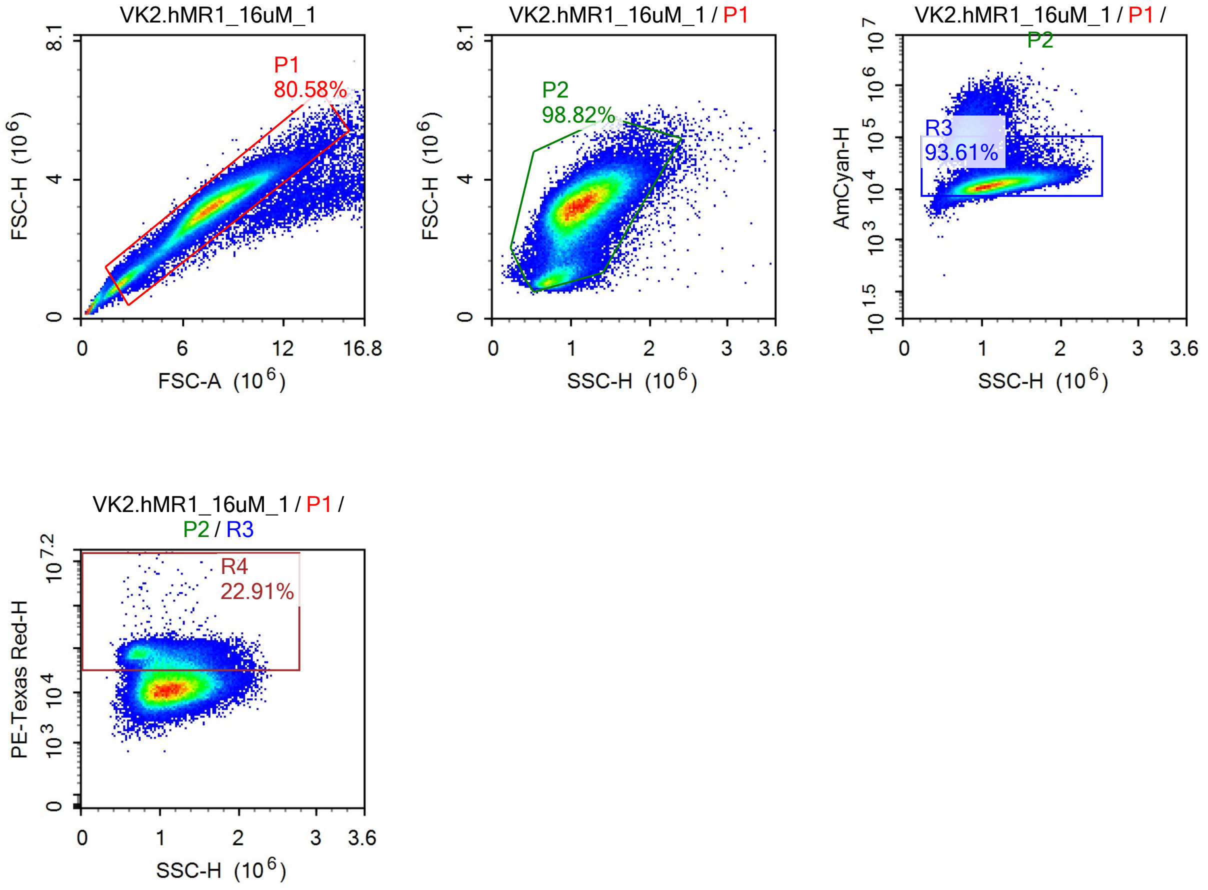


Sample Statistics of VK2.hMR1_16uM_1

| Gate | Count | % Parent | X | Y | Median X | Median Y |
|---|---|---|---|---|---|---|
| All | 123,869 | | | | | |
| P1 | 99,811 | 80.58% | FSC-A | FSC-H | 7,551,016 | 3,175,370 |
| P2 | 98,638 | 98.82% | SSC-H | FSC-H | 1,093,178 | 3,176,243 |
| R3 | 92,339 | 93.61% | SSC-H | AmCyan-H | 1,110,828 | 12,363 |
| R4 | 21,155 | 22.91% | SSC-H | PE-Texas Red-H | 1,047,460 | 58,530 |

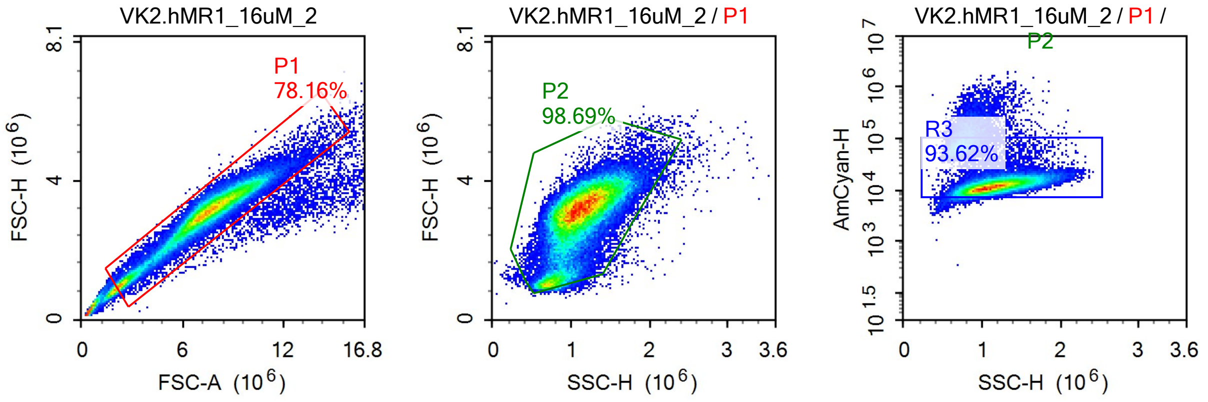

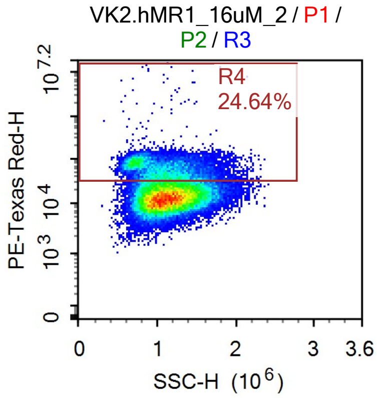


Sample Statistics of VK2.hMR1_16uM_2

| Gate | Count | % Parent | X | Y | Median X | Median Y |
|---|---|---|---|---|---|---|
| All | 75,746 | | | | | |
| P1 | 59,202 | 78.16% | FSC-A | FSC-H | 7,433,419 | 3,102,420 |
| P2 | 58,424 | 98.69% | SSC-H | FSC-H | 1,097,063 | 3,103,484 |
| R3 | 54,694 | 93.62% | SSC-H | AmCyan-H | 1,115,956 | 12,136 |
| R4 | 13,475 | 24.64% | SSC-H | PE-Texas Red-H | 1,069,452 | 58,614 |

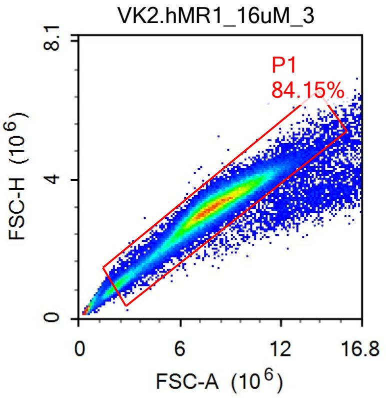


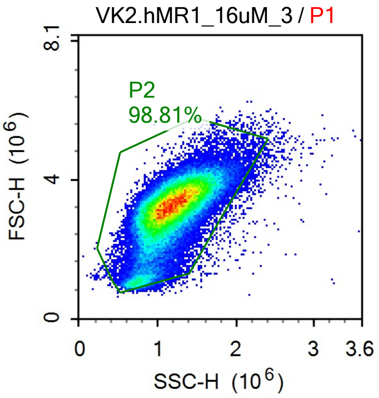


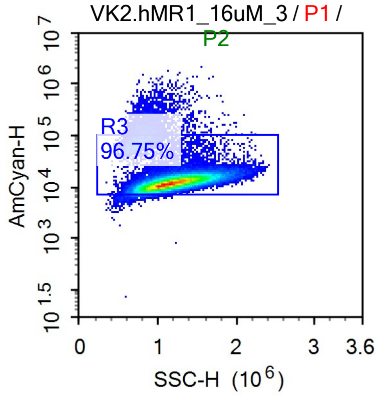


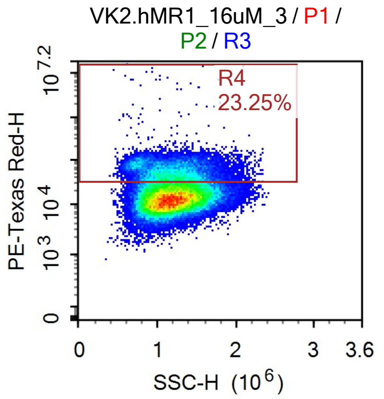

Sample Statistics of VK2.hMR1_16uM_3

| Gate | Count | % Parent | X | Y | Median X | Median Y |
|---|---|---|---|---|---|---|
| All | 85,821 | | | | | |
| P1 | 72,217 | 84.15% | FSC-A | FSC-H | 7,873,726 | 3,217,732 |
| P2 | 71,355 | 98.81% | SSC-H | FSC-H | 1,166,006 | 3,216,748 |
| R3 | 69,033 | 96.75% | SSC-H | AmCyan-H | 1,175,554 | 12,328 |
| R4 | 16,052 | 23.25% | SSC-H | PE-Texas Red-H | 1,186,769 | 54,379 |

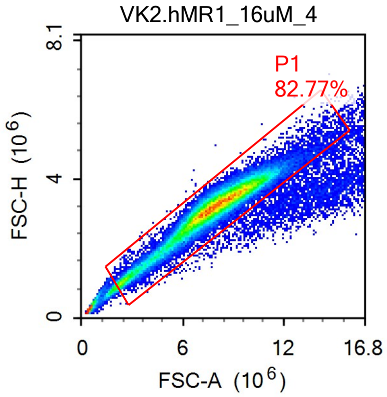


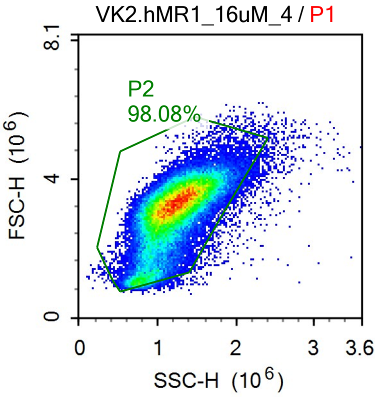


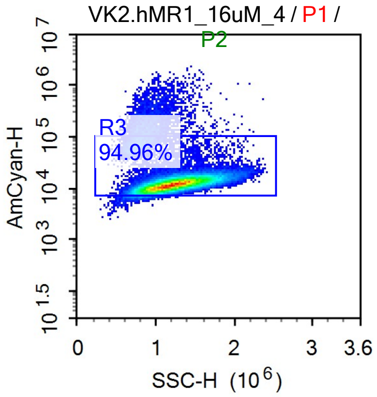


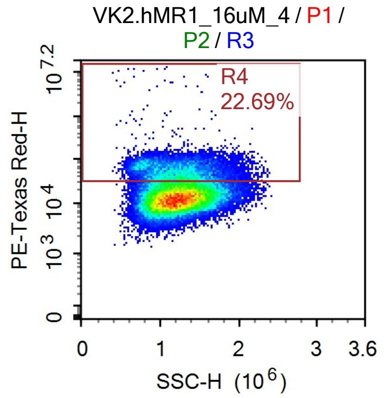


Sample Statistics of VK2.hMR1_16uM_4

| Gate | Count | % Parent | X | Y | Median X | Median Y |
|---|---|---|---|---|---|---|
| All | 73,987 | | | | | |
| P1 | 61,238 | 82.77% | FSC-A | FSC-H | 7,794,790 | 3,189,841 |
| P2 | 60,065 | 98.08% | SSC-H | FSC-H | 1,194,713 | 3,186,915 |
| R3 | 57,037 | 94.96% | SSC-H | AmCyan-H | 1,210,589 | 12,139 |
| R4 | 12,940 | 22.69% | SSC-H | PE-Texas Red-H | 1,214,260 | 53,478 |

### Report of Specimen9-VK2_Ctrl_1

Sample Name: Specimen9-VK2_Ctrl_1 | Run Time: 19-Mar-25 3:49 PM

Cytometer: NovoCyte Quanteon 621181110427 | Software: NovoExpress 1.6.0

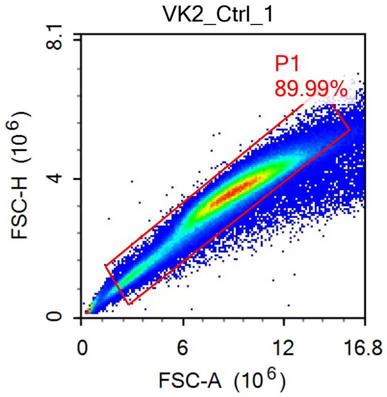


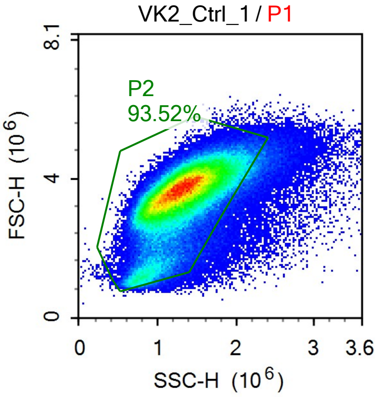


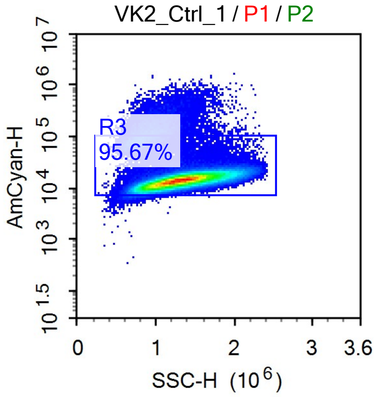


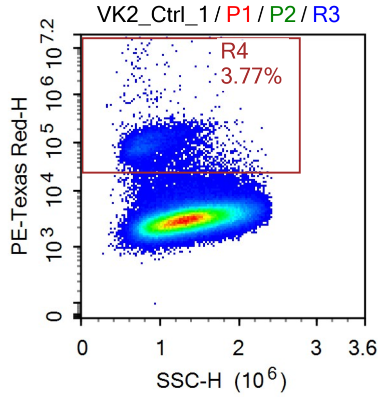


Sample Statistics

| Gate | Count | % Parent | X | Y | Median X | Median Y |
|---|---|---|---|---|---|---|
| All | 236,430 | | | | | |
| P1 | 212,752 | 89.99% | FSC-A | FSC-H | 8,879,873 | 3,587,677 |
| P2 | 198,959 | 93.52% | SSC-H | FSC-H | 1,302,124 | 3,575,605 |
| R3 | 190,352 | 95.67% | SSC-H | AmCyan-H | 1,314,958 | 14,195 |
| R4 | 7,181 | 3.77% | SSC-H | PE-Texas Red-H | 854,772 | 88,679 |

### Report of Specimen9-VK2_Ctrl_2

Sample Name: Specimen9-VK2_Ctrl_2 Run Time: 19-Mar-25 3:52 PM

Cytometer: NovoCyte Quanteon 621181110427 Software: NovoExpress 1.6.0

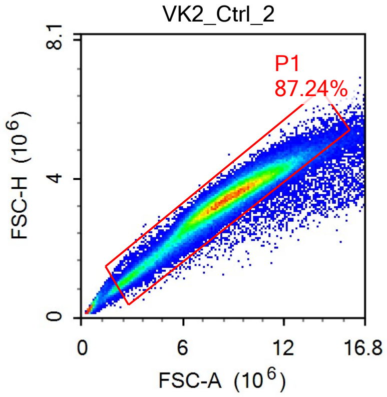


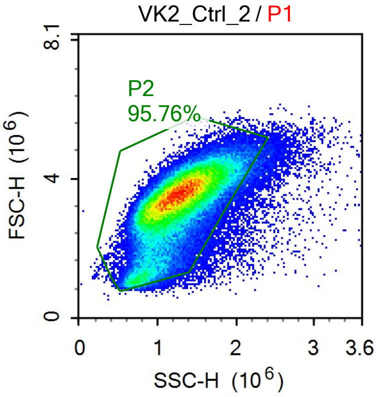


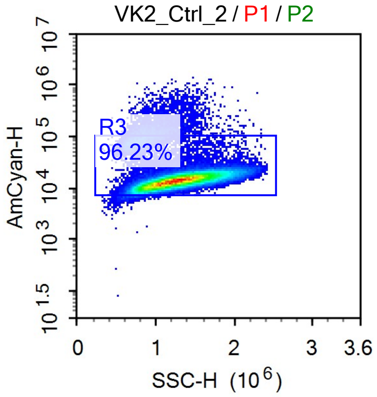


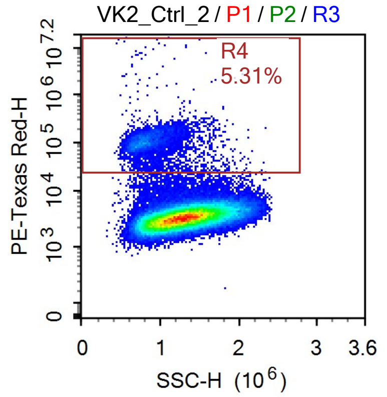


Sample Statistics

| Gate | Count | % Parent | X | Y | Median X | Median Y |
|---|---|---|---|---|---|---|
| All | 114,302 | | | | | |
| P1 | 99,716 | 87.24% | FSC-A | FSC-H | 8,551,798 | 3,407,710 |
| P2 | 95,490 | 95.76% | SSC-H | FSC-H | 1,256,015 | 3,400,918 |
| R3 | 91,893 | 96.23% | SSC-H | AmCyan-H | 1,267,227 | 14,111 |
| R4 | 4,884 | 5.31% | SSC-H | PE-Texas Red-H | 835,463 | 104,330 |

### Report of Specimen9-VK2_Ctrl_3

Sample Name: Specimen9-VK2_Ctrl_3 | Run Time: 19-Mar-25 3:55 PM

Cytometer: NovoCyte Quanteon 621181110427 | Software: NovoExpress 1.6.0

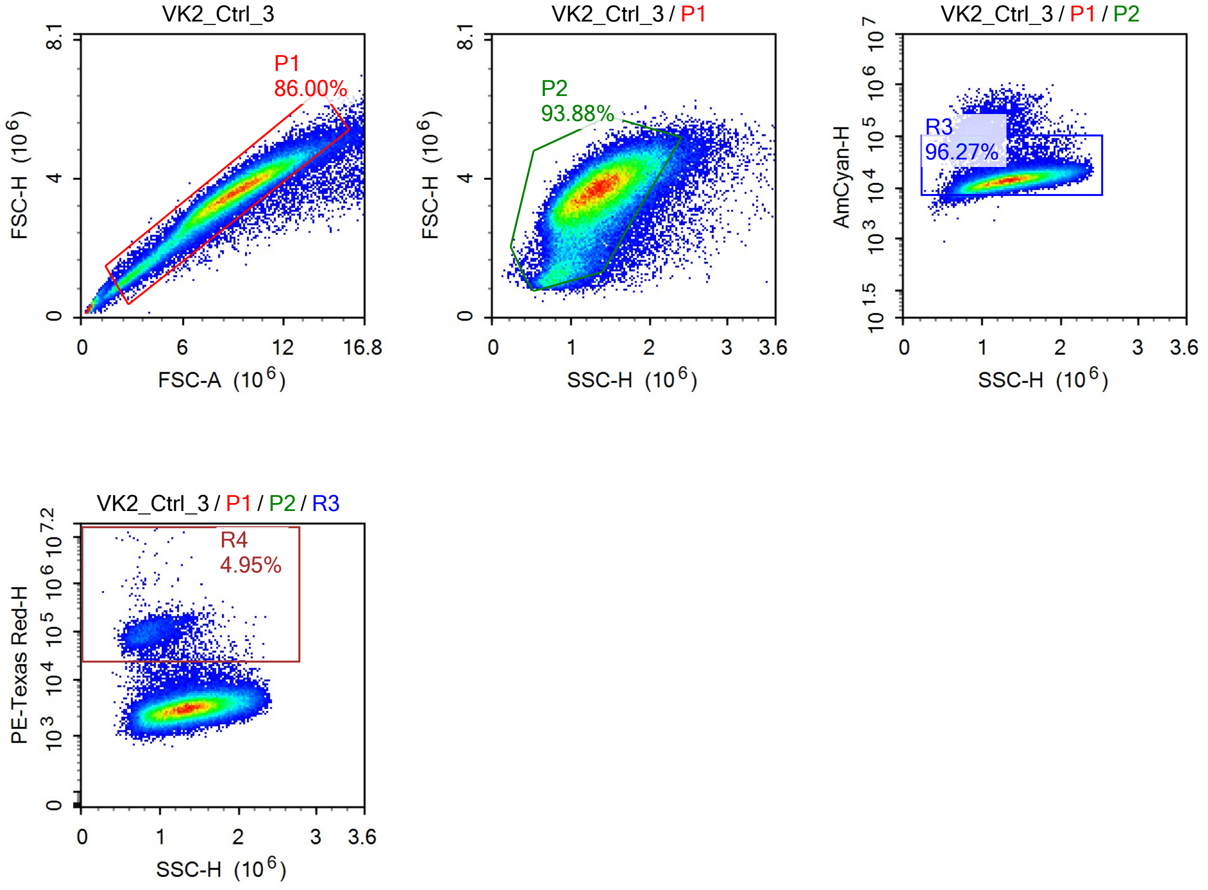


Sample Statistics

| Gate | Count | % Parent | X | Y | Median X | Median Y |
|---|---|---|---|---|---|---|
| All | 88,387 | | | | | |
| P1 | 76,013 | 86.00% | FSC-A | FSC-H | 8,865,887 | 3,522,486 |
| P2 | 71,362 | 93.88% | SSC-H | FSC-H | 1,314,128 | 3,511,782 |
| R3 | 68,701 | 96.27% | SSC-H | AmCyan-H | 1,325,149 | 14,125 |
| R4 | 3,400 | 4.95% | SSC-H | PE-Texas Red-H | 875,729 | 93,993 |

### Report of Specimen9-VK2_Ctrl_4

Sample Name: Specimen9-VK2_Ctrl_4 Run Time: 19-Mar-25 3:58 PM

Cytometer: NovoCyte Quanteon 621181110427 Software: NovoExpress 1.6.0

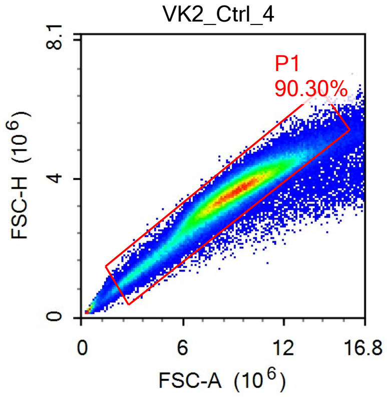


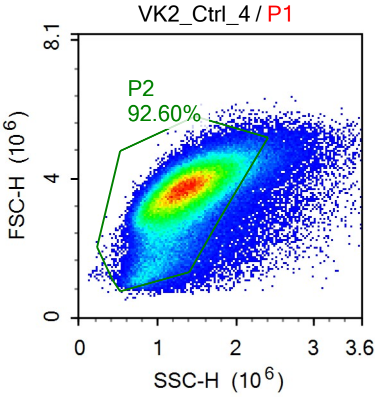


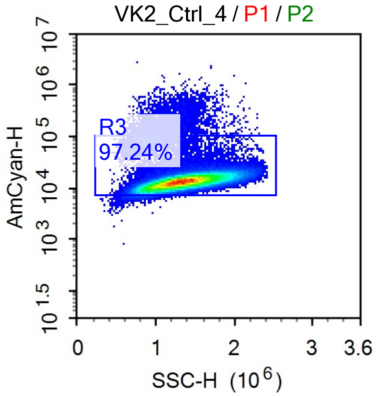


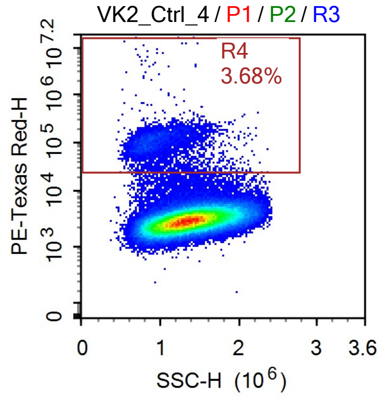


Sample Statistics

| Gate | Count | % Parent | X | Y | Median X | Median Y |
|---|---|---|---|---|---|---|
| All | 166,371 | | | | | |
| P1 | 150,229 | 90.30% | FSC-A | FSC-H | 8,951,205 | 3,596,502 |
| P2 | 139,117 | 92.60% | SSC-H | FSC-H | 1,334,975 | 3,585,551 |
| R3 | 135,273 | 97.24% | SSC-H | AmCyan-H | 1,342,082 | 13,339 |
| R4 | 4,974 | 3.68% | SSC-H | PE-Texas Red-H | 950,596 | 103,154 |

### Report of Specimen9-VK2.hMR1_Ctrl_1

Sample Name: Specimen9-VK2.hMR1_Ctrl_1 | Run Time: 19-Mar-25 4:01 PM

Cytometer: NovoCyte Quanteon 621181110427 | Software: NovoExpress 1.6.0

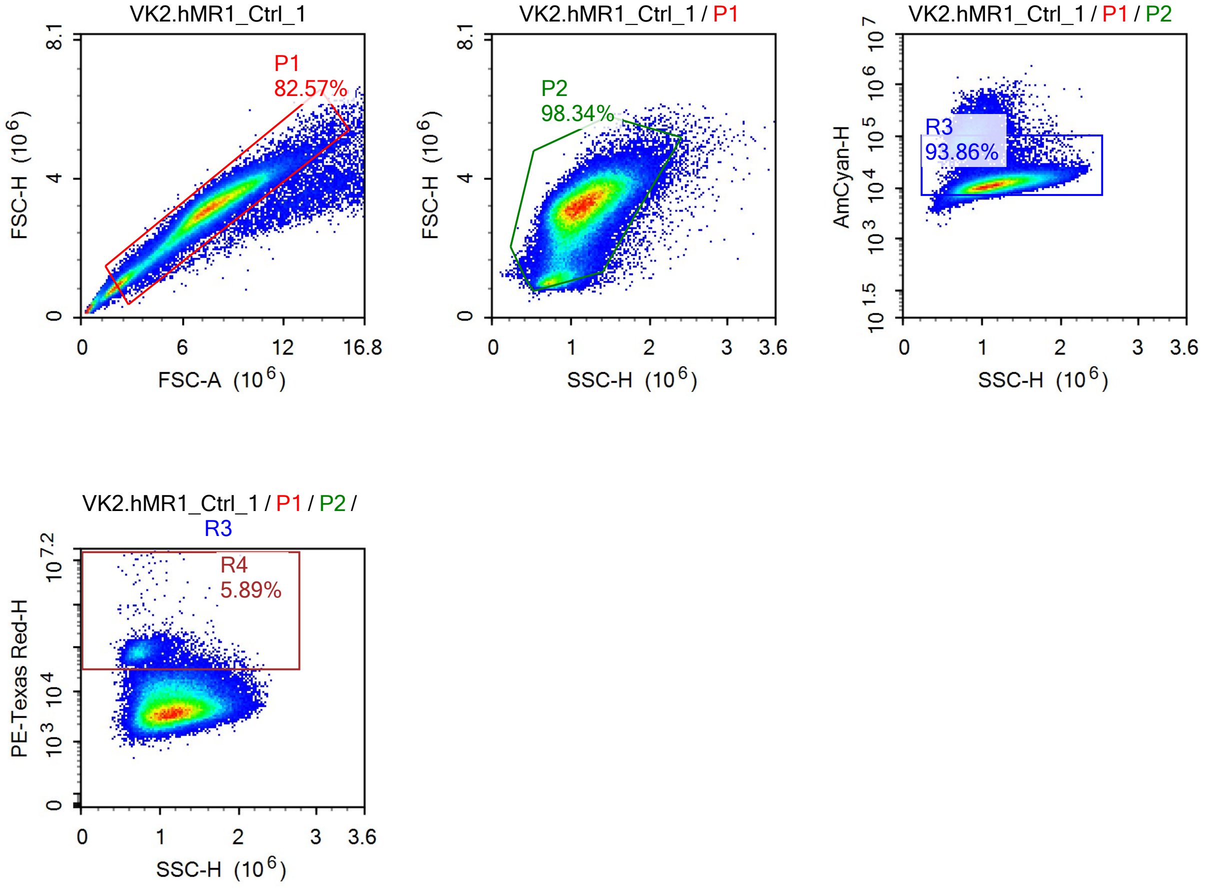


Sample Statistics

| Gate | Count | % Parent | X | Y | Median X | Median Y |
|---|---|---|---|---|---|---|
| All | 110,820 | | | | | |
| P1 | 91,504 | 82.57% | FSC-A | FSC-H | 7,464,061 | 3,110,558 |
| P2 | 89,983 | 98.34% | SSC-H | FSC-H | 1,118,753 | 3,112,641 |
| R3 | 84,460 | 93.86% | SSC-H | AmCyan-H | 1,136,985 | 11,837 |
| R4 | 4,974 | 5.89% | SSC-H | PE-Texas Red-H | 788,171 | 74,291 |

### Report of Specimen9-VK2.hMR1_Ctrl_2

Sample Name: Specimen9-VK2.hMR1_Ctrl_2  Run Time: 19-Mar-25 4:04 PM

Cytometer: NovoCyte Quanteon 621181110427  Software: NovoExpress 1.6.0

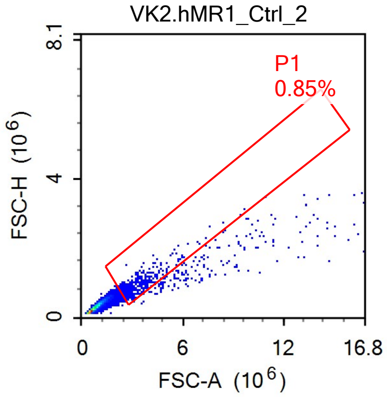


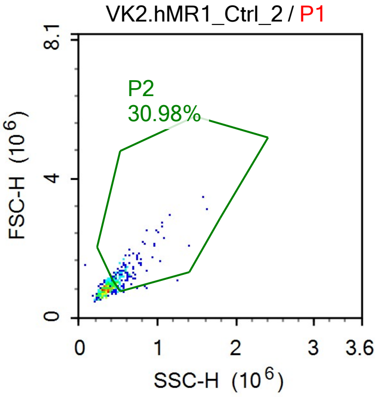


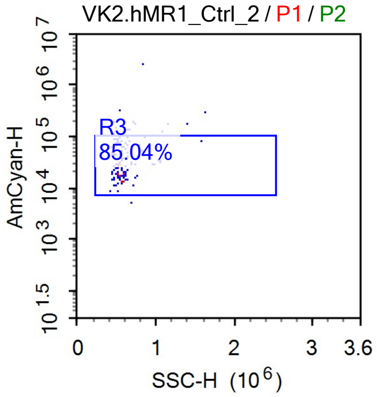


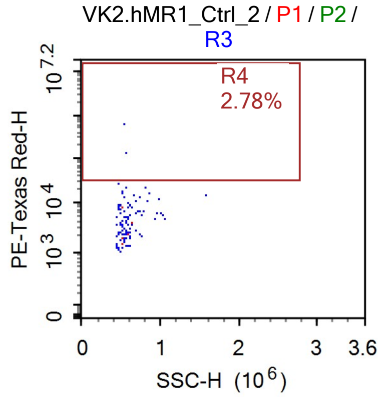


Sample Statistics

| Gate | Count | % Parent | X | Y | Median X | Median Y |
|---|---|---|---|---|---|---|
| All | 48,470 | | | | | |
| P1 | 410 | 0.85% | FSC-A | FSC-H | 2,996,505 | 871,143 |
| P2 | 127 | 30.98% | SSC-H | FSC-H | 552,866 | 1,258,282 |
| R3 | 108 | 85.04% | SSC-H | AmCyan-H | 540,164 | 29,515 |
| R4 | 3 | 2.78% | SSC-H | PE-Texas Red-H | 541,510 | 452,676 |

### Report of Specimen9-VK2.hMR1_Ctrl_3

| | |
|---|---|
| Sample Name: Specimen9-VK2.hMR1_Ctrl_3 | Run Time: 19-Mar-25 4:07 PM |
| Cytometer: NovoCyte Quanteon 621181110427 | Software: NovoExpress 1.6.0 |

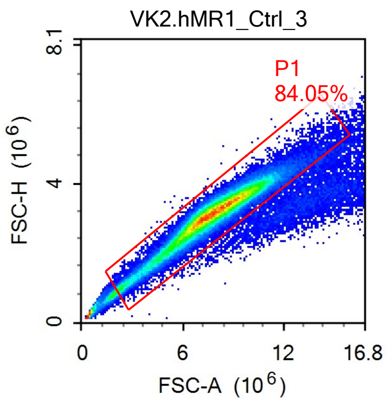


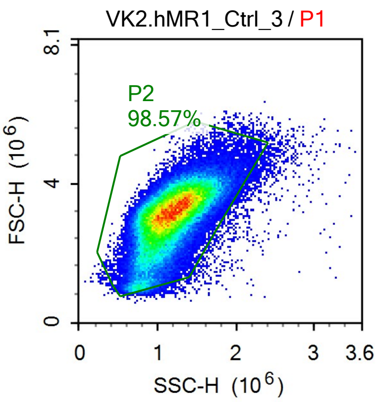


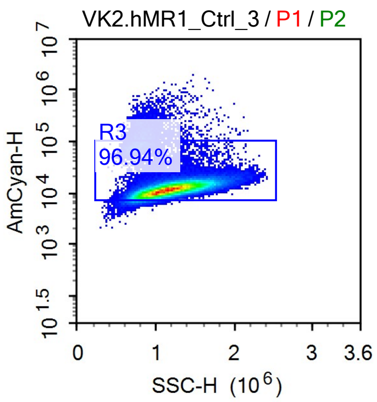


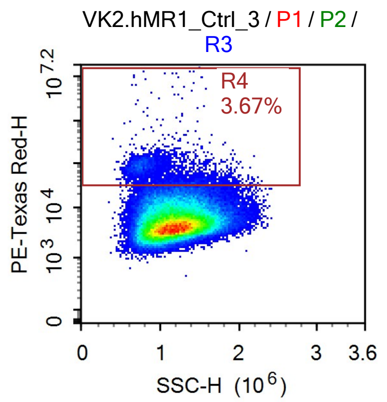


Sample Statistics

| Gate | Count | % Parent | X | Y | Median X | Median Y |
|---|---|---|---|---|---|---|
| All | 120,517 | | | | | |
| P1 | 101,294 | 84.05% | FSC-A | FSC-H | 7,684,813 | 3,129,962 |
| P2 | 99,841 | 98.57% | SSC-H | FSC-H | 1,157,231 | 3,128,508 |
| R3 | 96,789 | 96.94% | SSC-H | AmCyan-H | 1,167,389 | 11,914 |
| R4 | 3,549 | 3.67% | SSC-H | PE-Texas Red-H | 942,784 | 69,466 |

### Report of Specimen9-VK2.hMR1_Ctrl_4

Sample Name: Specimen9-VK2.hMR1_Ctrl_4 Run Time: 19-Mar-25 4:10 PM

Cytometer: NovoCyte Quanteon 621181110427 Software: NovoExpress 1.6.0

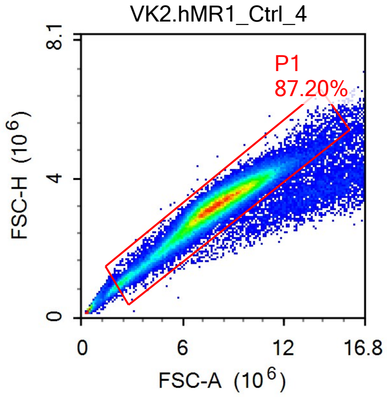


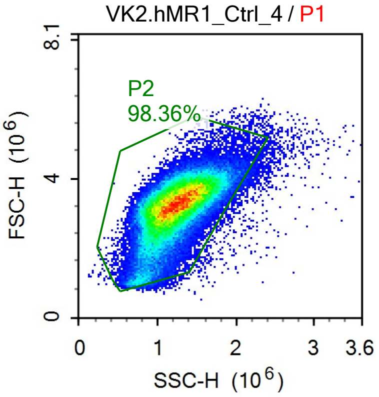


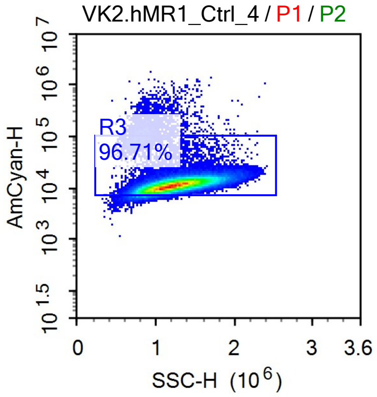


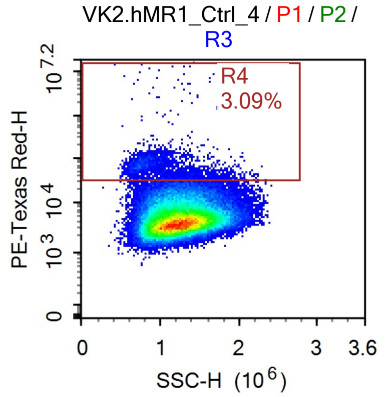


Sample Statistics

| Gate | Count | % Parent | X | Y | Median X | Median Y |
|---|---|---|---|---|---|---|
| All | 95,119 | | | | | |
| P1 | 82,943 | 87.20% | FSC-A | FSC-H | 7,940,226 | 3,239,076 |
| P2 | 81,581 | 98.36% | SSC-H | FSC-H | 1,213,323 | 3,234,310 |
| R3 | 78,898 | 96.71% | SSC-H | AmCyan-H | 1,224,096 | 11,485 |
| R4 | 2,436 | 3.09% | SSC-H | PE-Texas Red-H | 960,324 | 62,378 |

### Report of Specimen9-VK2_LGG_1

Sample Name: Specimen9-VK2_LGG_1 | Run Time: 19-Mar-25 4:13 PM

Cytometer: NovoCyte Quanteon 621181110427 | Software: NovoExpress 1.6.0

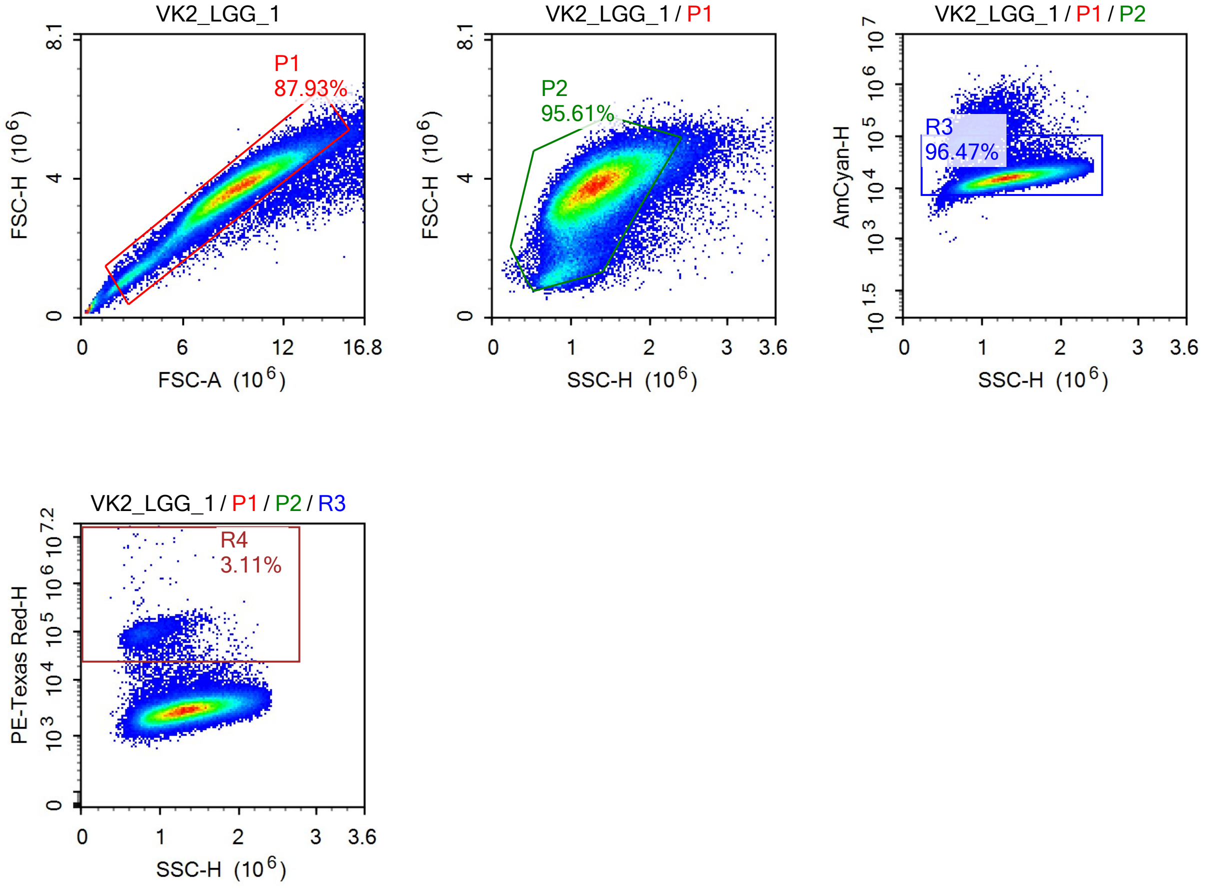


Sample Statistics

| Gate | Count | % Parent | X | Y | Median X | Median Y |
|---|---|---|---|---|---|---|
| All | 122,147 | | | | | |
| P1 | 107,409 | 87.93% | FSC-A | FSC-H | 9,145,303 | 3,703,138 |
| P2 | 102,699 | 95.61% | SSC-H | FSC-H | 1,298,473 | 3,688,976 |
| R3 | 99,077 | 96.47% | SSC-H | AmCyan-H | 1,309,038 | 15,651 |
| R4 | 3,080 | 3.11% | SSC-H | PE-Texas Red-H | 845,145 | 91,949 |

### Report of Specimen9-VK2_LGG_2

Sample Name: Specimen9-VK2_LGG_2 | Run Time: 19-Mar-25 4:15 PM

Cytometer: NovoCyte Quanteon 621181110427 | Software: NovoExpress 1.6.0

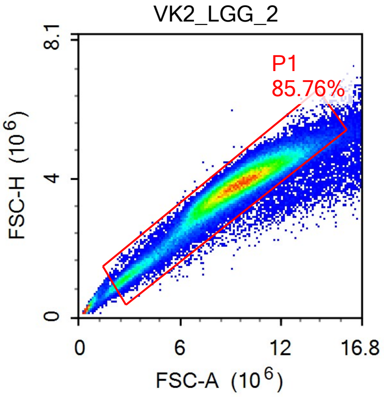


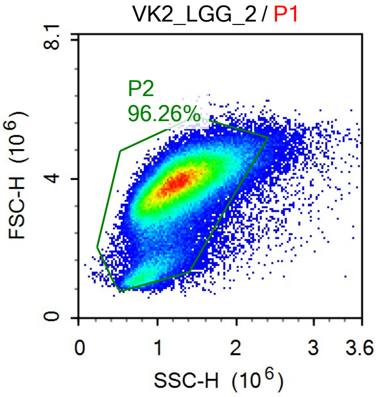


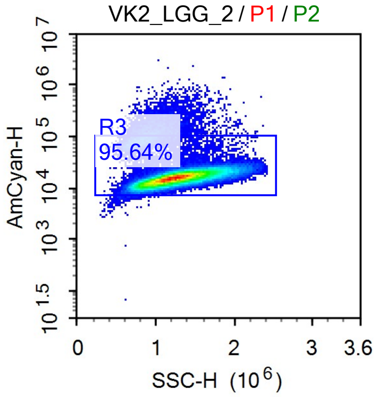


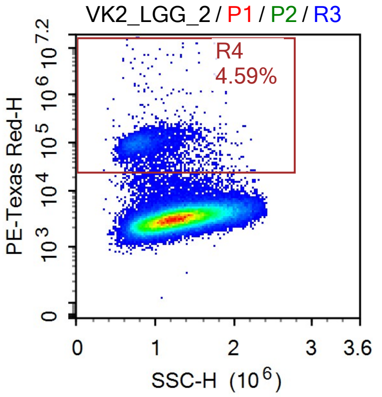


Sample Statistics

| Gate | Count | % Parent | X | Y | Median X | Median Y |
|---|---|---|---|---|---|---|
| All | 113,178 | | | | | |
| P1 | 97,060 | 85.76% | FSC-A | FSC-H | 9,320,344 | 3,790,748 |
| P2 | 93,432 | 96.26% | SSC-H | FSC-H | 1,242,899 | 3,782,672 |
| R3 | 89,359 | 95.64% | SSC-H | AmCyan-H | 1,253,649 | 15,911 |
| R4 | 4,104 | 4.59% | SSC-H | PE-Texas Red-H | 836,983 | 96,529 |

### Report of Specimen9-VK2_LGG_3

Sample Name: Specimen9-VK2_LGG_3 Run Time: 19-Mar-25 4:18 PM

Cytometer: NovoCyte Quanteon 621181110427 Software: NovoExpress 1.6.0

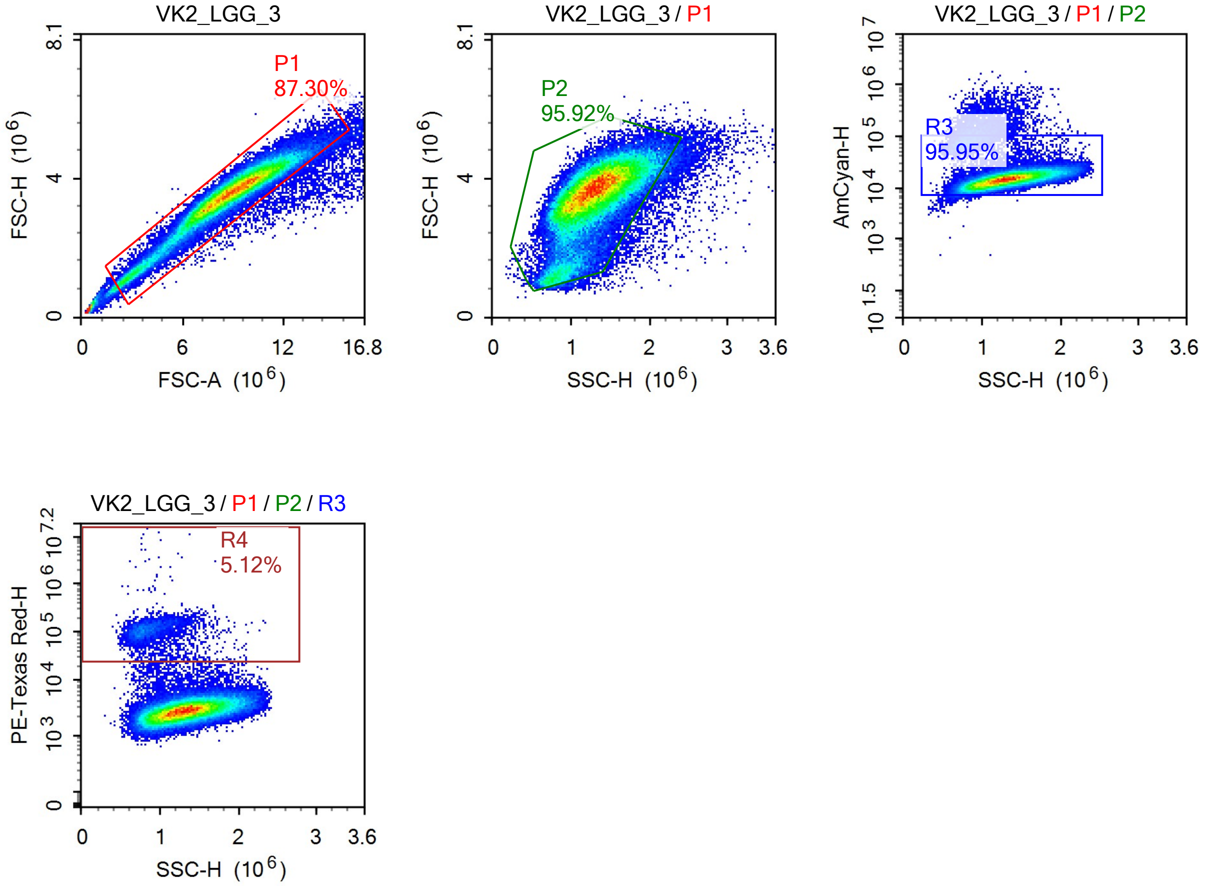


Sample Statistics

| Gate | Count | % Parent | X | Y | Median X | Median Y |
|---|---|---|---|---|---|---|
| All | 83,288 | | | | | |
| P1 | 72,712 | 87.30% | FSC-A | FSC-H | 8,859,694 | 3,593,505 |
| P2 | 69,744 | 95.92% | SSC-H | FSC-H | 1,279,219 | 3,581,043 |
| R3 | 66,916 | 95.95% | SSC-H | AmCyan-H | 1,291,268 | 14,799 |
| R4 | 3,427 | 5.12% | SSC-H | PE-Texas Red-H | 856,218 | 108,660 |

### Report of Specimen9-VK2_LGG_4

Sample Name: Specimen9-VK2_LGG_4 | Run Time: 19-Mar-25 4:21 PM

Cytometer: NovoCyte Quanteon 621181110427 | Software: NovoExpress 1.6.0

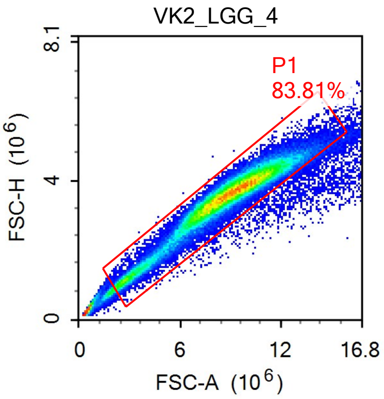


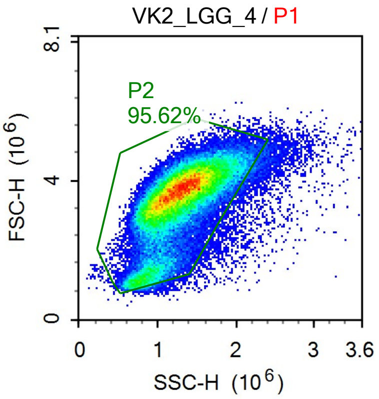


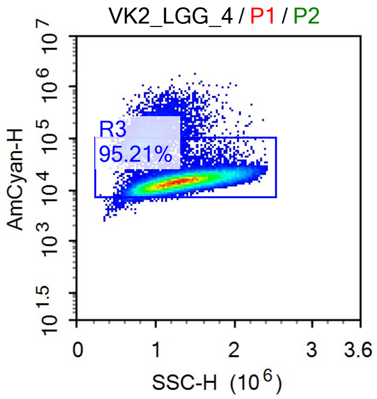


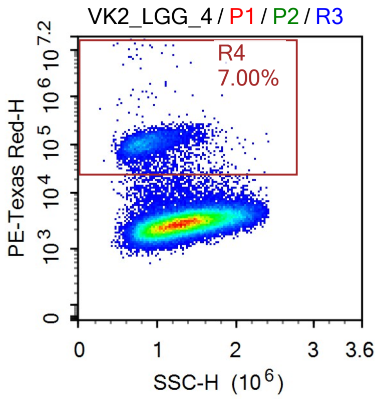


Sample Statistics

| Gate | Count | % Parent | X | Y | Median X | Median Y |
|---|---|---|---|---|---|---|
| All | 83,338 | | | | | |
| P1 | 69,846 | 83.81% | FSC-A | FSC-H | 8,842,944 | 3,600,152 |
| P2 | 66,789 | 95.62% | SSC-H | FSC-H | 1,262,145 | 3,591,099 |
| R3 | 63,587 | 95.21% | SSC-H | AmCyan-H | 1,277,723 | 14,730 |
| R4 | 4,448 | 7.00% | SSC-H | PE-Texas Red-H | 855,166 | 106,400 |

### Report of Specimen9-VK2.hMR1_LGG_1

Sample Name: Specimen9-VK2.hMR1_LGG_1 | Run Time: 19-Mar-25 4:24 PM

Cytometer: NovoCyte Quanteon 621181110427 | Software: NovoExpress 1.6.0

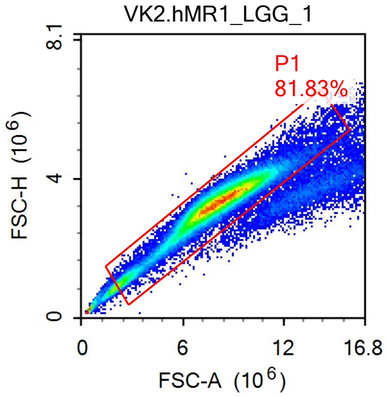


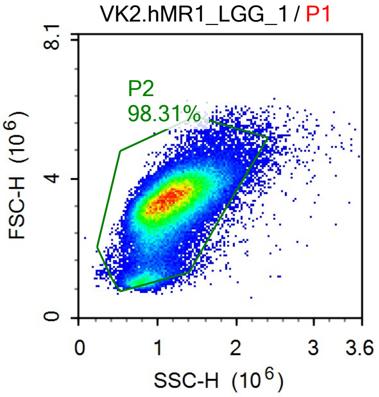


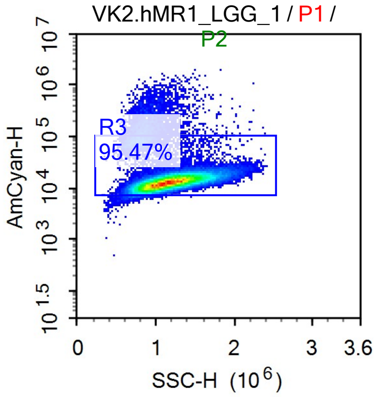


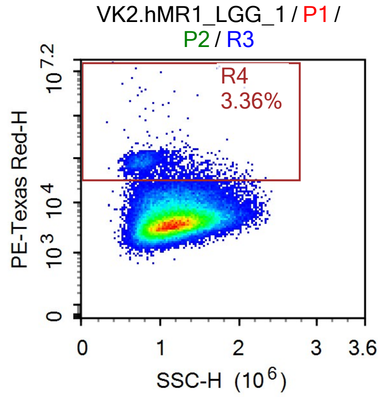


Sample Statistics

| Gate | Count | % Parent | X | Y | Median X | Median Y |
|---|---|---|---|---|---|---|
| All | 86,842 | | | | | |
| P1 | 71,060 | 81.83% | FSC-A | FSC-H | 8,054,100 | 3,327,143 |
| P2 | 69,861 | 98.31% | SSC-H | FSC-H | 1,144,386 | 3,325,115 |
| R3 | 66,693 | 95.47% | SSC-H | AmCyan-H | 1,157,133 | 13,203 |
| R4 | 2,240 | 3.36% | SSC-H | PE-Texas Red-H | 821,768 | 77,958 |

### Report of Specimen9-VK2.hMR1_LGG_2

Sample Name: Specimen9-VK2.hMR1_LGG_2 | Run Time: 19-Mar-25 4:27 PM

Cytometer: NovoCyte Quanteon 621181110427 | Software: NovoExpress 1.6.0

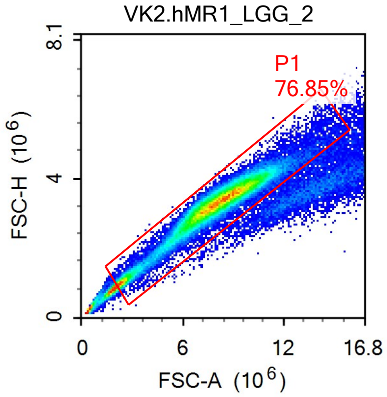


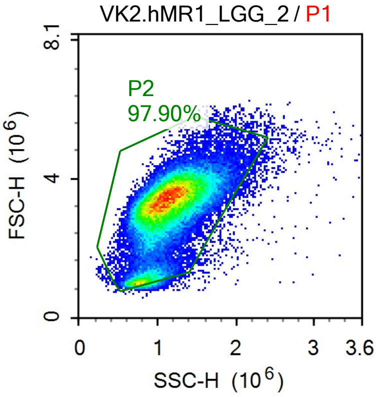


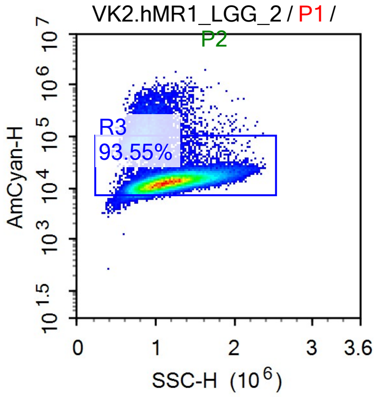


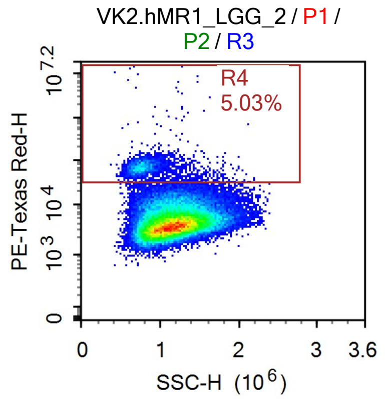


Sample Statistics

| Gate | Count | % Parent | X | Y | Median X | Median Y |
|---|---|---|---|---|---|---|
| All | 80,699 | | | | | |
| P1 | 62,016 | 76.85% | FSC-A | FSC-H | 8,005,536 | 3,302,596 |
| P2 | 60,716 | 97.90% | SSC-H | FSC-H | 1,118,008 | 3,302,219 |
| R3 | 56,799 | 93.55% | SSC-H | AmCyan-H | 1,135,033 | 13,244 |
| R4 | 2,859 | 5.03% | SSC-H | PE-Texas Red-H | 779,920 | 67,902 |

### Report of Specimen9-VK2.hMR1_LGG_3

Sample Name: Specimen9-VK2.hMR1_LGG_3 | Run Time: 19-Mar-25 4:30 PM

Cytometer: NovoCyte Quanteon 621181110427 | Software: NovoExpress 1.6.0

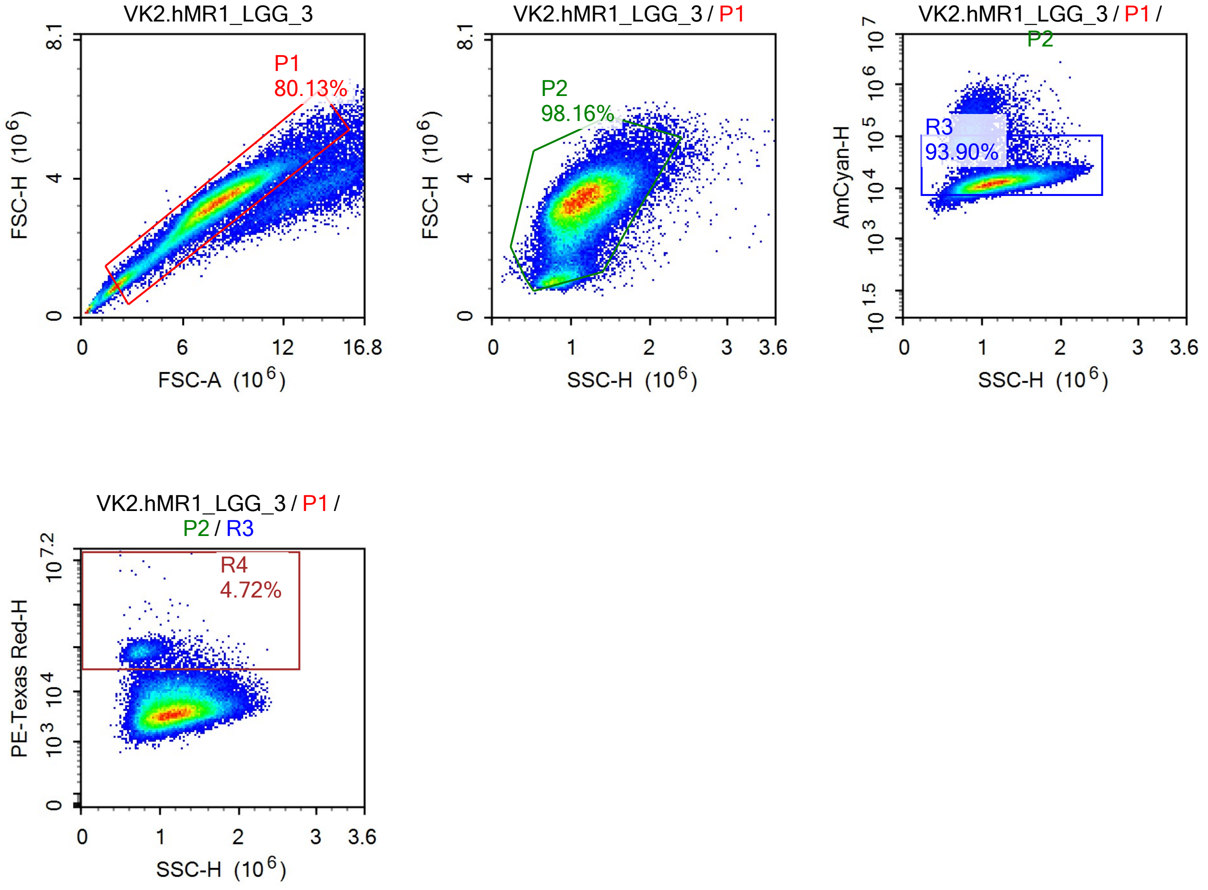


Sample Statistics

| Gate | Count | % Parent | X | Y | Median X | Median Y |
|---|---|---|---|---|---|---|
| All | 76,312 | | | | | |
| P1 | 61,151 | 80.13% | FSC-A | FSC-H | 7,862,102 | 3,266,130 |
| P2 | 60,027 | 98.16% | SSC-H | FSC-H | 1,127,765 | 3,263,521 |
| R3 | 56,367 | 93.90% | SSC-H | AmCyan-H | 1,145,658 | 12,826 |
| R4 | 2,662 | 4.72% | SSC-H | PE-Texas Red-H | 796,813 | 78,297 |

### Report of Specimen9-VK2.hMR1_LGG_4

Sample Name: Specimen9-VK2.hMR1_LGG_4 | Run Time: 19-Mar-25 4:32 PM

Cytometer: NovoCyte Quanteon 621181110427 | Software: NovoExpress 1.6.0

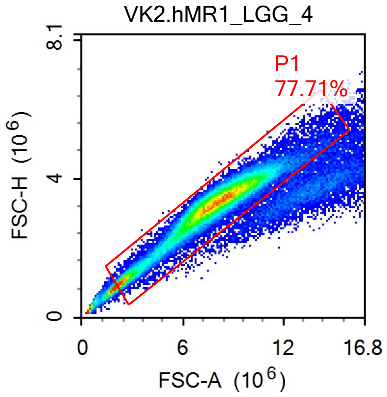


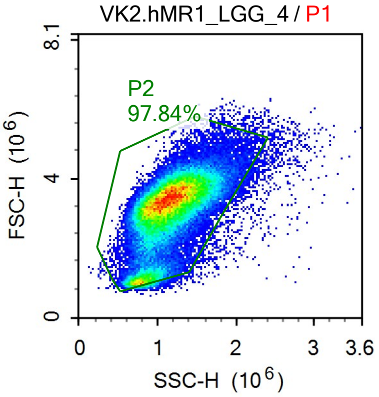


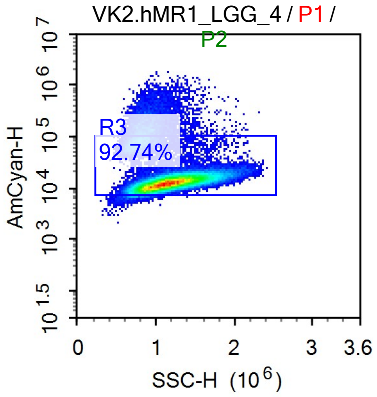


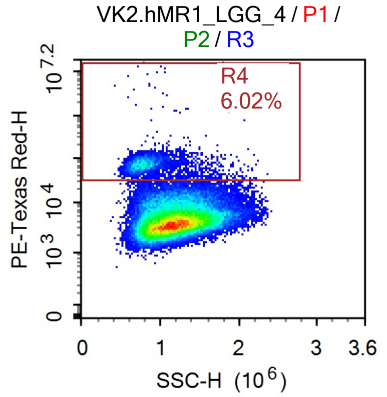


Sample Statistics

| Gate | Count | % Parent | X | Y | Median X | Median Y |
|---|---|---|---|---|---|---|
| All | 88,322 | | | | | |
| P1 | 68,633 | 77.71% | FSC-A | FSC-H | 7,834,561 | 3,274,430 |
| P2 | 67,148 | 97.84% | SSC-H | FSC-H | 1,132,345 | 3,272,088 |
| R3 | 62,274 | 92.74% | SSC-H | AmCyan-H | 1,152,421 | 12,938 |
| R4 | 3,752 | 6.02% | SSC-H | PE-Texas Red-H | 797,192 | 71,607 |

### Report of Specimen9-VK2_AMBV336_1

Sample Name: Specimen9-VK2_AMBV336_1

Run Time: 19-Mar-25 4:35 PM

Cytometer: NovoCyte Quanteon 621181110427

Software: NovoExpress 1.6.0

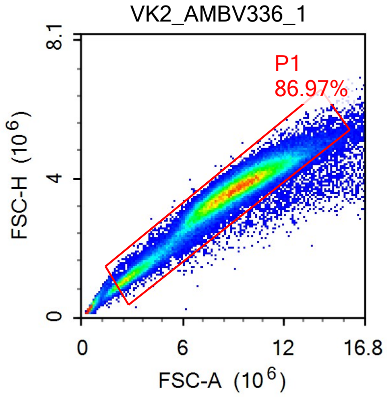


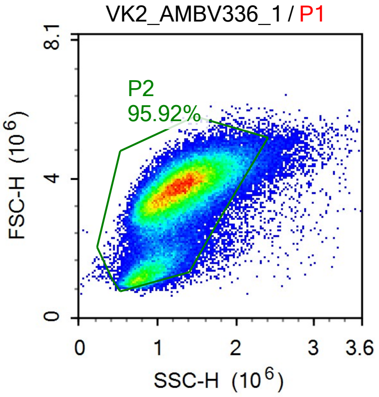


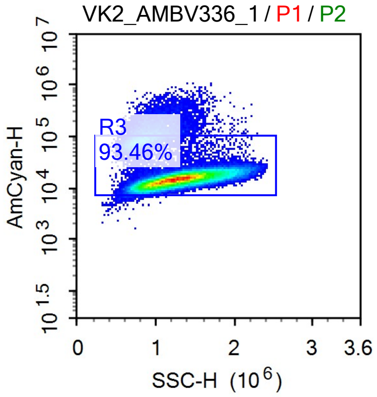


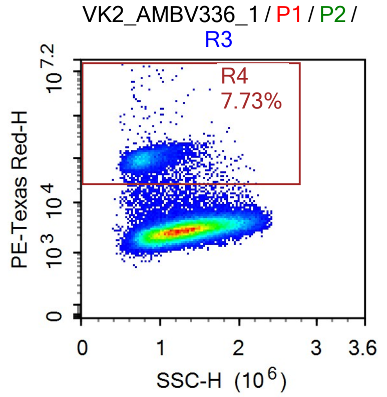


Sample Statistics

| Gate | Count | % Parent | X | Y | Median X | Median Y |
|---|---|---|---|---|---|---|
| All | 92,640 | | | | | |
| P1 | 80,570 | 86.97% | FSC-A | FSC-H | 8,809,750 | 3,612,630 |
| P2 | 77,282 | 95.92% | SSC-H | FSC-H | 1,237,389 | 3,600,111 |
| R3 | 72,224 | 93.46% | SSC-H | AmCyan-H | 1,258,022 | 15,047 |
| R4 | 5,584 | 7.73% | SSC-H | PE-Texas Red-H | 802,787 | 95,075 |

### Report of Specimen9-VK2_AMBV336_2

Sample Name: Specimen9-VK2_AMBV336_2 | Run Time: 19-Mar-25 4:38 PM

Cytometer: NovoCyte Quanteon 621181110427 | Software: NovoExpress 1.6.0

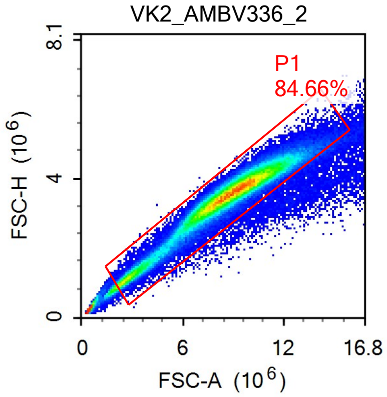


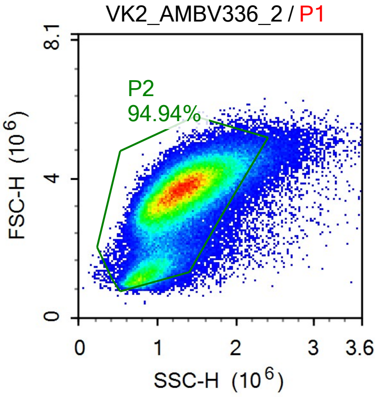


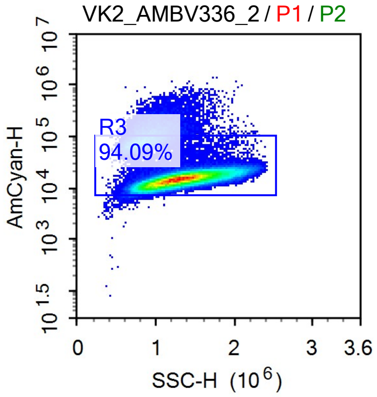


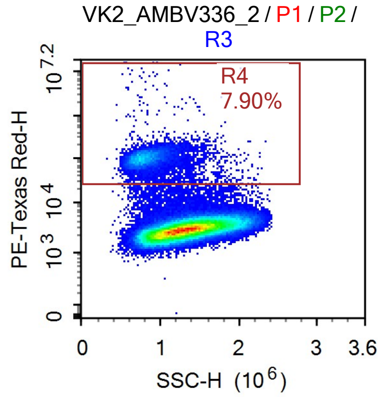


Sample Statistics

| Gate | Count | % Parent | X | Y | Median X | Median Y |
|---|---|---|---|---|---|---|
| All | 148,482 | | | | | |
| P1 | 125,709 | 84.66% | FSC-A | FSC-H | 8,734,182 | 3,539,809 |
| P2 | 119,350 | 94.94% | SSC-H | FSC-H | 1,267,718 | 3,525,008 |
| R3 | 112,295 | 94.09% | SSC-H | AmCyan-H | 1,285,940 | 15,187 |
| R4 | 8,866 | 7.90% | SSC-H | PE-Texas Red-H | 824,938 | 100,256 |

### Report of Specimen9-VK2_AMBV336_3

Sample Name: Specimen9-VK2_AMBV336_3 Run Time: 19-Mar-25 4:41 PM

Cytometer: NovoCyte Quanteon 621181110427 Software: NovoExpress 1.6.0

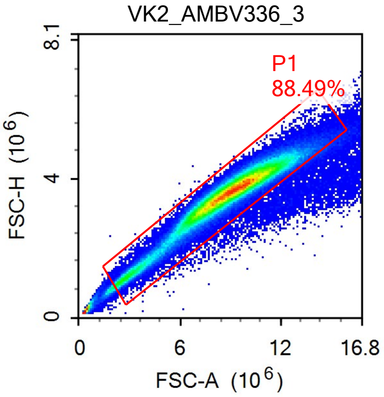


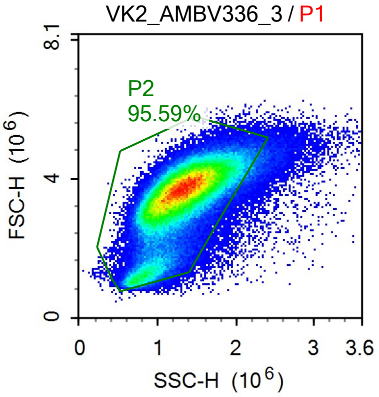


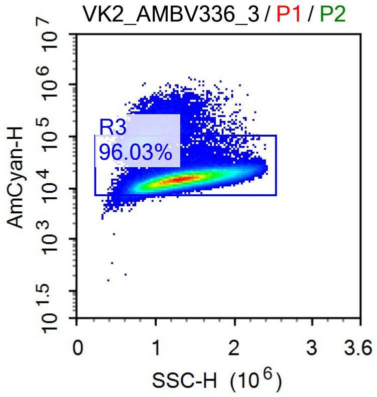


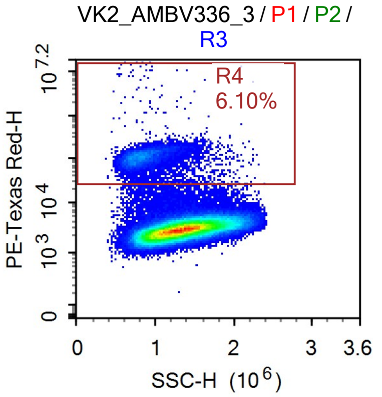


Sample Statistics

| Gate | Count | % Parent | X | Y | Median X | Median Y |
|---|---|---|---|---|---|---|
| All | 199,300 | | | | | |
| P1 | 176,363 | 88.49% | FSC-A | FSC-H | 8,922,351 | 3,604,883 |
| P2 | 168,578 | 95.59% | SSC-H | FSC-H | 1,295,085 | 3,587,765 |
| R3 | 161,878 | 96.03% | SSC-H | AmCyan-H | 1,307,231 | 15,026 |
| R4 | 9,871 | 6.10% | SSC-H | PE-Texas Red-H | 849,921 | 107,038 |

### Report of Specimen9-VK2_AMBV336_4

Sample Name: Specimen9-VK2_AMBV336_4 | Run Time: 19-Mar-25 5:01 PM

Cytometer: NovoCyte Quanteon 621181110427 | Software: NovoExpress 1.6.0

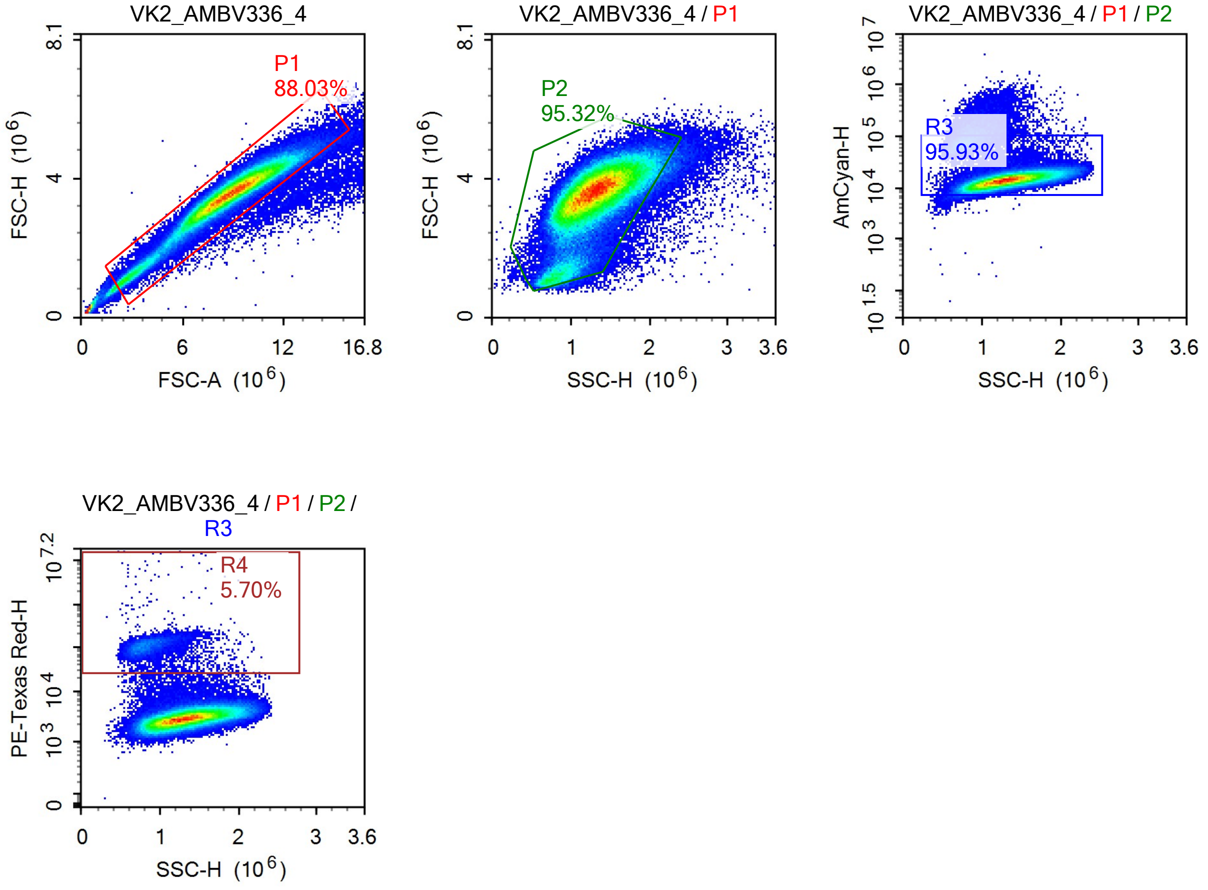


Sample Statistics

| Gate | Count | % Parent | X | Y | Median X | Median Y |
|---|---|---|---|---|---|---|
| All | 155,564 | | | | | |
| P1 | 136,943 | 88.03% | FSC-A | FSC-H | 8,802,005 | 3,565,066 |
| P2 | 130,528 | 95.32% | SSC-H | FSC-H | 1,288,307 | 3,547,654 |
| R3 | 125,210 | 95.93% | SSC-H | AmCyan-H | 1,302,226 | 14,259 |
| R4 | 7,143 | 5.70% | SSC-H | PE-Texas Red-H | 854,828 | 108,467 |

### Report of Specimen9-VK2.hMR1_AMBV336_

Sample Name: Specimen9-VK2.hMR1_AMBV336_1 Run Time: 19-Mar-25 5:04 PM

Cytometer: NovoCyte Quanteon 621181110427 Software: NovoExpress 1.6.0

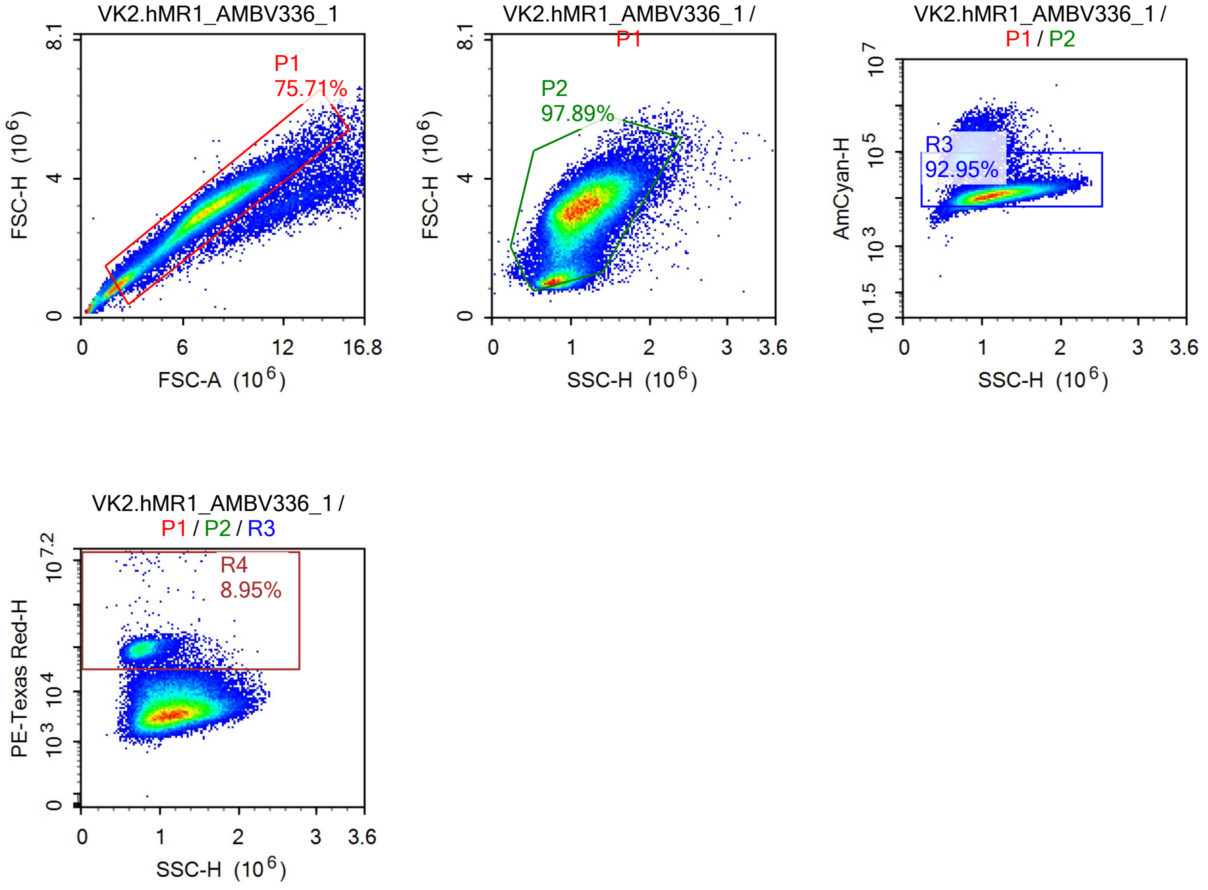


Sample Statistics

| Gate | Count | % Parent | X | Y | Median X | Median Y |
|---|---|---|---|---|---|---|
| All | 86,856 | | | | | |
| P1 | 65,757 | 75.71% | FSC-A | FSC-H | 7,267,949 | 3,012,623 |
| P2 | 64,370 | 97.89% | SSC-H | FSC-H | 1,101,514 | 3,019,189 |
| R3 | 59,831 | 92.95% | SSC-H | AmCyan-H | 1,123,232 | 12,826 |
| R4 | 5,355 | 8.95% | SSC-H | PE-Texas Red-H | 786,167 | 91,461 |

### Report of Specimen9-VK2.hMR1_AMBV336_

Sample Name: Specimen9-VK2.hMR1_AMBV336_2 Run Time: 19-Mar-25 5:07 PM

Cytometer: NovoCyte Quanteon 621181110427 Software: NovoExpress 1.6.0

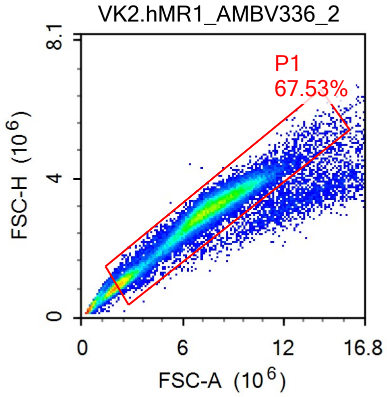


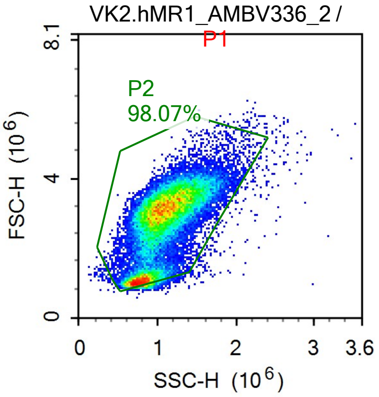


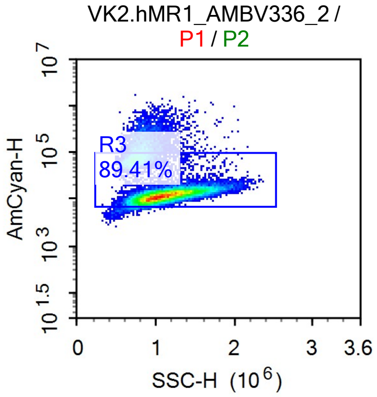


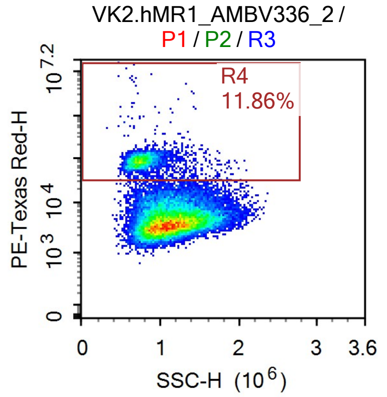


Sample Statistics

| Gate | Count | % Parent | X | Y | Median X | Median Y |
|---|---|---|---|---|---|---|
| All | 52,472 | | | | | |
| P1 | 35,435 | 67.53% | FSC-A | FSC-H | 7,037,553 | 2,953,829 |
| P2 | 34,751 | 98.07% | SSC-H | FSC-H | 1,045,420 | 2,964,692 |
| R3 | 31,071 | 89.41% | SSC-H | AmCyan-H | 1,075,594 | 12,806 |
| R4 | 3,684 | 11.86% | SSC-H | PE-Texas Red-H | 751,749 | 87,563 |

### Report of Specimen9-VK2.hMR1_AMBV336_

Sample Name: Specimen9-VK2.hMR1_AMBV336_3 Run Time: 19-Mar-25 5:10 PM

Cytometer: NovoCyte Quanteon 621181110427 Software: NovoExpress 1.6.0

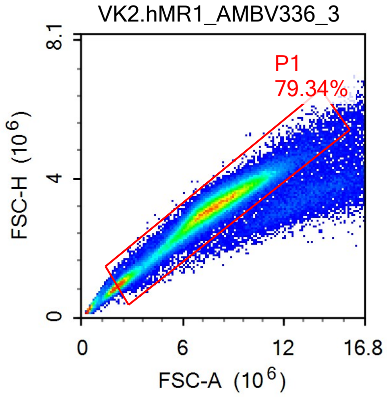


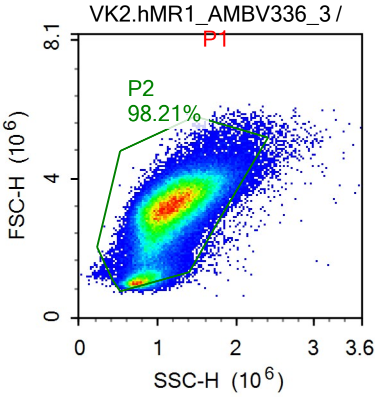


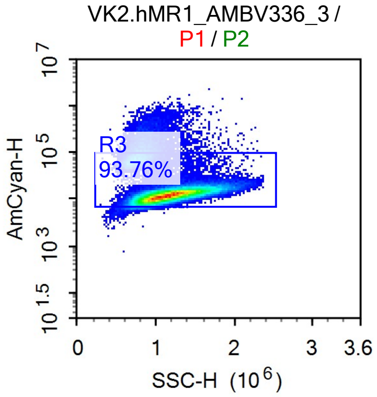


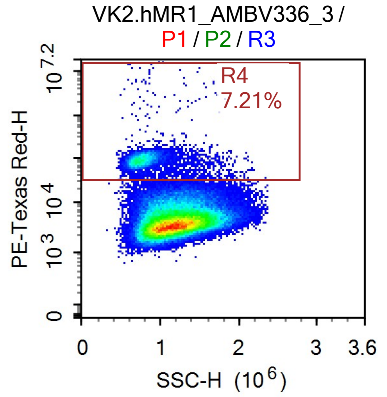


Sample Statistics

| Gate | Count | % Parent | X | Y | Median X | Median Y |
|---|---|---|---|---|---|---|
| All | 126,508 | | | | | |
| P1 | 100,372 | 79.34% | FSC-A | FSC-H | 7,608,183 | 3,129,788 |
| P2 | 98,573 | 98.21% | SSC-H | FSC-H | 1,135,944 | 3,128,204 |
| R3 | 92,426 | 93.76% | SSC-H | AmCyan-H | 1,156,599 | 12,909 |
| R4 | 6,668 | 7.21% | SSC-H | PE-Texas Red-H | 775,193 | 90,068 |

### Report of Specimen9-VK2.hMR1_AMBV336_

Sample Name: Specimen9-VK2.hMR1_AMBV336_4 | Run Time: 19-Mar-25 5:12 PM

Cytometer: NovoCyte Quanteon 621181110427 | Software: NovoExpress 1.6.0

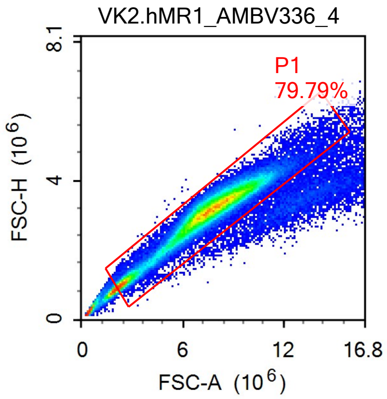


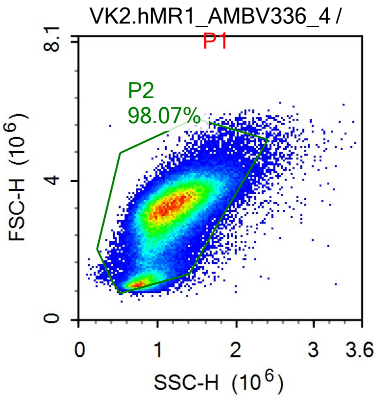


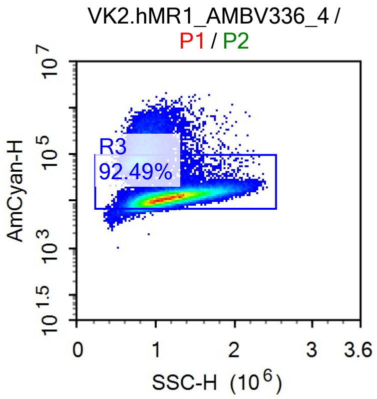


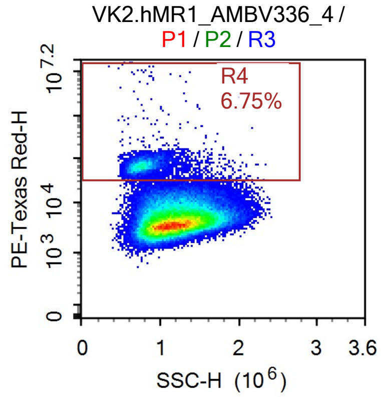


Sample Statistics

| Gate | Count | % Parent | X | Y | Median X | Median Y |
|---|---|---|---|---|---|---|
| All | 115,767 | | | | | |
| P1 | 92,367 | 79.79% | FSC-A | FSC-H | 7,483,276 | 3,127,722 |
| P2 | 90,585 | 98.07% | SSC-H | FSC-H | 1,131,298 | 3,127,765 |
| R3 | 83,780 | 92.49% | SSC-H | AmCyan-H | 1,156,123 | 12,483 |
| R4 | 5,654 | 6.75% | SSC-H | PE-Texas Red-H | 780,858 | 69,052 |

### Report of Specimen9-VK2_AMBV339_1

Sample Name: Specimen9-VK2_AMBV339_1 | Run Time: 19-Mar-25 5:14 PM

Cytometer: NovoCyte Quanteon 621181110427 | Software: NovoExpress 1.6.0

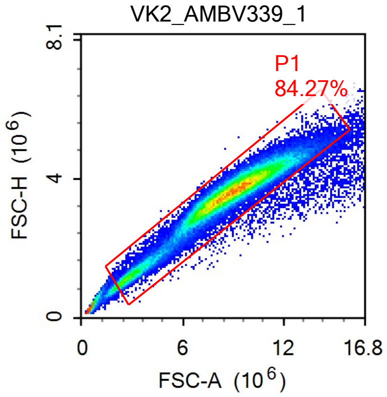


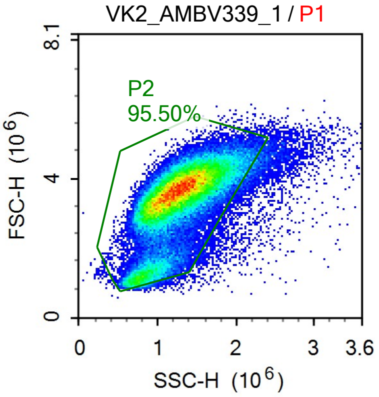


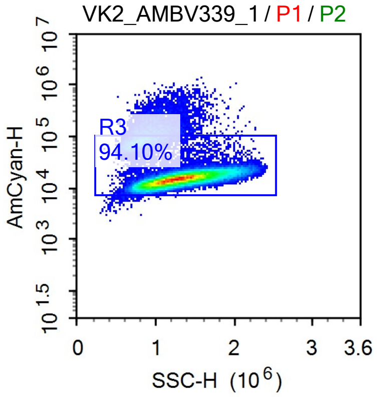


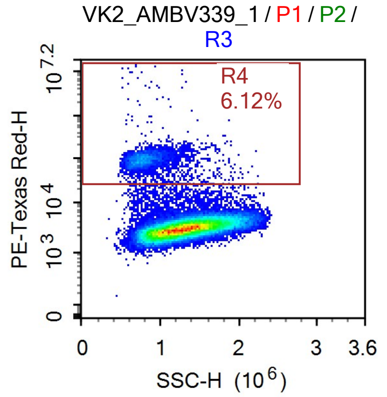


Sample Statistics

| Gate | Count | % Parent | X | Y | Median X | Median Y |
|---|---|---|---|---|---|---|
| All | 84,837 | | | | | |
| P1 | 71,488 | 84.27% | FSC-A | FSC-H | 8,807,462 | 3,596,674 |
| P2 | 68,269 | 95.50% | SSC-H | FSC-H | 1,257,905 | 3,582,933 |
| R3 | 64,240 | 94.10% | SSC-H | AmCyan-H | 1,275,515 | 15,260 |
| R4 | 3,933 | 6.12% | SSC-H | PE-Texas Red-H | 800,026 | 94,913 |

### Report of Specimen9-VK2_AMBV339_2

Sample Name: Specimen9-VK2_AMBV339_2 | Run Time: 19-Mar-25 5:16 PM
Cytometer: NovoCyte Quanteon 621181110427 | Software: NovoExpress 1.6.0

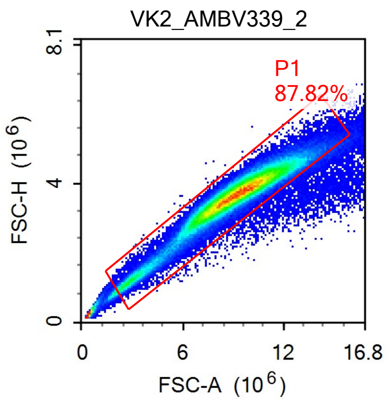


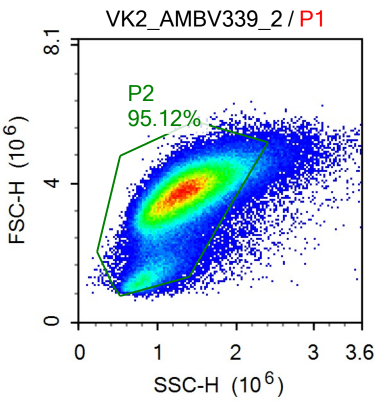


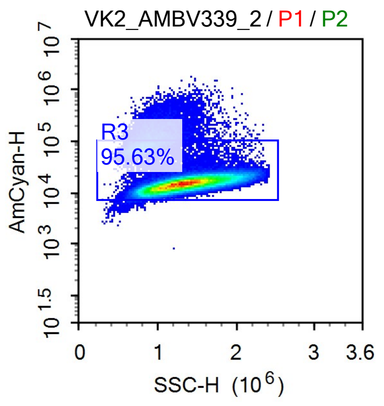


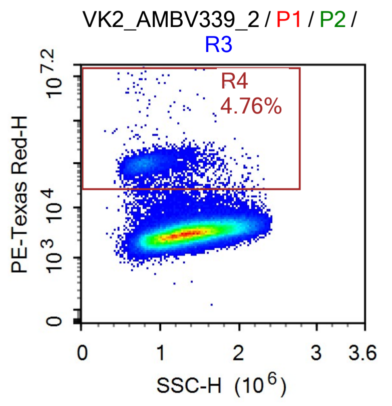


Sample Statistics

| Gate | Count | % Parent | X | Y | Median X | Median Y |
|---|---|---|---|---|---|---|
| All | 157,565 | | | | | |
| P1 | 138,367 | 87.82% | FSC-A | FSC-H | 9,128,234 | 3,678,816 |
| P2 | 131,616 | 95.12% | SSC-H | FSC-H | 1,308,204 | 3,657,370 |
| R3 | 125,870 | 95.63% | SSC-H | AmCyan-H | 1,321,666 | 15,199 |
| R4 | 5,994 | 4.76% | SSC-H | PE-Texas Red-H | 817,091 | 98,017 |

### Report of Specimen9-VK2_AMBV339_3

Sample Name: Specimen9-VK2_AMBV339_3 | Run Time: 19-Mar-25 5:18 PM

Cytometer: NovoCyte Quanteon 621181110427 | Software: NovoExpress 1.6.0

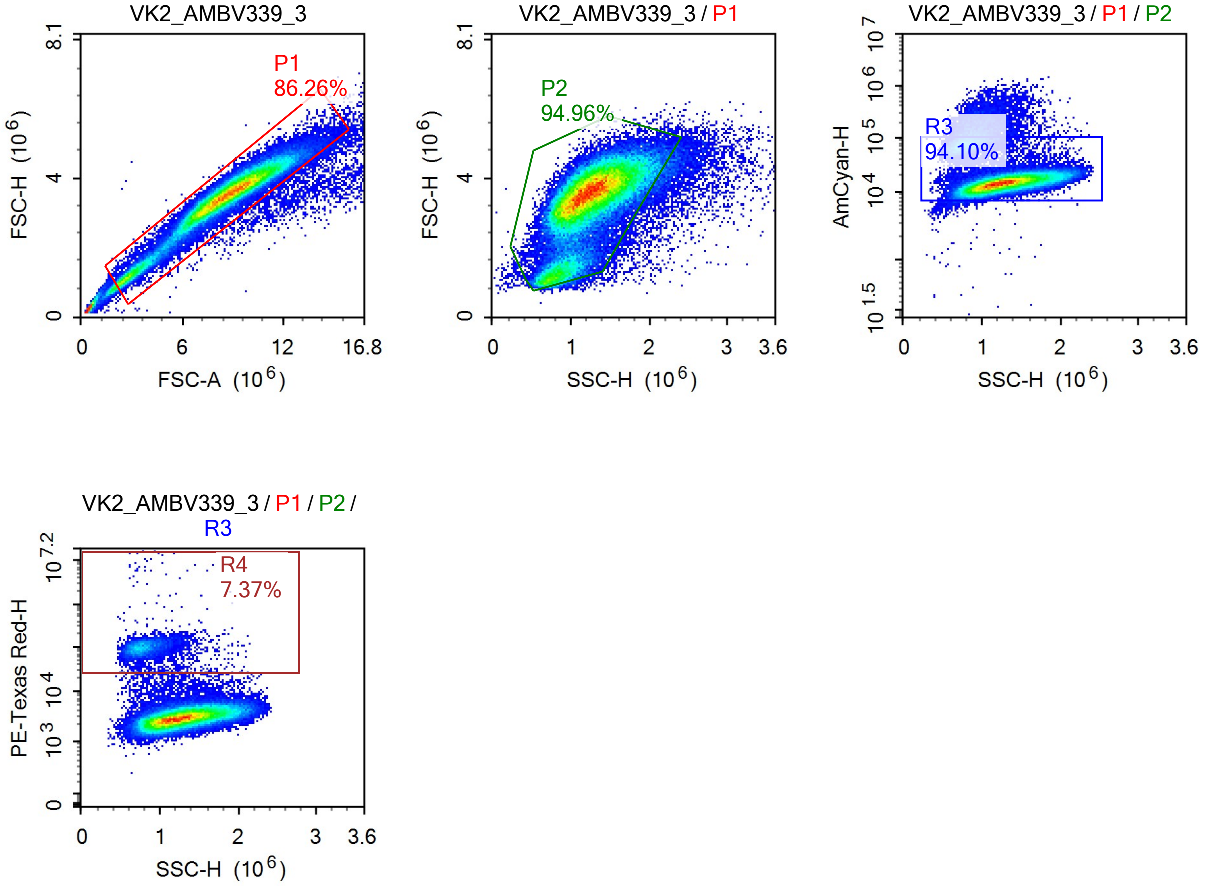


Sample Statistics

| Gate | Count | % Parent | X | Y | Median X | Median Y |
|---|---|---|---|---|---|---|
| All | 96,722 | | | | | |
| P1 | 83,432 | 86.26% | FSC-A | FSC-H | 8,598,250 | 3,508,558 |
| P2 | 79,228 | 94.96% | SSC-H | FSC-H | 1,225,679 | 3,494,444 |
| R3 | 74,553 | 94.10% | SSC-H | AmCyan-H | 1,243,158 | 14,864 |
| R4 | 5,492 | 7.37% | SSC-H | PE-Texas Red-H | 790,538 | 97,550 |

### Report of Specimen9-VK2_AMBV339_4

Sample Name: Specimen9-VK2_AMBV339_4 | Run Time: 19-Mar-25 5:21 PM

Cytometer: NovoCyte Quanteon 621181110427 | Software: NovoExpress 1.6.0

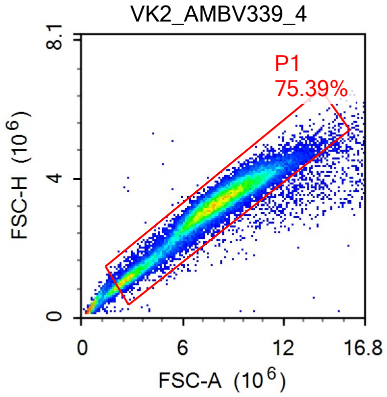


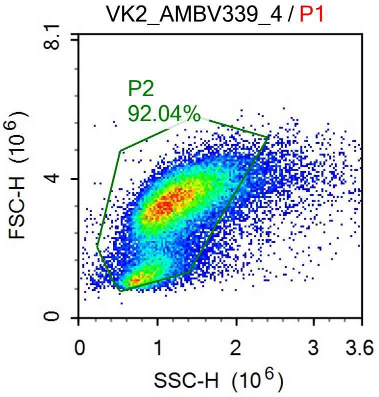


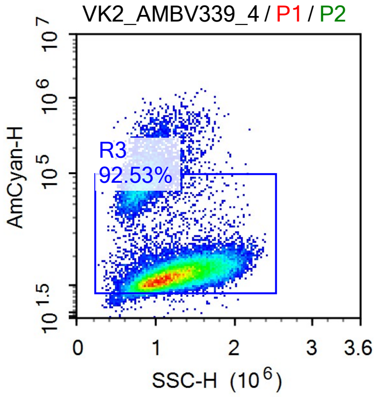


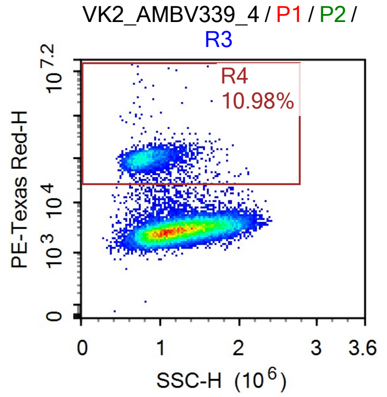


Sample Statistics

| Gate | Count | % Parent | X | Y | Median X | Median Y |
|---|---|---|---|---|---|---|
| All | 45,141 | | | | | |
| P1 | 34,033 | 75.39% | FSC-A | FSC-H | 7,715,882 | 3,200,911 |
| P2 | 31,325 | 92.04% | SSC-H | FSC-H | 1,154,114 | 3,188,625 |
| R3 | 28,985 | 92.53% | SSC-H | AmCyan-H | 1,174,134 | 13,331 |
| R4 | 3,183 | 10.98% | SSC-H | PE-Texas Red-H | 808,293 | 99,011 |

### Report of Specimen9-VK2.hMR1_AMBV339_

Sample Name: Specimen9-VK2.hMR1_AMBV339_1 Run Time: 19-Mar-25 5:23 PM

Cytometer: NovoCyte Quanteon 621181110427 Software: NovoExpress 1.6.0

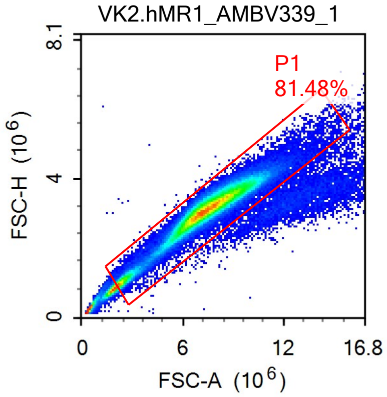


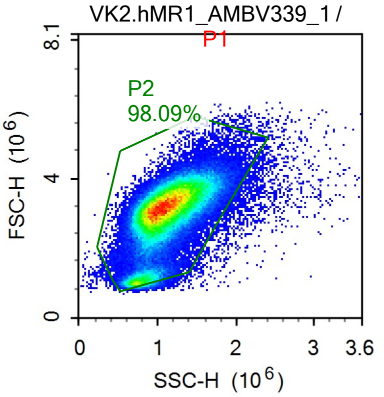


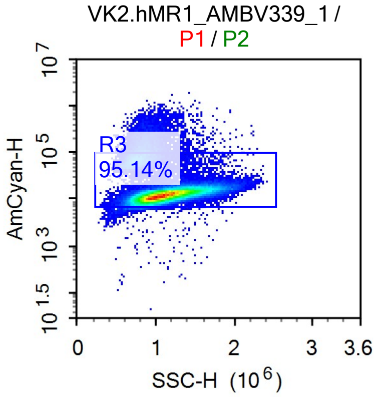


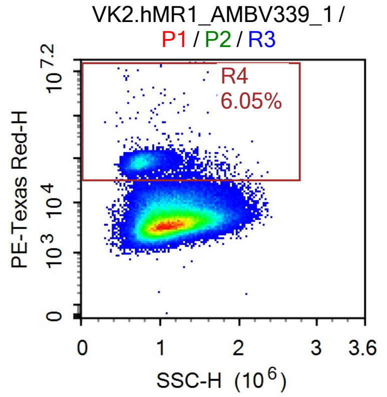


Sample Statistics

| Gate | Count | % Parent | X | Y | Median X | Median Y |
|---|---|---|---|---|---|---|
| All | 134,881 | | | | | |
| P1 | 109,906 | 81.48% | FSC-A | FSC-H | 7,394,257 | 3,128,506 |
| P2 | 107,806 | 98.09% | SSC-H | FSC-H | 1,091,235 | 3,129,387 |
| R3 | 102,571 | 95.14% | SSC-H | AmCyan-H | 1,104,107 | 12,823 |
| R4 | 6,205 | 6.05% | SSC-H | PE-Texas Red-H | 768,713 | 80,883 |

### Report of Specimen9-VK2.hMR1_AMBV339_

Sample Name: Specimen9-VK2.hMR1_AMBV339_2 | Run Time: 19-Mar-25 5:26 PM

Cytometer: NovoCyte Quanteon 621181110427 | Software: NovoExpress 1.6.0

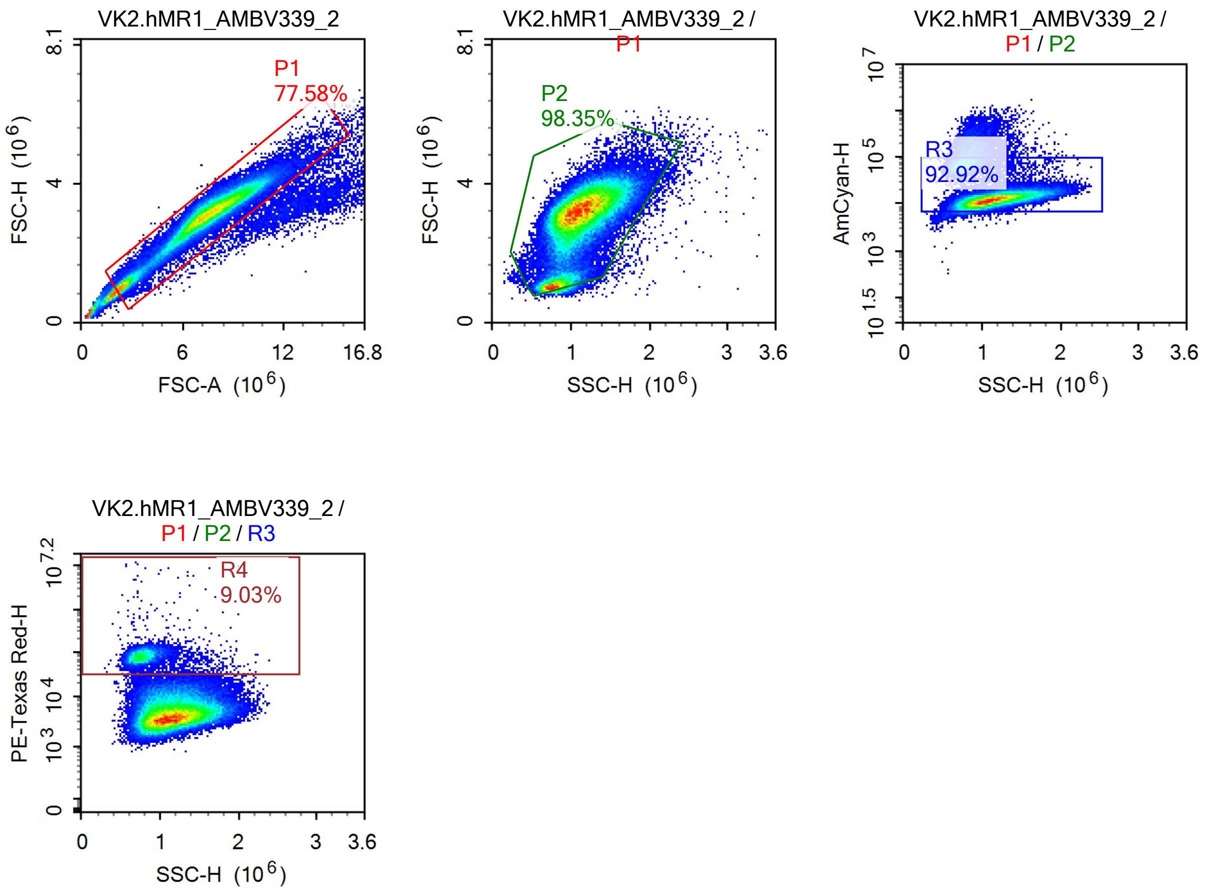


Sample Statistics

| Gate | Count | % Parent | X | Y | Median X | Median Y |
|---|---|---|---|---|---|---|
| All | 103,721 | | | | | |
| P1 | 80,464 | 77.58% | FSC-A | FSC-H | 7,365,231 | 3,088,601 |
| P2 | 79,137 | 98.35% | SSC-H | FSC-H | 1,093,111 | 3,093,364 |
| R3 | 73,532 | 92.92% | SSC-H | AmCyan-H | 1,112,821 | 13,074 |
| R4 | 6,637 | 9.03% | SSC-H | PE-Texas Red-H | 769,154 | 80,695 |

### Report of Specimen9-VK2.hMR1_AMBV339_

Sample Name: Specimen9-VK2.hMR1_AMBV339_3 — Run Time: 19-Mar-25 5:28 PM

Cytometer: NovoCyte Quanteon 621181110427 — Software: NovoExpress 1.6.0

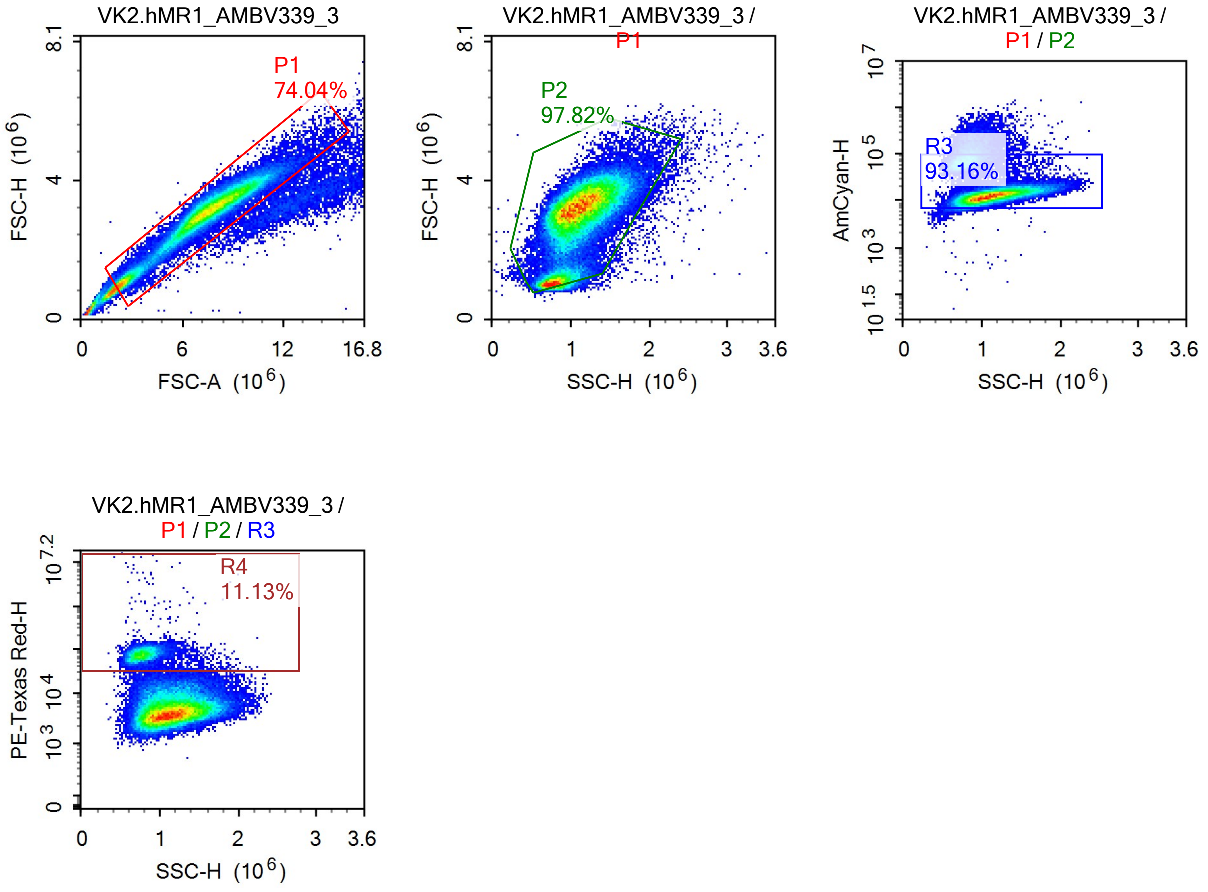


Sample Statistics

| Gate | Count | % Parent | X | Y | Median X | Median Y |
|---|---|---|---|---|---|---|
| All | 95,551 | | | | | |
| P1 | 70,749 | 74.04% | FSC-A | FSC-H | 7,374,406 | 3,081,183 |
| P2 | 69,205 | 97.82% | SSC-H | FSC-H | 1,077,944 | 3,087,776 |
| R3 | 64,472 | 93.16% | SSC-H | AmCyan-H | 1,094,372 | 13,670 |
| R4 | 7,177 | 11.13% | SSC-H | PE-Texas Red-H | 773,630 | 74,777 |

### Report of Specimen9-VK2.hMR1_AMBV339_

Sample Name: Specimen9-VK2.hMR1_AMBV339_4 Run Time: 19-Mar-25 5:30 PM

Cytometer: NovoCyte Quanteon 621181110427 Software: NovoExpress 1.6.0

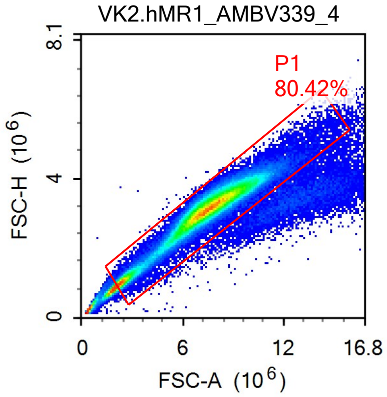


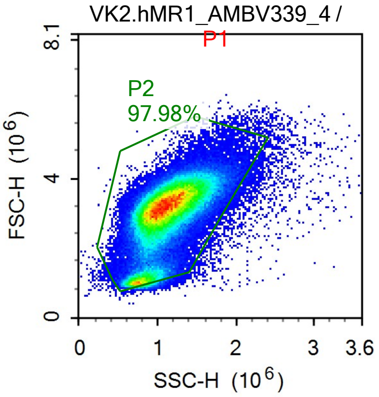


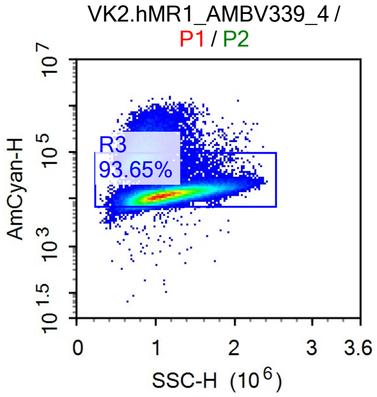


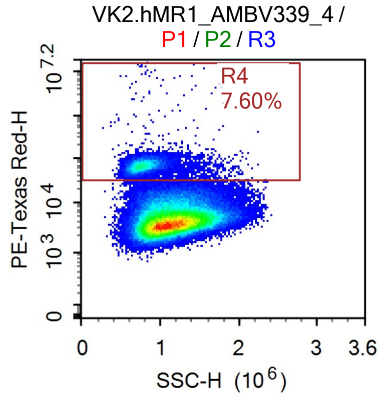


Sample Statistics

| Gate | Count | % Parent | X | Y | Median X | Median Y |
|---|---|---|---|---|---|---|
| All | 158,044 | | | | | |
| P1 | 127,095 | 80.42% | FSC-A | FSC-H | 7,384,728 | 3,123,634 |
| P2 | 124,522 | 97.98% | SSC-H | FSC-H | 1,090,682 | 3,123,951 |
| R3 | 116,621 | 93.65% | SSC-H | AmCyan-H | 1,107,537 | 12,918 |
| R4 | 8,867 | 7.60% | SSC-H | PE-Texas Red-H | 781,698 | 67,999 |

### Report of Specimen9-VK2_8uM_1

Sample Name: Specimen9-VK2_8uM_1 | Run Time: 19-Mar-25 5:32 PM

Cytometer: NovoCyte Quanteon 621181110427 | Software: NovoExpress 1.6.0

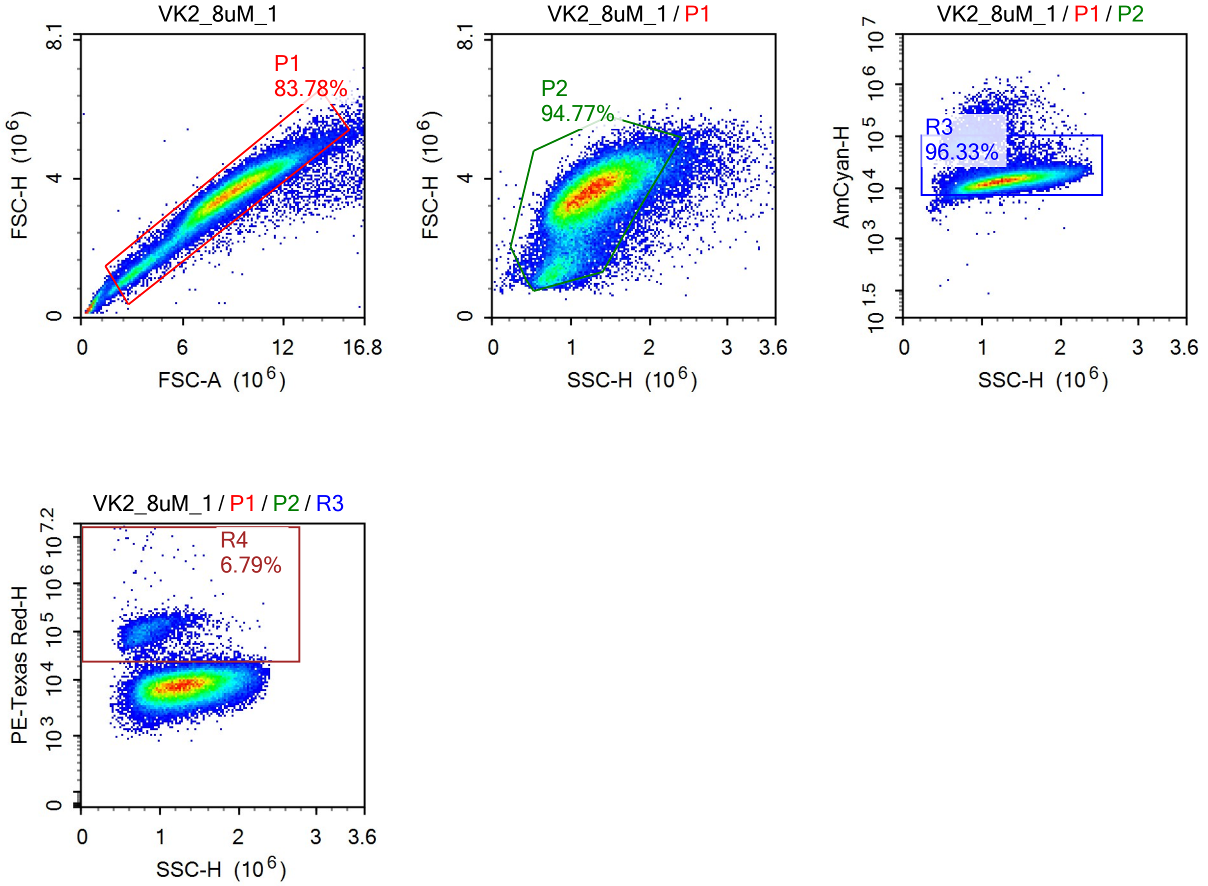


Sample Statistics

| Gate | Count | % Parent | X | Y | Median X | Median Y |
|---|---|---|---|---|---|---|
| All | 72,541 | | | | | |
| P1 | 60,778 | 83.78% | FSC-A | FSC-H | 8,705,908 | 3,544,727 |
| P2 | 57,600 | 94.77% | SSC-H | FSC-H | 1,251,396 | 3,529,520 |
| R3 | 55,486 | 96.33% | SSC-H | AmCyan-H | 1,263,665 | 13,906 |
| R4 | 3,765 | 6.79% | SSC-H | PE-Texas Red-H | 851,594 | 100,101 |

### Report of Specimen9-VK2_8uM_2

Sample Name: Specimen9-VK2_8uM_2 Run Time: 19-Mar-25 5:35 PM

Cytometer: NovoCyte Quanteon 621181110427 Software: NovoExpress 1.6.0

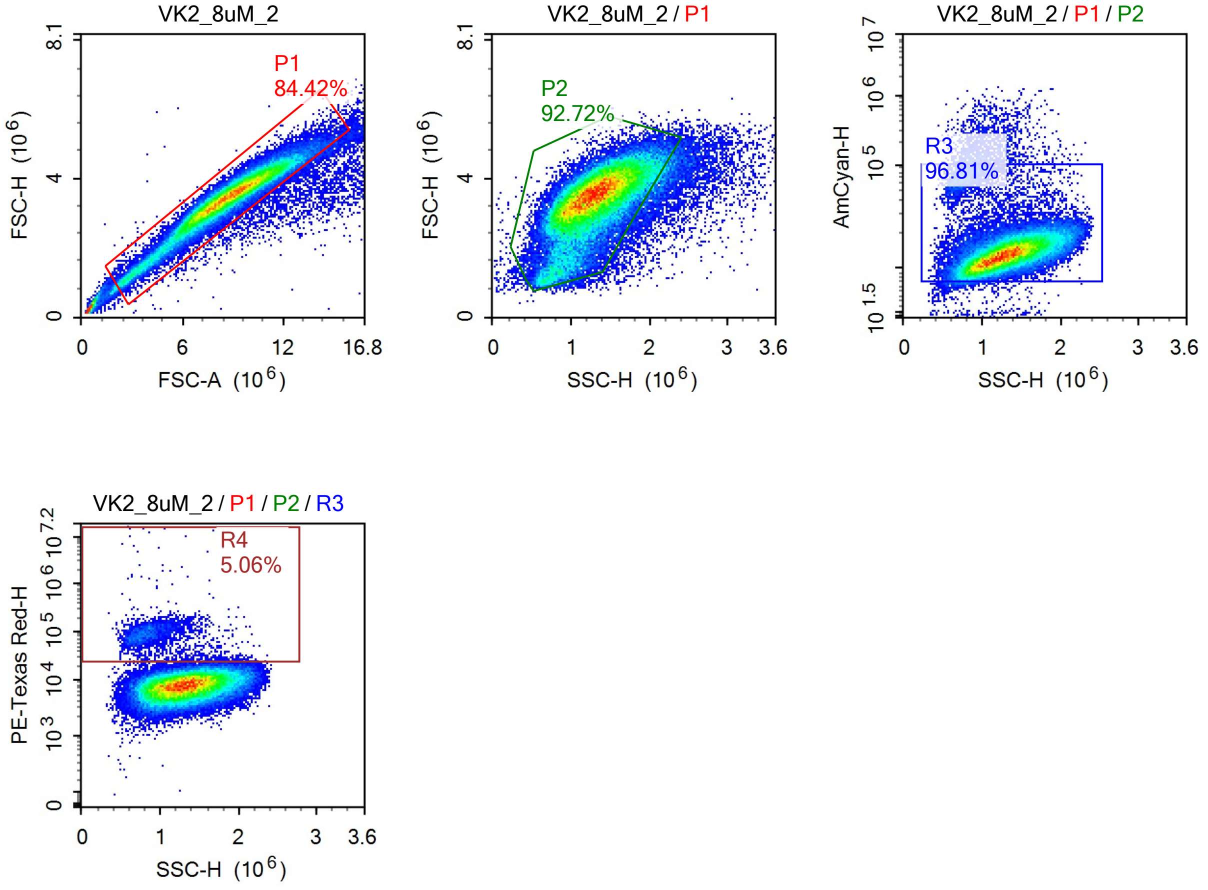


Sample Statistics

| Gate | Count | % Parent | X | Y | Median X | Median Y |
|---|---|---|---|---|---|---|
| All | 76,801 | | | | | |
| P1 | 64,834 | 84.42% | FSC-A | FSC-H | 8,668,046 | 3,455,183 |
| P2 | 60,115 | 92.72% | SSC-H | FSC-H | 1,283,807 | 3,433,937 |
| R3 | 58,196 | 96.81% | SSC-H | AmCyan-H | 1,293,369 | 13,833 |
| R4 | 2,942 | 5.06% | SSC-H | PE-Texas Red-H | 874,083 | 86,949 |

### Report of Specimen9-VK2_8uM_3

Sample Name: Specimen9-VK2_8uM_3

Run Time: 19-Mar-25 5:37 PM

Cytometer: NovoCyte Quanteon 621181110427

Software: NovoExpress 1.6.0

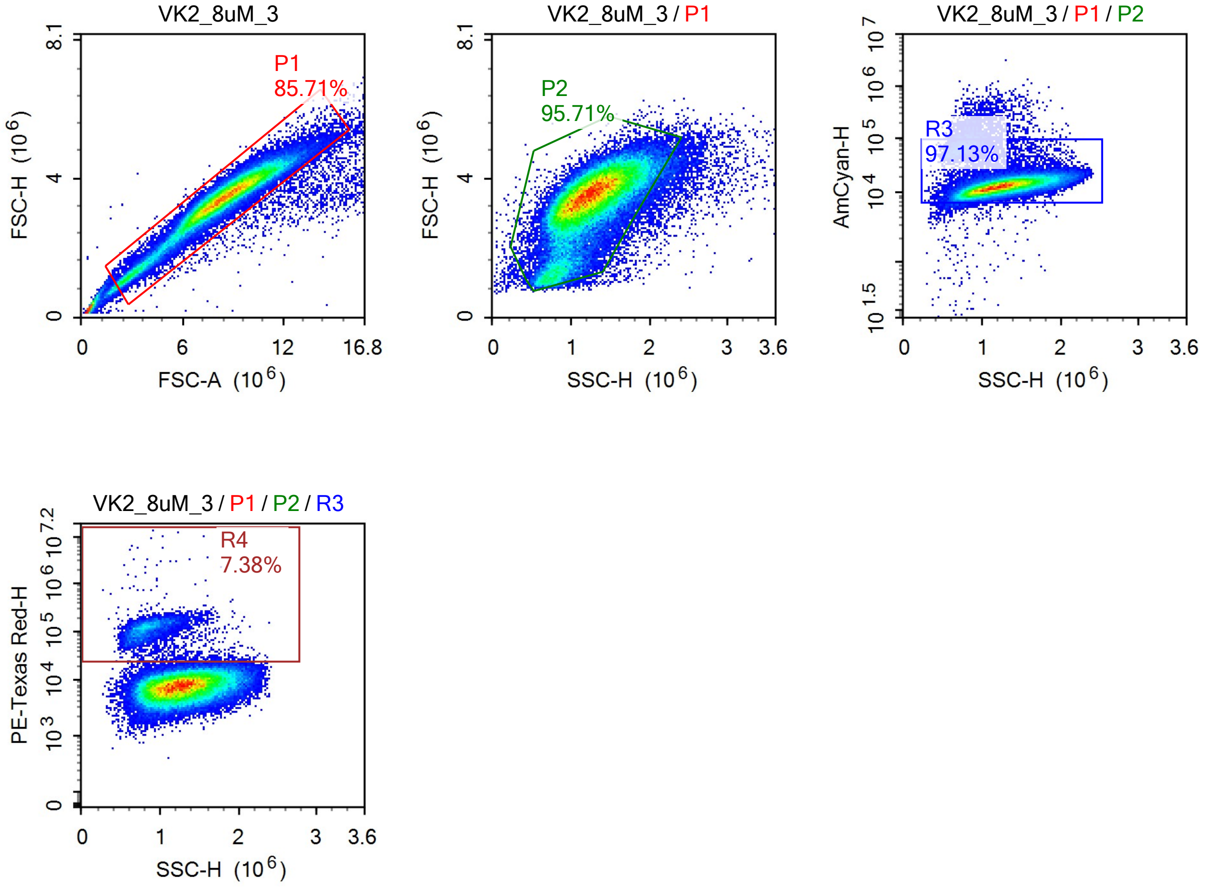


Sample Statistics

| Gate | Count | % Parent | X | Y | Median X | Median Y |
|---|---|---|---|---|---|---|
| All | 82,173 | | | | | |
| P1 | 70,428 | 85.71% | FSC-A | FSC-H | 8,405,141 | 3,447,312 |
| P2 | 67,407 | 95.71% | SSC-H | FSC-H | 1,232,131 | 3,435,778 |
| R3 | 65,472 | 97.13% | SSC-H | AmCyan-H | 1,241,265 | 13,449 |
| R4 | 4,835 | 7.38% | SSC-H | PE-Texas Red-H | 849,898 | 114,905 |

### Report of Specimen9-VK2_8uM_4

Sample Name: Specimen9-VK2_8uM_4 | Run Time: 19-Mar-25 5:39 PM

Cytometer: NovoCyte Quanteon 621181110427 | Software: NovoExpress 1.6.0

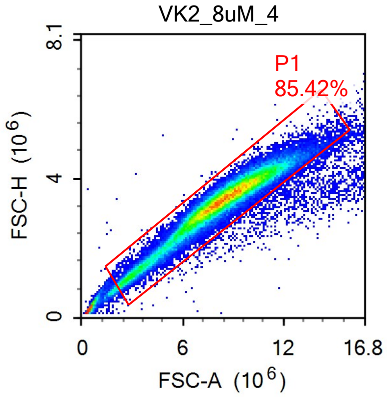


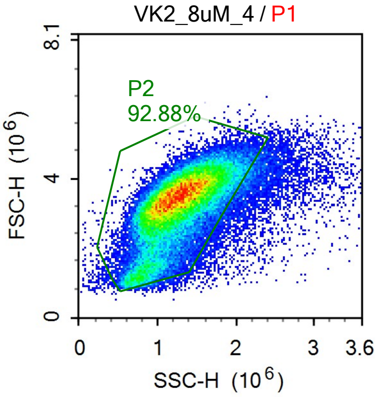


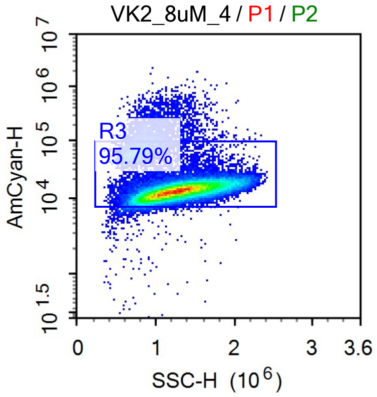


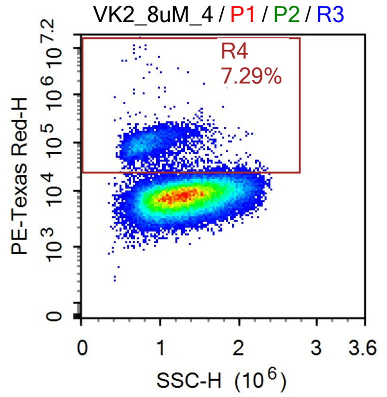


Sample Statistics

| Gate | Count | % Parent | X | Y | Median X | Median Y |
|---|---|---|---|---|---|---|
| All | 74,521 | | | | | |
| P1 | 63,659 | 85.42% | FSC-A | FSC-H | 8,283,757 | 3,368,992 |
| P2 | 59,126 | 92.88% | SSC-H | FSC-H | 1,250,567 | 3,351,860 |
| R3 | 56,638 | 95.79% | SSC-H | AmCyan-H | 1,263,226 | 13,412 |
| R4 | 4,131 | 7.29% | SSC-H | PE-Texas Red-H | 850,899 | 93,216 |

### Report of Specimen9-VK2.hMR1_8uM_1

Sample Name: Specimen9-VK2.hMR1_8uM_1 | Run Time: 19-Mar-25 5:41 PM

Cytometer: NovoCyte Quanteon 621181110427 | Software: NovoExpress 1.6.0

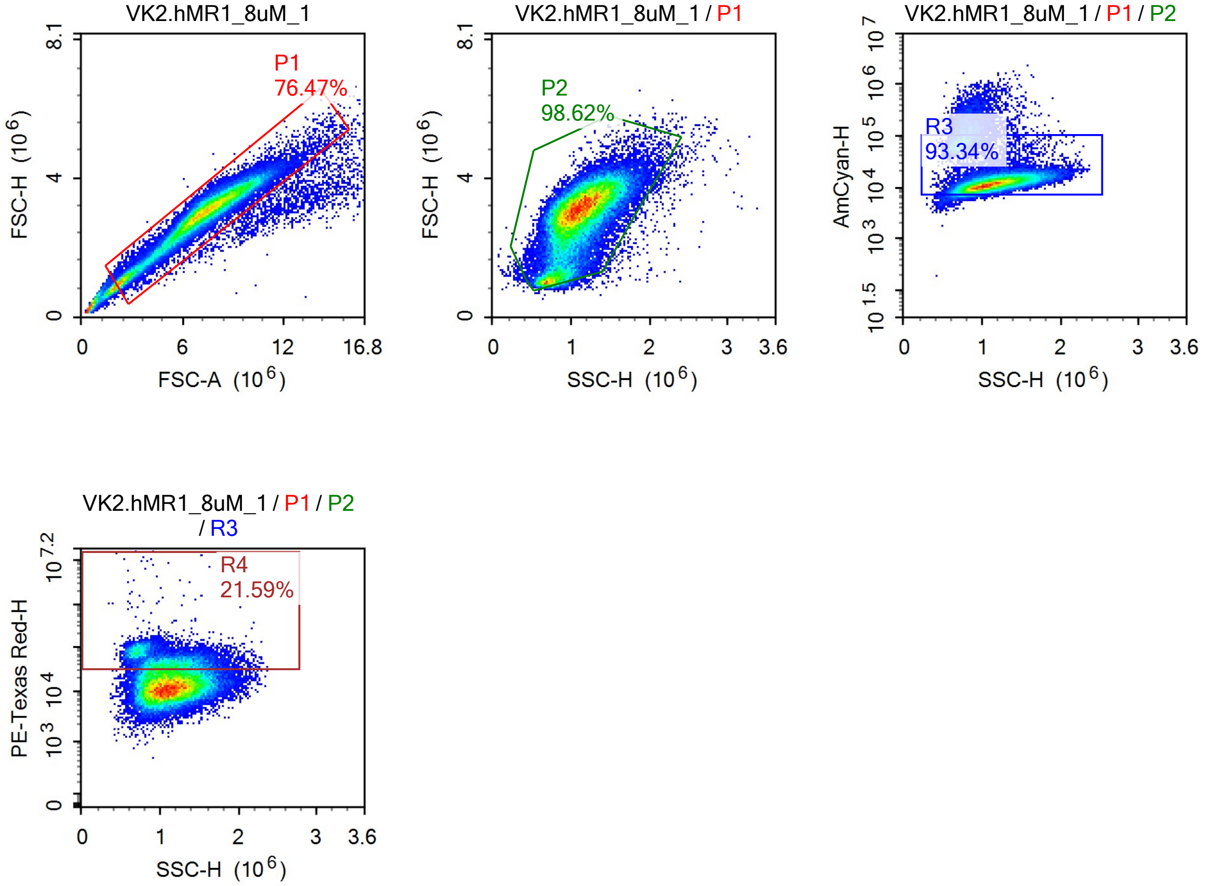


Sample Statistics

| Gate | Count | % Parent | X | Y | Median X | Median Y |
|---|---|---|---|---|---|---|
| All | 69,052 | | | | | |
| P1 | 52,804 | 76.47% | FSC-A | FSC-H | 7,155,727 | 2,982,166 |
| P2 | 52,077 | 98.62% | SSC-H | FSC-H | 1,079,640 | 2,984,785 |
| R3 | 48,609 | 93.34% | SSC-H | AmCyan-H | 1,099,276 | 11,962 |
| R4 | 10,495 | 21.59% | SSC-H | PE-Texas Red-H | 1,059,897 | 49,113 |

### Report of Specimen9-VK2.hMR1_8uM_2

Sample Name: Specimen9-VK2.hMR1_8uM_2 Run Time: 19-Mar-25 5:44 PM

Cytometer: NovoCyte Quanteon 621181110427 Software: NovoExpress 1.6.0

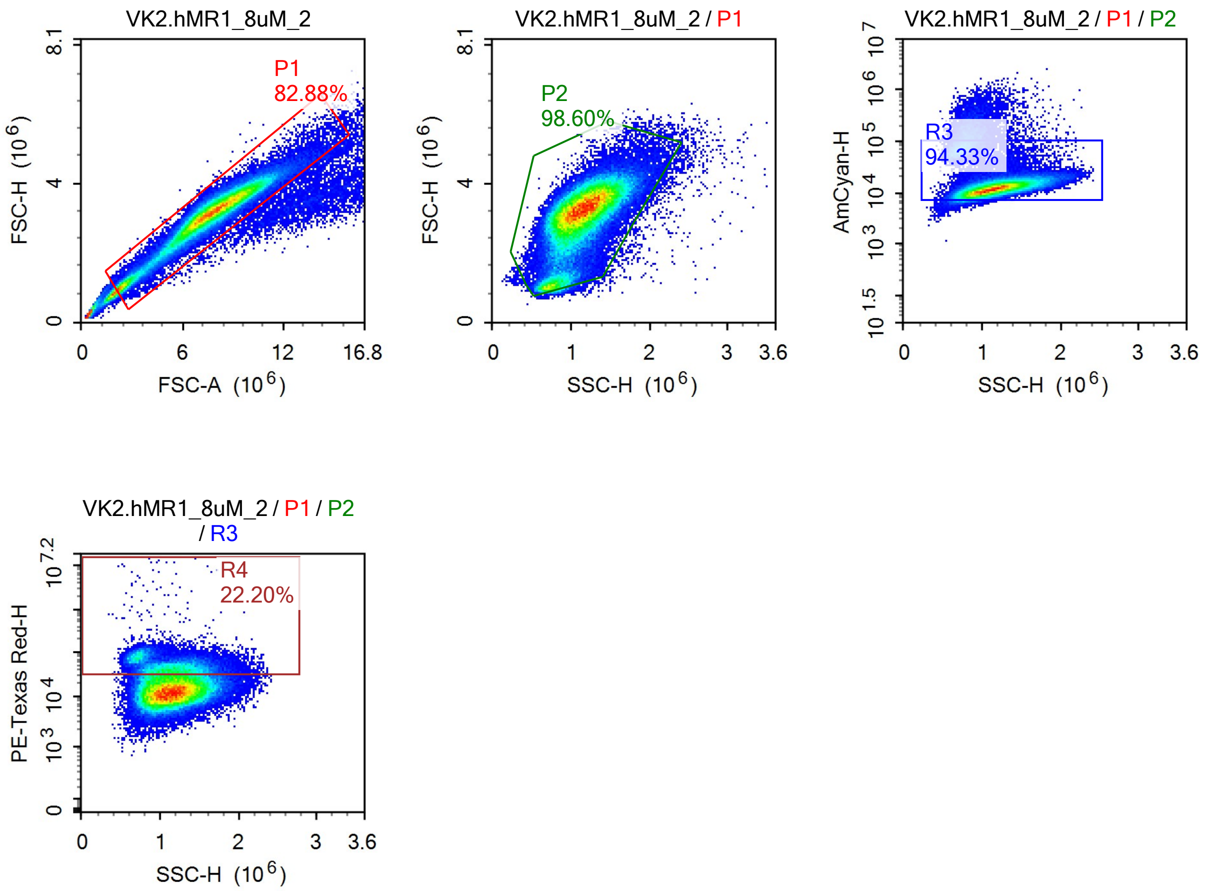


Sample Statistics

| Gate | Count | % Parent | X | Y | Median X | Median Y |
|---|---|---|---|---|---|---|
| All | 114,931 | | | | | |
| P1 | 95,259 | 82.88% | FSC-A | FSC-H | 7,661,931 | 3,158,153 |
| P2 | 93,923 | 98.60% | SSC-H | FSC-H | 1,117,269 | 3,158,415 |
| R3 | 88,600 | 94.33% | SSC-H | AmCyan-H | 1,134,857 | 12,376 |
| R4 | 19,666 | 22.20% | SSC-H | PE-Texas Red-H | 1,121,696 | 48,436 |

### Report of Specimen9-VK2.hMR1_8uM_3

Sample Name: Specimen9-VK2.hMR1_8uM_3 Run Time: 19-Mar-25 5:46 PM

Cytometer: NovoCyte Quanteon 621181110427 Software: NovoExpress 1.6.0

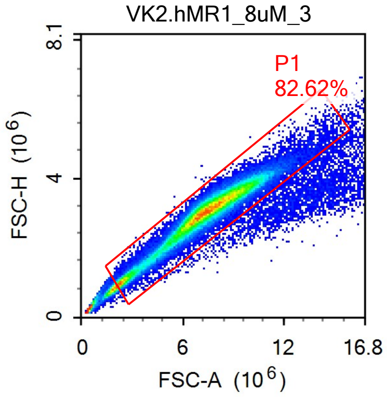


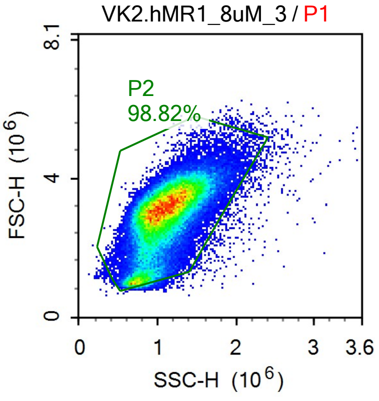


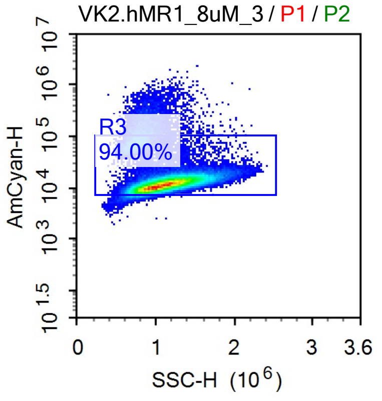


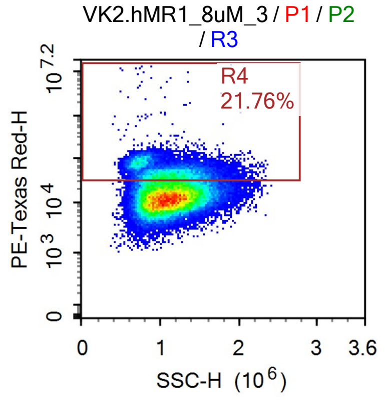


Sample Statistics

| Gate | Count | % Parent | X | Y | Median X | Median Y |
|---|---|---|---|---|---|---|
| All | 92,933 | | | | | |
| P1 | 76,782 | 82.62% | FSC-A | FSC-H | 7,343,198 | 3,047,619 |
| P2 | 75,879 | 98.82% | SSC-H | FSC-H | 1,076,430 | 3,049,286 |
| R3 | 71,325 | 94.00% | SSC-H | AmCyan-H | 1,093,983 | 11,992 |
| R4 | 15,523 | 21.76% | SSC-H | PE-Texas Red-H | 1,079,416 | 48,338 |

### Report of Specimen9-VK2.hMR1_8uM_4

Sample Name: Specimen9-VK2.hMR1_8uM_4 Run Time: 19-Mar-25 5:48 PM

Cytometer: NovoCyte Quanteon 621181110427 Software: NovoExpress 1.6.0

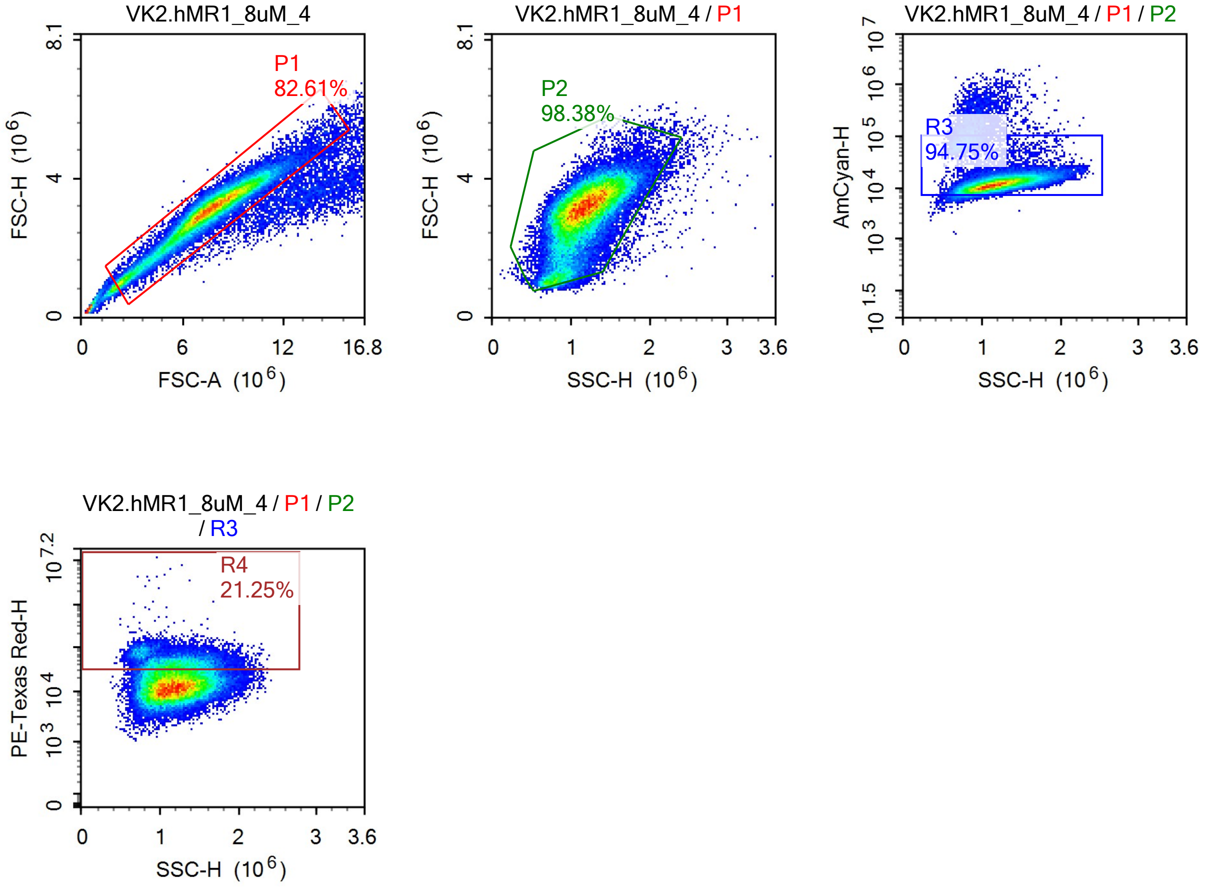


Sample Statistics

| Gate | Count | % Parent | X | Y | Median X | Median Y |
|---|---|---|---|---|---|---|
| All | 65,257 | | | | | |
| P1 | 53,912 | 82.61% | FSC-A | FSC-H | 7,572,097 | 3,103,816 |
| P2 | 53,040 | 98.38% | SSC-H | FSC-H | 1,152,443 | 3,102,489 |
| R3 | 50,257 | 94.75% | SSC-H | AmCyan-H | 1,167,760 | 12,007 |
| R4 | 10,680 | 21.25% | SSC-H | PE-Texas Red-H | 1,182,703 | 46,933 |

### Report of Specimen9-VK2_16uM_1

Sample Name: Specimen9-VK2_16uM_1

Run Time: 19-Mar-25 5:50 PM

Cytometer: NovoCyte Quanteon 621181110427

Software: NovoExpress 1.6.0

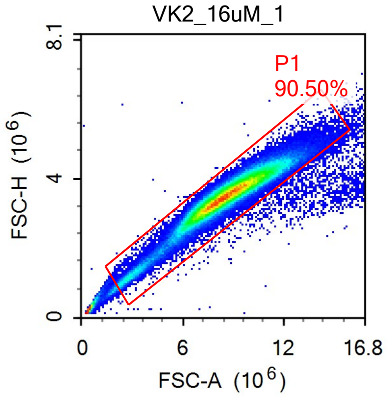


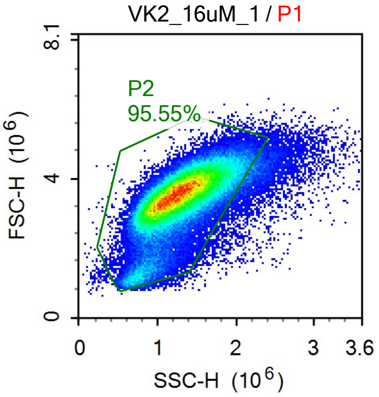


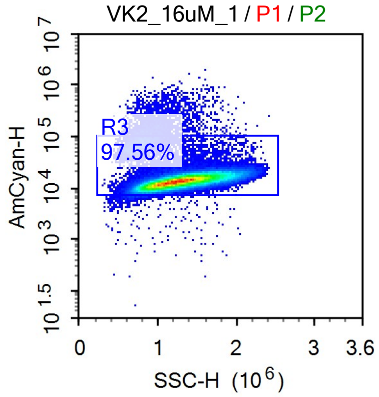


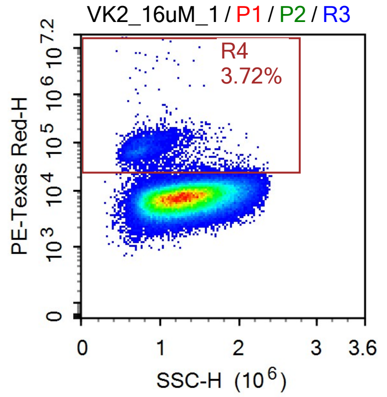


Sample Statistics

| Gate | Count | % Parent | X | Y | Median X | Median Y |
|---|---|---|---|---|---|---|
| All | 121,297 | | | | | |
| P1 | 109,776 | 90.50% | FSC-A | FSC-H | 8,608,862 | 3,531,274 |
| P2 | 104,892 | 95.55% | SSC-H | FSC-H | 1,273,352 | 3,514,047 |
| R3 | 102,335 | 97.56% | SSC-H | AmCyan-H | 1,280,717 | 13,761 |
| R4 | 3,804 | 3.72% | SSC-H | PE-Texas Red-H | 805,685 | 73,341 |

### Report of Specimen9-VK2_16uM_2

Sample Name: Specimen9-VK2_16uM_2 | Run Time: 19-Mar-25 5:53 PM

Cytometer: NovoCyte Quanteon 621181110427 | Software: NovoExpress 1.6.0

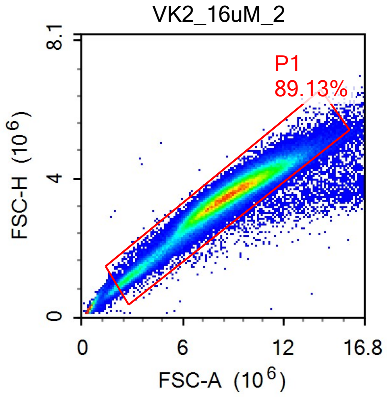


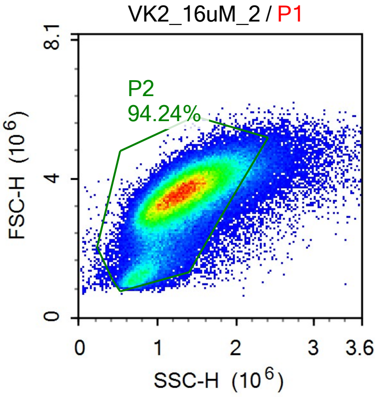


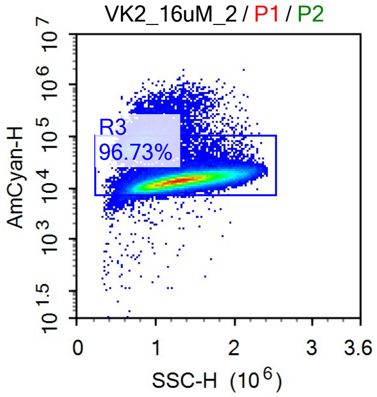


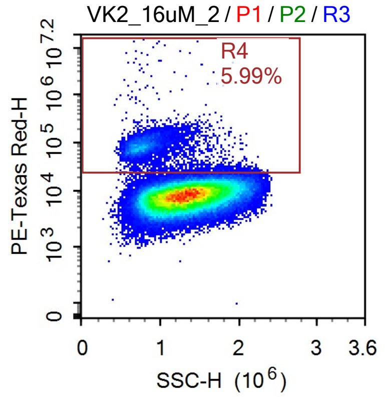


Sample Statistics

| Gate | Count | % Parent | X | Y | Median X | Median Y |
|---|---|---|---|---|---|---|
| All | 138,788 | | | | | |
| P1 | 123,699 | 89.13% | FSC-A | FSC-H | 8,679,775 | 3,512,208 |
| P2 | 116,575 | 94.24% | SSC-H | FSC-H | 1,292,612 | 3,488,345 |
| R3 | 112,761 | 96.73% | SSC-H | AmCyan-H | 1,302,101 | 14,239 |
| R4 | 6,756 | 5.99% | SSC-H | PE-Texas Red-H | 806,337 | 84,319 |

### Report of Specimen9-VK2_16uM_3

Sample Name: Specimen9-VK2_16uM_3 | Run Time: 19-Mar-25 5:55 PM

Cytometer: NovoCyte Quanteon 621181110427 | Software: NovoExpress 1.6.0

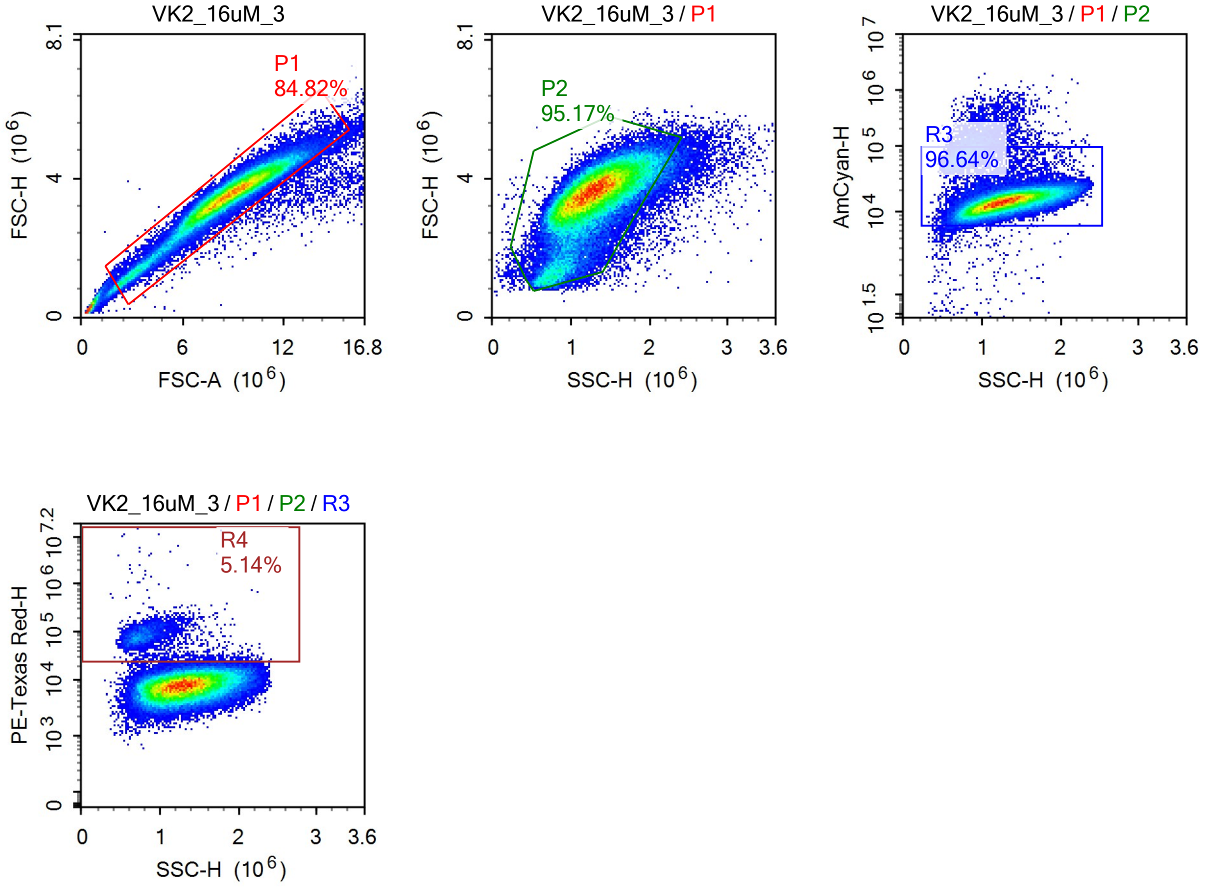


Sample Statistics

| Gate | Count | % Parent | X | Y | Median X | Median Y |
|---|---|---|---|---|---|---|
| All | 89,661 | | | | | |
| P1 | 76,050 | 84.82% | FSC-A | FSC-H | 8,768,270 | 3,541,852 |
| P2 | 72,379 | 95.17% | SSC-H | FSC-H | 1,272,147 | 3,525,819 |
| R3 | 69,948 | 96.64% | SSC-H | AmCyan-H | 1,282,324 | 14,277 |
| R4 | 3,593 | 5.14% | SSC-H | PE-Texas Red-H | 814,662 | 80,424 |

### Report of Specimen9-VK2_16uM_4

Sample Name: Specimen9-VK2_16uM_4 Run Time: 19-Mar-25 5:57 PM

Cytometer: NovoCyte Quanteon 621181110427 Software: NovoExpress 1.6.0

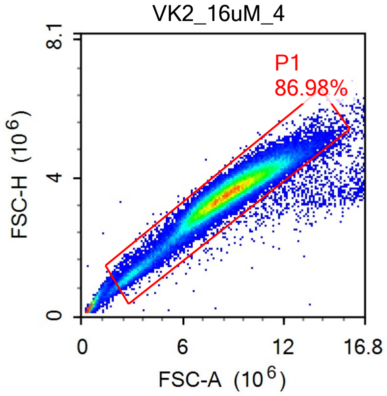


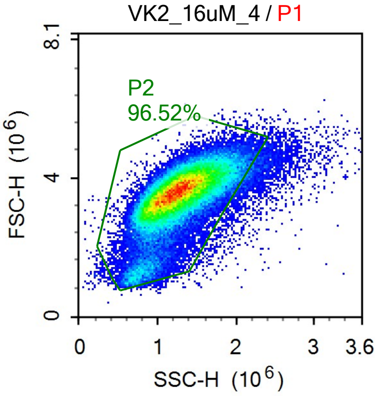


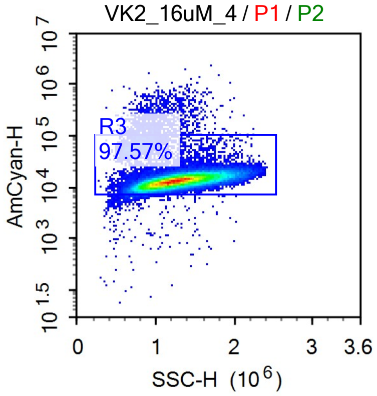


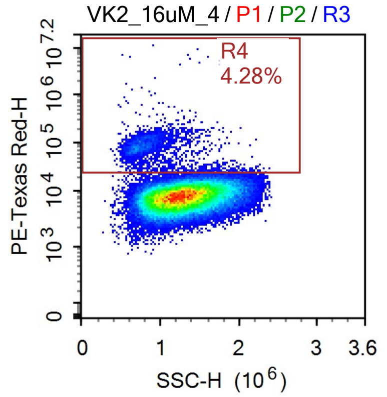


Sample Statistics

| Gate | Count | % Parent | X | Y | Median X | Median Y |
|---|---|---|---|---|---|---|
| All | 83,563 | | | | | |
| P1 | 72,684 | 86.98% | FSC-A | FSC-H | 8,672,274 | 3,564,755 |
| P2 | 70,156 | 96.52% | SSC-H | FSC-H | 1,256,278 | 3,552,946 |
| R3 | 68,452 | 97.57% | SSC-H | AmCyan-H | 1,263,443 | 13,518 |
| R4 | 2,933 | 4.28% | SSC-H | PE-Texas Red-H | 823,037 | 84,104 |

### Report of Specimen9-VK2.hMR1_16uM_1

Sample Name: Specimen9-VK2.hMR1_16uM_1 | Run Time: 19-Mar-25 6:00 PM

Cytometer: NovoCyte Quanteon 621181110427 | Software: NovoExpress 1.6.0

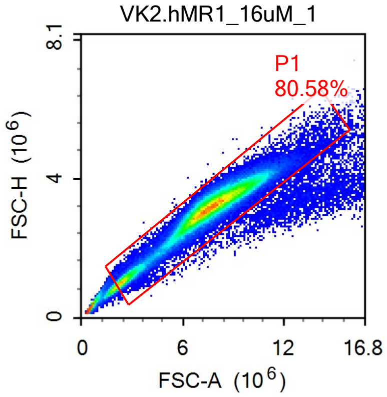


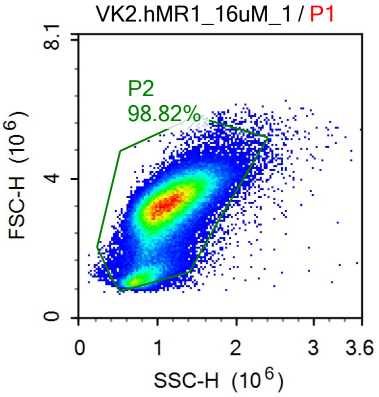


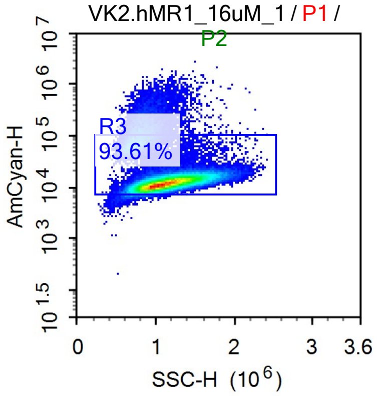


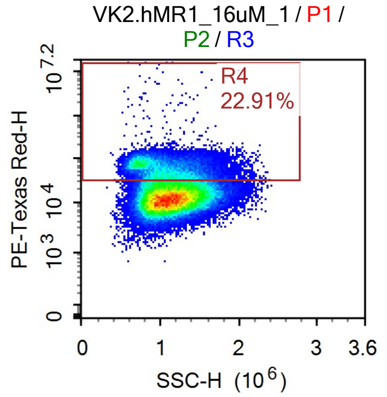


Sample Statistics

| Gate | Count | % Parent | X | Y | Median X | Median Y |
|---|---|---|---|---|---|---|
| All | 123,869 | | | | | |
| P1 | 99,811 | 80.58% | FSC-A | FSC-H | 7,551,016 | 3,175,370 |
| P2 | 98,638 | 98.82% | SSC-H | FSC-H | 1,093,178 | 3,176,243 |
| R3 | 92,339 | 93.61% | SSC-H | AmCyan-H | 1,110,828 | 12,363 |
| R4 | 21,155 | 22.91% | SSC-H | PE-Texas Red-H | 1,047,460 | 58,530 |

### Report of Specimen9-VK2.hMR1_16uM_2

Sample Name: Specimen9-VK2.hMR1_16uM_2 Run Time: 19-Mar-25 6:02 PM

Cytometer: NovoCyte Quanteon 621181110427 Software: NovoExpress 1.6.0

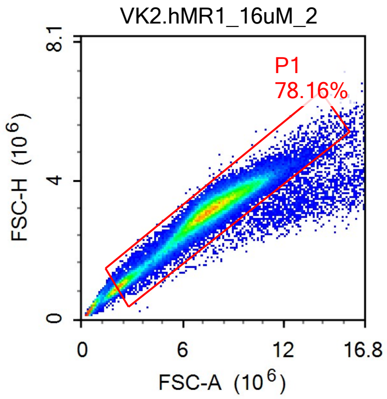


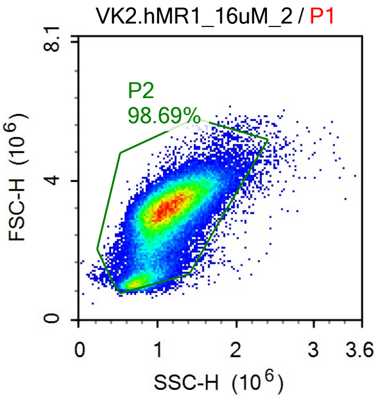


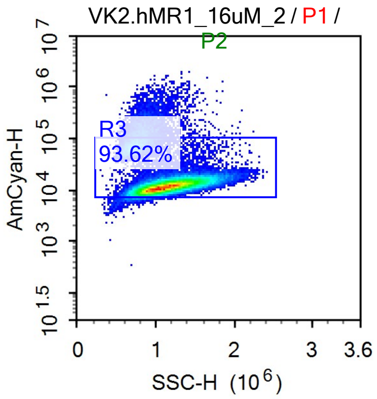


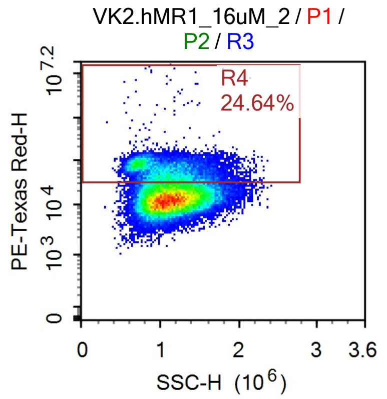


Sample Statistics

| Gate | Count | % Parent | X | Y | Median X | Median Y |
|---|---|---|---|---|---|---|
| All | 75,746 | | | | | |
| P1 | 59,202 | 78.16% | FSC-A | FSC-H | 7,433,419 | 3,102,420 |
| P2 | 58,424 | 98.69% | SSC-H | FSC-H | 1,097,063 | 3,103,484 |
| R3 | 54,694 | 93.62% | SSC-H | AmCyan-H | 1,115,956 | 12,136 |
| R4 | 13,475 | 24.64% | SSC-H | PE-Texas Red-H | 1,069,452 | 58,614 |

### Report of Specimen9-VK2.hMR1_16uM_3

Sample Name: Specimen9-VK2.hMR1_16uM_3 | Run Time: 19-Mar-25 6:04 PM

Cytometer: NovoCyte Quanteon 621181110427 | Software: NovoExpress 1.6.0

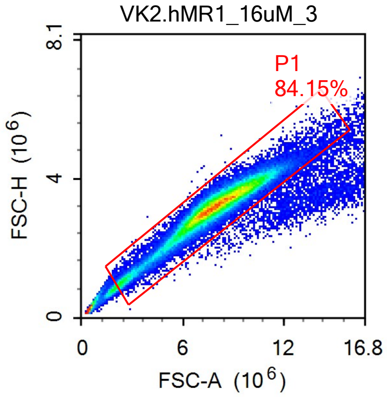


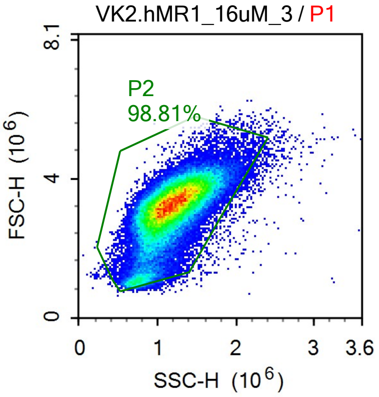


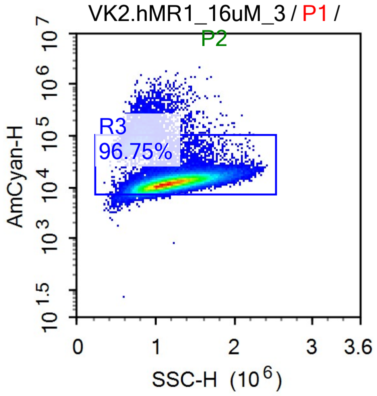


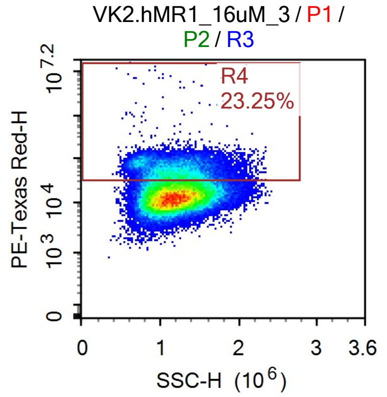


Sample Statistics

| Gate | Count | % Parent | X | Y | Median X | Median Y |
|---|---|---|---|---|---|---|
| All | 85,821 | | | | | |
| P1 | 72,217 | 84.15% | FSC-A | FSC-H | 7,873,726 | 3,217,732 |
| P2 | 71,355 | 98.81% | SSC-H | FSC-H | 1,166,006 | 3,216,748 |
| R3 | 69,033 | 96.75% | SSC-H | AmCyan-H | 1,175,554 | 12,328 |
| R4 | 16,052 | 23.25% | SSC-H | PE-Texas Red-H | 1,186,769 | 54,379 |

### Report of Specimen9-VK2.hMR1_16uM_4

Sample Name: Specimen9-VK2.hMR1_16uM_4 | Run Time: 19-Mar-25 6:07 PM

Cytometer: NovoCyte Quanteon 621181110427 | Software: NovoExpress 1.6.0

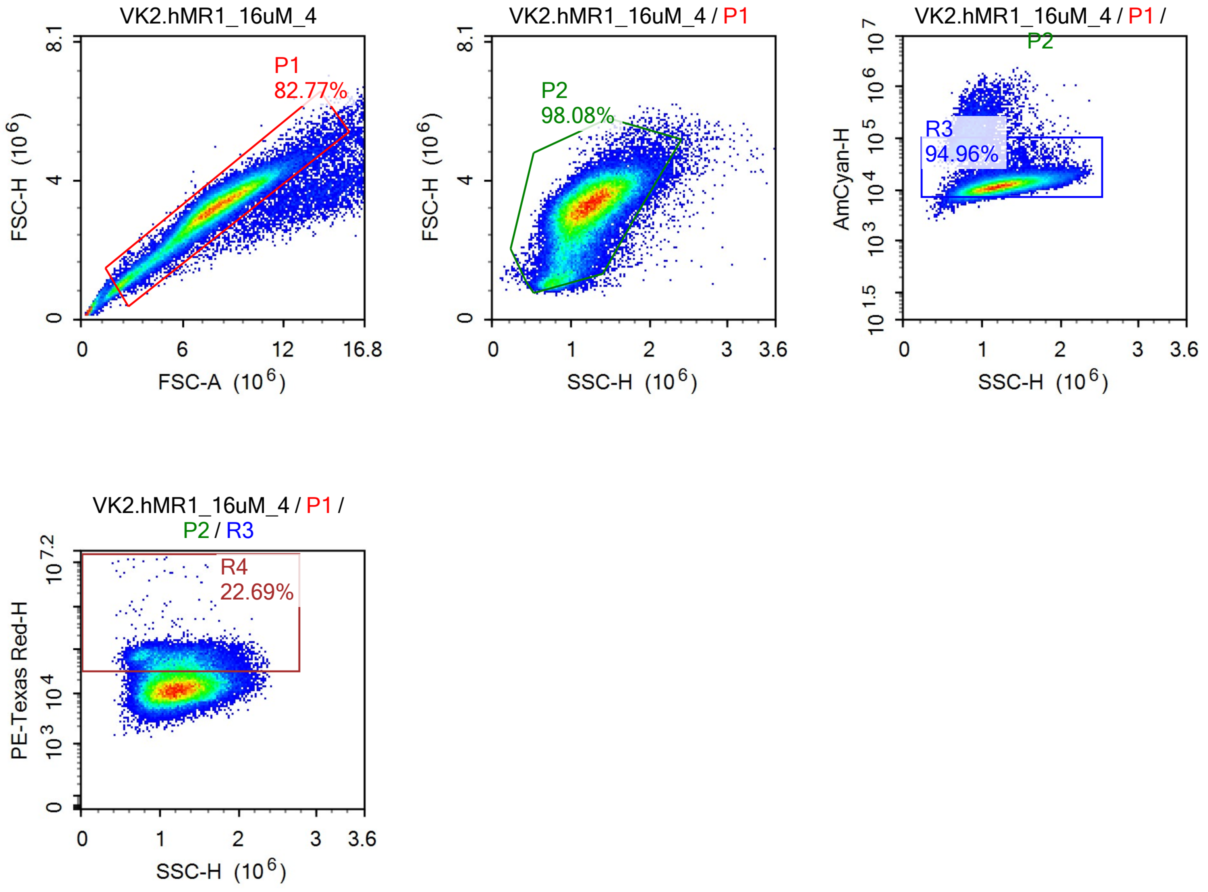


Sample Statistics

| Gate | Count | % Parent | X | Y | Median X | Median Y |
|---|---|---|---|---|---|---|
| All | 73,987 | | | | | |
| P1 | 61,238 | 82.77% | FSC-A | FSC-H | 7,794,790 | 3,189,841 |
| P2 | 60,065 | 98.08% | SSC-H | FSC-H | 1,194,713 | 3,186,915 |
| R3 | 57,037 | 94.96% | SSC-H | AmCyan-H | 1,210,589 | 12,139 |
| R4 | 12,940 | 22.69% | SSC-H | PE-Texas Red-H | 1,214,260 | 53,478 |
